# A single cell transcriptional roadmap for cardiopharyngeal fate diversification

**DOI:** 10.1101/150235

**Authors:** Wei Wang, Xiang Niu, Tim Stuart, Estelle Jullian, William Mauck, Robert G. Kelly, Rahul Satija, Lionel Christiaen

## Abstract

In vertebrates, multipotent progenitors located in the pharyngeal mesoderm form cardiomyocytes and branchiomeric head muscles, but the dynamic gene expression programs and mechanisms underlying cardiopharyngeal multipotency and heart vs. head muscle fate choices remain elusive. Here, we used single cell genomics in the simple chordate model Ciona, to reconstruct developmental trajectories forming first and second heart lineages, and pharyngeal muscle precursors, and characterize the molecular underpinnings of cardiopharyngeal fate choices. We show that FGF-MAPK signaling maintains multipotency and promotes the pharyngeal muscle fate, whereas signal termination permits the deployment of a pan-cardiac program, shared by the first and second lineages, to define heart identity. In the second heart lineage, a Tbx1/10-Dach pathway actively suppresses the first heart lineage program, conditioning later cell diversity in the beating heart. Finally, cross-species comparisons between Ciona and the mouse evoke the deep evolutionary origins of cardiopharyngeal networks in chordates.

---

Distinct cell types form multicellular animals and execute specialized functions within defined organs and systems, implying that individual cells within progenitor fields must acquire both organ-level and cell-type-specific identities. The mammalian heart comprises chamber-specific cardiomyocytes, various endocardial cell types, fibroblasts and smooth muscles^1^, and despite their specialized features, these cells share a cardiac identity. Popular models posit that heart cells emerge from multipotent cardiovascular progenitors^2^, implying that multipotent progenitors are first imbued with a cardiac identity, before producing a diversity of cell types. Consistent with this model, mammalian heart cells emerge primarily from *Mesp1*+ mesodermal progenitors^3–5^. However, lineage tracing and clonal analyses indicated that distinct compartments arise from separate progenitor pools, referred to as the first and second heart fields^6–11^. In addition, most early cardiac progenitors produce only one cell type^12^, and cell type segregation occurs early, possibly prior to commitment to a heart identity^13^. Moreover, derivatives of the second heart field (e.g. cardiomyocytes of the right ventricle and outflow tract) share a common origin with branchiomeric head muscles, in the cardiopharyngeal mesoderm^4,14–21^. The characteristics of multipotent cardiopharyngeal progenitors, and the mechanisms underlying early heart vs. pharyngeal/branchiomeric muscle fate choices remain largely elusive, and studies are partially hindered by the complexity of vertebrate embryos^21,22^.

The tunicate Ciona emerged as an innovative chordate model to study cardiopharyngeal development with unprecedented spatio-temporal resolution. In Ciona, invariant cell divisions produce distinct first and second heart lineages, and pharyngeal muscle precursors from defined multipotent cardiopharyngeal progenitors^23,24^ (Fig. 1a). Multipotent progenitors exhibit multilineage transcriptional priming, whereby conserved fate-specific determinants are transiently co-expressed, before regulatory cross-antagonisms partition the heart and pharyngeal muscle programs to their corresponding fate-restricted precursors^23–25^. Here, we characterised, with single cell resolution, the genome-wide characteristics and regulatory mechanisms governing cardiopharyngeal multipotency, early fate choices, and the establishment of cell diversity in the beating heart.

**Figure 1.**
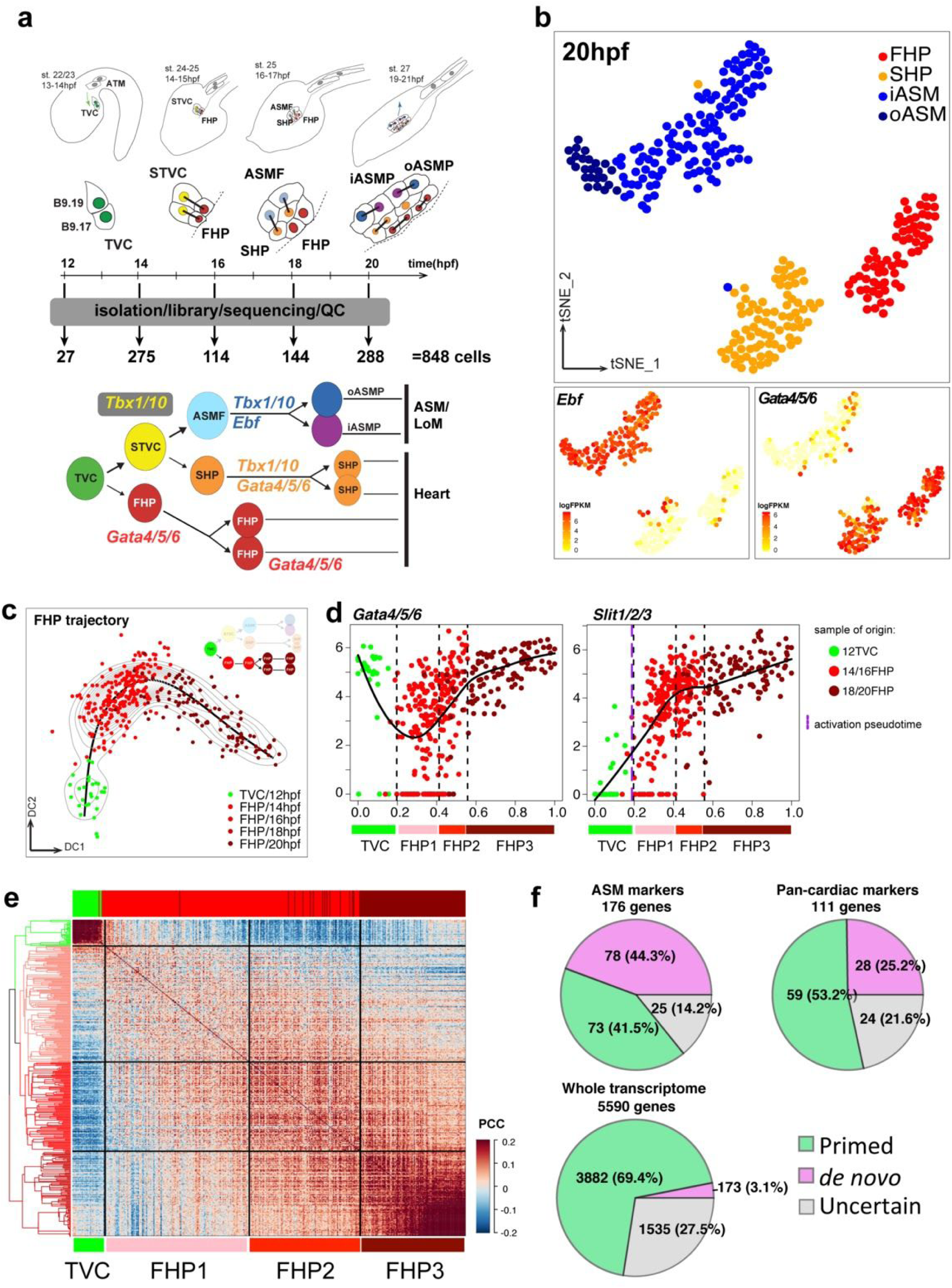
Cell clustering and pseudotemporal reconstruction of cardiopharyngeal developmental trajectories. (**a**) Early cardiopharyngeal development in *Ciona*, and sampling stages for scRNA-seq and established TVC lineage tree. Cardiopharyngeal lineage cells are shown for only one side and known cell-type-specific marker genes are indicated. st., stage according to ^72^; hpf, hours post-fertilization; TVC, trunk ventral cell; STVC, second trunk ventral cell; FHP, first heart precursor; ASMF, atrial siphon muscle founder cells; SHP, second heart precursor; iASMP, inner atrial siphon muscle precursor; oASMP, outer atrial siphon muscle precursor, according to ^73^; LoM, longitudinal muscles; QC, quality control. Dotted line: midline. (**b**) t-distributed Stochastic Neighbor Embedding (t-SNE) plots of 20 hpf scRNA-seq data showing distinct clusters of progenitor subtypes: FHP (red), SHP (orange), iASM (blue) and oASM(dark blue). Indicated marker gene expression levels are color-coded and shown on corresponding clusters. (**c**) 2D diffusion map of single cell transcriptomes for the FHP developmental trajectory. Black lines: principal curve; light gray contours: single cell density distribution. Color codes correspond to assigned cell identities following clustering at each time point. hpf, hours post-fertilization. Cartoon shows the cells used to reconstruct the unidirectional FHP trajectory. (**d**) Pseudotemporal expression profiles of indicated genes along the FHP trajectory. X-axis: normalized pseudotime as defined in (e), Y-axes: expression in log_2_ FPKM. Black lines indicate the smoothed expression. Black dashed lines indicate the transitions between predicted regulatory states, indicated and color-coded below, and purple dashed lines indicate calculated activation pseudotime. Dot colors refer to the sample of origin as indicated in (c). (**e**) Cross-correlation heatmaps to infer regulatory states along the FHP trajectory. Dendrogram (left) obtained from constrained hierarchical clustering. Top bars indicate the sample of origin with color codes as in (c). PCC, Pearson Correlation Coefficient. (**f**) Proportions of primed vs. *de novo* genes among defined categories of marker genes.

## RESULTS

### Single cell transcriptome profiling of early cardiopharyngeal lineages

To characterize gene expression changes underlying the transitions from multipotent progenitors to distinct fate-restricted precursors, we performed plate-based single cell RNA sequencing (scRNA-seq) with SMART-Seq2^26^ on cardiopharyngeal-lineage cells FACS-purified from synchronously developing embryos and larvae (Fig. 1a). We obtained 848 high-quality single cell transcriptomes from 5 time points covering early cardiopharyngeal development (Fig. 1a, Extended Data Fig. 1d). Using an unsupervised strategy^27^, we clustered single cell transcriptomes from each time point, and identified clusters according to known markers and previously established lineage information (Fig. 1a-b, Extended Data Fig. 1a, c). Focusing on fate-restricted cells isolated from post-hatching larvae (18 and 20 hours post-fertilization (hpf), FABA stages 26-28; Extended Data Table 1), we identified clusters of *Gata4/5/6*+ first and second heart precursors, and *Ebf*+ atrial siphon muscle (ASM) precursors^24,25,28^ (Fig. 1b). Differential expression analyses identified (1) ASM/pharyngeal muscle vs. pan-cardiac specific markers and (2) first vs. second heart precursor-specific markers (Extended Data Fig. 1b, Extended Data Table 2). The top 111 predicted pan-cardiac genes comprised established cardiac determinants, including *Gata4/5/6*, *Nk4/Nkx2-5*, and *Hand*, and we confirmed heart-specific expression by fluorescent *in situ* hybridization (FISH) for 19 candidate markers, including *Meis* and *Lrp4/8* (Extended Data Fig. 1e,g; Extended Data Table 2, 3). The pan-cardiac vs. pharyngeal muscle contrast dominated late cellular heterogeneity, but first and second heart precursor populations also segregated (Fig. 1b, Extended Data Fig. 1a), revealing 18 and 7 first- and second-lineage-specific markers, respectively (e.g. *Mmp21* and *Dach*; Extended Data Fig. 1f; Extended Data Tables 2,3). Our analyses thus uncovered specific programs activated in fate-committed progenitors, including both shared (‘pan-cardiac’) and first- vs. second-lineage-specific signatures for heart precursors.

To characterize gene expression dynamics, we ordered single cell transcriptomes from successive time points on pseudotemporal developmental projections^29,30^. Using the whole dataset while ignoring established clonality, we identified multipotent progenitors and separate cardiac and pharyngeal muscles branches (Extended Data Fig. 2). However, this unsupervised analysis failed to correctly distinguish the first and second heart lineages, likely because the shared pan-cardiac program dominates lineage-specific signatures (Extended Data Fig. 1b). Taking advantage of the invariant lineage (Fig. 1a), we combined cells corresponding to each branch, and created three unidirectional trajectories representing first- and second heart, and pharyngeal muscle lineages (Fig. 1c, Extended Data Fig. 3a-b). The distribution of cells along pseudotime axes corresponded to the time points of origin (Fig. 1c, Extended Data Fig. 3b, g; average Pearson Correlation Coefficient, PCC = 0.889), while providing higher-resolution insights into the gene expression dynamics. Validating this approach, *in silico* trajectories captured known lineage-specific expression changes of cardiopharyngeal regulators (Fig. 1d, Extended Data Fig. 3c-d; ^24,25,31^).

Developmental trajectories suggest a continuous process marked by gradual changes in gene expression. However, the latter occur preferentially in defined ‘pseudotime’ windows for multiple genes (Extended Data Fig. 3e-f), consistent with more abrupt biological transitions, such as cell divisions^31^. To identify possible switch-like discontinuities, we determined cell-to-cell cross-correlations along lineage-specific trajectories. Using constrained hierarchical clustering^32^, we identified 10 putative discrete regulatory states across the cardiopharyngeal trajectories, including two multipotent states, and eight successive transitions towards fate restriction (Fig. 1e, Extended Data Fig. 3h).

These successive transitions reveal underlying lineage-specific transcriptional dynamics. For example, the multipotent cardiopharyngeal progenitor state (aka TVC) differed markedly from subsequent cardiac states along the first cardiac trajectory (Fig. 1e), and gene expression mapping distinguished ‘primed’ and ‘*de novo*’-expressed heart markers, such as *Gata4/5/6* and *Slit1/2/3*, respectively (Fig. 1d). Conversely, primed pharyngeal muscle markers were downregulated along cardiac trajectories, as expected (Extended Data Fig. 3c, e-f; ^33,34^). Multilineage transcriptional priming is a hallmark of cardiopharyngeal multipotency that remained to be characterized globally^25^. Here, we estimated that 41% (73/176) of the pharyngeal-muscle-specific and 53% (59/111) of the pan-cardiac markers are already expressed in multipotent progenitors, indicating that lineage-specific maintenance of primed genes is a major determinant of cell-type-specific transcriptomes in the cardiopharyngeal lineage. Nevertheless, 88% (3,504/3,982) of late-expressedtranscripts, were already detected in multipotent progenitors (Fig. 1f), showing that *de novo* cell-type-specific gene activation contributes significantly to cell-type-specific programs (Fisher’s exact test, *P* < 2.2 × 10^−16^ for both the pan-cardiac- and ASM-specific gene sets, respectively).

We further explored the molecular basis for progression through regulatory states. As a proof of concept, we first focused on the pharyngeal muscle trajectory, for which the key regulators Hand-r, Tbx1/10 and Ebf have been characterized^24,25,28,31,35^ (Extended Data Figure 4). The first two regulatory states corresponded to successive generations of multipotent cardiopharyngeal progenitors (aka TVCs and STVCs, Fig. 1a, Extended Data Fig. 3g-i), confirming that asymmetric cell divisions provide the biological basis for these first transitions. To our surprise, the majority of newborn pharyngeal muscle precursors isolated from 16 hpf larvae clustered with multipotent progenitors isolated from 14hpf embryos (aka STVCs, Fig. 1a, Extended Data Fig. 3g-i), although they already expressed *Ebf* (Extended Data Fig. 4a), as previously observed by FISH^24,31^. This indicates that, although newborn pharyngeal muscle progenitors already express a key determinant, their transcriptome remains similar to their multipotent mother cells. Indeed, the pharyngeal muscle transcriptome is progressively remodelled as cells transition through successive states, involving both downregulation of primed cardiac markers and ‘*de novo*’ activation of pharyngeal muscle markers (Extended Data Fig. 4a-c, h-j). Moreover, systematic comparison with expression profiles following perturbations of Ebf function^25^ indicated that candidate Ebf target genes, including *Myogenic regulatory factor* (Mrf) (the *MyoD/Myf5* homolog), and *Myosin heavy chain 3* (*Mhc3*), are activated at later timepoints, consistent with Ebf-dependent transitions to committed pharyngeal muscle states^31^ (Extended Data Fig. 4a, d, e-g, k).

### Termination of FGF-MAPK signaling launches a pan-cardiac program for heart identity

Next we investigated gene expression changes underlying state transitions during cardiac specification. We focused on the first heart trajectory, which provided the largest pseudo-temporal range, to characterize the pan-cardiac program. Activation of ‘*de novo*’ pan-cardiac markers and down-regulation of primed pharyngeal muscle and multipotent-specific markers accounted for most gene expression changes (Fig. 2a-b, Extended Data Fig. 5a-d). These coordinated gene expression changes explained major transitions along the cardiac trajectories. For example, we used logistic regression to determine that *Slit1/2/3* is activated at the transition between the multipotent and FHP1 states, which we confirmed by FISH (Fig. 1d, Extended Data Fig. 1g). We identified a principal component (PC1), which correlated highly with pseudotime (PCC=0.96; Extended Data Fig. 5e), and used the PC1-loading of each gene to estimate the relative contribution of each class of markers to discrete regulatory states (Fig. 2c). This suggested that the multipotent state is primarily determined by the combined expression of TVC-specific genes, and primed cardiac and pharyngeal muscle markers. The TVC-to-FHP1 transition is marked by a sharp decline in TVC-specific gene expression, accompanied by down-regulation of primed ASM genes, upregulation of primed pan-cardiac genes, and activation of *de novo*-expressed pan-cardiac genes. In this regard, the FHP1 state may be considered a “transition state” between a multipotent TVC state, and the FHP2 state^36^. The latter is defined by the virtual absence of TVC-specific and primed ASM-specific transcripts, and high levels of both primed and *de novo*-expressed pan-cardiac markers, thus probably corresponding to a heart-specific state, whereas activation of cell-type/lineage-specific genes underlie the FHP2-to-FHP3 transition, and their expression helps define the first-lineage-specific state, FHP3, as is the case for *Mmp21* (Fig. 2d, Extended Data Fig. 5f).

**Figure 2.**
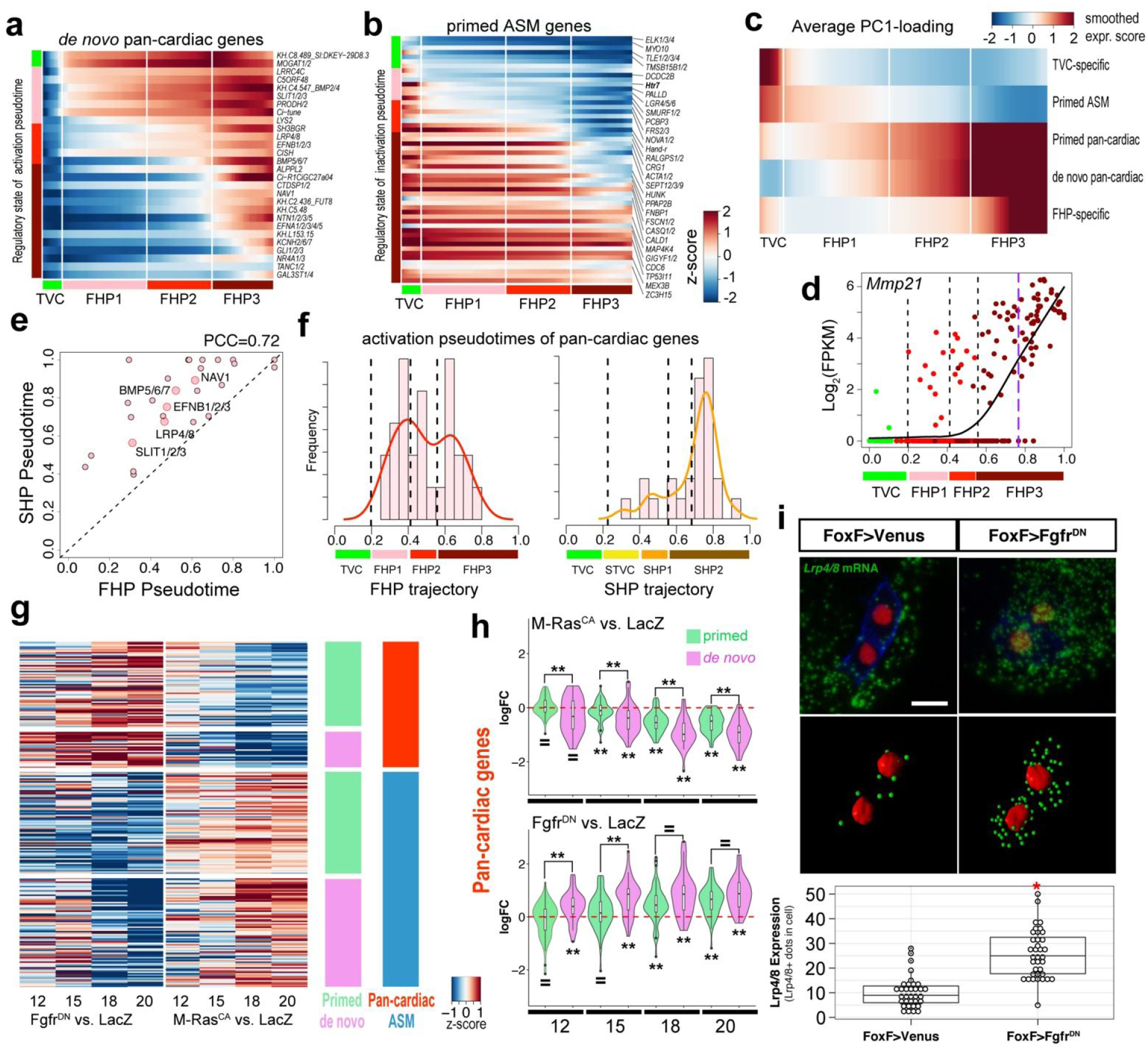
Characterization of the pan-cardiac gene expression program. (**a**) Heatmap of smoothed gene expression showing the successive activations of *de novo*-expressed pan-cardiac genes. (**b**) Heatmap of smoothed gene expression showing progressive depletion of primed ASM genes. For (a) and (b), white vertical lines indicate predicted regulatory states and color bar indicates relative expression level. (**c**) Average PC1-loading scores per indicated gene category, mapped onto the FHP trajectory. (**d**) Pseudotemporal expression profiles of *Mmp21* along the FHP trajectory. X-axis: normalized pseudotime. Black dashed lines: transitions between regulatory states; purple dashed line: calculated activation pseudotime. Dots colors refer to the sample of origin as in Fig. 1c. (**e**) Activation pseudotimes for *de novo*-expressed pan-cardiac genes along the FHP and SHP pseudotime axes. PCC: Pearson’s Correlation Coefficient. (**f**) Proportions of *de novo* pan-cardiac genes with calculated activation pseudotime in binned pseudotime windows along FHP and SHP trajectory respectively. (**g**) Log fold changes of primed/*de novo* ASM and pan-cardiac genes in Fgfr^DN^ vs. LacZ and M-Ras^CA^ vs. LacZ pairwise comparisons from FACS-purified cardiopharyngeal lineage cells obtained from 12, 15, 18 and 20 hpf larvae.hpf: hours post-fertilization. (**h**) Distributions of log_2_ fold changes in indicated conditions and time points relative to LacZ controls, and parsed by primed or *de novo*-expressed pan-cardiac genes. Summary statistics: results of t-test for significant difference from 0 are indicated below violin plots, results for KS tests for significant differences between “Primed” and “De novo” gene sets in each condition are indicated above violin plots. “=”, no difference, “**”, P-value <0.01. Details in Extended Data Table 4. (**i**) FGF/MAPK inhibition induces precocious expression of *Lrp4/8* in multipotent cardiopharyngeal progenitors (TVCs). *Lrp4/8* mRNAs (green) visualized by FISH. Anti beta-galactosidase antibody (red) marks TVC nuclei expressing *Mesp>nls::LacZ*. Mesp>hCD4::mCherry, revealed by anti mCherry antibody (blue), marks cell membranes. Anterior to the left. Scale bar, 10 μm. Imaris-processed images show *Lrp4/8* expression in the TVCs. Box plots represent the distributions of numbers of *Lrp4/8*+ dots per cell in indicated conditions. Bars in the box indicate the median value. * p<0.05 (Student’s t-test).

A true pan-cardiac program should unfold following similar dynamics in the first and second heart lineage, reflecting shared regulatory logic. Accordingly, we observed a striking agreement between the ordered activation pattern of individual genes along each trajectory (Fig. 2e), suggesting a remarkably conserved developmental program. Notably, the onset of each gene was consistently delayed in the second heart trajectory, starting with the STVC-to-SHP1 transition, as second heart precursors are born from a second generation of multipotent progenitors, ∼2h later than first heart precursors (Fig. 1a, 2e, f). Therefore, the *de novo* pan-cardiac program is tightly regulated and deployed in a reproducible cascade, whose onset is independently induced in the first and second heart lineages as they arise from multipotent progenitors.

We then sought to identify regulatory switches triggering the full pan-cardiac program in heart lineages. The FGF-MAPK signaling pathway is active and maintained specifically in multipotent cardiopharyngeal progenitors (TVCs and STVCs), and in early pharyngeal muscle precursors (ASMF), where it promotes the expression of *Hand-r*, *Tbx1/10*, and *Ebf.* By contrast, signaling is terminated in newborn first and second heart precursors^31^. We integrated sc- and bulk RNA-seq performed on FACS-purified cardiopharyngeal lineage cells following defined perturbations, and determined that FGF-MAPK signaling opposed pan-cardiac gene expression, while promoting the pharyngeal muscle program in swimming larvae (18 and 20 hpf, Fig. 2g-h, Extended Data Fig. 5j-m, Extended Data Table 4). At earlier stages (12 and 15 hpf), FGF-MAPK perturbations generally did not affect the expression of primed pan-cardiac genes in multipotent progenitors, whereas *de novo*-expressed genes were upregulated upon signaling inhibition by Fgfr^DN^ misexpression (Fig. 2g-h, Extended Data Fig. 5h-i,n-q; Extended Data Table 4). FISH assays further demonstrated that the *de novo*-expressed pan-cardiac marker *Lrp4/8* was upregulated in TVCs upon misexpression of Fgfr^DN^ (Fig. 2i). These analyses indicated that heart-lineage-specific termination of FGF-MAPK signaling permits the activation of *de novo*-expressed pan-cardiac genes, and subsequent heart fate specification, whereas ongoing FGF-MAPK signaling in cardiopharyngeal progenitors promotes multipotency both by maintaining the primed pharyngeal muscle program and by inhibiting the full deployment of the heart-specific program (Extended Data Fig. 12).

### Differences between cardiac lineages foster cellular diversity in the beating heart

A shared pan-cardiac gene program progressively defines the heart identity, but distinct precursors nevertheless clustered separately, revealing significant differences between the first and second heart lineages (Fig. 1b, Extended Data Fig. 1b; Extended Data Table 2). We mined the second heart trajectory to explore the development and evolution of the vertebrate second heart field (SHF), since the ascidian and vertebrate SHFs share regulatory inputs from *Nk4/Nkx2-5* and *Tbx1/10* orthologs^11,14,24,37–41^. Examining our list of markers distinguishing second and first heart precursors, we identified the *dachshund* homolog *Dach* as the only known transcription regulator^42–45^ (Extended Data Fig. 1f, 6a; Extended Data Table 2), and its upregulation as cells transitioned from a multipotent state suggested a role in specifying the second cardiac identity (Fig. 3a).

**Figure 3.**
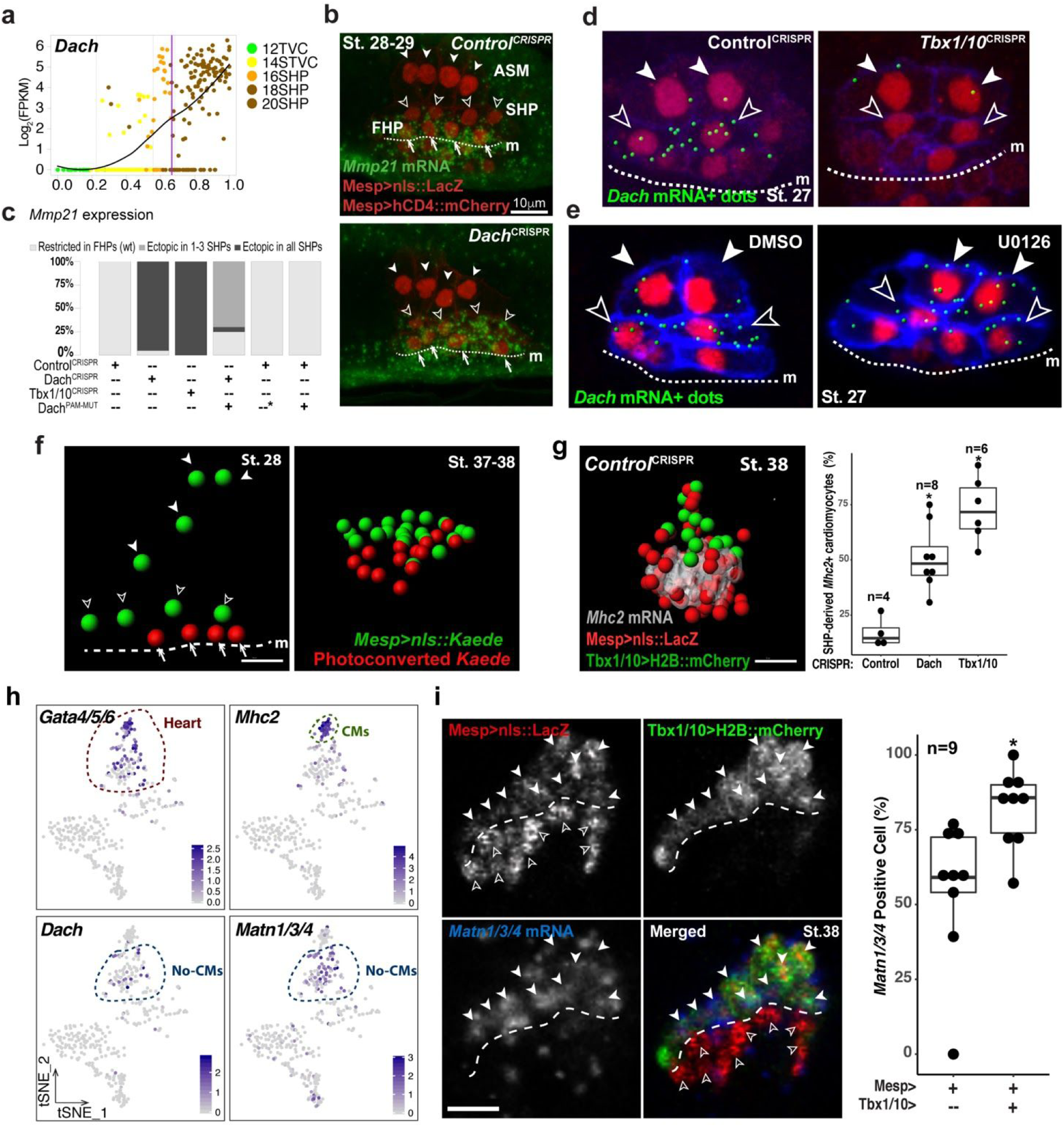
Transcriptional regulation of SHP fate specification. (**a**) *Dach* pseudotemporal expression pattern on the SHP trajectory. Black dashed lines: transitions between SHP regulatory states defined in Ext. Data Fig. 3h,j; purple dashed line: predicted induction time. (**b**) CRISPR/Cas9-mediated mutagenesis of *Dach* causes ectopic expression of the FHP-specific gene *Mmp21* in the SHPs. *Mmp21* mRNAs (green) detected by FISH. *Mesp>nls::LacZ*, revealed by anti beta-galactosidase antibody (red), marks the nuclei of cardiopharyngeal lineage cells. *Mesp>hCD4::mCherry*, revealed by anti-mCherry antibody (red), marks cell membranes. Open arrowheads, ASMPs; solid arrowheads, SHPs; arrows, FHPs. Dotted line: ventral midline. Anterior to the left. Scale bar, 10 μm. (**c**) Proportions of larvae showing indicated phenotypes in each experimental condition. Wt, wild-type, indicates *Mmp21* expression restricted to the FHPs. Dach^PAMmut^, Dach cDNA with wobble base mutation in the PAMs, driven by TVC-specific *Foxf* enhancer, renders the rescue construct (*Foxf>Dach^PAMmut^*) resistant to CRISPR. *, The Dach^PAMmut^ control was electroporated with a neutral *Foxf>Venus* construct. (**d**) *Tbx1/10* is required for *Dach* expression in SHP. B7.5-lineage-specific *Dach* expression visualized by FISH and shown as segmented *Dach*+ green dots (see Ext. Data Fig. 6 for raw images). *Mesp>nls::LacZ*, revealed with an anti beta-galactosidase antibody (red), labels nuclei of B7.5 lineage cells. *Mesp>hCD4::mCherry*, revealed by anti mCherry antibody (blue), marks cell membrane. Experiment performed in biological replicates. For each replicate, confocal stacks were acquired for 10 larvae in each condition. None of the 20 Tbx1/10^CRISPR^ larvae showed *Dach* expression in SHPs. Solid arrowheads, ASMFs, open arrowheads, SHPs. Anterior to the left. Scale bar, 10 μm. (**e**) FGF-MAPK signaling negatively regulates *Dach* expression in *Tbx1/10*+ ASMFs. Representative Imaris processed confocal stacks showing *Dach* expression in 18.5hpf (St. 27) larvae, following 3.5 hours treatments with DMSO (control) or U0126 (MEK inhibitor). Blocking MEK activity causes ectopic *Dach* expression in the ASMFs (solid arrowheads), in addition to its endogenous expression in the SHPs (open arrowheads) in 10/10 embryos analyzed. (**f**) Photoconversion and lineage tracing of TVC progeny. Segmented nuclei of live B7.5 lineage cells labelled with *Mesp>nls::Kaede::nls* (green). Nuclear Kaede photoconverted to red specifically in the FHPs in a 16hpf (St.25) larva (see Ext. Data Fig. 7 for raw images). The same animal is shown at 65hpf (St.37-38). Open arrowheads, ASMPs; solid arrowheads, SHPs; arrows FHPs; Dotted line: ventral midline. Anterior to the left. Scale bar, 10 μm. (**g**) *Dach* and *Tbx1/10* activities inhibit the production of *Mhc2*+ cardiomyocytes from the second heart lineage. Rendered segmented signals are shown (see Methods). Grey: *Mhc2* mRNA FISH. SHP-derived cells are labelled with the Tbx1/10 reporter, *3XT12>H2B::mCherry* revealed by an anti-mCherry antibody (green), B7.5 lineage cells are labelled with *Mesp>nls::LacZ* (red). Proportions of *Mhc2*+ cells among the 3xT12>H2B::mCherry+ SHP-derived cells in juvenile hearts in indicated experimental conditions. Scale bar, 10µm. (**h**) t-SNE plots of *Ciona* scRNA-seq data acquired in FACS-purified cardiopharyngeal lineage cells from ∼72hpf juveniles (St. 38), with expression pattern of *Gata4/5/6, Mhc2, Dach* and *Matn1/3/4* (see Ext. Data fig. 7f,g for clusters and top markers). The red dash line indicates two *Gata4/5/6*+ heart cell clusters. The green dash line indicates the cluster of *Mhc2*+ cardiomyocytes (CM). The blue dash line highlights the cluster of *Dach*+ and *Matn1/3/4*+ non-cardiomyocyte (No-CM) cells. (**i**) *Matn1/3/4* is enriched in the SHP-derived *Tbx1/10(3xT12)>H2B::mCherry*+ cells (green). *Matn1/3/4* mRNAs (Blue) are visualized by FISH. *Mesp>nls::LacZ*, revealed by anti beta-galactosidase antibody (red), marks the nuclei of cardiopharyngeal lineage cells. Solid arrowheads (outer-layer): mCherry+/beta-galactosidase+ SHP-derived cells positive for *Matn1/3/4* mRNA, open arrowheads (inner layer): mCherry-/beta-galactosidase+ cells negative for *Matn1/3/4* expression. Scale bar, 5µm. The images are X-Y cross sections of the juvenile heart. Corresponding boxplots with the proportions of *Matn1/3/4*+ cells among the Mesp>nls::LacZ+;3xT12>H2B::mCherry+ SHP-derived cells in juvenile hearts. There are significantly more *Matn1/3/4*+ cells among SHP-derived cells, compared to the Mesp>nls::LacZ+;Tbx1/10(3xT12)>H2B::mCherry-cells derived primarily from the first heart lineage. Median values are indicated in the boxplots. * p<0.05 (Student’s t-test)

*Dach1 and -2* have not been previously implicated in mammalian SHF development, but they belong to the conserved “retinal network”^46^, which comprises homologs of Six and Eya transcription factors that contribute to cardiopharyngeal development in the mouse^47,48^. Lineage-specific CRISPR/Cas9-mediated loss-of-function^49,50^, followed by gene expression assays, indicated that *Dach* is neither required for activation of the pharyngeal muscle marker *Ebf*, nor for its exclusion from the SHPs (Extended Data Fig. 6b). By contrast, loss of Dach function caused ectopic expression of the FHP marker *Mmp21* in the second heart precursors (Fig. 3b,c, Extended Data Fig. 6c). A CRISPR-resistant *Dach^PAMmut^* cDNA rescued the ectopic *Mmp21* expression in the second heart lineage, but did not abolish endogenous *Mmp21* expression in the first heart lineage (Fig. 3c, Extended Data Fig. 6c). Dach is thus both a marker and a key regulator of second heart lineage specification, which is required, but not sufficient, to prevent activation of the first-lineage-specific marker *Mmp21*.

Since the second heart lineage emerges from *Tbx1/10*+ multipotent progenitors^24^, we tested whether Tbx1/10 regulates *Dach* expression. CRISPR/Cas9-mediated lineage-specific loss of Tbx1/10 function^35^ inhibited *Dach* expression (Fig. 3d, Extended Data Fig. 6d), and caused ectopic activation of *Mmp21* (Fig. 3c, Extended Data Fig. 6b), indicating that Tbx1/10 promotes second heart lineage specification, in part by regulating *Dach* activation, in addition to its role in pharyngeal myogenesis.

Indeed, Tbx1/10 is also necessary in parallel to FGF-MAPK activity to activate *Ebf* and promote the pharyngeal muscle program^24,31,35^, in a manner similar to Tbx1 function in vertebrate branchiomeric myogenesis^51,52^. To explore the mechanism distinguishing between Tbx1/10 dual functions, we used the MEK/MAPKK inhibitor U0126, which inhibits *Ebf* expression^31^, and caused ectopic *Dach* activation in the lateral-most cardiopharyngeal cells that normally form *Ebf*+ pharyngeal muscle precursors (Fig. 3e). Moreover, Tbx1/10 misexpression and MEK/MAPKK inhibition synergized to cause precocious and ectopic *Dach* activation in cardiopharyngeal progenitors (Extended Data Fig. 6e). Taken together, these data indicate that termination of FGF-MAPK signaling in *Tbx1/10*+ cardiopharyngeal progenitors suffice to activate *Dach* expression and promote the second heart lineage identity, and demonstrate how distinct signaling environments can promote divergent regulatory programs in concert with Tbx1/10 expression.

First and second heart precursors share a common pan-cardiac signature, but initial molecular differences open the possibility that each lineage contributes differently to cardiogenesis. The beating *Ciona* heart is demonstrably simpler than its vertebrate counterpart; yet, diverse cell types form its single U-shaped compartment^53^. In post-metamorphic juveniles, the heart already beats, and double labeling with a *Mesp>nls::lacZ* reporter and the cardiac-specific *myosin heavy chain 2* (*Mhc2/Myh6*) marker showed that beta-galactosidase+; *Mhc2/Myh6*-cells surround *Mhc2/Myh6*+ cardiomyocytes^24,28^. Lineage tracing using the photoconvertible reporter Kaede^25^ indicated that first and second heart precursors derivatives remain within largely separate domains in juvenile hearts (Fig. 3f, Extended Data Fig. 7a). Specifically, first-lineage-derived cells form the inner layer of *Mhc2+* cardiomyocytes, whereas second-lineage-derived cells contribute to the outer layer of *Mhc2-* cells (Fig. 3g, Extended Data Fig. 7c, Extended Data Movie S1). Triple labeling and cell quantification using pan-cardiopharyngeal and second-heart-lineage-specific reporters, and the *Mhc2* probe, indicated that most *Mhc2*+ cardiomyocytes were located in the first-lineage-derived inner layer, whereas only ∼17% of second-lineage-derived cells express *Mhc2* in control juveniles (Fig. 3g). Thus, the first and second heart lineages contribute primarily *Mhc2/Myh6*+ cardiomyocytes and *Mhc2/Myh6*-cells to the beating juvenile heart, respectively.

To further characterize cellular diversity in the juvenile heart, we performed scRNA-seq on 386 FACS-purified cardiopharyngeal lineage cells dissociated from stage 38 juveniles, and identified clusters corresponding to pharyngeal muscle and cardiac lineages, including *Mhc2/Myh6*+ cardiomyocytes and an *Mhc2/Myh6*-population that expressed the second heart lineage markers *Dach* and *Matn* (Fig. 3h, Extended Data Fig. 7f-g). Triple labeling with *Mesp* and *Tbx1/10* reporters indicated that these *Dach+;Matn+* cells derived principally from the second heart precursors and formed the outer layer of the juvenile heart (Fig. 3i, Extended Data Fig. 7d), suggesting that cellular diversity in the beating heart emerges from the initial segregation of the first and second heart lineages. Consistent with this hypothesis, CRISPR/Cas9-mediated loss of Dach and Tbx1/10 early functions increased the proportions of *Mhc2/Myh6*+ cells in the SHP progeny (Fig. 3g, Extended Data Fig. 7b), demonstrating that early inhibition of the *Mmp21*+ first-lineage-specific program by the Tbx1/10-Dach pathway limits the potential of second heart lineage derivatives to differentiate into *Mhc2/Myh6*+ cardiomyocytes during organogenesis.

### Conserved cardiopharyngeal transcriptional signatures in chordates

Finally, we asked whether molecular features of cardiopharyngeal development are shared between Ciona and vertebrates. Recent scRNA-seq analysis of early mesodermal lineages in mice identified a population of pharyngeal mesoderm marked by high levels of both *Tbx1* and *Dach1* expression^54^ (Fig. 4a). Multicolor immunohistochemical staining revealed that Dach1 expression starts broadly in the pharyngeal mesoderm, and becomes restricted to second heart field cells in the dorsal pericardial wall, and to a defined population of outflow tract cells, both of which also express Isl1 (Fig. 4b, Extended Data Fig. 8a-b). Dach1 expression was excluded from the Nkx2.5+ ventricle, and absent from the Isl1+ skeletal muscle progenitor cells in the core mesoderm of the first and second pharyngeal arches (Fig. 4b, Extended Data Fig. 8a-b), in a manner reminiscent of *Dach* exclusion from the pharyngeal muscles in *Ciona* (Fig. 3d-e, Extended Data Fig. 1f).

**Figure 4.**
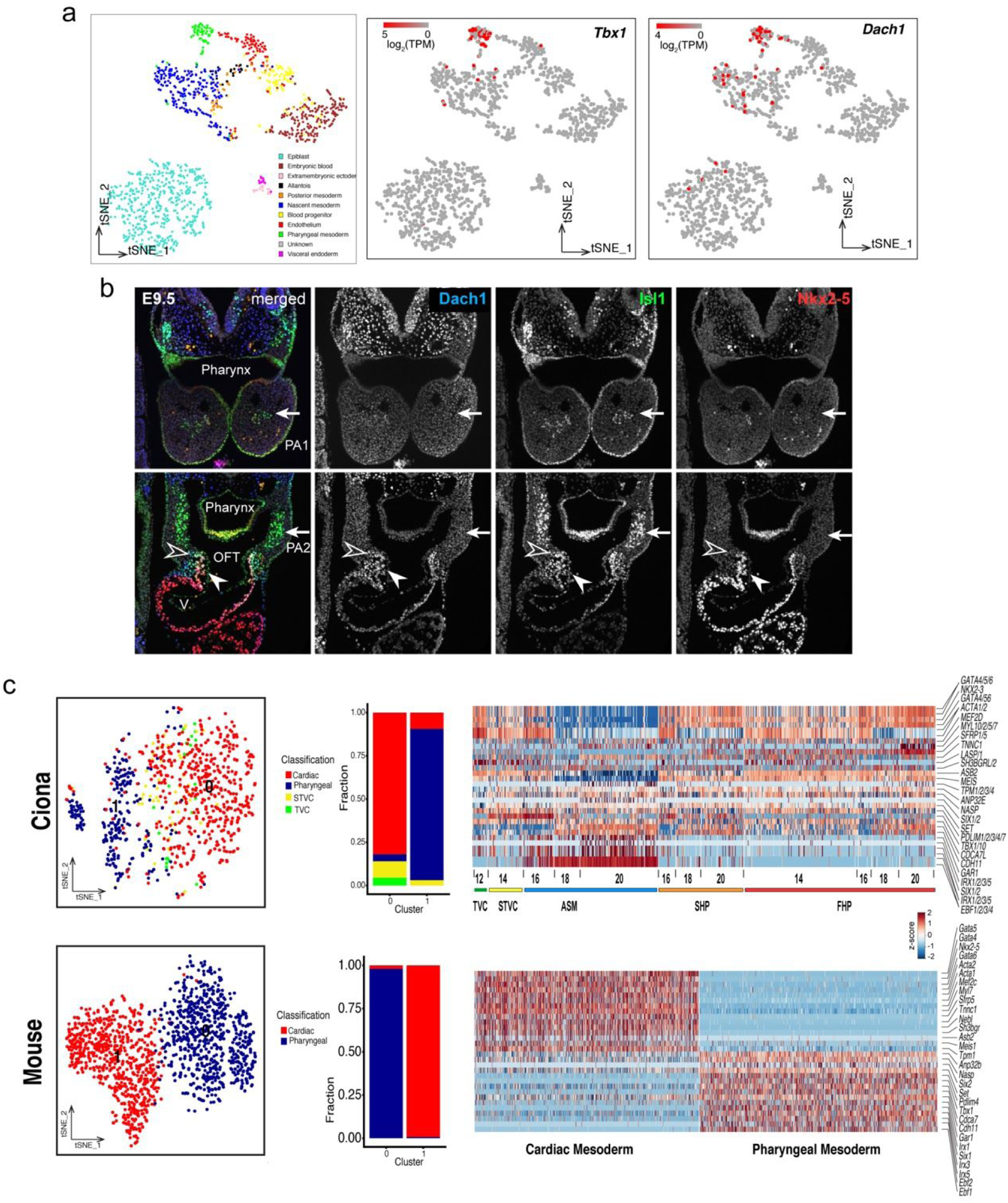
Conserved cardiopharyngeal programs in Ciona and in the Mouse. (**a**) t-SNE plots of mouse scRNA-seq data after Scialdone et al.^74^, with expression pattern of *Tbx1* and *Dach1*. Cluster identities are as determined in the original publication, with the pharyngeal mesoderm shown in green. (**b**) Expression patterns of Dach1, Islet1 and Nkx2.5 proteins in E9.5 mouse embryos. Arrows: Islet1+ head muscle progenitor cells in the mesodermal core of the first (PA1, top) and second (PA2, bottom) pharyngeal arches, showing absence of Dach1 and Nkx2.5 expression. Open arrowhead: Dach1+, Islet1+, Nkx2.5-second heart field cells in the dorsal pericardial wall, solid arrowhead: Triple Nkx2.5+, Dach1+, Islet1+ second heart field-derived cells in the outflow tract (OFT). Note the Nkx2.5+, Dach1-, Islet1-cells in the ventricle (V). (**c**) Aligned structure of Ciona and Mouse E8.25 cardiopharyngeal cells. tSNE plots showing the clustering of Ciona and Mouse E8.25 cardiopharyngeal cells, respectively, using conserved markers determined by canonical correlation (CC). Barplots indicate that original cell identities, defined in each species independently, as recovered in the clustering using conserved markers. Heatmaps show the single cell expression profiles for the top 30 conserved markers in each species, separately.

We extended the Ciona-to-mouse comparison of cardiopharyngeal transcriptomes using published scRNA-seq datasets^13,54,55^. Using canonical correlation analysis^56^, we identified genes that separated cardiac and pharyngeal mesoderm cells in both Ciona and E8.25 mouse embryos (Fig. 4c, Extended Data Table 5). We then used only the 30 best correlated genes to re-cluster scRNA-seq data independently from each species, and found that these markers sufficed to distinguish cardiac and pharyngeal muscle cells in either species, revealing a shared transcriptional program (Fig. 4c). We repeated this analysis using mouse datasets from earlier embryonic stages^13,54^, and consistently identified genes that separated cardiac and pharyngeal cells in both species, and were enriched in transcription factor- and DNA binding protein-coding genes (Extended Data Figs. 9a-b, 10a-b, 11). For instance, *Gata4* and *Ebf1* homologs were identified in all three comparisons as discriminating markers that separated cardiac and pharyngeal cells (Extended Data Fig. 10c, Extended Data Table 5). Overall, this analysis suggests that an evolutionary conserved transcriptional program, comprising homologs of *Ebf1*, *Gata4* and other regulatory genes, govern the heart vs. pharyngeal muscle fate choice in cardiopharyngeal mesoderm.

## DISCUSSION

Here, we present an extensive analysis of the transcriptome dynamics underlying early cardiopharyngeal development in a tractable chordate model. Using established clonal relationships to inform the reconstruction of developmental trajectories, we characterized essential features of the transcriptome dynamics underlying cardiopharyngeal multipotency and early fate specification. Multipotent cardiopharyngeal progenitors exhibit extensive multilineage transcriptional priming, although *de novo* gene activation significantly contributes to cell-type-specific transcriptomes, highlighting the importance of both transcriptional and post-transcriptional regulation in early heart vs. pharyngeal muscle fate choices.

Both first- and second-heart-lineage cells acquire a cardiac identity as they down-regulate multipotent progenitors markers and primed pharyngeal muscle genes, and deploy the full pan-cardiac program, entering this ‘transition state’^36^ upon termination of FGF-MAPK signaling. We propose that the dual functions of FGF-MAPK signaling, as observed during early cardiac specification in Ciona, are conserved in vertebrates considering that FGF-MAPK inputs are necessary to induce multipotent progenitors^57–59^, whereas signal termination is required for subsequent commitment to a heart fate and cardiomyocyte differentiation^60–65^.

Following commitment to a cardiac identity, first heart progenitors transition to an *Mmp21*+ state that precedes differentiation into *Mhc2/Myh6*+ cardiomyocytes in the beating heart. By contrast, second heart progenitors activate *Dach* in response to Tbx1/10 inputs inherited from their distinct multipotent mother cells. This Tbx1/10-Dach pathway inhibits *Mmp21* expression and the *Mhc2/Myh6*+ cardiomyocyte potential to foster heart cell diversity. As is the case in vertebrates^41,51,52,66–69^, Tbx1/10 plays a dual roles in branchiomeric/pharyngeal myogenesis^24,31,35^ and second heart lineage development, thus acting as a *bona fide* regulator of cardiopharyngeal multipotency. Moreover, the first and second heart precursors share a common cardiac identity but differ because they emerge from successive multipotent progenitors before and after the onset of *Tbx1/10* expression. Cellular diversity in the Ciona heart thus emerges by temporal patterning, as is the case for neuronal and muscle fates in *Drosophila*^70,71^. The proposal that first and second heart lineages form a coherent cardiac developmental unit, whereas distinct lineages contribute to cellular diversity within the heart, reconciles different views about the significance of the second heart field.

Finally, by leveraging recent computational methods for cross-species comparisons of single cell RNA-seq datasets, we identified shared markers of cardiopharyngeal regulatory states, while highlighting differences in expression dynamics, as seen for *Dach* homologs, thus illustrating the plasticity of gene regulatory networks controlling conserved developmental programs.

## Supporting information

ExtData_Movie1

ExtData_movie2

ExtData_Table1

ExtData_Table2

ExtData_Table3

ExtData_Table4

ExtData_Table5

ExtData_Table6

Markdown_Clustering

Markdown_Preprocessing

Markdown_Trajectory

## ACKNOWLEDGEMENTS

We are grateful to Florian Razy-Krajka for discussions and sharing reagents prior to publication. We thank Ashley Powers for help processing the single cell samples in the early phase of this study, Christopher Hafemeister and Andrew Butler for discussion and help on computational analyses. This project was funded by NIH/NHLBI R01 award HL108643 to L.C., trans-Atlantic network of excellence award 15CVD01 from the Leducq Foundation to R.K. and L.C., and an NIH New Innovator Award (DP2-HG-009623) to R.S.

## AUTHORS’ CONTRIBUTIONS

W.W. performed the *Ciona* experiments. E.J. performed the mouse experiments. X.N., T.S., and R.S. performed computational analyses. W.M.M., W.W., and R.S. performed the single cell RNA-seq experiments. W.W., X.N., R.K., R.S., and L.C. designed the experiments and analyses. W.W., X.N., R.S. and L.C. wrote the paper.

