## Supplementary Materials for "A single cell transcriptional roadmap for cardiopharyngeal fate diversification"

### CONTACT FOR REAGENT AND RESOURCE SHARING

Further information and requests for reagents may be directed to Lionel Christiaen and Rahul Satija.

### METHODS

#### Animals

All animal care and experiments were carried out in accord with current NIH guidelines. *Ciona robusta* adults were purchased from M-Rep (San Diego, CA), maintained in artificial seawater with constant illumination, and used for experiment within one week after arrival. CD1 mice were crossed to generate embryos that were staged using the day of the copulation plug as embryonic day (E) 0.5.

#### Isolation of gametes, fertilization, dechorionation, electroporation, and development

*Ciona* eggs and sperm were collected from at least two individual adult animals and kept separately in filtered artificial seawater (FASW) until fertilization. Eggs were mixed with activated sperm and incubated in FASW at room temperature for 5 min. Chorion and surrounding follicle cells were chemically removed with a pronase solution (FASW, 7.5mg/L sodium thioglycolate, 0.05% pronase, 0.042N NaOH), as described[1]. Fertilized and dechorionated eggs were electroporated as described[2], and cultured in FASW in agarose coated plastic Petri dishes at 18°C. The amount of fluorescent reporter DNA (*Mesp>nls::lacZ*, *Mesp>hCD4::mCherry*, *Mesp>tagRFP*, *MyoD905>EGFP* and *Hand-r>tagBFP*) for electroporation was typically 50 µg, except for *3XT12>H2B::mCherry* is 30 µg. In order to increase the survival rate of juveniles, healthy and fluorescent positive larvae were selected at St. 26-27 using Leica M205 FA fluorescence stereo microscopes and transferred to uncoated Petri dishes with fresh FASW. The larvae were cultured at 18 °C until harvest.

#### Dissociation and FACS

Sample dissociation and FACS were performed essentially as described[3–5]. Embryos and larvae were harvested at 12, 14, 16, 18 and 20hpf in 5ml borosilicate glass tubes (Fisher Scientific, Waltham, MA. Cat.No. 14-961-26) and washed with 2ml calcium- and magnesium-free artificial seawater (CMF-ASW: 449 mM NaCl, 33 mM $Na_2SO4$, 9 mM KCl, 2.15 mM $NaHCO_3$, 10 mM Tris-Cl pH 8.2, 2.5 mM EGTA). The tunic of juveniles was peeled off using BD PrecisionGlide™ Needles (REF 305115) under the dissecting microscope to facilitate the dissociation. Embryos and larvae were dissociated in 2ml 0.2% trypsin (w/v, Sigma, T- 4799) ASW by pipetting with glass Pasteur pipettes. The dissociation was stopped by adding 2ml filtered ice cold 0.05% BSA CMF-ASW. The dissociated cells were passed through 40µm cell-strainer and collected in 5ml polystyrene round-bottom tube (Corning Life Sciences, Oneonta, New York. REF 352235). Cells were collected by centrifugation at 800g for 3 min at 4°C, followed by two washes with ice cold 0.05% BSA CMF-ASW. Cell suspensions were filtered again through a 40µm cell-strainer and stored on ice. Following dissociation, cell suspensions were used for sorting within 1 hour.

B7.5 lineage cells were labeled by *Mesp>tagRFP* reporter. Contaminating B-line mesenchyme cells were counter-selected using *MyoD905>EGFP* as described[4–6]. The TVC-specific *Hand-r>tagBFP* reporter was used in a 3-color FACS scheme for positive co-selection of TVC-derived cells, in order to minimize the effects of mosaicism. Dissociated cell were loaded in a BD FACS Aria™ cell sorter. 488 nm laser, FITC filter was used for EGFP; 407 nm laser, 561 nm laser, DsRed filter was used for tagRFP and Pacific BlueTM filter was used for tagBFP. The nozzle size was 100 μm. tagRFP +, tagBFP + and EGFP – cells were collected for downstream RNA sequencing analysis.

#### Fluorescent *In Situ* Hybridization-Immunohistochemistry (FISH-IHC) in *Ciona* Larvae

FISH-IHC were performed essentially as described[4,7,8]. Embryos were harvested and fixed at desired developmental stages for 2h in 4% MEM-PFA and stored in 75% ethanol at −20°C. Antisense RNA probes were synthesized using either Gateway gene collections or amplified fragments of desired genes as templates (Extended Data Table 3). *In vitro* antisense RNA synthesis was performed using T7 RNA Polymerase (Roche, Cat. No. 10881767001) and DIG RNA Labeling Mix (Roche, Cat. No. 11277073910). Anti-Digoxigenin-POD Fab fragment (Roche, IN) was first used to detect the hybridized probes, then the signal were revealed using the Tyramide Signal Amplification (TSA) with Fluorescein TSA Plus Evaluation Kits (Perkin Elmer, MA). Anti–β-galactosidase monoclonal mouse antibody (Promega) was co-incubated with anti-mCherry polyclonal rabbit antibody (Bio Vision, Cat. No. 5993-100) for immunodetection of *Mesp>nls::lacZ* and *Mesp>hCD4::mCherry* products respectively. Goat anti-mouse secondary antibodies coupled with AlexaFluor-555 and AlexaFluor-633 were used to detect β-galactosidase-bound mouse antibodies and mCherry-bound rabbit antibodies after the TSA reaction. FISH samples were mounted in ProLong® Gold Antifade Mountant (ThermoFisher Scientific, Waltham, MA. Catalog number P36930).

#### Multicolor Immunohistochemical Staining in Mouse Embryo

After dissection, embryos were fixed for 30 minutes (E8.5 and E9.5) to 1 hour (E7.5 in decidua) in 4% paraformaldehyde, dehydrated and embedded in paraffin prior to sectioning at 10μm. Immunofluorescence was performed using standard protocols. Briefly, after rehydration, sections were treated for 15 minutes with antigen unmasking solution (H-3300, Vector laboratories). Slides were washed twice in 1xPBS Tween (0.05%) and incubated for one hour in TNB buffer (0.1M Tris-HCl pH 7.5, 0.15M NaCl, 0.5% Blocking reagent (Roche 11096176001)). Sections were incubated with primary antibodies for 36 hours in TNB using the following dilutions: Dach1 (1/100, Proteintech 10914-1-AP), Nkx2-5 (1/100, Santa Cruz sc-8697), Islet1 (1/100, DSHB 39.4D5 and 40.2D6). After three 5 minutes washes in 1xPBS Tween (0.05%) sections were incubated with secondary antibodies for one hour using Alexa 488, 568 and 647 (1/500, Invitrogen). Sections were counterstained with Hoechst (Sigma 33258), mounted using Fluoromount (Southern Biotech 0100-001) and imaged using a Zeiss Axio Imager Z1 with an Apotome module.

#### CRISPR/Cas9-Mediated Gene Knock-Down

Two guide RNAs targeting the third and the fifth exon of *Dach* with Fusi scores (http://crispor.tefor.net) 63 (sgDach1: AAAAGATTAAGCATCGCCC) and 64 (sgDach2: GAGCATTGCCATTGACGTG), respectively, were designed to mutagenize the *Dach* locus in the B7.5 lineage using CRISPR/Cas9 as described[9]. Two guide RNAs described by Tolkin et al[10]. (sgTbx1.6 TGCGGCTTCGGCTCCGTGG; sgTbx1.8 AACGAAAGATTGGTGGCCG), were used to mutagenize Tbx1/10 coding region. The efficiency of guide RNAs were evaluated using the peakshift method, as described[9]. Guide RNAs were expressed using the *Ciona robusta U6* promoter[11]. For each gene, two guide RNAs were used in combination with 25 *μ*g of each expression plasmid. 30 *μ*g of *Mesp>nls::Cas9::nls* plasmid were co-electroporated with guide RNA expression plasmids for B7.5 lineage-specific CRISPR/Cas9-mediated mutagenesis. Rescue of the *Dach* loss-of-function was achieved by TVC-specific overexpression of $Dach^{PAMmut}$ driven by a *Foxf* enhancer[12]. Point mutations (G303A and C852A) were introduced to the PAMs of sgDach1 and sgDach2 using an optimized QuikChange™ Site-directed Mutagenesis protocol. Two pairs of mutagenesis primers were designed using PrimerX website (http://www.bioinformatics.org/primerx/cgi-bin/DNA_1.cgi) as sgDACH_1_G303A_F: GATTAAGCATCGCCCCAGTCGTGTGCAACGTTG; sgDACH_1_G303A_R: CAACGTTGCACACGACTGGGGCGATGCTTAATC and sgDACH_2_C852A_F: CGTCGGGAATTCCACCCACGTCAATG; sgDACH_2_C852A_R: CATTGACGTGGGTGGAATTCCCGACG. The PCR mixture was prepared as 20 µl 5X HF buffer, 71 µl H2O, 3 µl 10 mM dNTP, 1.75 µl WT Dach plasmid at 15ng/µl, 2 µl 125 ng/µl top mutagenesis primer, 2 µl 125 ng/µl bottom mutagenesis primer, 1 µl Phusion® High-Fidelity DNA Polymerase (NEB M0530). PCR mixture were evenly distributed into 8 PCR tube-strip and PCR was performed with a denaturation at 95 °C for 4 min, followed by 18 cycles of (95 °C for 30 s, 50 to 72 °C (gradient) for 1 min, and 72 °C for 1 min) and a final extension at 72° for 5 min. The PCR products were pooled and 4 µl of DpnI was added directly to the tube, followed by the incubation at 37 °C for 2 hours. After the purification using QIAquick PCR Purification Kit (QIAGEN), the eluate are transformed into TOP10 cells. Successful mutagenesis was confirmed by sequencing.

#### Photoconversion and Lineage Tracing

Photoconversion and lineage tracing were performed as described[4]. Fertilized eggs were electroporated with 50 $\mu$g Mesp>nls:Kaede:nls to label the B7.5 lineage. Embryos were raised on agarose coated plastic Petri dishes in ASW at 18°C and transferred individually into Nunc™ MicroWell™ 96-Well Optical-Bottom plates (ThermoFisher Scientific, Waltham, MA. Supplier No. 164588) at 15hpf. Photoconversions were performed using the HC PL FLUOTAR 20×/0.50 objective on Leica Microsystems inverted TCS SP8 X confocal microscope, by shedding 405 nm UV light on ROI continuously for 2 min. Stack scanning of whole TVC lineage were documented at 16, 22.5, 40, 48 and 65 hpf.

#### Confocal Microscopy

Images were acquired with an inverted Leica TCS SP8 X confocal microscope, using HC PL APO 63×/1.30 objective. Z-stacks were acquired with 1 µm z-steps. Maximum projections were processed with maximum projection tools from the LEICA software LAS-AF.

#### Image Processing and Quantification

Confocal z stacks were processed using Imaris x64 8.4.1 (BitPlane). A region containing the TVC progeny was segmented first. For nuclei detection, the expected nucleus diameter was set at 2.5µm, Nucleus Threshold(Absolute Intensity) were calculated automatically by Imaris. The cell segmentation was carried out using “Detect Cell Boundary from Cell Membrane” function, the “Cell Smallest Diameter” was set as 5µm. The transcripts signal within the cell boundary was detected using “Vesicles Detection” function, the estimated diameter of dots was set as 1.44µm. To count the number of (Mhc2+, Matn+ or Dach+) cells in juvenile heart, a region of juvenile heart was segmented first, then the nuclei detection was performed as described above. A surface was created from the channel of FISH detention. Then the “Find Spots Close To Surface” function was used to define the nucleus with transcripts (the surface) closing to it using a 2µm threshold.

### Bulk RNA-seq Library Preparation, Sequencing and Analyses

200 to 800 cells were directly sorted in 100*μl* lysis buffer from the RNAqueous®-Micro Total RNA Isolation Kit (Ambion). For each condition, samples were obtained in biological duplicates. The total RNA extraction was performed following the manufacturer's instruction. The quality and quantity of total RNA was checked using Agilent RNA 6000 Pico Kit (Agilent) with Agilent 2100 Bioanalyzer. RNA samples with RNA Integrity Number (RIN)>8 were kept for downstream cDNA synthesis. 250-2000 pg of total RNA was loaded as template for cDNA synthesis using the SMART-Seq v4 Ultra Low Input RNA Kit (Clontech) with template switching technology. RNA-Seq Libraries were prepared and barcoded using Ovation® Ultralow System V2 1–16 (NuGen). Up to 6 barcoded samples were pooled in one lane of the flow cell and sequenced by Illumina Hi-Seq 2500. One direction and 50bp length reads were obtained from all the bulk RNA-seq libraries.

Sequencing reads were mapped to the *Ciona robusta* genome (Joined-scaffold (KH), http://ghost.zool.kyotou.ac.jp/datas/JoinedScaffold.zip) using TopHat 2.0.12[13,14] with parameter: --no-coverage-search. Cufflinks 2.2.0 [14] was used to calculate the Fragments Per Kilobase of transcript per Million mapped reads (FPKM). We used edgeR[15] to analyze differential gene expression in pairwise comparisons. Detailed summary statistics are provided in Ext. Data Table 4.

### Single Cell RNA-seq Library Preparation and Sequencing

Reverse transcription and cDNA amplification were carried out using modified Smart-seq2 protocol[16]. Single cells were sorted by FACS as described above into 96-well plates and collected in 3.4 µl RT buffer (0.5 µl 10µM 3' RT Primer ( 5' - AAG CAG TGG TAT CAA CGC AGA GTA C T30 VN - 3'), 0.5µl 10µM dNTP Mix, 0.5 µl 4U/µl RNase Inhibitor, 1 µl Maxima RT Buffer, 0,9 µl nuclease-free water) in each well. Plates were either stored at -80°C or processed immediately. Plates were incubated at 72°C for 3min and chilled on ice to denature the template RNA. 2µl RT reaction mixture (0.5µl 10um TSO primer (5′-AGACGTGTGCTCTTCCGATCTNNNNNrGrGrG-3′), 0.925µl 5M Betaine, 0.4 100mM MgCl2, 0.125µl 40U/ RNase inhibitor, 0.05µl 200 U/µl Maxima H Minus Reverse Transcriptase) were added to each well. Reverse transcription was carried out by incubating the plate at 42°C for 90 min, followed by 10 cycles of (50 °C for 2 min, 42 °C for 2 min) and heat inactivation at 70° for 15 min. 7µl PCR amplification mixture (0.25µl 10 µM PCR primer (5′AGACGTGTGCTCTTCCGATCT-3′), 6.25µl KAPA HIFI ReadyMix, 0.5µl nuclease-free water) were added to each well. PCR amplification was carried out with a denaturation at 98 °C for 3 min, followed by 21 cycles of (98 °C for 15 s, 67 °C for 20 s, and 72 °C for 6 min) and a final extension at 72° for 5 min. PCR products were purified by adding 10µl (0.8×) Agencourt AMPureXP SPRI beads (Beckman-Coulter) to each well, followed by 5min incubation and two washes with 100µl freshly prepared 70% Ethanol at room temperature. Purified cDNA were eluted in 20µl TE. The concentration of amplified cDNA was measured across the entire plate using Picogreen assays. The concentration of amplified cDNA was in a 0.5–2 ng/µl range. Fragment size distributions were checked for randomly selected wells with High-Sensitivity Bioanalyzer Chip (Agilent), the expected size average should be ~2 kb. For each sample, the amplified cDNA were normalized to a working concentration ranging from 0.1 to 0.2 ng/µl with TE buffer. 1.25 µl of diluted cDNA from each well were used for library preparation. Single cell libraries were prepared using the Nextera XT DNA Sample Kit (Illumina) according to manufacturer's instructions. After library amplification, 2.5µl from each well were pooled into a single 1.5-ml microcentrifuge tube, purified using Agencourt AMPure XP beads and eluted with 30µl TE buffer. 1µl purified library was used to measure the fragment size distribution using the

Agilent HS DNA BioAnalyzer chip and another 1µl of the purified library was loaded into Qubit fluorometer to estimate library concentration according to the manufacturer's instructions. Libraries were sequenced on an Illumina HiSeq 2500 sequencer to obtain paired-end 50bp reads.

### QUANTIFICATION AND STATISTICAL ANALYSIS

#### Read alignment and generation of gene expression matrix

For each demultiplexed bulk and single cell RNA-seq library, sequencing reads were mapped to the *Ciona robusta* genome (Joined-scaffold (KH), http://ghost.zool.kyotou.ac.jp/datas/JoinedScaffold.zip) using TopHat 2.0.12[13,14] with parameter: --no-coverage-search. Cufflinks 2.2.0 [14] was used to calculate the Fragments Per Kilobase of transcript per Million mapped reads (FPKM).

#### Preprocessing and batch effect removal

We adopted multiple quality control criteria to filter out low quality single cell transcriptomes. First, we only retained single cells that had more than 2,000 and less than 6,000 expressed genes, and genes that were detected in more than 3 cells. 1,182 out of 1,796 single cells and 14,864 out of 15,287 genes were retained. We used total reads and overall read mapping rates from TopHat output files to assess the quality of scRNA-seq. Cells with mapping rates less than 30% and total reads more than 2-million were removed (see Supplementary Note). 1,138 out of 1,182 cells passed the quality controls and were retained for downstream analyses.

Batch effects were identified by principal component analysis (PCA) using all detected genes. Principal components 2, 5 and 7 were dominated either by ribosomal genes or unannotated genes that showed strong expressions only in certain batches (Supplementary Note). These PC's were considered as batch effects created by sequencing and library preparation. For each gene *j*, its expression level $y_j$ was fitted by a linear mixed model of the total sum of latent batch effects ( $x_i$ ) and its real biological expression level ( $\varepsilon_j$ ) as the formula $y_j = \sum_i a_i x_i + \varepsilon_j$ , where $a_i$ denotes coefficient of batch effect $x_i$ . In our case, PCA rotation matrices of PC2, PC5 and PC7 served as batch effects and were regressed out by the above model (Supplementary Note).

Contaminating subpopulations were discovered upon clustering. 59 *Twist1*+ mesenchymal cells and 198 cells without previously identified lineage markers were detected in cluster 8, 9, 11 and 12 (Supplementary Note). These contaminating non-cardiopharyngeal lineage cells were removed before downstream analysis. 881 out of 1,138 single cells were retained for clustering and trajectory analysis.

#### Identification of variable genes and dimensional reduction analysis.

All scRNA-seq analyses were performed on each time point data individually. Downstream analysis followed the procedures of 'Seurat' R package v1.2 [17]; http://satijalab.org/seurat).

We first identified the set of genes that was most variable in 12, 14 and 20hpf single-cell data. We calculated the mean and dispersion (variance/mean) for each gene across all single cells, and placed genes into 20 bins based on their average expression. Within each bin, we then z-normalized the dispersion measure of all genes within the bin to identify genes whose expression values were highly variable even when compared to genes with similar average expression. We used a z-score cutoff of 2 for dispersion and average expression cutoff of 4 to identify highly variable genes. We then used those highly variable genes as input to the PCA to identify the primary data structures in 12, 14 and 20hpf data. For intermediate stages 16hpf and 18hpf, because of the known cell type similarity, we used cell type specific markers from 14hpf and 20hpf as input to PCA to obtain more robust dimensional reduction.

We extended the results of PCA analysis globally by projecting the PCA rotation matrix across the entire transcriptome. This additional projection allows us to identify other genes with strong PCA loadings that may not be included in our variable gene list. Statistically significant PCs were identified using a permutation test and independently confirmed using a modified resampling procedure[18] encoded in Seurat's 'jackStraw' function. Significant and biological meaningful PCs were retained for clustering and visualization. To visualize single cell data, we projected individual cells based on their PC scores onto a single two-dimensional map using t-distributed Stochastic Neighbor Embedding (t-SNE)[19].

### Single cell clustering and differential gene expression

Clustering of single cells was performed using the weighted shared nearest neighbor (SNN) graph-based clustering method[20]. To validate the legitimacy of clusters, we used 'ValidateClusters' function in Seurat, where we selected top 30 genes from significant PC's as defined above and utilized them to build a linear kernel SVM. The predictive accuracy of the SVM was assessed by repeated 5-fold cross validation. The accuracy cutoffs of 0.8 and 0.85 were used, and the merging of clusters was done based on the minimal connectivity from the SNN graph with a threshold of 0.001. Subsequently, TVC, STVC, FHP, SHP and ASM cells were identified from each time point data based on both known and newly identified markers (Extended Data Table 2). Specifically, for 12hpf data, no significant PC was identified due to the small and homogeneous TVC population. In 18hpf data, we also identified a cluster of 33 cells that expressed noticeable degree of both cardiac and ASM markers (Supplementary Note). We inferred this cluster of cells is possibly due to insufficient tissue-dissociation or sorting and sequencing errors. We removed these cells from further analysis.
The dataset can be mined through an online tool (ShinyApp) available at: (https://christiaenlab.shinyapps.io/tvc-lineage/) (e.g. test using the pan-cardiac and ASM markers GATA4/5/6 and EBF1/2/3/4)

To find markers differentially expressed among clusters, we used the same approach as in [21]. We used the binary classifier with ROC curve that was incorporated in Seurat's 'find.markers' function with parameters: test.use = 'roc', thresh.use = 1 and min.pct = 0.5, which selects genes that are expressed in more than 50% of single cells in the given cluster and with average expression larger than 1 $\log_2$(FPKM) for differential expression analysis. The selected genes were ranked based on AUCs from 0 to 1. The higher the AUC or power value the more differentially expressed the gene is for the given cluster. AUCs of 0.5 or below have limited predictive power.

### Single cell trajectory and transition state

We retrieved scRNA-seq data for each of the FHP, SHP, ASM trajectories by subsetting the master Seurat object containing all the single cell data. We adopted nonlinear dimensionality reduction technique of diffusion map[22–24], which reduces dimensionality through a random walking process, to identify developmental trajectory. We used the markers (power>0.3) of all cell types in each trajectory to calculate a cell-to-cell pairwise Euclidean distance matrix and used this matrix as input to diffusion map ('diffuse' function from 'diffusionMap' package). We retained only the first two diffusion map components as developmental trajectory for pseudotime analysis. Every cell was assigned to a pseudotime coordinate by fitting a principal curve[25] to the first two diffusion map components. Pseudotime was determined by the unit-speed arc-length parameterization of each cell on principal curve and normalized to [0,1] range.

After identification of pseudotime, we selected genes that are dynamically expressed across the pseudotime using 'aic' function in the 'locfit' package. Genes expressed in more than 50% of single cells with mean expression level greater than 2 log(FPKM) were considered as expressed in each cell type. For every expressed gene, we built two local polynomial models: a null model with degree 0 that assumes the gene expression stays constant along pseudotime and an alternative model with degree 2 that assumes gene expression changes along pseudotime[26]. We evaluated these two model using Akaike Information Criterion (AIC) to calculate the AIC score differences as AIC(degree=2) – AIC(degree = 0). Genes with AIC score differences lower than -5 were considered to favor the alternative model to be dynamically expressed in pseudotime space.

We then used these dynamic genes to subdivide the pseudotime space into distinct regulatory states separated by discrete transitions. First, we built a cell-to-cell cross-correlation matrix based on dynamic gene expressions for each trajectory. Constrained hierarchical clustering tree, which maintains pseudotime ordering, was built with the CONISS algorithm[27] using 'chclust' function from 'rioja' package based on the cross-correlation matrix. We used gap statistics to empirically determine the number of clusters (k) considered as the transition states along pseudotime. Briefly, we start with k = 1 (no transition), and increase it to 10. If the $k^{th}$ gap statistics $Gap_k$ increases less than 10% of the previous $k\text{-}1^{th}$ gap statistics $Gap_{k-1}$ then we consider the current k as the suitable number of transition states.

### Primed and *de novo* gene expression

Before defining primed and *de novo* genes, we first unbiasedly clustered the temporal gene expression patterns of both pan cardiac and ASM markers across all three trajectories. With k=2 of 'kmeans' clustering, we identified two groups of gene expression patterns in both cardiac and ASM genes that mimicked the primed and *de novo* patterns. Then we performed more stringent selection to define primed and *de novo* cardiac/ASM marker genes. We first identified all pan-cardiac and ASM markers (power>0.5) using the 18hpf and 20hpf datasets. Then we defined primed genes as genes that were expressed in more than 50% of single cells in both the multipotent progenitors (12hpf TVCs) and the fate restricted cells (18/20hpf FHPs/SHPs/iASMs/oASMs). Similarly, we defined *de novo*-expressed genes as expressed in less than 25% of single cells in the multipotent progenitors (12hpf TVCs) but expressed in more than 50% of single cells in the fate restricted cells (18/20hpf FHPs/SHPs/iASMs/oASMs). The genes expressed between 25%-50% of single cells were classified as ambiguous. The progenitor genes were defined as genes that were expressed in more than 50% of single cells in 12hpf TVCs but less than 25% of single cells in any of 18/20hpf FHP/SHP/iASM/oASM clusters. FHP/SHP specific markers were defined as genes that only belonged to FHP/SHP group but not to pan-cardiac or ASM gene set.

To visualize the smoothed pseudotime expression pattern and predict the induction time of a given gene, we smoothed the expression profiles along the pseudotime axis using local polynomial fit ('loess' function) with degree of smoothing equals to 0.75. Gene induction time was predicted based on the smoothed pseudotime expression profile using a logistic regression model. Gene expressions were first normalized to [0,1] range. Normalized expression values that were smaller than 0.5 were considered in 'off' state and bigger or equal to 0.5 were considered in 'on' state. We used this binary state notation and pseudotime coordinates to train a logistic model ('glm' function with family=binomial(link='logit')) to predict the on/off state of a given gene. The induction time was determined as the closest pseudotime coordinate to 0.5. Genes were then subdivided into two groups, either turning on or turning off, and sorted by their induction pseudotime.

To quantify the relative contribution of multipotent progenitor, primed ASM/cardiac, *de novo* ASM/cardiac and FHP/SHP specific genes, we performed principal component analysis using these groups of genes defined using above criteria on FHP and SHP trajectories separately. We observed that principal component 1 (PC1) strongly correlated with the defined pseudotime (PCCs>0.9). This allowed us to used PC1 loadings of each gene to calculate the contribution of each group as a scaled score using the formula: $G = {X_g}^T * {PCA^{rot}}_{[\bullet,1]}$. The $X_g$ represents scaled expression matrix of group $g$ where each column is a single cell and each row is a gene from group $g$. The ${PCA^{rot}}_{[\bullet,1]}$ represents the PC1 variable loadings which is the first column of the loading matrix. The G is the scaled score of gene group $g$. For heatmap visualization, we smoothed this score along the pseudotime axis using local polynomial fit ('loess' function) with degree of smoothing equals to 0.75.

#### Mouse single cell RNA-seq data

Mouse single cell data shown in Fig.4a were retrieved from the following website: http://gastrulation.stemcells.cam.ac.uk/scialdone2016.[26] The count matrix were transformed into log Transcripts per Million (logTPM). The variable genes, PCA and tSNE analysis were performed as described above. The visualization was based on the clustering results of the original paper. The mouse single cell data used for canonical correlation analysis were retrieved from the following papers and websites: http://singlecell.stemcells.cam.ac.uk/mesp1[28] and https://marionilab.cruk.cam.ac.uk/organogenesis.[29]

#### Canonical correlation analysis

Ciona and mouse genes were subset to those with known orthologs[30] and the best blast hits (using the cutoff as e-value<0.01 and qcovs>30) in both species . In cases where one gene in Ciona was duplicated in mouse, the corresponding gene was duplicated in the ciona gene expression matrix (Extended Data Table 6). Each dataset was first independently clustered and marker genes identified that were expressed in a subset of the clusters using a Wilcoxon rank sum test. Scaled and centered log-normalized expression values for common marker genes between both species were then used as the input gene set for a canonical correlation analysis between the two species[31]. The top canonical correlation vector that separated cardiac from pharyngeal muscle cells across species was then identified, and the top genes 30 that contributed most to this canonical correlation vector identified. The scaled and centered expression values for these 30 genes were then used to computed a 2-dimensional tSNE embedding for each species separately [32]. Clusters in this 30-gene space were identified using an unsupervised graph-based approach, as described previously [33–35]. Gene set enrichment analysis was performed on a non-redundant list of top CC genes for each mouse experiment using Panther[36], with the set of all gene orthologs between Ciona and mouse used as the background gene set.

### DATA AND SOFTWARE AVAILABILITY

The accession number for the expression data reported in this paper is GEO: GSE99846. The expression data and the code/Rmarkdown files for the analyses reported in this paper are available at https://github.com/stevexniu/single-cell-ciona.

### EXTENDED DATA FIGURES

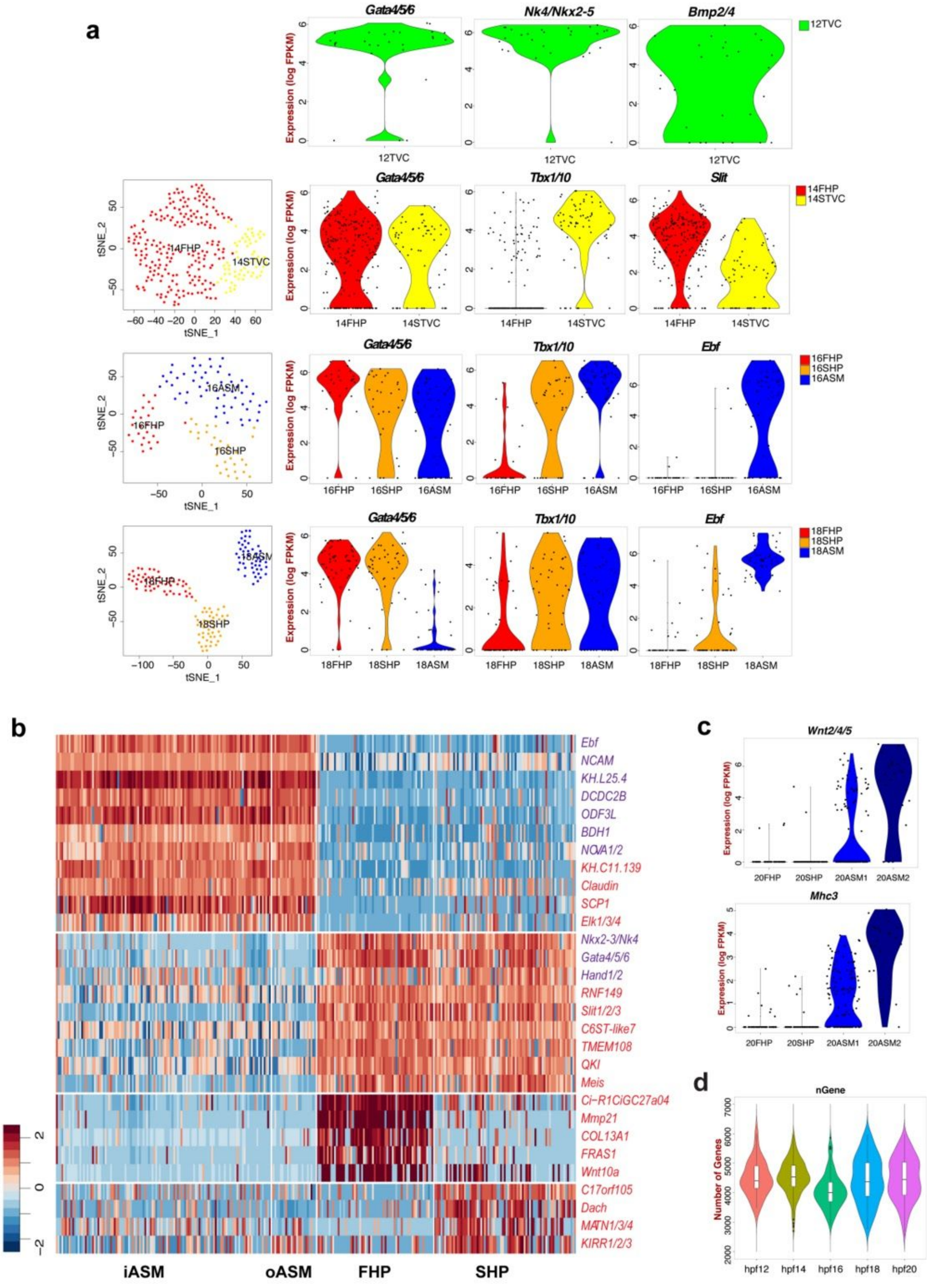
a
Gata4/5/6
Nk4/Nkx2-5
Bmp2/4
12TVC
Expression (log FPKM)
Gata4/5/6
Tbx1/10
Slit
14FHP
14STVC
tSNE_1
tSNE_2
Gata4/5/6
Tbx1/10
Ebf
16FHP
16SHP
16ASM
Gata4/5/6
Tbx1/10
Ebf
18FHP
18SHP
18ASM
b
Ebf
NCAM
KH.L25.4
DCDC2B
ODF3L
BDH1
NOVA1/2
KH.C11.139
Claudin
SCP1
Elk1/3/4
Nkx2-3/Nk4
Gata4/5/6
Hand1/2
RNF149
Slit1/2/3
C6ST-like7
TMEM108
QKI
Meis
Ci-R1CiGC27a04
Mmp21
COL13A1
FRAS1
Wnt10a
C17orf105
Dach
MATN1/3/4
KIRR1/2/3
Z-score
iASM
oASM
FHP
SHP
c
Wnt2/4/5
Mhc3
20FHP
20SHP
20ASM1
20ASM2
d
nGene
Number of Genes
hpf12
hpf14
hpf16
hpf18
hpf20

**Extended Data Figure 1| Cell clustering and novel cell-type-specific markers.**

(**a**) t-SNE plots showing clusters of individual transcriptomes from 14, 16, and 18hpf samples (left), and violin plots showing expression of indicated marker genes in defined clusters. (t-SNE is not shown for the 12 hpf data because it consists of a homogeneous population of TVCs). Color codes for cell identities as in Fig. 1a.

(**b**) Expression heatmap of 20 hpf single cell transcriptomes showing top predicted differentially expressed marker genes across different cell types. Blue names: previously known ASM and heart markers, red names: newly discovered markers.

(**c**) Violin plots showing expression of known outer Atrial Siphon Muscle Precursors (oASMP) markers in 20hpf scRNA-seq dataset as per Razy-Krajka et al.[4].

(**d**) Number of genes detected by scRNA-seq in samples obtained from larvae dissociated at different time points (12, 14, 16, 18, 20 hpf).

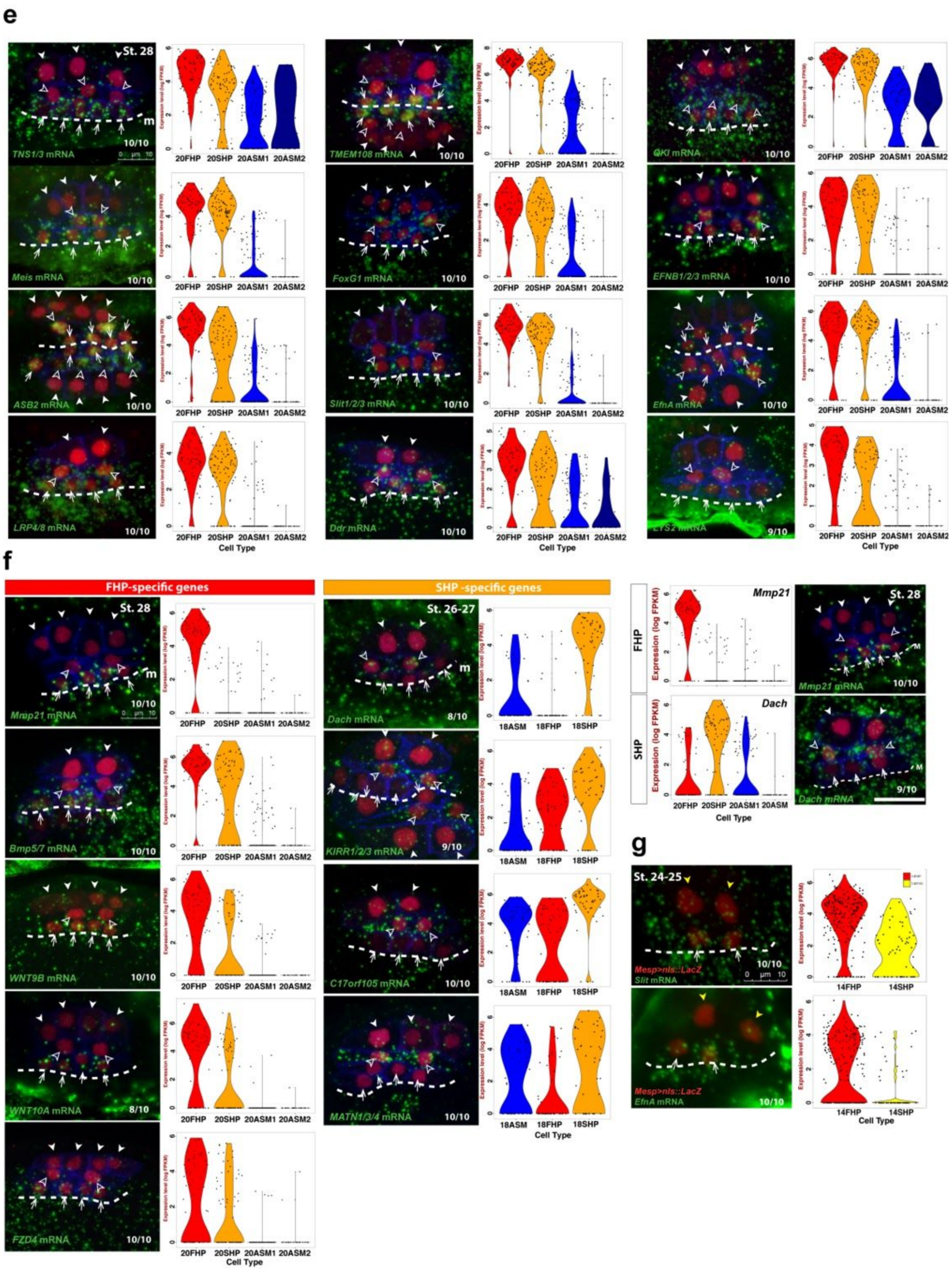


(**e**) Violin plots and FISH validation for indicated predicted pan-cardiac genes.
(**f**) Violin plots and FISH validation for indicated predicted FHP and SHP specific genes.
(**g**) Violin plots and FISH validation for indicated early *de novo* pan-cardiac markers genes in st.24-25 (14hpf) embryos.

(e-g) mRNAs are visualized by whole mount fluorescent in situ hybridization (green). Nuclei of TVCs are marked by *Mesp>nls::LacZ* and revealed by anti beta-galactosidase antibody (red). Mesp-driven hCD4::mCherry accumulates in the cell membrane and was revealed by anti mCherry antibody (blue). Anterior to the left. Scale bar, 10 µm. Solid arrowheads, ASM; open arrowheads, SHPs; arrows, FHPs; M, midline (dotted line). The numbers of embryos showing the demonstrated gene expression pattern and numbers of the observed embryos are indicated at the right bottom corner of each image.

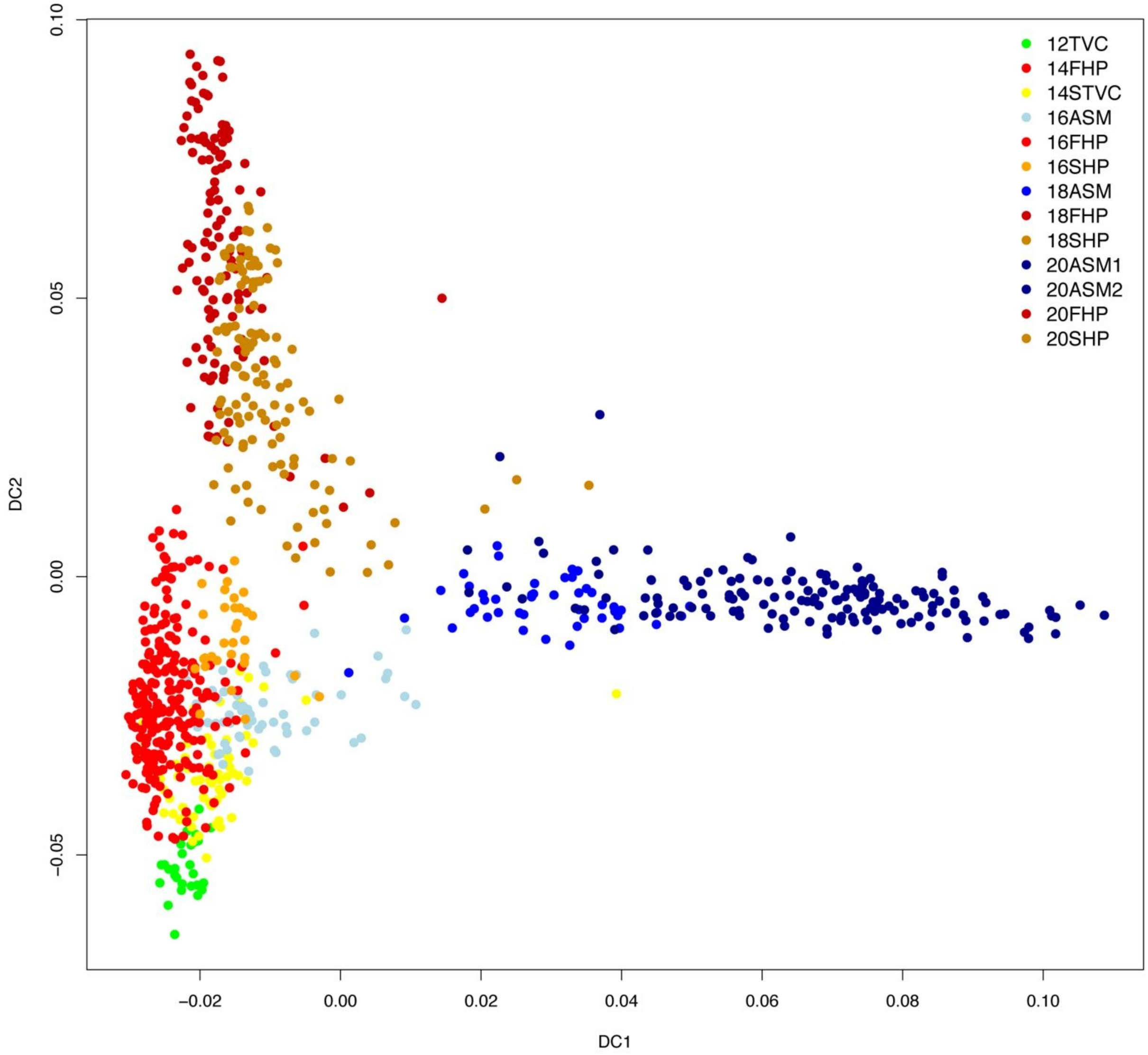


**Extended Data Figure 2| Diffusion maps showing the cardiopharyngeal specification trajectories in *Ciona*.**

Color-coded cell identities as defined by unsupervised clustering from larvae dissociated at different time points (12, 14, 16, 18, 20 hpf) (see Ext. Data Fig. 1a). DC: Diffusion Coordinate. By pooling all the Ciona cardiopharyngeal cells together, the diffusion map successfully captured the splitted trajectories of pharyngeal vs. cardiac fates, but failed to recover the branching points between the first and second heart lineages.

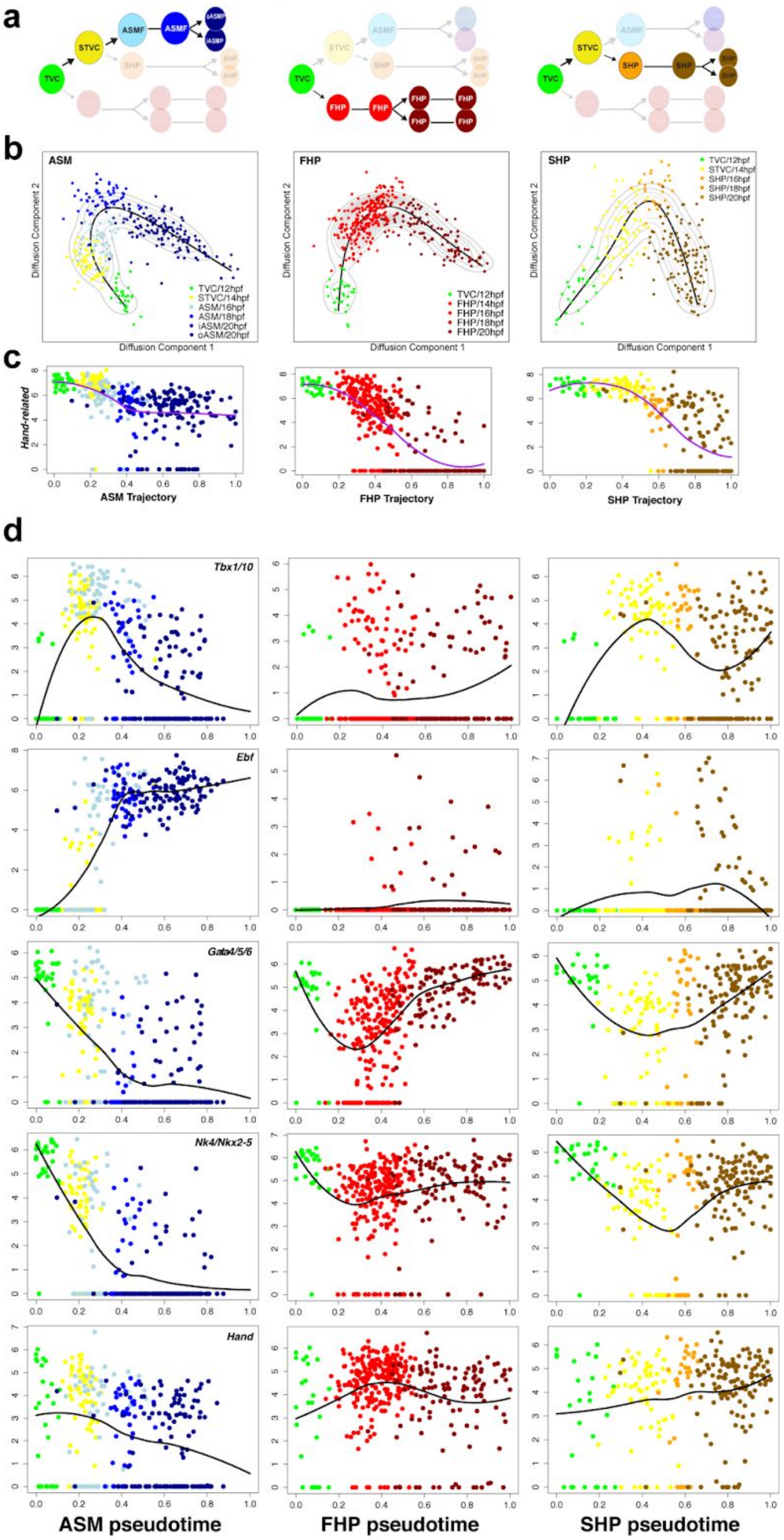
a
TVC
STVC
ASMF
oASMP
iASMP
SHP
FHP
b
ASM
FHP
SHP
Diffusion Component 1
Diffusion Component 2
TVC/12hpf
STVC/14hpf
ASM/16hpf
ASM/18hpf
iASM/20hpf
oASM/20hpf
FHP/14hpf
FHP/16hpf
FHP/18hpf
FHP/20hpf
SHP/16hpf
SHP/18hpf
SHP/20hpf
c
Hand-related
ASM Trajectory
FHP Trajectory
SHP Trajectory
d
Tbx1/10
Ebf
Gata4/5/6
Nk4/Nkx2-5
Hand
ASM pseudotime
FHP pseudotime
SHP pseudotime

**Extended Data Figure 3| Pseudotemporal reconstruction of cardiopharyngeal developmental trajectories based on established clonal relationships.**

(**a**) Cartoon shows the cells used to reconstruct the three unidirectional cardiopharyngeal progenitor trajectories.

(**b**) 2D diffusion maps of single cell transcriptomes for three developmental trajectories. Black lines indicate principal curves and light gray contours indicate single cell density distribution along principal curves. Color codes correspond to cell identities assigned by clustering at each time point (Ext. Data Fig. 1a). Hpf, hours post-fertilization.

(**c**) Expression of the primed ASM marker *Hand-related/Hand-r* on three trajectories. The purple lines show smoothed expression profiles. Y-axis: $\log_2$(FPKM). X-axis, normalized pseudotime along the principal curve (shown in (b)). Colors codes as in (b).

(**d**) Pseudotemporal expression profiles of indicated known marker genes along the ASM, FHP and SHP trajectories. Y-axes: expression in $\log_2$(FPKM). Black lines indicate the smoothed expression.

e

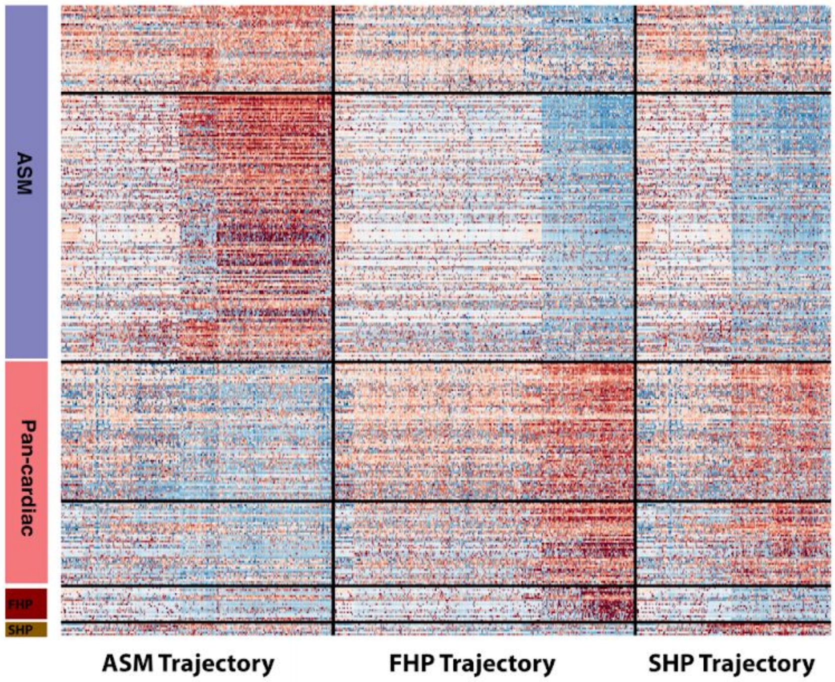


f

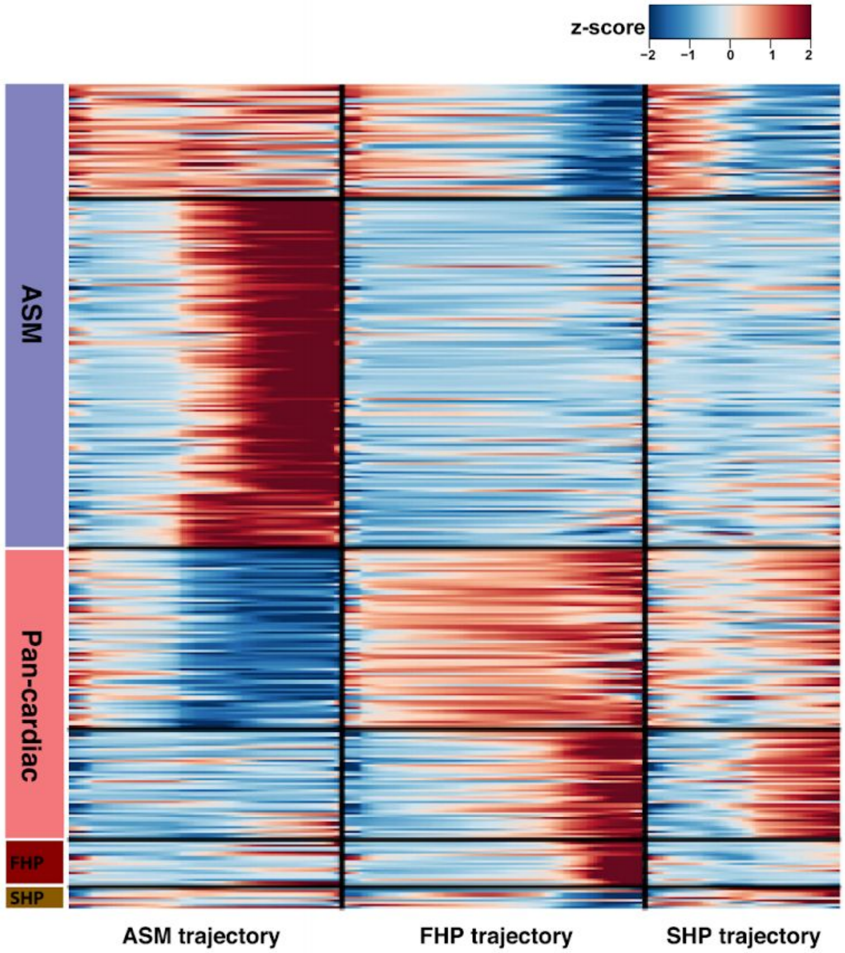




(**e**) Raw scaled expression heatmap of temporal expression pattern of indicated cell-type-specific marker genes on ASM, FHP and SHP trajectory.

(**f**) Heatmap of smoothed temporal expression pattern of indicated cell type specific marker genes on ASM, FHP and SHP trajectory corresponding to (e). Expressions are grouped by k-means clustering.

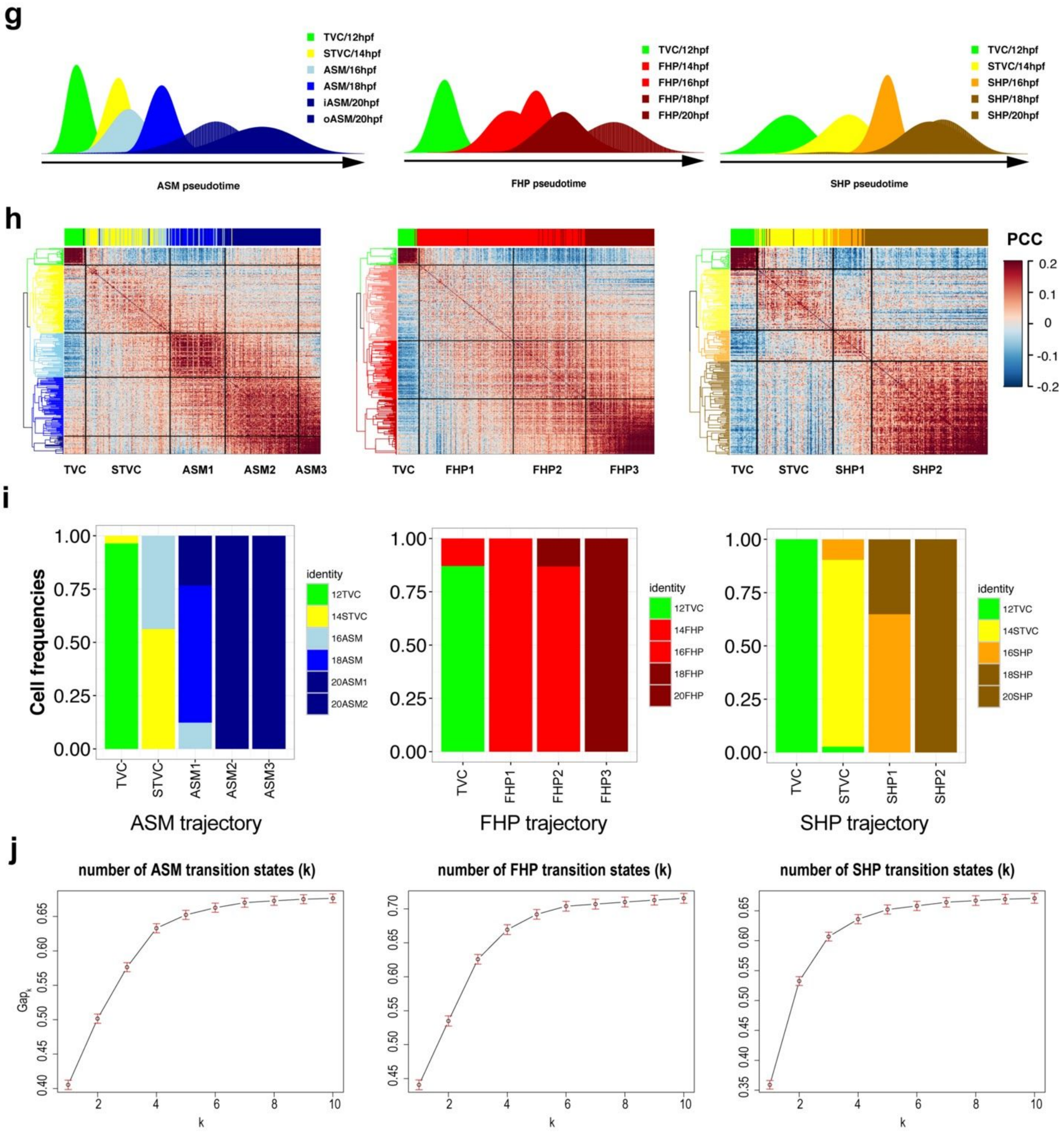


(**g**) Distribution of identified cell types isolated at defined time points along the ASM, FHP and SHP trajectories. This shows the agreement between the time series and developmental progression, but also that cells isolated from a given time points are not all at the same developmental "pseudotime".

(**h**) Cross-correlation heatmaps showing potential regulatory states along three trajectories. The left side dendrograms indicate the results of constrained hierarchical clustering. The top bars indicate the sample of origin for each cell ordered along the pseudotime axes. Color codes for top bars as in (g). PCC:Pearson's Correlation Coefficient.

(**i**) Bar plots showing relative cell identity composition for each regulatory states identified on ASM, FHP and SHP trajectories. Note that numbers of 16ASM cells clustering with the 'STVC' state in the ASM trajectory, and indicating that these cells retain most STVC characteristics and have not yet activated the ASM-specific program (see text for the details).
(**j**) Gap statistics plots used to determine the number of transition states in each trajectory.

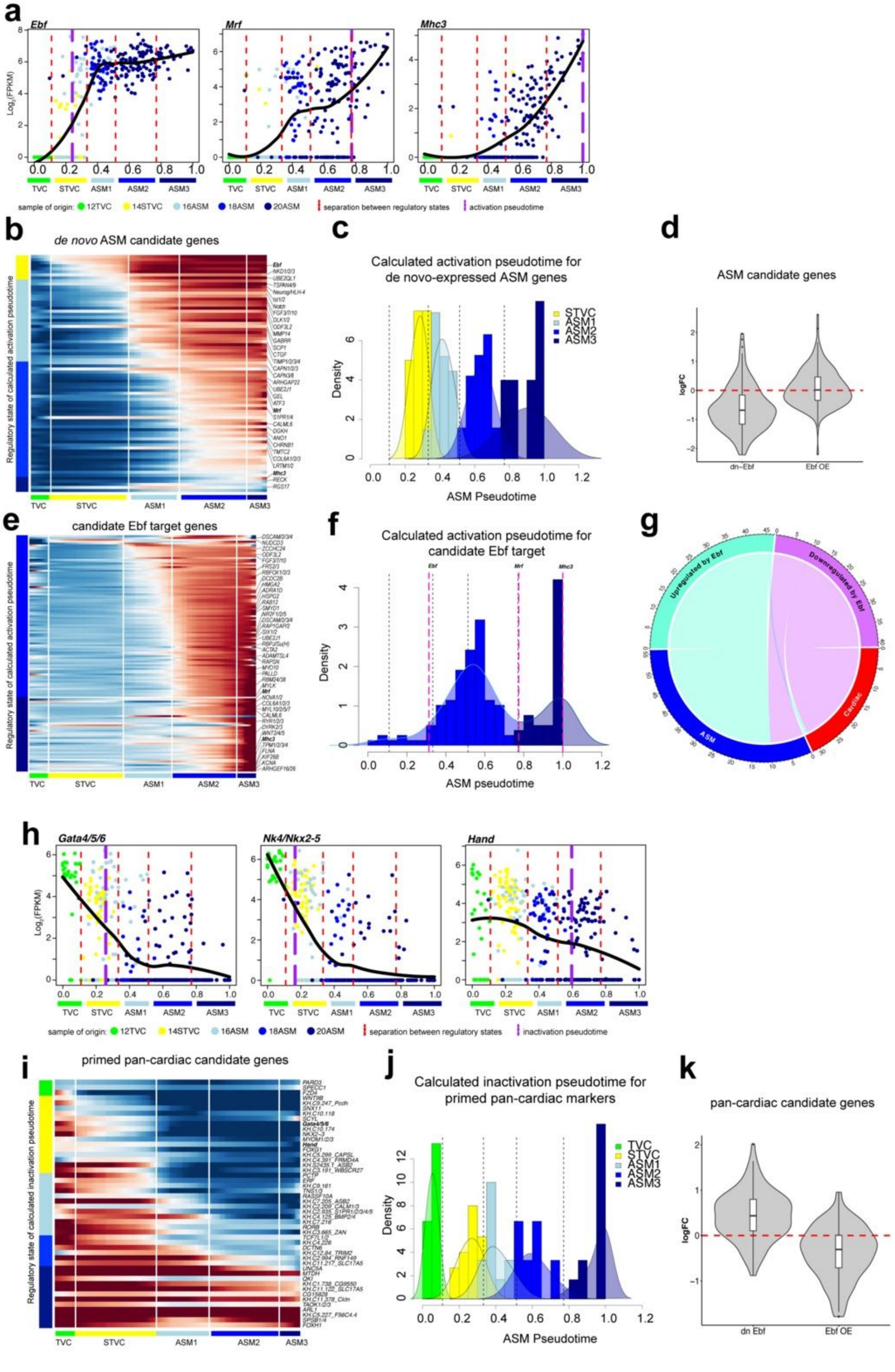

a
Ebf
Mrf
Mhc3
Log2(FPKM)
TVC STVC ASM1 ASM2 ASM3
sample of origin: 12TVC 14STVC 16ASM 18ASM 20ASM
separation between regulatory states
activation pseudotime
b
de novo ASM candidate genes
Regulatory state of calculated activation pseudotime
c
Calculated activation pseudotime for de novo-expressed ASM genes
Density
ASM Pseudotime
d
ASM candidate genes
logFC
dn-Ebf
Ebf OE
e
candidate Ebf target genes
f
Calculated activation pseudotime for candidate Ebf target
ASM pseudotime
g
Upregulated by Ebf
Downregulated by Ebf
ASM
Cardiac
h
Gata4/5/6
Nk4/Nkx2-5
Hand
inactivation pseudotime
i
primed pan-cardiac candidate genes
Regulatory state of calculated inactivation pseudotime
j
Calculated inactivation pseudotime for primed pan-cardiac markers
k
pan-cardiac candidate genes
dn Ebf
Ebf OE

**Extended Data Figure 4| Transcriptional regulation of ASM fate specification.**

(**a**) Pseudotemporal expression profiles of indicated genes along the ASM trajectory. X-axis: normalized pseudotime as defined in Ext. Data Fig. 3. Y-axes: expression in $log_2$(FPKM). Black lines indicate the smoothed expression. Red dashed lines indicate the transitions between predicted regulatory states, indicated and color-coded as in Ext. Data Fig. 3, and purple dashed lines indicate calculated activation pseudotime.

(**b**) Heatmap of smoothed pseudotemporal expression profiles showing successive activation of candidate *de novo* ASM genes. White vertical lines mark transitions between indicated regulatory states along the ASM trajectory. Colored bars on the left indicate the regulatory state of calculate activation pseudotime, showing that most ASM candidates turn on in the ASM1 (light blue) and ASM2 (navy blue) states.

(**c**) Histogram with density lines showing predicted induction time of *de novo* ASM genes with corresponding ASM regulatory states (Separated by black dashed lines).

(**d**) Violin plots showing the $log_2$(fold change) of all candidate ASM-specific genes in response to defined perturbations of Ebf function, a dominant-negative (dn-Ebf) and Ebf precocious and misexpression (Ebf-OE) as described in Razy-Krajka et al.[4].

(**e**) Heatmap of smoothed pseudotemporal expression profiles for candidate ASM-specific Ebf target genes defined in Razy-Krajka et al.[4], and showing two main waves of activation in regulatory states ASM2 and ASM3 (left colored bars). The white vertical lines mark transitions between indicated regulatory states.

(**f**) Histogram with density lines showing predicted activation pseudotime of candidate Ebf target genes with corresponding ASM regulatory states (separated by black dashed lines). Purple lines indicate predicted *Ebf*, *Mrf* and *Mhc3* induction time respectively.

(**g**) Chord diagram showing mutual enrichment of ASM vs. Cardiac genes among candidate target genes activated or inhibited by Ebf, respectively. Note that Ebf is predicted to downregulate a few ASM candidate genes, which are primed and quickly downregulated after the onset of Ebf (e.g. *Hand-r*, [4]).

(**h**) Gene plots showing the progressive depletion of the indicated primed pan-cardiac genes along the ASM trajectory. X-axis: normalized pseudotime as defined in Ext. Data Fig. 3. Y-axes: expression in $log_2$(FPKM). Black lines indicate the smoothed expression. Red dashed lines indicate transitions between indicated regulatory states and purple dashed lines indicate predicted inactivation pseudotime.

(**i**) Heatmap of smoothed pseudotemporal expression profiles showing progressive depletion of primed pan-cardiac genes along the ASM trajectory, with calculated pseudotime of inactivation mapped onto discrete regulatory states indicated on the left colored bars.

(**j**) Histogram with density lines showing predicted inactivation pseudotime of primed cardiac genes with corresponding ASM regulatory states (separated by black dashed lines).

(**k**) Violin plots showing the log fold changes of pan-cardiac genes corresponding to Ebf perturbation.

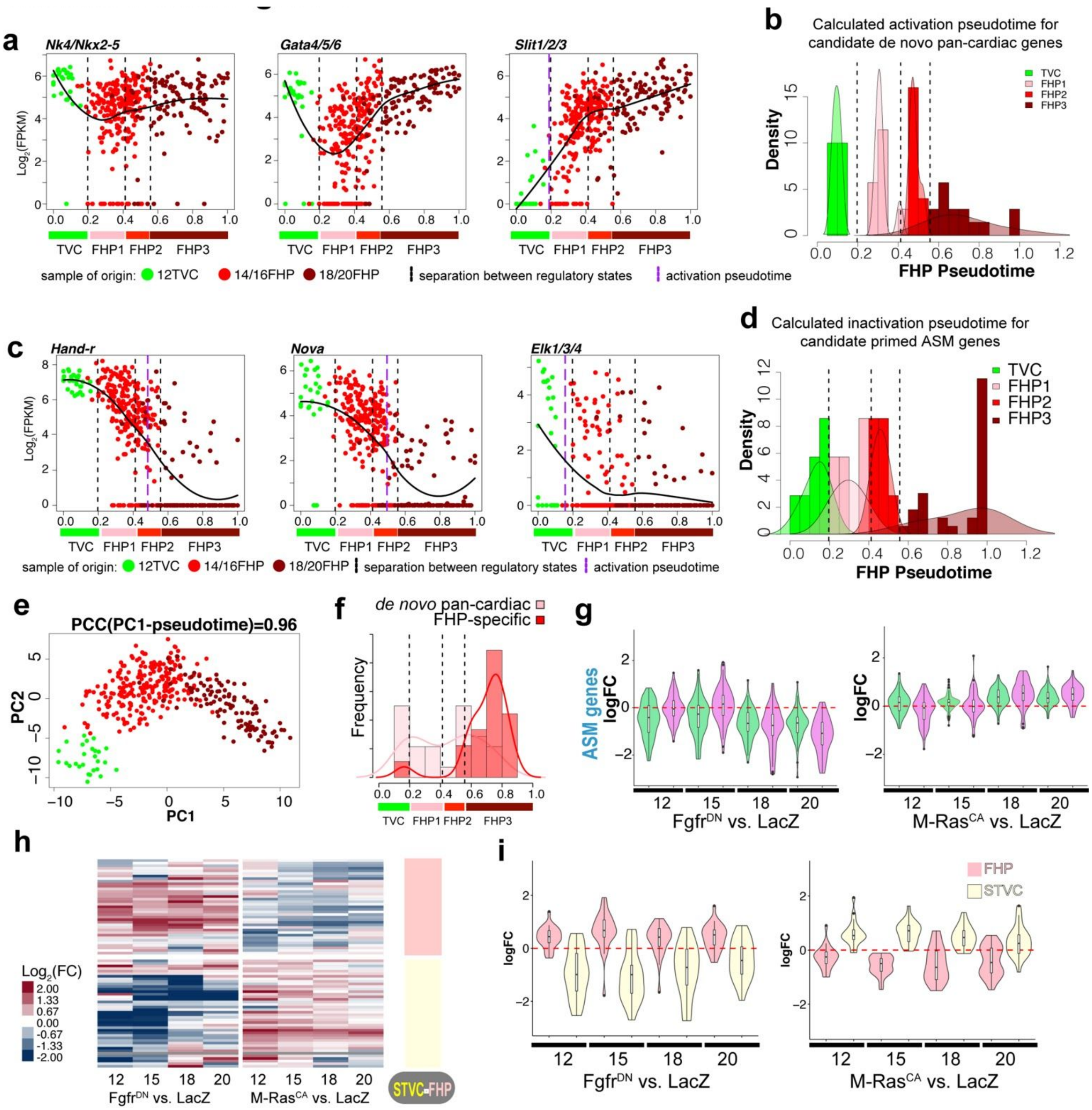


**Extended Data Figure 5| Transcriptional regulation of FHP fate specification.**

(**a**) Pseudotemporal expression plots showing the activation of the representative pan-cardiac genes on FHP trajectory. X-axis: normalized pseudotime as defined in Fig. 1c, Y-axes: expression in $\log_2$(FPKM). Black lines indicate the smoothed expression. Black dashed lines indicate the transitions between predicted regulatory states, indicated and color-coded below, and purple dashed lines indicate calculated activation pseudotime. Dots colors refer to the sample of origin as indicated in Fig. 1a.

(**b**) Density plots showing the number of *de novo* pan-cardiac genes with calculated activation pseudotime in binned pseudotime windows along FHP trajectories. The black dashed lines indicate the regulatory states.

(**c**) Pseudotemporal expression plots showing the depletion of the representative primed ASM genes on FHP trajectory. X-axis: normalized pseudotime as defined in Fig. 1c, Y-axes: expression in $\log_2$(FPKM). Black lines indicate the smoothed expression. Black dashed lines indicate the transitions between predicted regulatory states, indicated and color-coded below, and purple dashed lines indicate calculated inactivation pseudotime. Dots colors refer to the sample of origin as indicated in Fig. 1c.

(**d**) Histogram with density lines showing predicted inactivation pseudotime of primed ASM genes with corresponding FHP regulatory states (separated by black dashed lines).
(**e**) Principal component analysis showing correlation between PC1 and pseudotime.
(**f**) Density plots showing the number of *de novo* pan-cardiac and FHP-specific genes with calculated activation pseudotime in binned pseudotime windows along FHP trajectory. The black dashed lines indicate the regulatory states. FHP-specific genes tend to be activated later than pan-cardiac markers.
(**g**) Violin plots showing the log fold changes of ASM genes in indicated conditions and time points relative to LacZ controls.
(**h**) Heatmap showing $\log_2$(fold change) of early FHP and STVC genes (identified in the 14hpf scRNA-seq data) in indicated conditions and time points relative to LacZ controls.
(**i**) Violin plots showing log fold changes for defined categories of genes (FHP and STVC markers identified in the 14hpf scRNA-seq data) in indicated conditions and time points relative to LacZ controls.
(**j-q**) Bar plots representing the gene set enrichment statistics reported in Ext. Data Table 4. In all panels, only ASM and pan-cardiac markers (defined by scRNA-seq, as in Fig. 1) are shown but all expressed genes ($\log_2$(CPM)>3) were considered for the statistics. Responses to indicated perturbations of FGF-MAPK signaling at defined time points were categorized into "up-regulated", "down-regulated" or "no change" using EdgeR summary statistics based on biological duplicate experiments, using arbitrary cutoffs [$|\log_2$(fold change)$| > 1$ and P-value <0.1]. For each panel, plots on the left show the expected and observed numbers of genes in indicated categories, plots on the right show enrichment as $\log_2$(observed/expected). Asterisks indicate significant enrichment, as determined by hypergeometric tests: ***, $p < 0.001$; **, $p<0.01$; *, $p<0.05$. Detailed reports are available in Extended Data Table 4.

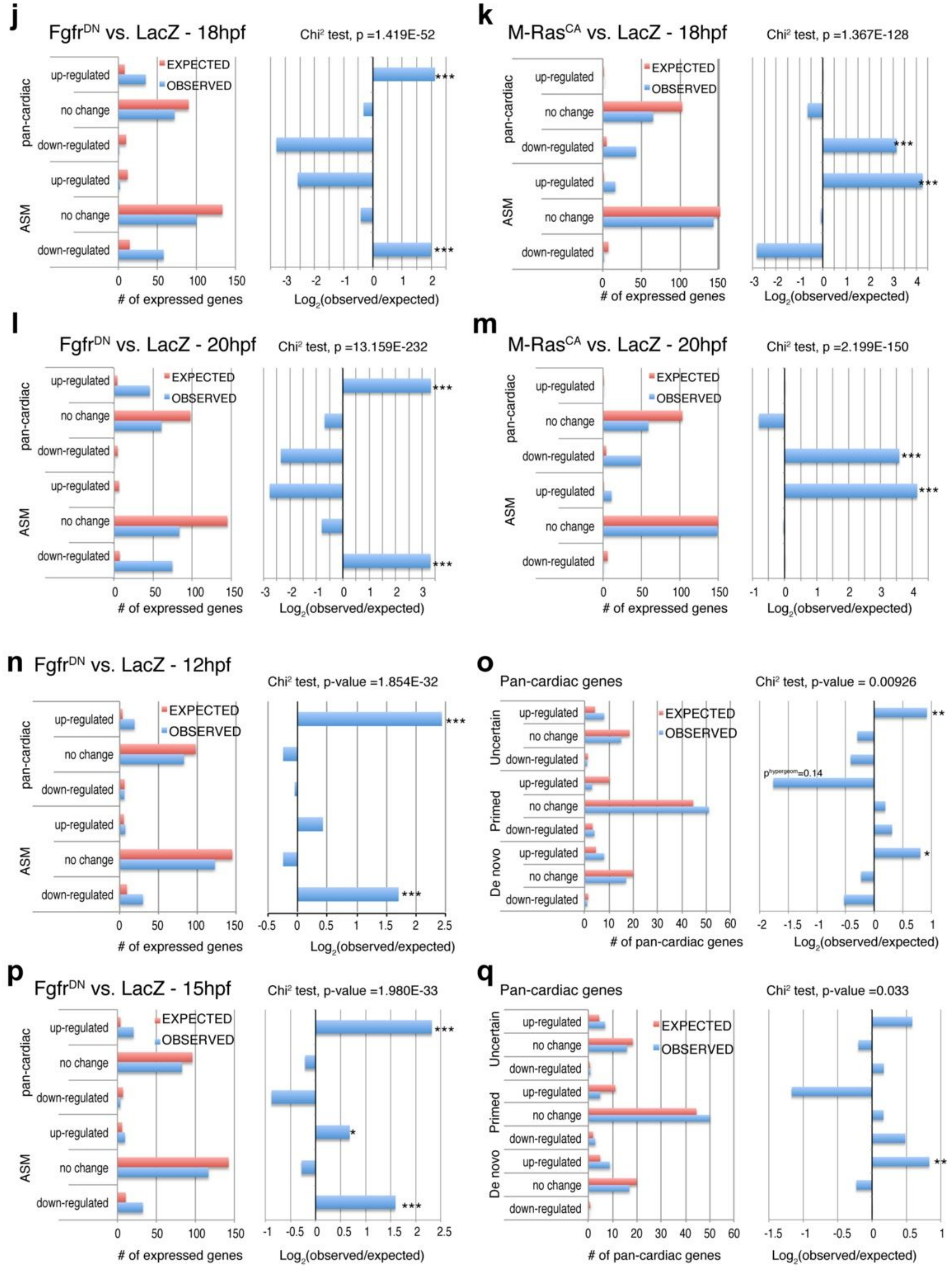


(**j-m**) At late times points (cardiopharyngeal lineage cells sorted from 18 or 20 hpf larvae), inhibition of

FGF-MAPK signaling up-regulated more pan-cardiac markers and down-regulated more ASM markers than expected by chance (j,l), whereas gain of FGF-MAPK function down-regulated more pan-cardiac markers and up-regulated more ASM markers than expected by chance. These results are consistent with the hypothesis that FGF-MAPK promotes the ASM program while inhibiting the pan-cardiac program.
(**n-q**) At earlier time points, when multipotent cardiopharyngeal progenitors are present in the control samples, loss of FGF-MAPK function down-regulated more ASM markers than expected by chance, which is consistent with the notion that expression of primed ASM markers already depends on FGF-MAPK signaling in multipotent progenitors. By contrast, loss of FGF-MAPK function upregulated more of the *de novo* expressed (and uncertain) pan-cardiac markers than expected by chance, which is consistent with our hypothesis that FGF-MAPK activity in multipotent progenitors prevents the activation of de novo-expressed pan-cardiac markers.

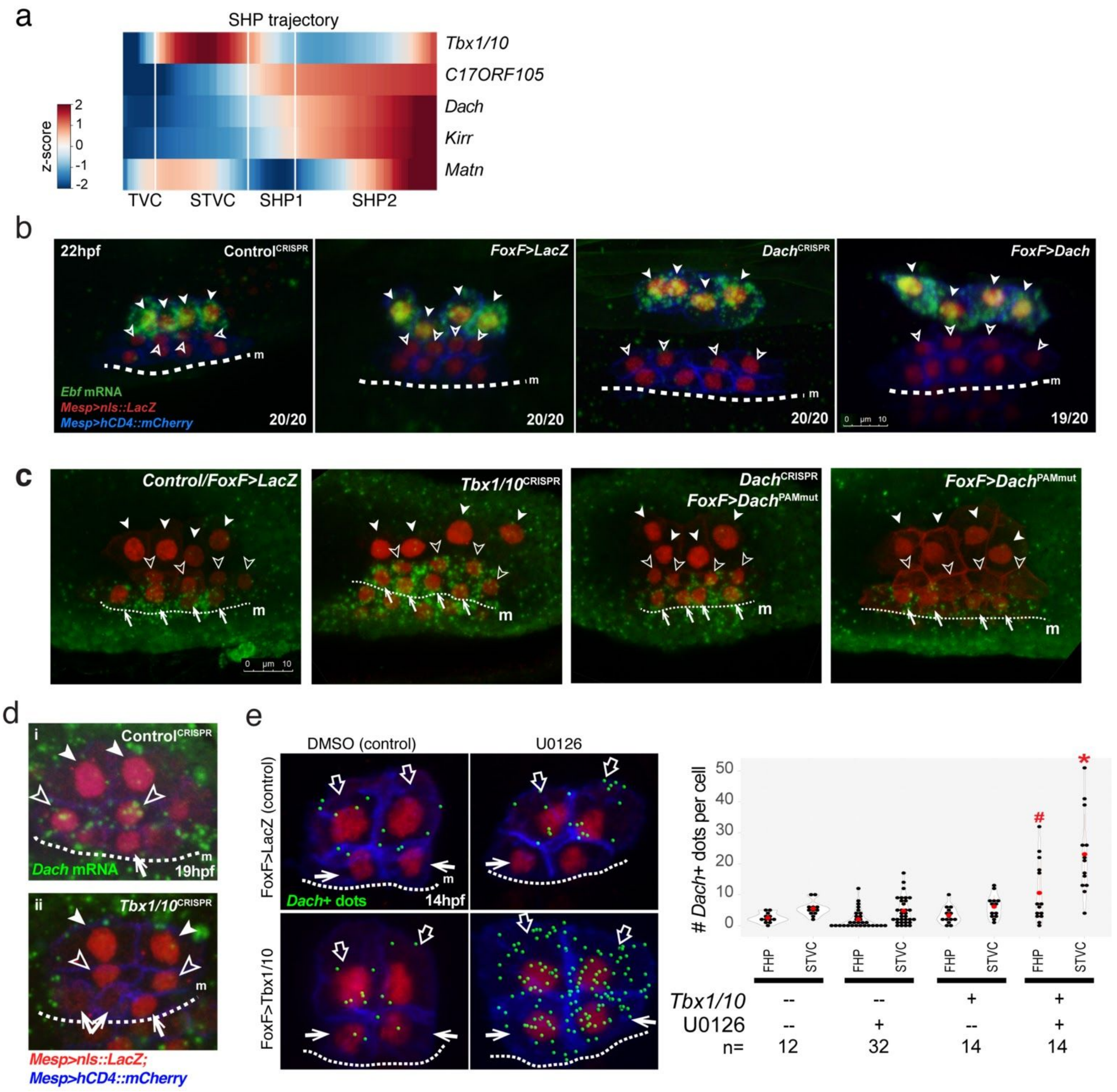


**Extended Data Figure 6| Dach regulates the SHP fate specification.**

(**a**) Heatmap of smoothed gene expression showing *Tbx1/10* activation and SHP-specific genes, including *Dach*, on the SHP trajectory. White vertical lines indicate transitions between predicted SHP regulatory states.

(**b**) *Ebf* expression specifically in the ASMPs at 22hpf is not altered by perturbations of Dach function. Conditions indicated on top-right corner of each panel. Ebf mRNAs are visualized by whole mount fluorescent in situ hybridization (green). Nuclei of TVC-derived cells progeny are marked by Mesp>nls::LacZ and revealed by anti beta-galactosidase antibody (red). Mesp-driven hCD4::mCherry accumulates in the cell membrane and revealed by anti mCherry antibody (blue). Numbers indicate observed/total scored. Solid arrowheads: ASMPs, open arrowheads: SHPs, dotted line: midline (m).

(**c**) Dach and Tbx1/10 are required to prevent expression of the FHP-specific gene *Mmp21* in the SHPs. Conditions indicated on top-right corner of each panel. *Mmp21* mRNAs are visualized by whole mount fluorescent in situ hybridization (green). Nuclei of TVC-derived cells progeny are marked by Mesp>nls::LacZ and revealed by anti beta-galactosidase antibody (red). Mesp-driven hCD4::mCherry accumulates in the cell membrane and revealed by anti mCherry antibody (red). Open arrowheads,

ASMPs; solid arrowheads, SHPs; arrows, FHPs. Dotted line: ventral midline. Anterior to the left. Scale bar, 10 µm.

(**d**) Tbx1/10 is required for *Dach* expression in SHP. *Dach* expression is visualized by *in situ* hybridization (green). Nuclei of B7.5 lineage cells are labelled by *Mesp>nls::LacZ* and revealed with an anti beta-galactosidase antibody (red). Mesp-driven hCD4::mCherry accumulates at the cell membrane as revealed by anti mCherry antibody (Blue). Solid arrowheads, ASMFs, open arrowheads, SHPs; arrows, FHPs. Anterior to the left, stages indicated in hpf. Scale bar, 10 µm. (**i**) endogenous expression of *Dach* in SHPs in a larvae electroporated with *Mesp>Cas9* and a control single guide RNA-expressing construct ($CRISPR^{Control}$ ). (ii) endogenous *Dach* expression is inhibited upon CRISPR/Cas9-mediated loss of Tbx1/10 function. Experiment performed in biological replicates. For each replicate, confocal stacks were acquired for 10 larvae in each condition. None of the 20 Tbx1/10 CRISPR larvae showed *Dach* expression in SHPs.

(**e**) Overexpression of Tbx1/10 and blocking of FGF-MAPK induce precocious expression of *Dach* in all of B7.5 lineage cells at 14hpf. Imaris-processed images showing *Dach* expression (green dots) in the TVC-derived cells of 14hpf embryos. Open arrows: STVCs, arrows: FHPs; dotted line: midline (m).

(**f**) Violin plots corresponding to (e) and representing the counts of *Dach*+ dots per cell. Red dots indicate the mean value in each condition. Vertical red lines indicate mean +/- SD. * $p<0.05$; # $p<0.1$ (Student's t-test).

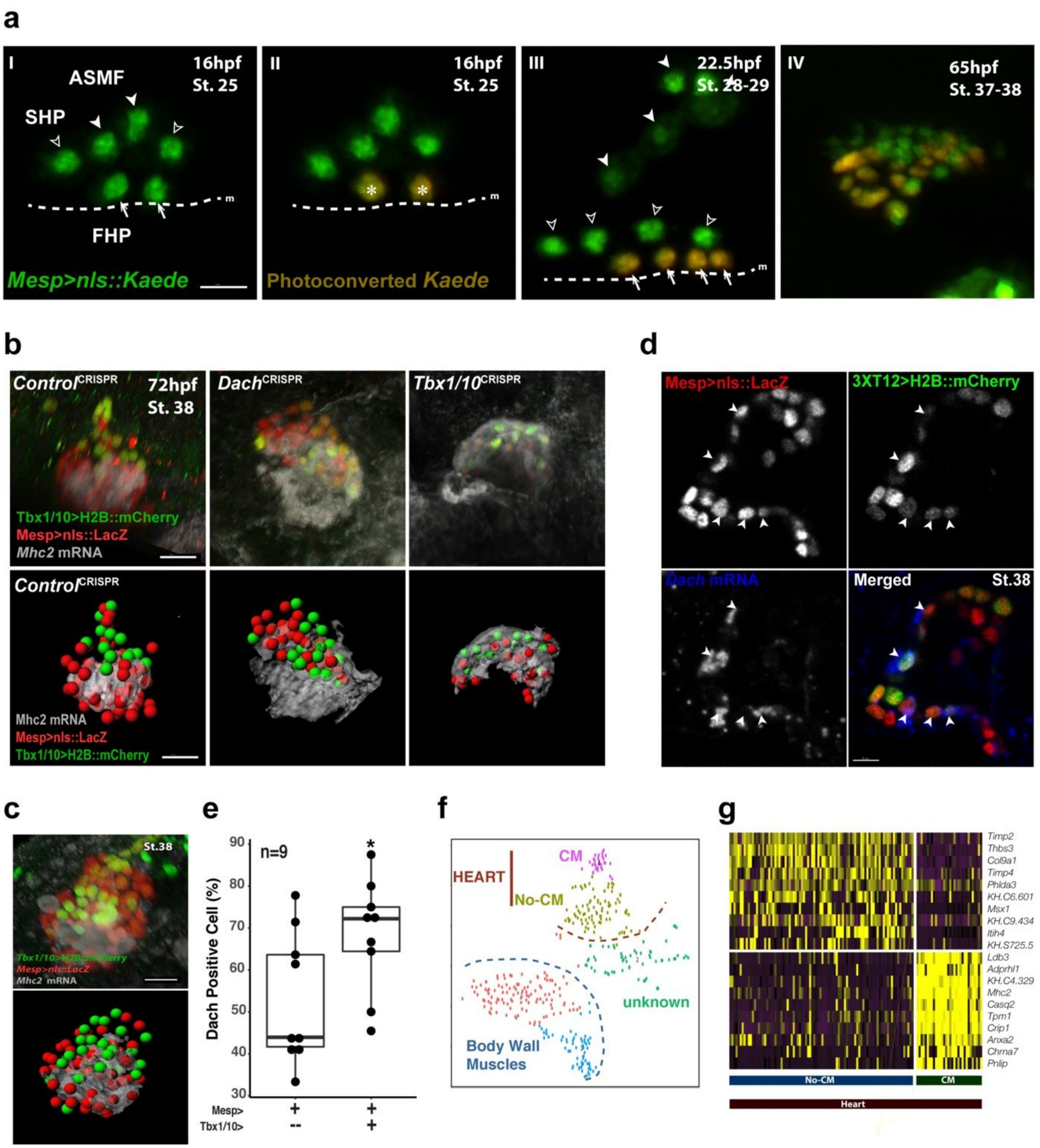


### Extended Data Figure 7| The juvenile heart

(**a**) Photoconversion and lineage tracing of TVC progeny. Nucleus of live B7.5 lineage cells are labelled with *Mesp>nls::Kaede::nls* (green). Nuclear Kaede photoconverted from green (I) to red (II) specifically in the FHPs in a 16hpf larva. The same animal is shown at successive time points (indicated as hours post-fertilization, hpf). I' to IV' show segmented nuclei. Open arrowheads, ASMPs; solid arrowheads, SHPs; arrows, FHPs, Dotted line: midline.

(**b**) *Dach* and *Tbx1/10* is required to limit the proportion of SHP-derived cells forming *Mhc2*+ cardiomyocytes in juveniles. Grey: *Mhc2* mRNA visualized by *in situ* hybridization. SHP-derived cells are labelled with *3XT12>H2B::mCherry* (green), B7.5 lineage cells are labelled with *Mesp>nls::LacZ* (red). Upper-panels : The original confocal images; lower-panels: The Imaris processed images for better illustration (method).

(**c**) The wild-type Juvenile heart (St. 38). Raw confocal data (left) and Imaris processed image (right) showing Mhc2 expression (grey) primarily in FHP-derived cells marked by the B7.5 lineage marker Mesp>nls::LacZ (red), but not by the STVC-specific marker Tbx1/10(3xT12)>H2B::mCherry (green), which marks the SHP-derived cells. Scale bar, 10µm.
(**d**) *Dach* is enriched in the Tbx1/10(3xT12)>H2B::mCherry+ cells (green), which marks the SHP-derived cells. *Dach* mRNAs are visualized by whole mount fluorescent in situ hybridization (Blue). Nuclei of TVC-derived cells progeny are marked by Mesp>nls::LacZ and revealed by anti beta-galactosidase antibody (red). Solid arrowheads: Tbx1/10(3xT12)>H2B::mCherry+; Mesp>nls::LacZ+ SHP-derived cells has *Dach* expression. Scale bar, 5µm. Note that the images are X - Y cross section of the juvenile heart.
(**e**) Boxplots corresponding to (d) with the proportions of *Dach* cells among the Mesp>nls::LacZ+;Tbx1/10(3xT12)>H2B::mCherry+ SHP-derived cells and Mesp>nls::LacZ+;Tbx1/10(3xT12)>H2B::mCherry- cells in juvenile hearts. Note the *Dach* is significantly enriched in SHP-derived cells, comparing to the percent of *Dach*+ cells in the Mesp>nls::LacZ+;Tbx1/10(3xT12)>H2B::mCherry- cells derived from both FHPs and SHPs (note: a fraction of SHP-derived cells are not labelled by Tbx1/10(3xT12)>H2B::mCherry due to the mosaicism) . Bars in the box indicate the median value in each condition. * $p<0.05$ (Student's t-test).
(**f**) The t-distributed Stochastic Neighbor Embedding (t-SNE) plots of st. 38 juvenile scRNA-seq data of FACS purified TVC-derived cells. Clusters "CM" and "No-CM" form the juvenile heart.
(**g**) Expression heatmap of single cell transcriptomes showing top predicted differentially expressed marker genes across the st. 38 juvenile heart.

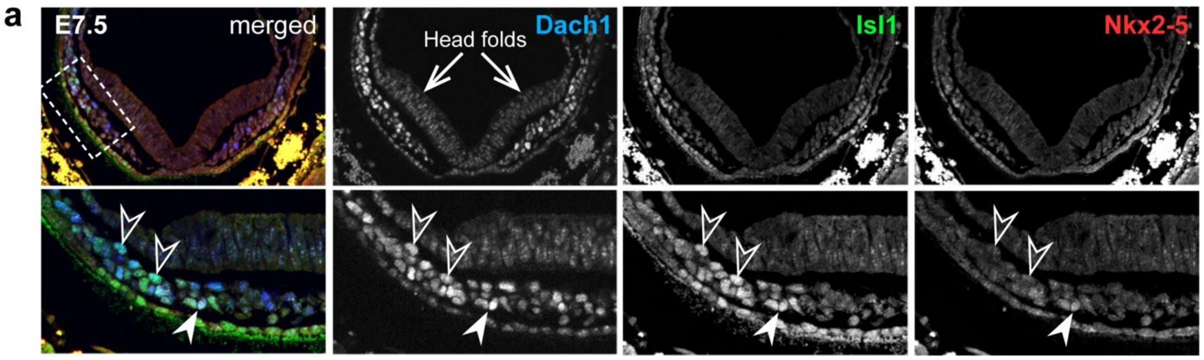


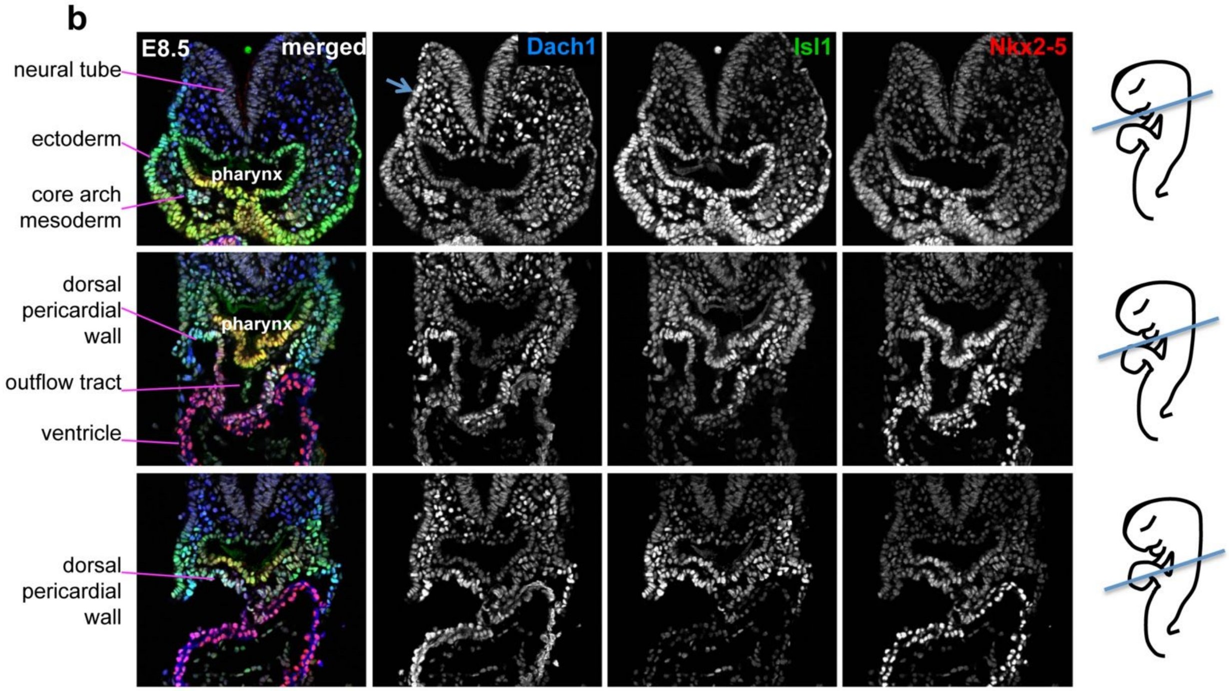


**Extended Data Figure 8| Dach1 expression in murine cardiopharyngeal mesoderm.**
(**a**) Expression patterns of Dach1, Islet1 and Nkx2.5 proteins in E7.5 embryos as indicated. Bottom panels are close-up views of the regions boxed with a dotted line. Arrows: head folds, Open arrowheads: double Dach1+, Islet1+ pharyngeal mesoderm cell, solid arrowhead: Nkx2.5 expression in a Dach1+, Islet1+ cell.
(**b**) Dach1, Isl1 and Nkx2-5 protein expression detected by immunofluorescence in E8.5 mouse embryos at distinct levels in the pharyngeal mesoderm and developing heart. Note the broad Dach1 expression, which overlaps with Isl1 in second heart field cells in the dorsal pericardial wall.

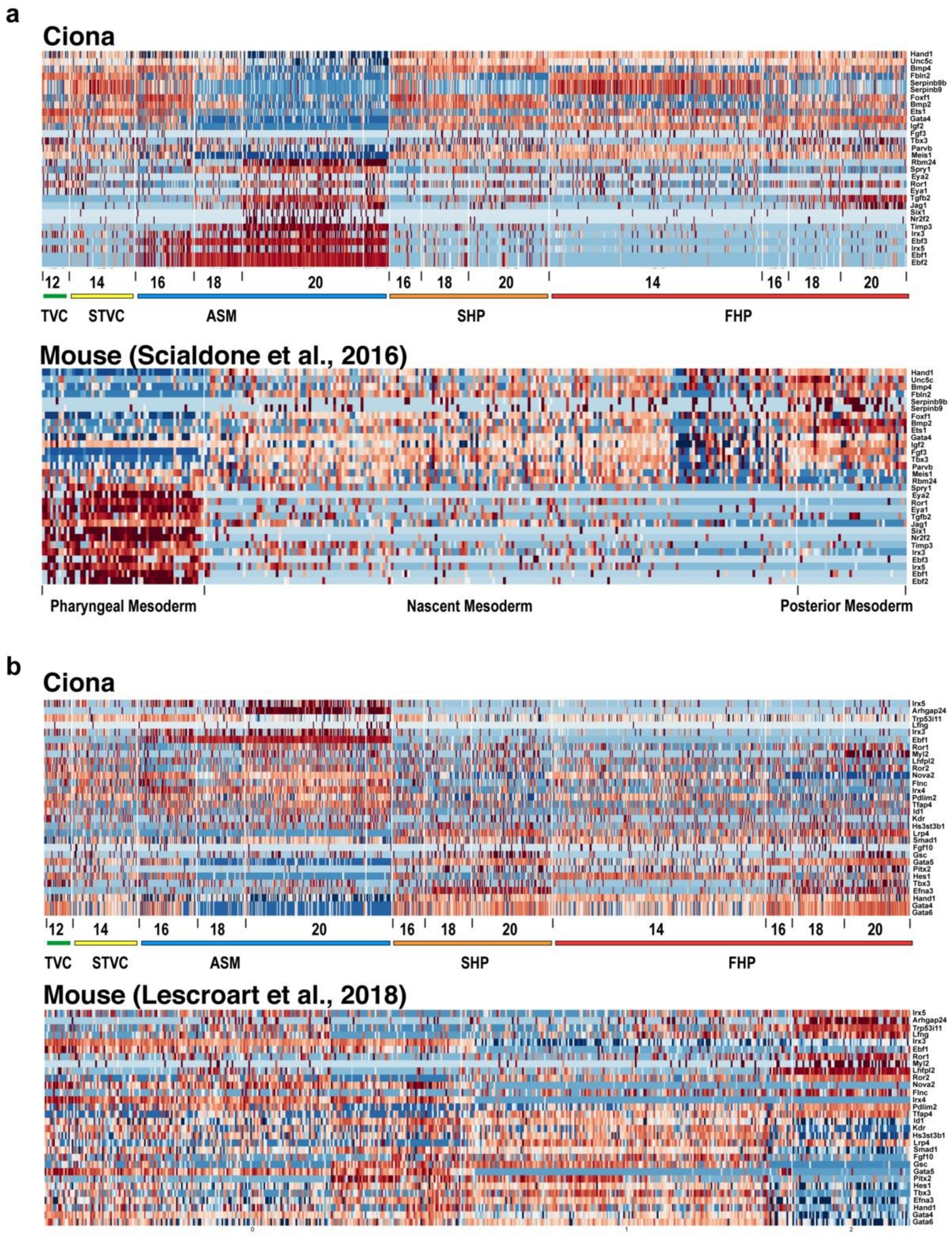


**Extended Data Figure 9| A conserved cardiopharyngeal program shared between Ciona and the mouse.**

Expression heatmaps of the top 30 genes of canonical correlation vector that contributed most to the heart vs. pharyngeal mesoderm separation in Ciona and the mouse. Detailed informations are provided in Extended Data Table 5.
(**a**) Top 30 genes identified from the canonical correlation analysis performed on Ciona and Mouse E6.50-E7.25 single cell RNAseq data from Scialdone et al.[26].
(**b**) Top 30 genes identified from the canonical correlation analysis performed on Ciona and Mouse Mesp+ E6.75- E7.50 single cell RNAseq data from Lescroart et al.[28].

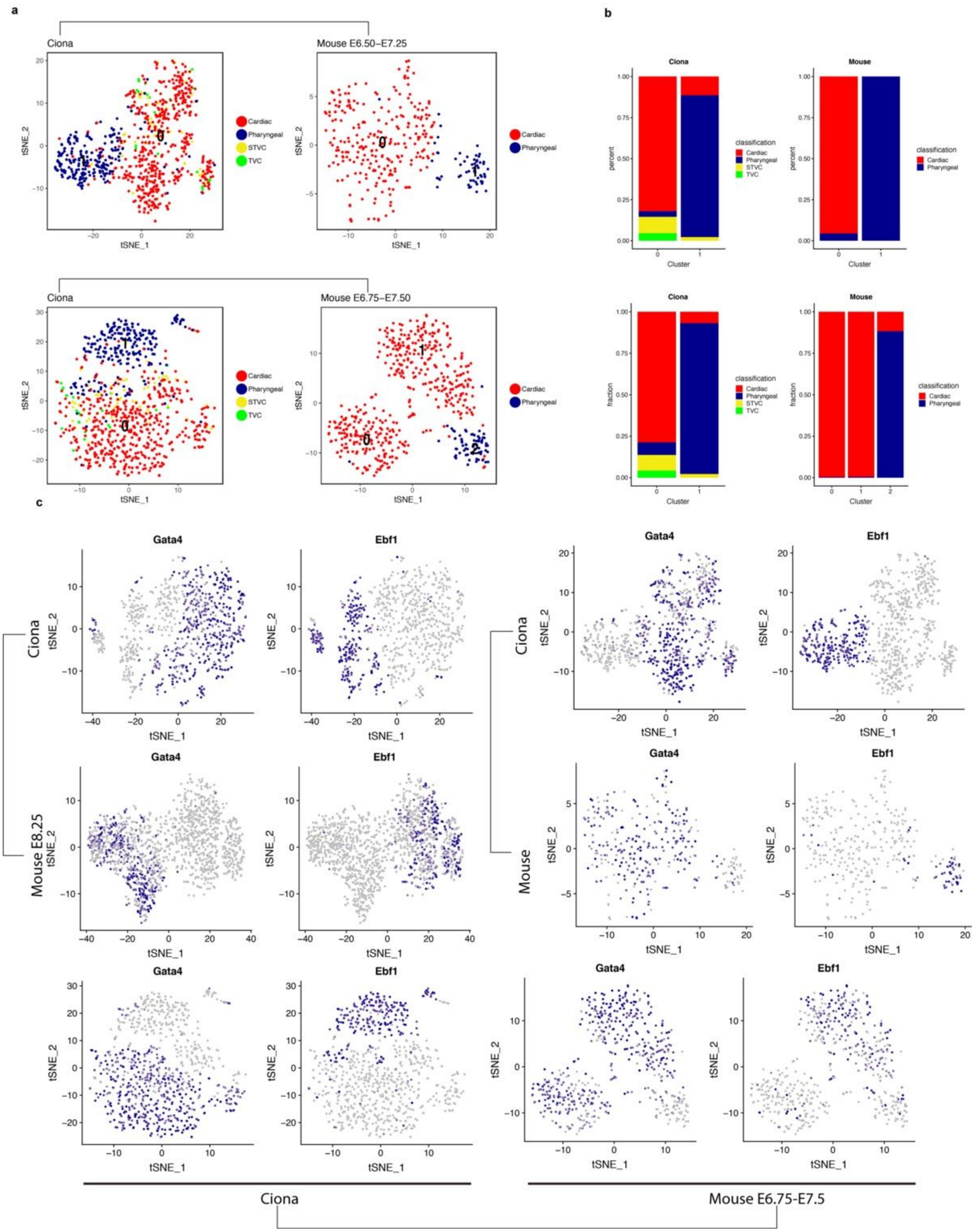


**Extended Data Figure 10|**

(**a**) tSNE plots of Ciona and Mouse scRNA-seq data computed by the top 30 genes of canonical correlation vector that contributed most to the separation of cardiopharyngeal identities in both species. The scaled and centered expression values for these 30 genes were then used to computed a 2-dimension tSNE

embedding for each species separately. Cell classifications are as determined in the original studies of Ciona and Mouse.
(**b**) Box plot showing the classified contribution to each clusters in Ciona and Mouse.
(**c**) tSNE plots of of Ciona and Mouse scRNA-seq data as described in (a), with the expression patterns of *Ebf1* and *Gata4*.

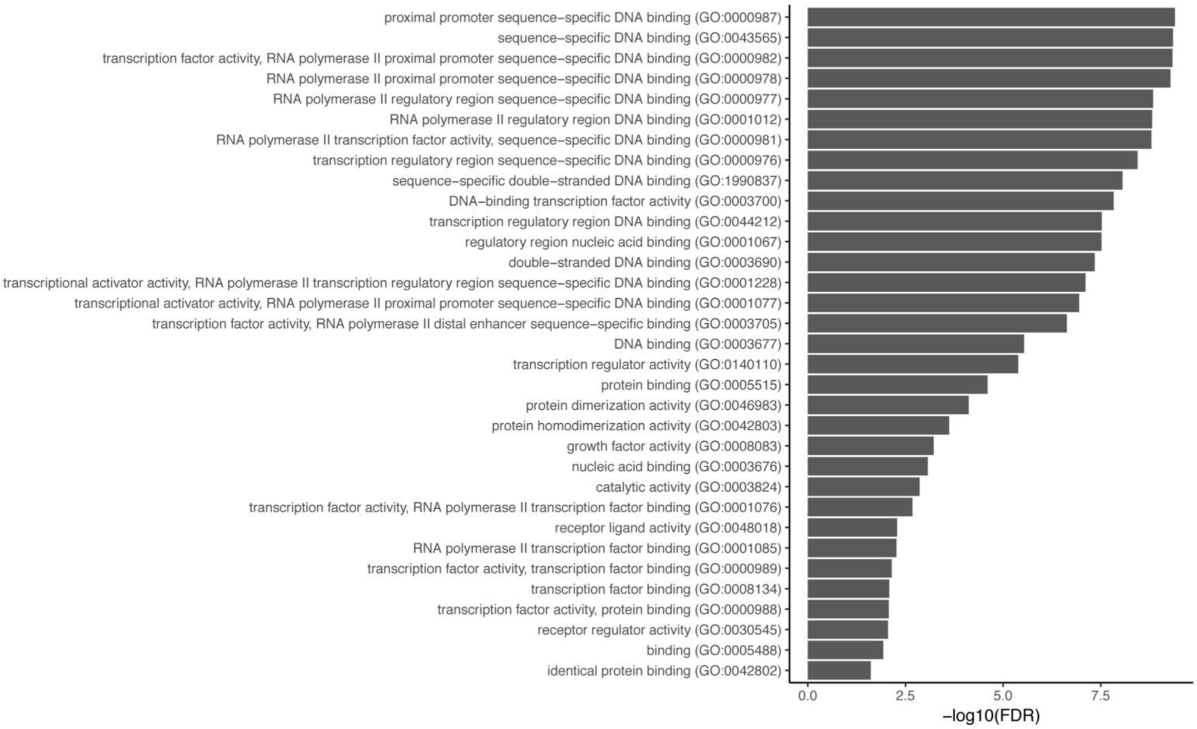


**Extended Data Figure 11| Gene ontology enrichment analysis of conserved cardiopharyngeal markers.**
The top CC gene set is significantly enriched in genes encoding transcription factors and DNA binding proteins.

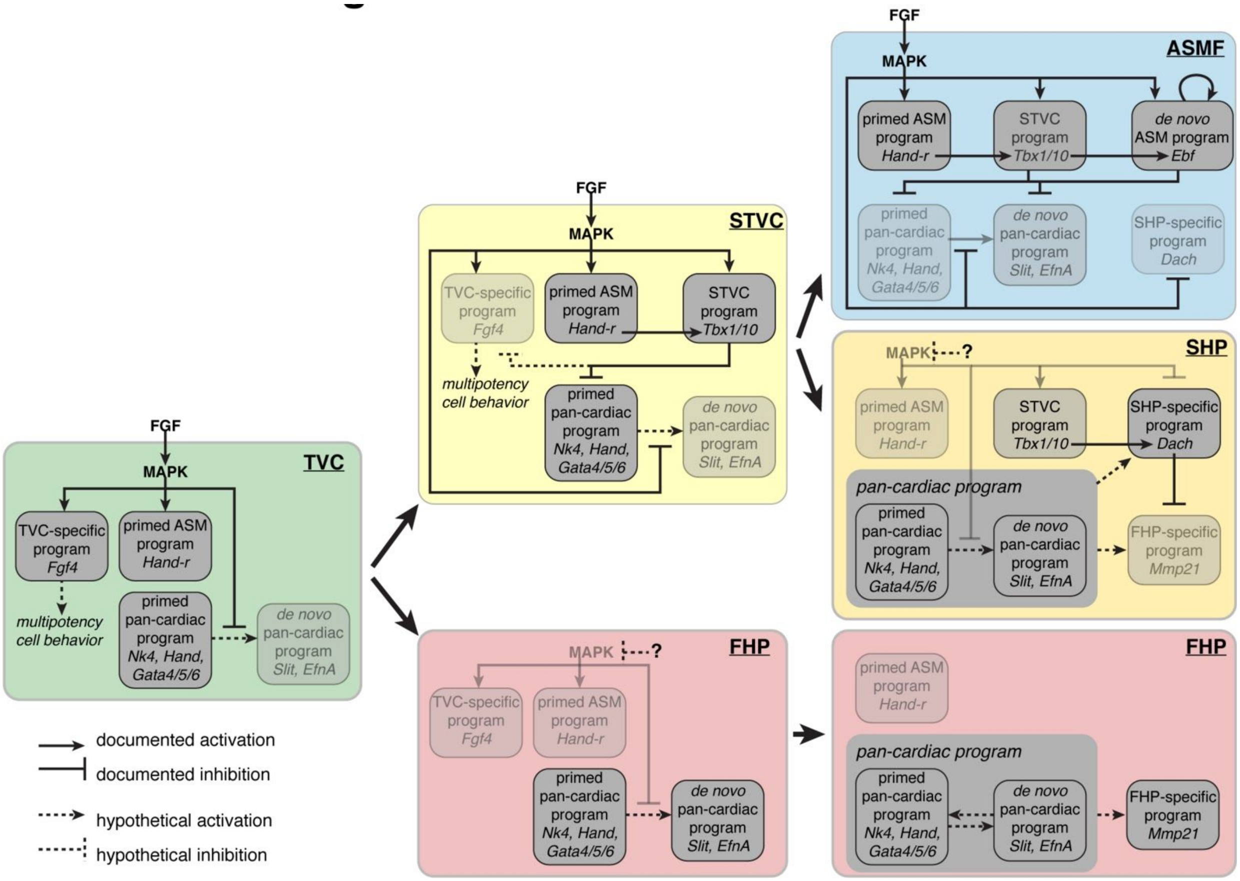


**Extended Data Figure 12| Summary model.**

Summary model showing the maintenance and progressive restriction of FGF-MAPK signaling in the multipotent progenitors (TVCs, trunk ventral cells, and STVCs, second trunk ventral cells) and atrial siphon muscle founder cells (ASMFs). Inhibition of MAPK activity permit the deployment of *de novo*-expressed pan-cardiac genes in both cardiac lineages (FHP, first heart precursors, and SHP, second heart precursors). FHPs specifically activate genes like *Mmp21*, and later produce most *Mhc2*+ cardiomyocytes, whereas SHPs descend from *Tbx1/10*+ multipotent progenitors, and thus activate *Dach*, which contributes to inhibiting the FHP-specific program. See discussion and Razy-Krajka et al., 2018[37] for details.)

**Extended Data Table 1. Developmental stages in Ciona.**
Hours after fertilization (hpf) at 18 degree centigrade to FABA[38] stage conversion.

**Extended Data Table 2. Top predicted differentially expressed marker genes across different cardiopharyngeal cell types.**
The first 7 tabs list candidate marker genes for indicated cell identities: TVC-specific, 12 hours post-fertilization (hpf), defined by anti-correlation with late-expressed genes in each trajectory; FHP(14hpf) and STVC(14hpf) defined by differential expression from the 14hpf scRNA-seq dataset; ASM, Pan-cardiac, FHP and SHP defined by differential gene expression from the 18 and 20hpf scRNA-seq dataset as described.
This table also includes the Ebf downstream target genes predicted by Razy-Krajka et. al[4] . In these tabs, "Foxf>Ebf " refers to a misexpression construct, FoxF>Ebf::WRPW to misexpression of a dominant-negative form of Ebf, with the DNA binding domain fused to the WRPW repressor motif.

**Extended Data Table 3. Validation of cell-type-specific gene expression by FISH assays for selected candidate cardiopharyngeal marker genes.**
Clone ID and Position in cDNA libraries as available on the ANISEED[30] database are indicated.

**Extended Data Table 4. Bulk RNA-seq upon the perturbation of FGF-MAPK signaling.**
TVC-specific Handr and Foxf enhancers were used to over-express $Fgfr^{DN}$ or $M\text{-}Ras^{CA}$, respectively, in order to inhibit or enhance the activity of the FGF-MAPK signaling in the TVCs and their progeny[37]. Average $Log_2$(FPKM) values from two biological replicates of RNA-seq were used to calculate the $log_2$(fold change) of the gene expression level compared to control cells, which were transfected with equal amounts of *Hand-r>* or *Foxf>LacZ* plasmids. The first tab provides summary statistics obtained by edgeR, alongside cell identities and primed/de novo assignments inferred from scRNA-seq.
The "Fig2h.SUMMARY.STATS" tab provide the summary statistics for t and Kolmogorov-Smirnov (KS) tests performed on the data presented in Fig.2h.
The "EXPRESSED.GENES.CLASSIF" tab summarized the classifications using for subsequent enrichment analyses presented in subsequent tabs, named after the perturbation being tested, and summarized in Extended Data Figure 5j-q.

**Extended Data Table 5. Canonical correlation (CC) vector of genes separated cardiac and pharyngeal mesoderm cells in Ciona and Mouse.**
Gene loading and predicted cell-type-specific enrichment are indicated for each gene in the main "cardiopharyngeal" CCs, in each one of the three comparisons (Ciona dataset compared to selected mouse datasets from Scialdone et al., 2016; Ibarra-Soria et al., 2018, or Lescroart et al., 2018).

**Extended Data Table 6. Ciona to Mouse gene name dictionary.**
Including orthologs and Best Blast Hits (See methods for details).

**Extended Data Movie 1. Animated 3D projection of St. 38 juvenile heart.**
the Mesp>nls::LacZ+; Tbx1/10(3xT12)>H2B::mCherry+ (colocalized red and green): SHP-derived cells. Mesp>nls::LacZ+;Tbx1/10(3xT12)>H2B::mCherry- (red): cells derived from both FHPs and SHPs * (* a fraction of SHPs are not labelled by Tbx1/10(3xT12)>H2B::mCherry due to the mosaicism). The SHP-derived cells form the outer layer of the heart compartment, with FHP-derived cells attached inside.

**Extended Data Movie 2. 3D animation of Mesp>LacZ reporter labeled cells in St.38 juvenile.**
Mesp>nls::LacZ+ (red) labeled Heart, ASM, LoM and ATM cells. 3xT12>H2B::mCherry+ (green) labeled SHP derived heart cells and ASM derived atrial siphon ring.
