## Supplementary material for "A single cell transcriptional roadmap for cardiopharyngeal fate diversification": Markdown_Clustering

### Clustering Analysis

*Xiang Niu May 11, 2016*

1. Load packages and Data

```
library(Seurat)
library(RColorBrewer)
source("functions.R")
load("hpfall.remv1.Robj")
load("hpfall.remv2.Robj")
col = colorRampPalette(rev(brewer.pal(n = 10, name = "RdBu")))
```

2. hpf12

```
# Subset data from preprocessed Seurat object
hpf12 = subsetData(hpfall.remv2, which.cells(hpfall.remv2, "hpf12"), do.scale = F)
hpf12
```

```
## An object of class seurat in project allhpf
##  14864 genes across 27 samples.
```

```
# Find variable gene with 4 < Average expression and Dispersion > 2
hpf12 = mean.var.plot(hpf12)
```

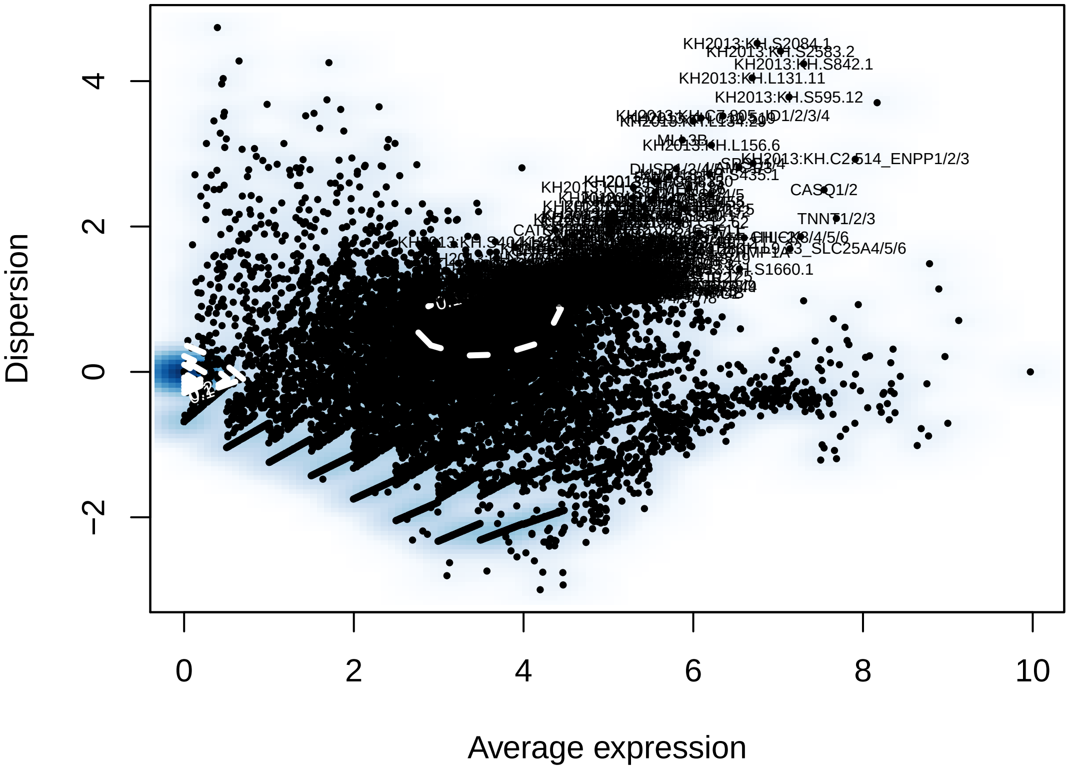

```r
length(hpf12@var.genes)
```

```
## [1] 428
```

```r
# Run a PCA using variable gene list
hpf12 = pca(hpf12, do.print = F)
pcScree(hpf12, hpf12@var.genes, 10)
```

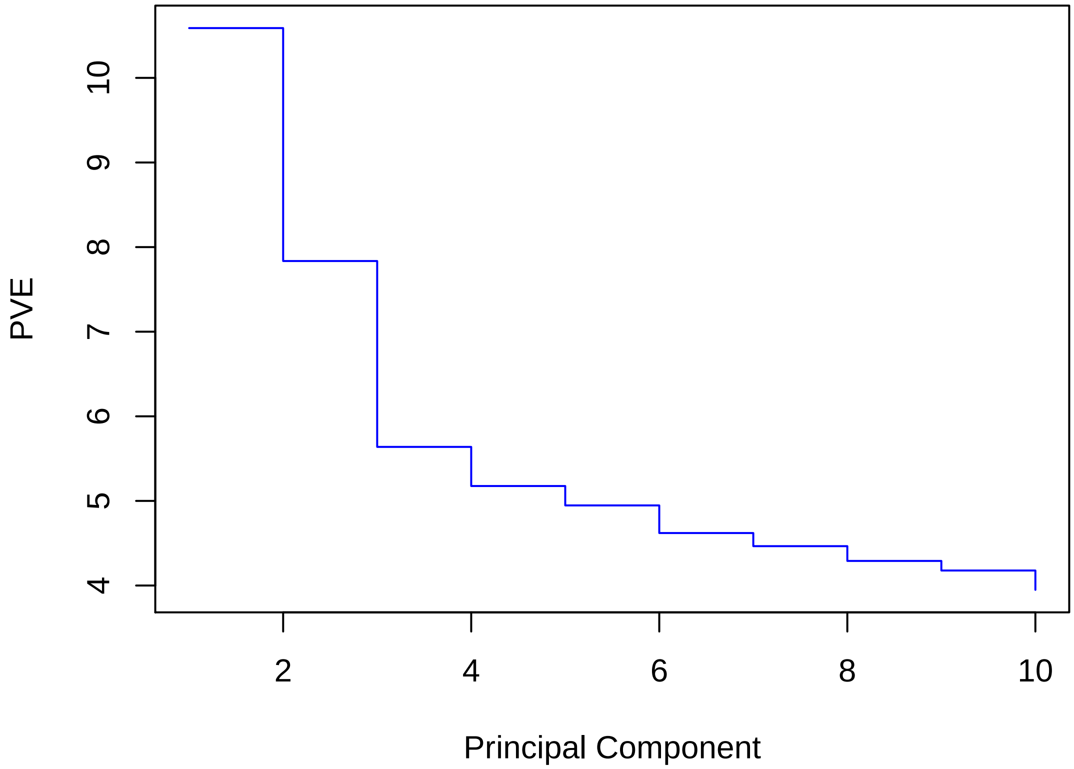


```r
pcHeatmap(hpf12, 1, do.balanced = T, col.use = col)
```

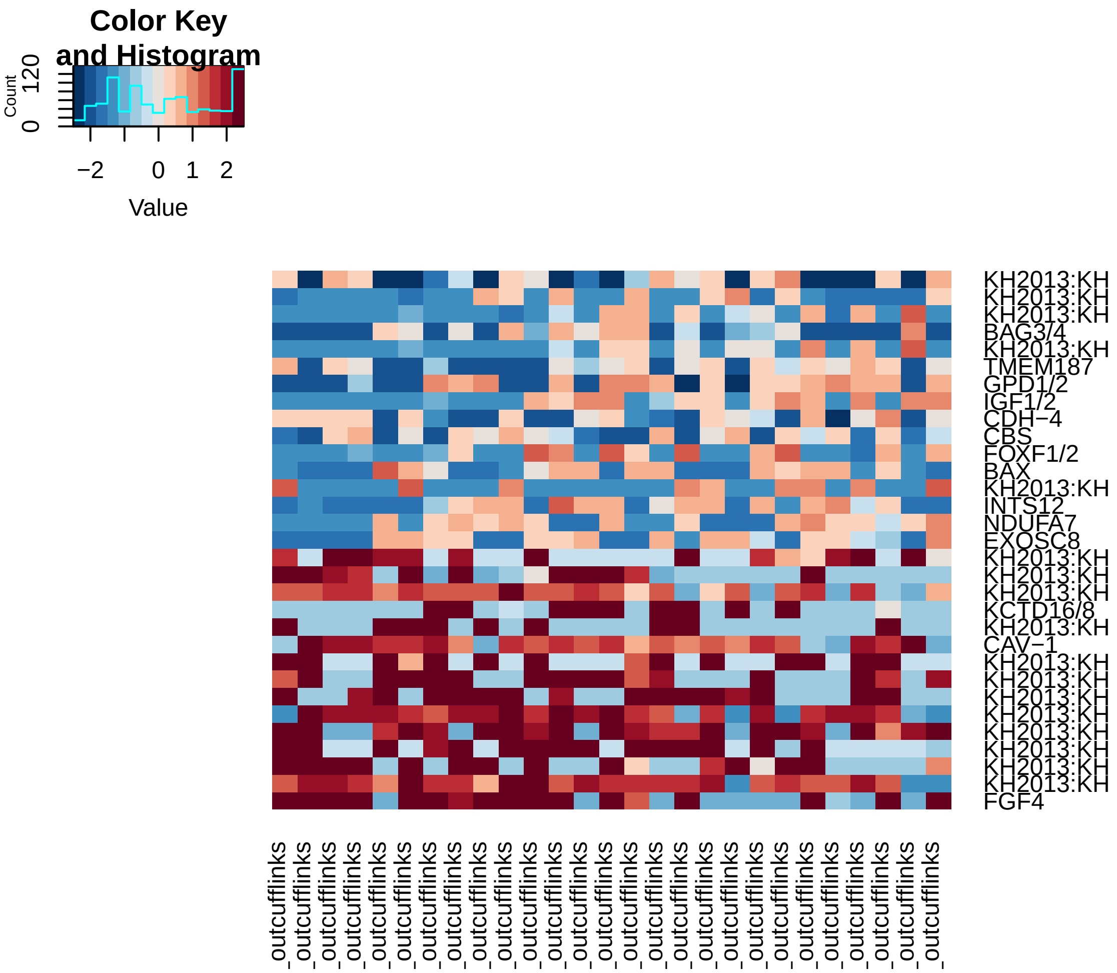


```r
pcHeatmap(hpf12, 2, do.balanced = T, col.use = col)
```

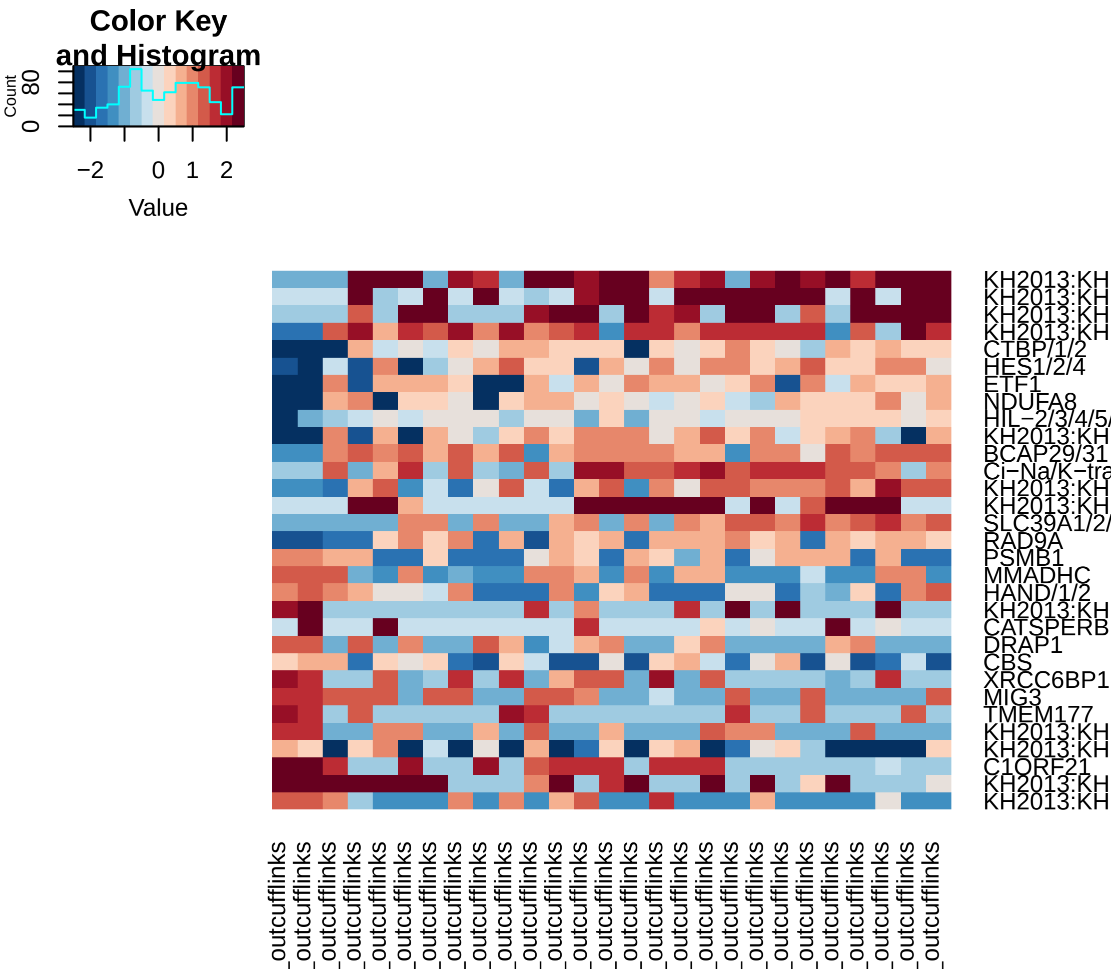


```
pcHeatmap(hpf12, 3, do.balanced = T, col.use = col)
```

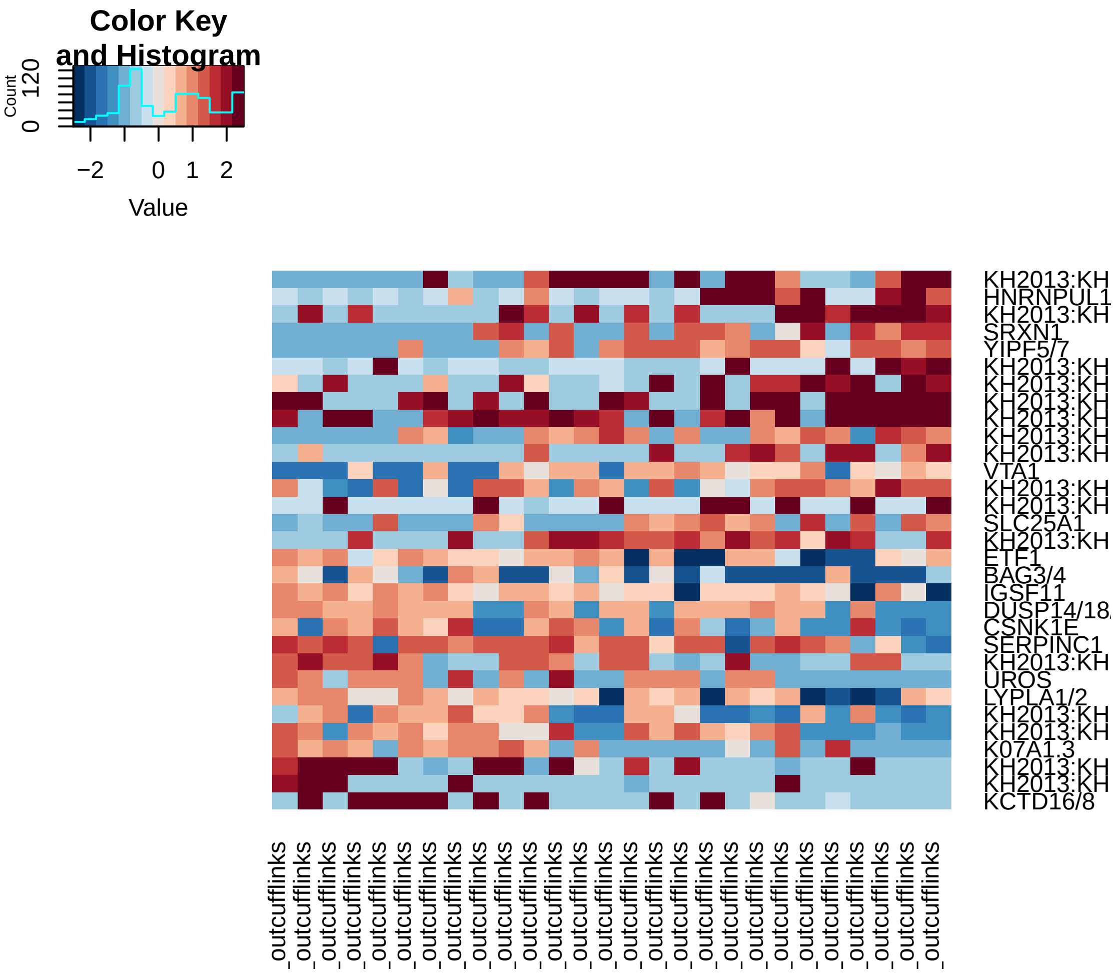


```r
pca.plot(hpf12, 1, 2)
```

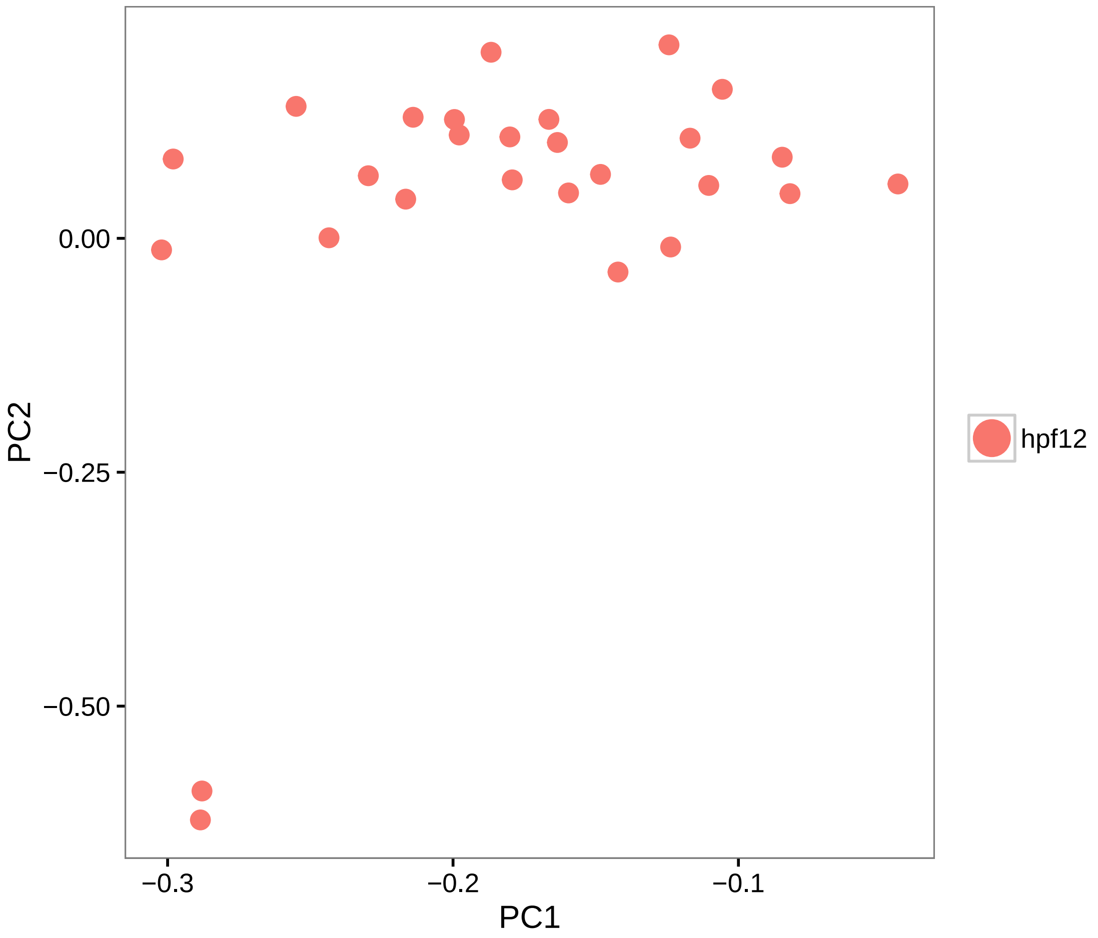


```
pca.plot(hpf12, 1, 3)
```

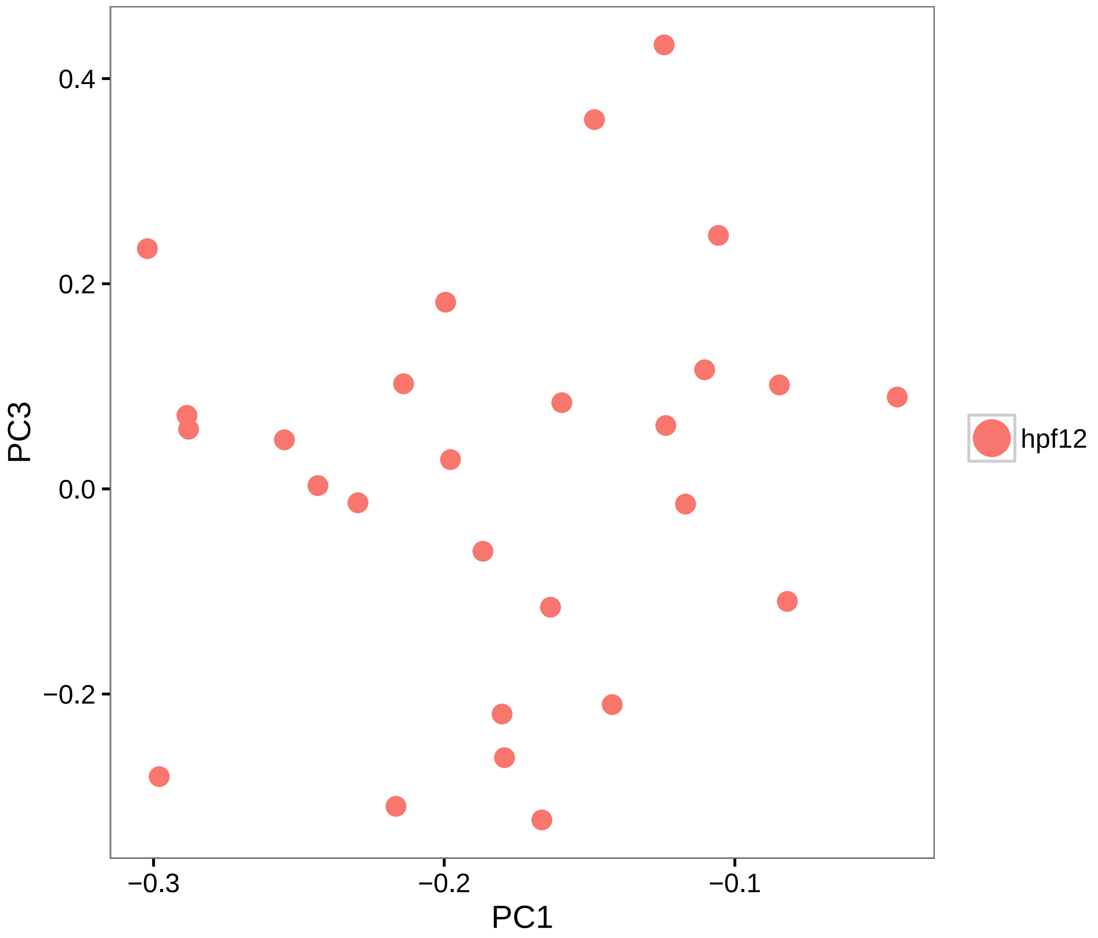


```
# Calculate PCA scores for all genes (PCA projection)
hpf12 = project.pca(hpf12, do.print = F)

# Visualize the full projected PCA, which now includes new genes which were
# not previously (use.full=TRUE)
pcHeatmap(hpf12, 1, use.full = T, do.balanced = T, col.use = col)
```

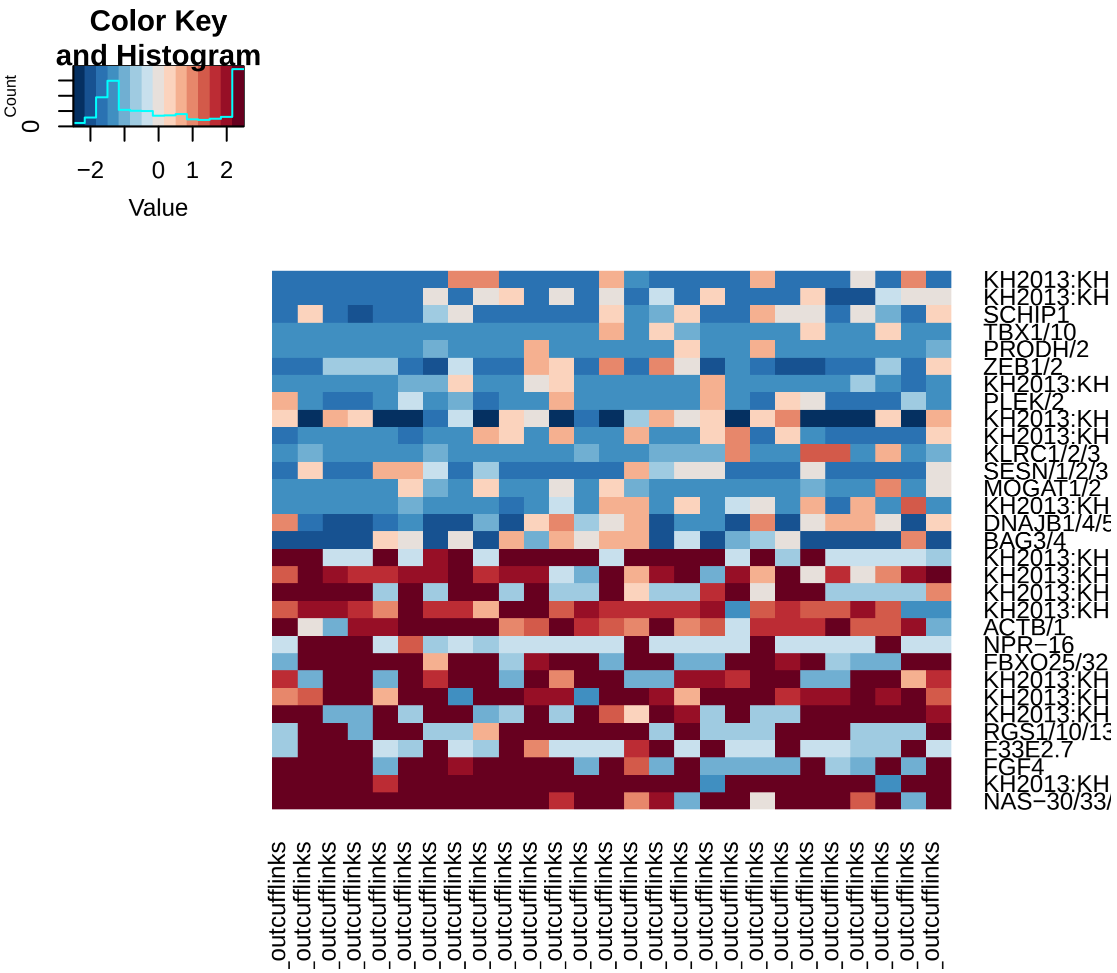


```r
pcHeatmap(hpf12, 2, use.full = T, do.balanced = T, col.use = col)
```

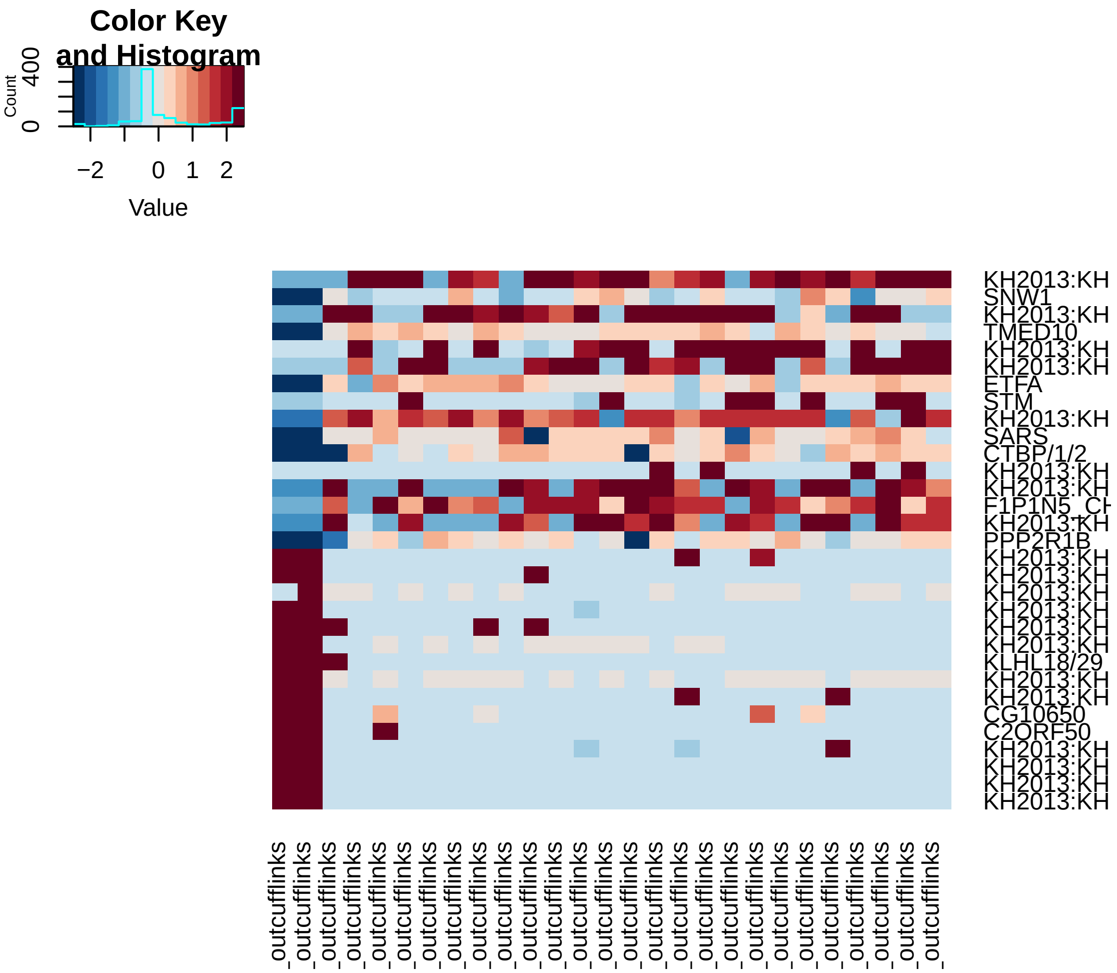


```
pcHeatmap(hpf12, 3, use.full = T, do.balanced = T, col.use = col)
```

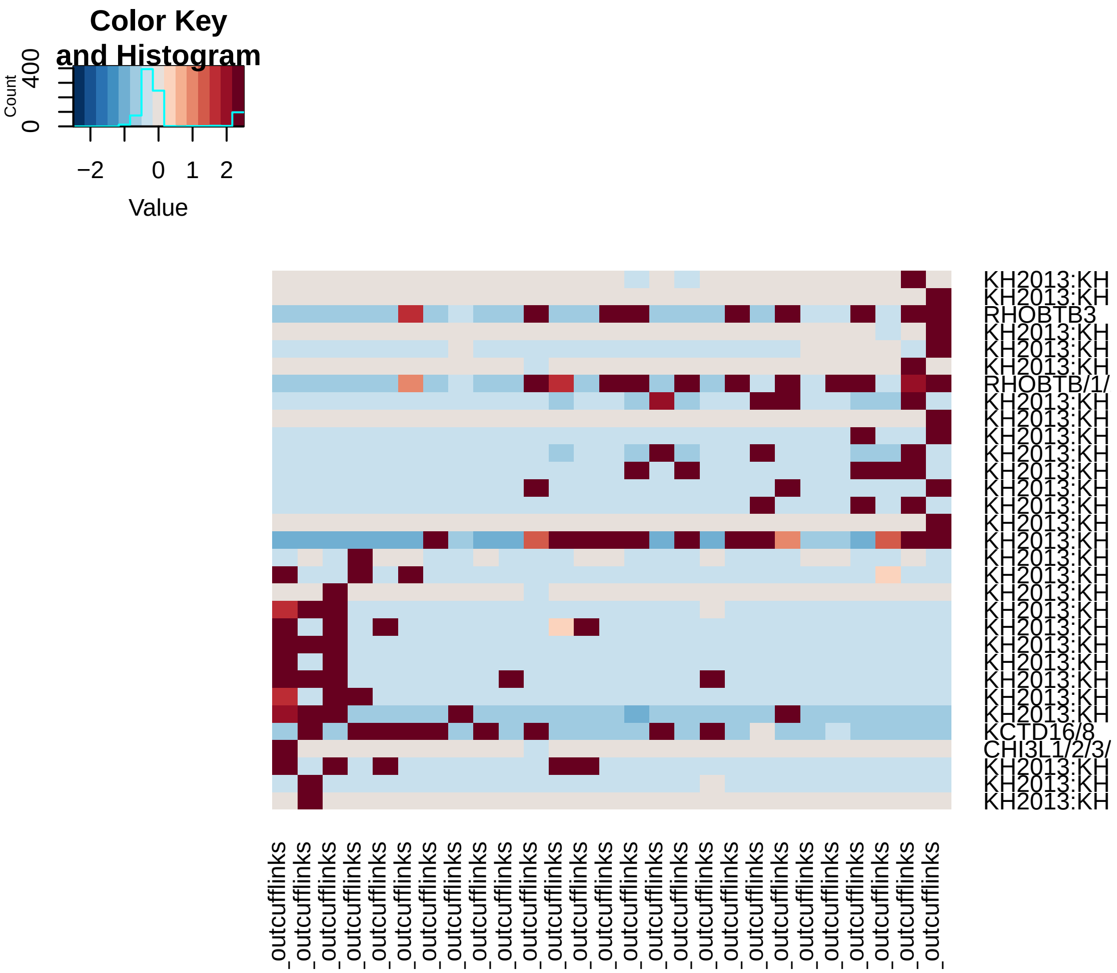


```
# Do 200 random samplings to find significant genes, each time randomly
# permute 1% of genes This returns a 'p-value' for each gene in each PC,
# based on how likely the gene/PC score woud have been observed by chance
# Note that in this case we get the same result with 200 or 1000 samplings,
# so we do 200 here for expediency (20 PCs is used for significant test)
hpf12 = jackStraw(hpf12, num.replicate = 200, do.print = F, num.pc = 20)

# The jackStraw plot compares the distribution of P-values for each PC with
# a uniform distribution (dashed line) 'Significant' PCs will have a strong
# enrichment of genes with low p-values (solid curve above dashed line)
jackStrawPlot.new(hpf12, PCs = 1:12)
```

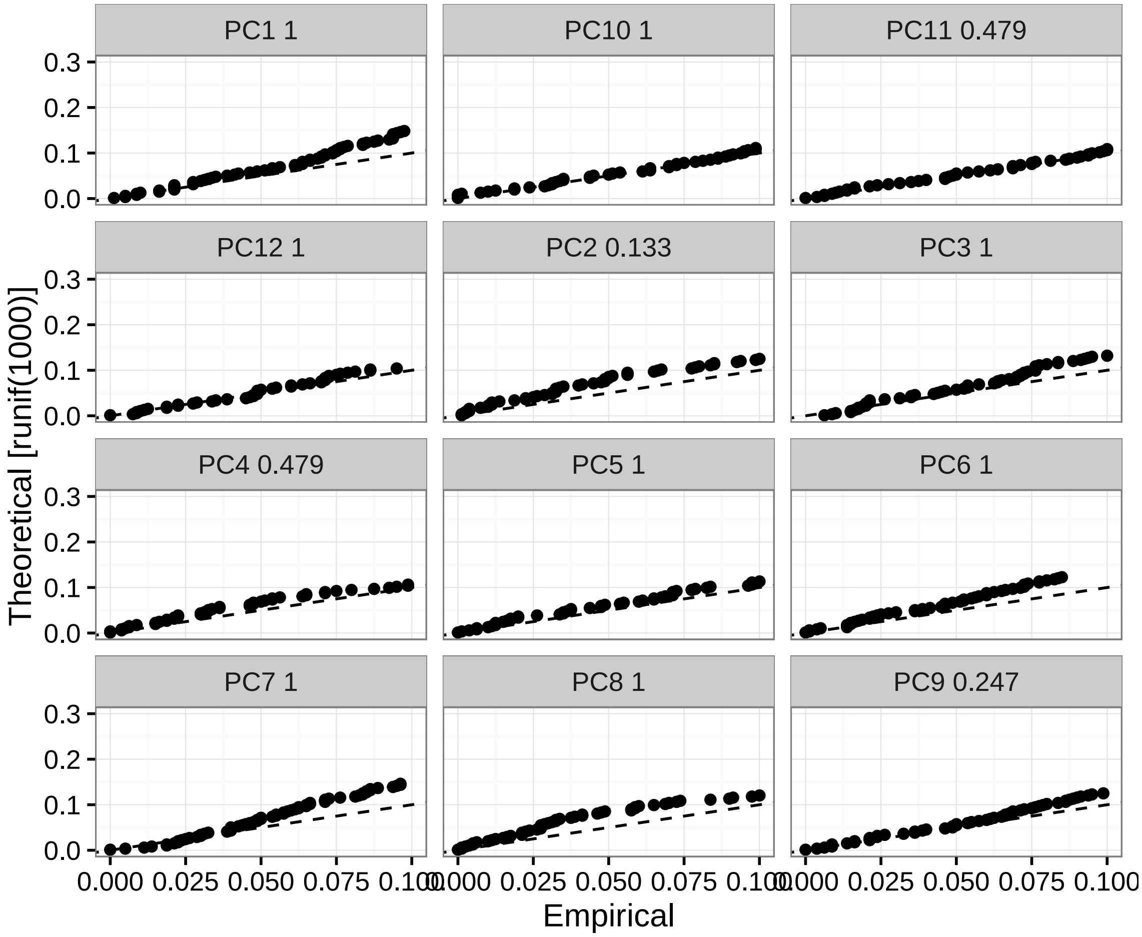


```r
# jackStraw plots show that none of PCs are significant enough, which
# suggests homogenous population.

# With preliminary studies these cells are 12hpf TVC cells
hpf12 = set.ident(hpf12, ident.use = "12TVC")

# Visualize known TVC cell markers
vlnPlot(hpf12, c("GATA4/5/6", "HAND/2", "HAND1/2", "NKX2-3"))
```

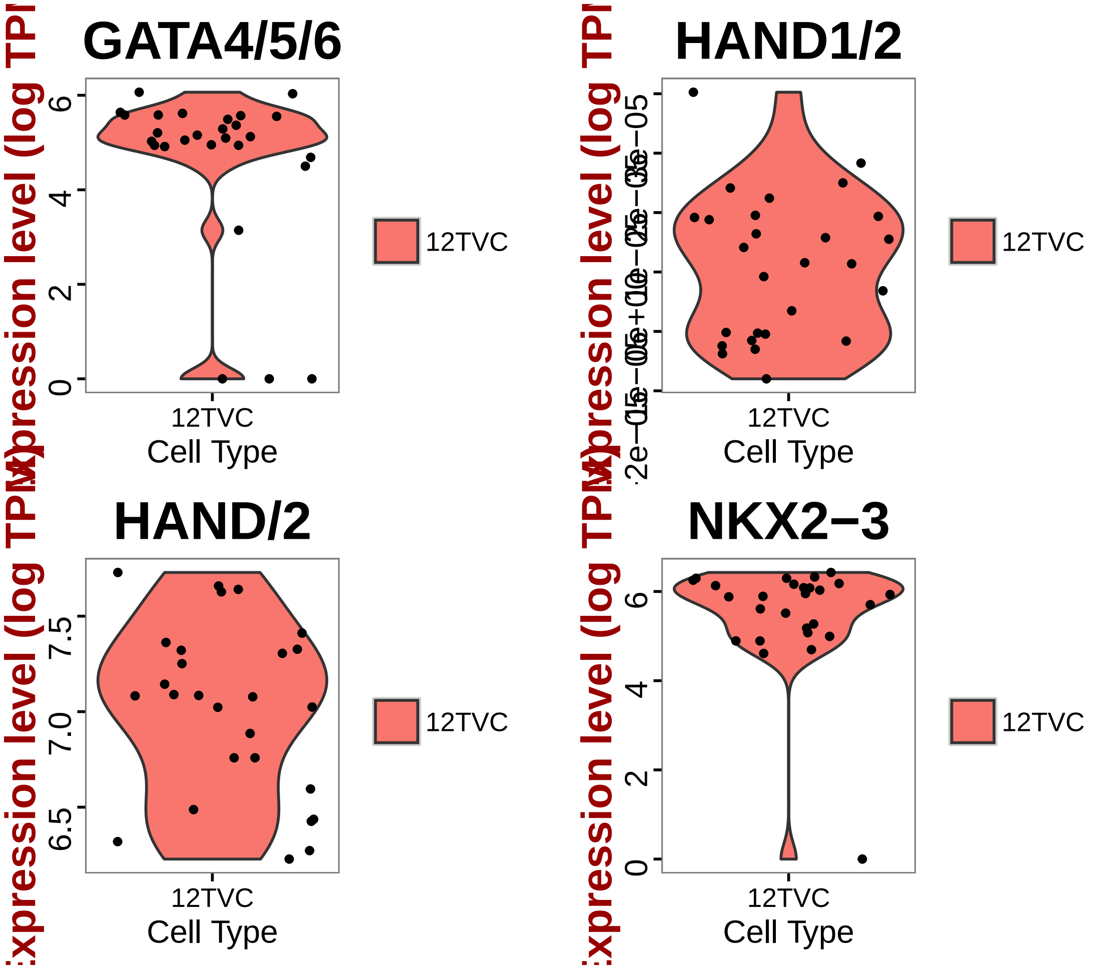


```r
# Write cell names into text files
write.table(hpf12@cell.names, file = "12TVCCells.txt", sep = "\t")

# Find 12TVC marker with 12 contamination cells
mesen.cell = as.character(unlist(read.table("mesen.cellname.txt")))
contam.cell = as.character(unlist(read.table("contam.cellname.txt")))
contam.name = grep("hpf12", c(mesen.cell, contam.cell), value = T)
hpf12.new = subsetData(hpfall.remv1, which.cells(hpfall.remv1, "hpf12"), do.scale = F)
hpf12.new = set.ident(hpf12.new, hpf12@cell.names, "12TVC")
hpf12.new = set.ident(hpf12.new, contam.name, "12Contam")
tvc.marker = find.markers(hpf12.new, "12TVC", "12Contam", thresh.use = 1, test.use = "ro
    min.pct = 0.5)
head(tvc.marker[order(tvc.marker$myAUC, decreasing = T), ], 20)
```

```
##                               myAUC avg_diff power pct.1 pct.2
## TCEAL3/5/6                    0.950 1.307940 0.900 1.000 0.959
## SMURF1/2                      0.946 2.091587 0.892 1.000 0.306
## DCBLD2                        0.946 1.868379 0.892 1.000 0.633
```

```
## KH2013:KH.C8.743           0.944 3.169373 0.888 0.926 0.143
## NAF1                       0.941 1.630562 0.882 1.000 0.510
## FARP1/2                    0.940 1.512813 0.880 1.000 0.633
## KH2013:KH.C2.118_ENPP1/2/3 0.933 1.845469 0.866 1.000 0.306
## NOLC1                      0.933 1.433875 0.866 1.000 0.755
## KH2013:KH.C2.514_ENPP1/2/3 0.925 1.807612 0.850 1.000 0.592
## KH2013:KH.C7.205_ASB2      0.923 1.886513 0.846 1.000 0.327
## KH2013:KH.L153.32_PPP1R9A  0.923 1.222093 0.846 1.000 0.694
## NOP5/58                    0.923 1.676462 0.846 0.963 0.633
## UCHL1/3/4                  0.922 1.401090 0.844 1.000 0.571
## KH2013:KH.S2435.1_ASB2     0.921 1.701182 0.842 1.000 0.245
## GPX7/8                     0.917 1.600426 0.834 1.000 0.490
## DDX56                      0.915 1.689561 0.830 1.000 0.347
## KH2013:KH.C3.665_ZAN       0.915 1.392634 0.830 0.963 0.245
## SMG5                       0.914 1.680871 0.828 0.963 0.469
## KH2013:KH.L152.12_GAS2     0.912 1.503341 0.824 1.000 0.224
## KH2013:KH.L170.61_RRP9     0.911 1.619069 0.822 1.000 0.571
```

```
write.table(tvc.marker, "TVC.markers.txt", row.names = T, sep = "\t")

# Draw a heatmap of all cells for these marker genes
vlnPlot(hpf12, c("GATA4/5/6", "HAND/1/2", "NKX2-3", "FZD4"), cols.use = "green")
```

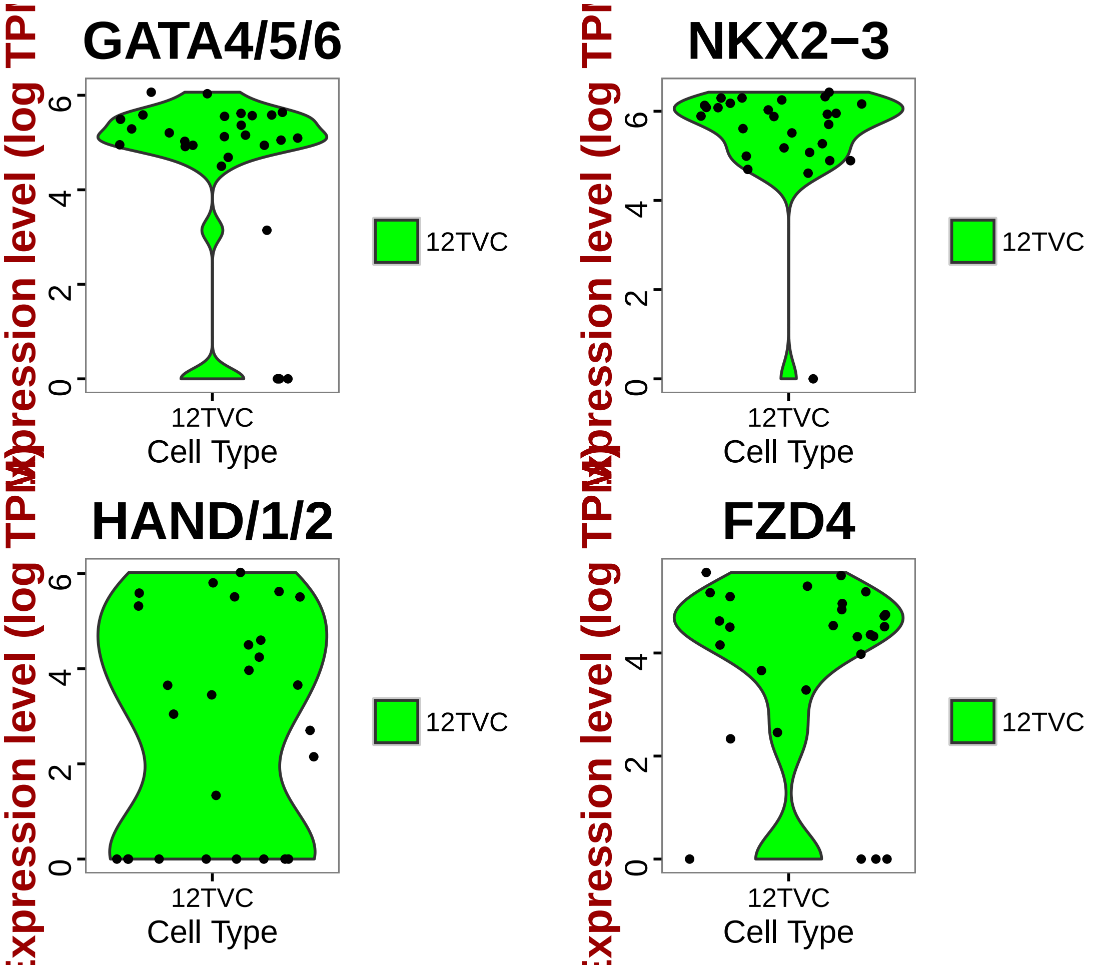


```
doHeatMap(hpf12, genes.use = rownames(head(tvc.marker[order(tvc.marker$myAUC,
    decreasing = T), ], 20)), remove.key = TRUE, slim.col.label = T, cex.col = 1.2,
    col.use = col, draw.line = F)
```

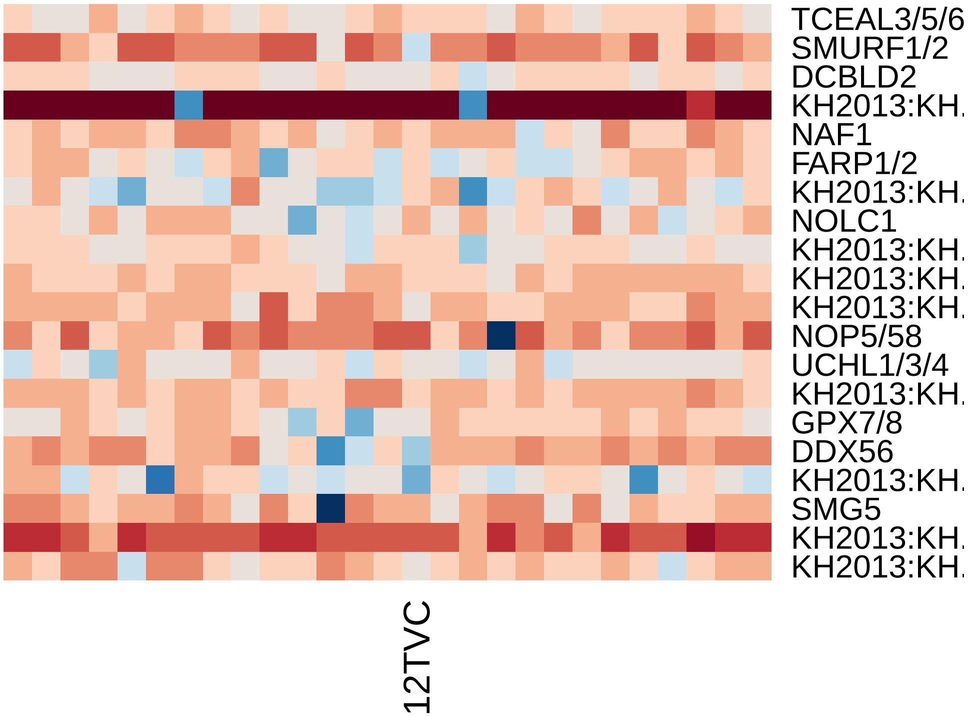


3. hpf14

```r
# Subset data from preprocessed Seurat object
hpf14 = subsetData(hpfall.remv2, which.cells(hpfall.remv2, "hpf14"), do.scale = F)
hpf14
```

```
## An object of class seurat in project allhpf
##  14864 genes across 275 samples.
```

```r
# Find variable gene with 4 < Average expression and Dispersion > 2
hpf14 = mean.var.plot(hpf14)
```

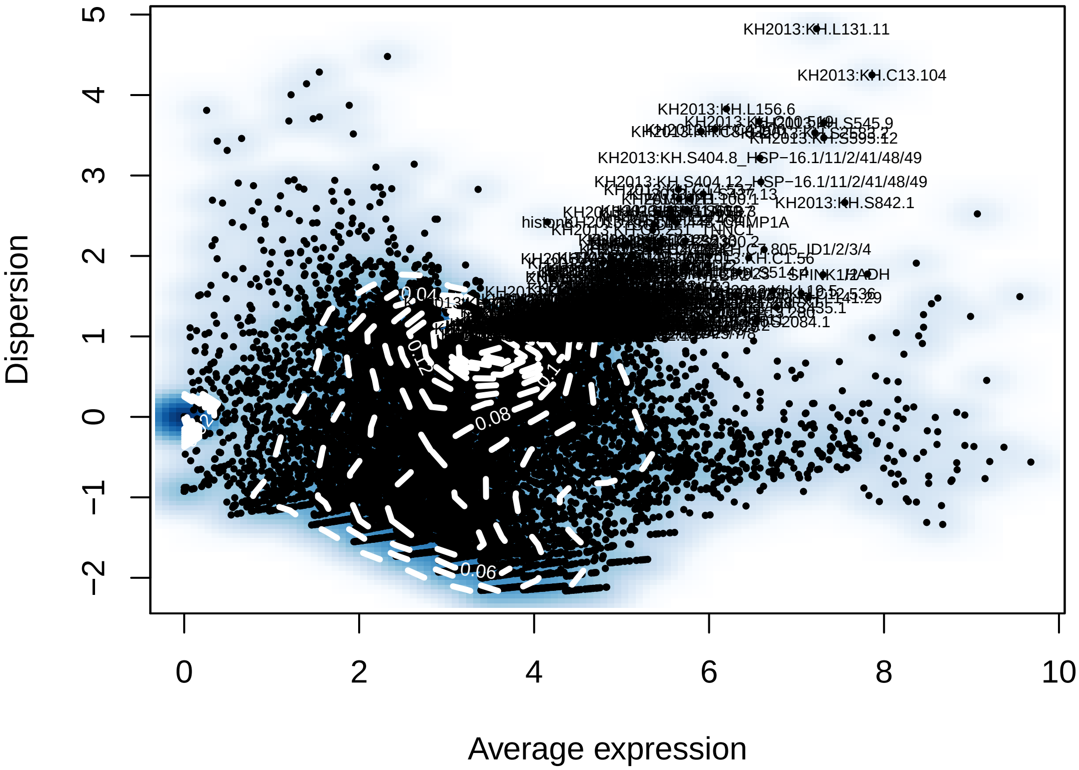


```r
length(hpf14@var.genes)
```

```
## [1] 384
```

```r
# Run a PCA using variable gene list
hpf14 = pca(hpf14, do.print = F)
pcScree(hpf14, hpf14@var.genes, 10)
```

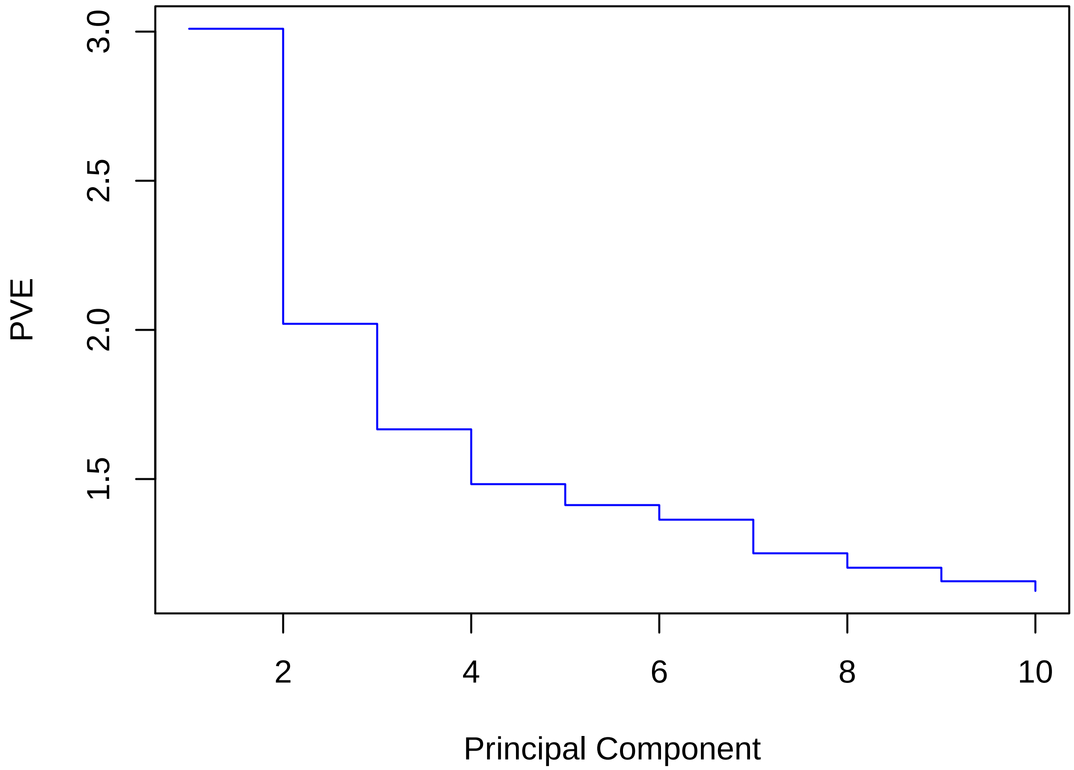


```
pcHeatmap(hpf14, 1, do.balanced = T, col.use = col)
```

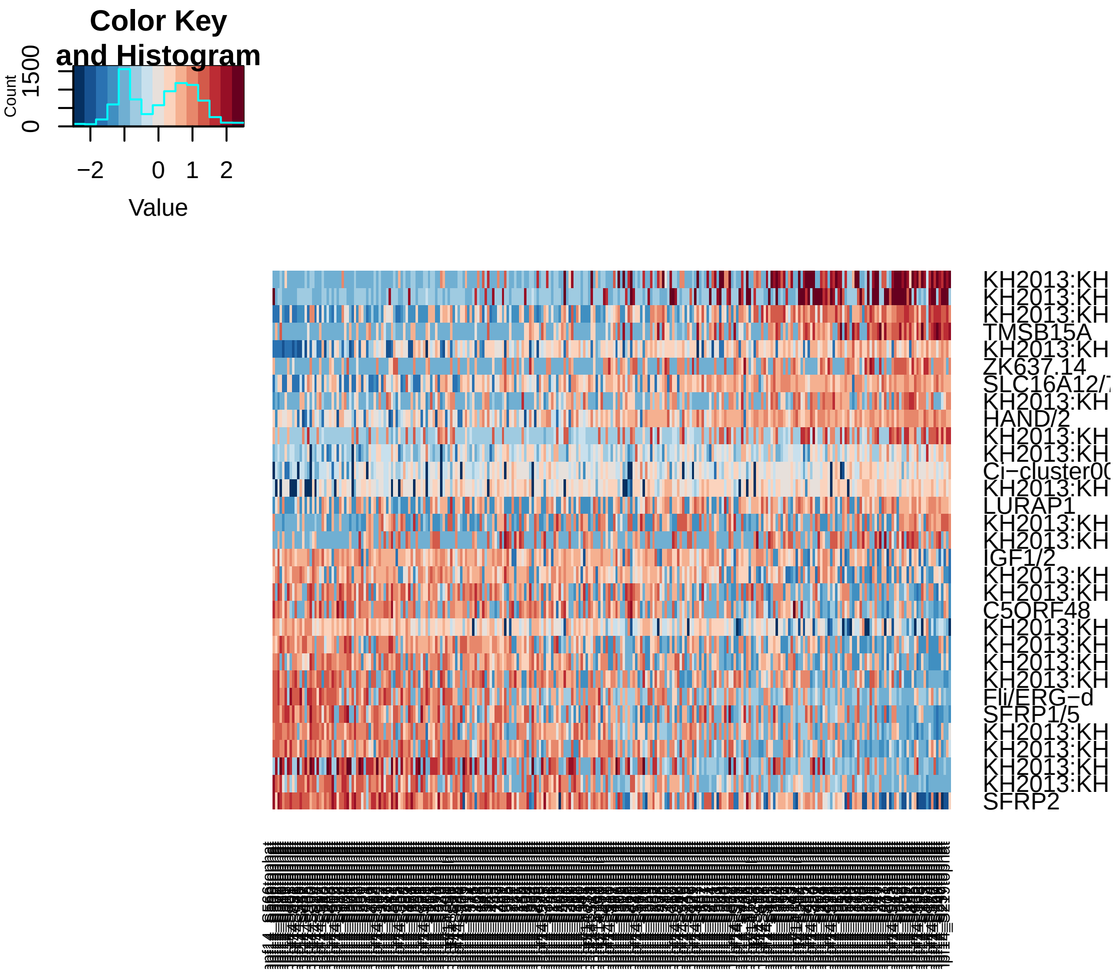


```r
pcHeatmap(hpf14, 2, do.balanced = T, col.use = col)
```

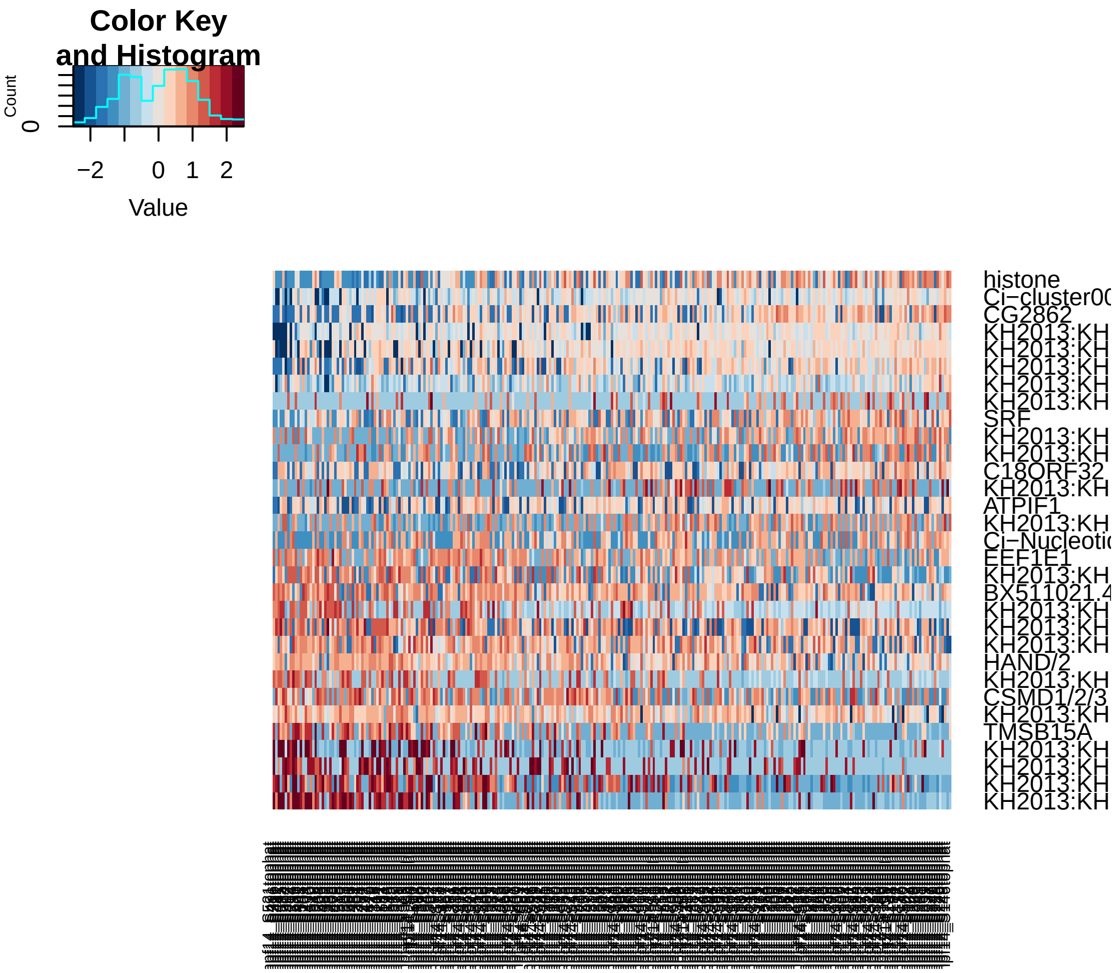


```r
pcHeatmap(hpf14, 3, do.balanced = T, col.use = col)
```

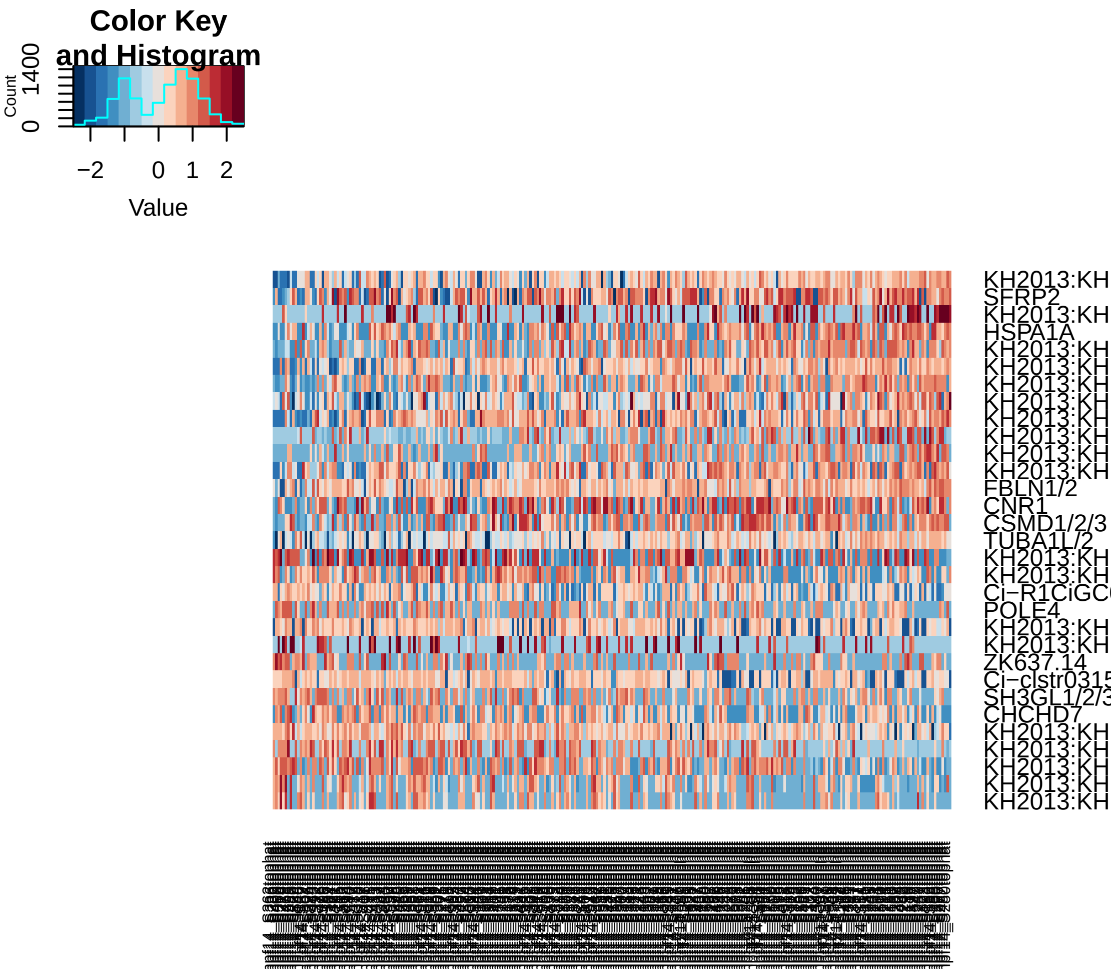


```r
pca.plot(hpf14, 1, 2)
```

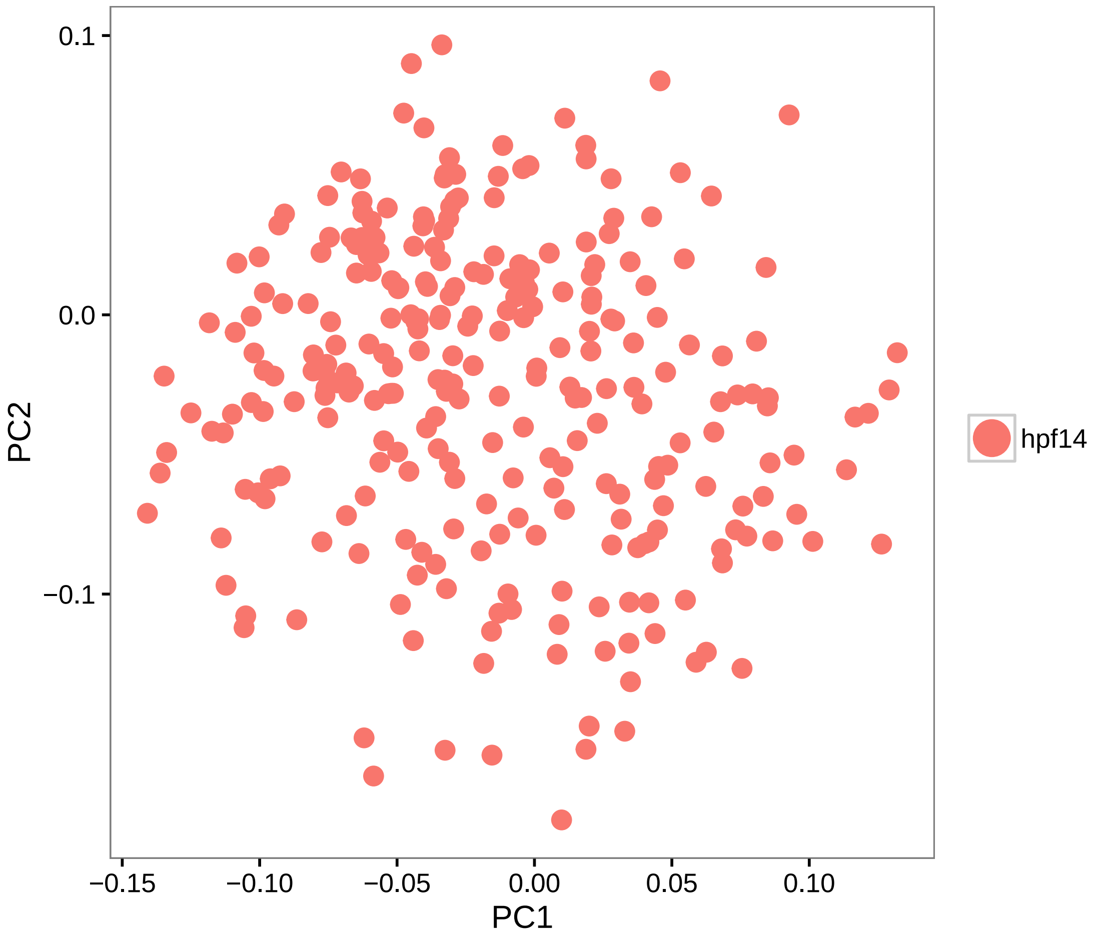


```
pca.plot(hpf14, 1, 3)
```

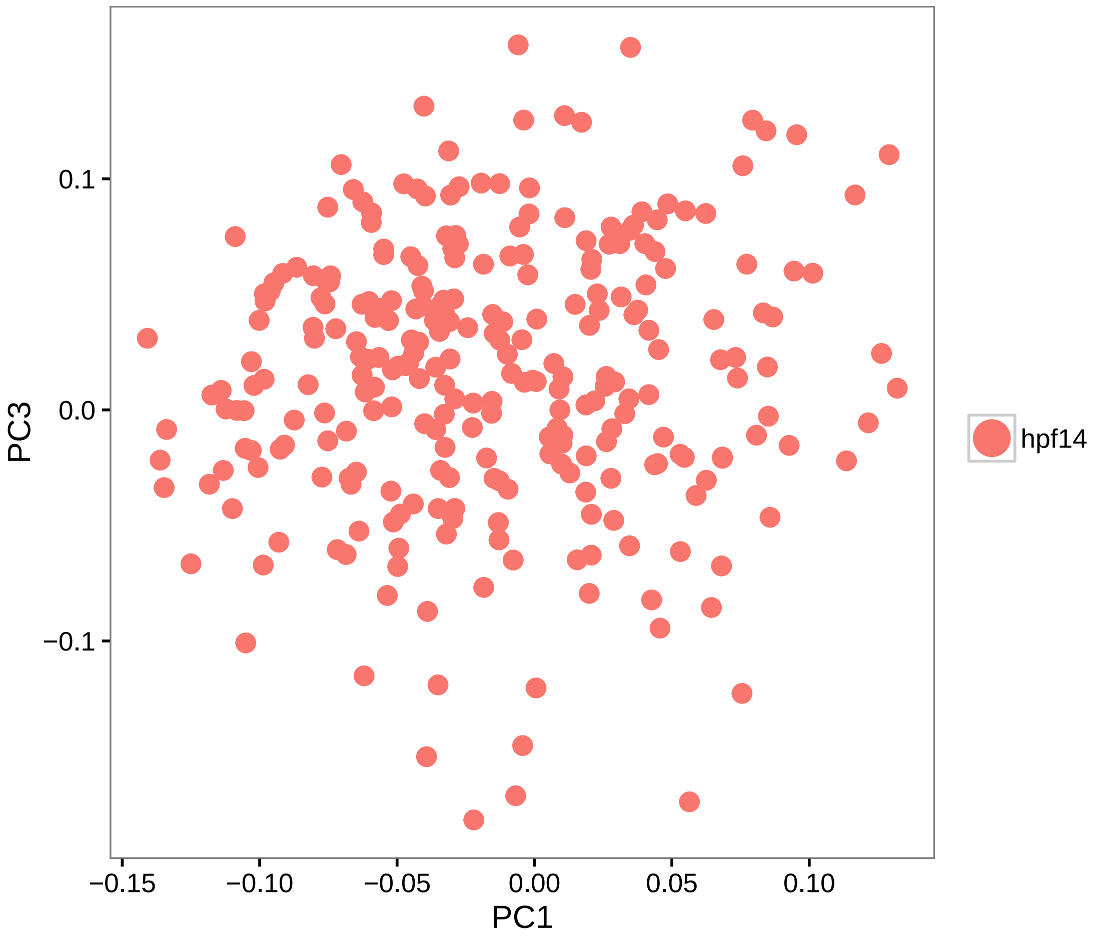


```
# Calculate PCA scores for all genes (PCA projection)
hpf14 = project.pca(hpf14, do.print = F)

# Visualize the full projected PCA, which now includes new genes which were
# not previously (use.full=TRUE)
pcHeatmap(hpf14, 1, use.full = T, do.balanced = T, col.use = col)
```

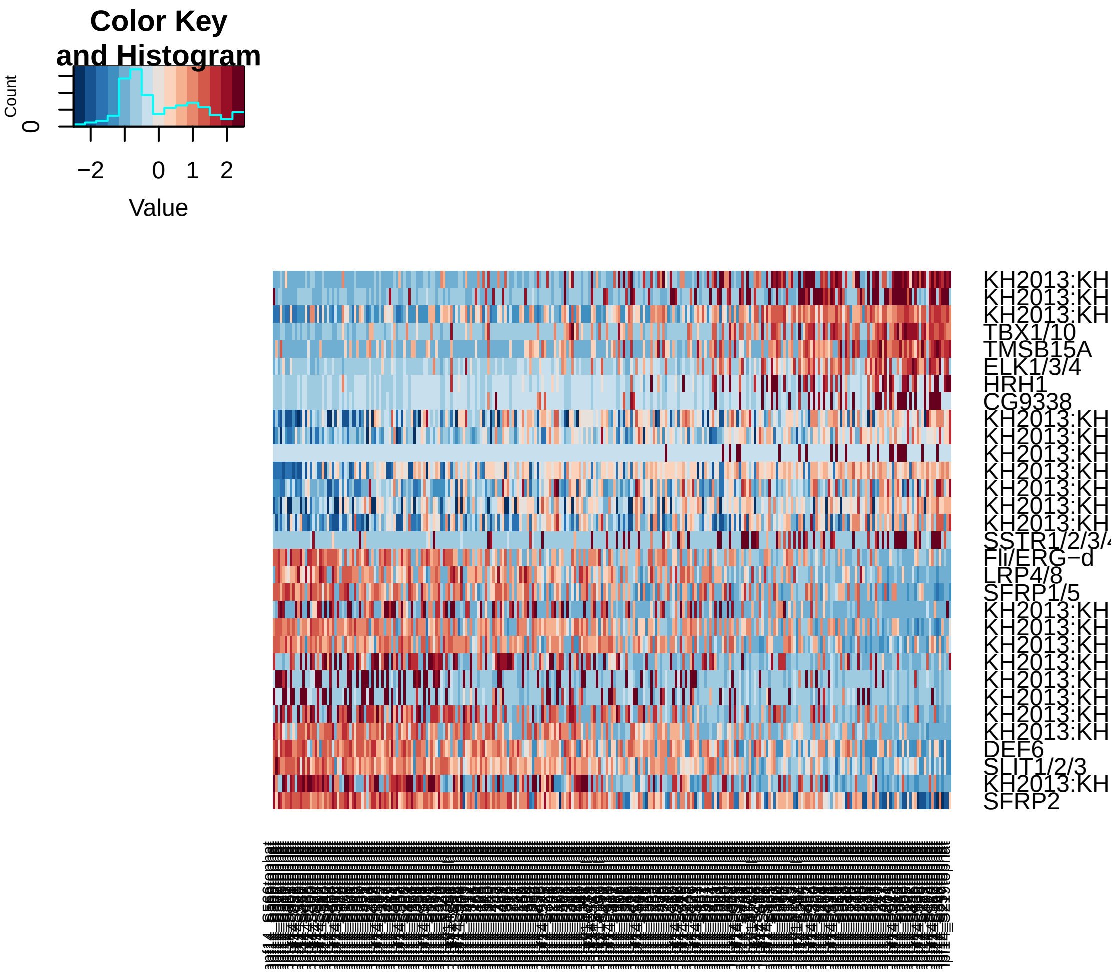


```r
pcHeatmap(hpf14, 2, use.full = T, do.balanced = T, col.use = col)  #technical
```

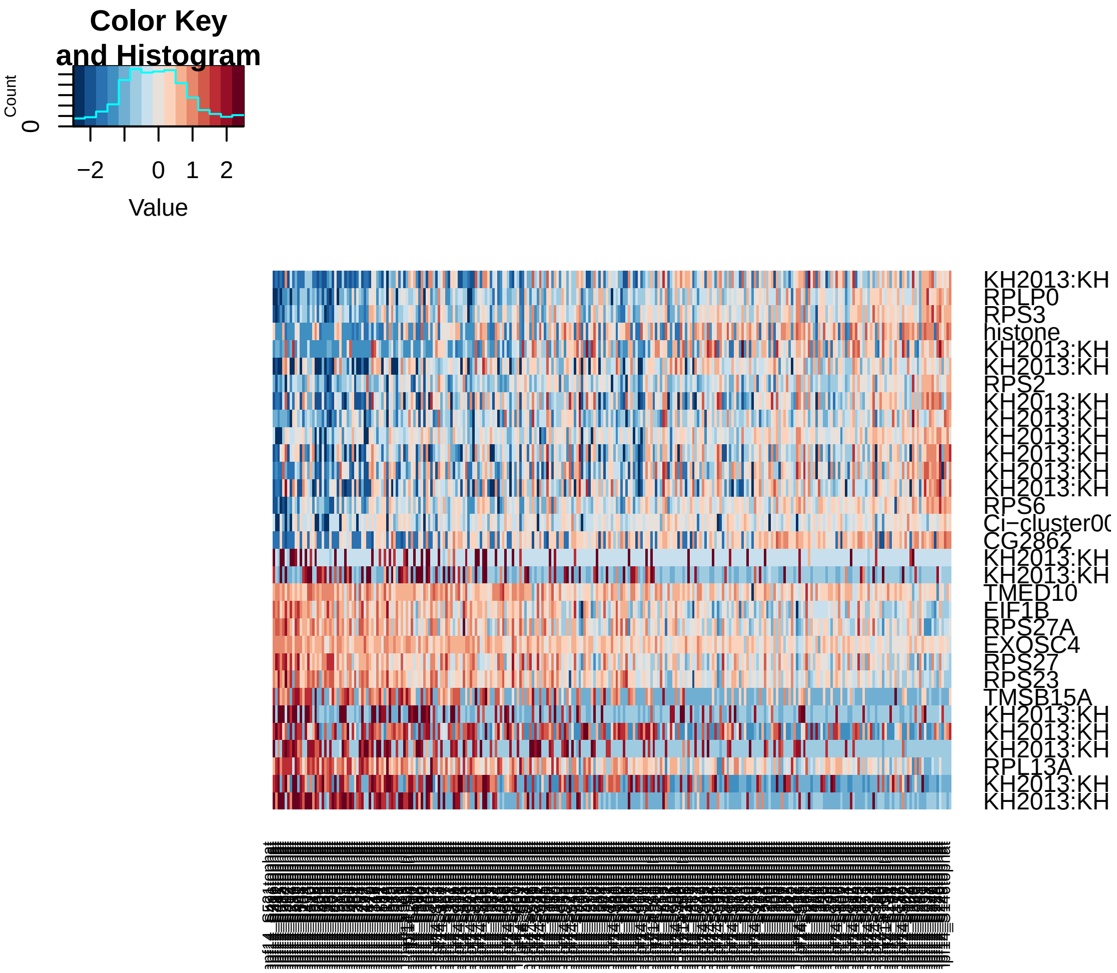


```
pcHeatmap(hpf14, 3, use.full = T, do.balanced = T, col.use = col)  #technical
```

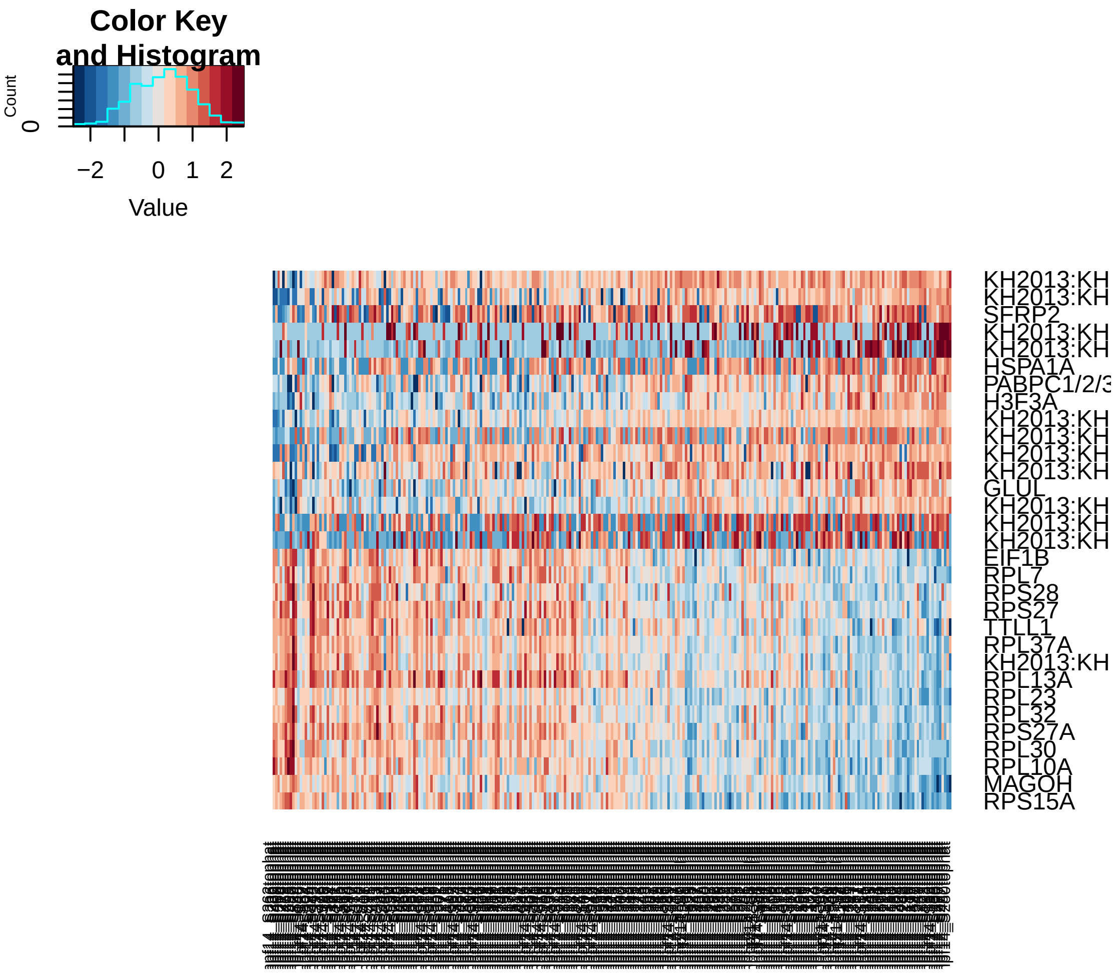


```
pcHeatmap(hpf14, 4, use.full = T, do.balanced = T, col.use = col)
```

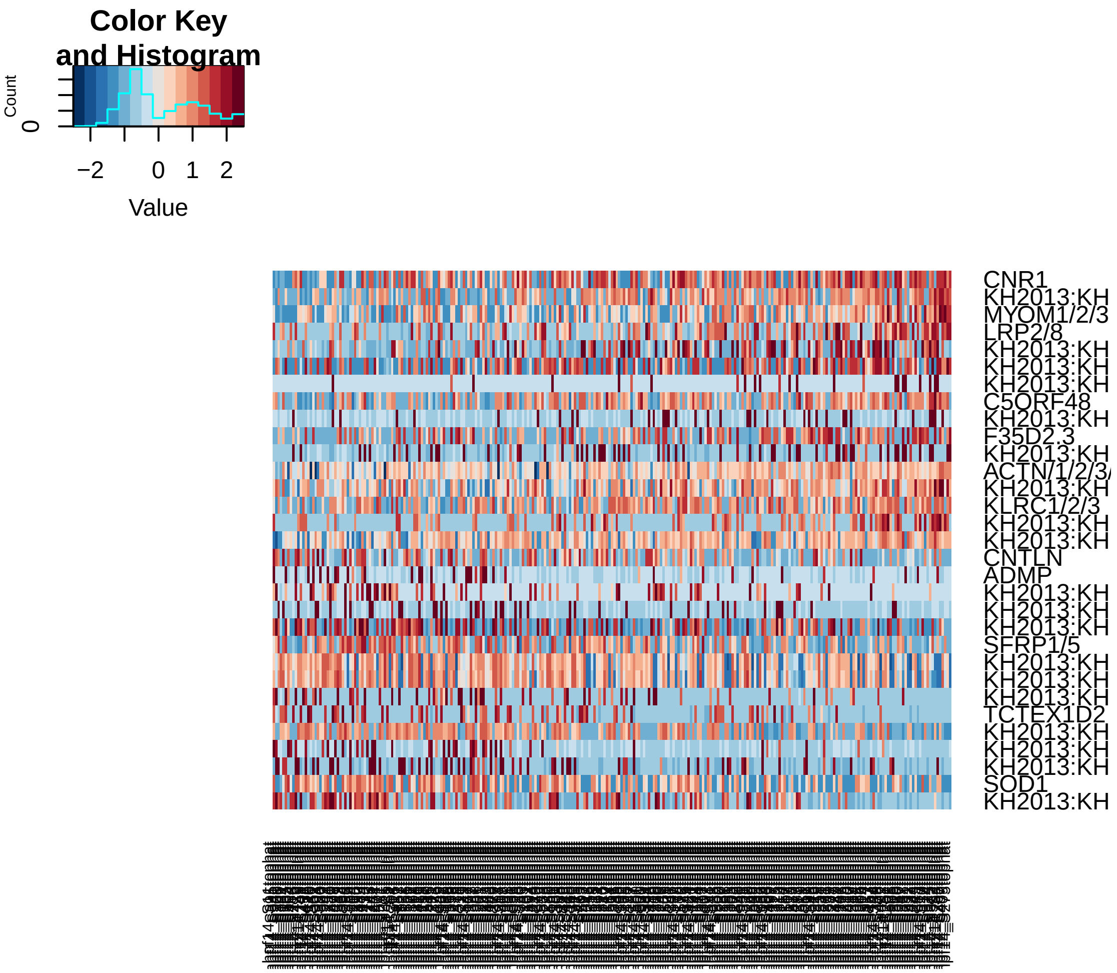


```r
pcHeatmap(hpf14, 5, use.full = T, do.balanced = T, col.use = col)
```

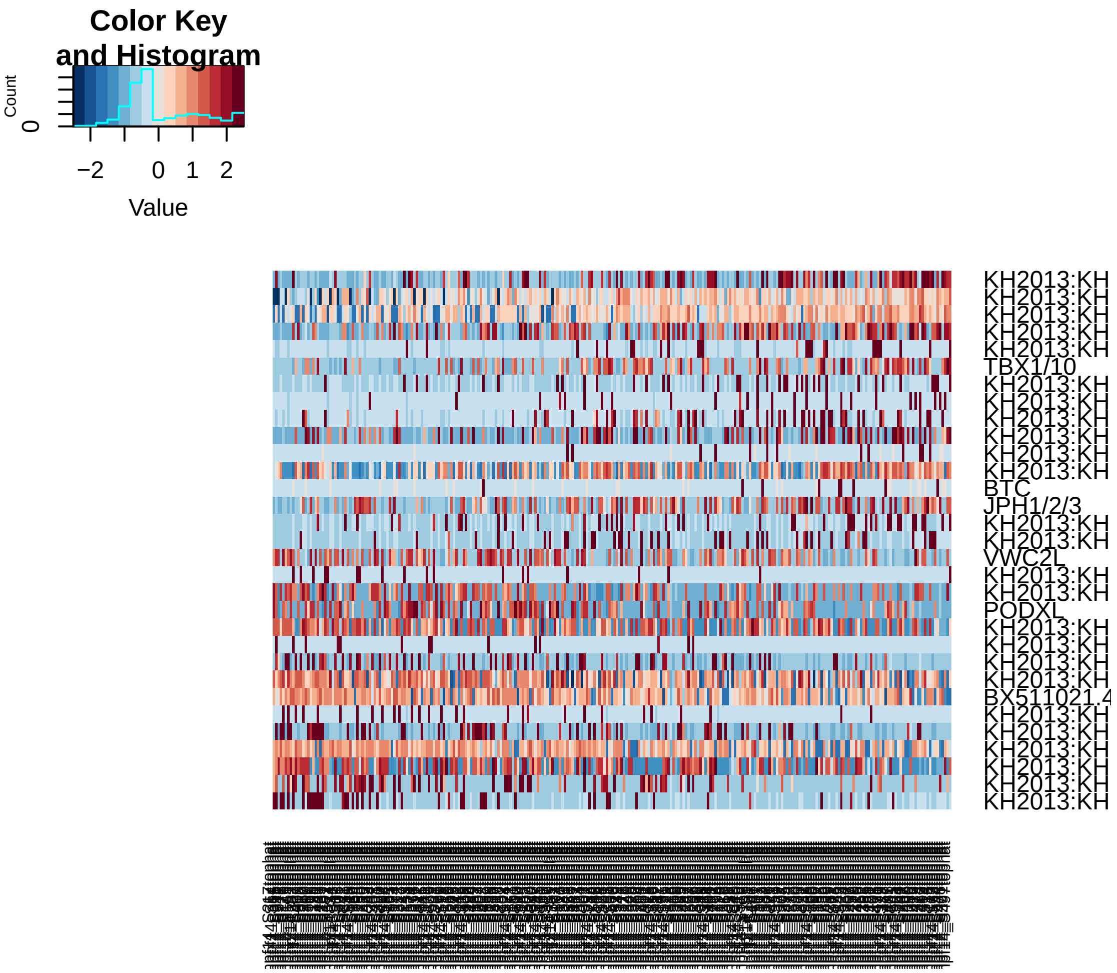


```r
# Do 200 random samplings to find significant genes, each time randomly
# permute 1% of genes This returns a 'p-value' for each gene in each PC,
# based on how likely the gene/PC score woud have been observed by chance
hpf14 = jackStraw(hpf14, num.replicate = 200, do.print = F)

# The jackStraw plot compares the distribution of P-values for each PC with
# a uniform distribution (dashed line) 'Significant' PCs will have a strong
# enrichment of genes with low p-values (solid curve above dashed line)
jackStrawPlot.new(hpf14, PCs = 1:12)
```

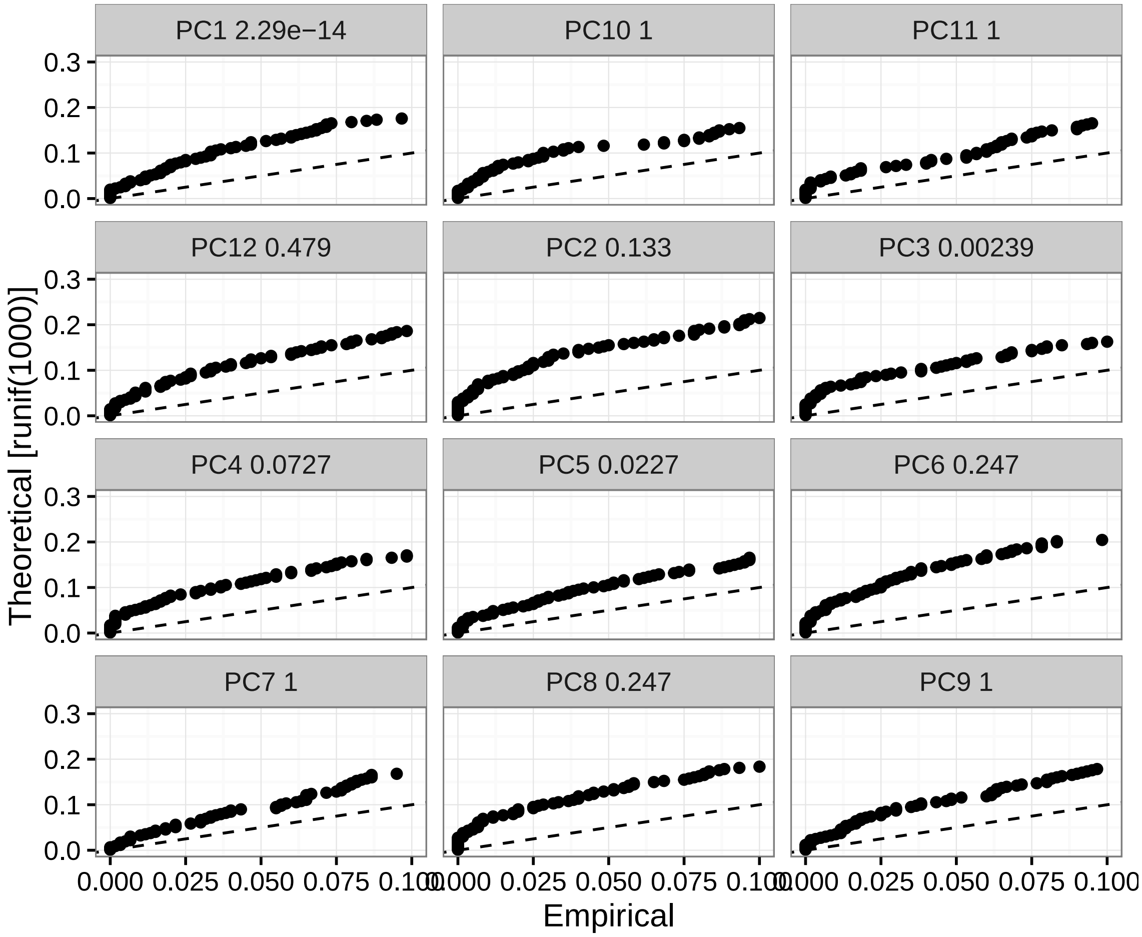


```
# In this case only PC1 is strongly significant and PC5 is significant, PC3
# though significant contains technical genes

# Run tSNE using significant PCs as input (spectral tSNE), we get distinct
# point clouds
hpf14 = run_tsne(hpf14, max_iter = 2000, dims.use = c(1, 5))
tsne.plot(hpf14, do.label = T, label.pt.size = 1)
```

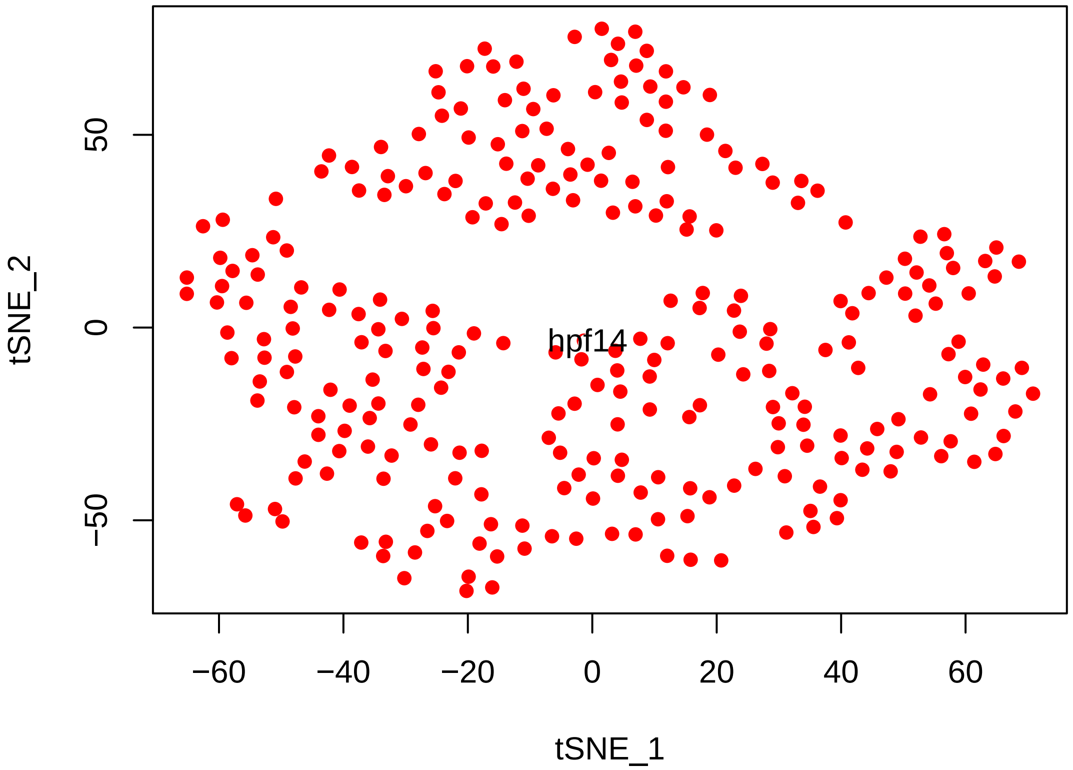


```r
# Find cell clusters using Modularity optimization cluster detection.
hpf14 = FindClusters(hpf14, pc.use = c(1, 5), do.modularity = T, resolution = 1,
    prune.SNN = 0.1, print.output = 0, k.param = 20)
```

```
## [1] "SNN : processed 69 cells"
## [1] "SNN : processed 138 cells"
## [1] "SNN : processed 206 cells"
## [1] "SNN : processed 275 cells"
```

```r
tsne.plot(hpf14, do.label = T, label.pt.size = 1)
```

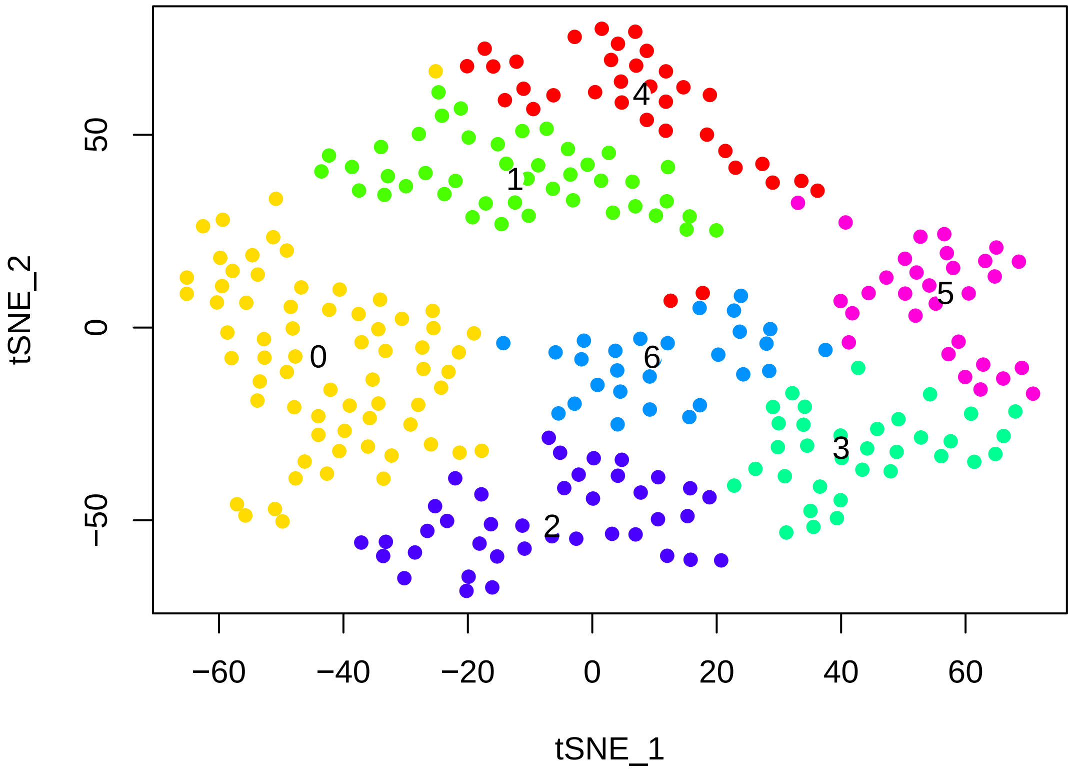


```r
# The validity of the clusters can be validated using a classification
# scheme based on linear SVMs.(In this case cutoff od 0.86 is selected to
# optimize clustering)
hpf14 = BuildSNN(hpf14, pc.use = c(1, 5), do.sparse = F, k.param = 20)
```

```
## [1] "SNN : processed 69 cells"
## [1] "SNN : processed 138 cells"
## [1] "SNN : processed 206 cells"
## [1] "SNN : processed 275 cells"
```

```r
hpf14 = ValidateClusters(hpf14, pc.use = c(1, 5), min.connectivity = 0.001,
    acc.cutoff = 0.85)
```

```
## [1] "  0% complete --- merge clusters 6 and 4, classification accuracy of 0.7087"
## [1] " 23% complete --- merge clusters 5 and 3, classification accuracy of 0.7495"
## [1] " 38% complete --- merge clusters 0 and 2, classification accuracy of 0.8438"
## [1] " 54% complete --- merge clusters 4 and 1, classification accuracy of 0.7940"
## [1] " 85% complete --- merge clusters 2 and 1, classification accuracy of 0.7865"
## [1] "100% complete --- started with 7 clusters, 2 clusters remaining"
```

```r
tsne.plot(hpf14, do.label = T, label.pt.size = 1)
```

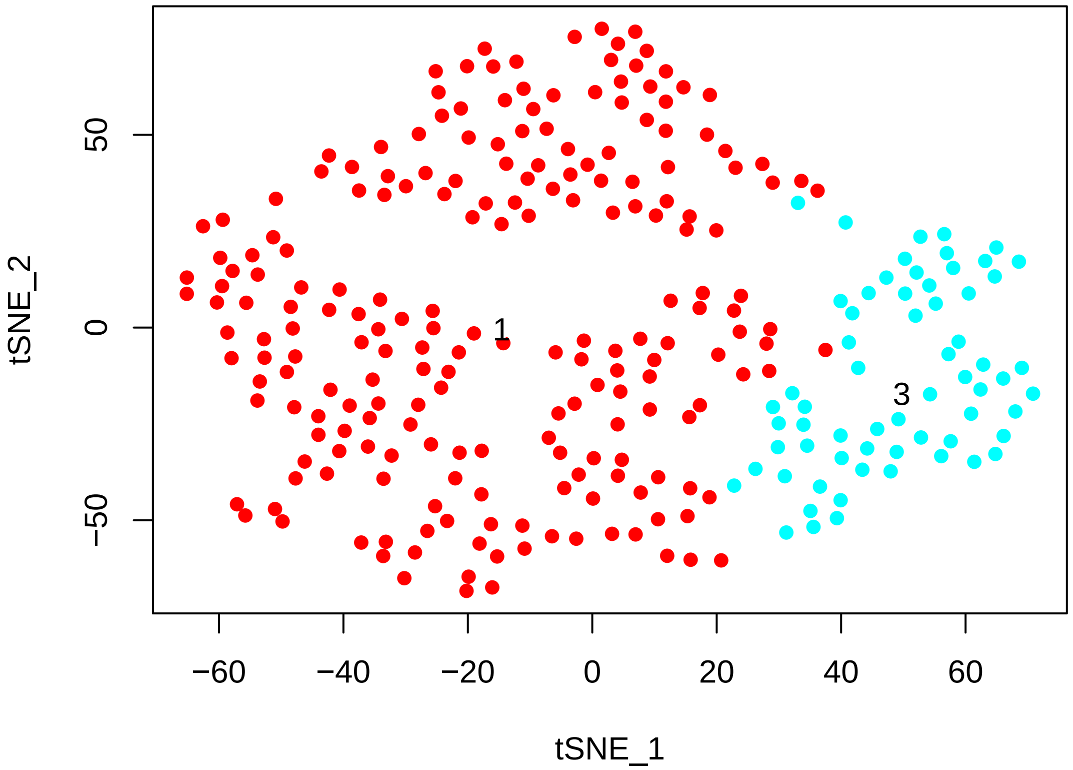


```
# Find cluster markers using ROC test with thresh.use = 1, min.pct = 0.5 The
# ROC test returns the 'classification power' for any individual marker
# (ranging from 0 - random, to 1 - perfect). Though not a statistical test,
# it is often very useful for finding clean markers.
cl1_14.markers = find.markers(hpf14, 1, thresh.use = 1, test.use = "roc", min.pct = 0.5)
head(cl1_14.markers[order(cl1_14.markers$myAUC, decreasing = T), ], 20)
```

```
##                              myAUC avg_diff power pct.1 pct.2
## SLIT1/2/3                    0.797 1.550038 0.594 0.886 0.625
## LRP4/8                       0.761 1.619001 0.522 0.692 0.234
## DEF6                         0.749 1.325675 0.498 0.829 0.562
## SFRP2                        0.739 1.183835 0.478 0.872 0.641
## KH2013:KH.S1159.1_FLI1       0.731 1.277835 0.462 0.725 0.406
## KH2013:KH.C13.152_F14B6.6    0.729 1.495670 0.458 0.654 0.266
## COL13A1                      0.727 1.442988 0.454 0.645 0.250
## PDE5A                        0.704 1.457533 0.408 0.621 0.312
## VWC2L                        0.703 1.425926 0.406 0.559 0.156
## Fli/ERG-d                    0.702 1.543703 0.404 0.611 0.281
## LYS2                         0.702 1.173341 0.404 0.720 0.469
## KH2013:KH.C11.362            0.698 1.068338 0.396 0.725 0.391
## KH2013:KH.C4.547_BMP2/4      0.694 1.079053 0.388 0.706 0.422
## C5ORF48                      0.691 1.172332 0.382 0.682 0.406
## EFNB1/2/3                    0.690 1.372033 0.380 0.578 0.250
```

```
## LRRC42                           0.682 1.202552 0.364 0.616 0.328
## KH2013:KH.C2.245_B3GNT2/7/9      0.682 1.169769 0.364 0.611 0.344
## RAP1GAP/2                        0.671 1.000794 0.342 0.635 0.391
## KH2013:KH.C7.787_SELP            0.666 1.267515 0.332 0.517 0.219
## KH2013:KH.C3.52_EFNA1/2/3/4/5 0.649 1.747759 0.298 0.540 0.328
```

```
# Visualize known markers with a violin plot
vlnPlot(hpf14, c("TBX1/10", "HAND/2"))
```

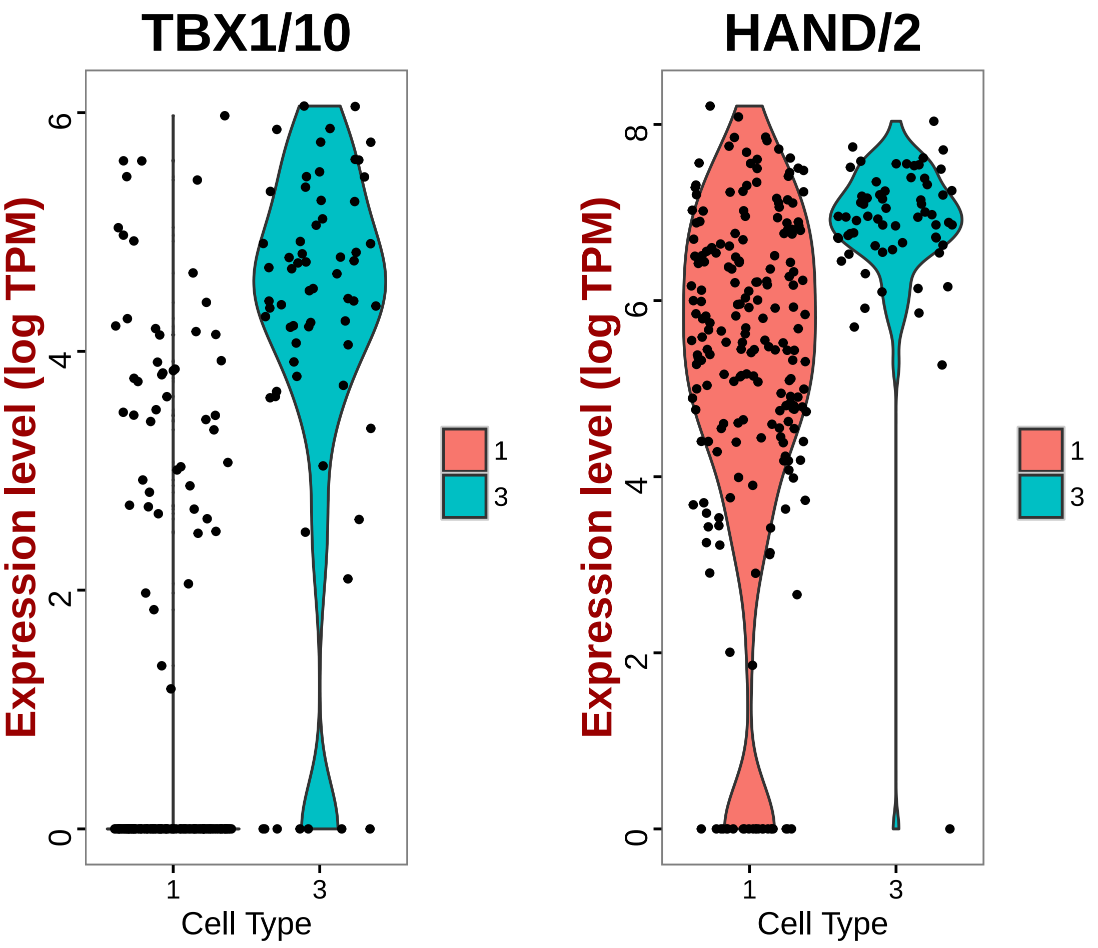


```
# Based on prelimenary studies on TBX1/10- and HAND/2+ expression in TVC
# lineage, cluster 12 is FHP cells

# Visualize new markers with a violin plot
vlnPlot(hpf14, c("LRP4/8", "SLIT1/2/3"))
```

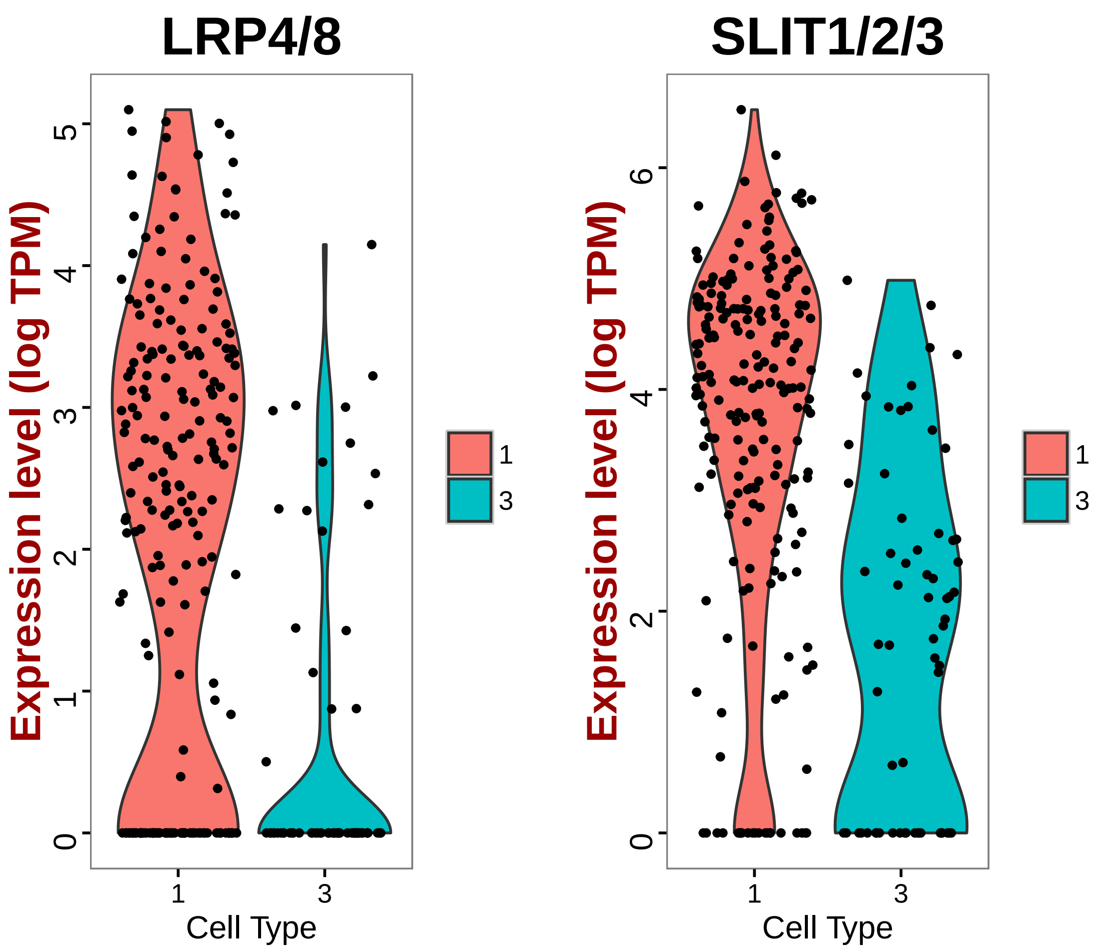


```
# Find markers for cluster 4
cl3_14.markers = find.markers(hpf14, 3, thresh.use = 1, test.use = "roc", min.pct = 0.5)
head(cl3_14.markers[order(cl3_14.markers$myAUC, decreasing = T), ], 20)
```

```
##                              myAUC avg_diff power pct.1 pct.2
## KH2013:KH.S555.1_HTR7        0.895 1.787203 0.790 0.984 0.555
## TBX1/10                      0.884 1.995667 0.768 0.891 0.237
## KH2013:KH.C4.404             0.866 2.276980 0.732 0.859 0.223
## TMSB15A                      0.860 2.237798 0.720 0.891 0.355
## ELK1/3/4                     0.819 2.005421 0.638 0.766 0.194
## HRH1                         0.771 2.110109 0.542 0.594 0.057
## ZEB1/2                       0.769 1.113927 0.538 0.938 0.872
## KH2013:KH.C3.696             0.758 1.277699 0.516 0.781 0.341
## KH2013:KH.C4.125_BMP2/4      0.750 1.278512 0.500 0.781 0.436
## ZK637.14                     0.742 1.315203 0.484 0.781 0.351
## KH2013:KH.C10.203            0.735 1.301948 0.470 0.688 0.256
## IRX4/6                       0.725 1.142938 0.450 0.734 0.384
## ITPKA                        0.722 1.384907 0.444 0.750 0.464
## FOXF1/2                      0.714 1.146914 0.428 0.734 0.422
```

```
## KH2013:KH.S1012.2          0.710 1.454288 0.420 0.578 0.190
## KH2013:KH.L108.33          0.709 1.591887 0.418 0.531 0.118
## W02G9.4                    0.687 1.177161 0.374 0.594 0.246
## SSTR1/2/3/4/5              0.686 1.393247 0.372 0.516 0.161
## KH2013:KH.C1.953_CG32702   0.670 1.115838 0.340 0.609 0.322
## KH2013:KH.C9.692_GABRR1/2/3 0.669 1.146391 0.338 0.531 0.209
```

```
# Visualize known markers with a violin plot
vlnPlot(hpf14, c("HAND/2", "TBX1/10"))
```

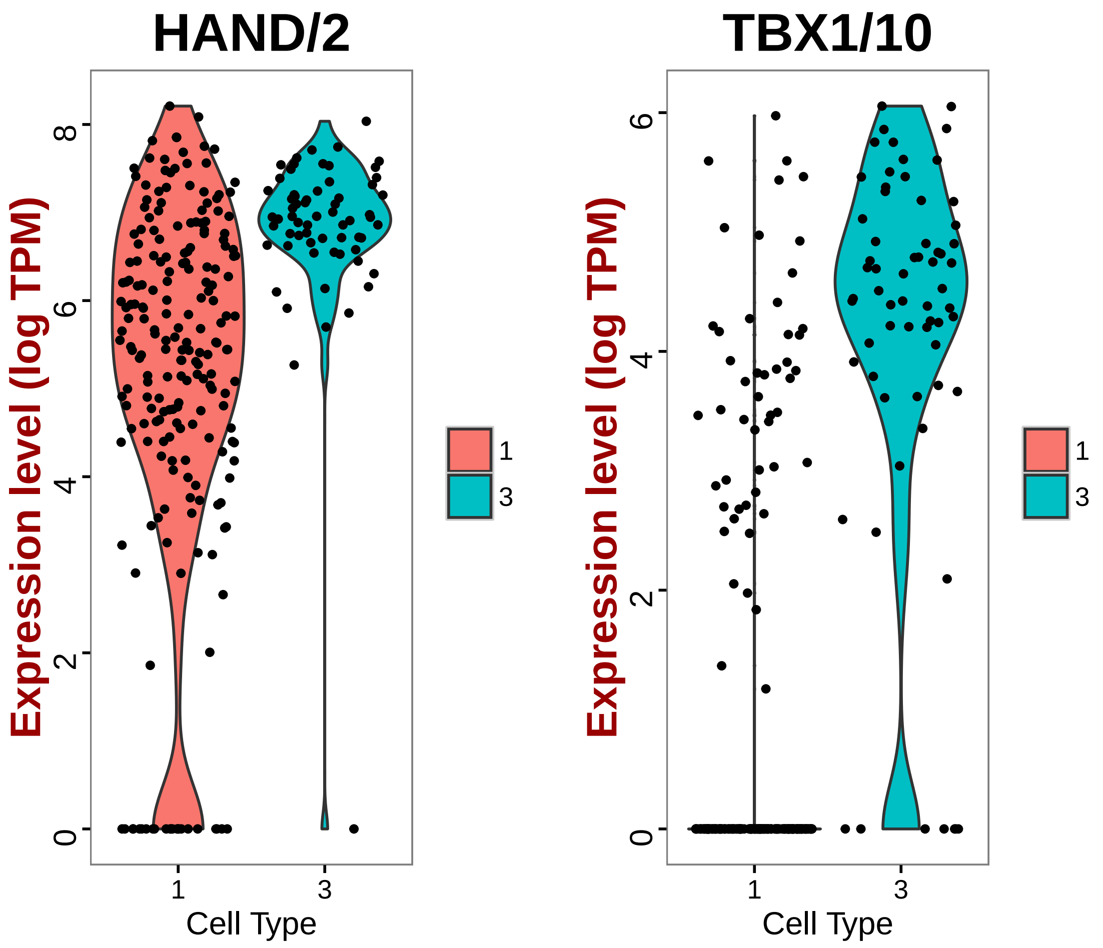


```
# Based on preliminary studies these TBX1/10- HAND/2+ cells are STVCs

# Visualize new markers with a violin plot
vlnPlot(hpf14, c("KH2013:KH.S555.1_HTR7", "TMSB15A"))
```

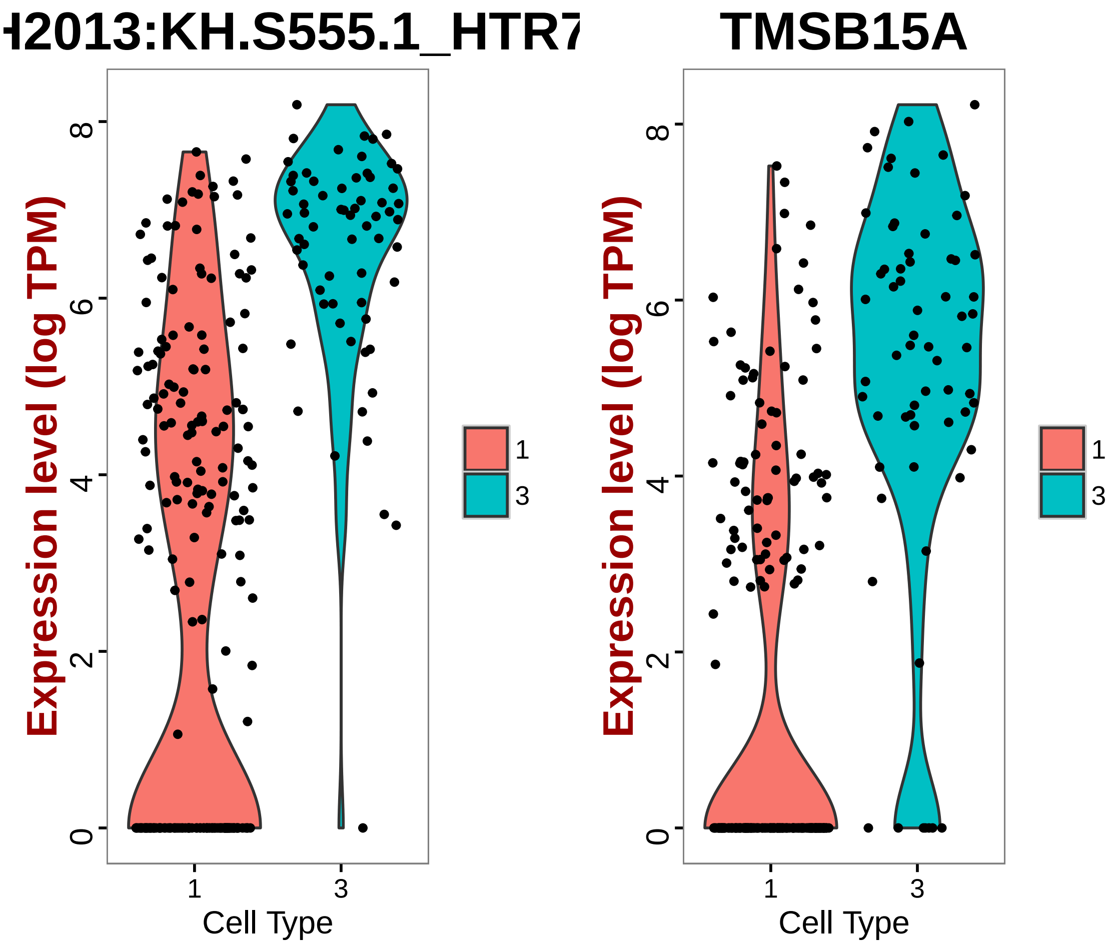


```
# Write cell names into text files
write.table(which.cells(hpf14, 1), file = "14FHPCells.txt", sep = "\t")
write.table(which.cells(hpf14, 3), file = "14STVCCells.txt", sep = "\t")

# Rename cluster identities
hpf14 = rename.ident(hpf14, 1, "14FHP")
hpf14 = rename.ident(hpf14, 3, "14STVC")

# Visualize tSNE used color scheme FHP-red, STVC-yellow
tsne.plot(hpf14, do.label = T, label.pt.size = 1, label.cex.text = 1.2, label.cols.use =
    "yellow"))
```

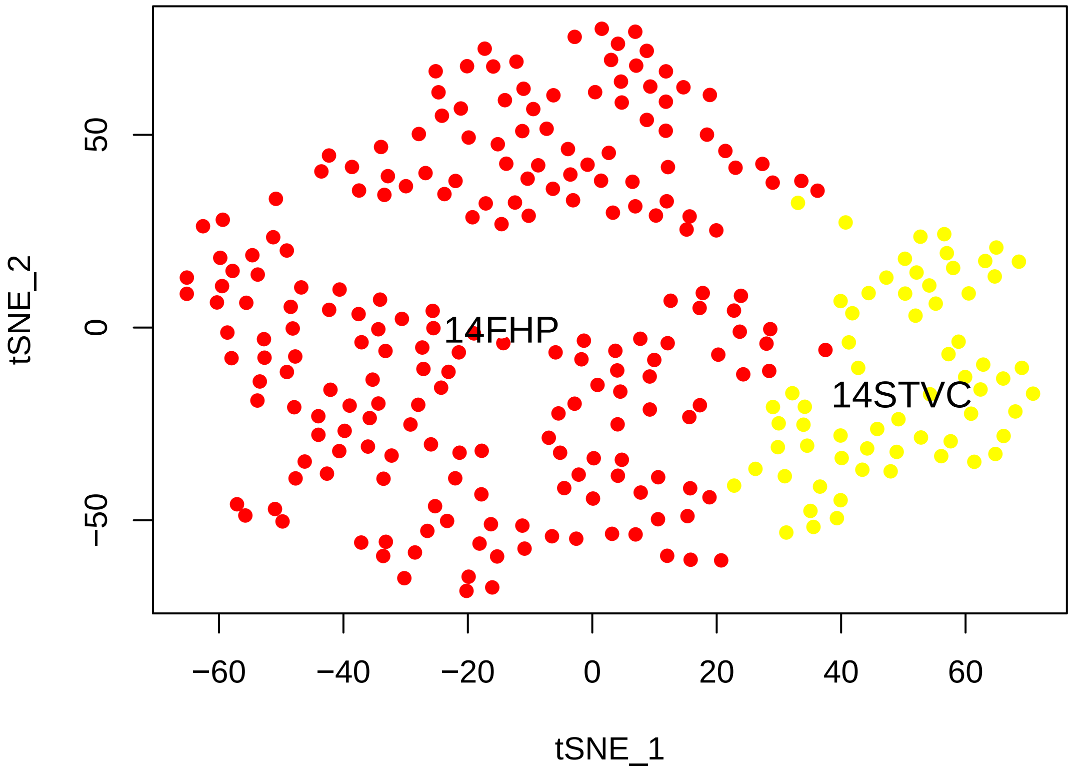


```
# Store FHP markers in text file
FHP_14.markers = cl1_14.markers
head(FHP_14.markers[order(FHP_14.markers$myAUC, decreasing = T), ], 20)
```

```
##                                myAUC avg_diff power pct.1 pct.2
## SLIT1/2/3                      0.797 1.550038 0.594 0.886 0.625
## LRP4/8                         0.761 1.619001 0.522 0.692 0.234
## DEF6                           0.749 1.325675 0.498 0.829 0.562
## SFRP2                          0.739 1.183835 0.478 0.872 0.641
## KH2013:KH.S1159.1_FLI1         0.731 1.277835 0.462 0.725 0.406
## KH2013:KH.C13.152_F14B6.6      0.729 1.495670 0.458 0.654 0.266
## COL13A1                        0.727 1.442988 0.454 0.645 0.250
## PDE5A                          0.704 1.457533 0.408 0.621 0.312
## VWC2L                          0.703 1.425926 0.406 0.559 0.156
## Fli/ERG-d                      0.702 1.543703 0.404 0.611 0.281
## LYS2                           0.702 1.173341 0.404 0.720 0.469
## KH2013:KH.C11.362              0.698 1.068338 0.396 0.725 0.391
## KH2013:KH.C4.547_BMP2/4        0.694 1.079053 0.388 0.706 0.422
## C5ORF48                        0.691 1.172332 0.382 0.682 0.406
## EFNB1/2/3                      0.690 1.372033 0.380 0.578 0.250
## LRRC42                         0.682 1.202552 0.364 0.616 0.328
## KH2013:KH.C2.245_B3GNT2/7/9    0.682 1.169769 0.364 0.611 0.344
## RAP1GAP/2                      0.671 1.000794 0.342 0.635 0.391
```

```
## KH2013:KH.C7.787_SELP         0.666 1.267515 0.332 0.517 0.219
## KH2013:KH.C3.52_EFNA1/2/3/4/5 0.649 1.747759 0.298 0.540 0.328
```

```
write.table(FHP_14.markers, file = "FHP_14.markers.txt", sep = "\t")

# Store STVC markers in text file
STVC_14.markers = cl3_14.markers
head(STVC_14.markers[order(STVC_14.markers$myAUC, decreasing = T), ], 20)
```

```
##                               myAUC avg_diff power pct.1 pct.2
## KH2013:KH.S555.1_HTR7        0.895 1.787203 0.790 0.984 0.555
## TBX1/10                      0.884 1.995667 0.768 0.891 0.237
## KH2013:KH.C4.404             0.866 2.276980 0.732 0.859 0.223
## TMSB15A                      0.860 2.237798 0.720 0.891 0.355
## ELK1/3/4                     0.819 2.005421 0.638 0.766 0.194
## HRH1                         0.771 2.110109 0.542 0.594 0.057
## ZEB1/2                       0.769 1.113927 0.538 0.938 0.872
## KH2013:KH.C3.696             0.758 1.277699 0.516 0.781 0.341
## KH2013:KH.C4.125_BMP2/4      0.750 1.278512 0.500 0.781 0.436
## ZK637.14                     0.742 1.315203 0.484 0.781 0.351
## KH2013:KH.C10.203            0.735 1.301948 0.470 0.688 0.256
## IRX4/6                       0.725 1.142938 0.450 0.734 0.384
## ITPKA                        0.722 1.384907 0.444 0.750 0.464
## FOXF1/2                      0.714 1.146914 0.428 0.734 0.422
## KH2013:KH.S1012.2            0.710 1.454288 0.420 0.578 0.190
## KH2013:KH.L108.33            0.709 1.591887 0.418 0.531 0.118
## W02G9.4                      0.687 1.177161 0.374 0.594 0.246
## SSTR1/2/3/4/5                0.686 1.393247 0.372 0.516 0.161
## KH2013:KH.C1.953_CG32702     0.670 1.115838 0.340 0.609 0.322
## KH2013:KH.C9.692_GABRR1/2/3 0.669 1.146391 0.338 0.531 0.209
```

```
write.table(STVC_14.markers, file = "STVC_14.markers.txt", sep = "\t")

# Visualize markers of different clusters using violin plot and feature plot
genes.viz.14 = c("LRP4/8", "SLIT1/2/3", "TBX1/10", "KH2013:KH.S555.1_HTR7")
feature.plot(hpf14, genes.viz.14, pt.size = 1)
```

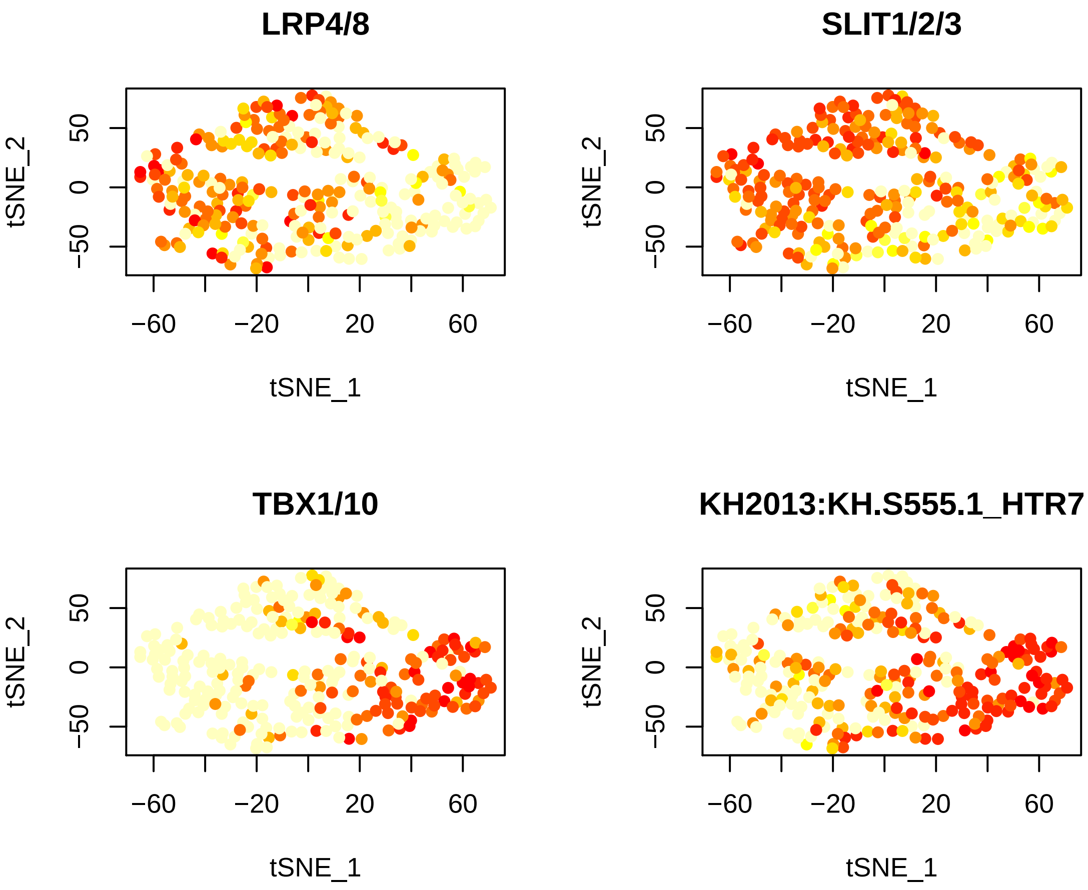


```
vlnPlot(hpf14, genes.viz.14, col = c("red", "yellow"))
```

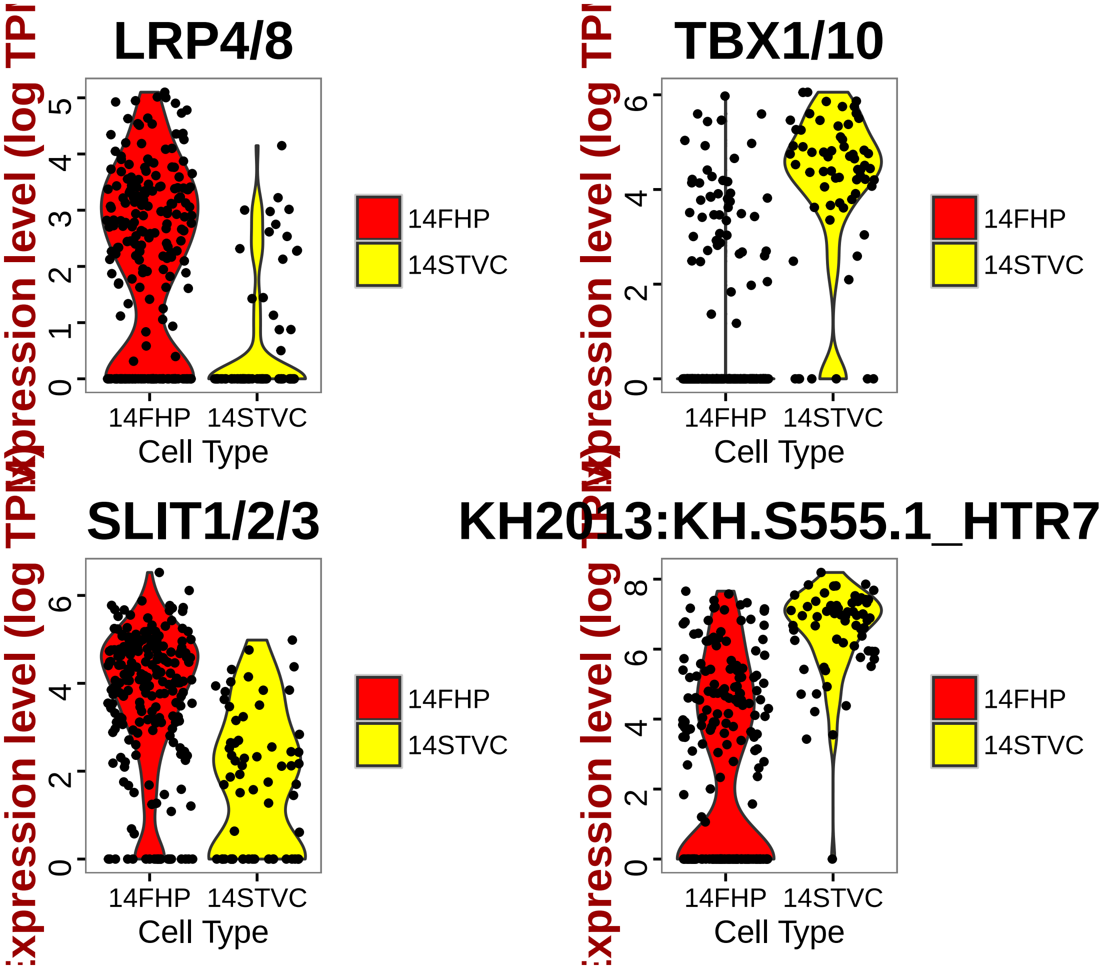


```
# Select markers for plotting on a Heatmap (top 20 positive markers with
# high classification power)
marker14FHP = rownames(FHP_14.markers[order(FHP_14.markers$myAUC, decreasing = T)[1:20],
    ])
marker14STVC = rownames(STVC_14.markers[order(STVC_14.markers$myAUC, decreasing = T)[1:2
    ])
marker.14 = c(marker14FHP, marker14STVC)

# Draw a heatmap of all cells for these marker genes
doHeatMap(hpf14, genes.use = marker.14, remove.key = TRUE, slim.col.label = T,
    cex.col = 1.2, col.use = col, draw.line = F)
```

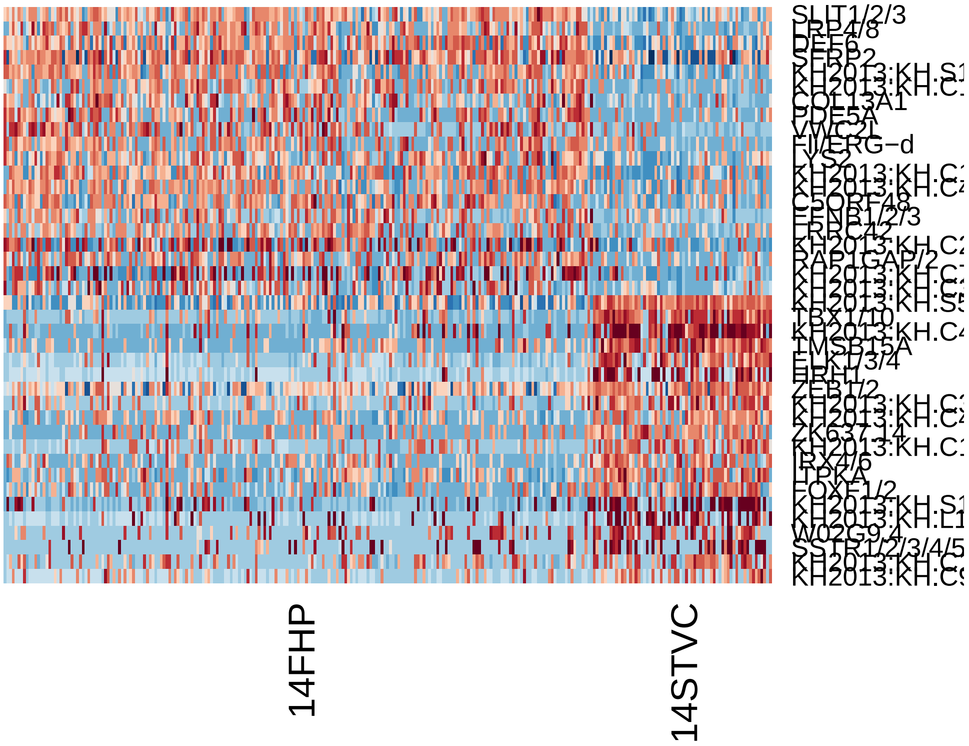


4. hpf20

```
# Subset data from preprocessed Seurat object
hpf20 = subsetData(hpfall.remv2, which.cells(hpfall.remv2, "hpf20"), do.center = F,
    do.scale = F)
hpf20
```

```
## An object of class seurat in project allhpf
##  14864 genes across 288 samples.
```

```
# Find variable gene with 4 < Average expression and Dispersion > 2
hpf20 = mean.var.plot(hpf20)
```

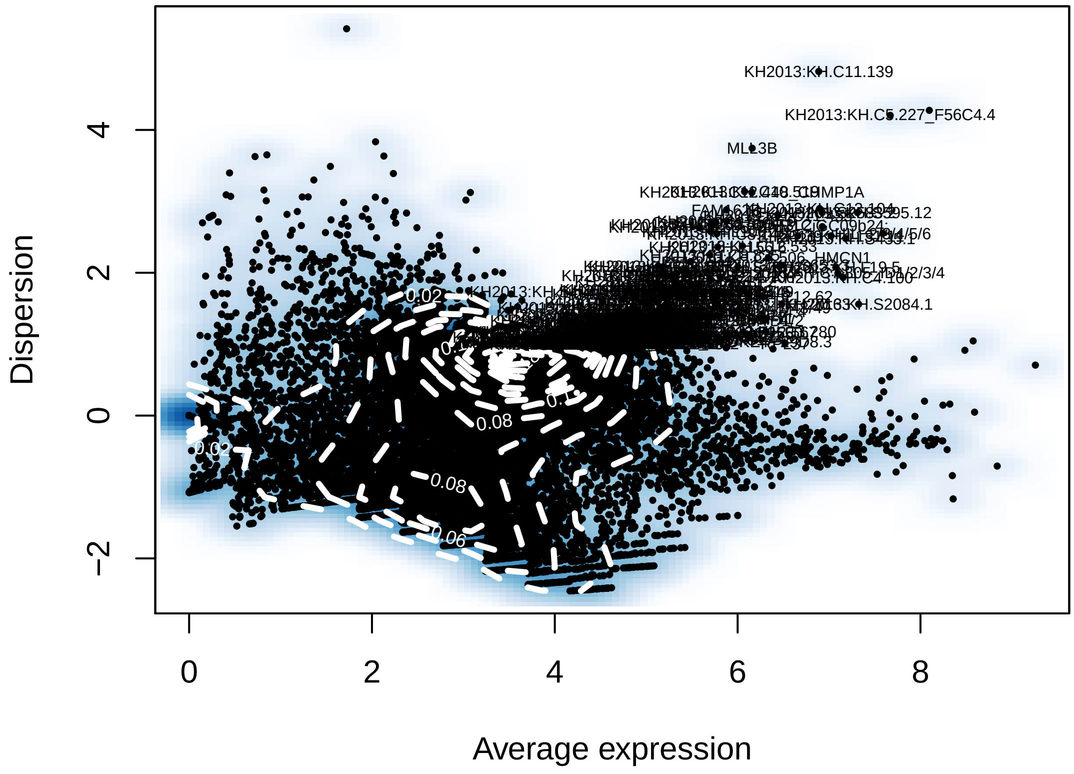


```r
length(hpf20@var.genes)
```

```
## [1] 293
```

```r
# Run a PCA using variable gene list
hpf20 = pca(hpf20, do.print = F)
pcScree(hpf20, hpf20@var.genes, 10)
```

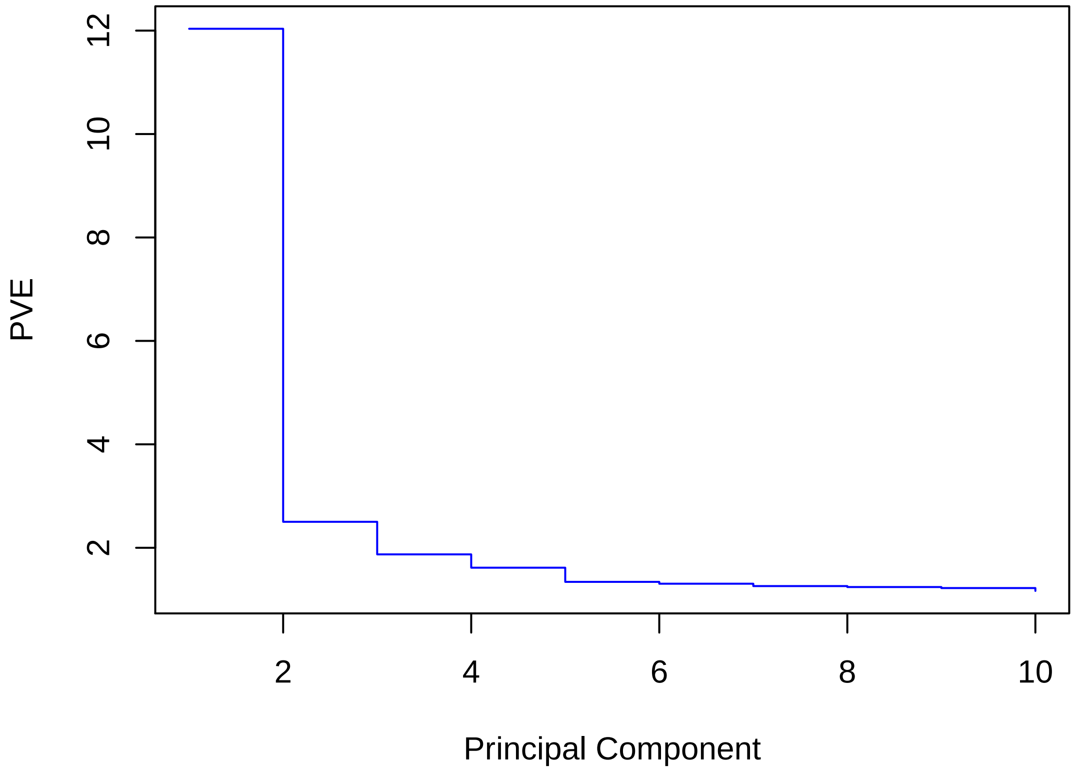


```r
pcHeatmap(hpf20, 1, do.balanced = T, col.use = col)
```

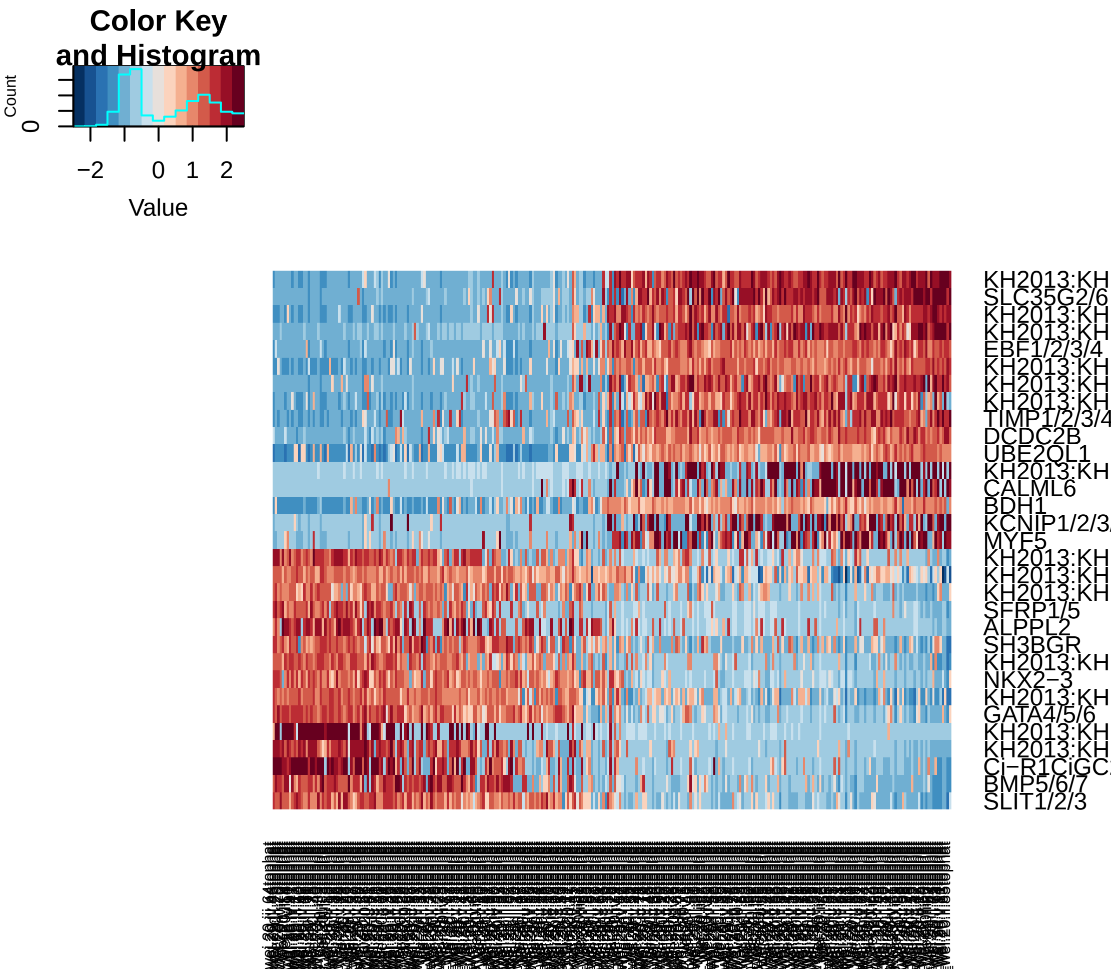


```r
pcHeatmap(hpf20, 2, do.balanced = T, col.use = col)
```

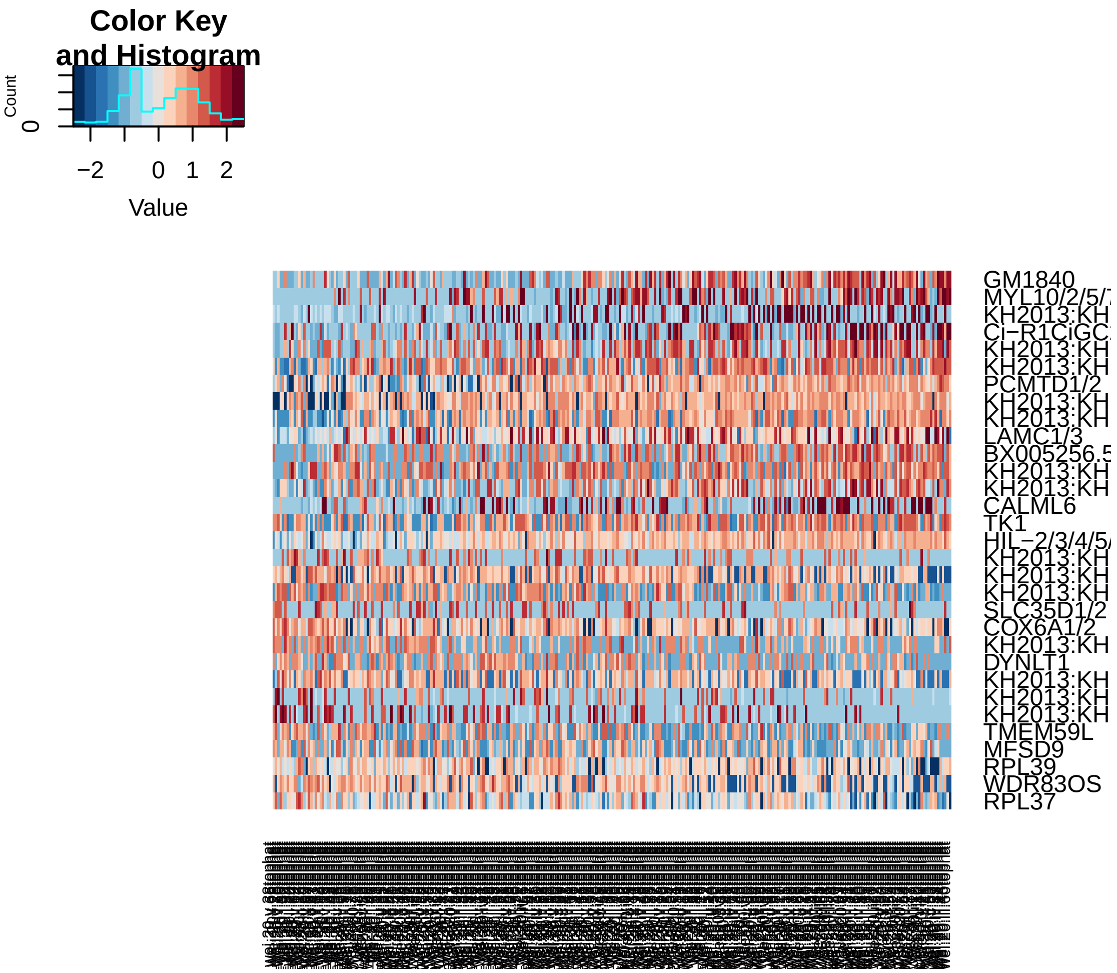


```r
pcHeatmap(hpf20, 3, do.balanced = T, col.use = col)
```

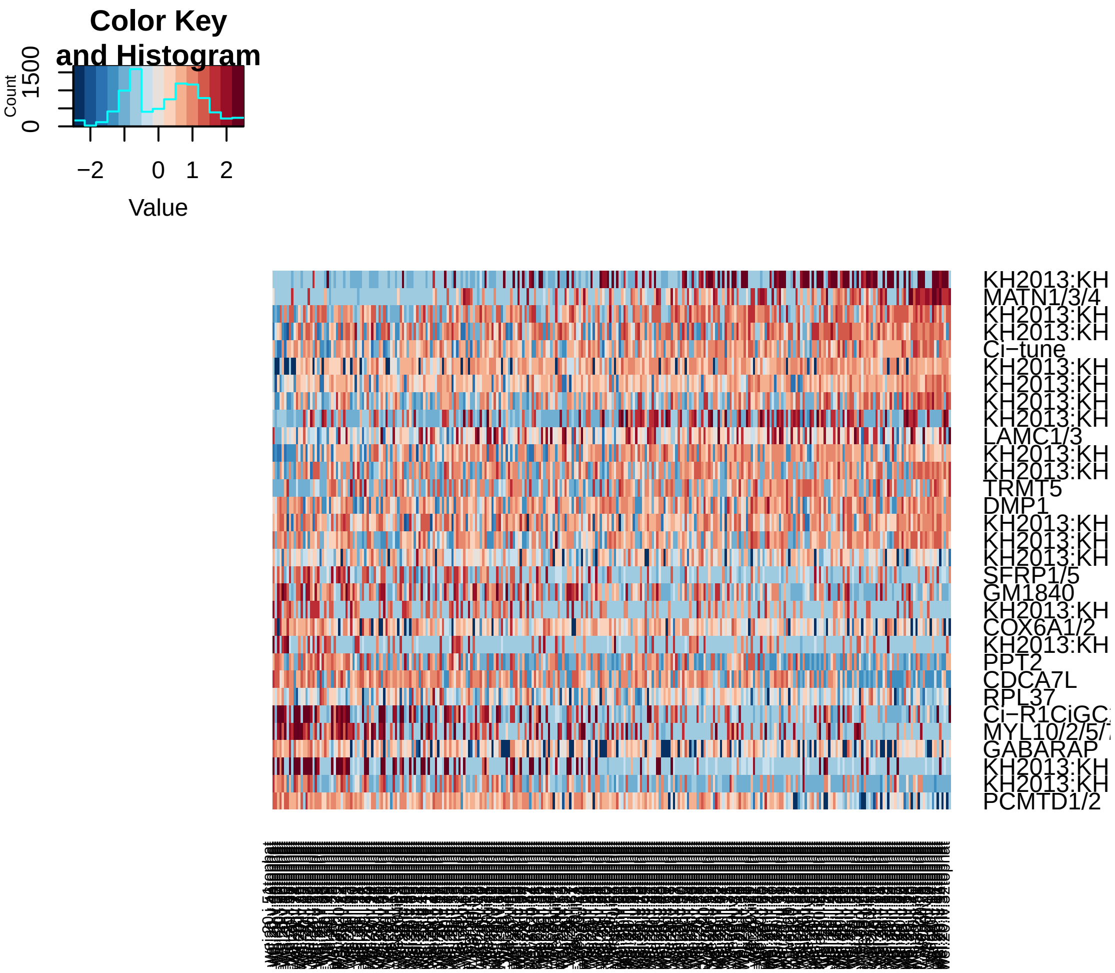


```
pca.plot(hpf20, 1, 2)
```

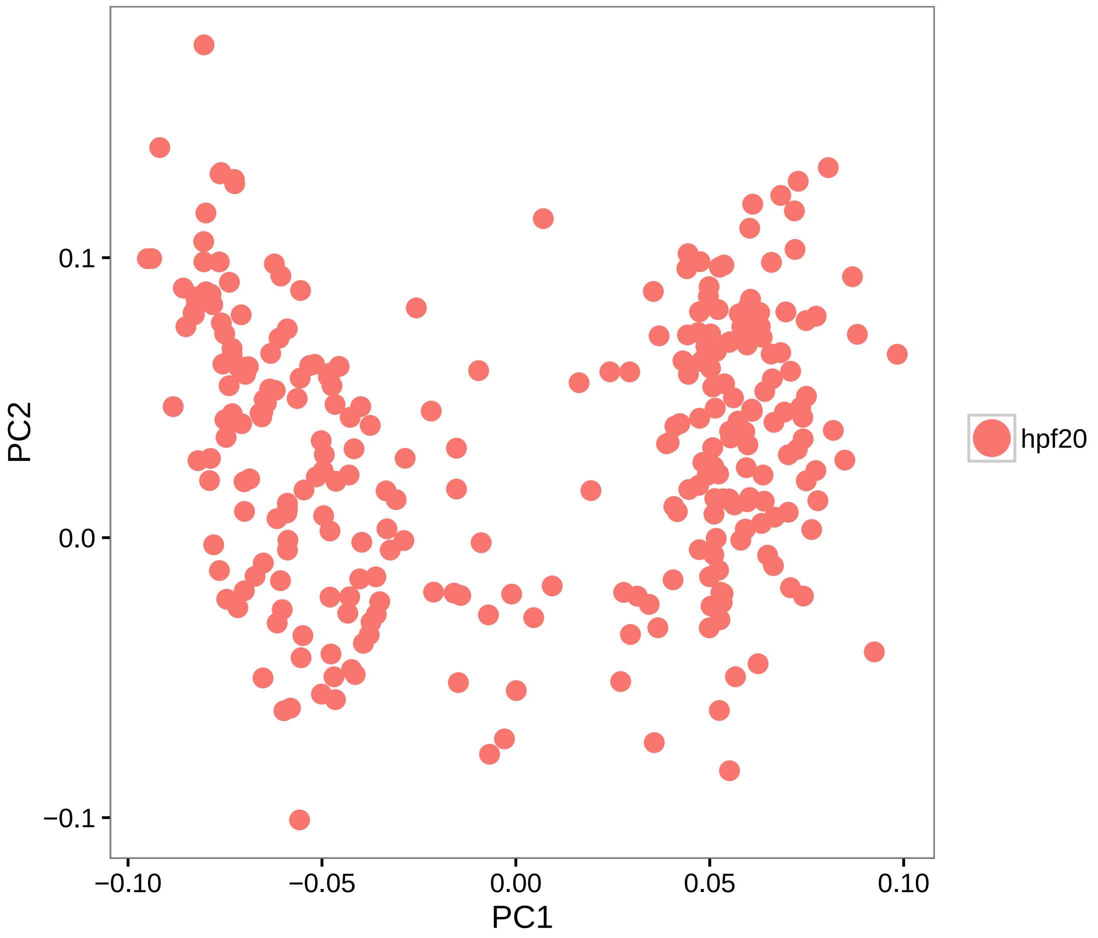


```r
pca.plot(hpf20, 1, 3)
```

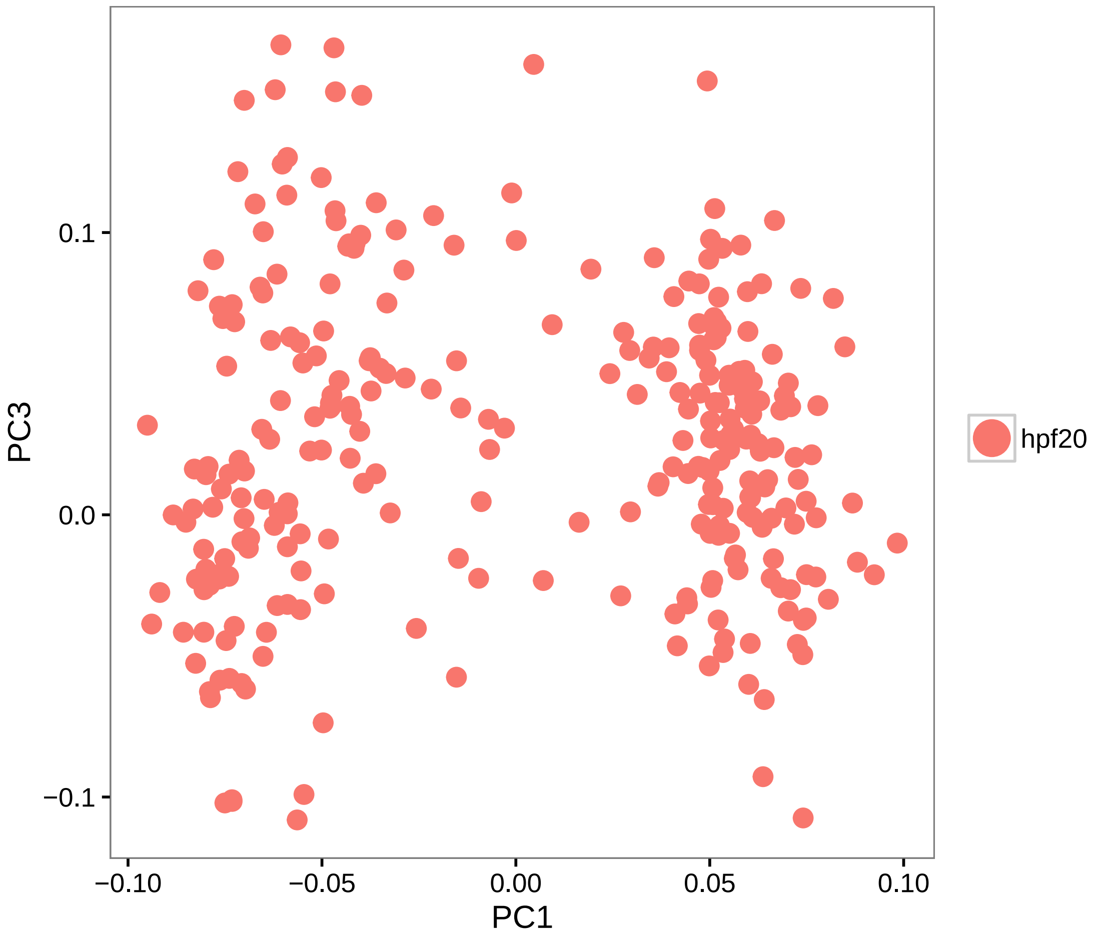


```
# Calculate PCA scores for all genes (PCA projection)
hpf20 = project.pca(hpf20, do.print = F)

# Visualize the full projected PCA, which now includes new genes which were
# not previously (use.full=TRUE)
pcHeatmap(hpf20, 1, use.full = T, do.balanced = T, col.use = col)
```

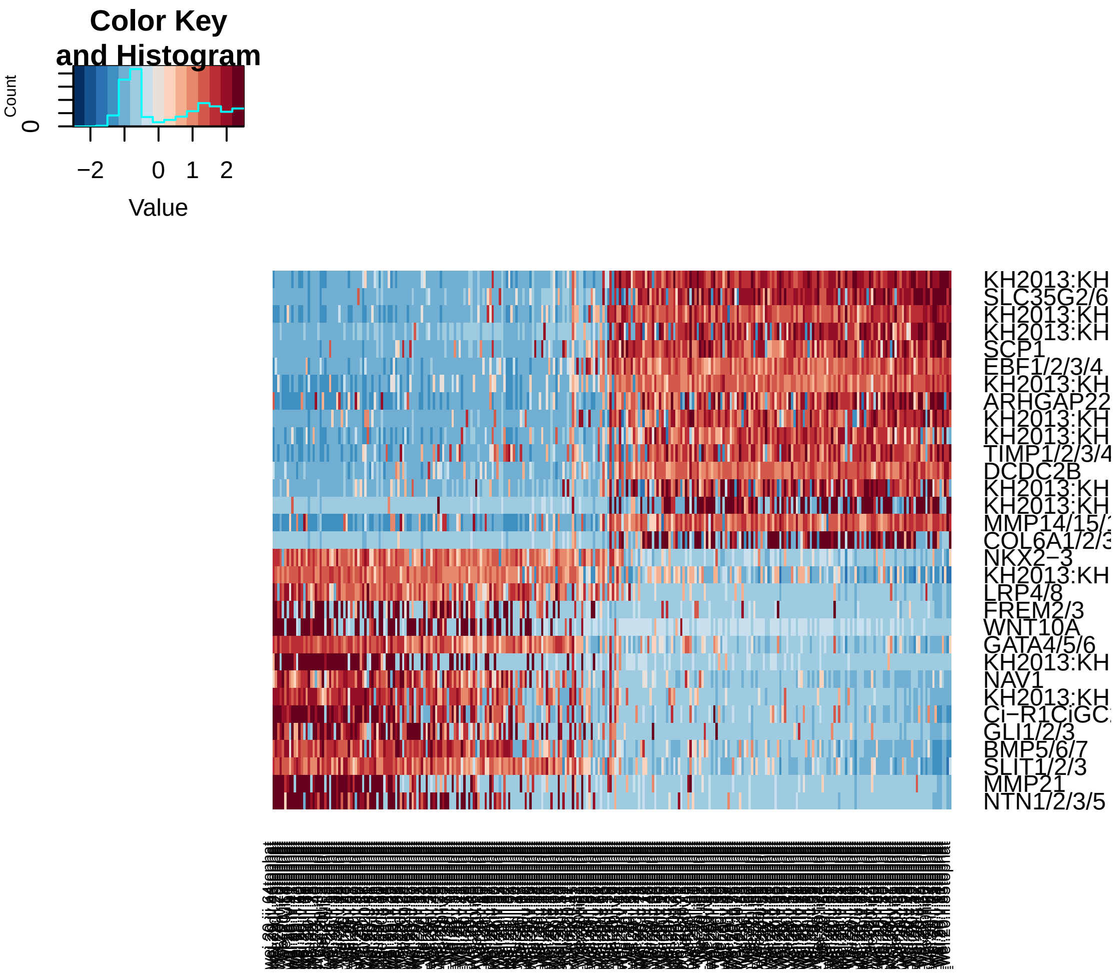


```
pcHeatmap(hpf20, 2, use.full = T, do.balanced = T, col.use = col)  #technical
```

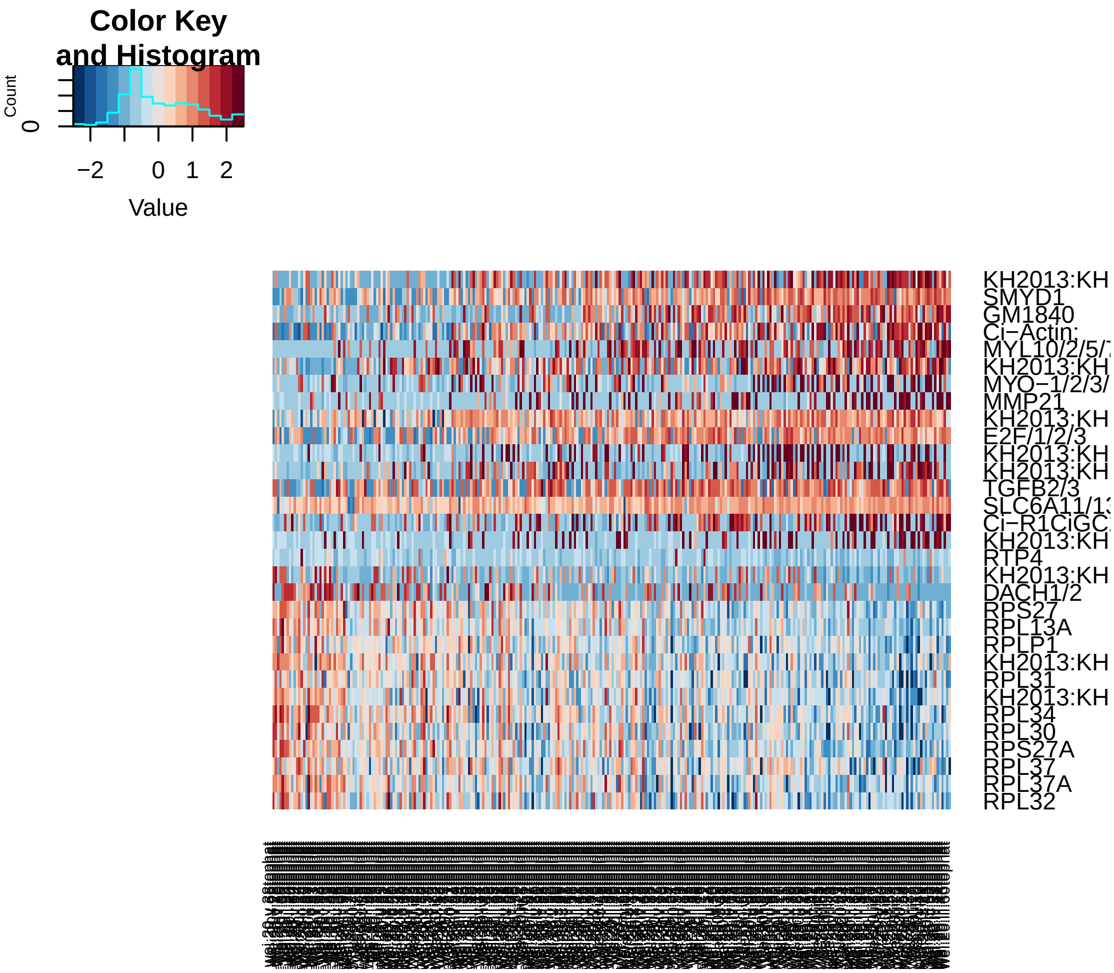


```
pcHeatmap(hpf20, 3, use.full = T, do.balanced = T, col.use = col)
```

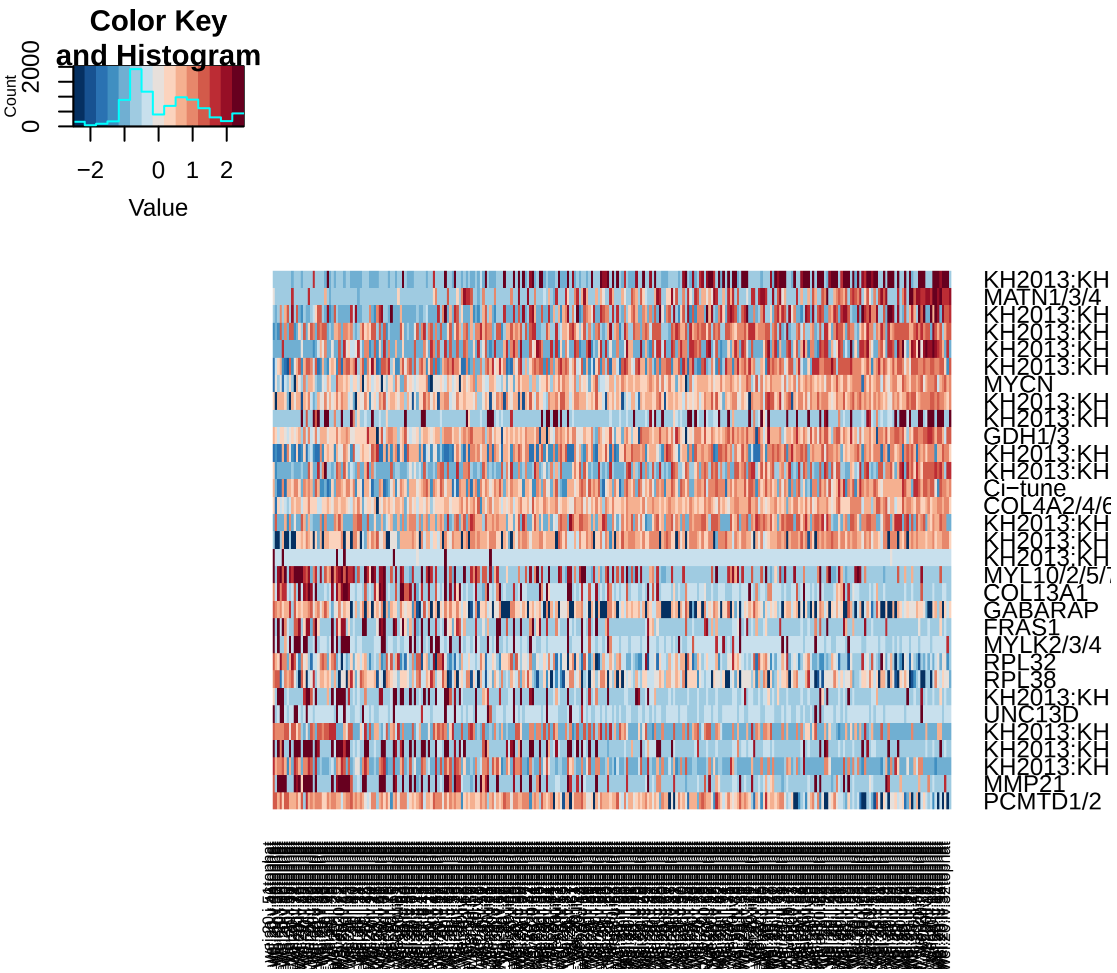


```r
# Do 200 random samplings to find significant genes, each time randomly
# permute 1% of genes This returns a 'p-value' for each gene in each PC,
# based on how likely the gene/PC score woud have been observed by chance
hpf20 = jackStraw(hpf20, num.replicate = 200, do.print = F)

# The jackStraw plot compares the distribution of P-values for each PC with
# a uniform distribution (dashed line) 'Significant' PCs will have a strong
# enrichment of genes with low p-values (solid curve above dashed line)
jackStrawPlot.new(hpf20, PCs = 1:12)
```

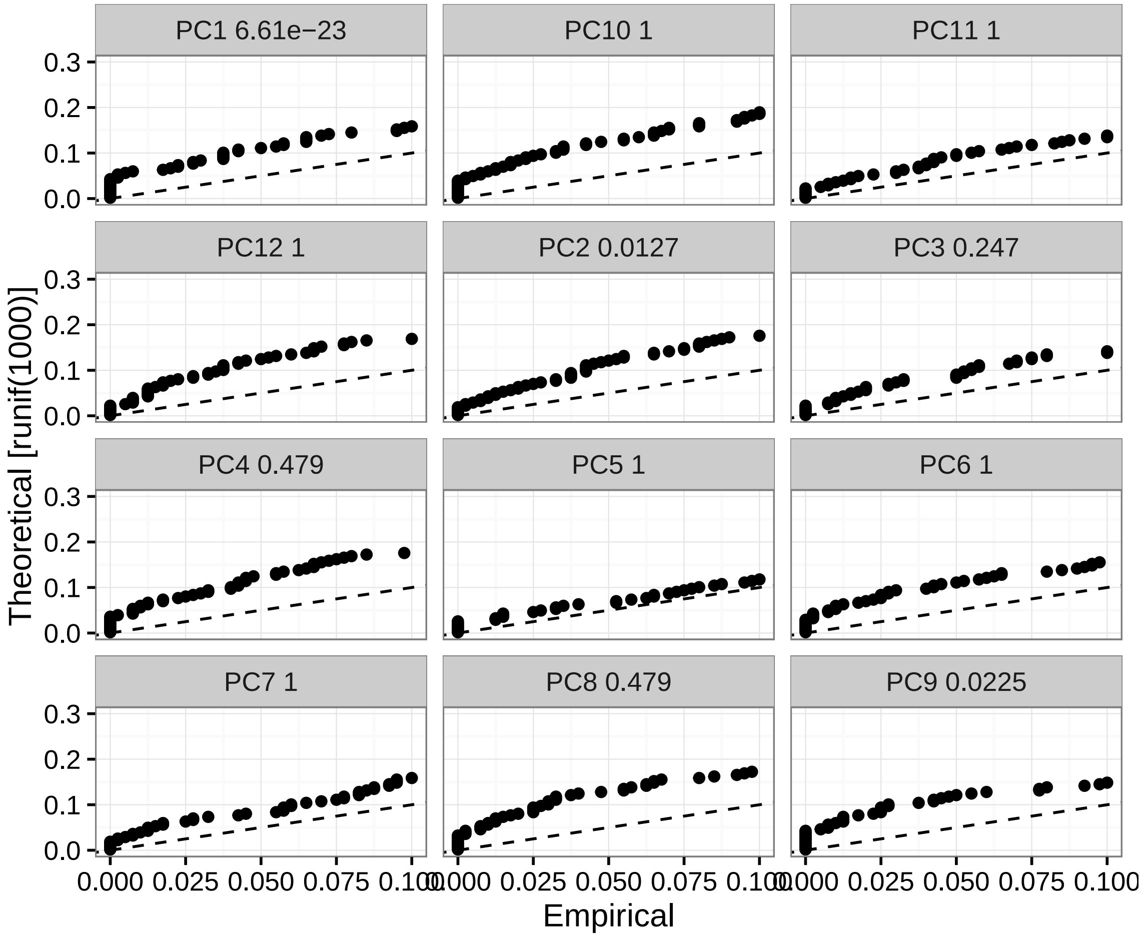


```r
# In this case only PC1 is strongly significant, PC2 is significant but
# contain technical genes

# Select 300 genes from PC1 and rerun PCA
good.genes20 = pcTopGenes(hpf20, 1, 300, T, T)

# Run a PCA using selected gene list
hpf20 = pca(hpf20, pc.genes = good.genes20, do.print = F)
pcScree(hpf20, good.genes20, 10)
```

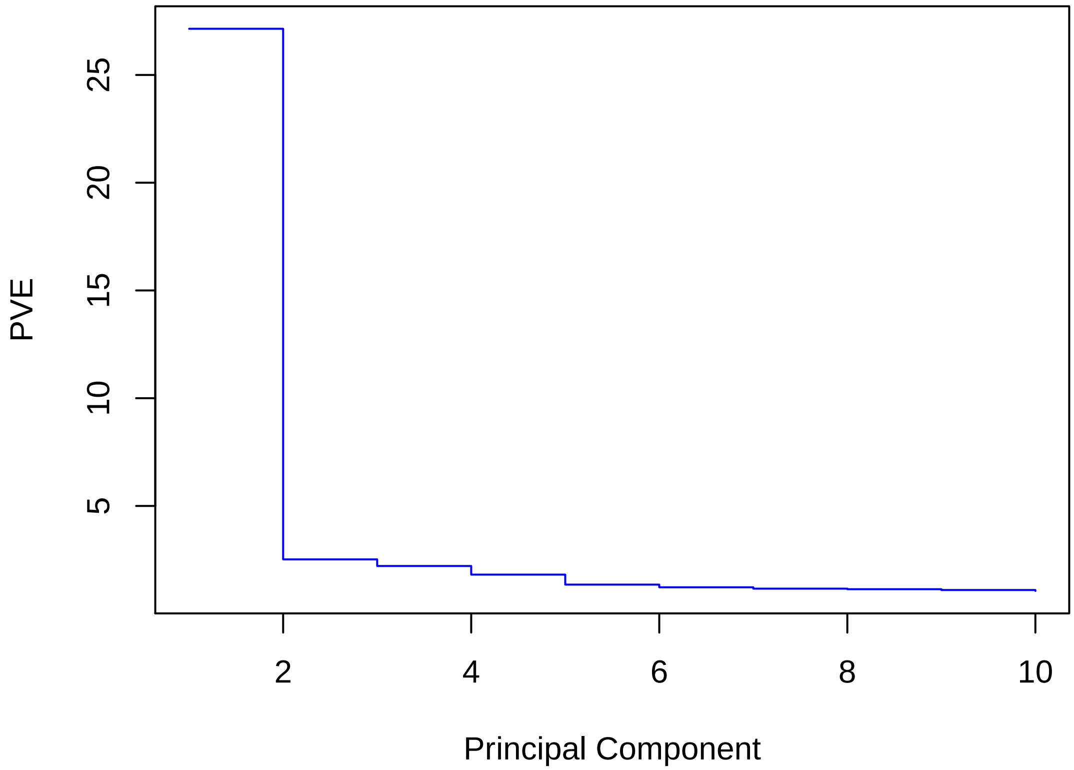


```
pcHeatmap(hpf20, 1, do.balanced = T, col.use = col)
```

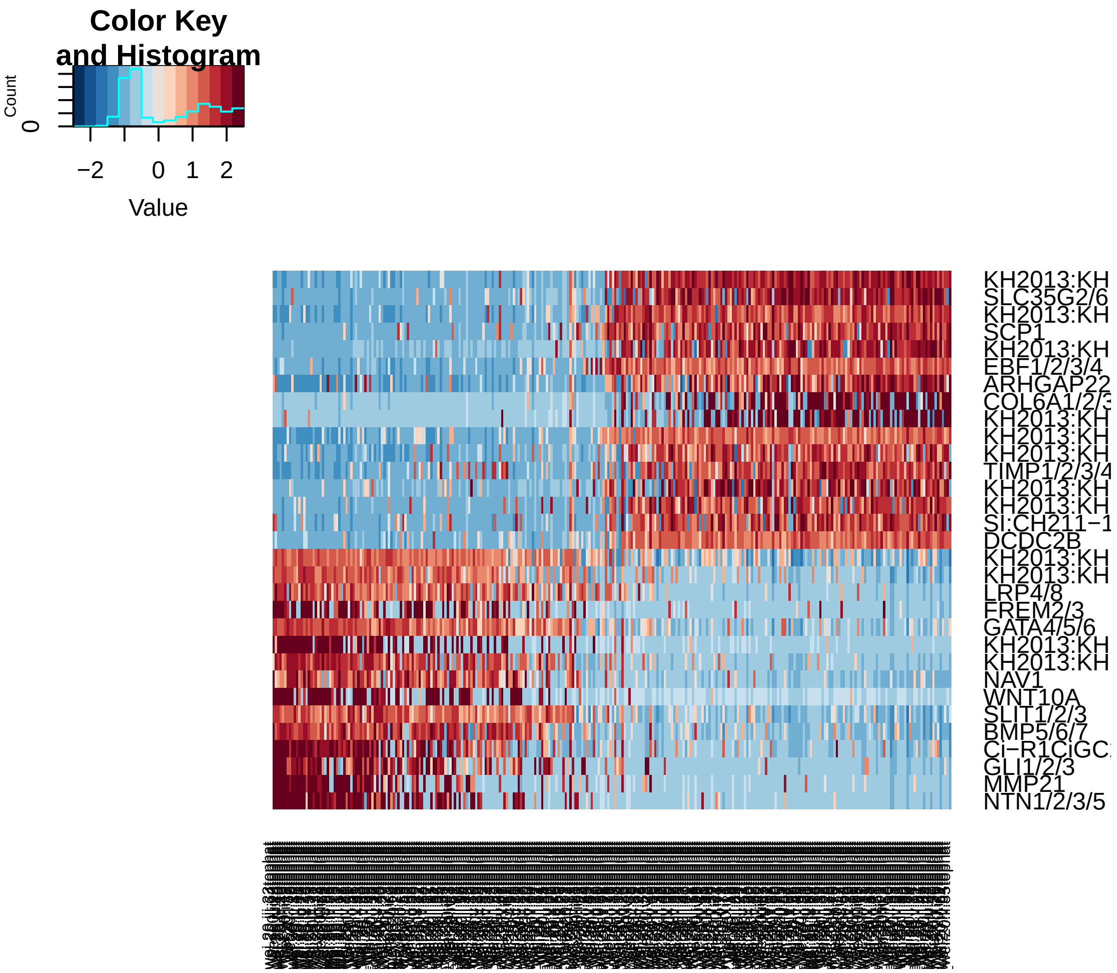


```r
pcHeatmap(hpf20, 2, do.balanced = T, col.use = col)
```

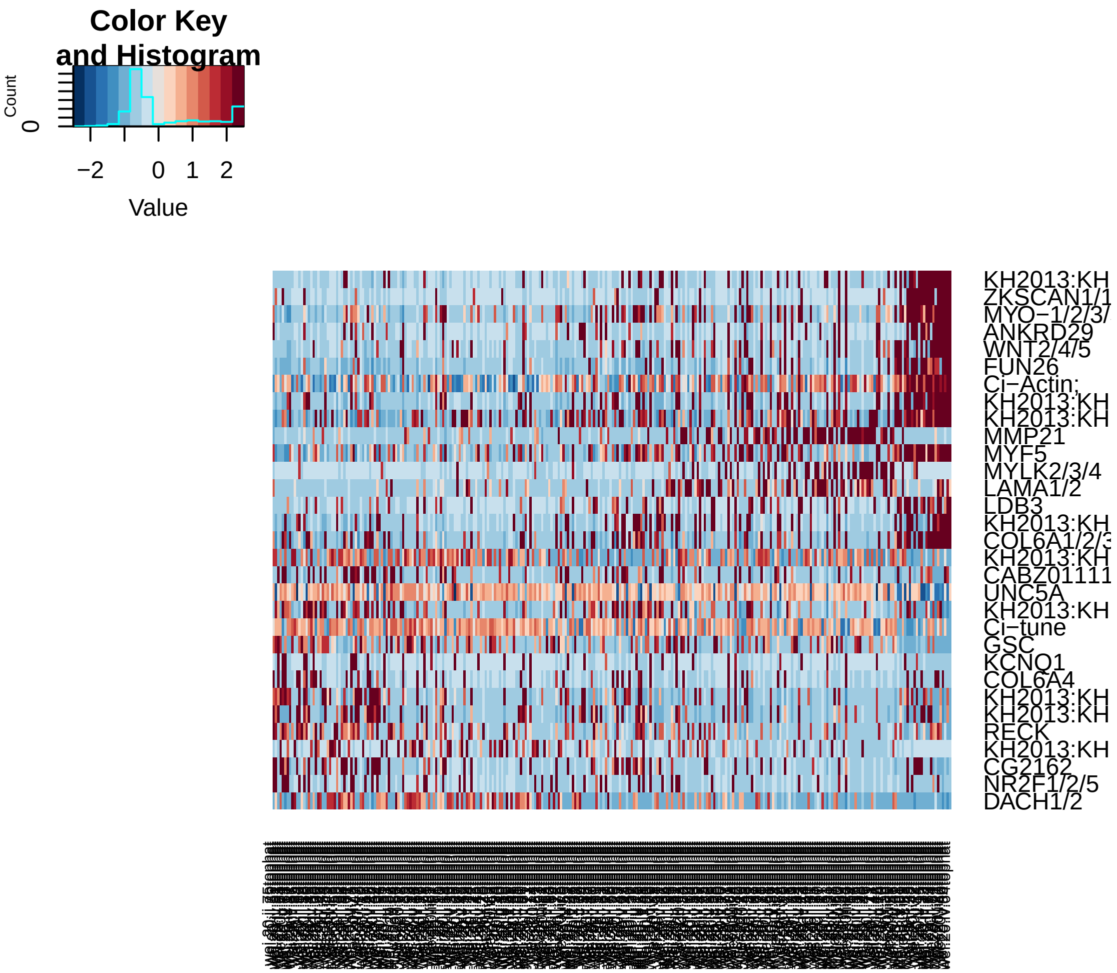


```r
pcHeatmap(hpf20, 3, do.balanced = T, col.use = col)
```

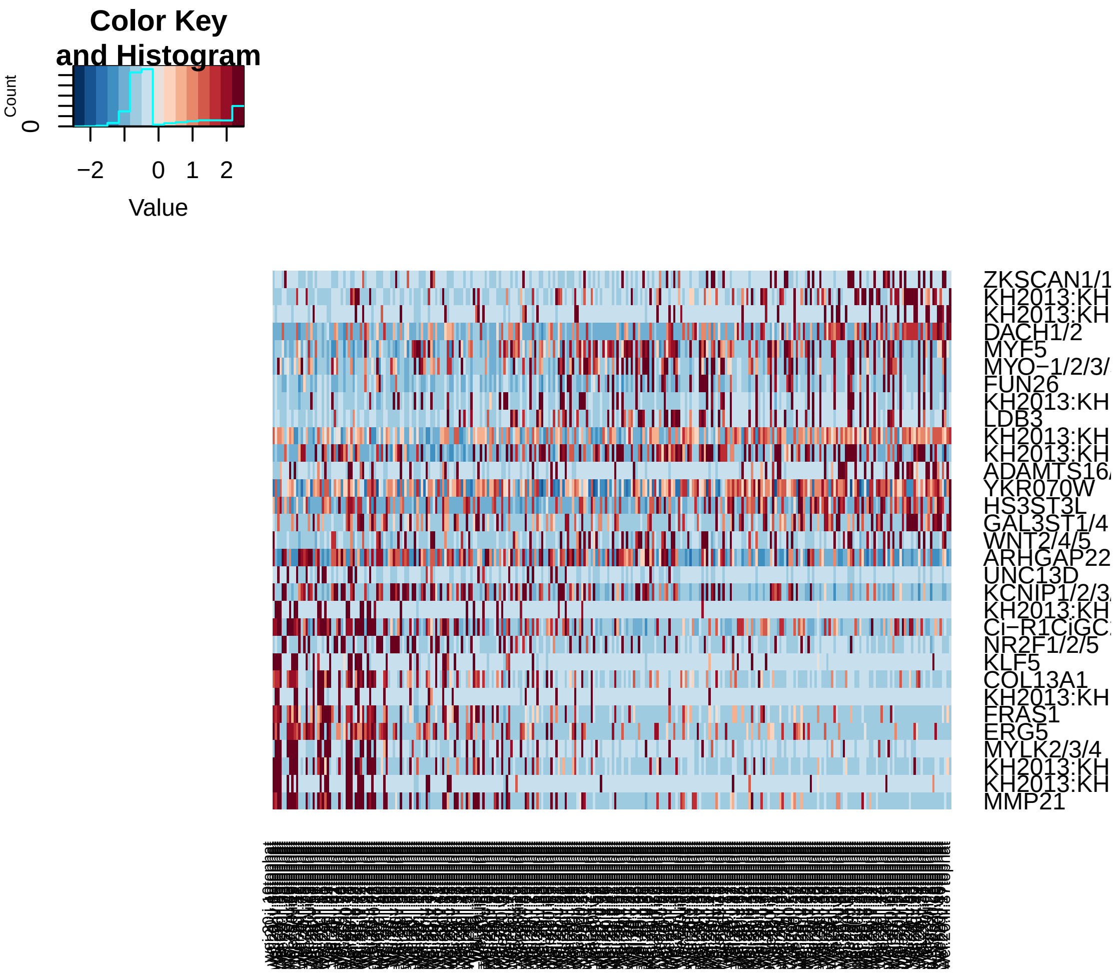


```
pca.plot(hpf20, 1, 2)
```

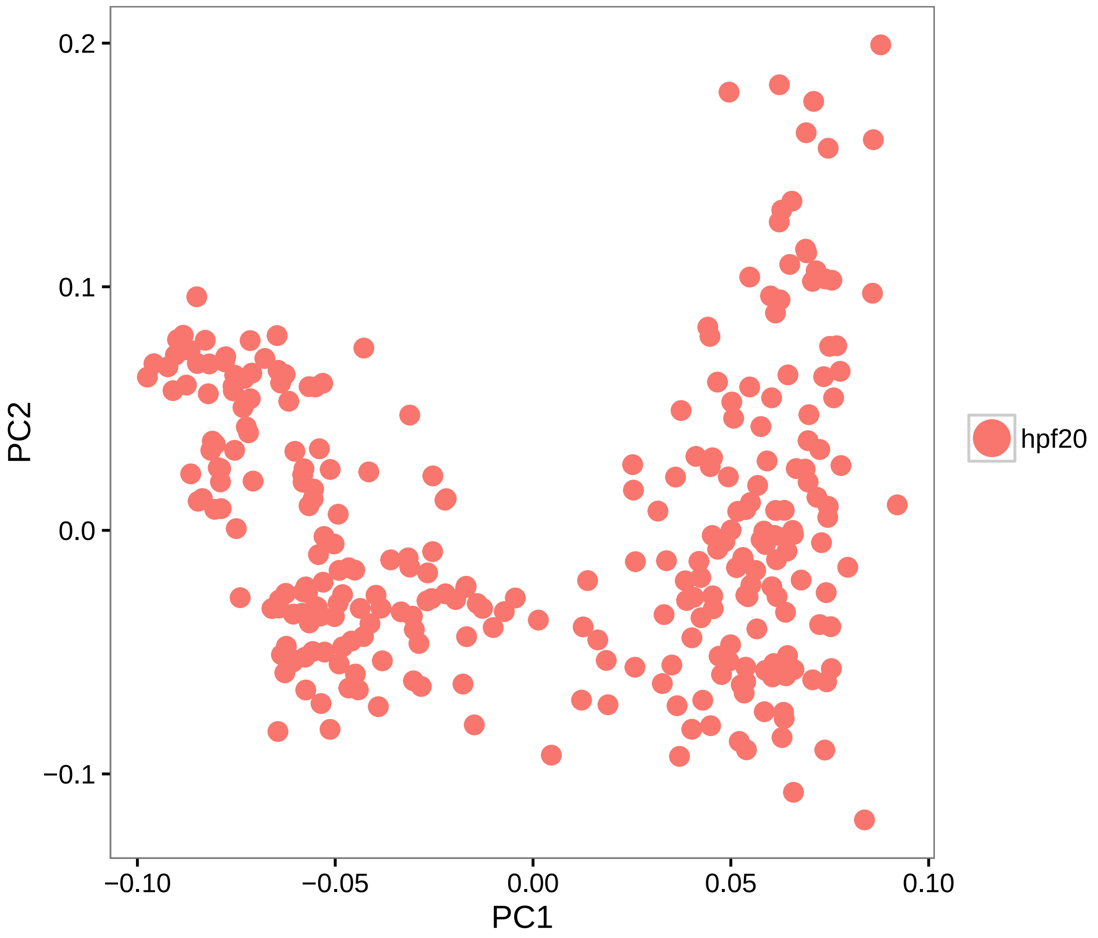


```
pca.plot(hpf20, 1, 3)
```

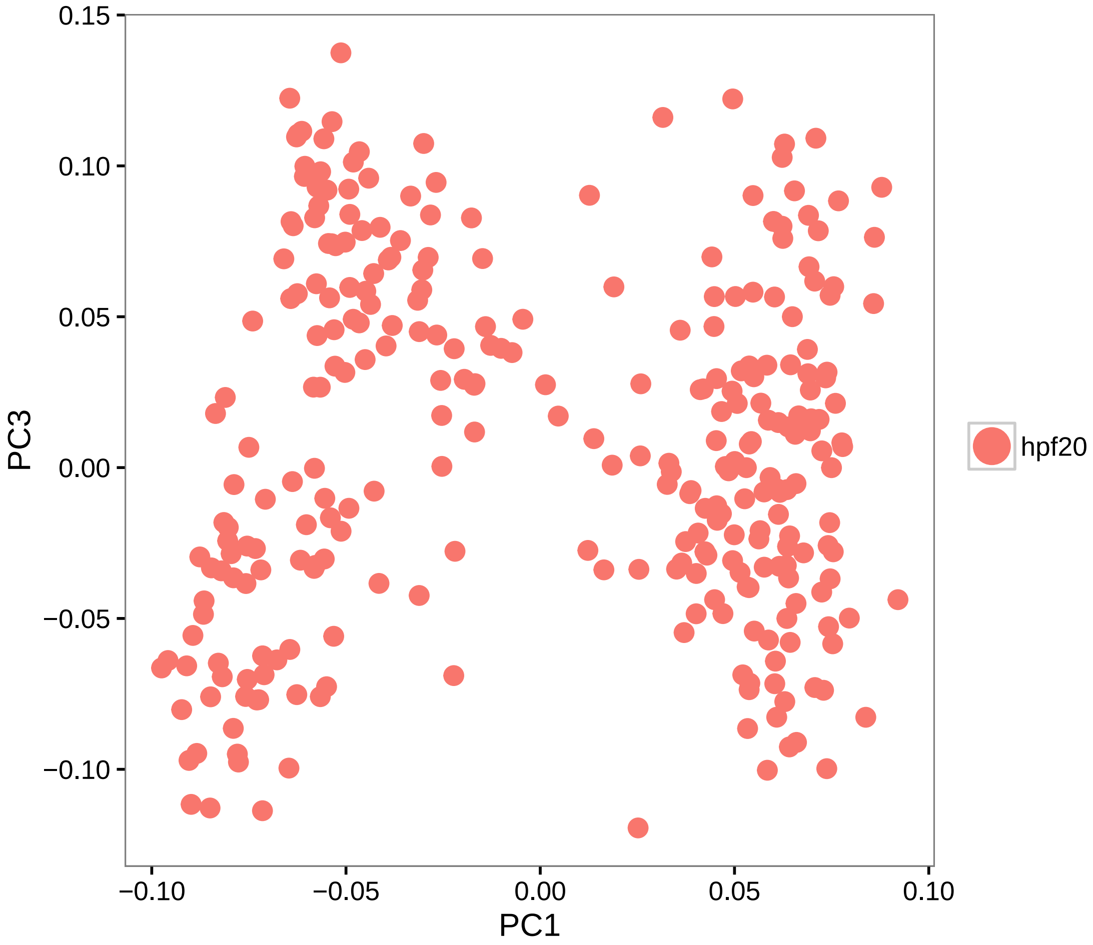


```r
# Calculate PCA scores for all genes (PCA projection)
hpf20 = project.pca(hpf20, do.print = F)

# Visualize the full projected PCA, which now includes new genes which were
# not previously (use.full=TRUE)
pcHeatmap(hpf20, 1, use.full = T, do.balanced = T, col.use = col)
```

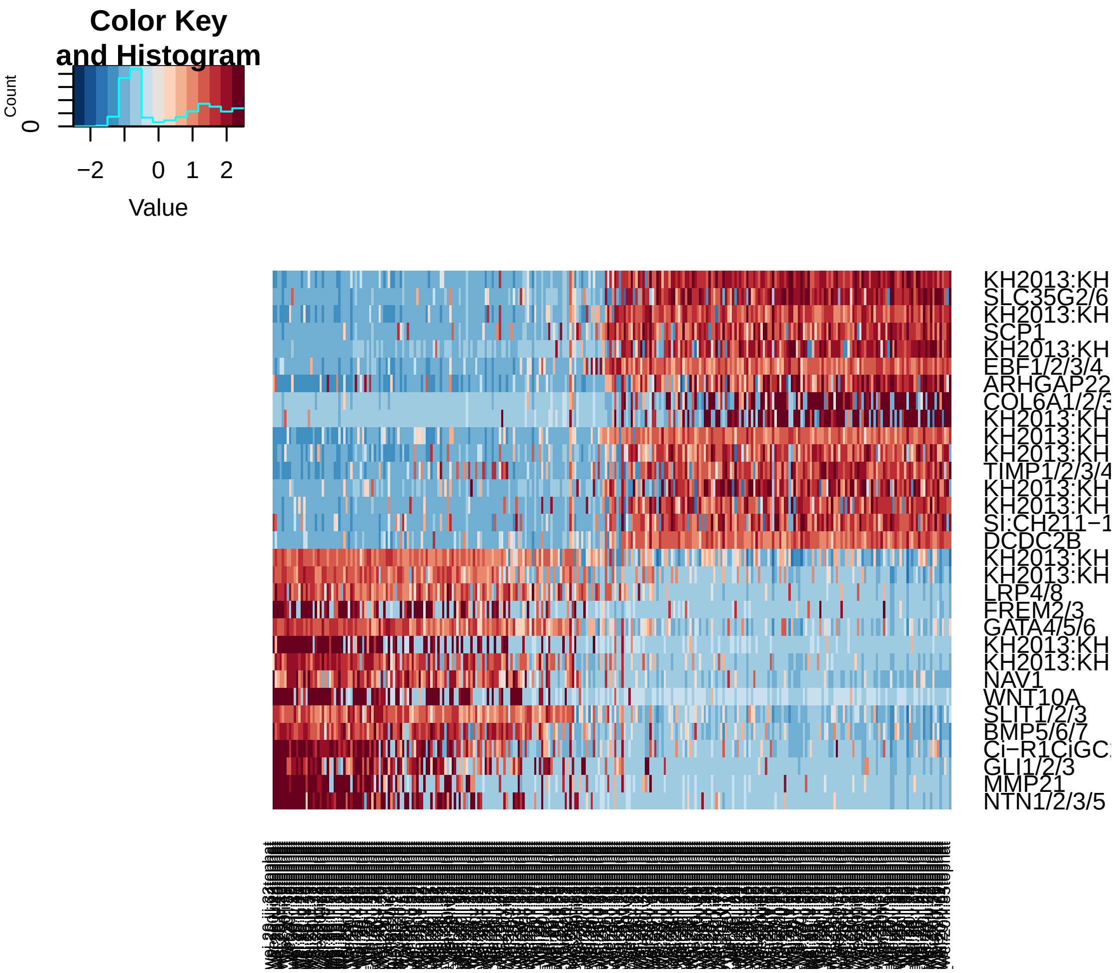


```r
pcHeatmap(hpf20, 2, use.full = T, do.balanced = T, col.use = col)
```

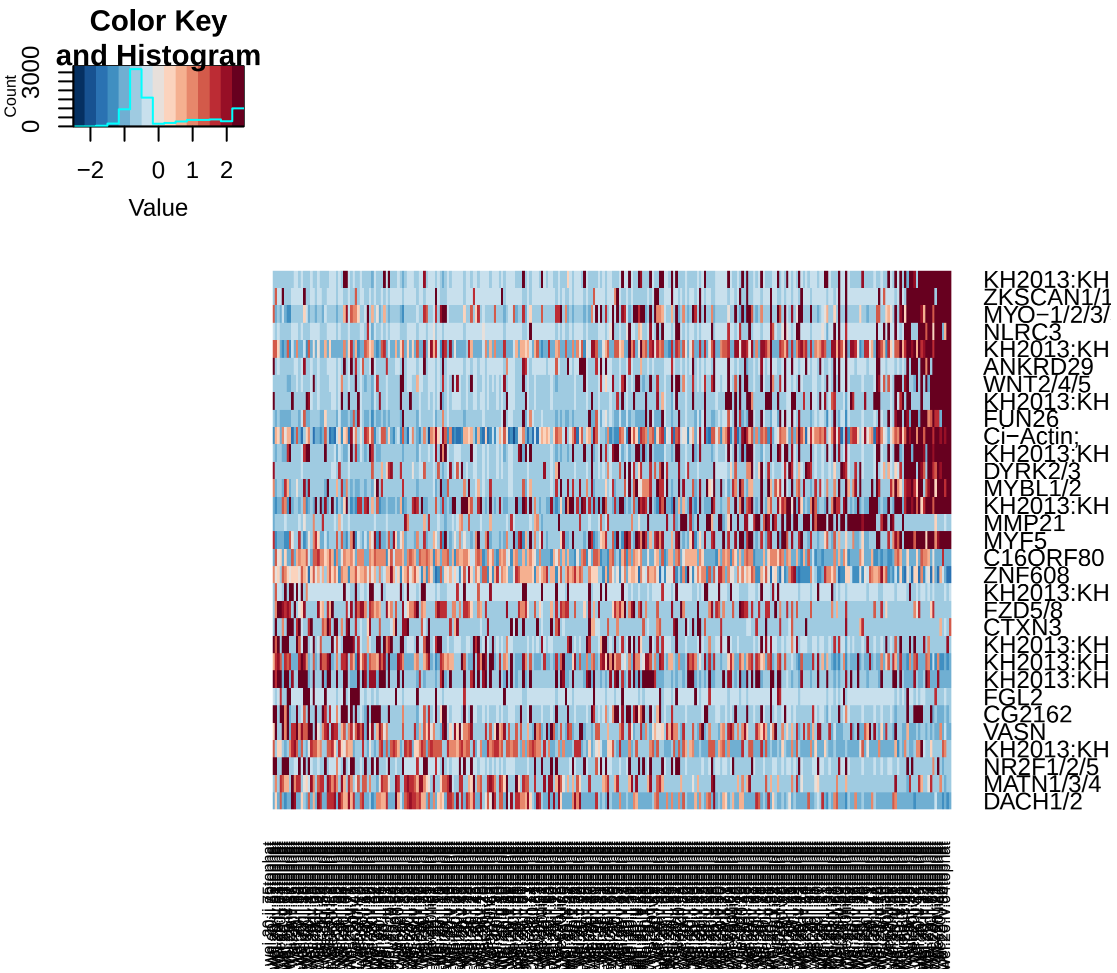


```r
pcHeatmap(hpf20, 3, use.full = T, do.balanced = T, col.use = col)
```

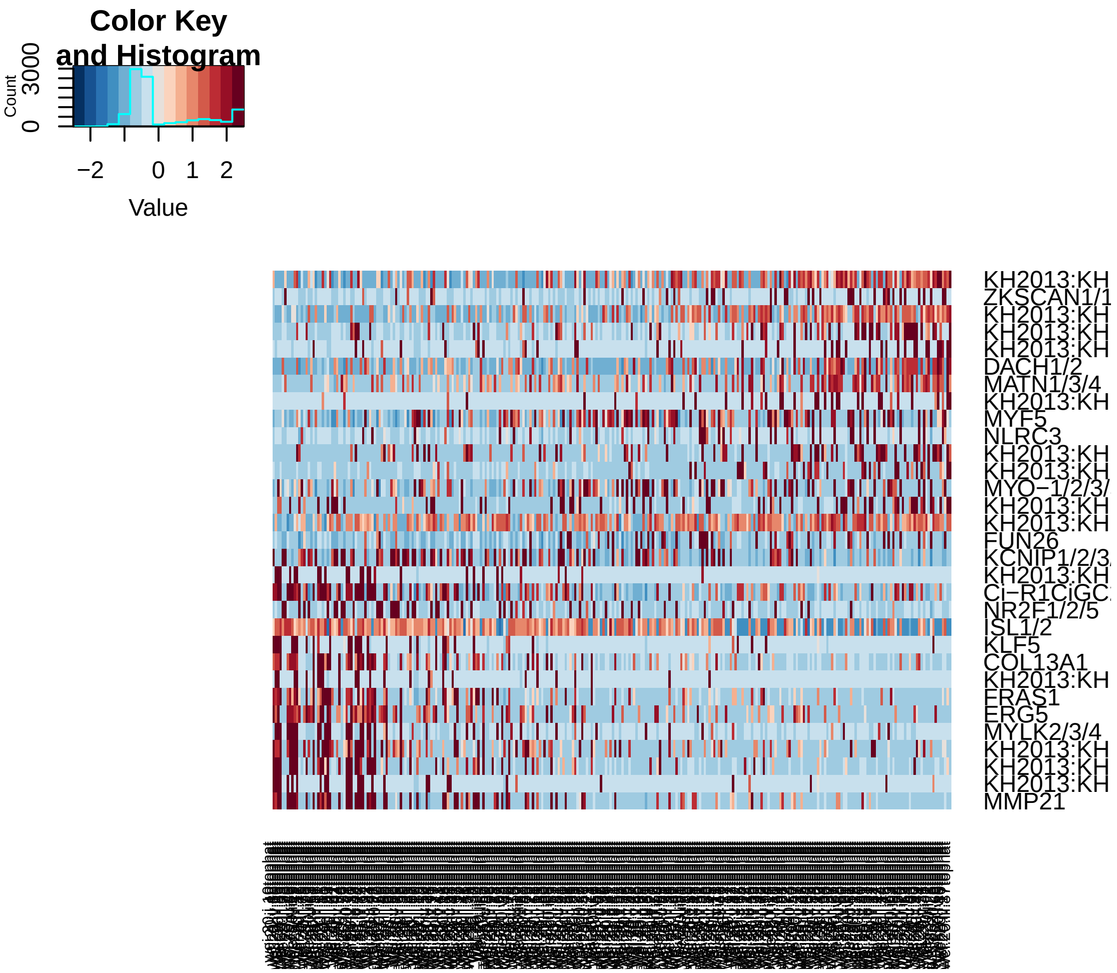


```r
# Do 200 random samplings to find significant genes, each time randomly
# permute 1% of genes This returns a 'p-value' for each gene in each PC,
# based on how likely the gene/PC score woud have been observed by chance
hpf20 = jackStraw(hpf20, num.replicate = 200, do.print = F)

# The jackStraw plot compares the distribution of P-values for each PC with
# a uniform distribution (dashed line) 'Significant' PCs will have a strong
# enrichment of genes with low p-values (solid curve above dashed line)
jackStrawPlot.new(hpf20, PCs = 1:12)
```

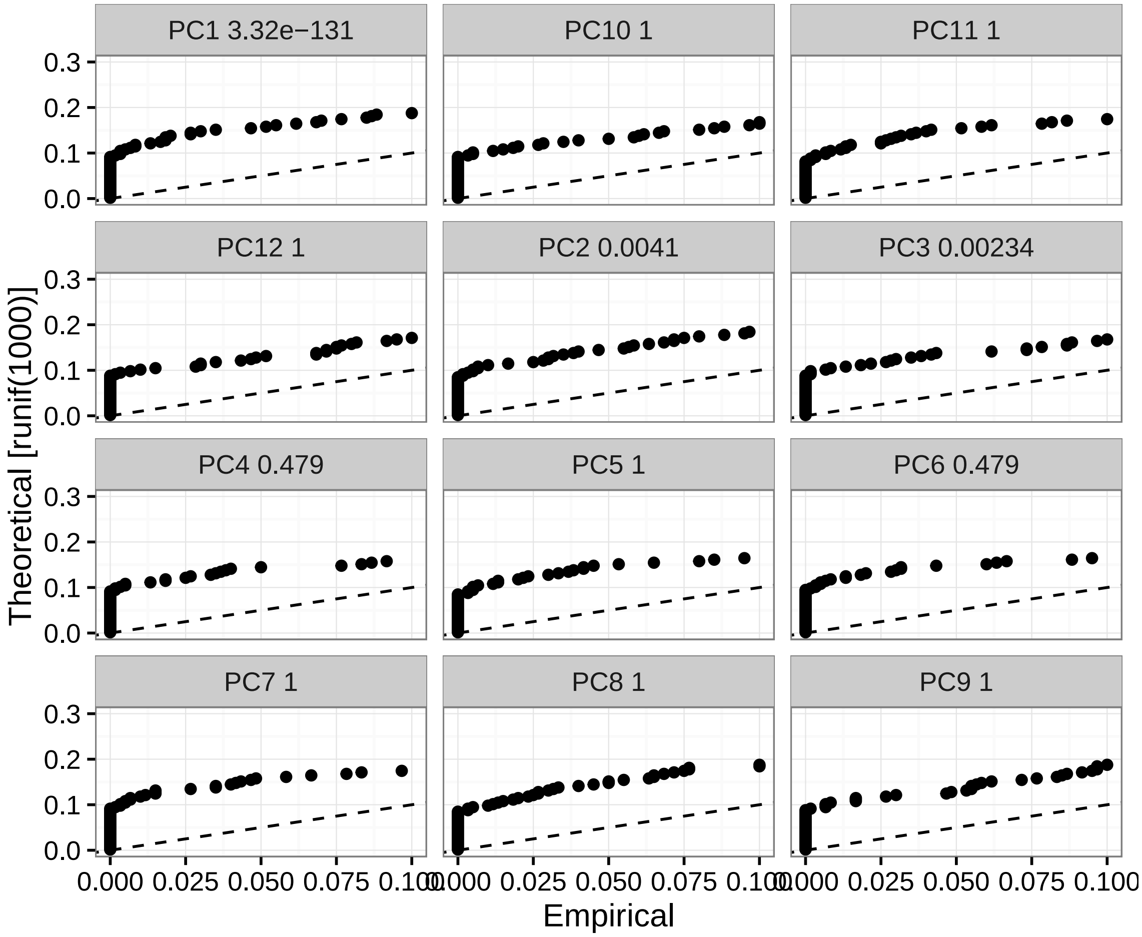


```
# In this case PC1-3 are significant

# Run tSNE using significant PCs as input (spectral tSNE), we get distinct
# point clouds
hpf20 = run_tsne(hpf20, max_iter = 2000, dims.use = 1:3)
tsne.plot(hpf20, do.label = T, label.pt.size = 1)
```

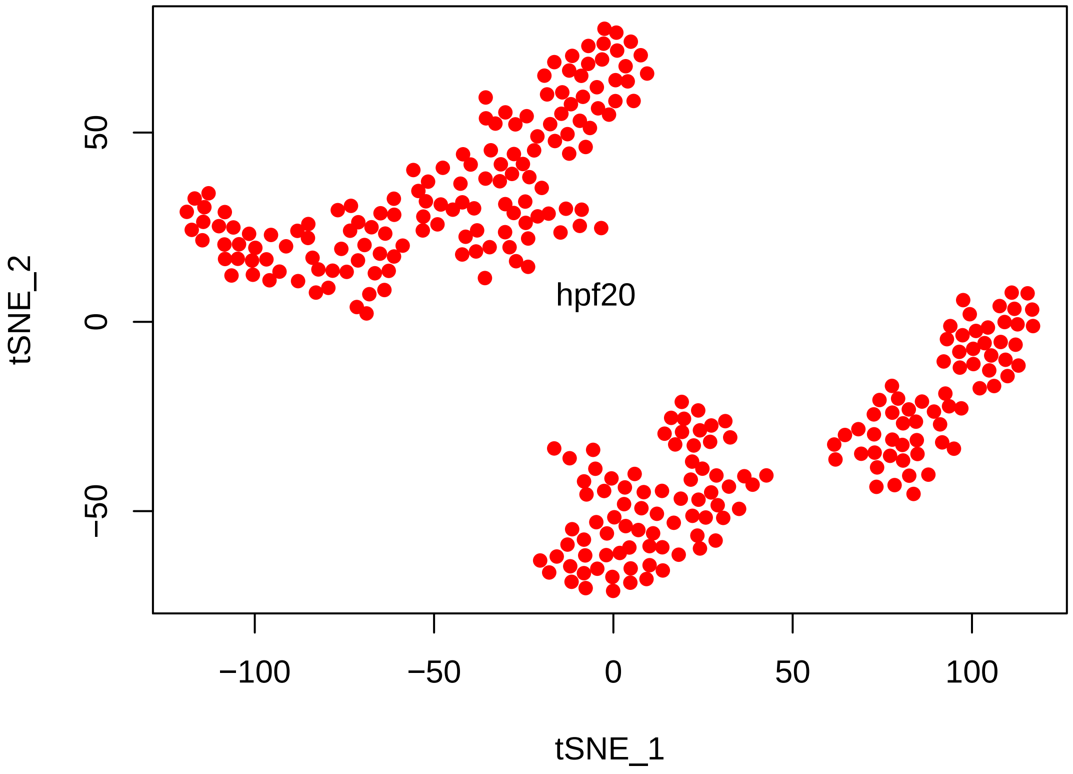


```r
# Find cell clusters using Modularity optimization cluster detection.
hpf20 = FindClusters(hpf20, pc.use = 1:3, do.modularity = T, resolution = 1,
    prune.SNN = 0.1, print.output = 0, k.param = 20)
```

```
## [1] "SNN : processed 72 cells"
## [1] "SNN : processed 144 cells"
## [1] "SNN : processed 216 cells"
## [1] "SNN : processed 288 cells"
```

```r
tsne.plot(hpf20, do.label = T, label.pt.size = 1)
```

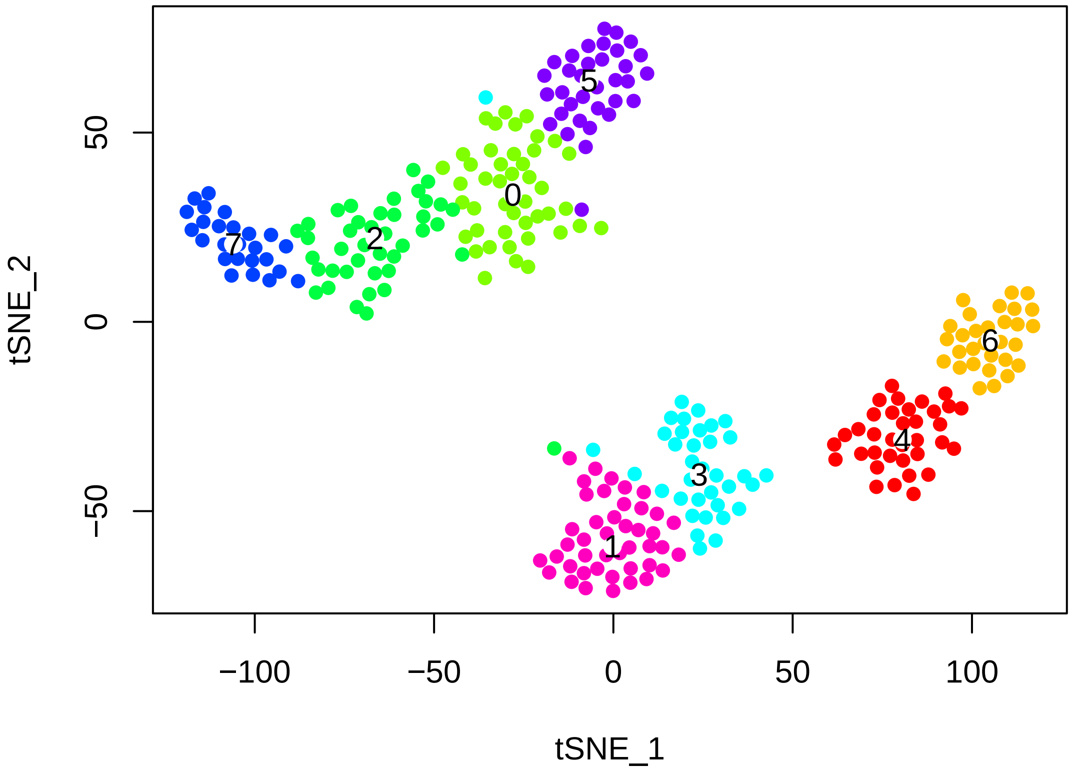


```r
# The validity of the clusters can be validated using a classification
# scheme based on linear SVMs.
hpf20 = BuildSNN(hpf20, pc.use = 1:3, do.sparse = T, k.param = 20)
```

```
## [1] "SNN : processed 72 cells"
## [1] "SNN : processed 144 cells"
## [1] "SNN : processed 216 cells"
## [1] "SNN : processed 288 cells"
```

```r
hpf20 = ValidateClusters(hpf20, pc.use = 1:3, min.connectivity = 0.001, acc.cutoff = 0.8
```

```
## [1] "  0% complete --- merge clusters 0 and 5, classification accuracy of 0.6994"
## [1] " 14% complete --- merge clusters 1 and 3, classification accuracy of 0.7552"
## [1] " 57% complete --- merge clusters 5 and 2, classification accuracy of 0.8306"
## [1] " 57% complete --- merge clusters 4 and 6, classification accuracy of 0.8324"
## [1] "100% complete --- started with 8 clusters, 4 clusters remaining"
```

```r
tsne.plot(hpf20, do.label = T, label.pt.size = 1)
```

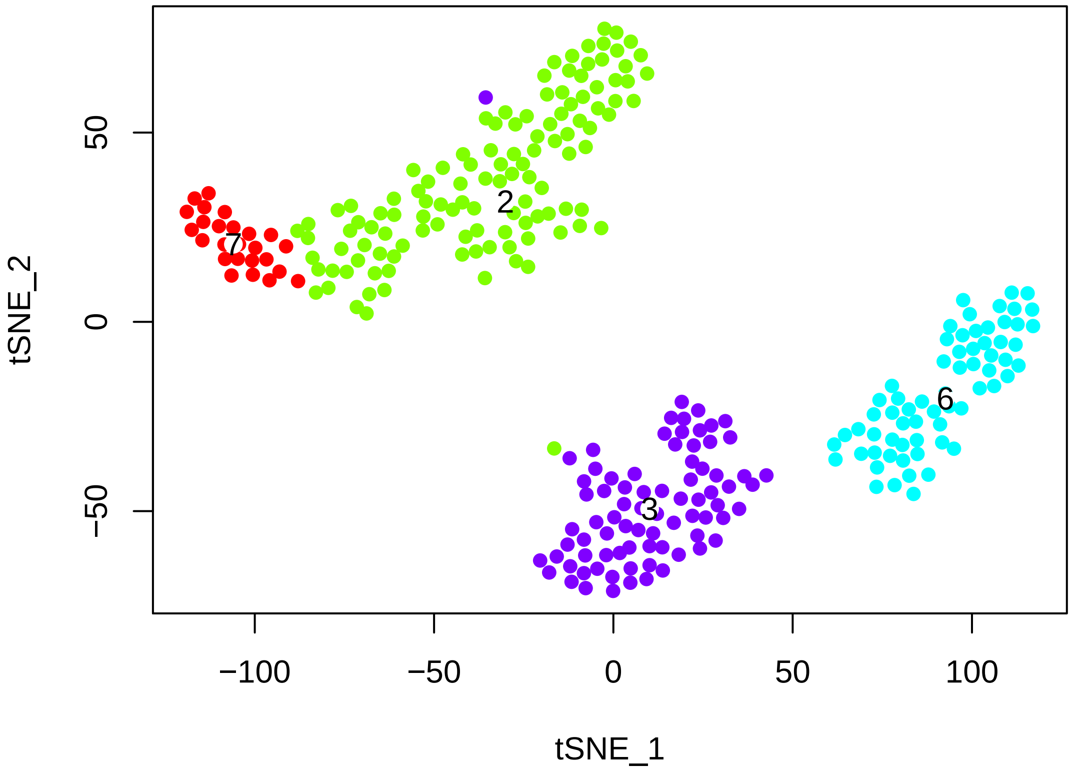


```r
# Find cluster markersusing ROC test with thresh.use = 1, min.pct = 0.5 The
# ROC test returns the 'classification power' for any individual marker
# (ranging from 0 - random, to 1 - perfect). Though not a statistical test,
# it is often very useful for finding clean markers.  Find markers for
# cluster 2
cl2_20.markers = find.markers(hpf20, 2, thresh.use = 1, test.use = "roc", min.pct = 0.5)
head(cl2_20.markers[order(cl2_20.markers$myAUC, decreasing = T), ], 20)
```

```
##                            myAUC avg_diff power pct.1 pct.2
## KH2013:KH.C11.139          0.930 2.042404 0.860 0.992 0.337
## KH2013:KH.L25.4            0.918 1.888086 0.836 0.992 0.231
## KH2013:KH.C5.62_ODF3L2     0.899 1.562319 0.798 0.983 0.278
## KH2013:KH.L18.10_THSD7A    0.894 1.571177 0.788 0.983 0.331
## SCP1                       0.883 1.728818 0.766 0.958 0.278
## KH2013:KH.C4.506_HMCN1     0.882 1.291218 0.764 1.000 0.858
## KH2013:KH.S1269.1          0.880 1.388814 0.760 0.992 0.533
## SLC35G2/6                  0.875 1.853114 0.750 0.899 0.225
## UBE2QL1                    0.874 1.422110 0.748 0.966 0.432
## KH2013:KH.C6.201           0.870 1.898804 0.740 0.874 0.183
## MMP14/15/16/24             0.868 1.387104 0.736 0.975 0.355
## ARHGAP22/24/25             0.868 1.743081 0.736 0.891 0.260
## FRS2/3                     0.866 1.408759 0.732 0.975 0.609
## EBF1/2/3/4                 0.865 1.253061 0.730 1.000 0.314
```

```
## DCDC2B                      0.863 1.184517 0.726 0.983 0.325
## BDH1                        0.862 1.240573 0.724 0.983 0.396
## NOVA1/2                     0.856 1.161588 0.712 1.000 0.462
## Ci-R1CiGC09b24;             0.853 1.216660 0.706 0.983 0.888
## TIMP1/2/3/4                 0.852 1.920971 0.704 0.882 0.349
## HLH-4                       0.845 1.462400 0.690 0.950 0.544
```

```r
# Visualize known markers with a violin plot
vlnPlot(hpf20, c("NKX2-3", "EBF1/2/3/4", "GATA4/5/6"))
```

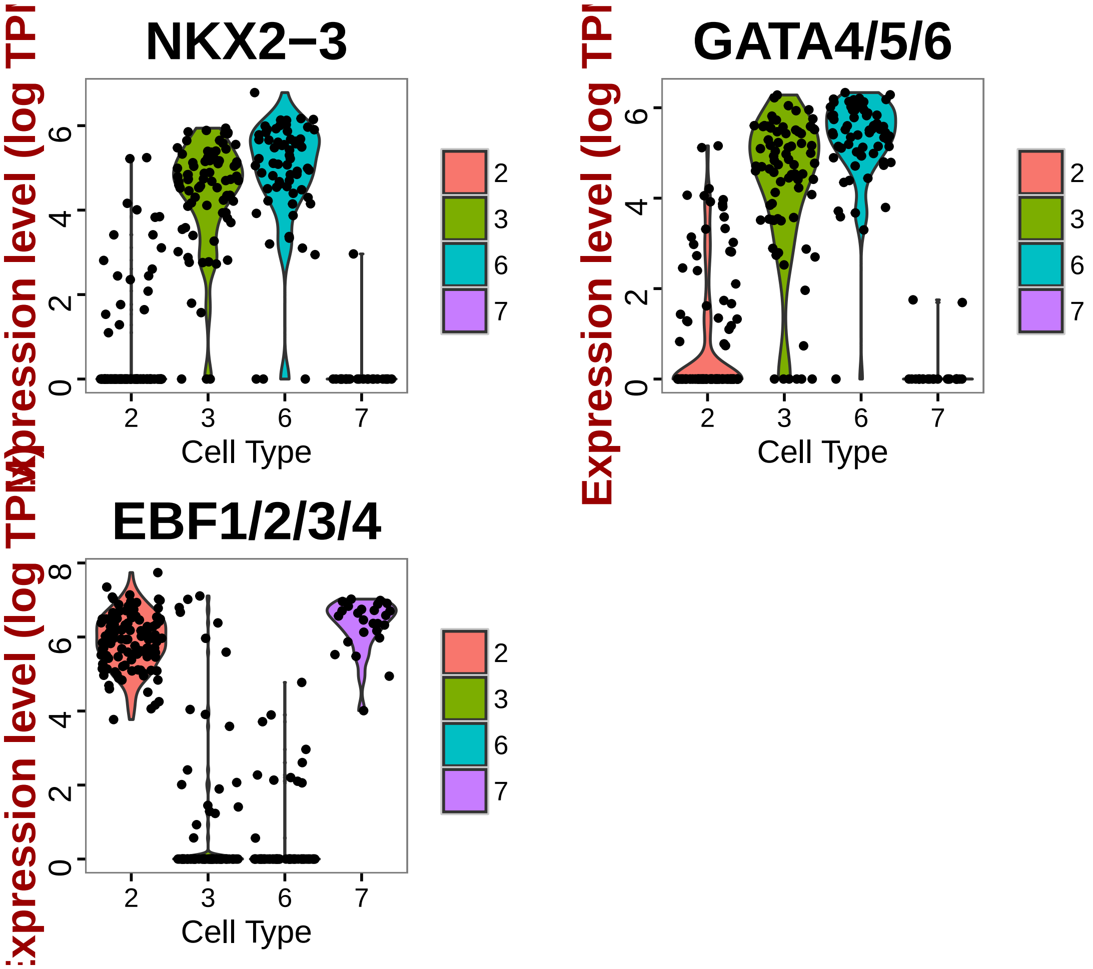


```r
# Based on preliminary studies, these EBF1/2/3/4+ NKX- GATA- cells are ASM1
# cells

# Visualize new markers with a violin plot
vlnPlot(hpf20, c("KH2013:KH.C4.506_HMCN1", "KH2013:KH.C11.139"))
```

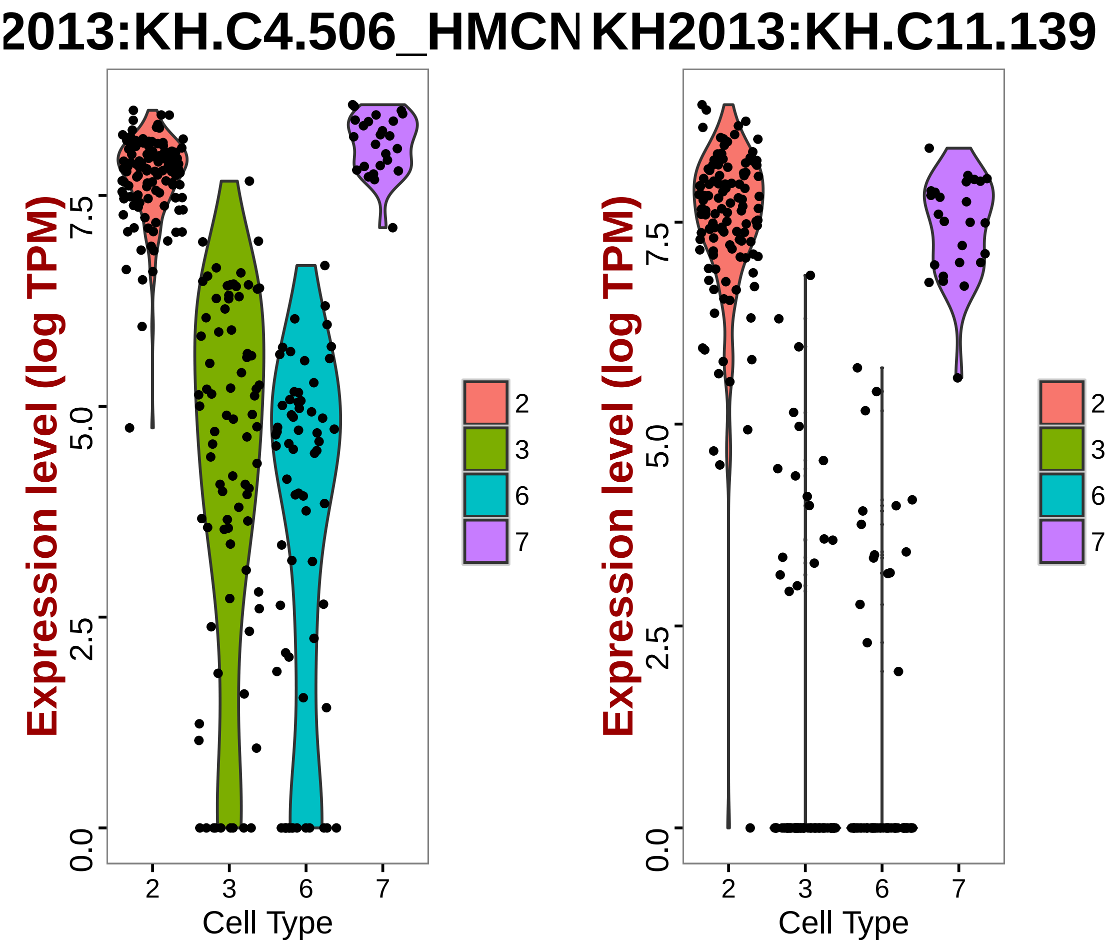


```r
# Find markers for cluster 3
cl3_20.markers = find.markers(hpf20, 3, thresh.use = 1, test.use = "roc", min.pct = 0.5)
head(cl3_20.markers[order(cl3_20.markers$myAUC, decreasing = T), ], 20)
```

```
##                              myAUC  avg_diff power pct.1 pct.2
## KH2013:KH.C1.638_C17ORF105 0.842  1.727786 0.684 0.861 0.325
## KH2013:KH.C4.125_BMP2/4    0.841  1.652993 0.682 0.937 0.517
## DACH1/2                    0.788  1.930722 0.576 0.734 0.278
## KH2013:KH.C4.226           0.787  1.372513 0.574 0.911 0.574
## KH2013:KH.C2.209_CALM1/3   0.775  1.004900 0.550 0.924 0.646
## FOXF1/2                    0.767  1.249064 0.534 0.873 0.512
## YKR070W                    0.750  1.143960 0.500 0.899 0.699
## KH2013:KH.L153.109         0.745  2.030674 0.490 0.595 0.124
## SNX11                      0.739  1.187810 0.478 0.747 0.297
## KH2013:KH.S1887.1          0.716  1.458843 0.432 0.620 0.254
## KH2013:KH.C11.667          0.714  1.038638 0.428 0.886 0.727
## PCTP                       0.704  1.024480 0.408 0.709 0.368
## MATN1/3/4                  0.695  1.452748 0.390 0.633 0.368
## HS3ST3L                    0.685  1.075695 0.370 0.671 0.378
```

```
## TBX1/10                    0.681  1.420956 0.362 0.633 0.373
## KANK1/2/3/4                0.640  1.073334 0.280 0.595 0.411
## KH2013:KH.C9.40_GSTO1/2    0.615  1.051532 0.230 0.506 0.335
## PIGT                       0.502 -5.064610 0.004 0.570 0.569
## RBM15B                     0.433 -3.685868 0.134 0.646 0.789
## PSMD14                     0.430 -3.980281 0.140 0.835 0.928
```

```r
# Visualize known markers with a violin plot
vlnPlot(hpf20, c("NKX2-3", "EBF1/2/3/4", "GATA4/5/6", "TBX1/10"))
```

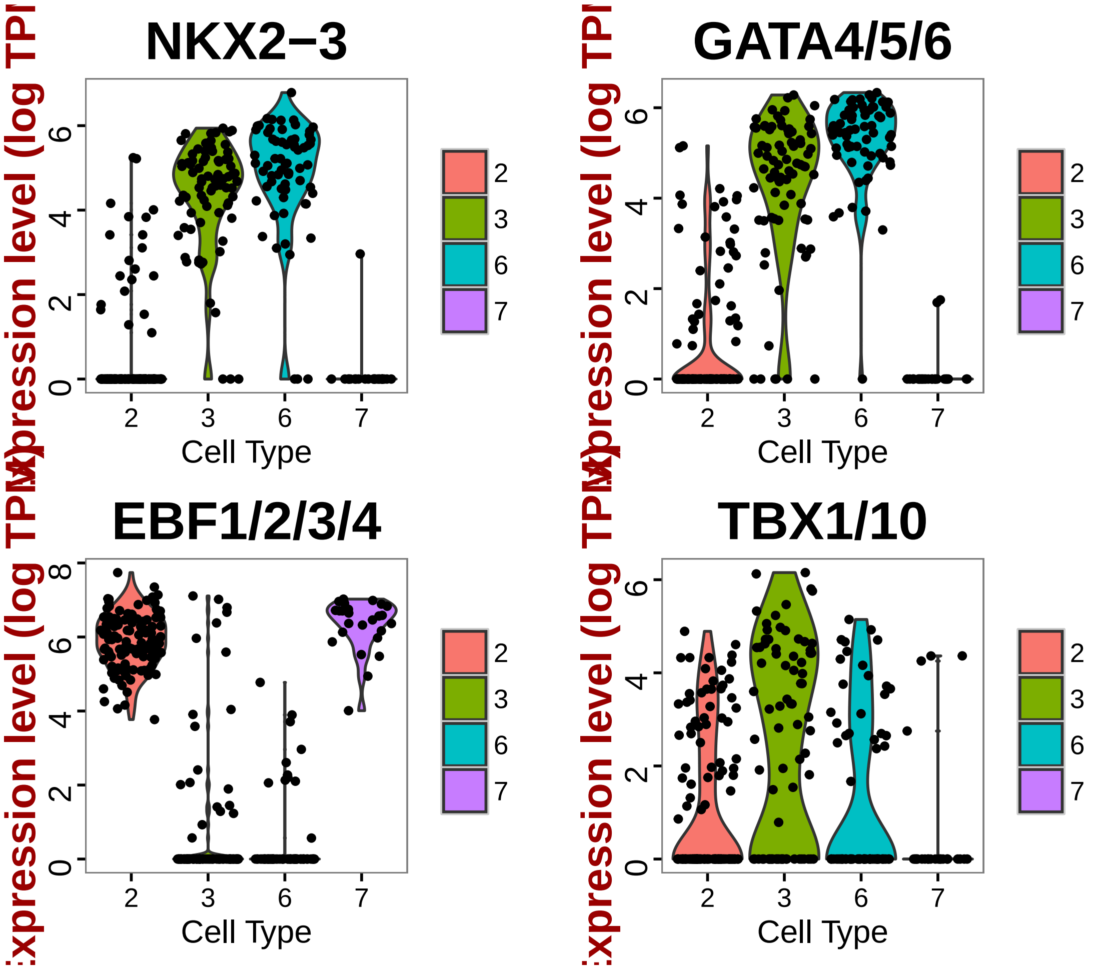


```r
# Based on preliminary studies, these EBF1/2/3/4- NKX+ GATA- TBX1/10+ cells
# are SHP cells

# Visualize new markers with a violin plot
vlnPlot(hpf20, c("KH2013:KH.C1.638_C17ORF105", "DACH1/2"))
```

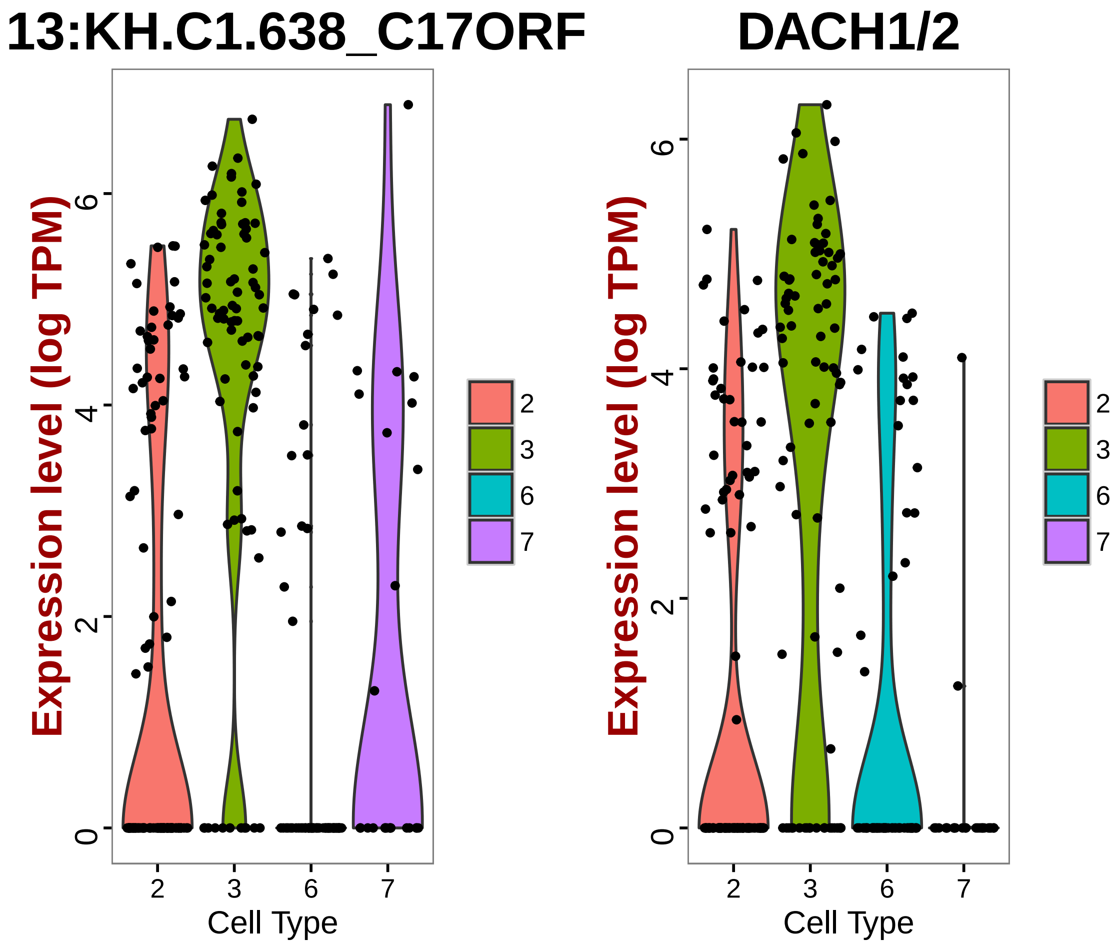


```r
# Find markers for cluster 6
cl6_20.markers = find.markers(hpf20, 6, thresh.use = 1, test.use = "roc", min.pct = 0.5)
head(cl6_20.markers[order(cl6_20.markers$myAUC, decreasing = T), ], 20)
```

```
##                                                myAUC avg_diff power pct.1
## Ci-R1CiGC27a04                                 0.962 3.284688 0.924 0.954
## KH2013:KH.C4.547_BMP2/4                        0.951 2.195473 0.902 0.985
## KH2013:KH.C5.227_F56C4.4                       0.944 1.991633 0.888 1.000
## KH2013:KH.C11.378_CLDN1/10/14/19/2/3/4/5/6/7/9 0.937 1.539702 0.874 1.000
## KH2013:KH.C1.479_NEB                           0.927 1.680541 0.854 1.000
## MMP21                                          0.927 3.901756 0.854 0.877
## FRAS1                                          0.911 2.683945 0.822 0.831
## SFRP1/5                                        0.910 2.269762 0.820 0.923
## KH2013:KH.C10.203                              0.909 2.466237 0.818 0.923
## COL13A1                                        0.907 3.395970 0.814 0.846
## KH2013:KH.C1.953_CG32702                       0.906 2.177912 0.812 0.938
## KH2013:KH.C7.205_ASB2                          0.904 1.818015 0.808 0.969
## NTN1/2/3/5                                     0.902 2.350696 0.804 0.892
## KH2013:KH.C2.994_RNF149                        0.900 1.278227 0.800 1.000
```

```
## KH2013:KH.C3.665_ZAN                          0.896 1.455140 0.792 0.985
## MTDH                                          0.895 1.022437 0.790 1.000
## C5ORF48                                       0.894 2.399851 0.788 0.892
## GATA4/5/6                                     0.891 1.513036 0.782 0.985
## KH2013:KH.C3.191_WBSCR27                      0.886 1.950294 0.772 0.938
## KH2013:KH.C1.738_CG9550                       0.881 1.235709 0.762 0.985
##                                               pct.2
## Ci-R1CiGC27a04                                0.238
## KH2013:KH.C4.547_BMP2/4                       0.381
## KH2013:KH.C5.227_F56C4.4                      0.973
## KH2013:KH.C11.378_CLDN1/10/14/19/2/3/4/5/6/7/9 0.946
## KH2013:KH.C1.479_NEB                          0.605
## MMP21                                         0.117
## FRAS1                                         0.117
## SFRP1/5                                       0.193
## KH2013:KH.C10.203                             0.247
## COL13A1                                       0.108
## KH2013:KH.C1.953_CG32702                      0.309
## KH2013:KH.C7.205_ASB2                         0.430
## NTN1/2/3/5                                    0.166
## KH2013:KH.C2.994_RNF149                       0.857
## KH2013:KH.C3.665_ZAN                          0.628
## MTDH                                          0.991
## C5ORF48                                       0.229
## GATA4/5/6                                     0.471
## KH2013:KH.C3.191_WBSCR27                      0.435
## KH2013:KH.C1.738_CG9550                       0.897
```

```r
# Visualize known markers with a violin plot
vlnPlot(hpf20, c("NKX2-3", "EBF1/2/3/4", "TBX1/10"))
```

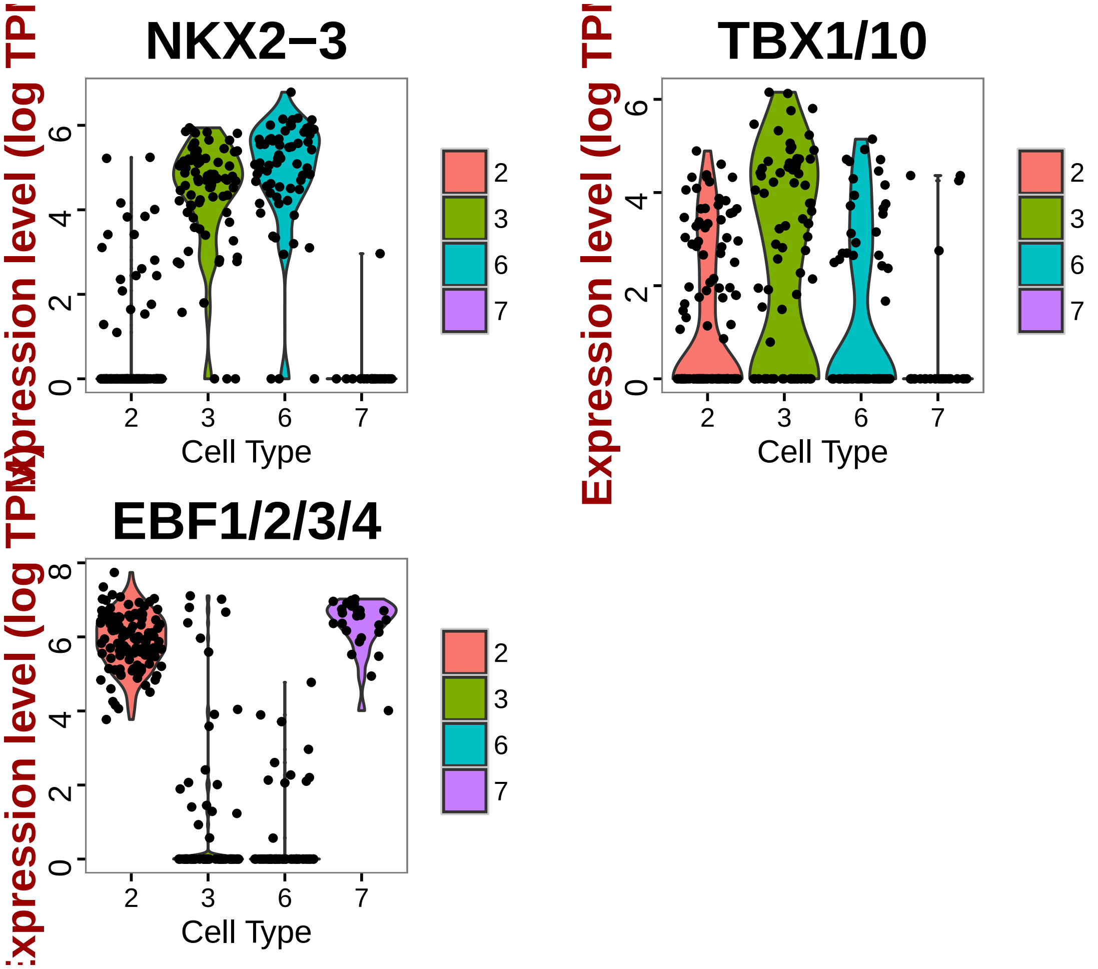


```r
# Confirmed with theses markers cluster 5 NKX+ EBF1/2/3/4- TBX1/10- cells
# are FHP cells

# Visualize new markers with a violin plot
vlnPlot(hpf20, c("MMP21", "FRAS1"))
```

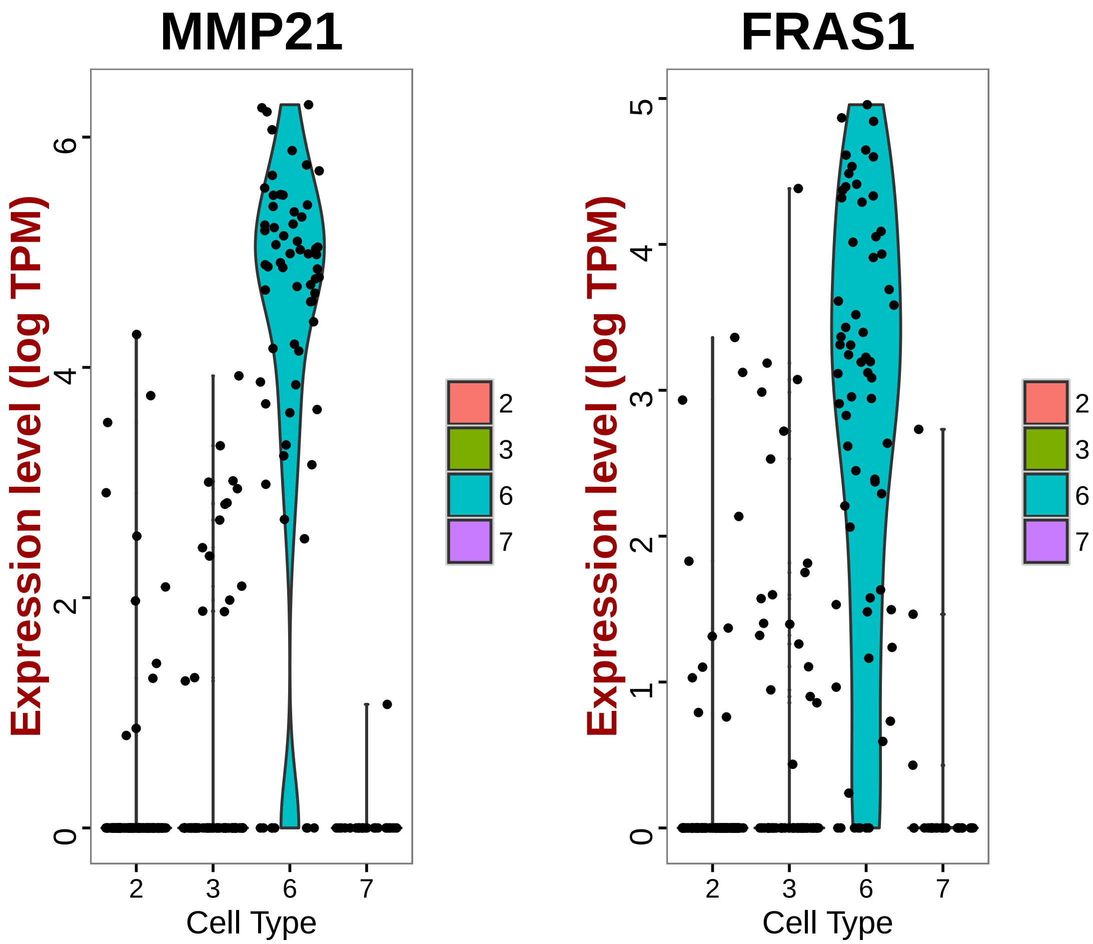


```
# Find markers for cluster 7
cl7_20.markers = find.markers(hpf20, 7, thresh.use = 1, test.use = "roc", min.pct = 0.5)
head(cl7_20.markers[order(cl7_20.markers$myAUC, decreasing = T), ], 20)
```

```
##                          myAUC avg_diff power pct.1 pct.2
## MYF5                     0.936 2.237196 0.872  1.00 0.403
## KH2013:KH.C8.649_ACTA1/2 0.927 2.296572 0.854  1.00 0.589
## DCDC2B                   0.914 1.380557 0.828  1.00 0.559
## MYO-1/2/3/5              0.914 2.429771 0.828  0.92 0.278
## CALML6                   0.905 2.418975 0.810  0.96 0.304
## FUN26                    0.905 2.611697 0.810  0.88 0.106
## RBM24/38                 0.902 1.965112 0.804  0.96 0.384
## SMYD1                    0.902 1.604893 0.804  1.00 0.802
## MYO10                    0.899 1.617349 0.798  1.00 0.441
## Ci-Actin;                0.894 2.092404 0.788  0.96 0.738
## KH2013:KH.C4.506_HMCN1   0.887 1.052516 0.774  1.00 0.909
## KH2013:KH.C5.522_SLC6A13 0.887 1.453291 0.774  1.00 0.471
## ZKSCAN1/14/17/2/3/6      0.885 3.396235 0.770  0.80 0.080
## KH2013:KH.C12.669_PHEX   0.883 1.820194 0.766  0.96 0.407
```

```
## KH2013:KH.C7.476_CHRNB1     0.880 1.952671 0.760  0.92 0.304
## KH2013:KH.C12.419_TSPAN4/9 0.878 1.252526 0.756  1.00 0.821
## KH2013:KH.C1.509           0.876 2.059997 0.752  0.88 0.205
## KH2013:KH.S643.6_CDKN1B    0.876 1.562963 0.752  1.00 0.612
## RBFOX1/2/3                 0.875 1.347928 0.750  1.00 0.863
## KH2013:KH.C5.404_MBL1/2    0.875 2.442091 0.750  0.84 0.114
```

```r
# Visualize known markers with a violin plot
vlnPlot(hpf20, c("NKX2-3", "EBF1/2/3/4", "TBX1/10"))
```

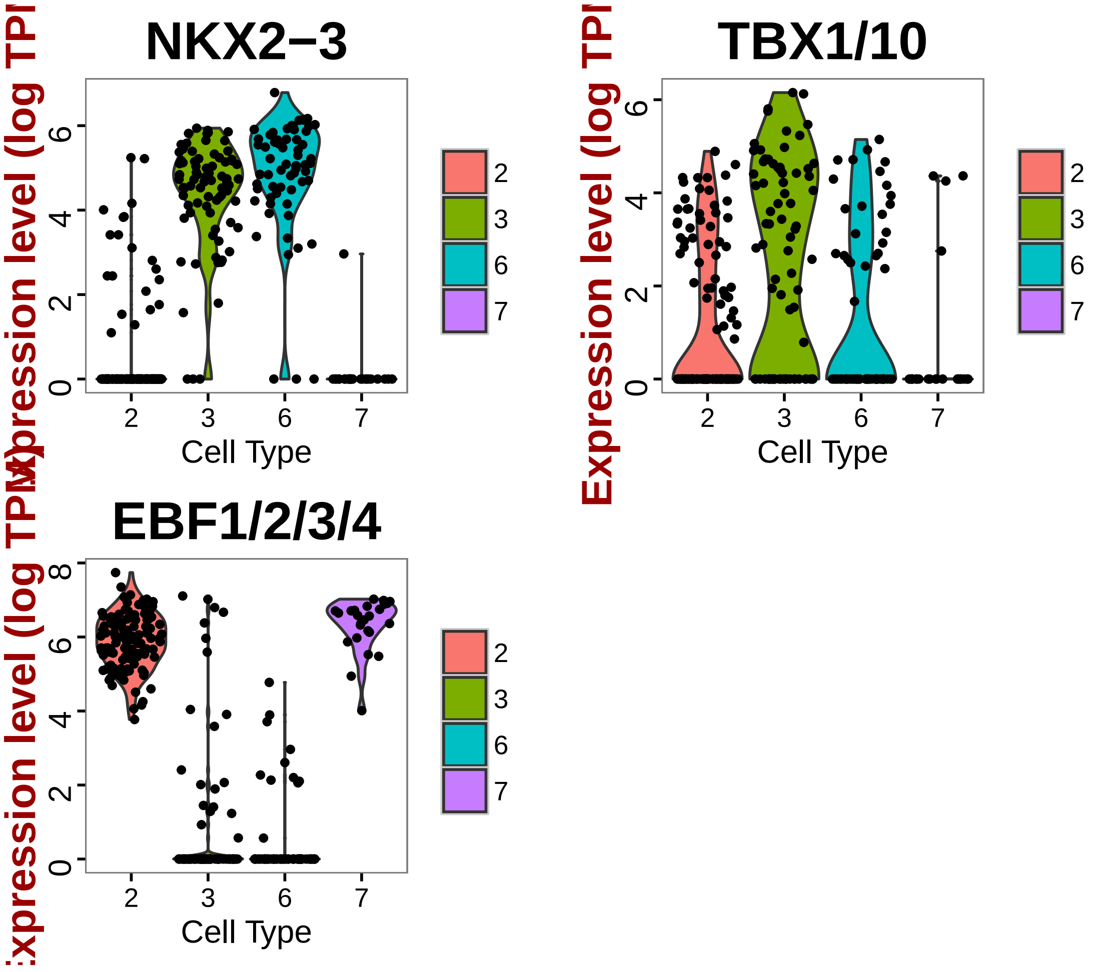


```r
# Confirmed with theses markers cluster 5 NKX+ EBF1/2/3/4- TBX1/10- cells
# are ASM2 cells

# Visualize new markers with a violin plot
vlnPlot(hpf20, c("MYF5", "CALML6"))
```

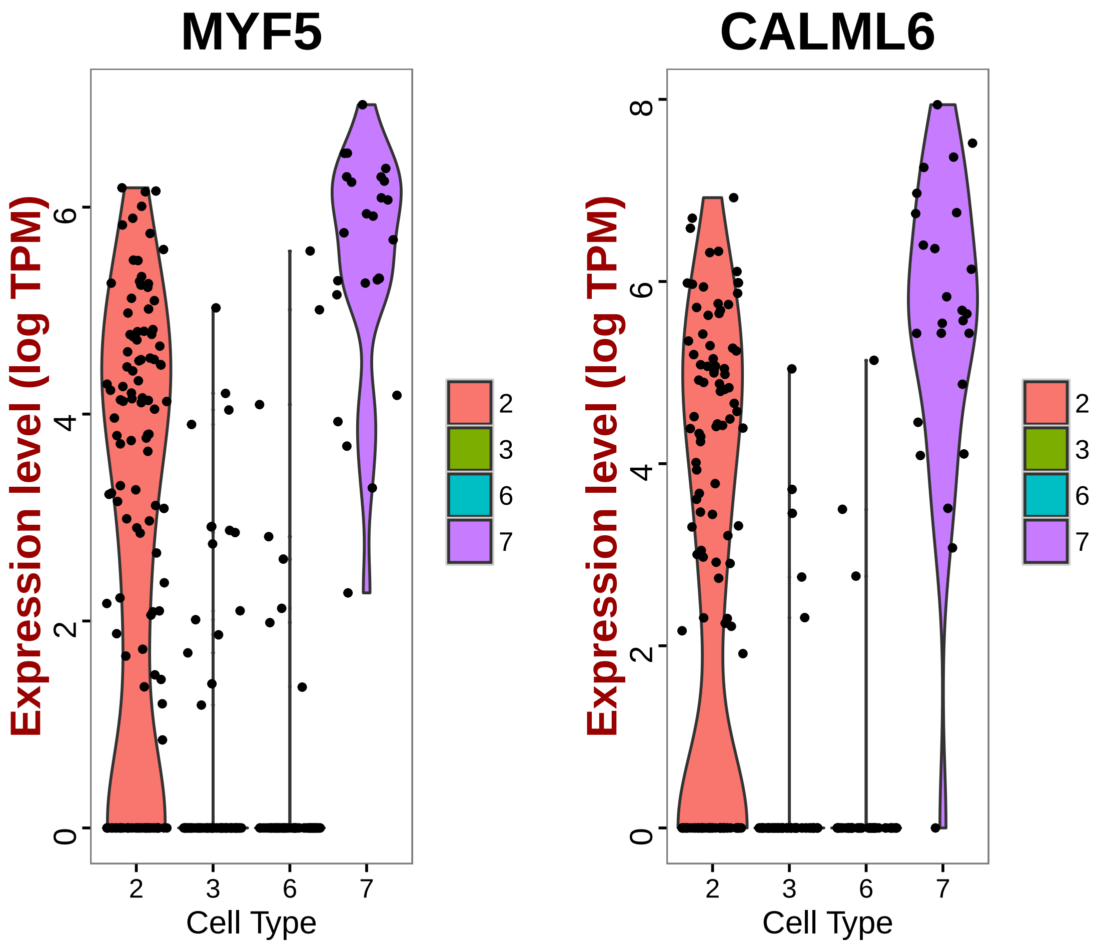


```
# MYF5 is a more differentiated marker, this group of ASM is more
# deferentiated ASM cells.

# Write cell names into text files
write.table(which.cells(hpf20, 2), file = "20ASM1Cells.txt", sep = "\t")
write.table(which.cells(hpf20, 3), file = "20SHPCells.txt", sep = "\t")
write.table(which.cells(hpf20, 6), file = "20FHPCells.txt", sep = "\t")
write.table(which.cells(hpf20, 7), file = "20ASM2Cells.txt", sep = "\t")

# Rename cluster identities
hpf20 = rename.ident(hpf20, 2, "20ASM1")
hpf20 = rename.ident(hpf20, 3, "20SHP")
hpf20 = rename.ident(hpf20, 6, "20FHP")
hpf20 = rename.ident(hpf20, 7, "20ASM2")

# Visualize tSNE used color scheme FHP-red, SHP-orange, ASM-blue
tsne.plot(hpf20, do.label = T, label.pt.size = 1, label.cex.text = 1.2, label.cols.use =
    "blue", "red", "orange"))
```

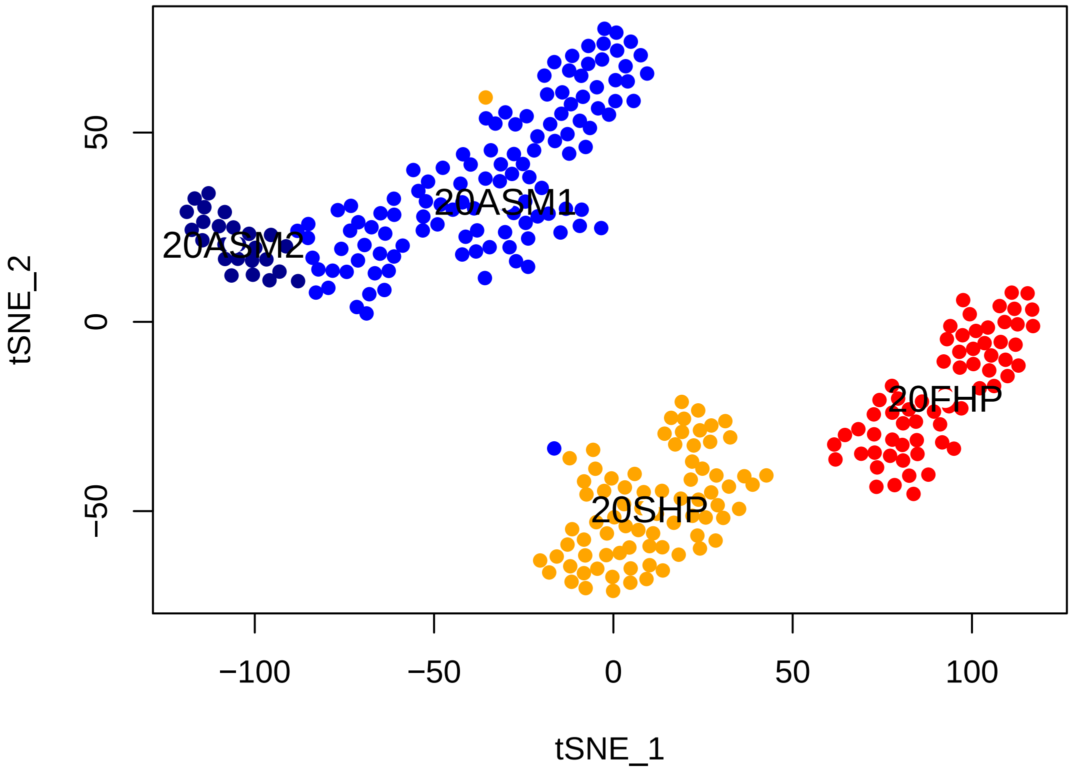


```
# Store FHP markers in text file
FHP_20.markers = cl6_20.markers
head(FHP_20.markers[order(FHP_20.markers$myAUC, decreasing = T), ], 20)
```

```
##                                                myAUC avg_diff power pct.1
## Ci-R1CiGC27a04                                 0.962 3.284688 0.924 0.954
## KH2013:KH.C4.547_BMP2/4                        0.951 2.195473 0.902 0.985
## KH2013:KH.C5.227_F56C4.4                       0.944 1.991633 0.888 1.000
## KH2013:KH.C11.378_CLDN1/10/14/19/2/3/4/5/6/7/9 0.937 1.539702 0.874 1.000
## KH2013:KH.C1.479_NEB                           0.927 1.680541 0.854 1.000
## MMP21                                          0.927 3.901756 0.854 0.877
## FRAS1                                          0.911 2.683945 0.822 0.831
## SFRP1/5                                        0.910 2.269762 0.820 0.923
## KH2013:KH.C10.203                              0.909 2.466237 0.818 0.923
## COL13A1                                        0.907 3.395970 0.814 0.846
## KH2013:KH.C1.953_CG32702                       0.906 2.177912 0.812 0.938
## KH2013:KH.C7.205_ASB2                          0.904 1.818015 0.808 0.969
## NTN1/2/3/5                                     0.902 2.350696 0.804 0.892
## KH2013:KH.C2.994_RNF149                        0.900 1.278227 0.800 1.000
## KH2013:KH.C3.665_ZAN                           0.896 1.455140 0.792 0.985
## MTDH                                           0.895 1.022437 0.790 1.000
## C5ORF48                                        0.894 2.399851 0.788 0.892
## GATA4/5/6                                      0.891 1.513036 0.782 0.985
```

```
## KH2013:KH.C3.191_WBSCR27                      0.886 1.950294 0.772 0.938
## KH2013:KH.C1.738_CG9550                       0.881 1.235709 0.762 0.985
##                                               pct.2
## Ci-R1CiGC27a04                                0.238
## KH2013:KH.C4.547_BMP2/4                       0.381
## KH2013:KH.C5.227_F56C4.4                      0.973
## KH2013:KH.C11.378_CLDN1/10/14/19/2/3/4/5/6/7/9 0.946
## KH2013:KH.C1.479_NEB                          0.605
## MMP21                                         0.117
## FRAS1                                         0.117
## SFRP1/5                                       0.193
## KH2013:KH.C10.203                             0.247
## COL13A1                                       0.108
## KH2013:KH.C1.953_CG32702                      0.309
## KH2013:KH.C7.205_ASB2                         0.430
## NTN1/2/3/5                                    0.166
## KH2013:KH.C2.994_RNF149                       0.857
## KH2013:KH.C3.665_ZAN                          0.628
## MTDH                                          0.991
## C5ORF48                                       0.229
## GATA4/5/6                                     0.471
## KH2013:KH.C3.191_WBSCR27                      0.435
## KH2013:KH.C1.738_CG9550                       0.897
```

```
write.table(FHP_20.markers, file = "FHP_20.markers.txt", sep = "\t")

# Store SHP markers in text file
SHP_20.markers = cl3_20.markers
head(SHP_20.markers[order(SHP_20.markers$myAUC, decreasing = T), ], 20)
```

```
##                              myAUC  avg_diff power pct.1 pct.2
## KH2013:KH.C1.638_C17ORF105 0.842  1.727786 0.684 0.861 0.325
## KH2013:KH.C4.125_BMP2/4    0.841  1.652993 0.682 0.937 0.517
## DACH1/2                    0.788  1.930722 0.576 0.734 0.278
## KH2013:KH.C4.226           0.787  1.372513 0.574 0.911 0.574
## KH2013:KH.C2.209_CALM1/3   0.775  1.004900 0.550 0.924 0.646
## FOXF1/2                    0.767  1.249064 0.534 0.873 0.512
## YKR070W                    0.750  1.143960 0.500 0.899 0.699
## KH2013:KH.L153.109         0.745  2.030674 0.490 0.595 0.124
## SNX11                      0.739  1.187810 0.478 0.747 0.297
## KH2013:KH.S1887.1          0.716  1.458843 0.432 0.620 0.254
## KH2013:KH.C11.667          0.714  1.038638 0.428 0.886 0.727
## PCTP                       0.704  1.024480 0.408 0.709 0.368
## MATN1/3/4                  0.695  1.452748 0.390 0.633 0.368
```

```
## HS3ST3L                      0.685  1.075695 0.370 0.671 0.378
## TBX1/10                      0.681  1.420956 0.362 0.633 0.373
## KANK1/2/3/4                  0.640  1.073334 0.280 0.595 0.411
## KH2013:KH.C9.40_GSTO1/2      0.615  1.051532 0.230 0.506 0.335
## PIGT                         0.502 -5.064610 0.004 0.570 0.569
## RBM15B                       0.433 -3.685868 0.134 0.646 0.789
## PSMD14                       0.430 -3.980281 0.140 0.835 0.928
```

```
write.table(SHP_20.markers, file = "SHP_20.markers.txt", sep = "\t")

# Store ASM1 markers in text file
ASM1_20.markers = cl2_20.markers
head(ASM1_20.markers[order(ASM1_20.markers$myAUC, decreasing = T), ], 20)
```

```
##                           myAUC avg_diff power pct.1 pct.2
## KH2013:KH.C11.139         0.930 2.042404 0.860 0.992 0.337
## KH2013:KH.L25.4           0.918 1.888086 0.836 0.992 0.231
## KH2013:KH.C5.62_ODF3L2    0.899 1.562319 0.798 0.983 0.278
## KH2013:KH.L18.10_THSD7A   0.894 1.571177 0.788 0.983 0.331
## SCP1                      0.883 1.728818 0.766 0.958 0.278
## KH2013:KH.C4.506_HMCN1    0.882 1.291218 0.764 1.000 0.858
## KH2013:KH.S1269.1         0.880 1.388814 0.760 0.992 0.533
## SLC35G2/6                 0.875 1.853114 0.750 0.899 0.225
## UBE2QL1                   0.874 1.422110 0.748 0.966 0.432
## KH2013:KH.C6.201          0.870 1.898804 0.740 0.874 0.183
## MMP14/15/16/24            0.868 1.387104 0.736 0.975 0.355
## ARHGAP22/24/25            0.868 1.743081 0.736 0.891 0.260
## FRS2/3                    0.866 1.408759 0.732 0.975 0.609
## EBF1/2/3/4                0.865 1.253061 0.730 1.000 0.314
## DCDC2B                    0.863 1.184517 0.726 0.983 0.325
## BDH1                      0.862 1.240573 0.724 0.983 0.396
## NOVA1/2                   0.856 1.161588 0.712 1.000 0.462
## Ci-R1CiGC09b24;           0.853 1.216660 0.706 0.983 0.888
## TIMP1/2/3/4               0.852 1.920971 0.704 0.882 0.349
## HLH-4                     0.845 1.462400 0.690 0.950 0.544
```

```
write.table(ASM1_20.markers, file = "ASM1_20.markers.txt", sep = "\t")

# Store ASM2 markers in text file
ASM2_20.markers = cl7_20.markers
head(ASM2_20.markers[order(ASM2_20.markers$myAUC, decreasing = T), ], 20)
```

```
##                              myAUC avg_diff power pct.1 pct.2
```

```
## MYF5                         0.936 2.237196 0.872  1.00 0.403
## KH2013:KH.C8.649_ACTA1/2     0.927 2.296572 0.854  1.00 0.589
## DCDC2B                       0.914 1.380557 0.828  1.00 0.559
## MYO-1/2/3/5                  0.914 2.429771 0.828  0.92 0.278
## CALML6                       0.905 2.418975 0.810  0.96 0.304
## FUN26                        0.905 2.611697 0.810  0.88 0.106
## RBM24/38                     0.902 1.965112 0.804  0.96 0.384
## SMYD1                        0.902 1.604893 0.804  1.00 0.802
## MYO10                        0.899 1.617349 0.798  1.00 0.441
## Ci-Actin;                    0.894 2.092404 0.788  0.96 0.738
## KH2013:KH.C4.506_HMCN1       0.887 1.052516 0.774  1.00 0.909
## KH2013:KH.C5.522_SLC6A13     0.887 1.453291 0.774  1.00 0.471
## ZKSCAN1/14/17/2/3/6          0.885 3.396235 0.770  0.80 0.080
## KH2013:KH.C12.669_PHEX       0.883 1.820194 0.766  0.96 0.407
## KH2013:KH.C7.476_CHRNB1      0.880 1.952671 0.760  0.92 0.304
## KH2013:KH.C12.419_TSPAN4/9 0.878 1.252526 0.756  1.00 0.821
## KH2013:KH.C1.509             0.876 2.059997 0.752  0.88 0.205
## KH2013:KH.S643.6_CDKN1B      0.876 1.562963 0.752  1.00 0.612
## RBFOX1/2/3                   0.875 1.347928 0.750  1.00 0.863
## KH2013:KH.C5.404_MBL1/2      0.875 2.442091 0.750  0.84 0.114
```

```
write.table(ASM2_20.markers, file = "ASM2_20.markers.txt", sep = "\t")

# Find pan Heart Progenitor markers
panHP_20.markers = find.markers(hpf20, c("20SHP", "20FHP"), thresh.use = 1,
    test.use = "roc", min.pct = 0.5)
head(panHP_20.markers[order(panHP_20.markers$myAUC, decreasing = T), ], 20)
```

```
##                                      myAUC avg_diff power pct.1
## KH2013:KH.C2.994_RNF149              0.977 2.285073 0.954 0.993
## SLIT1/2/3                            0.961 3.358580 0.922 0.965
## NKX2-3                               0.959 3.261362 0.918 0.958
## KH2013:KH.C1.738_CG9550              0.953 1.858233 0.906 0.993
## KH2013:KH.C3.665_ZAN                 0.951 3.369149 0.902 0.951
## GATA4/5/6                            0.951 3.224164 0.902 0.944
## QKI                                  0.933 1.992776 0.866 0.986
## KH2013:KH.C10.174                    0.927 2.971638 0.854 0.917
## KH2013:KH.C2.514_ENPP1/2/3           0.927 1.030617 0.854 1.000
## LRP4/8                               0.915 2.558266 0.830 0.889
## UNC5A                                0.913 1.610891 0.826 1.000
## KH2013:KH.C9.650_SULT1ST1/2/3/4      0.910 1.510445 0.820 0.993
## KH2013:KH.C2.209_CALM1/3             0.903 2.226478 0.806 0.944
## KH2013:KH.C4.547_BMP2/4              0.884 3.020256 0.768 0.840
## CG15828                              0.882 1.072115 0.764 1.000
```

```
## KH2013:KH.C11.378_CLDN1/10/14/19/2/3/4/5/6/7/9 0.880 1.799816 0.760 0.972
## NAV1                                            0.879 2.707348 0.758 0.799
## KH2013:KH.C2.935_S1PR1/2/3/4/5                  0.875 1.985909 0.750 0.910
## KH2013:KH.C3.716_EFNA1/2/3/4/5                  0.872 2.300622 0.744 0.868
## FOXG1                                           0.872 2.046786 0.744 0.889
##                                                 pct.2
## KH2013:KH.C2.994_RNF149                         0.785
## SLIT1/2/3                                       0.229
## NKX2-3                                          0.146
## KH2013:KH.C1.738_CG9550                         0.840
## KH2013:KH.C3.665_ZAN                            0.465
## GATA4/5/6                                       0.229
## QKI                                             0.771
## KH2013:KH.C10.174                               0.243
## KH2013:KH.C2.514_ENPP1/2/3                      1.000
## LRP4/8                                          0.111
## UNC5A                                           0.757
## KH2013:KH.C9.650_SULT1ST1/2/3/4                 0.861
## KH2013:KH.C2.209_CALM1/3                        0.500
## KH2013:KH.C4.547_BMP2/4                         0.194
## CG15828                                         0.972
## KH2013:KH.C11.378_CLDN1/10/14/19/2/3/4/5/6/7/9 0.944
## NAV1                                            0.118
## KH2013:KH.C2.935_S1PR1/2/3/4/5                  0.410
## KH2013:KH.C3.716_EFNA1/2/3/4/5                  0.285
## FOXG1                                           0.312
```

```r
write.table(panHP_20.markers, file = "panHP_20.markers.txt", sep = "\t")

# Find SHP specific markers that distinguish tow heart progenitors FHP and
# SHP
SHPspecific_20.markers = find.markers(hpf20, "20SHP", "20FHP", thresh.use = 1,
    test.use = "roc", min.pct = 0.5)
head(SHPspecific_20.markers[order(SHPspecific_20.markers$myAUC, decreasing = T),
    ], 20)
```

```
##                                          myAUC avg_diff power pct.1 pct.2
## KH2013:KH.C1.638_C17ORF105               0.861 2.080418 0.722 0.861 0.246
## DACH1/2                                  0.788 1.960855 0.576 0.734 0.292
## MATN1/3/4                                0.786 3.205297 0.572 0.633 0.138
## PPAP2B                                   0.784 1.536951 0.568 0.886 0.585
## KH2013:KH.C10.172                        0.724 1.018103 0.448 0.759 0.369
## TIMP4                                    0.695 1.525484 0.390 0.608 0.277
## SPINK1/2                                 0.689 1.225729 0.378 0.570 0.215
```

```
## KH2013:KH.L154.26_CSMD2                    0.686 1.969010 0.372 0.557 0.277
## MIG3                                       0.684 1.056078 0.368 0.570 0.231
## KANK1/2/3/4                                0.684 1.139440 0.368 0.595 0.262
## NVL                                        0.680 1.009940 0.360 0.747 0.554
## KH2013:KH.L153.109                         0.680 1.170249 0.360 0.595 0.246
## C12ORF52                                   0.679 1.332137 0.358 0.557 0.292
## KH2013:KH.S1269.1                          0.672 1.295684 0.344 0.582 0.292
## KH2013:KH.S1140.1_GPM6B                    0.670 1.050835 0.340 0.671 0.415
## TBX1/10                                    0.661 1.134796 0.322 0.633 0.385
## GPR89B                                     0.656 1.013249 0.312 0.519 0.231
## KH2013:KH.S404.8_HSP-16.1/11/2/41/48/49 0.654 1.208633 0.308 0.747 0.523
## MXRA7                                      0.653 1.006081 0.306 0.759 0.600
## COPS2                                      0.646 1.083980 0.292 0.519 0.292
```

```r
write.table(SHPspecific_20.markers, file = "SHPspecific_20.markers.txt", sep = "\t")

# Find FHP specific markers that distinguish two heart progenitors FHP and
# SHP
FHPspecific_20.markers = find.markers(hpf20, "20FHP", "20SHP", thresh.use = 1,
    test.use = "roc", min.pct = 0.5)
head(FHPspecific_20.markers[order(FHPspecific_20.markers$myAUC, decreasing = T),
    ], 20)
```

```
##                                          myAUC avg_diff power pct.1 pct.2
## Ci-R1CiGC27a04                           0.945 2.550143 0.890 0.954 0.443
## KH2013:KH.C5.227_F56C4.4                 0.922 1.583096 0.844 1.000 0.962
## MMP21                                    0.920 3.614448 0.840 0.877 0.203
## COL13A1                                  0.891 2.810263 0.782 0.846 0.190
## FRAS1                                    0.887 2.287808 0.774 0.831 0.190
## KH2013:KH.C4.547_BMP2/4                  0.887 1.342081 0.774 0.985 0.722
## KH2013:KH.C10.203                        0.857 1.758780 0.714 0.923 0.418
## SMYD1                                    0.848 1.757263 0.696 0.923 0.506
## SMTN                                     0.838 1.238689 0.676 0.985 0.848
## BX005256.5                               0.837 1.667243 0.674 0.938 0.519
## C5ORF48                                  0.836 1.655992 0.672 0.892 0.418
## SFRP1/5                                  0.835 1.369541 0.670 0.923 0.405
## NTN1/2/3/5                               0.829 1.383314 0.658 0.892 0.380
## KH2013:KH.C1.1083_EPHA10/2/3/4/5/6/7/8 0.827 2.507890 0.654 0.708 0.089
## LAMA1/2                                  0.825 1.628374 0.650 0.738 0.165
## ISL1/2                                   0.819 1.549760 0.638 0.846 0.342
## FOXP1/2/4                                0.814 2.022667 0.628 0.769 0.203
## KH2013:KH.C1.953_CG32702                 0.807 1.211560 0.614 0.938 0.646
## KH2013:KH.C12.244                        0.801 1.641349 0.602 0.785 0.266
## KH2013:KH.C3.191_WBSCR27                 0.796 1.211239 0.592 0.938 0.620
```

```r
write.table(FHPspecific_20.markers, file = "FHPspecific_20.markers.txt", sep = "\t")

# Find ASM1 specific markers that distinguish two heart progenitors ASM1 and
# ASM2
ASM1specific_20.markers = find.markers(hpf20, "20ASM1", "20ASM2", thresh.use = 1,
    test.use = "roc", min.pct = 0.5)
head(ASM1specific_20.markers[order(ASM1specific_20.markers$myAUC, decreasing = T),
    ], 20)
```

```
##                             myAUC avg_diff power pct.1 pct.2
## GPX7/8                      0.765 1.262169 0.530 0.882  0.64
## SFXN2                       0.739 1.422001 0.478 0.739  0.44
## LATS1/2                     0.737 1.317006 0.474 0.697  0.32
## UNC5A                       0.728 1.247983 0.456 0.815  0.48
## KH2013:KH.C10.340_DHRS11 0.715 1.095224 0.430 0.672  0.28
## FZD5/8                      0.705 1.552283 0.410 0.555  0.20
## KH2013:KH.S2635.1_FNBP1  0.703 1.304549 0.406 0.681  0.48
## KH2013:KH.C2.646            0.699 1.046240 0.398 0.765  0.60
## RASSF10A                    0.697 1.149363 0.394 0.521  0.16
## VASN                        0.684 1.168025 0.368 0.504  0.12
## Ci-tune                     0.678 1.178515 0.356 0.689  0.40
## MRRF                        0.677 1.570648 0.354 0.580  0.32
## CD2BP2                      0.674 1.258361 0.348 0.706  0.60
## KH2013:KH.C1.29_EIF4EBP1 0.673 1.261134 0.346 0.639  0.40
## RGS17/18/19/20              0.655 1.096347 0.310 0.571  0.32
## ARRDC2/3                    0.654 1.148108 0.308 0.538  0.28
## HAND/1/2                    0.645 1.345731 0.290 0.521  0.36
## KH2013:KH.L124.2            0.645 2.411661 0.290 0.966  1.00
## CHCHD7                      0.645 1.266356 0.290 0.563  0.32
## KH2013:KH.C2.209_CALM1/3 0.636 1.193868 0.272 0.538  0.32
```

```r
write.table(ASM1specific_20.markers, file = "ASM1specific_20.markers.txt", sep = "\t")

# Find ASM2 specific markers that distinguish two heart progenitors ASM2 and
# ASM1
ASM2specific_20.markers = find.markers(hpf20, "20ASM2", "20ASM1", thresh.use = 1,
    test.use = "roc", min.pct = 0.5)
head(ASM2specific_20.markers[order(ASM2specific_20.markers$myAUC, decreasing = T),
    ], 20)
```

```
##                             myAUC avg_diff power pct.1 pct.2
## KH2013:KH.C8.649_ACTA1/2 0.886 1.832513 0.772  1.00 0.706
## MYF5                        0.873 1.562098 0.746  1.00 0.697
```

```
## SMYD1                      0.873 1.325876 0.746  1.00 0.933
## ZKSCAN1/14/17/2/3/6        0.869 2.751014 0.738  0.80 0.151
## FUN26                      0.864 1.904252 0.728  0.88 0.235
## Ci-Actin;                  0.861 1.712992 0.722  0.96 0.782
## MYO-1/2/3/5                0.845 1.814388 0.690  0.92 0.563
## DYRK2/3                    0.842 2.433072 0.684  0.84 0.252
## KH2013:KH.C12.669_PHEX     0.839 1.439534 0.678  0.96 0.597
## KH2013:KH.C7.476_CHRNB1    0.830 1.470585 0.660  0.92 0.454
## KH2013:KH.C5.404_MBL1/2    0.826 1.706676 0.652  0.84 0.235
## CALML6                     0.820 1.671514 0.640  0.96 0.605
## KH2013:KH.S489.1_ARF1/3    0.819 1.012200 0.638  1.00 0.765
## CCDC66                     0.818 1.216917 0.636  0.96 0.647
## RBM24/38                   0.817 1.232162 0.634  0.96 0.748
## KH2013:KH.S643.6_CDKN1B    0.809 1.113191 0.618  1.00 0.815
## MYBL1/2                    0.803 1.214419 0.606  0.92 0.437
## KH2013:KH.C1.509           0.799 1.285231 0.598  0.88 0.445
## LGALS4/8/9                 0.797 1.557405 0.594  0.80 0.445
## NLRC3                      0.790 1.435796 0.580  0.60 0.160
```

```
write.table(ASM2specific_20.markers, file = "ASM2specific_20.markers.txt", sep = "\t")

# Find ASM markers
ASM_20.markers = find.markers(hpf20, c("20ASM1", "20ASM2"), "20FHP", thresh.use = 1,
    test.use = "roc", min.pct = 0.5)
head(ASM_20.markers[order(ASM_20.markers$myAUC, decreasing = T), ], 20)
```

```
##                            myAUC avg_diff power pct.1 pct.2
## EBF1/2/3/4                 0.999 4.550177 0.998 1.000 0.154
## KH2013:KH.C4.506_HMCN1     0.997 3.159464 0.994 1.000 0.785
## KH2013:KH.C11.139          0.994 4.919472 0.988 0.993 0.231
## KH2013:KH.L25.4            0.994 4.655595 0.988 0.993 0.062
## DCDC2B                     0.989 4.127543 0.978 0.986 0.154
## KH2013:KH.C5.62_ODF3L2     0.988 4.192618 0.976 0.986 0.108
## KH2013:KH.S1269.1          0.983 3.046282 0.966 0.993 0.292
## BDH1                       0.981 3.236993 0.962 0.986 0.215
## NOVA1/2                    0.980 2.940057 0.960 1.000 0.215
## SCP1                       0.978 4.539122 0.956 0.958 0.046
## KH2013:KH.C8.892_TUB1/3    0.967 2.799581 0.934 0.993 0.508
## KH2013:KH.C5.522_SLC6A13 0.962 3.947985 0.924 0.931 0.092
## UBE2QL1                    0.957 2.591103 0.914 0.972 0.369
## KH2013:KH.L18.10_THSD7A    0.956 2.568935 0.912 0.972 0.231
## ELK1/3/4                   0.949 3.539293 0.898 0.917 0.108
## PCBP3                      0.948 2.392518 0.896 0.965 0.169
## SLC35G2/6                  0.942 4.200869 0.884 0.903 0.108
```

```
## KH2013:KH.L154.26_CSMD2  0.942 3.077734 0.884 0.938 0.277
## TIMP1/2/3/4              0.941 4.461382 0.882 0.903 0.123
## MMP14/15/16/24           0.940 2.208247 0.880 0.979 0.308
```

```r
# ASM_20.markers=find.markers(hpf20,c('20ASM1','20ASM2'),thresh.use =
# 1,test.use = 'roc',min.pct = 0.5)
# head(ASM_20.markers[order(ASM_20.markers$myAUC,decreasing = T),],20)
write.table(ASM_20.markers, file = "ASM_20.markers.txt", sep = "\t")

# Visualize markers of different clusters using violin plot and feature plot
hpf20@ident = factor(hpf20@ident, ordered = T, levels = c("20FHP", "20SHP",
    "20ASM1", "20ASM2"))
genes.viz.20 = c("DACH1/2", "MYF5", "EBF1/2/3/4", "MMP21")
feature.plot(hpf20, genes.viz.20, pt.size = 0.8)
```

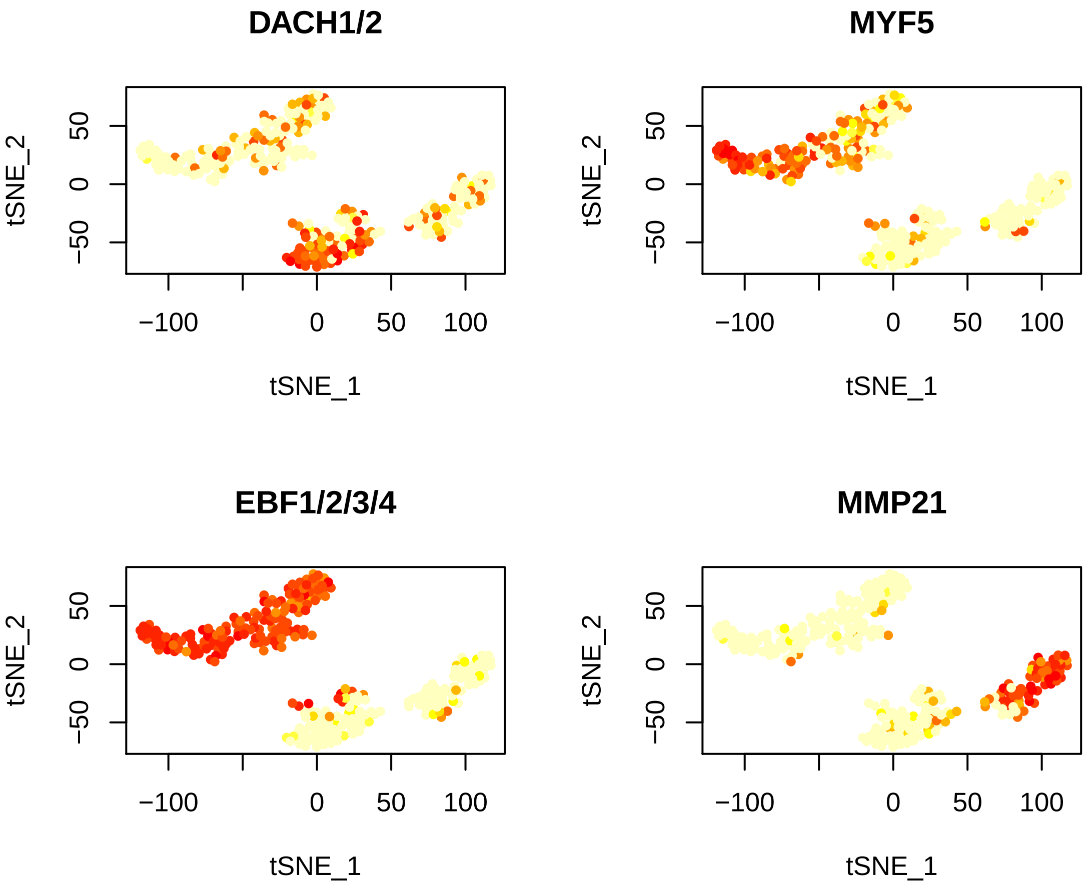


```r
vlnPlot(hpf20, genes.viz.20, cols.use = c("red", "orange", "blue", "blue4"))
```

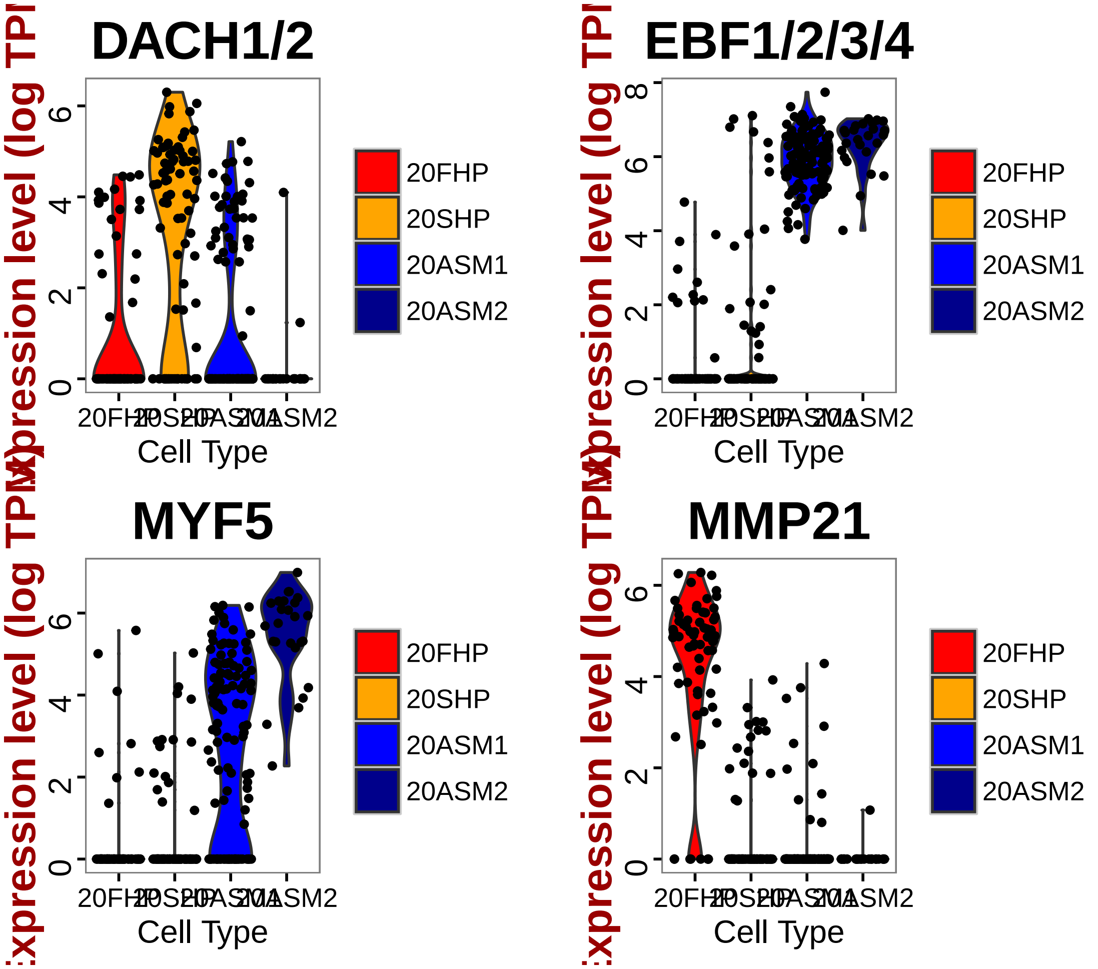


```
# Select markers for plotting on a Heatmap (top 10 positive markers with
# high classification power(myAUC))
marker20FHP = rownames(FHP_20.markers[order(FHP_20.markers$myAUC, decreasing = T)[1:15],
    ])
marker20SHP = rownames(SHP_20.markers[order(SHP_20.markers$myAUC, decreasing = T)[1:15],
    ])
marker20ASM1 = rownames(ASM1_20.markers[order(ASM1_20.markers$myAUC, decreasing = T)[1:1
    ])
marker20ASM2 = rownames(ASM2_20.markers[order(ASM2_20.markers$myAUC, decreasing = T)[1:1
    ])
marker.20 = c(marker20FHP, marker20SHP, marker20ASM1, marker20ASM2)

panHeart.select = rownames(subset(panHP_20.markers, power > 0.6 & avg_diff >
    0))
asm.select = rownames(subset(ASM_20.markers, power > 0.6 & avg_diff > 0))

# Draw a heatmap of all cells for these marker genes
doHeatMap(hpf20, genes.use = marker.20, remove.key = F, slim.col.label = T,
    draw.line = T, cex.col = 1.2, col.use = col)
```

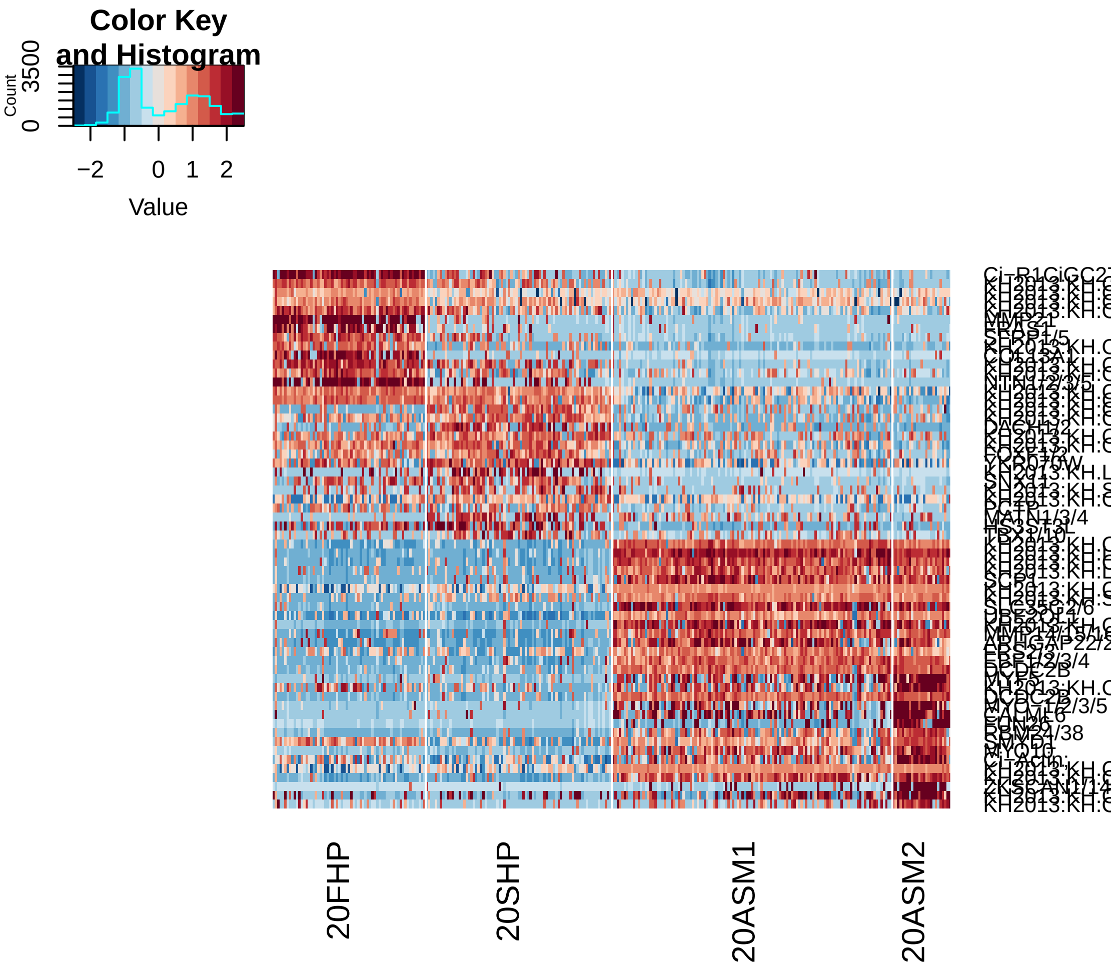


```r
doHeatMap(hpf20, genes.use = c(panHeart.select, asm.select), remove.key = F,
    slim.col.label = T, draw.line = T, cex.col = 1.2, col.use = col)
```

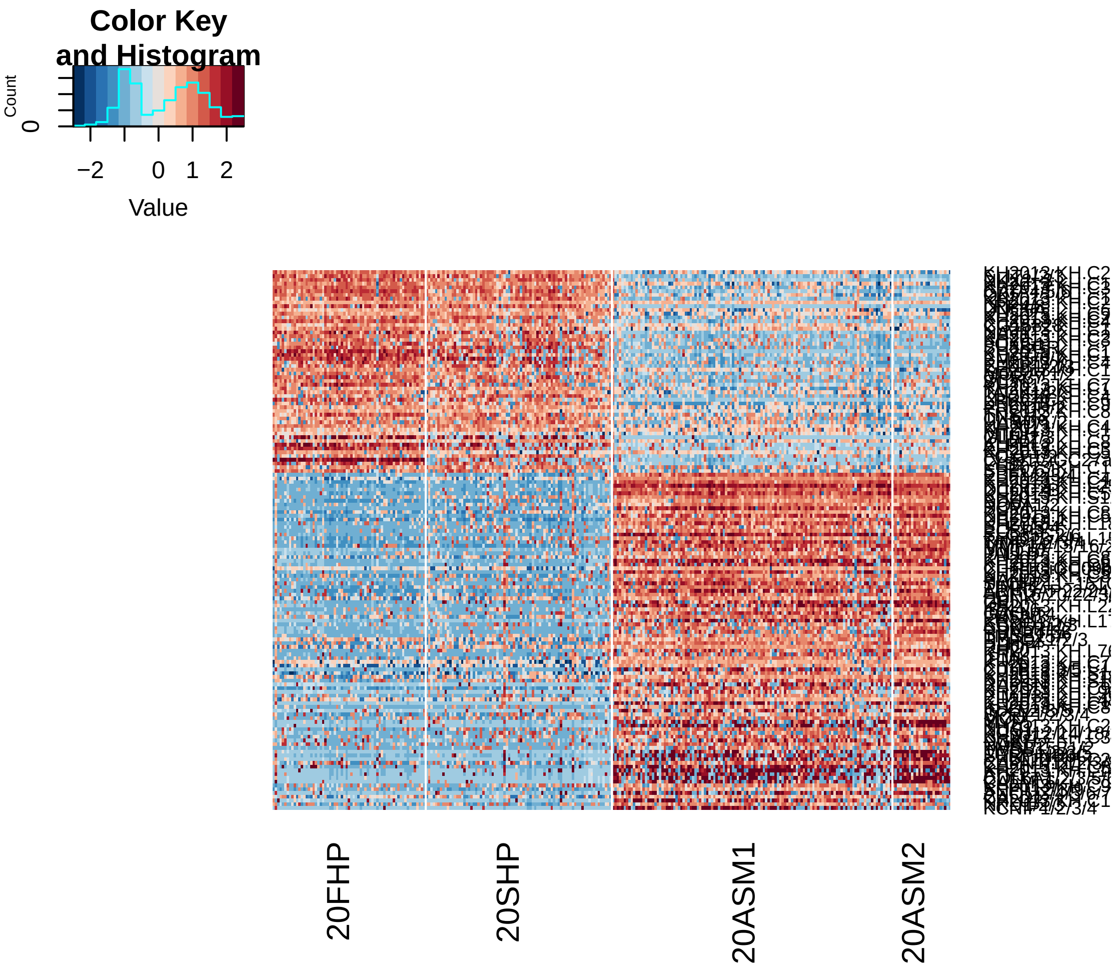


5. hpf18

```
# Subset data from preprocessed Seurat object
hpf18 = subsetData(hpfall.remv2, which.cells(hpfall.remv2, "hpf18"), do.center = F,
    do.scale = F)
hpf18
```

```
## An object of class seurat in project allhpf
##  14864 genes across 177 samples.
```

```
# Based on hpf12 hpf14 and hpf20 data, we have successfully uncovered all
# the TVC lineage types: TVC, STVC, FHP, SHP, ASM Based on preliminary
# studies, hpf18 contains three TVC lineage cell types: FHP, SHP and ASM
# Therefore we can run PCA using 20hpf cell markers (20FHP, 20SHP, 20ASM)
# with power > 0.5 and postive expressions
marker20FHP.use = rownames(subset(FHP_20.markers, power > 0.5 & avg_diff > 0))
marker20SHP.use = rownames(subset(SHP_20.markers, power > 0.5 & avg_diff > 0))
marker20ASM1.use = rownames(subset(ASM1_20.markers, power > 0.5 & avg_diff >
```

```r
    0))
marker20ASM2.use = rownames(subset(ASM2_20.markers, power > 0.5 & avg_diff >
    0))
marker.20.use = unique(c(marker20FHP.use, marker20SHP.use, marker20ASM1.use,
    marker20ASM2.use))
length(marker.20.use)
```

```
## [1] 200
```

```r
hpf18 = pca(hpf18, pc.genes = marker.20.use, do.print = F)
pcScree(hpf18, marker.20.use, 10)
```

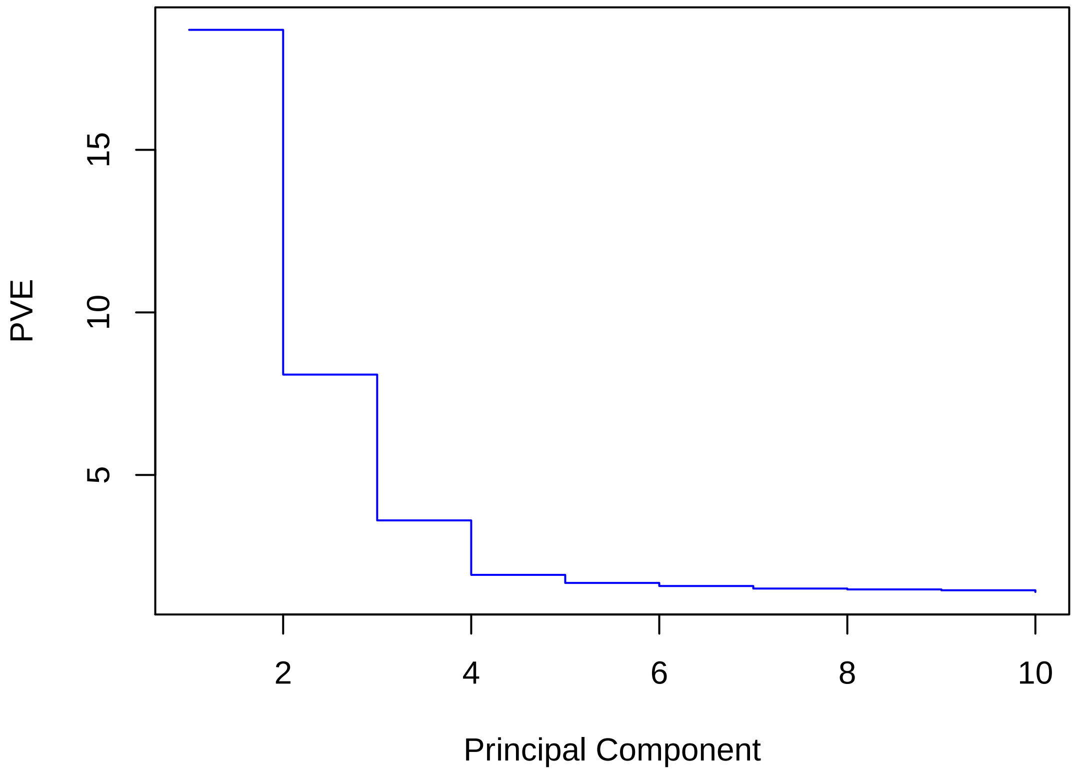


```r
pcHeatmap(hpf18, 1, do.balanced = T, col.use = col)
```

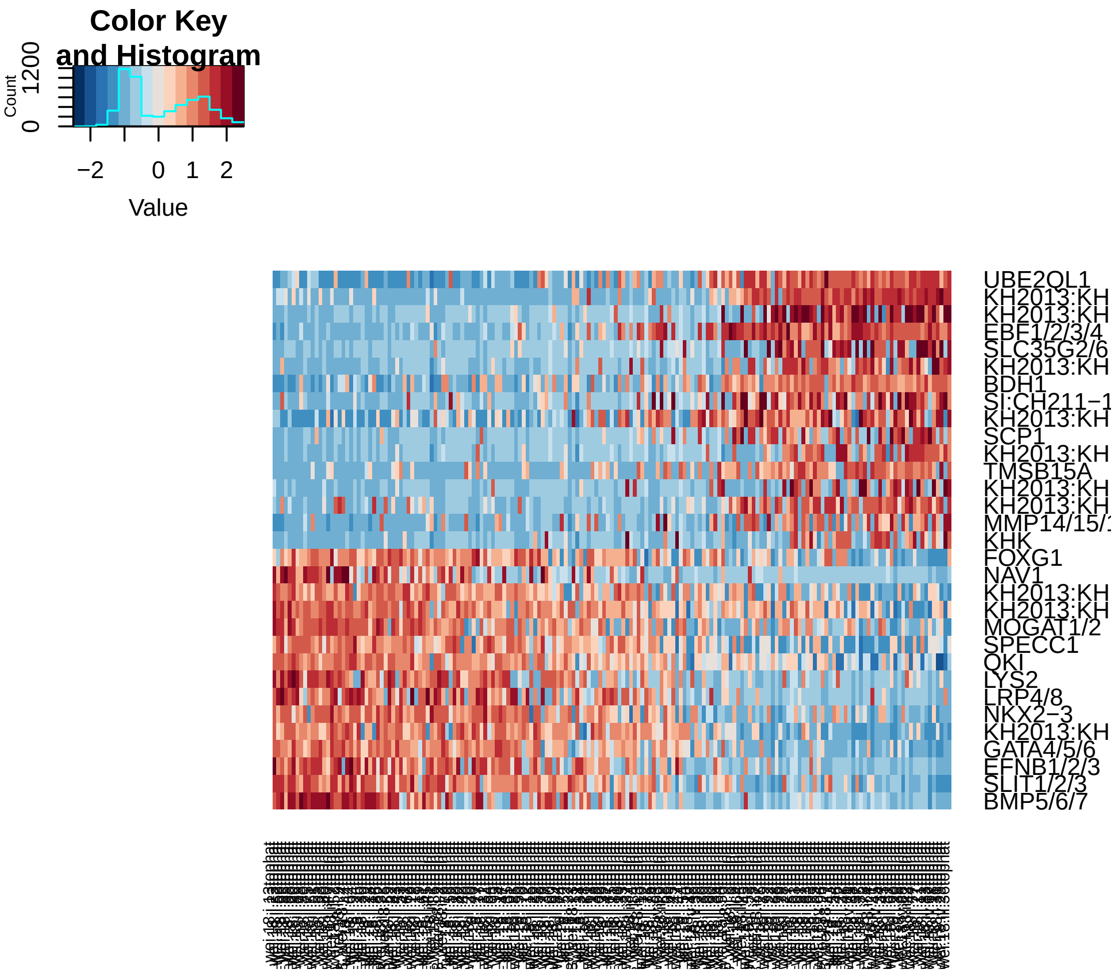


```
pcHeatmap(hpf18, 2, do.balanced = T, col.use = col)
```

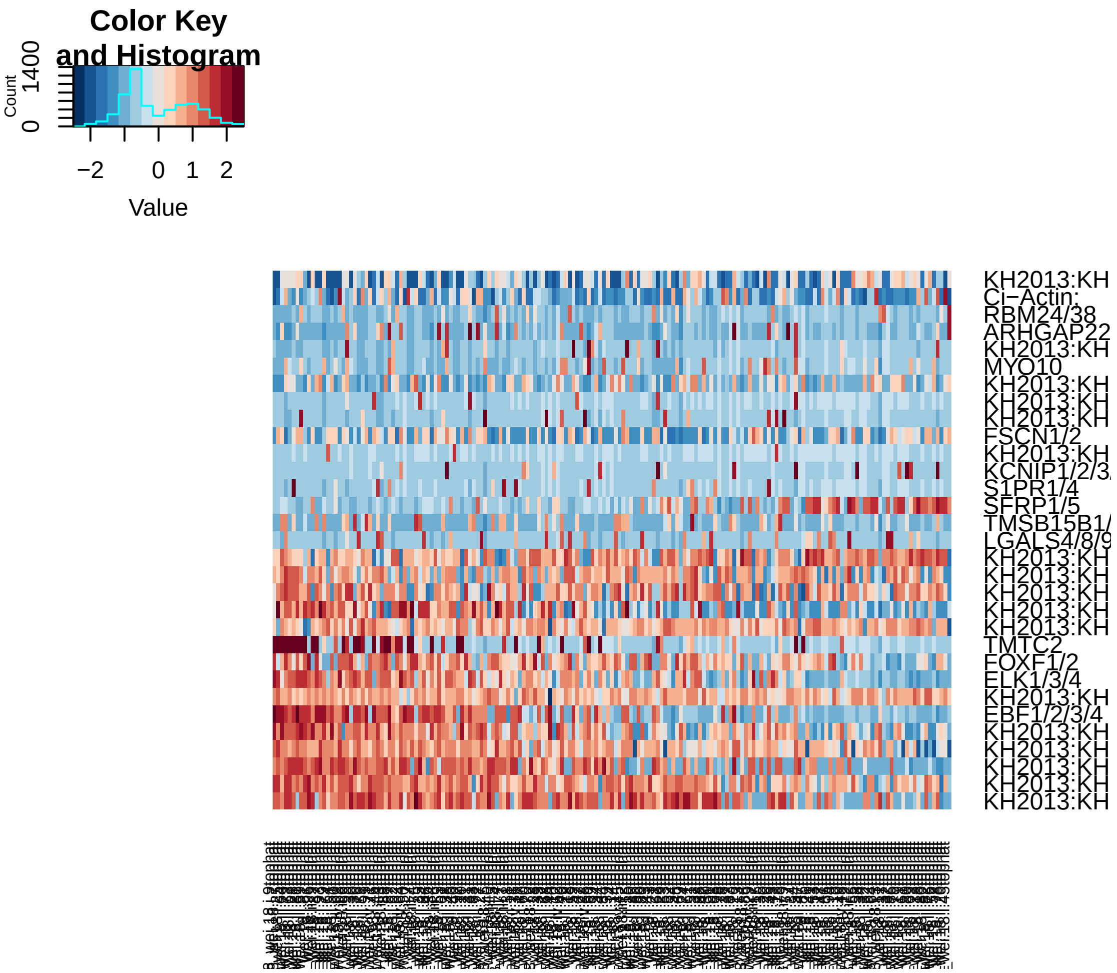


```r
pcHeatmap(hpf18, 3, do.balanced = T, col.use = col)
```

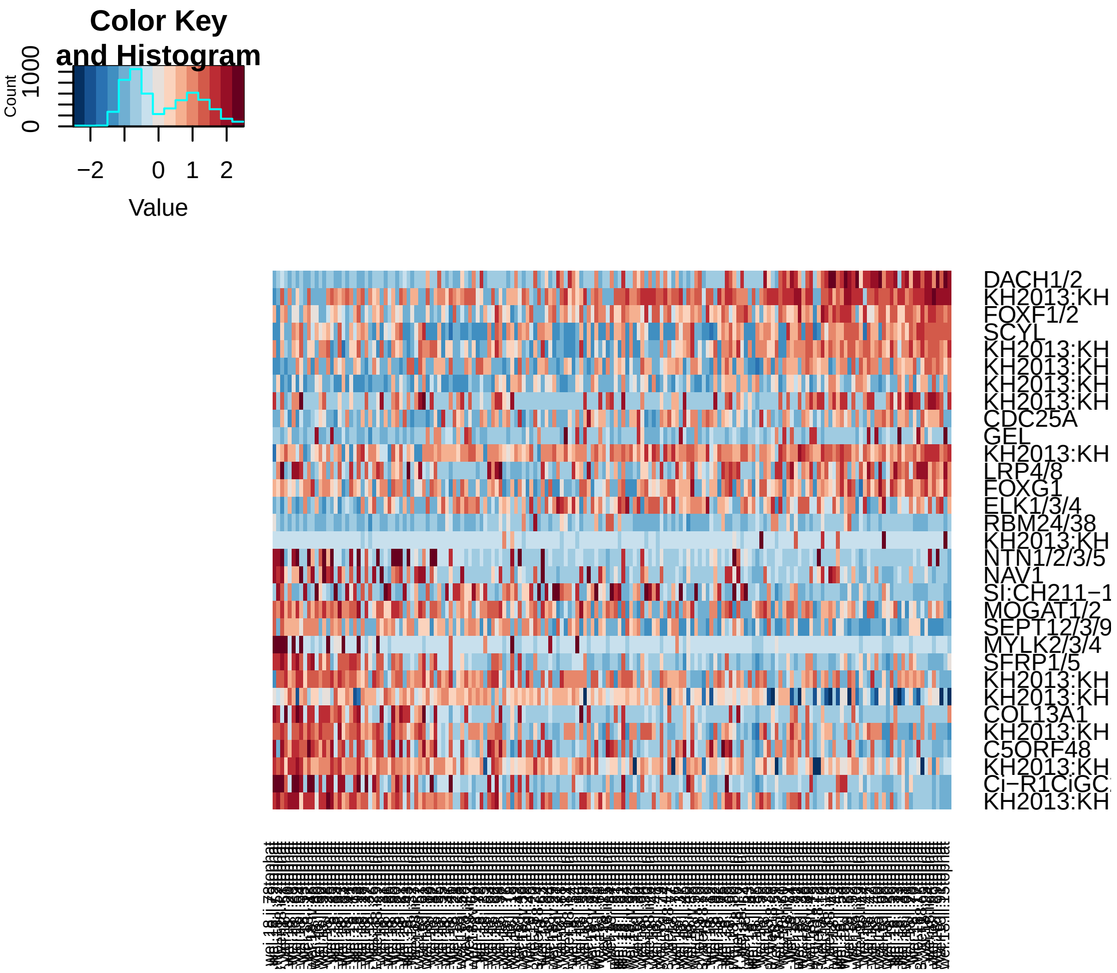


```r
pca.plot(hpf18, 1, 2)
```

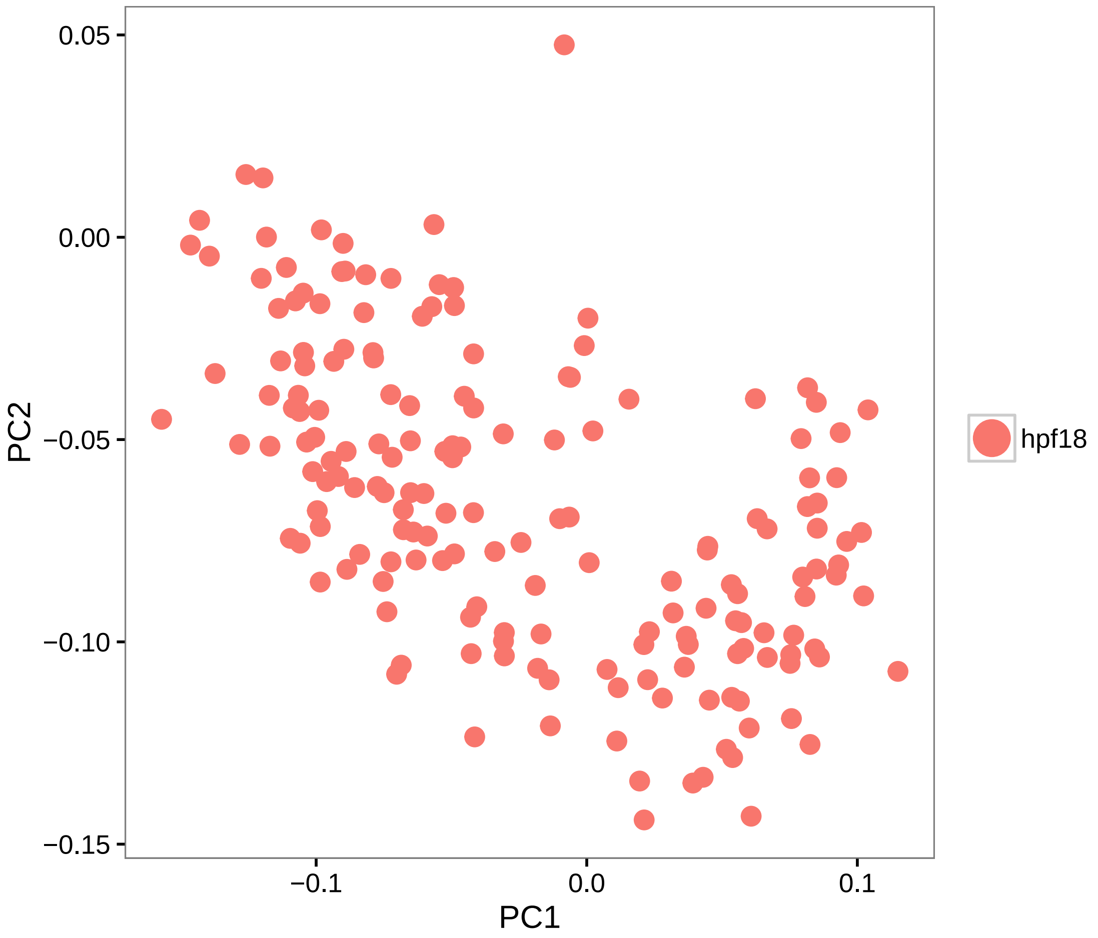


```
pca.plot(hpf18, 1, 3)
```

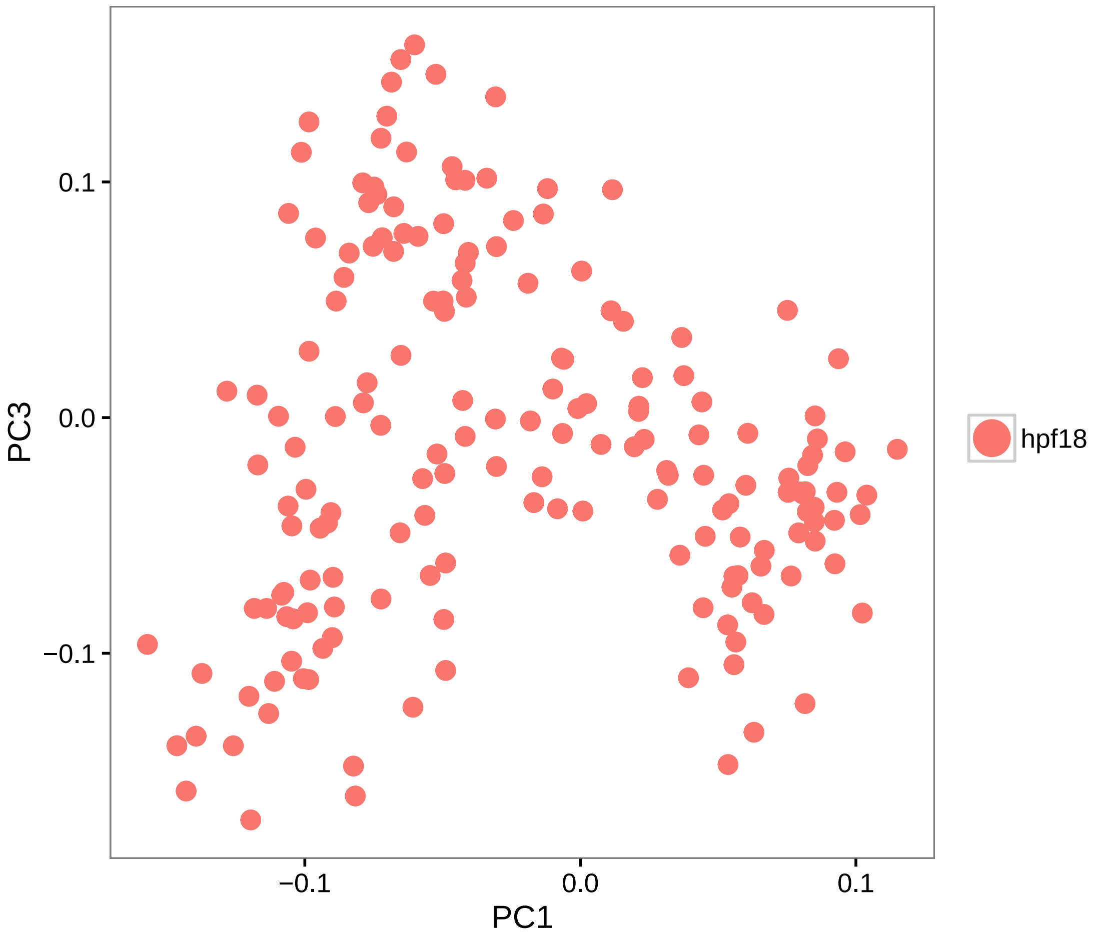


```
# Calculate PCA scores for all genes (PCA projection)
hpf18 = project.pca(hpf18, do.print = F)

# Visualize the full projected PCA, which now includes new genes which were
# not previously (use.full=TRUE)
pcHeatmap(hpf18, 1, use.full = T, do.balanced = T, col.use = col)
```

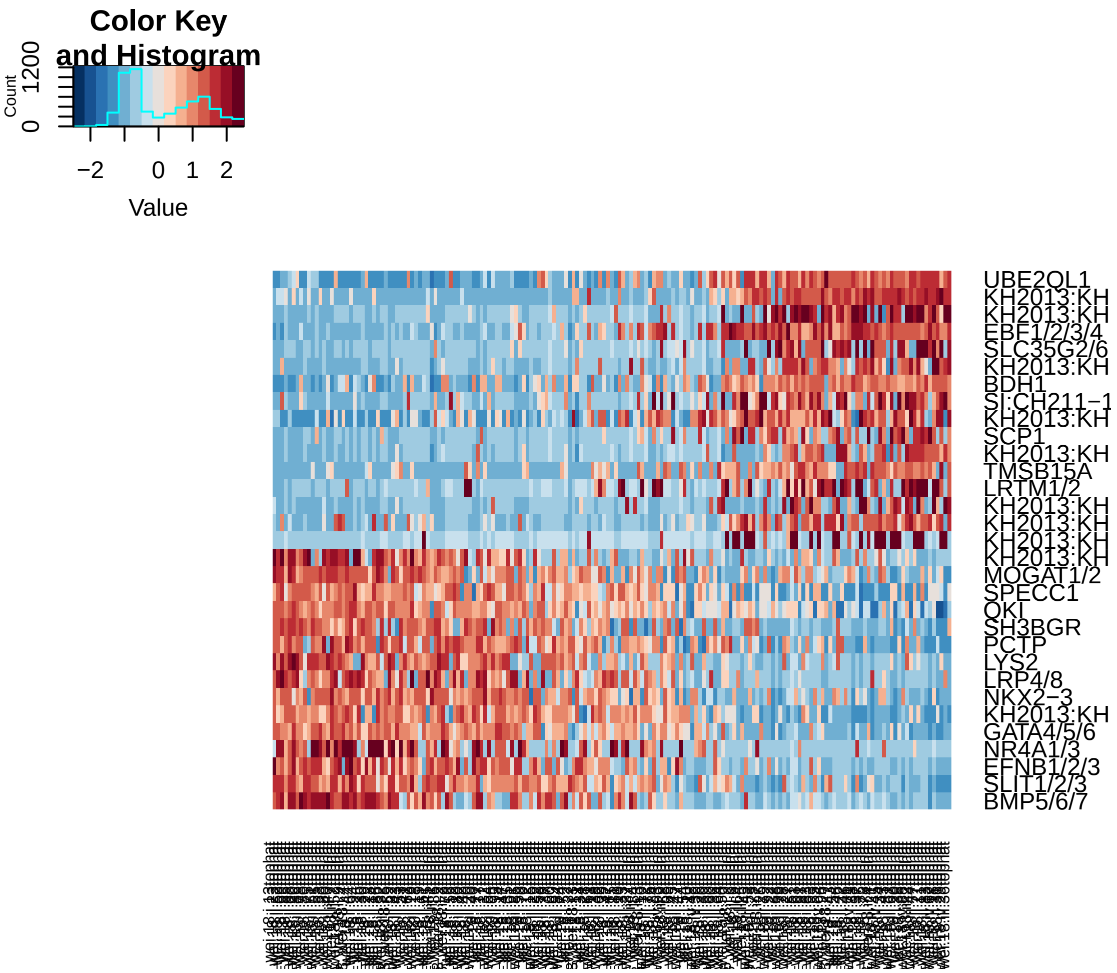


```
pcHeatmap(hpf18, 2, use.full = T, do.balanced = T, col.use = col)  #technical
```

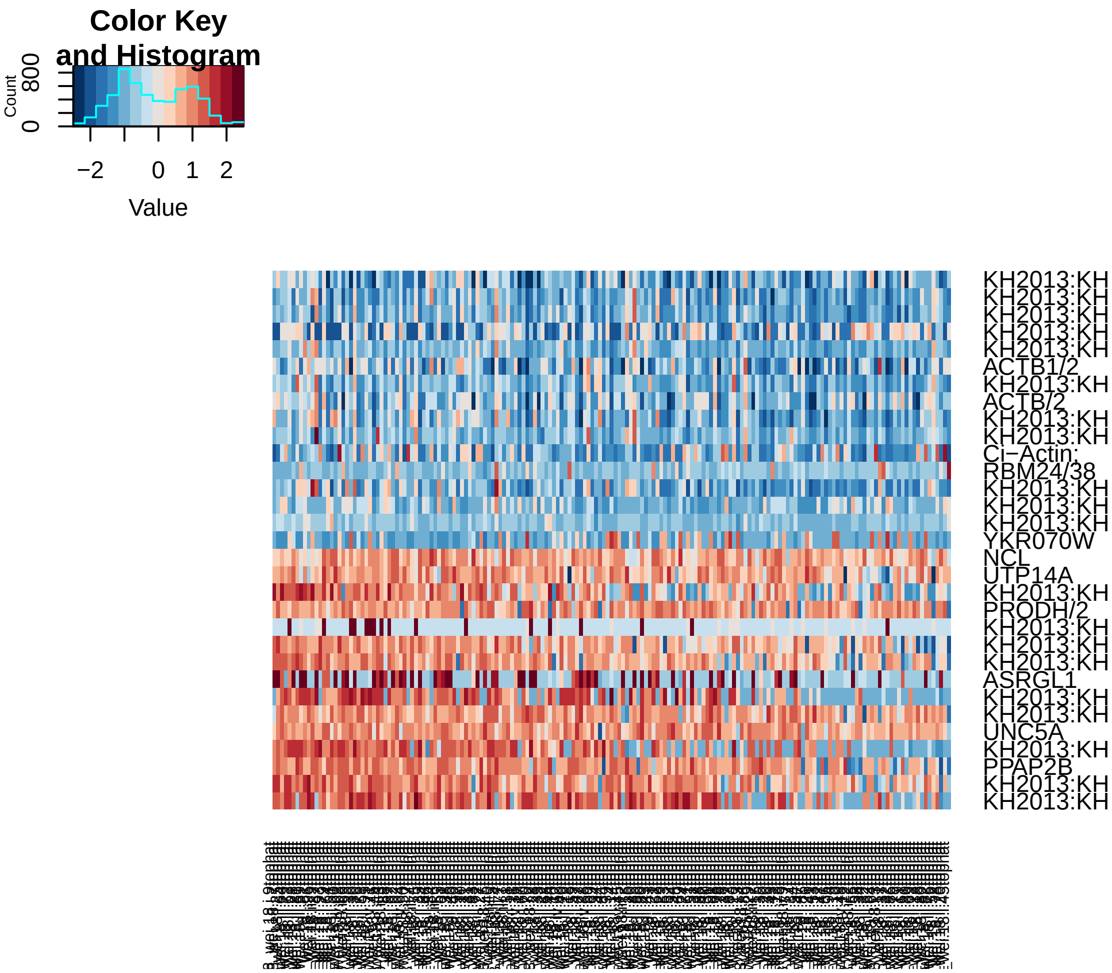


```r
pcHeatmap(hpf18, 3, use.full = T, do.balanced = T, col.use = col)
```

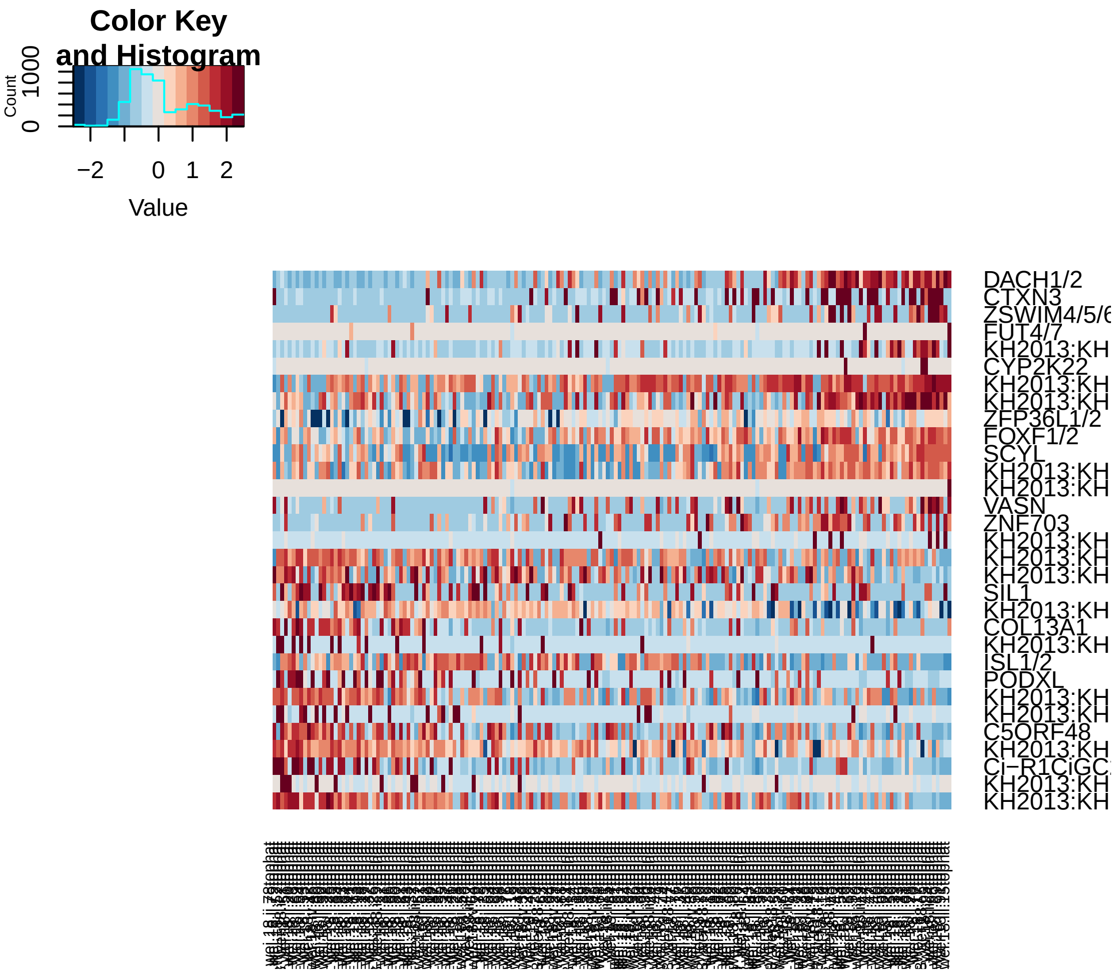


```r
# Do 200 random samplings to find significant genes, each time randomly
# permute 1% of genes This returns a 'p-value' for each gene in each PC,
# based on how likely the gene/PC score woud have been observed by chance
hpf18 = jackStraw(hpf18, num.replicate = 200, do.print = F)

# The jackStraw plot compares the distribution of P-values for each PC with
# a uniform distribution (dashed line) 'Significant' PCs will have a strong
# enrichment of genes with low p-values (solid curve above dashed line)
jackStrawPlot.new(hpf18, PCs = 1:12)
```

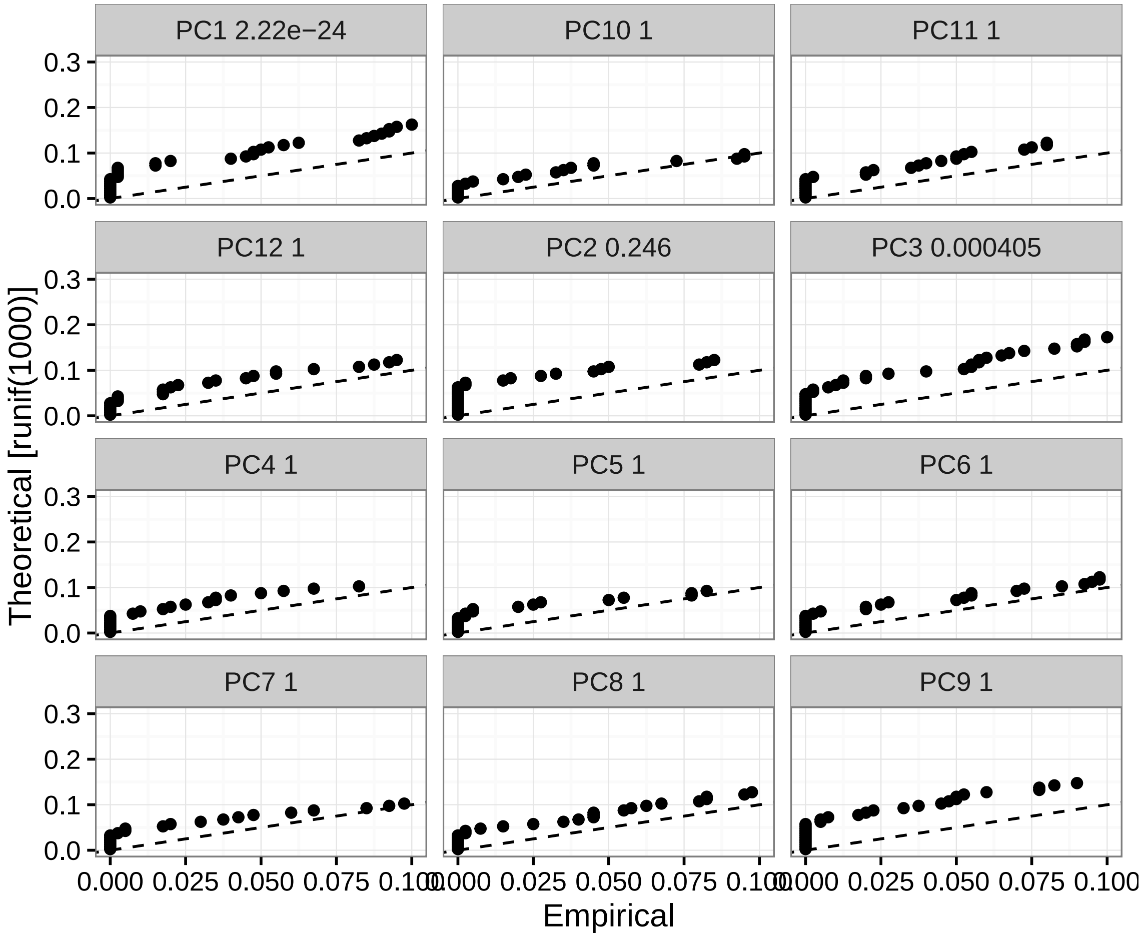


```r
# In this case only PC1 and PC3 are strongly significant.

# Run tSNE using significant PCs as input (spectral tSNE), we get distinct
# point clouds
hpf18 = run_tsne(hpf18, max_iter = 2000, dims.use = c(1, 3))
tsne.plot(hpf18, do.label = T, label.pt.size = 1)
```

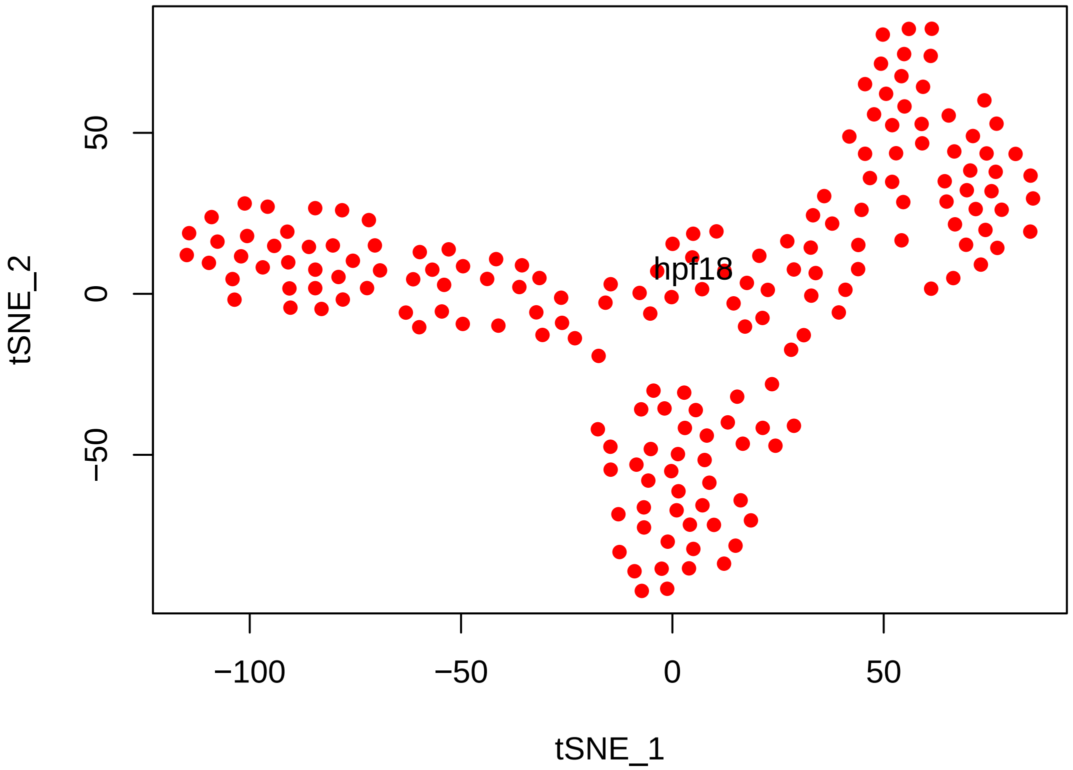


```r
# Find cell clusters using Modularity optimization cluster detection.
hpf18 = FindClusters(hpf18, pc.use = c(1, 3), do.modularity = T, resolution = 1,
    prune.SNN = 0.1, print.output = 0, k.param = 20, k.scale = floor(177/20))
```

```
## [1] "SNN : processed 44 cells"
## [1] "SNN : processed 88 cells"
## [1] "SNN : processed 133 cells"
## [1] "SNN : processed 177 cells"
```

```r
tsne.plot(hpf18, do.label = T, label.pt.size = 1)
```

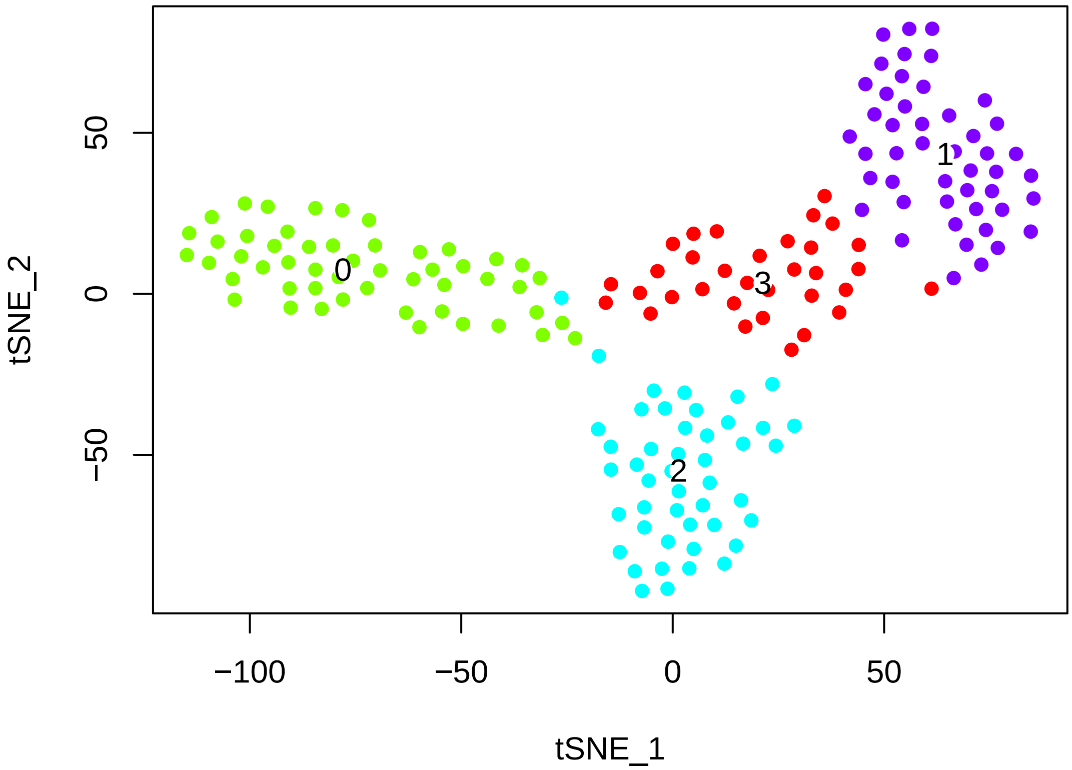


```
# The validity of the clusters can be validated using a classification
# scheme based on linear SVMs.
hpf18 = BuildSNN(hpf18, pc.use = c(1, 3), do.sparse = T, k.param = 20, k.scale = floor(1
```

```
## [1] "SNN : processed 44 cells"
## [1] "SNN : processed 88 cells"
## [1] "SNN : processed 133 cells"
## [1] "SNN : processed 177 cells"
```

```
hpf18 = ValidateClusters(hpf18, pc.use = c(1, 3), min.connectivity = 0.001,
    acc.cutoff = 0.85)
```

```
## [1] "100% complete --- started with 4 clusters, 4 clusters remaining"
```

```
tsne.plot(hpf18, do.label = T, label.pt.size = 1)

# Find cluster markersusing ROC test with thresh.use = 1, min.pct = 0.5 The
# ROC test returns the 'classification power' for any individual marker
# (ranging from 0 - random, to 1 - perfect). Though not a statistical test,
# it is often very useful for finding clean markers.  Find markers for
# cluster 0
cl0_18.markers = find.markers(hpf18, 0, thresh.use = 1, test.use = "roc", min.pct = 0.5)
head(cl0_18.markers[order(cl0_18.markers$myAUC, decreasing = T), ], 20)
```

```
##                              myAUC avg_diff power pct.1 pct.2
## NAV1                         0.906 2.416886 0.812 0.882 0.151
## KH2013:KH.C3.665_ZAN         0.905 1.528765 0.810 0.980 0.825
## MOGAT1/2                     0.899 1.491851 0.798 1.000 0.579
## BMP5/6/7                     0.890 1.943625 0.780 0.941 0.270
## KH2013:KH.C4.547_BMP2/4      0.881 1.778261 0.762 0.961 0.437
## KH2013:KH.C5.227_F56C4.4     0.862 1.196972 0.724 1.000 0.960
## KH2013:KH.C1.953_CG32702     0.847 1.778144 0.694 0.902 0.508
## LAMA1/2                      0.835 1.664197 0.670 0.588 0.063
## SFRP1/5                      0.833 3.061270 0.666 0.745 0.151
## COL13A1                      0.828 2.536370 0.656 0.725 0.119
## Ci-R1CiGC27a04               0.821 3.809702 0.642 0.686 0.095
## FZD4                         0.821 1.729355 0.642 0.784 0.159
## LYS2                         0.814 1.294364 0.628 0.863 0.405
## EFNB1/2/3                    0.805 1.119200 0.610 0.922 0.397
## KH2013:KH.C7.695_CYP24A1     0.798 1.849781 0.596 0.863 0.429
## KH2013:KH.C10.203            0.786 1.130693 0.572 0.902 0.651
## NTN1/2/3/5                   0.782 2.734596 0.564 0.588 0.048
## RAC1/2                       0.781 1.024299 0.562 0.941 0.611
## C5ORF48                      0.778 1.347507 0.556 0.784 0.341
## SPECC1                       0.769 1.041143 0.538 0.980 0.698
```

```r
# Visualize known markers with a violin plot
vlnPlot(hpf18, c("NKX2-3", "EBF1/2/3/4", "TBX1/10", "DACH1/2"))
```

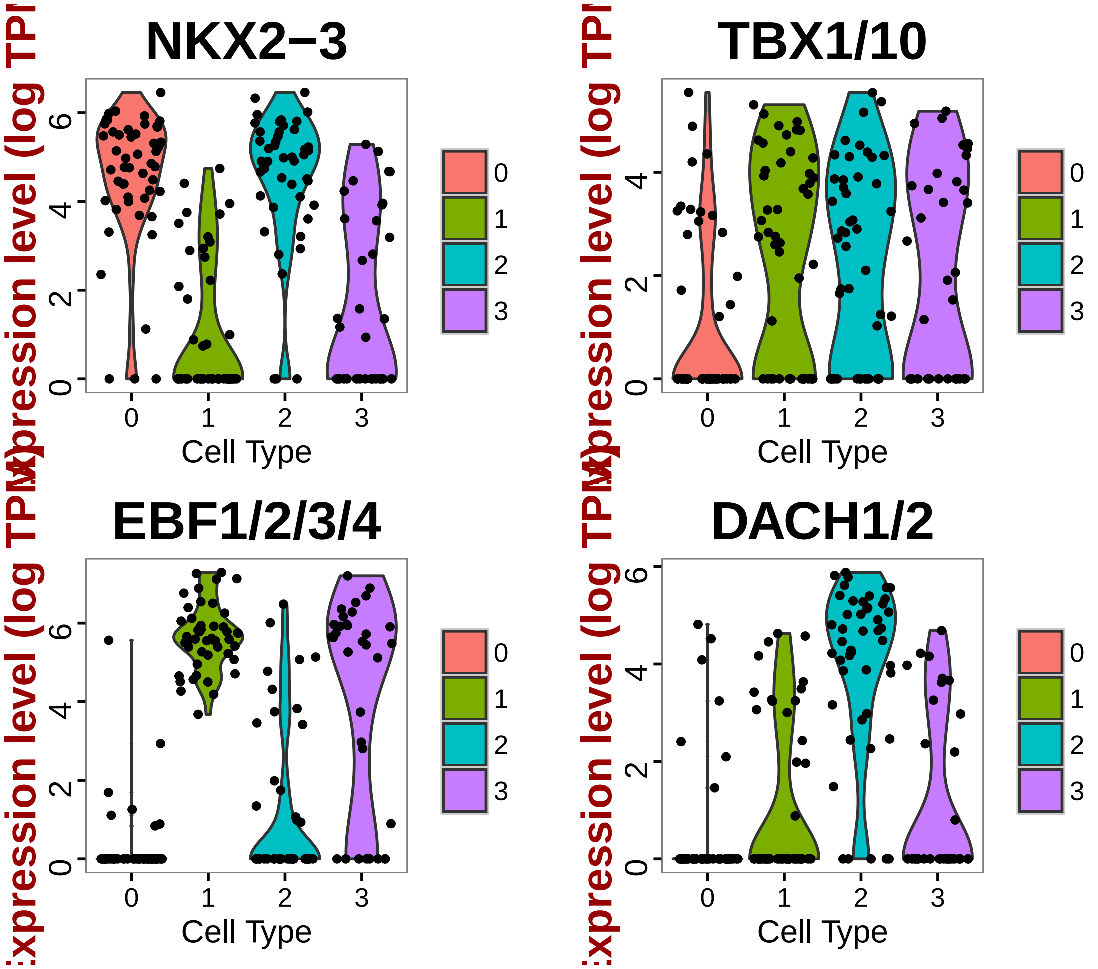


```
# Based on EBF1/2/3/4- and NKX+, DACH1/2-, these are FHP cells

# Visualize new markers with a violin plot
vlnPlot(hpf18, c("NAV1", "BMP5/6/7"))
```

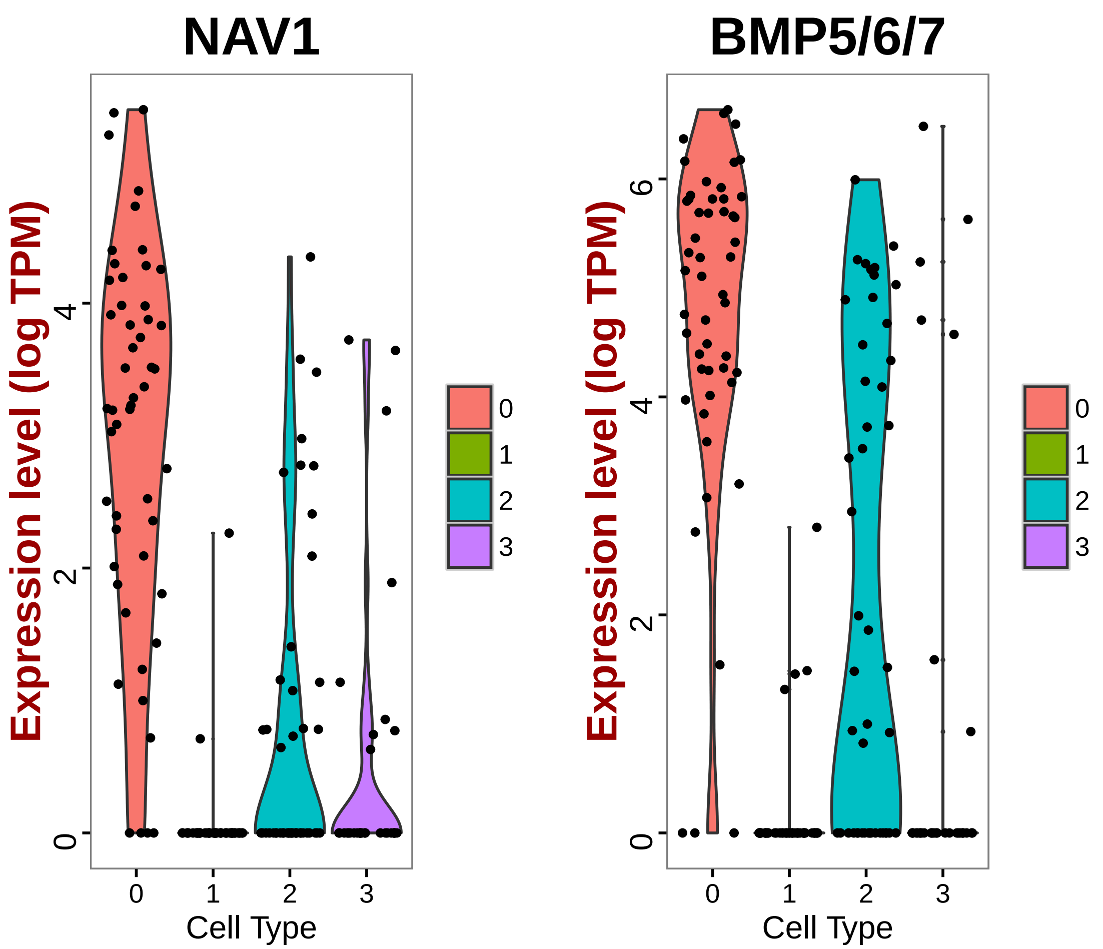


```r
# Find markers for cluster 1
cl1_18.markers = find.markers(hpf18, 1, thresh.use = 1, test.use = "roc", min.pct = 0.5)
head(cl1_18.markers[order(cl1_18.markers$myAUC, decreasing = T), ], 20)
```

```
##                                myAUC avg_diff power pct.1 pct.2
## KH2013:KH.C11.139              0.980 3.608234 0.960 0.979 0.254
## UBE2QL1                        0.951 2.206912 0.902 1.000 0.354
## BDH1                           0.931 1.935226 0.862 0.979 0.446
## KH2013:KH.C4.506_HMCN1         0.928 1.584676 0.856 1.000 0.808
## KH2013:KH.C5.127_FMO1          0.909 2.870722 0.818 0.872 0.108
## KH2013:KH.C5.62_ODF3L2         0.904 2.904233 0.808 0.872 0.108
## KH2013:KH.L18.10_THSD7A        0.893 1.859869 0.786 0.872 0.231
## KH2013:KH.C10.209              0.882 1.205272 0.764 1.000 0.777
## EBF1/2/3/4                     0.875 1.459145 0.750 1.000 0.331
## Ci-R1CiGC09b24;                0.869 1.385696 0.738 1.000 0.785
## KH2013:KH.S1269.1              0.866 1.686455 0.732 0.915 0.331
## KH2013:KH.C8.782_LPHN1/2/3     0.859 1.144203 0.718 1.000 0.692
## SI:CH211-15101.2               0.856 1.477531 0.712 0.894 0.254
## KH2013:KH.C7.805_ID1/2/3/4     0.855 1.197553 0.710 0.979 0.815
```

```
## SLC35G2/6                  0.851 2.414261 0.702 0.766 0.100
## DCDC2B                     0.847 1.681428 0.694 0.872 0.300
## KH2013:KH.L25.4            0.845 3.078121 0.690 0.723 0.062
## KH2013:KH.S511.3           0.839 1.126590 0.678 0.979 0.523
## TMSB15A                    0.838 1.960725 0.676 0.872 0.346
## KH2013:KH.L154.26_CSMD2    0.834 1.168114 0.668 0.957 0.577
```

```
# Visualize known markers with a violin plot
vlnPlot(hpf18, c("NKX2-3", "EBF1/2/3/4", "TBX1/10", "DACH1/2"))
```

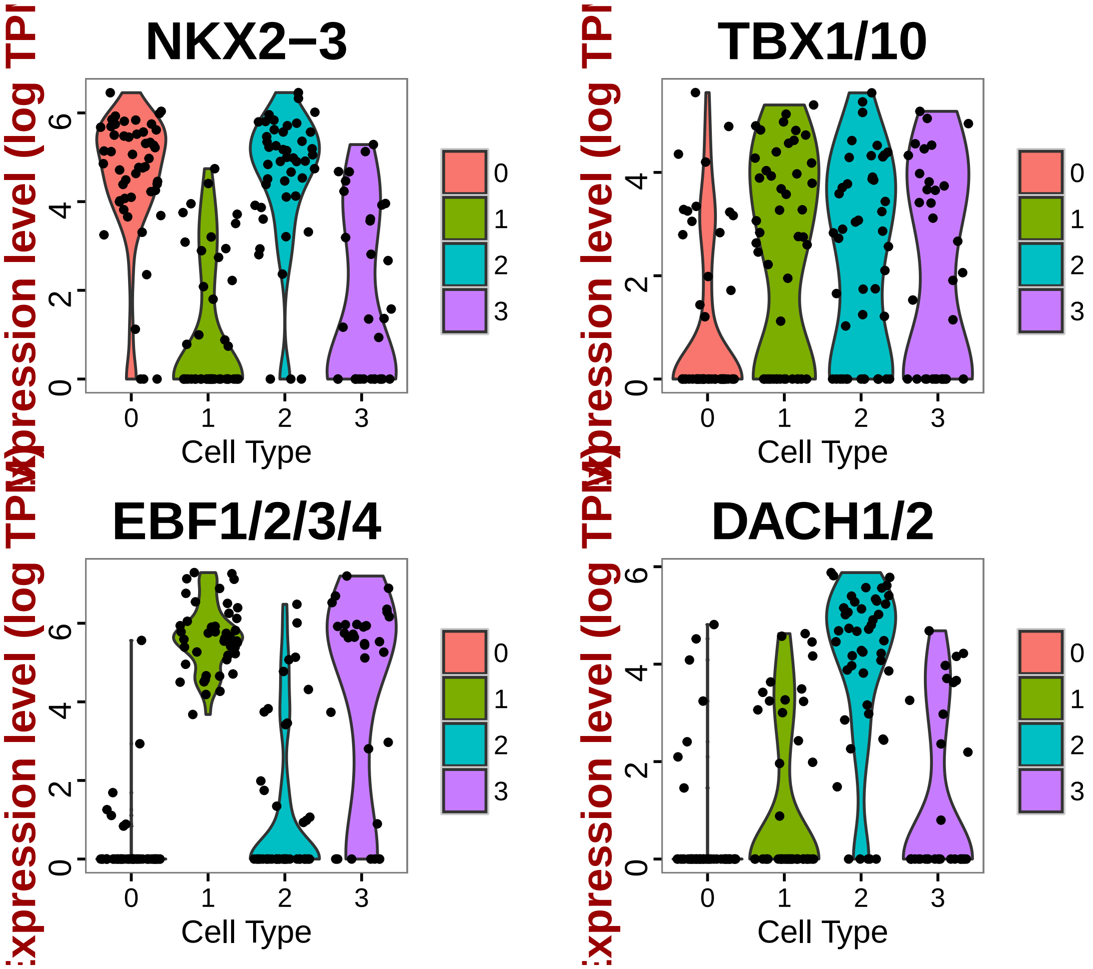


```
# Based on previous discoveries NKX- EBF+ TBX+ DACH- cells are ASM

# Visualize new markers with a violin plot
vlnPlot(hpf18, c("BDH1", "UBE2QL1"))
```

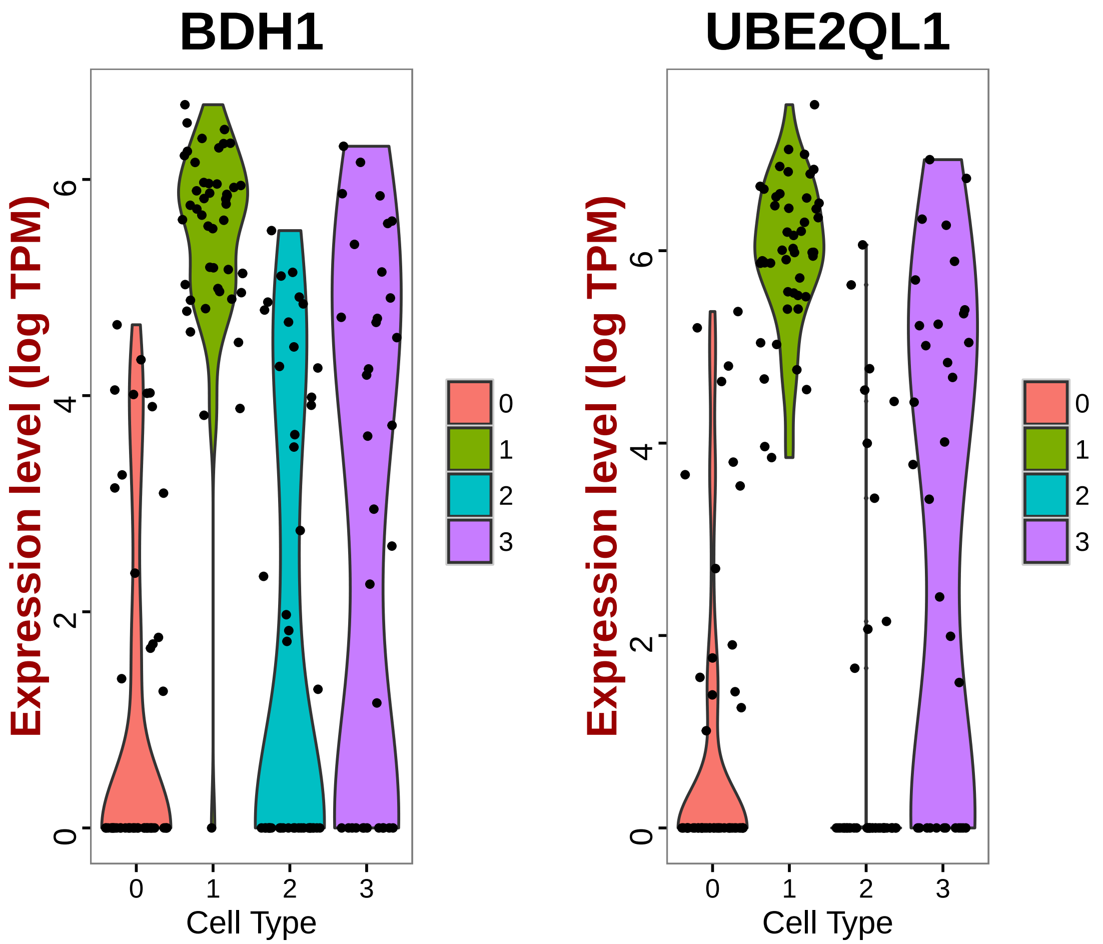


```
# Find markers for cluster 2
cl2_18.markers = find.markers(hpf18, 2, thresh.use = 1, test.use = "roc", min.pct = 0.5)
head(cl2_18.markers[order(cl2_18.markers$myAUC, decreasing = T), ], 20)
```

```
##                                   myAUC  avg_diff power pct.1 pct.2
## DACH1/2                           0.885  2.322664 0.770 0.891 0.252
## KH2013:KH.C1.638_C17ORF105        0.841  1.244018 0.682 0.935 0.740
## KH2013:KH.C10.174                 0.832  1.082488 0.664 1.000 0.580
## SCYL                              0.797  1.327073 0.594 0.891 0.527
## KH2013:KH.C10.172                 0.771  1.475241 0.542 0.848 0.557
## FOXF1/2                           0.738  1.061960 0.476 0.913 0.763
## SUDS3                             0.732  1.072577 0.464 0.913 0.718
## LENG8                             0.700  1.019048 0.400 0.630 0.389
## KH2013:KH.C3.716_EFNA1/2/3/4/5 0.699  1.004073 0.398 0.870 0.618
## KH2013:KH.C6.44_KCNK10/2/4        0.692  1.027052 0.384 0.674 0.298
## KH2013:KH.C3.299                  0.691  1.150896 0.382 0.587 0.252
## DLG5                              0.675  1.005763 0.350 0.630 0.321
## KH2013:KH.S725.5                  0.670  1.190847 0.340 0.652 0.382
## KH2013:KH.C2.147                  0.659  1.120100 0.318 0.522 0.244
```

```
## KH2013:KH.L108.49            0.638  1.017853 0.276 0.543 0.290
## MATN1/3/4                    0.628  1.046960 0.256 0.652 0.481
## FIR1                         0.620  1.138595 0.240 0.543 0.336
## ODF2                         0.589  1.003972 0.178 0.565 0.458
## MICALL1/2                    0.532 -1.529114 0.064 0.565 0.481
## SMEK1/2                      0.529  1.923853 0.058 0.565 0.534
```

```r
# Visualize known markers with a violin plot
vlnPlot(hpf18, c("NKX2-3", "EBF1/2/3/4", "TBX1/10", "DACH1/2"))
```

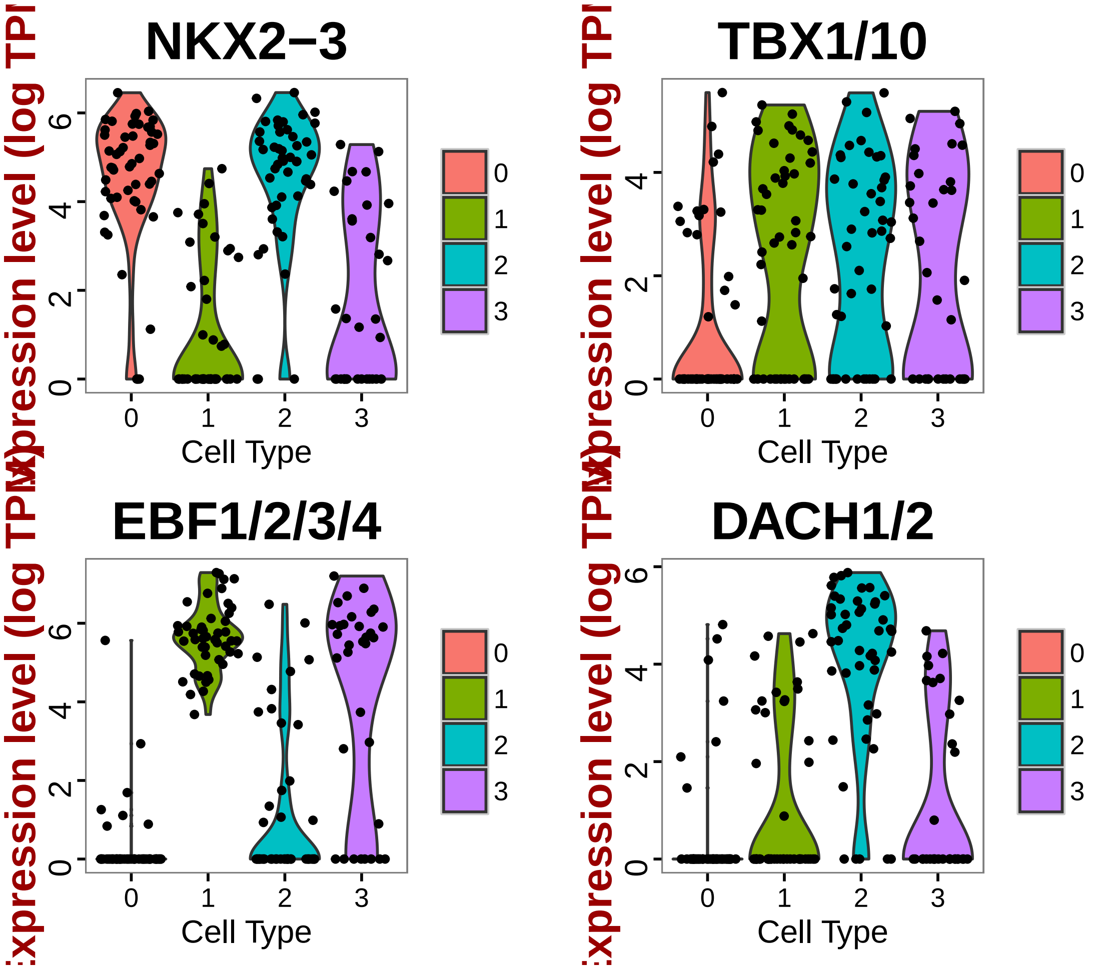


```r
# Based on preliminary discoveries these NKX+ EBF1/2/3/4- TBX+ DACH1/2+
# cells are SHP's

# Visualize new markers with a violin plot
vlnPlot(hpf18, c("KH2013:KH.C10.174", "SCYL"))
```

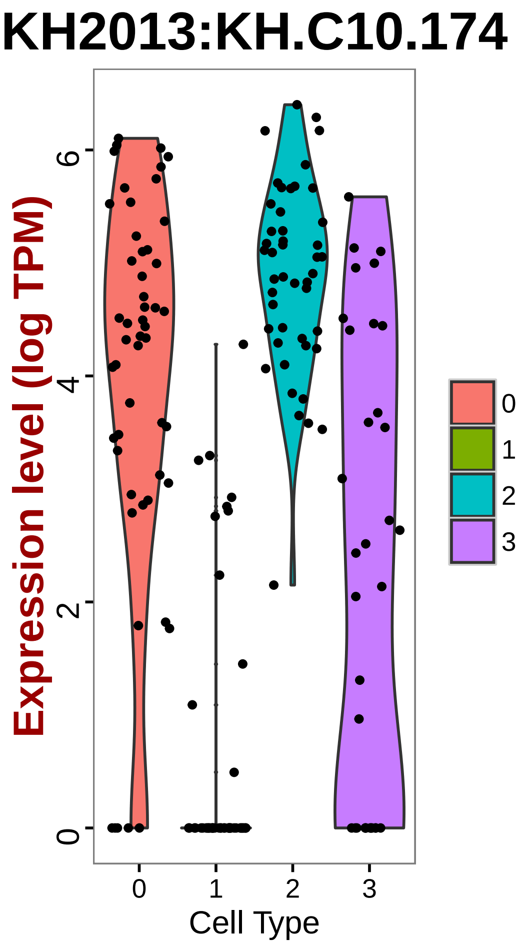


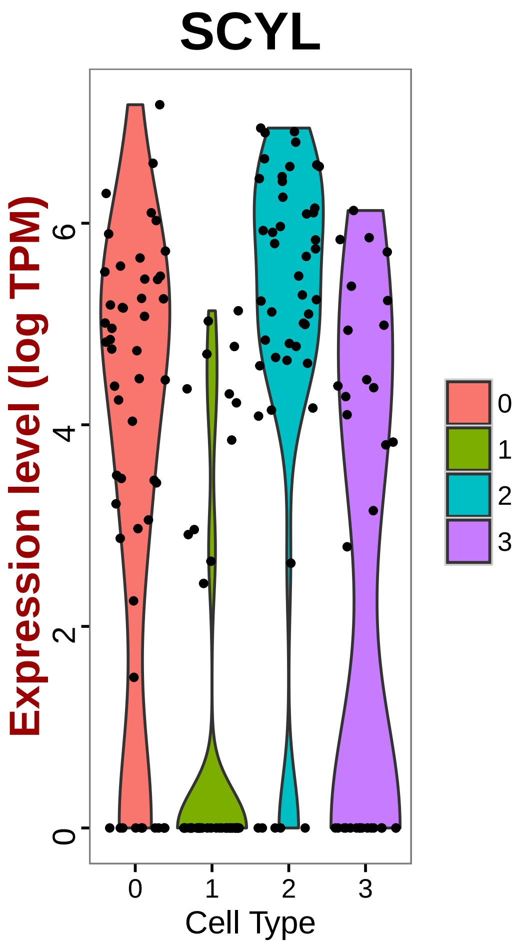


```r
# Find markers for cluster 3
cl3_18.markers = find.markers(hpf18, 3, thresh.use = 1, test.use = "roc", min.pct = 0.5)
head(cl3_18.markers[order(cl3_18.markers$myAUC, decreasing = T), ], 20)
```

```
##                           myAUC  avg_diff power pct.1 pct.2
## TMEM144                   0.652  1.230936 0.304 0.515 0.243
## LGR4/5/6                  0.643  1.121841 0.286 0.727 0.542
## BECN1                     0.640  1.131217 0.280 0.576 0.319
## SLC6A1/5                  0.621  1.602470 0.242 0.636 0.361
## PARP2                     0.613  1.028850 0.226 0.576 0.396
## KH2013:KH.C9.182_AADAT    0.585  1.062296 0.170 0.515 0.438
## ITPKA                     0.544  1.161089 0.088 0.515 0.535
## MICALL1/2                 0.535 -1.558863 0.070 0.576 0.486
## PIGT                      0.489 -4.667773 0.022 0.545 0.556
## RBM15B                    0.473 -2.221725 0.054 0.697 0.792
## SMEK1/2                   0.471 -2.776749 0.058 0.515 0.549
## KH2013:KH.C1.953_CG32702 0.405 -1.000614 0.190 0.576 0.632
## RARG                      0.392 -1.025694 0.216 0.455 0.549
## TTC19                     0.369 -1.209717 0.262 0.273 0.514
```

```
## KH2013:KH.C8.190         0.358 -2.057851 0.284 0.333 0.549
## GPR37                    0.354 -1.300561 0.292 0.424 0.583
## KH2013:KH.C7.695_CYP24A1 0.340 -1.290304 0.320 0.333 0.604
## SLIT1/2/3                0.318 -1.109890 0.364 0.545 0.743
## RBMS1/2/3                0.302 -1.117102 0.396 0.394 0.701
## NKX2-3                   0.291 -1.361200 0.418 0.515 0.729
```

```r
# Visualize known markers with a violin plot
vlnPlot(hpf18, c("NKX2-3", "EBF1/2/3/4", "TBX1/10", "DACH1/2"))
```

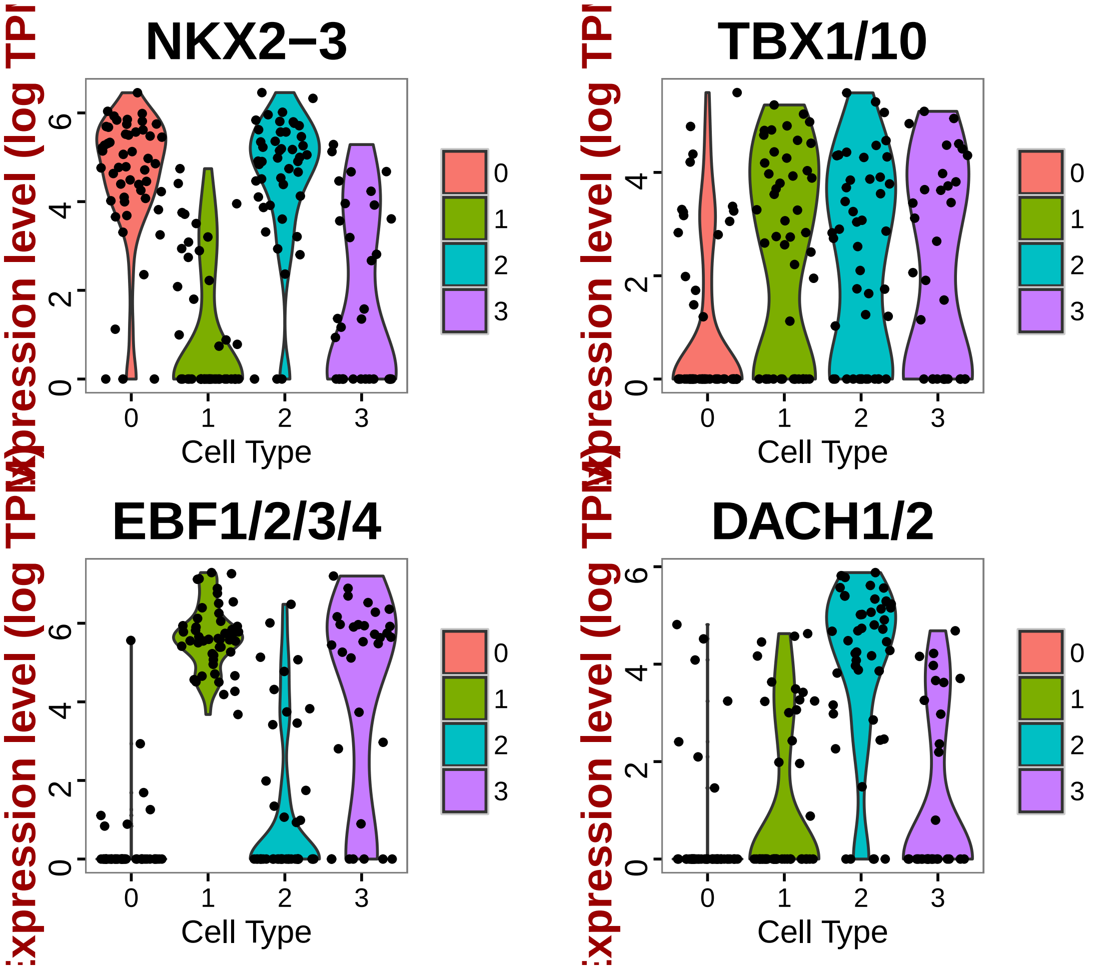


```r
# Based on preliminary discoveries these NKX- EBF1/2/3/4+ TBX+ DACH1/2-
# cells look like ASM but has few highly expressed markers

# Visualize new markers with a violin plot
vlnPlot(hpf18, c("TMEM144", "LGR4/5/6"))
```

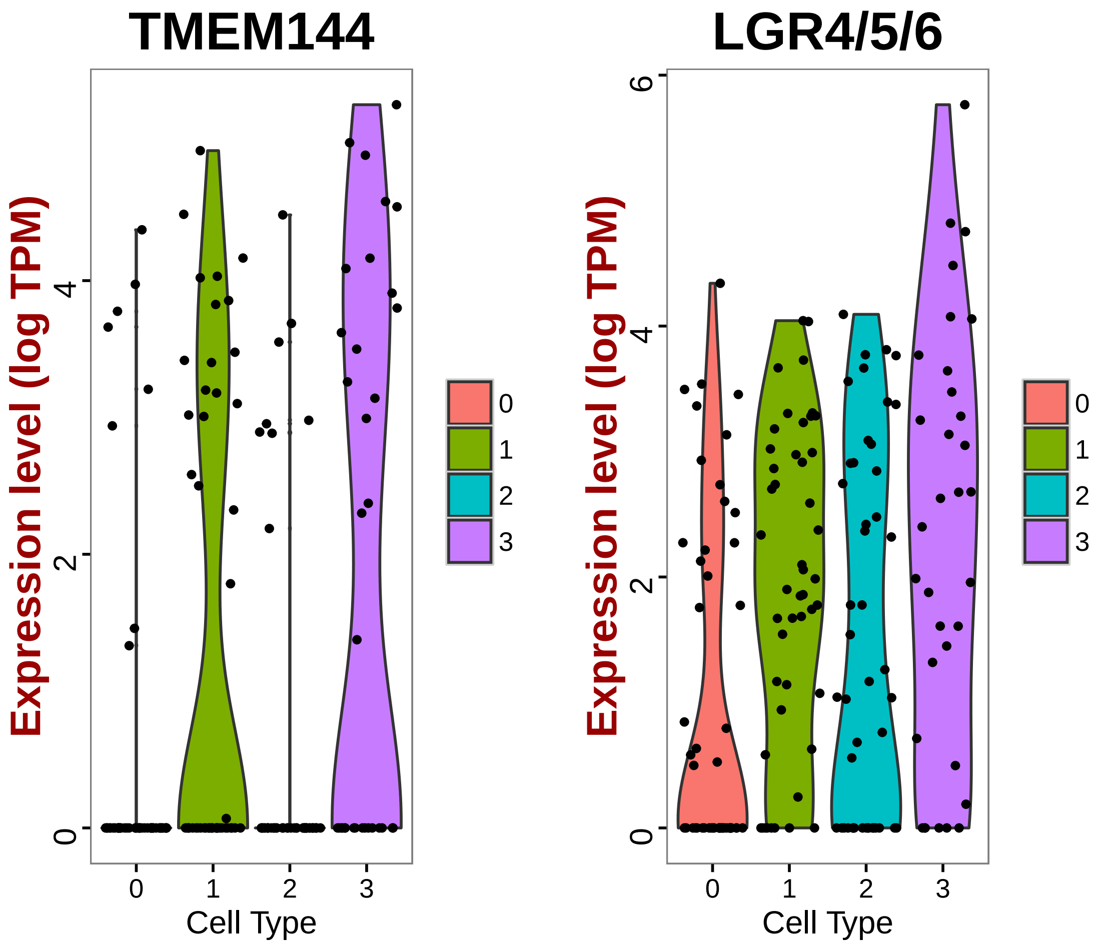


```r
# Plot hpf20 markers
doHeatMap(hpf18, genes.use = marker.20, slim.col.label = T, remove.key = T,
    rowsep = seq(0, 60, 15), col.use = col)
```

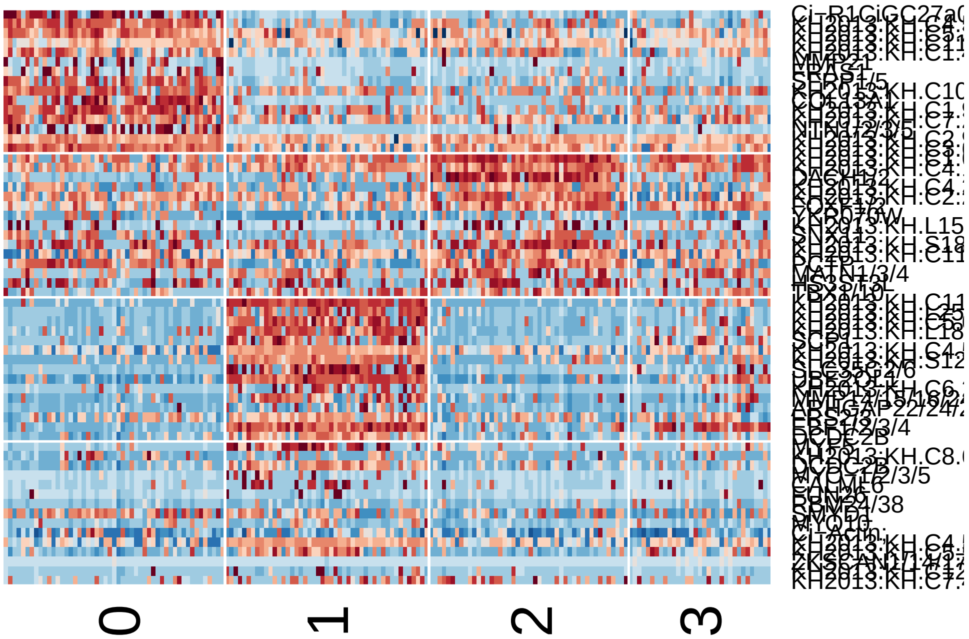


```r
# Based on the heatmap this group of cells do not express 20ASM2
# markers(later differentiated markers), also it tend to express some heart
# progenitor markers.  Named Unknown for late analysis

# Write cell names and markers into text files
write.table(which.cells(hpf18, 0), file = "18FHPCells.txt", sep = "\t")
write.table(which.cells(hpf18, 1), file = "18ASMCells.txt", sep = "\t")
write.table(which.cells(hpf18, 2), file = "18SHPCells.txt", sep = "\t")

# Rename cluster identities
hpf18 = rename.ident(hpf18, 0, "18FHP")
hpf18 = rename.ident(hpf18, 1, "18ASM")
hpf18 = rename.ident(hpf18, 2, "18SHP")
hpf18 = rename.ident(hpf18, 3, "18Unknown")

# Visualize tSNE used color scheme FHP-red, SHP-orange, ASM-blue
tsne.plot(hpf18, do.label = T, label.pt.size = 1, label.cex.text = 1.2, label.cols.use =
    "red", "orange", "blue"))
```

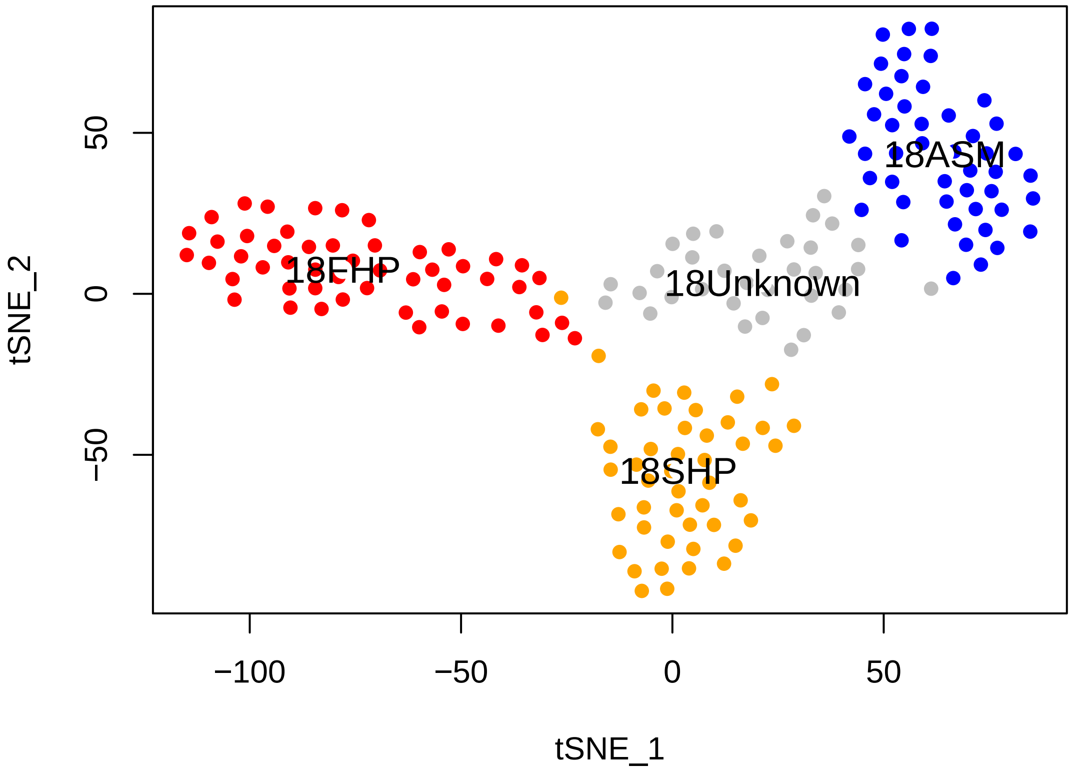


```r
# Store FHP markers in text file
FHP_18.markers = find.markers(hpf18, "18FHP", c("18SHP", "18ASM"), thresh.use = 1,
    test.use = "roc", min.pct = 0.5)
head(FHP_18.markers[order(FHP_18.markers$myAUC, decreasing = T), ], 20)
```

```
##                            myAUC avg_diff power pct.1 pct.2
## MOGAT1/2                   0.921 1.695660 0.842 1.000 0.591
## NAV1                       0.909 2.469266 0.818 0.882 0.151
## KH2013:KH.C3.665_ZAN       0.909 1.565571 0.818 0.980 0.817
## KH2013:KH.C5.227_F56C4.4 0.892 1.683649 0.784 1.000 0.946
## BMP5/6/7                   0.891 2.057759 0.782 0.941 0.301
## KH2013:KH.C4.547_BMP2/4    0.886 1.917017 0.772 0.961 0.452
## KH2013:KH.C1.953_CG32702 0.849 1.780812 0.698 0.902 0.484
## LAMA1/2                    0.838 1.650821 0.676 0.588 0.054
## SFRP1/5                    0.828 2.941007 0.656 0.745 0.172
## COL13A1                    0.826 2.513739 0.652 0.725 0.118
## Ci-R1CiGC27a04             0.824 4.002483 0.648 0.686 0.086
## C5ORF48                    0.821 2.357489 0.642 0.784 0.280
## FZD4                       0.821 1.810412 0.642 0.784 0.183
## KH2013:KH.C10.203          0.804 1.277857 0.608 0.902 0.645
## LYS2                       0.803 1.212652 0.606 0.863 0.430
## EFNB1/2/3                  0.798 1.092338 0.596 0.922 0.387
## KH2013:KH.C11.362          0.792 2.079359 0.584 0.784 0.355
```

```
## NTN1/2/3/5                0.786 2.737731 0.572 0.588 0.043
## KH2013:KH.C7.695_CYP24A1 0.786 1.784354 0.572 0.863 0.462
## WNT9B                    0.763 1.601544 0.526 0.725 0.301
```

```
write.table(FHP_18.markers, file = "FHP_18.markers.txt", sep = "\t")

# Store SHP markers in text file
SHP_18.markers = find.markers(hpf18, "18SHP", c("18FHP", "18ASM"), thresh.use = 1,
    test.use = "roc", min.pct = 0.5)
head(SHP_18.markers[order(SHP_18.markers$myAUC, decreasing = T), ], 20)
```

```
##                            myAUC avg_diff power pct.1 pct.2
## DACH1/2                    0.892 2.438529 0.784 0.891 0.224
## KH2013:KH.C1.638_C17ORF105 0.860 1.393115 0.720 0.935 0.714
## SCYL                       0.794 1.296783 0.588 0.891 0.531
## FOXF1/2                    0.790 1.416604 0.580 0.913 0.704
## KH2013:KH.C10.172          0.769 1.482743 0.538 0.848 0.582
## SPSB1/4                    0.753 1.110795 0.506 0.978 0.796
## SUDS3                      0.749 1.180373 0.498 0.913 0.704
## ZNF703                     0.719 1.080167 0.438 0.652 0.245
## KH2013:KH.C6.44_KCNK10/2/4 0.704 1.173353 0.408 0.674 0.296
## KH2013:KH.C3.232           0.703 1.013660 0.406 0.674 0.347
## CCNB3                      0.692 1.069960 0.384 0.717 0.408
## KH2013:KH.C3.299           0.688 1.224314 0.376 0.587 0.276
## DLG5                       0.681 1.004789 0.362 0.630 0.316
## KH2013:KH.C2.147           0.671 1.243503 0.342 0.522 0.224
## KH2013:KH.S725.5           0.668 1.163315 0.336 0.652 0.398
## MATN1/3/4                  0.649 1.204521 0.298 0.652 0.439
## KH2013:KH.C14.406          0.638 1.101730 0.276 0.717 0.531
## KH2013:KH.C12.536          0.630 1.082419 0.260 0.848 0.786
## histone                    0.629 1.006188 0.258 0.739 0.673
## KH2013:KH.S1660.1          0.591 1.010002 0.182 0.739 0.684
```

```
write.table(SHP_18.markers, file = "SHP_18.markers.txt", sep = "\t")

# Store ASM markers in text file
ASM_18.markers = find.markers(hpf18, "18ASM", c("18SHP", "18FHP"), thresh.use = 1,
    test.use = "roc", min.pct = 0.5)
head(ASM_18.markers[order(ASM_18.markers$myAUC, decreasing = T), ], 20)
```

```
##                            myAUC avg_diff power pct.1 pct.2
## KH2013:KH.C11.139          0.984 4.588292 0.968 0.979 0.196
## UBE2QL1                    0.984 3.208404 0.968 1.000 0.258
```

```
## EBF1/2/3/4                 0.971 2.877694 0.942 1.000 0.196
## BDH1                       0.964 2.570204 0.928 0.979 0.381
## KH2013:KH.C4.506_HMCN1     0.942 1.738256 0.884 1.000 0.794
## Ci-R1CiGC09b24;            0.939 2.065758 0.878 1.000 0.753
## KH2013:KH.C10.209          0.931 1.544434 0.862 1.000 0.732
## KH2013:KH.C5.127_FMO1      0.928 4.482840 0.856 0.872 0.082
## KH2013:KH.C5.62_ODF3L2     0.919 3.552450 0.838 0.872 0.072
## SI:CH211-15101.2           0.917 2.834658 0.834 0.894 0.155
## KH2013:KH.C8.782_LPHN1/2/3 0.910 1.483750 0.820 1.000 0.670
## KH2013:KH.L18.10_THSD7A    0.910 2.158503 0.820 0.872 0.175
## KH2013:KH.S1269.1          0.890 1.958529 0.780 0.915 0.289
## KH2013:KH.L154.26_CSMD2    0.889 1.614030 0.778 0.957 0.536
## TMSB15A                    0.886 2.831017 0.772 0.872 0.247
## KH2013:KH.S511.3           0.882 1.513598 0.764 0.979 0.454
## SLC35G2/6                  0.877 4.418579 0.754 0.766 0.041
## KH2013:KH.C7.805_ID1/2/3/4 0.874 1.352515 0.748 0.979 0.804
## ELK1/3/4                   0.864 1.368382 0.728 0.979 0.423
## KH2013:KH.S555.1_HTR7      0.862 1.327817 0.724 0.979 0.443
```

```
write.table(ASM_18.markers, file = "ASM_18.markers.txt", sep = "\t")

# Find pan Heart Progenitor markers
panHP_18.markers = find.markers(hpf18, c("18SHP", "18FHP"), "18ASM", thresh.use = 1,
    test.use = "roc", min.pct = 0.5)
head(panHP_18.markers[order(panHP_18.markers$myAUC, decreasing = T), ], 20)
```

```
##                                 myAUC avg_diff power pct.1 pct.2
## KH2013:KH.C10.174               0.952 3.337930 0.904 0.948 0.213
## SLIT1/2/3                       0.949 2.523519 0.898 0.979 0.255
## GATA4/5/6                       0.946 3.202716 0.892 0.938 0.191
## QKI                             0.931 2.260126 0.862 0.979 0.723
## NKX2-3                          0.928 2.720123 0.856 0.938 0.298
## SPECC1                          0.916 2.230076 0.832 0.979 0.489
## PCTP                            0.910 2.477431 0.820 0.938 0.362
## TNS1/3                          0.901 2.394328 0.802 0.928 0.255
## EFNB1/2/3                       0.900 3.684817 0.800 0.825 0.064
## KH2013:KH.C8.80_SI:DKEY-29D8.3  0.895 1.182032 0.790 1.000 0.872
## LRP4/8                          0.892 2.625513 0.784 0.825 0.085
## KH2013:KH.C8.489_SI:DKEY-29D8.3 0.887 2.125379 0.774 0.928 0.617
## KH2013:KH.C2.994_RNF149         0.885 1.044521 0.770 1.000 0.979
## BMP5/6/7                        0.877 4.532942 0.754 0.742 0.085
## KH2013:KH.C2.935_S1PR1/2/3/4/5  0.875 1.766753 0.750 0.969 0.468
## SH3BGR                          0.873 2.986558 0.746 0.804 0.128
## KH2013:KH.C3.665_ZAN            0.868 1.999101 0.736 0.959 0.702
```

```
## SCYL                               0.855 2.519750 0.710 0.835 0.255
## IGF1/2                             0.854 2.586108 0.708 0.794 0.149
## LYS2                               0.850 2.201142 0.700 0.773 0.191
```

```r
write.table(panHP_18.markers, file = "panHP_18.markers.txt", sep = "\t")

# Find SHP specific markers that distinguish tow heart progenitors FHP and
# SHP
SHPspecific_18.markers = find.markers(hpf18, "18SHP", "18FHP", thresh.use = 1,
    test.use = "roc", min.pct = 0.5)
head(SHPspecific_18.markers[order(SHPspecific_18.markers$myAUC, decreasing = T),
    ], 20)
```

```
##                               myAUC avg_diff power pct.1 pct.2
## DACH1/2                       0.911 2.783039 0.822 0.891 0.137
## KH2013:KH.C1.638_C17ORF105    0.878 1.574147 0.756 0.935 0.588
## GDH1/3                        0.807 1.282798 0.614 0.978 0.824
## ZFP318                        0.797 1.205806 0.594 0.870 0.588
## KH2013:KH.L154.26_CSMD2       0.794 1.907616 0.588 0.783 0.314
## KH2013:KH.C9.692_GABRR1/2/3 0.768 2.048822 0.536 0.717 0.333
## TUBA1L/2                      0.767 1.329805 0.534 0.935 0.941
## FOXF1/2                       0.761 1.189351 0.522 0.913 0.765
## ZNF703                        0.757 1.547560 0.514 0.652 0.157
## ELK1/3/4                      0.746 1.261431 0.492 0.652 0.216
## KH2013:KH.C10.172             0.731 1.248617 0.462 0.848 0.725
## PCBP3                         0.720 1.574723 0.440 0.630 0.314
## MATN1/3/4                     0.708 1.603016 0.416 0.652 0.294
## NOTCH1/4                      0.708 2.316566 0.416 0.522 0.118
## KH2013:KH.C9.208_TTN          0.705 1.052186 0.410 0.630 0.314
## ZNF367                        0.702 1.026594 0.404 0.652 0.275
## PRKD1/2/3                     0.697 1.127657 0.394 0.587 0.353
## CCNB3                         0.696 1.039161 0.392 0.717 0.373
## NOVA1/2                       0.693 1.657795 0.386 0.609 0.314
## KH2013:KH.C14.406             0.689 1.426562 0.378 0.717 0.471
```

```r
write.table(SHPspecific_18.markers, file = "SHPspecific_18.markers.txt", sep = "\t")

# Find FHP specific markers that distinguish tow heart progenitors FHP and
# SHP
FHPspecific_18.markers = find.markers(hpf18, "18FHP", "18SHP", thresh.use = 1,
    test.use = "roc", min.pct = 0.5)
head(FHPspecific_18.markers[order(FHPspecific_18.markers$myAUC, decreasing = T),
    ], 20)
```

```
##                            myAUC avg_diff power pct.1 pct.2
## MOGAT1/2                   0.866 1.227366 0.732 1.000 0.696
## KH2013:KH.C1.953_CG32702 0.865 1.958750 0.730 0.902 0.391
## NAV1                       0.862 1.946789 0.724 0.882 0.283
## KH2013:KH.C5.227_F56C4.4 0.859 1.443269 0.718 1.000 0.935
## KH2013:KH.C3.665_ZAN       0.844 1.114215 0.688 0.980 0.935
## KH2013:KH.C4.547_BMP2/4    0.841 1.478739 0.682 0.961 0.565
## BMP5/6/7                   0.814 1.381609 0.628 0.941 0.522
## COL13A1                    0.813 2.370790 0.626 0.725 0.174
## Ci-R1CiGC27a04             0.809 3.468222 0.618 0.686 0.152
## C5ORF48                    0.808 2.160508 0.616 0.784 0.326
## LAMA1/2                    0.806 1.428195 0.612 0.588 0.109
## KH2013:KH.C10.203          0.802 1.238416 0.604 0.902 0.587
## SFRP1/5                    0.801 2.553373 0.602 0.745 0.283
## NTN1/2/3/5                 0.769 2.193801 0.538 0.588 0.087
## KH2013:KH.C3.21_HSPG2      0.751 2.223697 0.502 0.549 0.130
## FZD4                       0.751 1.169858 0.502 0.784 0.370
## KH2013:KH.C11.362          0.749 1.660434 0.498 0.784 0.478
## KH2013:KH.C7.695_CYP24A1 0.738 1.434126 0.476 0.863 0.587
## DSEL                       0.734 1.339817 0.468 0.667 0.239
## KH2013:KH.C7.649_ENPP7     0.723 1.007389 0.446 0.765 0.435
```

```
write.table(FHPspecific_18.markers, file = "FHPspecific_18.markers.txt", sep = "\t")

# Find gene expression percentage among single cells
hpf18.pct = cluster.alpha(hpf18, thresh.min = 0)
# Find top 50 pan heart marker expression percentage in 18Unknown cells
sum(hpf18.pct[rownames(head(panHP_18.markers[order(panHP_18.markers$myAUC, decreasing =
    ], 50)), 4] > 0.5)/50
```

```
## [1] 0.7
```

```
# Unknown cells have 70% top 50 pan heart marker expressed
sum(hpf18.pct[rownames(head(panHP_18.markers[order(panHP_18.markers$myAUC, decreasing =
    ], 50)), c(1, 3)] > 0.5)/100
```

```
## [1] 0.98
```

```
# FHP and SHP cells have only 98% top 50 pan heart marker expressed
sum(hpf18.pct[rownames(head(panHP_18.markers[order(panHP_18.markers$myAUC, decreasing =
    ], 50)), 2] > 0.5)/100
```

```
## [1] 0.21
```

```r
# ASM cells have 21% top 50 pan heart marker expressed based on pan heart
# markers the Unknwon cells express more heart progenitors markers

# Find top 50 pan heart marker expression percentage in 18Unknown cells
sum(hpf18.pct[rownames(head(ASM_18.markers[order(ASM_18.markers$myAUC, decreasing = T),
    ], 50)), 4] > 0.5)/50
```

```
## [1] 0.64
```

```r
# Unknown cells have 74% top 50 ASM marker expressed
sum(hpf18.pct[rownames(head(ASM_18.markers[order(ASM_18.markers$myAUC, decreasing = T),
    ], 50)), c(1, 3)] > 0.5)/100
```

```
## [1] 0.3
```

```r
# FHP and SHP cells have only 30% top 50 ASM marker expressed
sum(hpf18.pct[rownames(head(ASM_18.markers[order(ASM_18.markers$myAUC, decreasing = T),
    ], 50)), 2] > 0.5)/50
```

```
## [1] 1
```

```r
# ASM cells have 100% top 50 ASM marker expressed based on ASM markers the
# Unknwon cells express more ASM progenitors markers

# Heat map of 20hpf markers also demonstrates that the 18Unknown cells tend
# to have both heart and muscle progenitor characteristics
doHeatMap(hpf18, genes.use = marker.20, slim.col.label = T, remove.key = T,
    rowsep = seq(0, 60, 15), col.use = col)
```

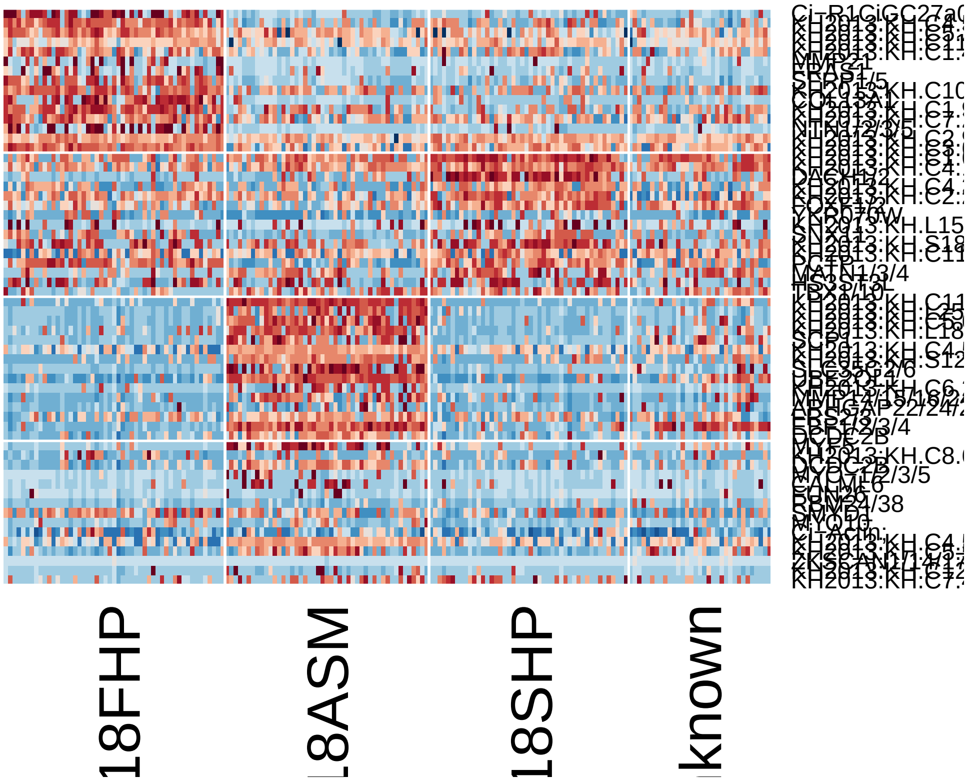


```r
# Based on available published data EBF1/2/3/4 and GATA4/5/6 inhibiting each
# other during the fate determination of ASM and SHP. No cells were
# previously reported to have both EBF and GATA expression in TVC lineage
# (FISH data).
genePlot(hpf18, cell.ids = which.cells(hpf18, "18Unknown"), "EBF1/2/3/4", "GATA4/5/6")
abline(v = 2, h = 2, lwd = 3, lty = 2)
```

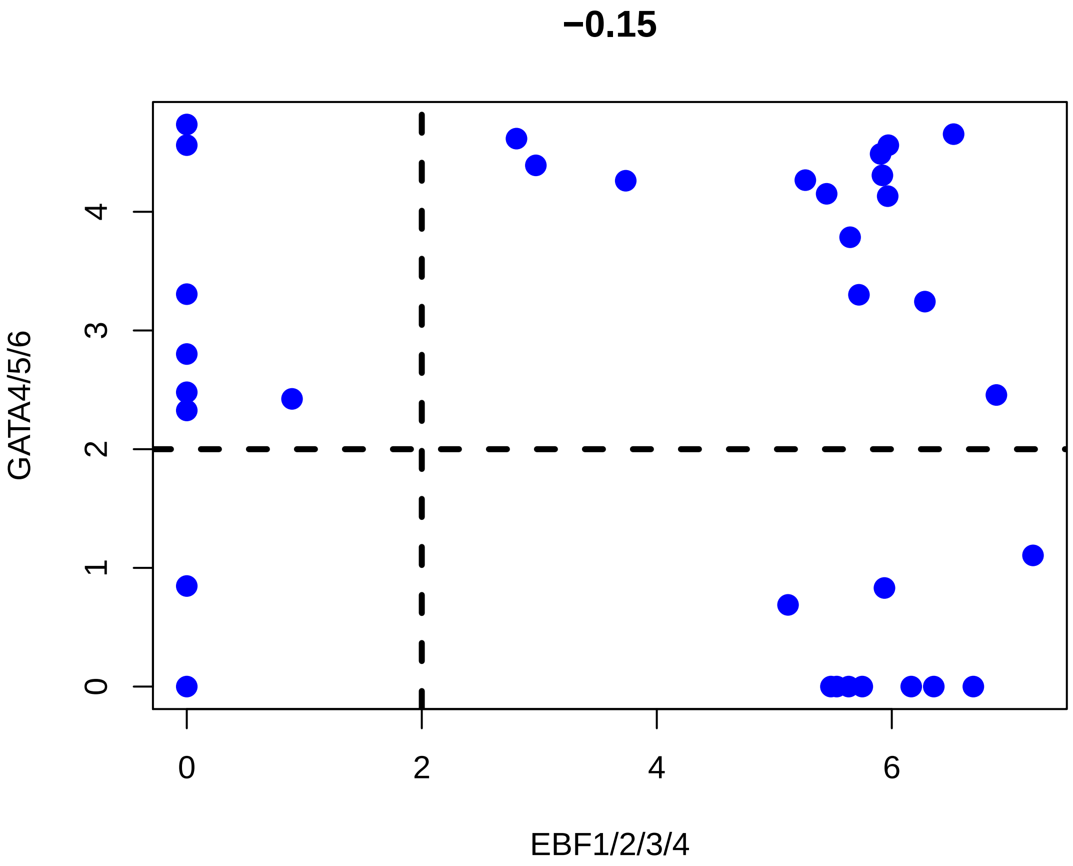


```r
# There are 14 out of 33 cells (42%) have good expression level(2 logFPKM)
# of both genes.  Thus, upon reasoning above, we conclude that this cluster
# contains cells that contradict to experimental discoveries which has a
# high probability to be doublet cells due to technical error.

# Filter the cells based on reasoning above
hpf18.new = subsetData(hpf18, which.cells(hpf18, c("18ASM", "18SHP", "18FHP")),
    do.scale = F)
tsne.plot(hpf18.new, do.label = T, cols.use = c("red", "orange", "blue"))
```

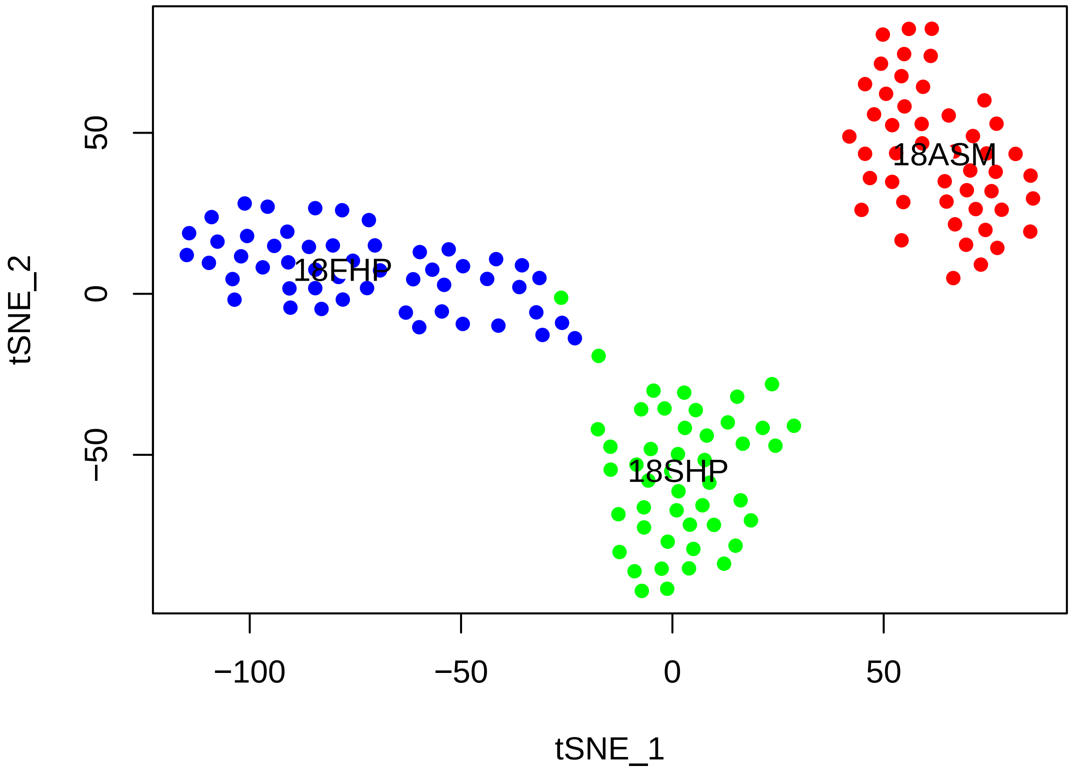


```r
# Visualize markers of different clusters using violin plot and feature plot
hpf18.new@ident = factor(hpf18.new@ident, ordered = T, levels = c("18FHP", "18SHP",
    "18ASM"))
genes.viz.18 = c("DACH1/2", "NKX2-3", "EBF1/2/3/4", "SFRP1/5")
feature.plot(hpf18.new, genes.viz.18, pt.size = 0.8)
```

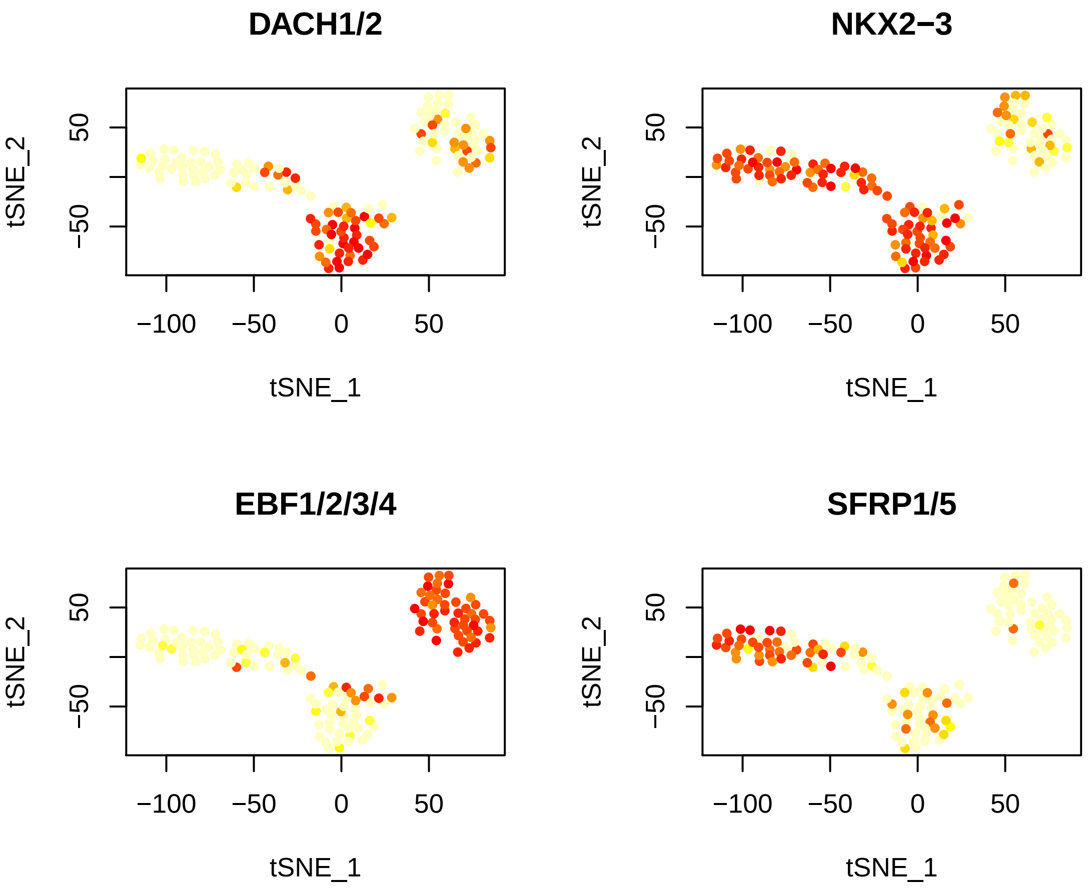


```r
vlnPlot(hpf18.new, genes.viz.18, cols.use = c("red", "orange", "blue"))
```

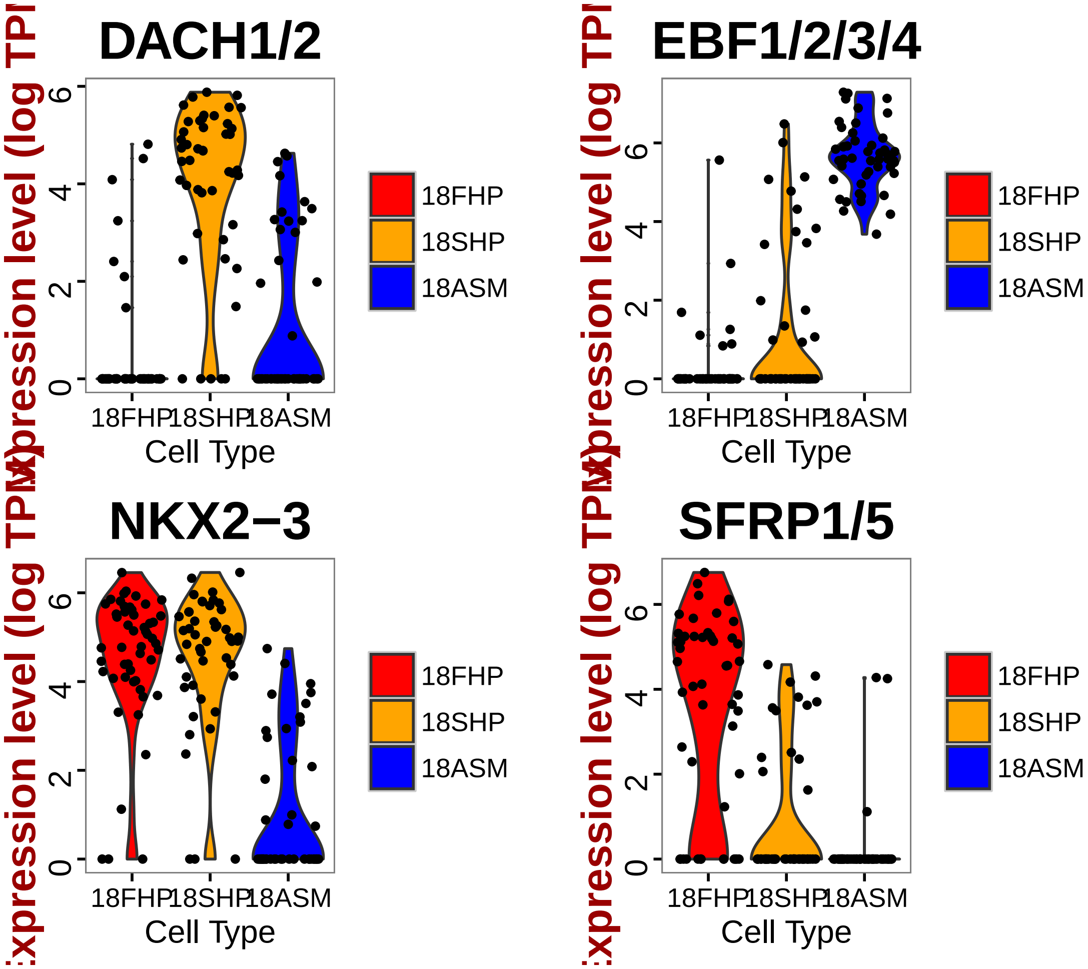


```
# Select markers for plotting on a Heatmap (top 10 positive markers with
# high classfication power(myAUC))
marker18FHP = rownames(FHP_18.markers[order(FHP_18.markers$myAUC, decreasing = T)[1:15],
    ])
marker18SHP = rownames(SHP_18.markers[order(SHP_18.markers$myAUC, decreasing = T)[1:15],
    ])
marker18ASM = rownames(ASM_18.markers[order(ASM_18.markers$myAUC, decreasing = T)[1:15],
    ])
marker.18 = c(marker18FHP, marker18SHP, marker18ASM)

# Draw a heatmap of all cells for these marker genes
doHeatMap(hpf18.new, genes.use = marker.18, remove.key = TRUE, slim.col.label = T,
    cex.col = 1.2, col.use = col)
```

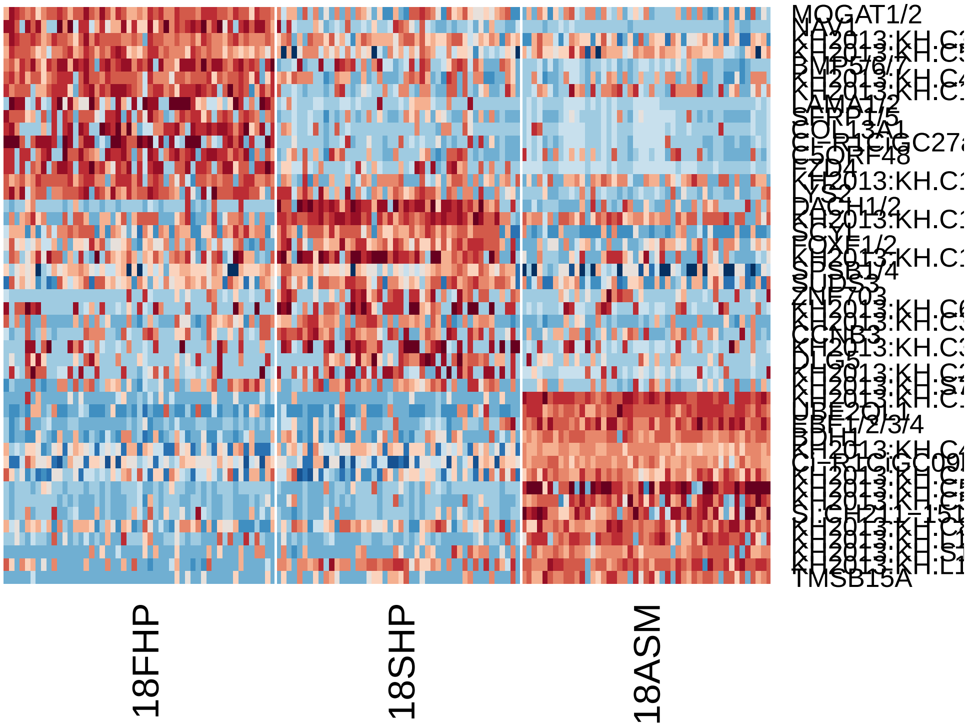


6. hpf16

```r
# Subset data from preprocessed Seurat object
hpf16 = subsetData(hpfall.remv2, which.cells(hpfall.remv2, "hpf16"), do.scale = F)
hpf16
```

```
## An object of class seurat in project allhpf
##  14864 genes across 114 samples.
```

```r
# Based on hpf12 hpf14 hpf18 and hpf20 data, we have successfully uncovered
# all the TVC lineage types: TVC, STVC, FHP, SHP, ASM Based on preliminary
# studies, hpf16 contains three TVC lineage cell types: FHP, late STVC,
# early SHP and early ASM which are divided from early STVC. Therefore hpf16
# is an intermediate stage between hpf14 and hpf18, we can run PCA with both
# hpf14 (14STVC,14FHP) and hpf18 cell markers (18FHP, 18SHP, 18ASM).
# Power>0.4 is used to obtain more markers from hpf14 cells.
marker20FHP.use = rownames(subset(FHP_20.markers, power > 0.5 & avg_diff > 0))
marker20SHP.use = rownames(subset(SHP_20.markers, power > 0.5 & avg_diff > 0))
marker20ASM1.use = rownames(subset(ASM1_20.markers, power > 0.5 & avg_diff >
    0))
marker20ASM2.use = rownames(subset(ASM2_20.markers, power > 0.5 & avg_diff >
    0))
marker14FHP.use = rownames(subset(FHP_14.markers, power > 0.4 & avg_diff > 0))
marker14STVC.use = rownames(subset(STVC_14.markers, power > 0.4 & avg_diff >
```

```r
    0))
marker.16.use = unique(c(marker20FHP.use, marker20SHP.use, marker20ASM1.use,
    marker20ASM2.use, marker14FHP.use, marker14STVC.use))
length(marker.16.use)
```

```
## [1] 217
```

```r
# Run a PCA using marker list
hpf16 = pca(hpf16, pc.genes = marker.16.use, do.print = F)
pcScree(hpf16, marker.16.use, 10)
```

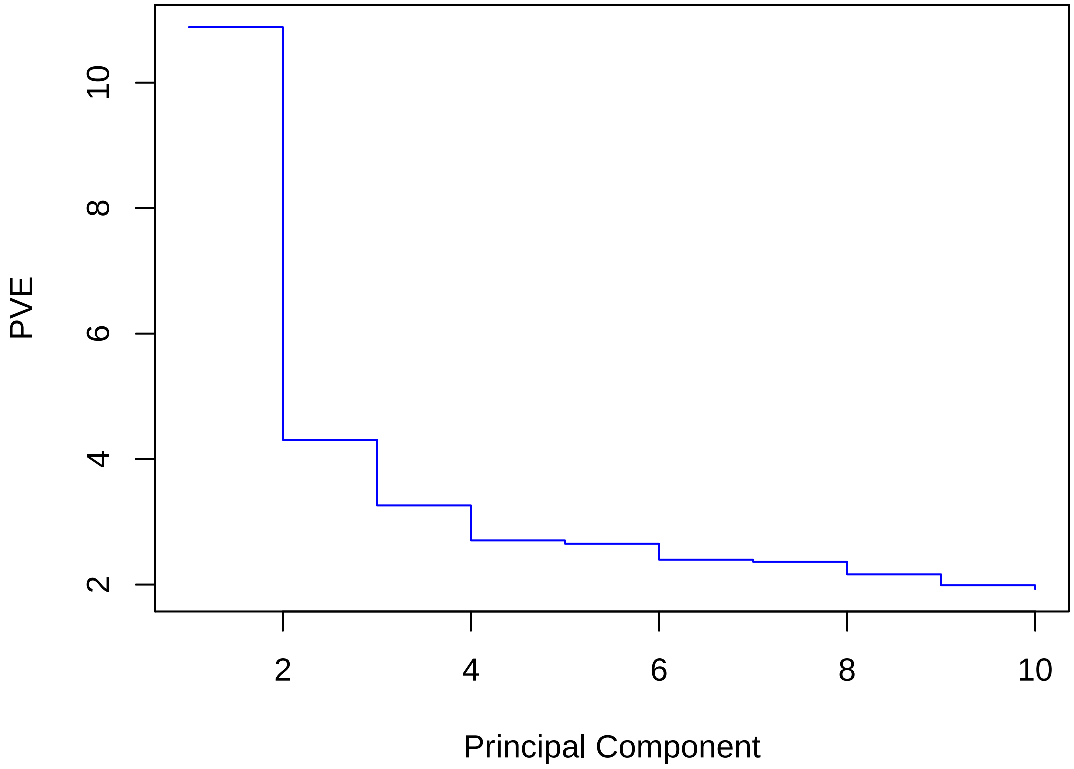


```r
pcHeatmap(hpf16, 1, do.balanced = T, col.use = col)
```

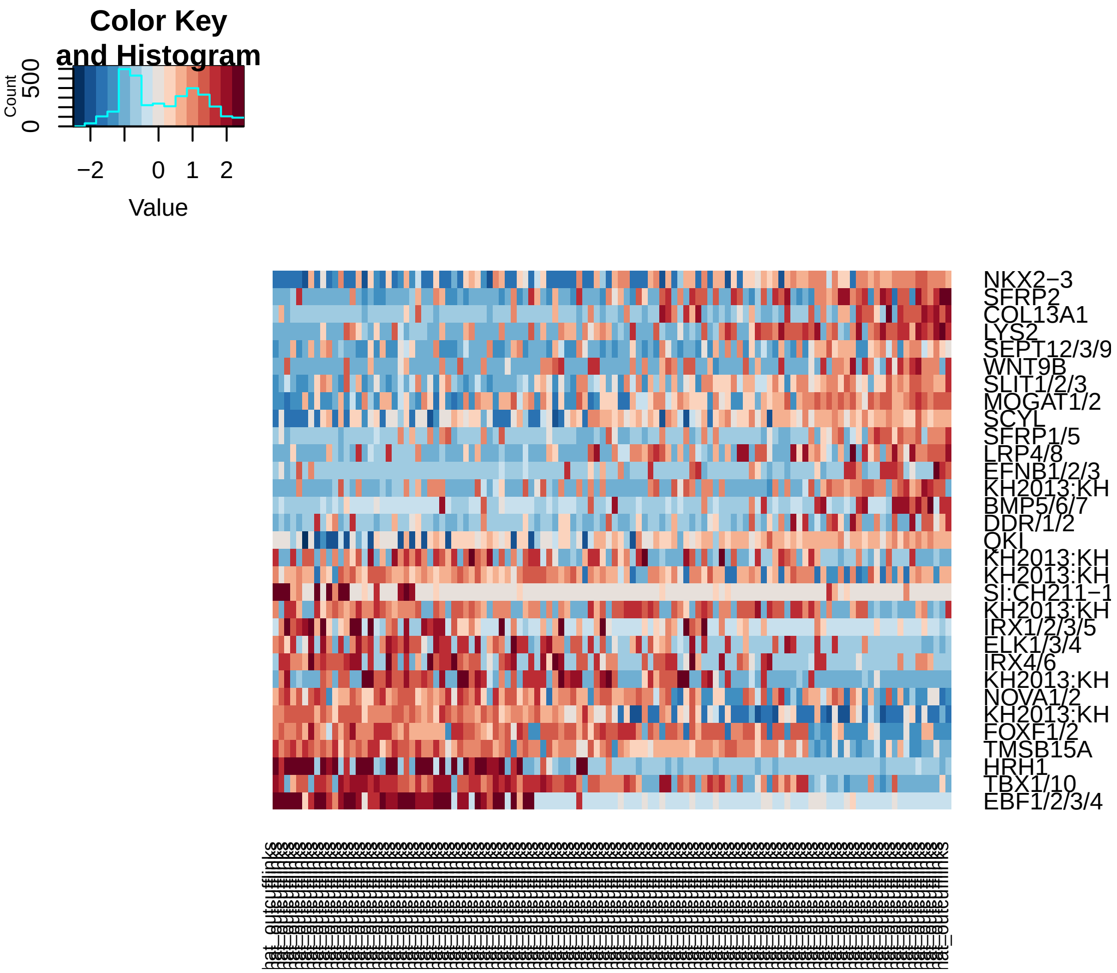


```
pcHeatmap(hpf16, 2, do.balanced = T, col.use = col)
```

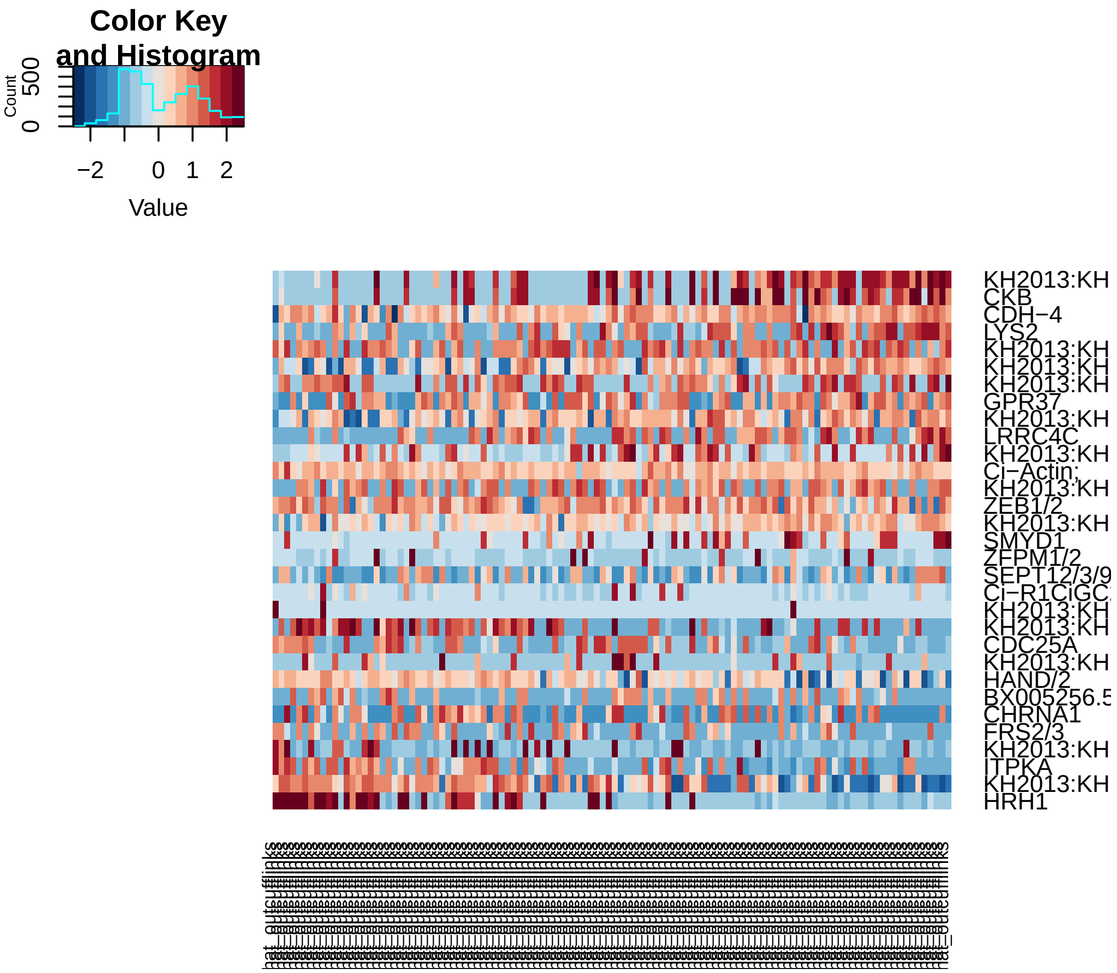


```r
pcHeatmap(hpf16, 3, do.balanced = T, col.use = col)
```

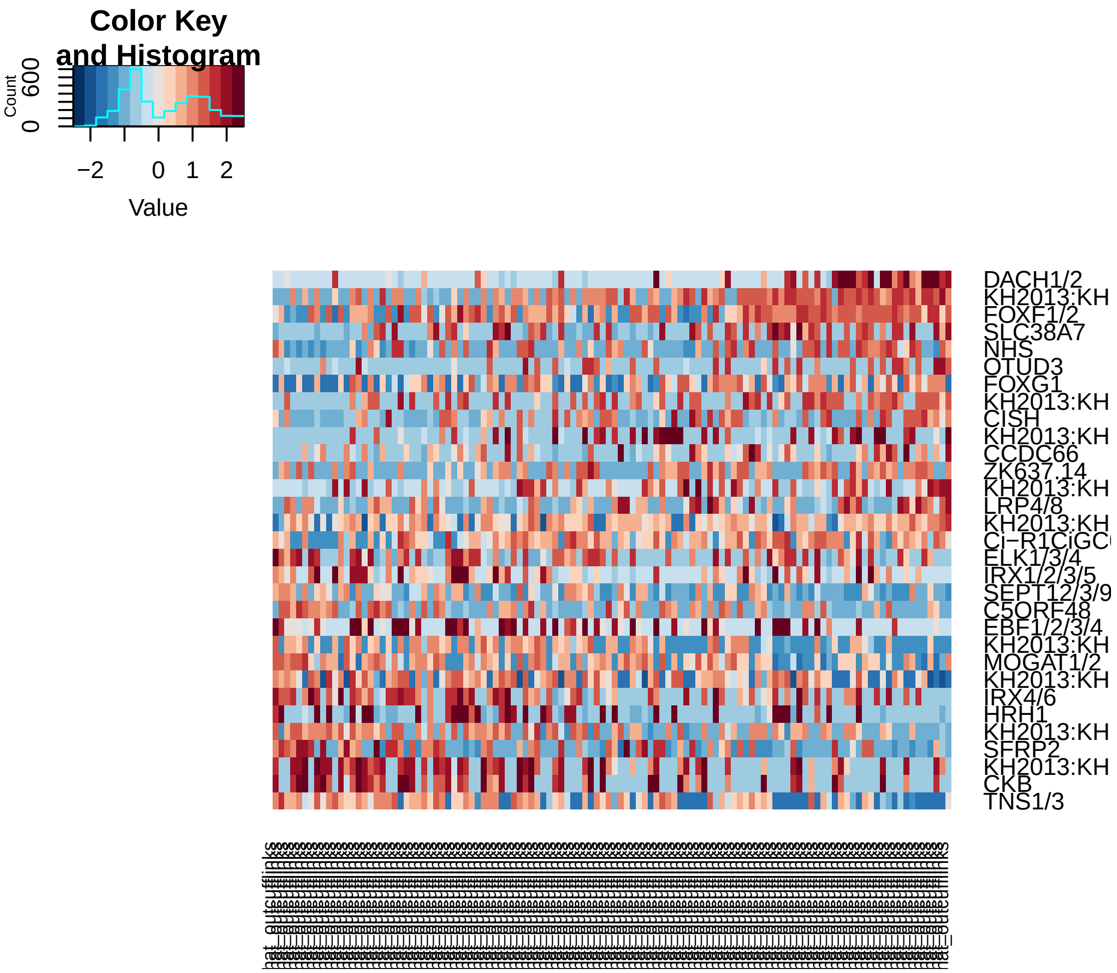


```r
pca.plot(hpf16, 1, 2)
```

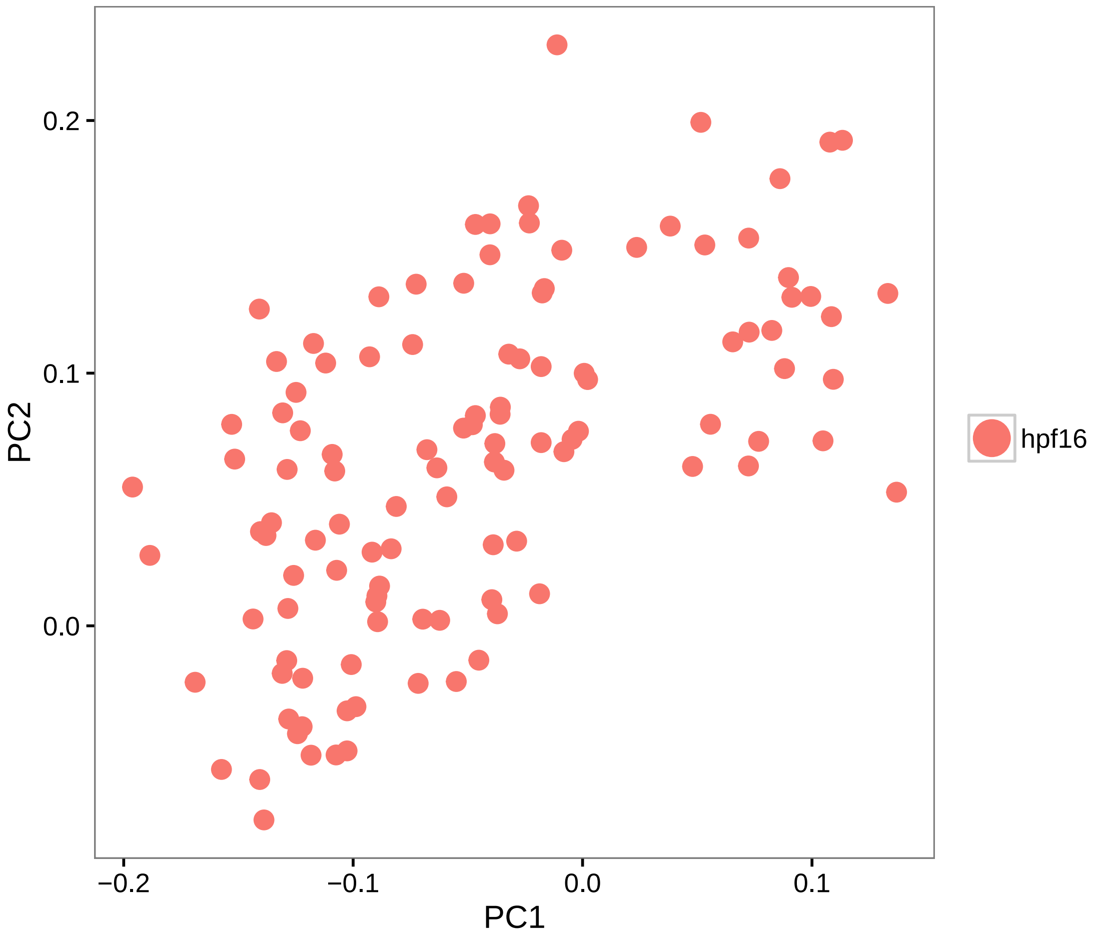


```r
pca.plot(hpf16, 1, 3)
```

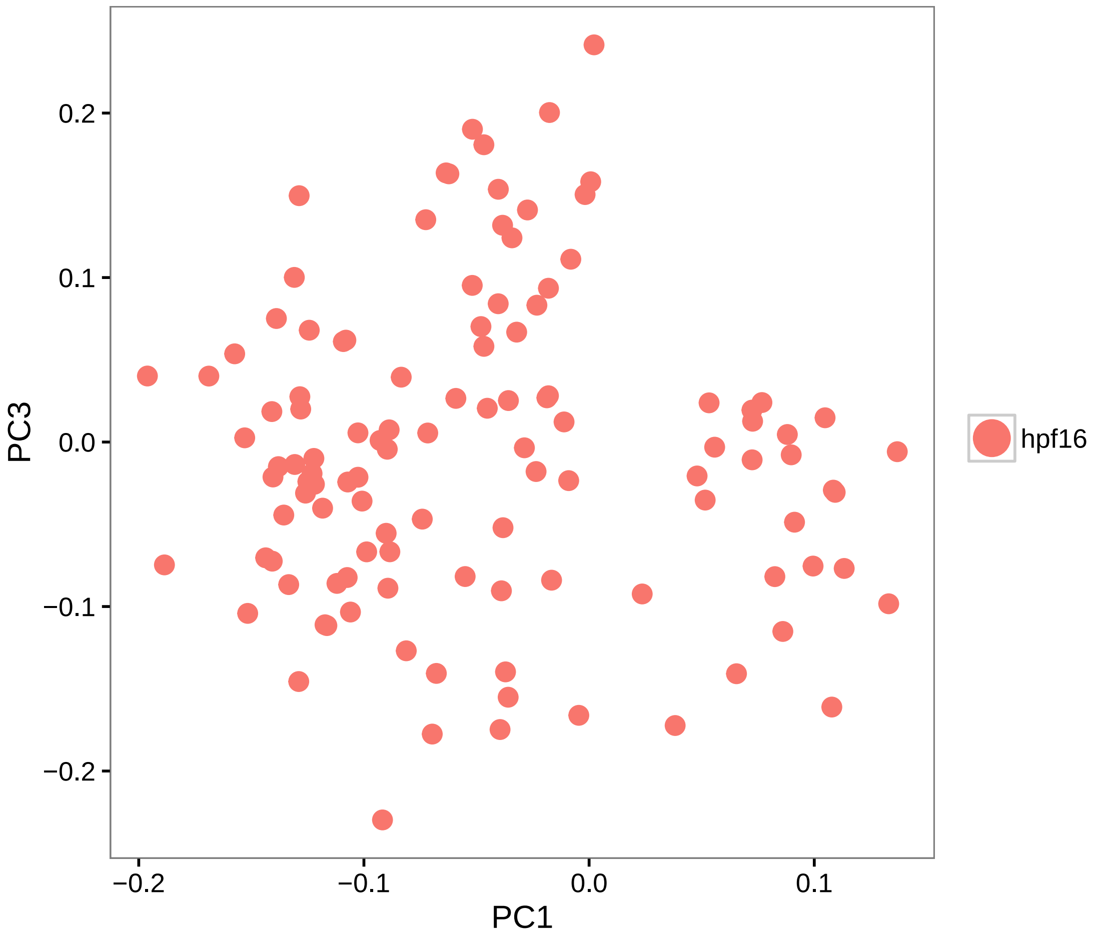


```r
# Calculate PCA scores for all genes (PCA projection)
hpf16 = project.pca(hpf16, do.print = F)

# Visualize the full projected PCA, which now includes new genes which were
# not previously (use.full=TRUE)
pcHeatmap(hpf16, 1, use.full = T, do.balanced = T, col.use = col)
```

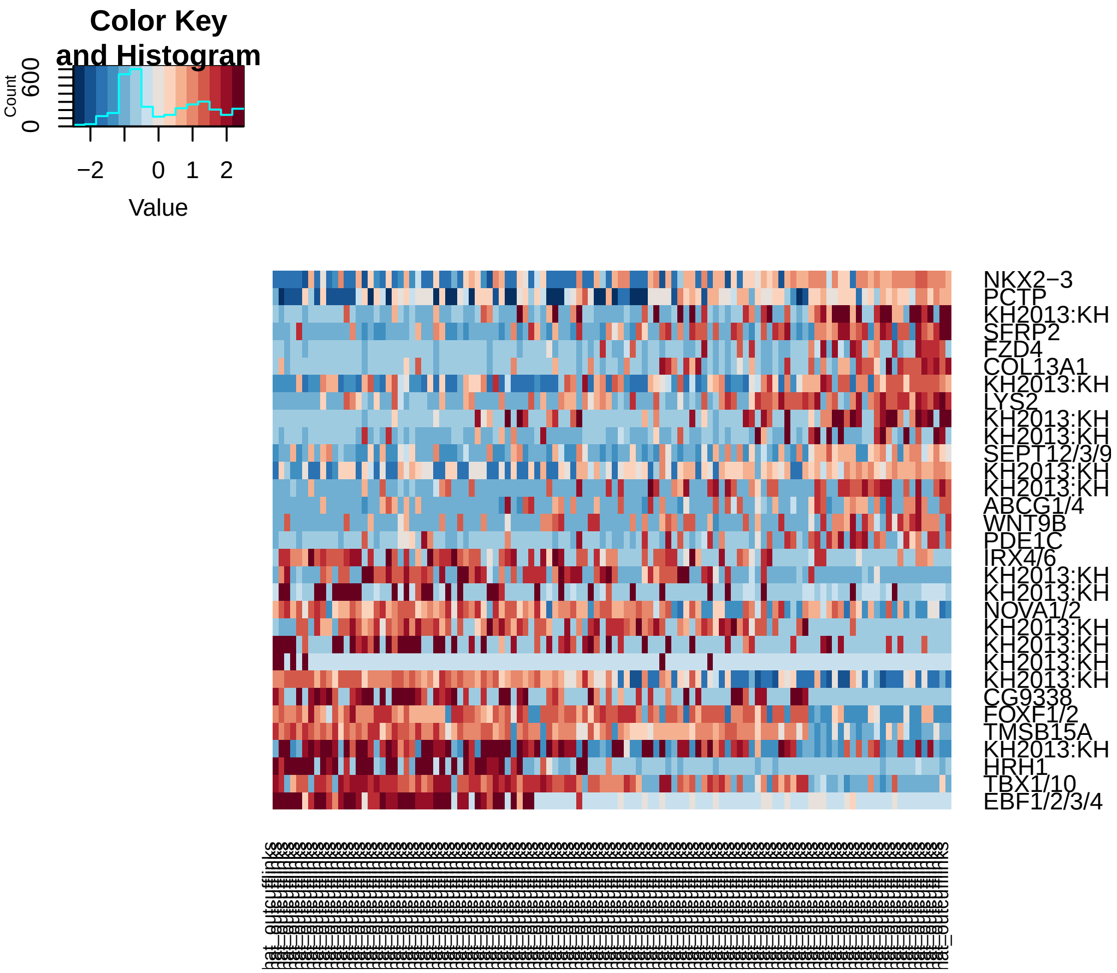


```r
pcHeatmap(hpf16, 2, use.full = T, do.balanced = T, col.use = col)  #technical
```

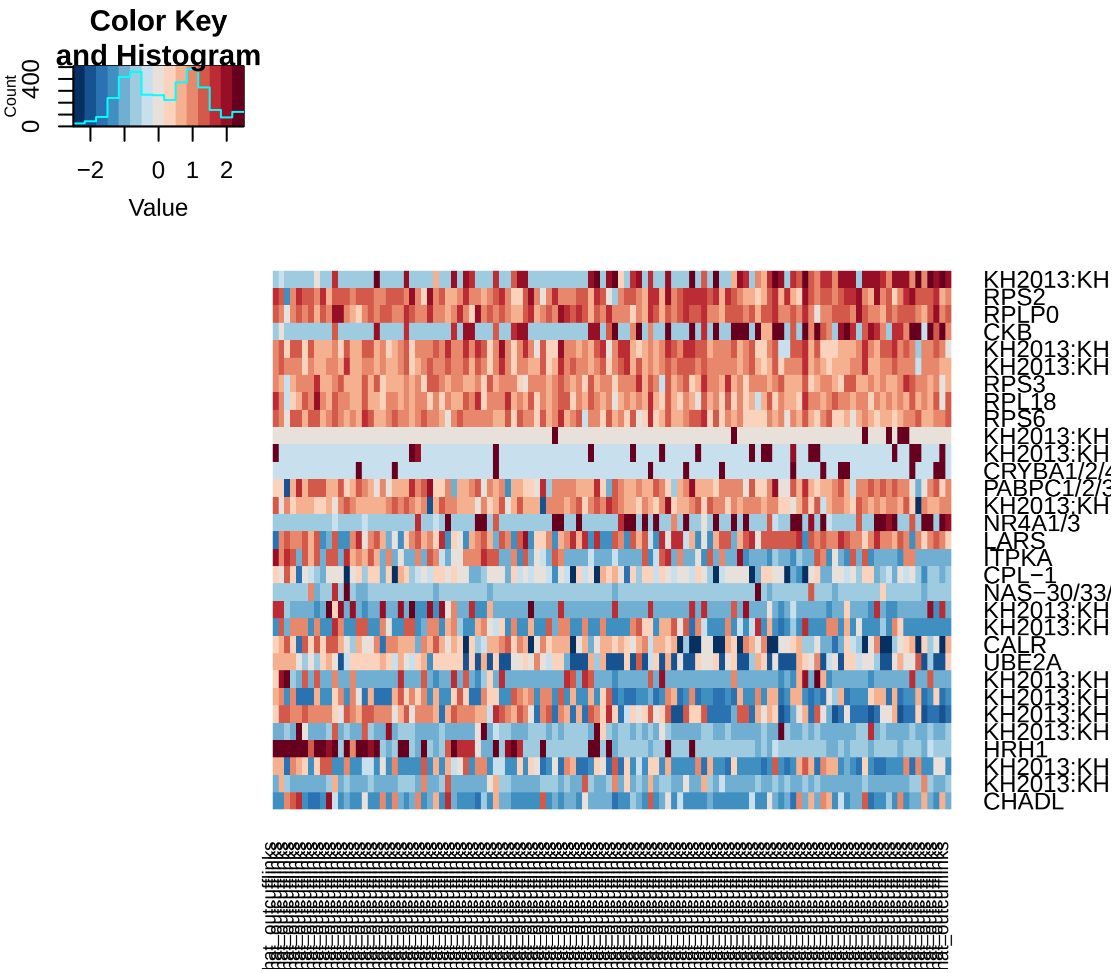


```
pcHeatmap(hpf16, 3, use.full = T, do.balanced = T, col.use = col)
```

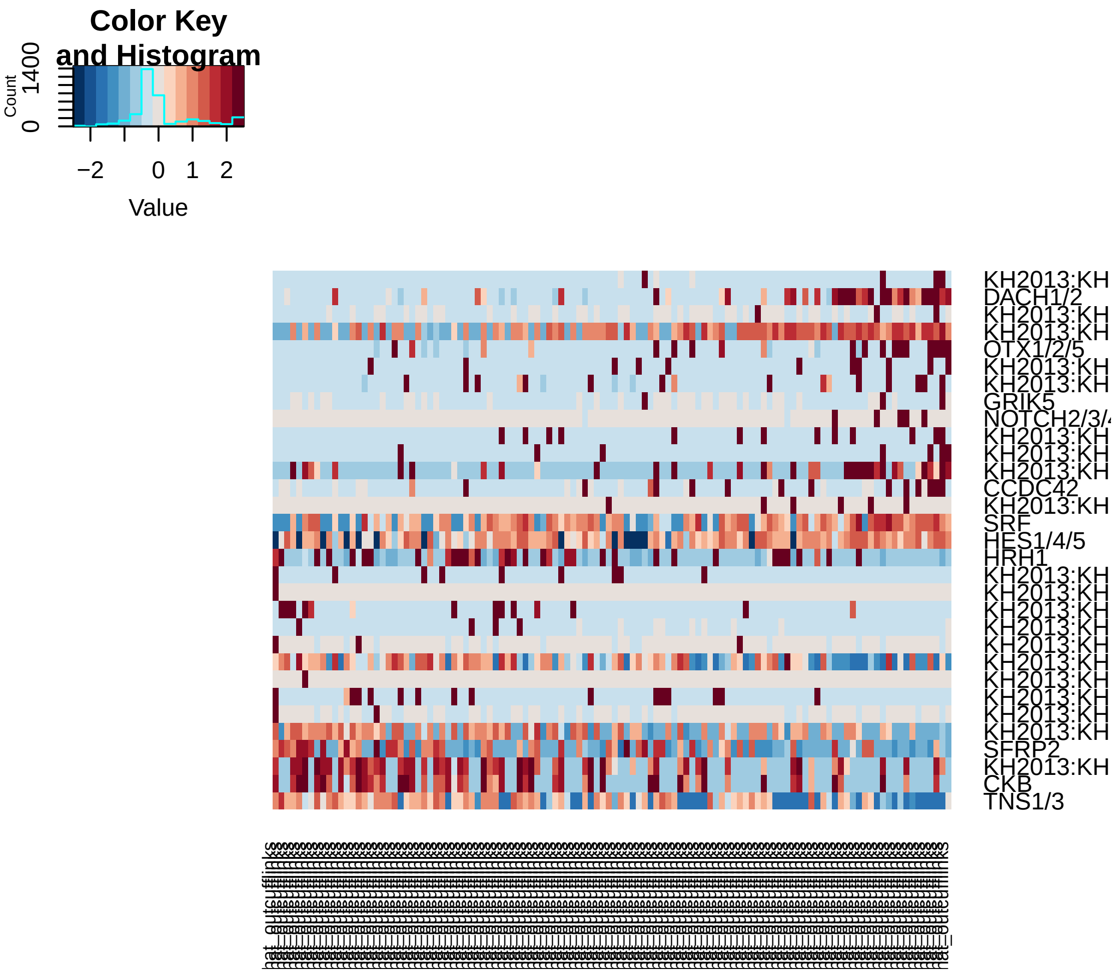


```r
# Do 200 random samplings to find significant genes, each time randomly
# permute 1% of genes This returns a 'p-value' for each gene in each PC,
# based on how likely the gene/PC score woud have been observed by chance
# Note that in this case we get the same result with 200 or 1000 samplings,
# so we do 200 here for expediency. Due to a lightly lower PCA input,
# permutation frequency is set to 0.02.
hpf16 = jackStraw(hpf16, num.replicate = 200, do.print = F)

# The jackStraw plot compares the distribution of P-values for each PC with
# a uniform distribution (dashed line) 'Significant' PCs will have a strong
# enrichment of genes with low p-values (solid curve above dashed line)
jackStrawPlot.new(hpf16, PCs = 1:12)
```

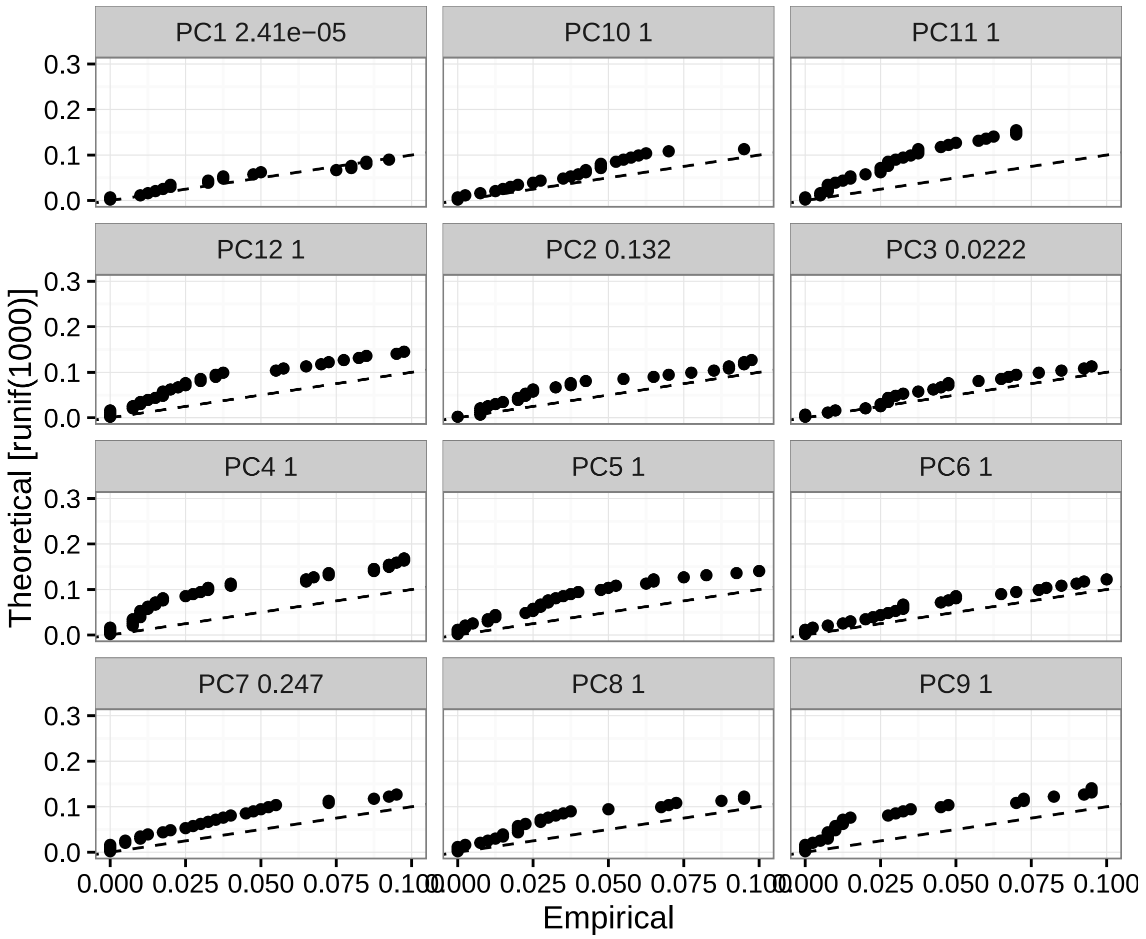


```
# In this case PC1 and PC3 are significant

# Run tSNE using significant PCs as input (spectral tSNE), we get distinct
# point clouds
hpf16 = run_tsne(hpf16, max_iter = 2000, dims.use = c(1, 3))
tsne.plot(hpf16, do.label = T, label.pt.size = 1)
```

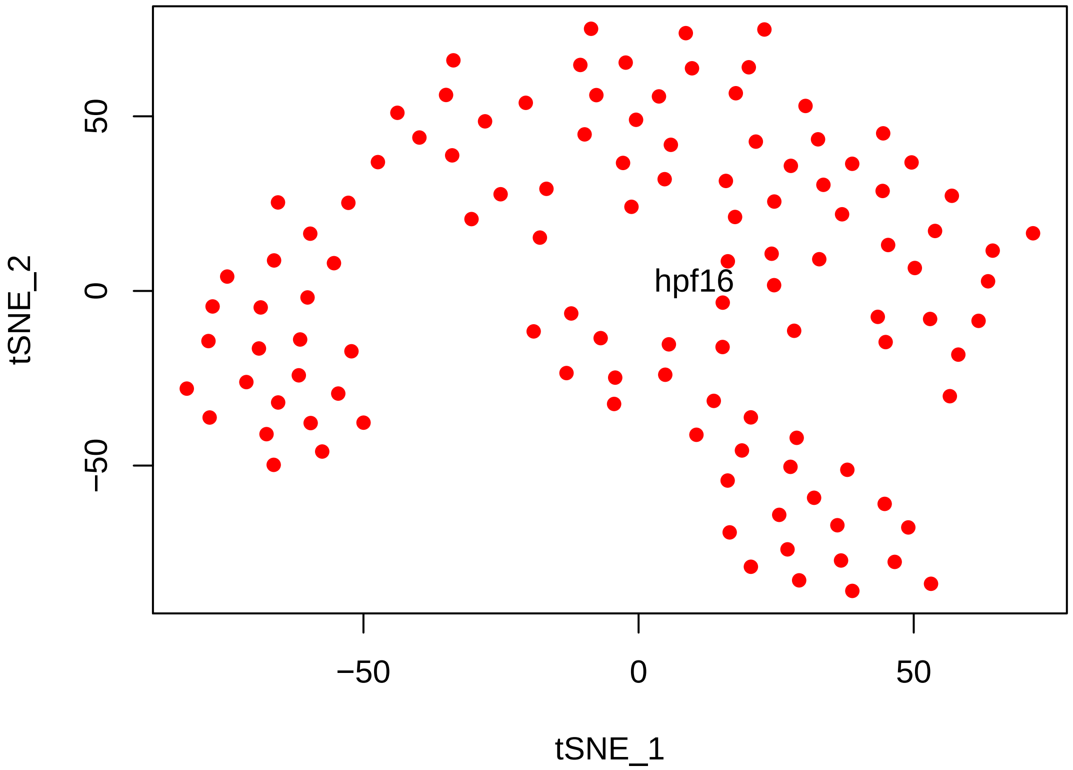


```
# Find cell clusters using Modularity optimization cluster detection.  hpf16
# dataset contain fewer cells than the others a k.param=10.
hpf16 = FindClusters(hpf16, pc.use = c(1, 3), do.modularity = T, resolution = 1,
    prune.SNN = 0.1, print.output = 0, k.param = 20, k.scale = floor(114/20))
```

```
## [1] "SNN : processed 28 cells"
## [1] "SNN : processed 57 cells"
## [1] "SNN : processed 86 cells"
## [1] "SNN : processed 114 cells"
```

```
tsne.plot(hpf16, do.label = T, label.pt.size = 1)
```

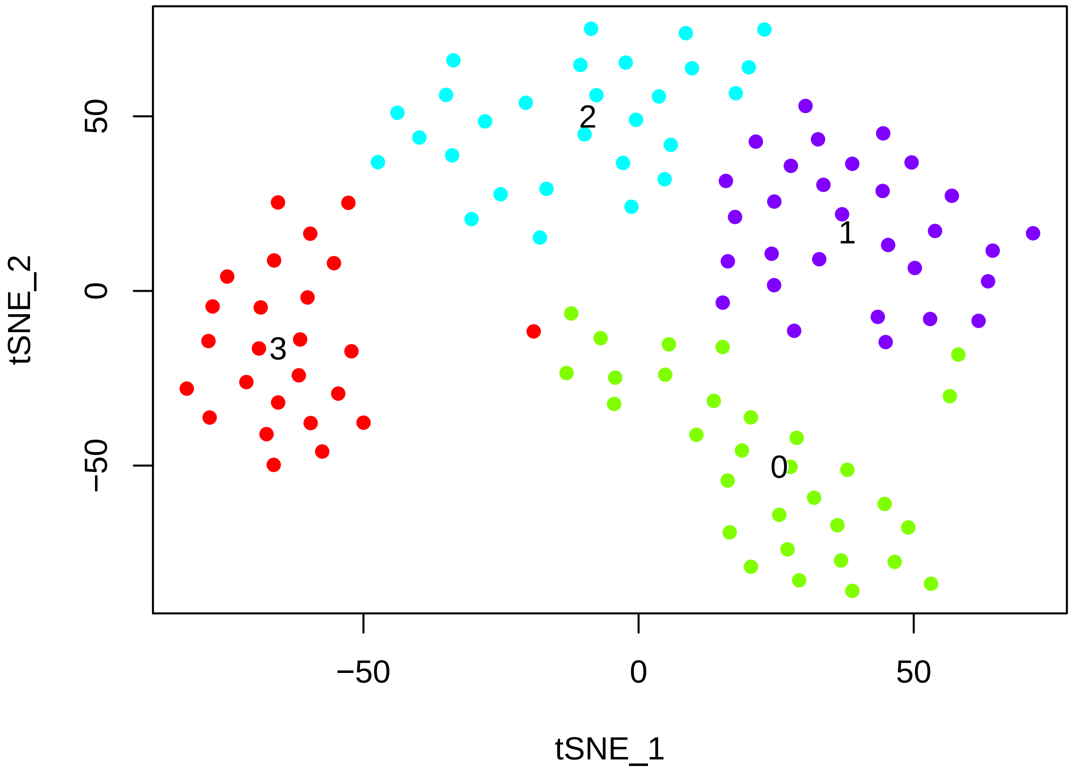


```
# The validity of the clusters can be validated using a classification
# scheme based on linear SVMs.
hpf16 = BuildSNN(hpf16, pc.use = c(1, 3), do.sparse = T, k.param = 20, k.scale = floor(1
```

```
## [1] "SNN : processed 28 cells"
## [1] "SNN : processed 57 cells"
## [1] "SNN : processed 86 cells"
## [1] "SNN : processed 114 cells"
```

```
hpf16 = ValidateClusters(hpf16, pc.use = c(1, 3), min.connectivity = 0.001,
    acc.cutoff = 0.85)
```

```
## [1] "  0% complete --- merge clusters 1 and 2, classification accuracy of 0.7733"
## [1] "100% complete --- started with 4 clusters, 3 clusters remaining"
```

```
tsne.plot(hpf16, do.label = T, label.pt.size = 1)
```

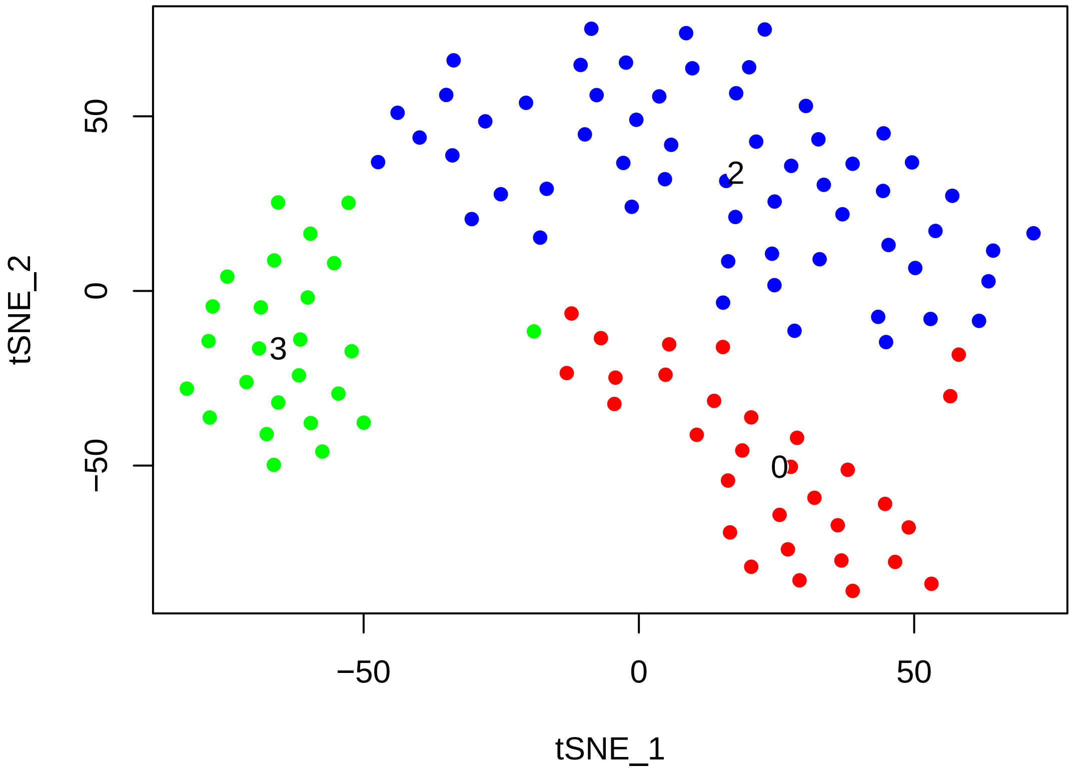


```r
# Find cluster markersusing ROC test with thresh.use = 1, min.pct = 0.5 The
# ROC test returns the 'classification power' for any individual marker
# (ranging from 0 - random, to 1 - perfect). Though not a statistical test,
# it is often very useful for finding clean markers.  Find markers for
# cluster 0
cl0_16.markers = find.markers(hpf16, 0, thresh.use = 1, test.use = "roc", min.pct = 0.5)
head(cl0_16.markers[order(cl0_16.markers$myAUC, decreasing = T), ], 20)
```

```
##                              myAUC  avg_diff power pct.1 pct.2
## KH2013:KH.C1.638_C17ORF105   0.812  1.121174 0.624 0.935 0.578
## DACH1/2                      0.803  2.475778 0.606 0.710 0.133
## KH2013:KH.C9.692_GABRR1/2/3  0.740  1.120309 0.480 0.839 0.518
## KH2013:KH.C3.616_GNRHR/1/2/4 0.722  1.529193 0.444 0.613 0.193
## SFR1                         0.704  1.048728 0.408 0.710 0.434
## ORC1                         0.701  1.215804 0.402 0.645 0.289
## GM1840                       0.698  1.469439 0.396 0.613 0.265
## KH2013:KH.C10.172            0.691  1.726626 0.382 0.548 0.193
## ZGC:174877                   0.685  1.114684 0.370 0.645 0.349
## IKBKG                        0.684  1.433664 0.368 0.581 0.301
## KH2013:KH.L150.4             0.661  1.021943 0.322 0.581 0.349
## PPT2                         0.654  1.024035 0.308 0.613 0.349
## USP48                        0.649  1.120253 0.298 0.581 0.458
## KH2013:KH.C6.155_FLRT1/2/3   0.621  1.114470 0.242 0.581 0.410
```

```
## RBM15B                        0.581 -1.461119 0.162 0.774 0.590
## KH2013:KH.L124.2              0.526  1.806418 0.052 0.806 0.916
## ZNRD1                         0.507 -3.447701 0.014 0.516 0.494
## KH2013:KH.L39.6               0.473 -4.277293 0.054 0.452 0.542
## SMEK1/2                       0.461 -1.636758 0.078 0.613 0.602
## KH2013:KH.C8.649_ACTA1/2      0.390 -2.190225 0.220 0.355 0.590
```

```r
# Visualize known markers with a violin plot
vlnPlot(hpf16, c("NKX2-3", "EBF1/2/3/4", "TBX1/10", "DACH1/2"))
```

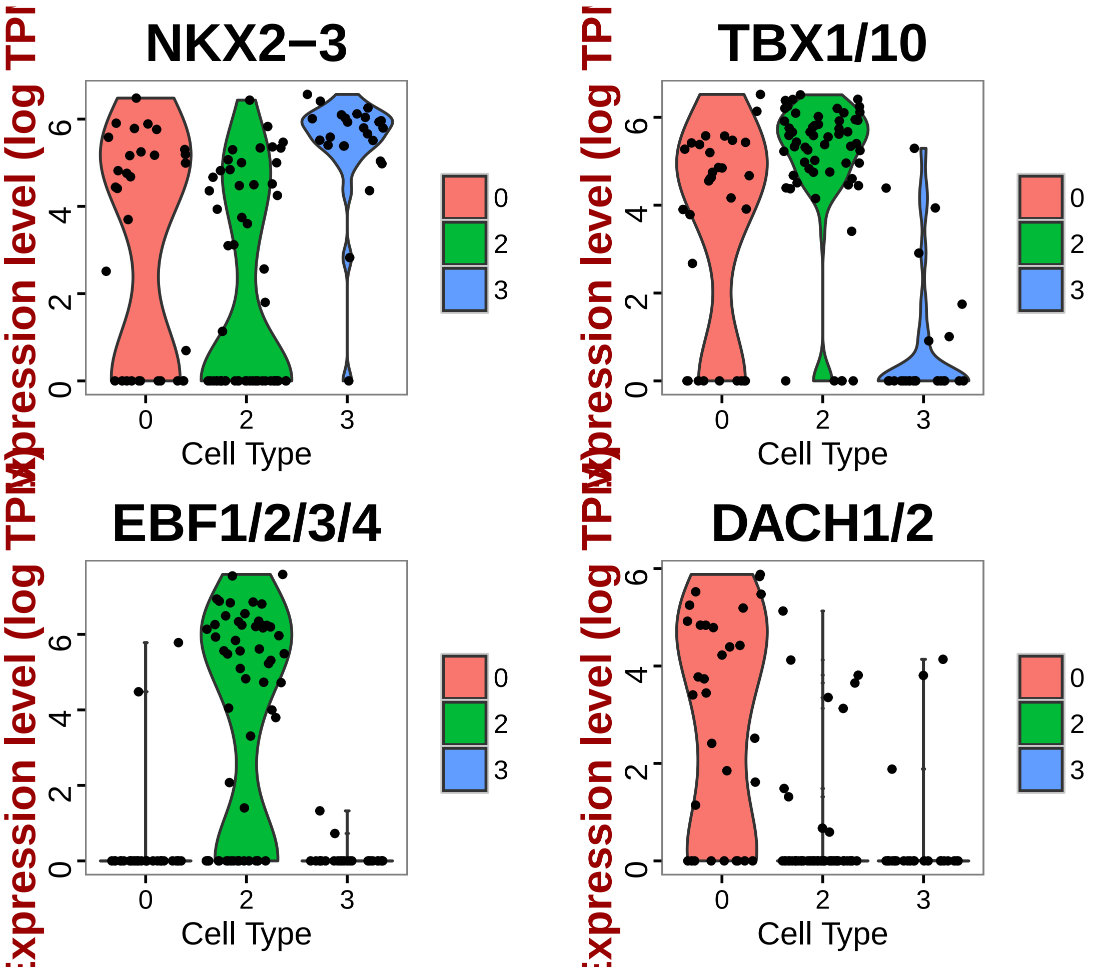


```r
# Based on preliminary studies these NKX2-3+ EBF1/2/3/4- DACH1/2+ TBX1/10+
# cells are SHPs

# Find markers for cluster 2
cl2_16.markers = find.markers(hpf16, 2, thresh.use = 1, test.use = "roc", min.pct = 0.5)
head(cl2_16.markers[order(cl2_16.markers$myAUC, decreasing = T), ], 20)
```

```
##                              myAUC avg_diff power pct.1 pct.2
```

```
## KH2013:KH.S555.1_HTR7  0.939 2.307248 0.878 0.983 0.464
## KH2013:KH.C1.21_CG2781 0.852 1.103323 0.704 1.000 0.946
## TBX1/10                0.820 1.194245 0.640 0.914 0.500
## EBF1/2/3/4             0.817 3.644445 0.634 0.672 0.054
## CDC6                   0.816 1.009440 0.632 1.000 0.929
## HAND/2                 0.810 1.079622 0.620 0.966 0.786
## ZCCHC24                0.797 1.156534 0.594 0.931 0.536
## HRH1                   0.796 3.209113 0.592 0.603 0.018
## SMURF1/2               0.787 1.502168 0.574 0.828 0.536
## TMSB15A                0.784 1.133905 0.568 0.966 0.679
## ELK1/3/4               0.749 1.134942 0.498 0.741 0.286
## IRX4/6                 0.735 1.253873 0.470 0.759 0.357
## KH2013:KH.C4.404       0.733 1.737132 0.466 0.690 0.304
## KH2013:KH.L134.29      0.724 1.222385 0.448 0.741 0.446
## IRX1/2/3/5             0.705 1.109414 0.410 0.569 0.214
## NOX5                   0.696 1.016322 0.392 0.638 0.286
## KIAA0513               0.693 1.031799 0.386 0.759 0.500
## KH2013:KH.C9.223_CDC42 0.689 1.056894 0.378 0.914 0.821
## ITPKA                  0.666 1.211642 0.332 0.586 0.339
## KH2013:KH.C12.239      0.647 1.110297 0.294 0.534 0.304
```

```r
# Visualize known markers with a violin plot
vlnPlot(hpf16, c("NKX2-3", "EBF1/2/3/4", "DACH1/2", "TBX1/10"))
```

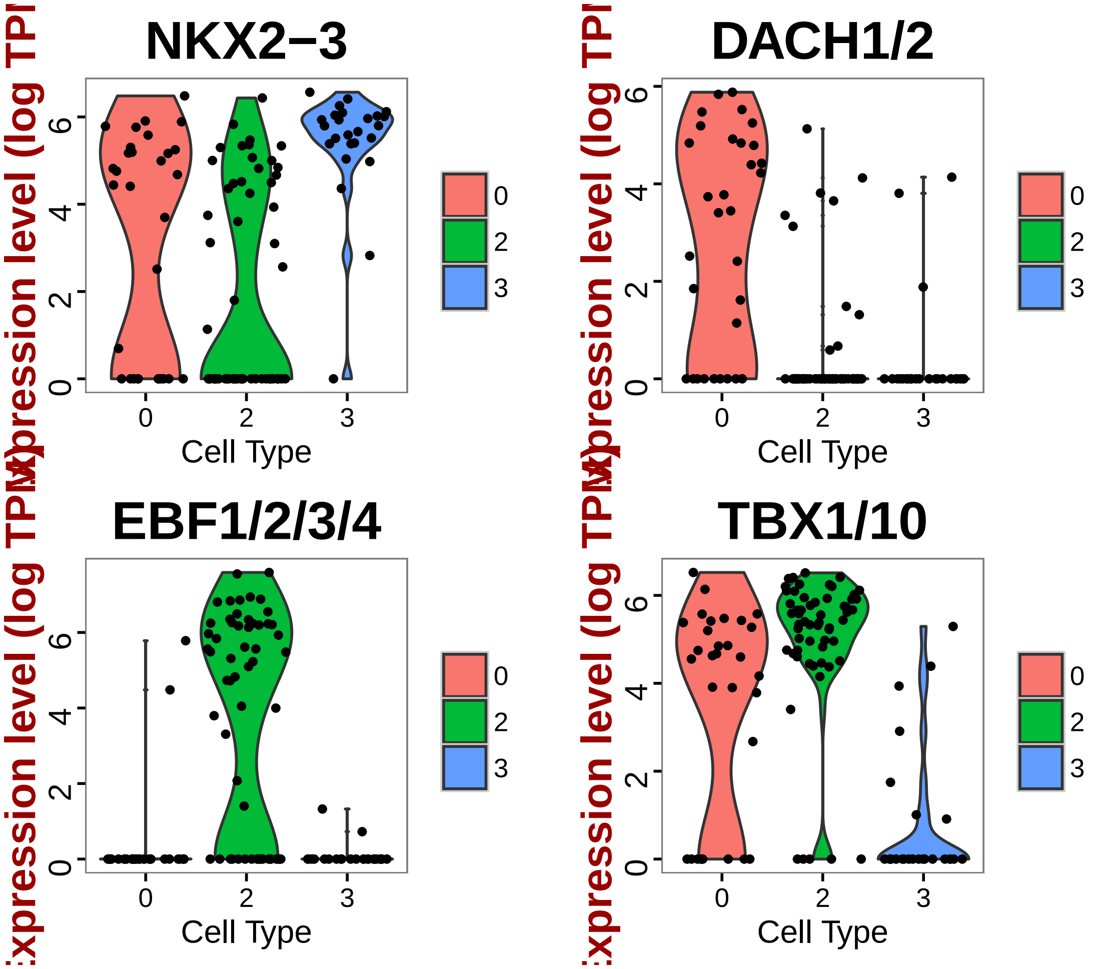


```r
# Based on preliminary studies these NKX2-3- EBF1/2/3/4+ DACH1/2+ TBX1/10+
# cells are ASM

# Find markers for cluster 3
cl3_16.markers = find.markers(hpf16, 3, thresh.use = 1, test.use = "roc", min.pct = 0.5)
head(cl3_16.markers[order(cl3_16.markers$myAUC, decreasing = T), ], 20)
```

```
##                                                                 myAUC
## KH2013:KH.C5.227_F56C4.4                                        0.950
## SMTN                                                            0.931
## MOGAT1/2                                                        0.919
## NKX2-3                                                          0.879
## KH2013:KH.C9.40_GSTO1/2                                         0.873
## SI:DKEY-79F11.5                                                 0.852
## KH2013:KH.C9.770_RGR                                            0.851
## SCYL                                                            0.831
## KH2013:KH.C8.489_SI:DKEY-29D8.3                                 0.830
## KH2013:KH.C4.547_BMP2/4                                         0.829
```

```
## KH2013:KH.C9.247_BX119910.1/10/11/12/13/14/16/2/3/4/5/6/7/8/9 0.828
## CDH-4                                                          0.827
## SFRP1/5                                                        0.826
## TIMP4                                                          0.820
## KH2013:KH.C2.22                                                0.818
## SFRP2                                                          0.818
## SLIT1/2/3                                                      0.816
## LYS2                                                           0.816
## KH2013:KH.C4.376_UNC93A                                        0.806
## COL13A1                                                        0.792
##                                                                avg_diff
## KH2013:KH.C5.227_F56C4.4                                       1.269902
## SMTN                                                           1.785546
## MOGAT1/2                                                       1.568086
## NKX2-3                                                         1.329046
## KH2013:KH.C9.40_GSTO1/2                                        1.365326
## SI:DKEY-79F11.5                                                1.537301
## KH2013:KH.C9.770_RGR                                           1.117639
## SCYL                                                           1.030414
## KH2013:KH.C8.489_SI:DKEY-29D8.3                                1.243672
## KH2013:KH.C4.547_BMP2/4                                        1.630799
## KH2013:KH.C9.247_BX119910.1/10/11/12/13/14/16/2/3/4/5/6/7/8/9 1.348407
## CDH-4                                                          1.034349
## SFRP1/5                                                        1.934746
## TIMP4                                                          1.211017
## KH2013:KH.C2.22                                                1.290851
## SFRP2                                                          1.820400
## SLIT1/2/3                                                      1.306153
## LYS2                                                           1.452199
## KH2013:KH.C4.376_UNC93A                                        1.210027
## COL13A1                                                        2.009872
##                                                                power pct.1
## KH2013:KH.C5.227_F56C4.4                                       0.900  1.00
## SMTN                                                           0.862  1.00
## MOGAT1/2                                                       0.838  1.00
## NKX2-3                                                         0.758  0.96
## KH2013:KH.C9.40_GSTO1/2                                        0.746  1.00
## SI:DKEY-79F11.5                                                0.704  0.92
## KH2013:KH.C9.770_RGR                                           0.702  1.00
## SCYL                                                           0.662  1.00
## KH2013:KH.C8.489_SI:DKEY-29D8.3                                0.660  0.96
## KH2013:KH.C4.547_BMP2/4                                        0.658  0.88
## KH2013:KH.C9.247_BX119910.1/10/11/12/13/14/16/2/3/4/5/6/7/8/9 0.656  0.92
## CDH-4                                                          0.654  1.00
## SFRP1/5                                                        0.652  0.80
```

```
## TIMP4                                                    0.640  0.92
## KH2013:KH.C2.22                                          0.636  0.92
## SFRP2                                                    0.636  0.84
## SLIT1/2/3                                                0.632  0.96
## LYS2                                                     0.632  0.84
## KH2013:KH.C4.376_UNC93A                                  0.612  0.72
## COL13A1                                                  0.584  0.72
##                                                          pct.2
## KH2013:KH.C5.227_F56C4.4                                 1.000
## SMTN                                                     0.438
## MOGAT1/2                                                 0.618
## NKX2-3                                                   0.506
## KH2013:KH.C9.40_GSTO1/2                                  0.652
## SI:DKEY-79F11.5                                          0.685
## KH2013:KH.C9.770_RGR                                     0.708
## SCYL                                                     0.742
## KH2013:KH.C8.489_SI:DKEY-29D8.3                          0.742
## KH2013:KH.C4.547_BMP2/4                                  0.337
## KH2013:KH.C9.247_BX119910.1/10/11/12/13/14/16/2/3/4/5/6/7/8/9 0.494
## CDH-4                                                    0.944
## SFRP1/5                                                  0.191
## TIMP4                                                    0.438
## KH2013:KH.C2.22                                          0.438
## SFRP2                                                    0.315
## SLIT1/2/3                                                0.584
## LYS2                                                     0.404
## KH2013:KH.C4.376_UNC93A                                  0.236
## COL13A1                                                  0.180
```

```r
# Visualize known markers with a violin plot
vlnPlot(hpf16, c("NKX2-3", "EBF1/2/3/4", "TBX1/10", "DACH1/2"))
```

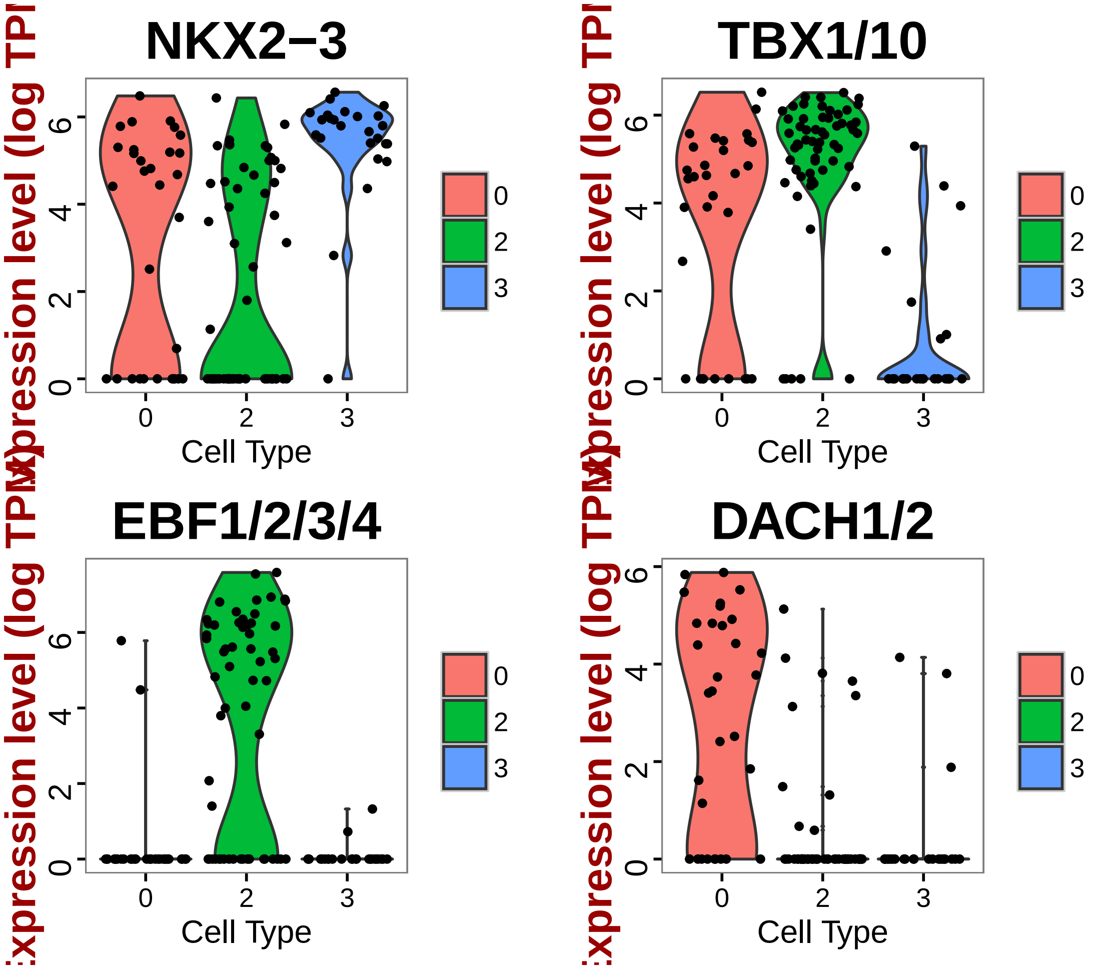


```
# Based known markers, this group is FHP

# Write cell names into text files
write.table(which.cells(hpf16, 0), file = "16SHPCells.txt", sep = "\t")
write.table(which.cells(hpf16, 2), file = "16ASMCells.txt", sep = "\t")
write.table(which.cells(hpf16, 3), file = "16FHPCells.txt", sep = "\t")

# Rename cluster identities
hpf16 = rename.ident(hpf16, 0, "16SHP")
hpf16 = rename.ident(hpf16, 2, "16ASM")
hpf16 = rename.ident(hpf16, 3, "16FHP")

# Visualize tSNE used color scheme
tsne.plot(hpf16, do.label = T, label.pt.size = 1, label.cex.text = 1.2, label.cols.use =
    "red", "blue"))
```

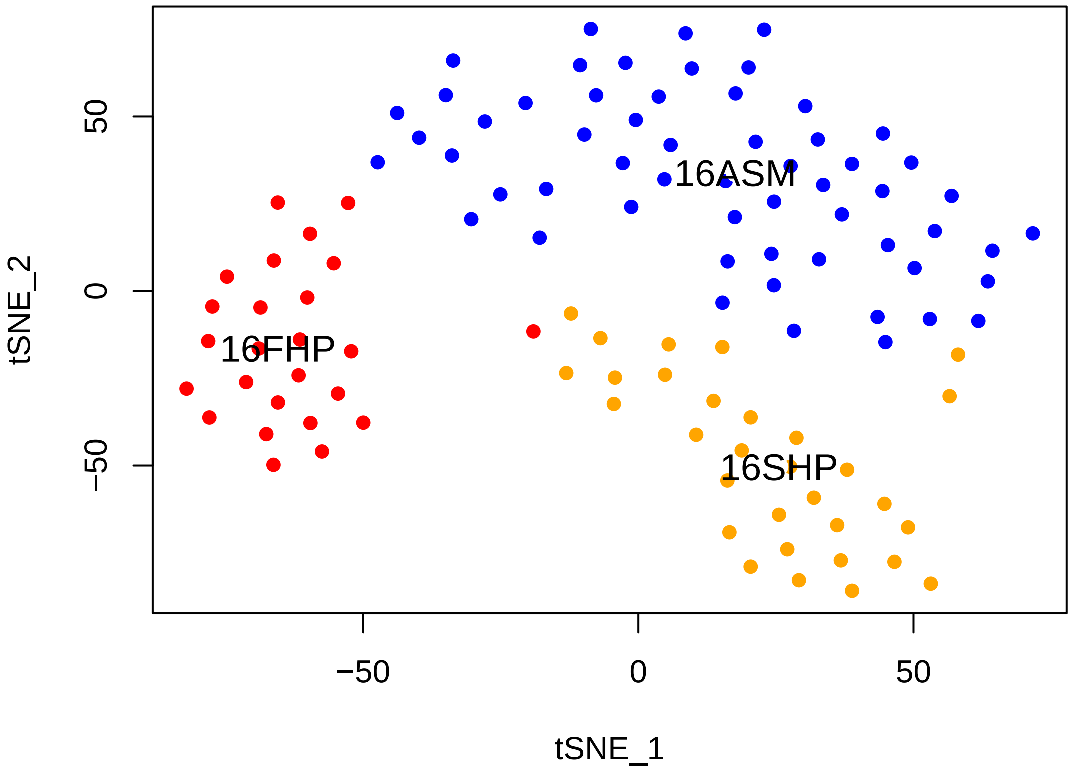


```r
# Visualize markers of different clusters using violin plot and feature plot
hpf16@ident = factor(hpf16@ident, ordered = T, levels = c("16FHP", "16SHP",
    "16ASM"))
genes.viz.16 = c("DACH1/2", "TBX1/10", "NKX2-3", "EBF1/2/3/4")
feature.plot(hpf16, genes.viz.16, pt.size = 0.8)
```

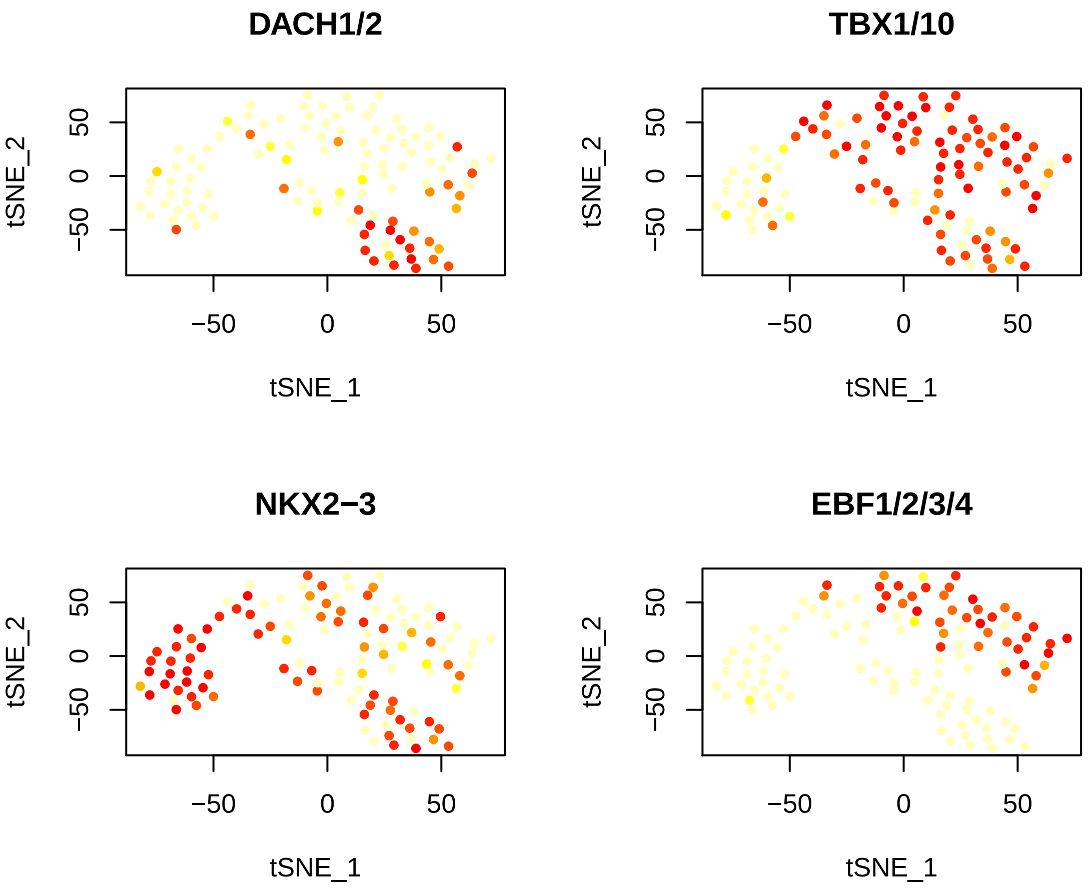


```
vlnPlot(hpf16, genes.viz.16, cols.use = c("red", "orange", "blue"))
```

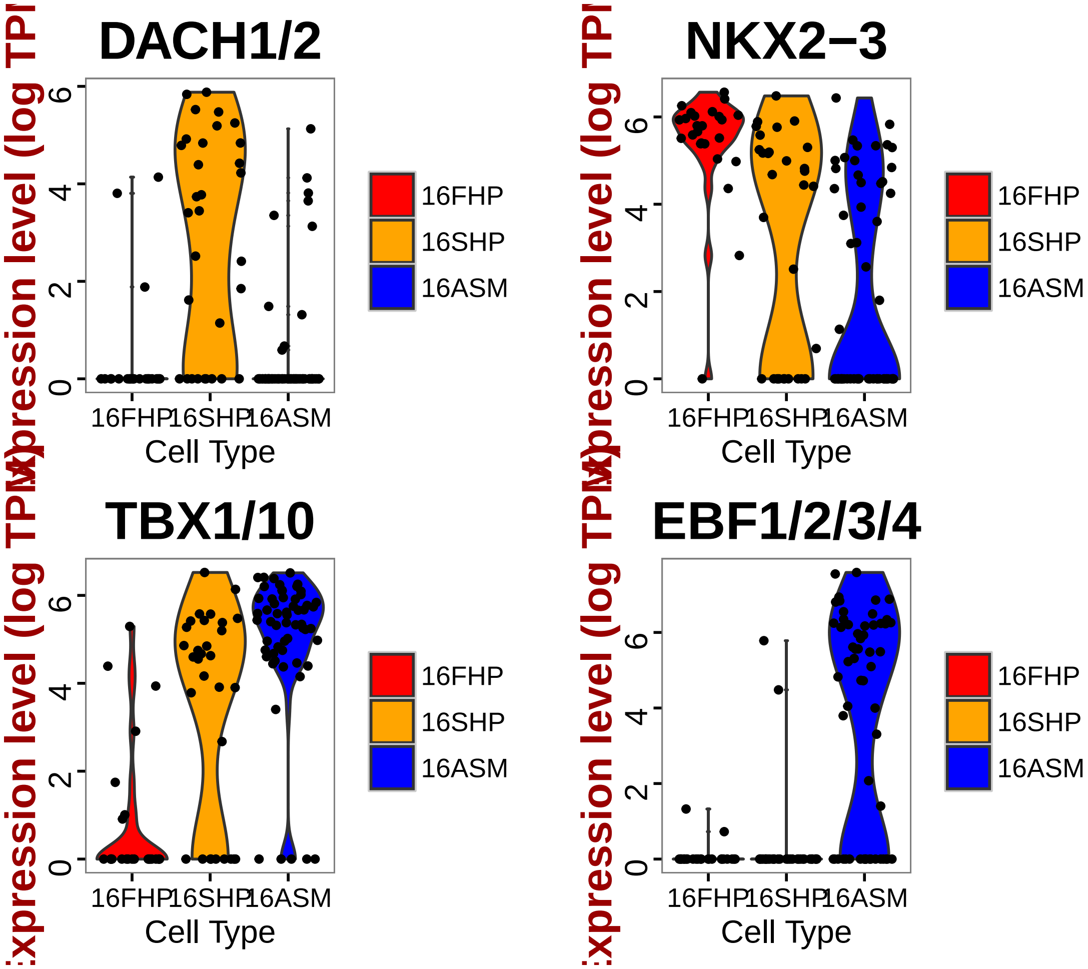


```r
# Store FHP markers in text file
FHP_16.markers = cl3_16.markers
head(FHP_16.markers[order(FHP_16.markers$myAUC, decreasing = T), ], 20)
```

```
##                                                                  myAUC
## KH2013:KH.C5.227_F56C4.4                                          0.950
## SMTN                                                              0.931
## MOGAT1/2                                                          0.919
## NKX2-3                                                            0.879
## KH2013:KH.C9.40_GSTO1/2                                           0.873
## SI:DKEY-79F11.5                                                   0.852
## KH2013:KH.C9.770_RGR                                              0.851
## SCYL                                                              0.831
## KH2013:KH.C8.489_SI:DKEY-29D8.3                                   0.830
## KH2013:KH.C4.547_BMP2/4                                           0.829
## KH2013:KH.C9.247_BX119910.1/10/11/12/13/14/16/2/3/4/5/6/7/8/9 0.828
## CDH-4                                                             0.827
## SFRP1/5                                                           0.826
```

```
## TIMP4                                                          0.820
## KH2013:KH.C2.22                                                0.818
## SFRP2                                                          0.818
## SLIT1/2/3                                                      0.816
## LYS2                                                           0.816
## KH2013:KH.C4.376_UNC93A                                        0.806
## COL13A1                                                        0.792
##                                                                avg_diff
## KH2013:KH.C5.227_F56C4.4                                       1.269902
## SMTN                                                           1.785546
## MOGAT1/2                                                       1.568086
## NKX2-3                                                         1.329046
## KH2013:KH.C9.40_GSTO1/2                                        1.365326
## SI:DKEY-79F11.5                                                1.537301
## KH2013:KH.C9.770_RGR                                           1.117639
## SCYL                                                           1.030414
## KH2013:KH.C8.489_SI:DKEY-29D8.3                                1.243672
## KH2013:KH.C4.547_BMP2/4                                        1.630799
## KH2013:KH.C9.247_BX119910.1/10/11/12/13/14/16/2/3/4/5/6/7/8/9 1.348407
## CDH-4                                                          1.034349
## SFRP1/5                                                        1.934746
## TIMP4                                                          1.211017
## KH2013:KH.C2.22                                                1.290851
## SFRP2                                                          1.820400
## SLIT1/2/3                                                      1.306153
## LYS2                                                           1.452199
## KH2013:KH.C4.376_UNC93A                                        1.210027
## COL13A1                                                        2.009872
##                                                                power pct.1
## KH2013:KH.C5.227_F56C4.4                                       0.900  1.00
## SMTN                                                           0.862  1.00
## MOGAT1/2                                                       0.838  1.00
## NKX2-3                                                         0.758  0.96
## KH2013:KH.C9.40_GSTO1/2                                        0.746  1.00
## SI:DKEY-79F11.5                                                0.704  0.92
## KH2013:KH.C9.770_RGR                                           0.702  1.00
## SCYL                                                           0.662  1.00
## KH2013:KH.C8.489_SI:DKEY-29D8.3                                0.660  0.96
## KH2013:KH.C4.547_BMP2/4                                        0.658  0.88
## KH2013:KH.C9.247_BX119910.1/10/11/12/13/14/16/2/3/4/5/6/7/8/9 0.656  0.92
## CDH-4                                                          0.654  1.00
## SFRP1/5                                                        0.652  0.80
## TIMP4                                                          0.640  0.92
## KH2013:KH.C2.22                                                0.636  0.92
## SFRP2                                                          0.636  0.84
```

```
## SLIT1/2/3                                                     0.632  0.96
## LYS2                                                          0.632  0.84
## KH2013:KH.C4.376_UNC93A                                       0.612  0.72
## COL13A1                                                       0.584  0.72
##                                                               pct.2
## KH2013:KH.C5.227_F56C4.4                                      1.000
## SMTN                                                          0.438
## MOGAT1/2                                                      0.618
## NKX2-3                                                        0.506
## KH2013:KH.C9.40_GSTO1/2                                       0.652
## SI:DKEY-79F11.5                                               0.685
## KH2013:KH.C9.770_RGR                                          0.708
## SCYL                                                          0.742
## KH2013:KH.C8.489_SI:DKEY-29D8.3                               0.742
## KH2013:KH.C4.547_BMP2/4                                       0.337
## KH2013:KH.C9.247_BX119910.1/10/11/12/13/14/16/2/3/4/5/6/7/8/9 0.494
## CDH-4                                                         0.944
## SFRP1/5                                                       0.191
## TIMP4                                                         0.438
## KH2013:KH.C2.22                                               0.438
## SFRP2                                                         0.315
## SLIT1/2/3                                                     0.584
## LYS2                                                          0.404
## KH2013:KH.C4.376_UNC93A                                       0.236
## COL13A1                                                       0.180
```

```r
write.table(FHP_16.markers, file = "FHP_16.markers.txt", sep = "\t")

# Store SHP markers in text file
SHP_16.markers = cl0_16.markers
head(SHP_16.markers[order(SHP_16.markers$myAUC, decreasing = T), ], 20)
```

```
##                             myAUC  avg_diff power pct.1 pct.2
## KH2013:KH.C1.638_C17ORF105   0.812  1.121174 0.624 0.935 0.578
## DACH1/2                      0.803  2.475778 0.606 0.710 0.133
## KH2013:KH.C9.692_GABRR1/2/3  0.740  1.120309 0.480 0.839 0.518
## KH2013:KH.C3.616_GNRHR/1/2/4 0.722  1.529193 0.444 0.613 0.193
## SFR1                         0.704  1.048728 0.408 0.710 0.434
## ORC1                         0.701  1.215804 0.402 0.645 0.289
## GM1840                       0.698  1.469439 0.396 0.613 0.265
## KH2013:KH.C10.172            0.691  1.726626 0.382 0.548 0.193
## ZGC:174877                   0.685  1.114684 0.370 0.645 0.349
## IKBKG                        0.684  1.433664 0.368 0.581 0.301
## KH2013:KH.L150.4             0.661  1.021943 0.322 0.581 0.349
```

```
## PPT2                          0.654  1.024035 0.308 0.613 0.349
## USP48                         0.649  1.120253 0.298 0.581 0.458
## KH2013:KH.C6.155_FLRT1/2/3    0.621  1.114470 0.242 0.581 0.410
## RBM15B                        0.581 -1.461119 0.162 0.774 0.590
## KH2013:KH.L124.2              0.526  1.806418 0.052 0.806 0.916
## ZNRD1                         0.507 -3.447701 0.014 0.516 0.494
## KH2013:KH.L39.6               0.473 -4.277293 0.054 0.452 0.542
## SMEK1/2                       0.461 -1.636758 0.078 0.613 0.602
## KH2013:KH.C8.649_ACTA1/2      0.390 -2.190225 0.220 0.355 0.590
```

```
write.table(SHP_16.markers, file = "SHP_16.markers.txt", sep = "\t")

# Store ASM markers in text file
ASM_16.markers = cl2_16.markers
head(ASM_16.markers[order(ASM_16.markers$myAUC, decreasing = T), ], 20)
```

```
##                         myAUC avg_diff power pct.1 pct.2
## KH2013:KH.S555.1_HTR7   0.939 2.307248 0.878 0.983 0.464
## KH2013:KH.C1.21_CG2781 0.852 1.103323 0.704 1.000 0.946
## TBX1/10                 0.820 1.194245 0.640 0.914 0.500
## EBF1/2/3/4              0.817 3.644445 0.634 0.672 0.054
## CDC6                    0.816 1.009440 0.632 1.000 0.929
## HAND/2                  0.810 1.079622 0.620 0.966 0.786
## ZCCHC24                 0.797 1.156534 0.594 0.931 0.536
## HRH1                    0.796 3.209113 0.592 0.603 0.018
## SMURF1/2                0.787 1.502168 0.574 0.828 0.536
## TMSB15A                 0.784 1.133905 0.568 0.966 0.679
## ELK1/3/4                0.749 1.134942 0.498 0.741 0.286
## IRX4/6                  0.735 1.253873 0.470 0.759 0.357
## KH2013:KH.C4.404        0.733 1.737132 0.466 0.690 0.304
## KH2013:KH.L134.29       0.724 1.222385 0.448 0.741 0.446
## IRX1/2/3/5              0.705 1.109414 0.410 0.569 0.214
## NOX5                    0.696 1.016322 0.392 0.638 0.286
## KIAA0513                0.693 1.031799 0.386 0.759 0.500
## KH2013:KH.C9.223_CDC42 0.689 1.056894 0.378 0.914 0.821
## ITPKA                   0.666 1.211642 0.332 0.586 0.339
## KH2013:KH.C12.239       0.647 1.110297 0.294 0.534 0.304
```

```
# Find HP markers (pan cardiac marker)
HP_16.markers = find.markers(hpf16, c("16SHP", "16FHP"), thresh.use = 1, test.use = "roc
    min.pct = 0.5)
head(HP_16.markers[order(HP_16.markers$myAUC, decreasing = T), ], 20)
```

```
##                                   myAUC avg_diff power pct.1 pct.2
```

```
## QKI                              0.845 1.109346 0.690 0.982 0.776
## SCYL                             0.833 1.217014 0.666 0.964 0.638
## SLIT1/2/3                        0.805 1.300099 0.610 0.893 0.448
## KH2013:KH.C8.489_SI:DKEY-29D8.3 0.789 1.261880 0.578 0.893 0.690
## HAND/1/2                         0.786 1.013146 0.572 0.893 0.586
## LRP4/8                           0.784 1.805954 0.568 0.696 0.224
## LYS2                             0.774 1.779484 0.548 0.714 0.293
## NKX2-3                           0.759 1.255571 0.518 0.768 0.448
## IGF1/2                           0.733 1.277706 0.466 0.786 0.500
## SI:DKEY-79F11.5                  0.715 1.158875 0.430 0.839 0.638
## KH2013:KH.C9.608_PDE9A           0.710 1.064685 0.420 0.643 0.276
## KH2013:KH.C3.716_EFNA1/2/3/4/5  0.706 1.252546 0.412 0.714 0.379
## TANC1/2                          0.704 1.290037 0.408 0.518 0.207
## KH2013:KH.L119.21_POLA1          0.686 1.000919 0.372 0.696 0.431
## SMTN                             0.685 1.142035 0.370 0.679 0.448
## ZGC:174877                       0.684 1.254427 0.368 0.589 0.276
## LMOD1/3                          0.683 1.012676 0.366 0.607 0.241
## USP48                            0.665 1.094911 0.330 0.607 0.379
## KH2013:KH.C13.152_F14B6.6        0.661 1.103362 0.322 0.571 0.328
## KH2013:KH.C3.52_EFNA1/2/3/4/5   0.661 1.102856 0.322 0.571 0.259
```

```r
write.table(HP_16.markers, file = "panHP_16.markers.txt", sep = "\t")

# Find SHP specific markers that distinguish two heart progenitors FHP and
# SHP
SHPspecific_16.markers = find.markers(hpf16, "16SHP", "16FHP", thresh.use = 1,
    test.use = "roc", min.pct = 0.5)
head(SHPspecific_16.markers[order(SHPspecific_16.markers$myAUC, decreasing = T),
    ], 20)
```

```
##                               myAUC avg_diff power pct.1 pct.2
## TMSB15A                       0.937 2.553345 0.874 0.968  0.32
## PCMTD1/2                      0.929 1.542177 0.858 1.000  0.88
## FOXF1/2                       0.926 2.612649 0.852 0.903  0.32
## KH2013:KH.C9.692_GABRR1/2/3   0.897 3.262190 0.794 0.839  0.08
## KH2013:KH.C1.638_C17ORF105    0.835 1.377923 0.670 0.935  0.40
## CHID1                         0.828 1.637985 0.656 0.806  0.28
## ZNF703                        0.813 2.656630 0.626 0.710  0.12
## KH2013:KH.S1660.1             0.812 1.157955 0.624 1.000  0.76
## DACH1/2                       0.810 2.692742 0.620 0.710  0.12
## E2F/1/2/3                     0.801 1.314878 0.602 0.903  0.60
## SFR1                          0.794 2.042680 0.588 0.710  0.28
## GDH1/3                        0.788 1.844341 0.576 0.742  0.36
## TBX1/10                       0.783 2.133826 0.566 0.710  0.24
```

```
## KH2013:KH.C4.230_TAGLN/2/3   0.775 1.300167 0.550 0.806  0.36
## histone                      0.772 1.009498 0.544 0.871  0.52
## PRMT7                        0.772 1.300034 0.544 0.839  0.48
## KH2013:KH.C3.616_GNRHR/1/2/4 0.743 1.895749 0.486 0.613  0.16
## ZK637.14                     0.743 1.290958 0.486 0.710  0.24
## ITPKA                        0.735 2.323179 0.470 0.548  0.08
## KH2013:KH.C5.329             0.734 1.284407 0.468 0.710  0.36
```

```
write.table(SHPspecific_16.markers, file = "SHPspecific_16.markers.txt", sep = "\t")

# Find FHP specific markers that distinguish two heart progenitors FHP and
# SHP
FHPspecific_16.markers = find.markers(hpf16, "16FHP", "16SHP", thresh.use = 1,
    test.use = "roc", min.pct = 0.5)
head(FHPspecific_16.markers[order(FHPspecific_16.markers$myAUC, decreasing = T),
    ], 20)
```

```
##                                                             myAUC
## MOGAT1/2                                                    0.954
## SMTN                                                        0.934
## KH2013:KH.C5.227_F56C4.4                                    0.932
## KH2013:KH.C9.770_RGR                                        0.883
## KH2013:KH.C9.40_GSTO1/2                                     0.858
## KH2013:KH.C7.205_ASB2                                       0.853
## KH2013:KH.C2.668_SLC38A9                                    0.850
## SFRP1/5                                                     0.843
## TIMP4                                                       0.830
## KH2013:KH.C4.457_UNC93A                                     0.827
## SI:DKEY-79F11.5                                             0.826
## CDH-4                                                       0.826
## SFRP2                                                       0.815
## KH2013:KH.C9.247_BX119910.1/10/11/12/13/14/16/2/3/4/5/6/7/8/9 0.815
## KH2013:KH.C4.547_BMP2/4                                     0.808
## KH2013:KH.C2.22                                             0.806
## COL13A1                                                     0.801
## SEPT12/3/9                                                  0.797
## KH2013:KH.C11.362                                           0.780
## TNS1/3                                                      0.772
##                                                             avg_diff
## MOGAT1/2                                                    1.931942
## SMTN                                                        1.823235
## KH2013:KH.C5.227_F56C4.4                                    1.122933
## KH2013:KH.C9.770_RGR                                        1.359233
## KH2013:KH.C9.40_GSTO1/2                                     1.277879
```

```
## KH2013:KH.C7.205_ASB2                                          1.045074
## KH2013:KH.C2.668_SLC38A9                                       2.120019
## SFRP1/5                                                        2.235711
## TIMP4                                                          1.119125
## KH2013:KH.C4.457_UNC93A                                        1.331659
## SI:DKEY-79F11.5                                                1.317059
## CDH-4                                                          1.039662
## SFRP2                                                          2.048199
## KH2013:KH.C9.247_BX119910.1/10/11/12/13/14/16/2/3/4/5/6/7/8/9 1.313579
## KH2013:KH.C4.547_BMP2/4                                        1.472345
## KH2013:KH.C2.22                                                1.218241
## COL13A1                                                        2.130965
## SEPT12/3/9                                                     1.331829
## KH2013:KH.C11.362                                              1.416327
## TNS1/3                                                         1.102989
##                                                                power pct.1
## MOGAT1/2                                                       0.908  1.00
## SMTN                                                           0.868  1.00
## KH2013:KH.C5.227_F56C4.4                                       0.864  1.00
## KH2013:KH.C9.770_RGR                                           0.766  1.00
## KH2013:KH.C9.40_GSTO1/2                                        0.716  1.00
## KH2013:KH.C7.205_ASB2                                          0.706  1.00
## KH2013:KH.C2.668_SLC38A9                                       0.700  0.88
## SFRP1/5                                                        0.686  0.80
## TIMP4                                                          0.660  0.92
## KH2013:KH.C4.457_UNC93A                                        0.654  0.72
## SI:DKEY-79F11.5                                                0.652  0.92
## CDH-4                                                          0.652  1.00
## SFRP2                                                          0.630  0.84
## KH2013:KH.C9.247_BX119910.1/10/11/12/13/14/16/2/3/4/5/6/7/8/9 0.630  0.92
## KH2013:KH.C4.547_BMP2/4                                        0.616  0.88
## KH2013:KH.C2.22                                                0.612  0.92
## COL13A1                                                        0.602  0.72
## SEPT12/3/9                                                     0.594  0.84
## KH2013:KH.C11.362                                              0.560  0.88
## TNS1/3                                                         0.544  0.88
##                                                                pct.2
## MOGAT1/2                                                       0.613
## SMTN                                                           0.419
## KH2013:KH.C5.227_F56C4.4                                       1.000
## KH2013:KH.C9.770_RGR                                           0.645
## KH2013:KH.C9.40_GSTO1/2                                        0.710
## KH2013:KH.C7.205_ASB2                                          0.871
## KH2013:KH.C2.668_SLC38A9                                       0.323
## SFRP1/5                                                        0.161
```

```
## TIMP4                                                                 0.355
## KH2013:KH.C4.457_UNC93A                                               0.161
## SI:DKEY-79F11.5                                                       0.774
## CDH-4                                                                 0.935
## SFRP2                                                                 0.355
## KH2013:KH.C9.247_BX119910.1/10/11/12/13/14/16/2/3/4/5/6/7/8/9 0.484
## KH2013:KH.C4.547_BMP2/4                                               0.323
## KH2013:KH.C2.22                                                       0.548
## COL13A1                                                               0.194
## SEPT12/3/9                                                            0.258
## KH2013:KH.C11.362                                                     0.452
## TNS1/3                                                                0.548
```

```
write.table(FHPspecific_16.markers, file = "FHPspecific_16.markers.txt", sep = "\t")

# Select markers for plotting on a Heatmap (top 10 positive markers with
# high discriminatory power) (5 markers are shown for STVC and SHP due to
# less markers)
marker16FHP = rownames(FHP_16.markers[order(FHP_16.markers$myAUC, decreasing = T)[1:15],
    ])
marker16SHP = rownames(SHP_16.markers[order(SHP_16.markers$myAUC, decreasing = T)[1:15],
    ])
marker16ASM = rownames(ASM_16.markers[order(ASM_16.markers$myAUC, decreasing = T)[1:15],
    ])
marker.16 = c(marker16FHP, marker16SHP, marker16ASM)

# Redefine ASM markers
ASM_16.markers = find.markers(hpf16, "16ASM", "16FHP", thresh.use = 1, test.use = "roc",
    min.pct = 0.5)
head(ASM_16.markers[order(ASM_16.markers$myAUC, decreasing = T), ], 20)
```

```
##                              myAUC avg_diff power pct.1 pct.2
## KH2013:KH.S555.1_HTR7        0.968 3.169129 0.936 0.983  0.40
## TMSB15A                      0.953 3.156753 0.906 0.966  0.32
## TBX1/10                      0.929 2.827890 0.858 0.914  0.24
## CDC6                         0.921 1.599529 0.842 1.000  0.84
## TP53I11                      0.890 1.224223 0.780 1.000  0.72
## FOXF1/2                      0.888 2.343817 0.776 0.897  0.32
## ZCCHC24                      0.881 1.848557 0.762 0.931  0.40
## SELT/2                       0.876 1.038634 0.752 0.983  0.96
## KH2013:KH.C1.21_CG2781       0.872 1.198799 0.744 1.000  0.92
## HAND/2                       0.859 1.436427 0.718 0.966  0.64
## E2F/1/2/3                    0.845 1.243666 0.690 0.914  0.60
## SLC16A12/7                   0.839 1.432206 0.678 0.931  0.64
```

```
## EBF1/2/3/4                   0.823 5.630515 0.646 0.672  0.04
## KH2013:KH.C9.692_GABRR1/2/3 0.818 2.464266 0.636 0.707  0.08
## KH2013:KH.C4.404             0.817 3.428063 0.634 0.690  0.08
## SMURF1/2                     0.817 1.927637 0.634 0.828  0.52
## KH2013:KH.L134.29            0.809 2.316999 0.618 0.741  0.28
## ELK1/3/4                     0.807 1.756193 0.614 0.741  0.16
## HRH1                         0.802 4.432899 0.604 0.603  0.00
## CG9338                       0.802 4.212724 0.604 0.603  0.00
```

```r
write.table(ASM_16.markers, file = "ASM_16.markers.txt", sep = "\t")

# Draw a heatmap of all cells for these marker genes
doHeatMap(hpf16, genes.use = marker.16, remove.key = TRUE, slim.col.label = T,
    cex.col = 1.2, col.use = col)
```

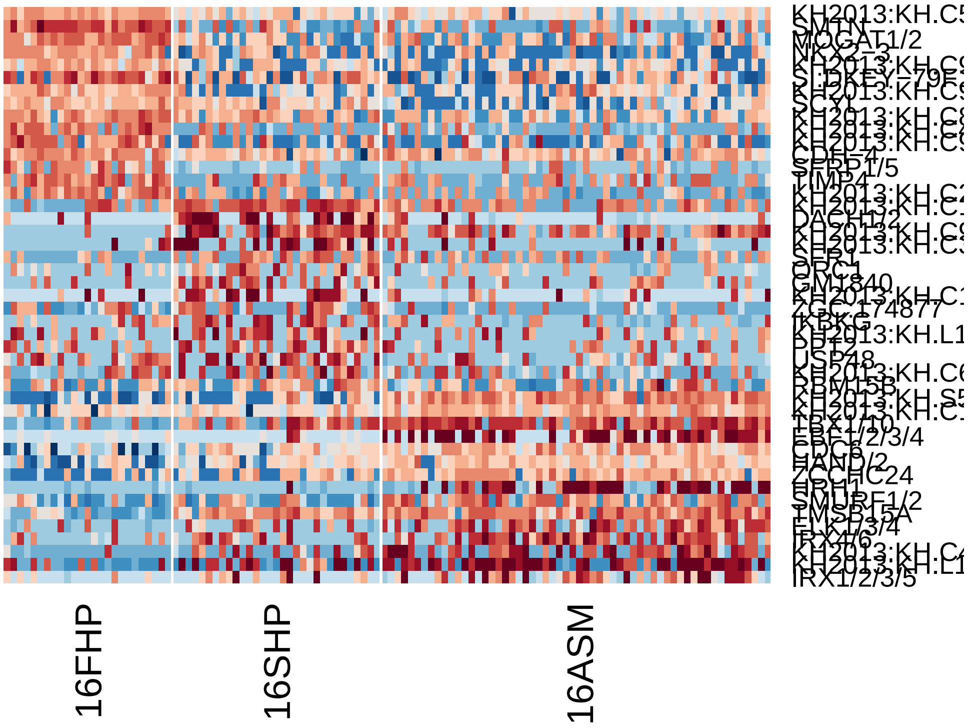


```r
# Store clustering information into a master Seurat object
hpfall.cluster = hpfall.remv2
hpfall.cluster = set.ident(hpfall.cluster, which.cells(hpfall.cluster, "hpf12"),
    hpf12@ident)
hpfall.cluster = set.ident(hpfall.cluster, which.cells(hpfall.cluster, "hpf14"),
    hpf14@ident)
hpfall.cluster = set.ident(hpfall.cluster, which.cells(hpfall.cluster, "hpf16"),
    hpf16@ident)
hpfall.cluster = set.ident(hpfall.cluster, which.cells(hpfall.cluster, "hpf18"),
```

```
    hpf18@ident)
hpfall.cluster = set.ident(hpfall.cluster, which.cells(hpfall.cluster, "hpf20"),
    hpf20@ident)

boxPlot.FPKM(hpfall.cluster, "reads", name.y = "Total Reads", name.x = "", ratio.plot =
```

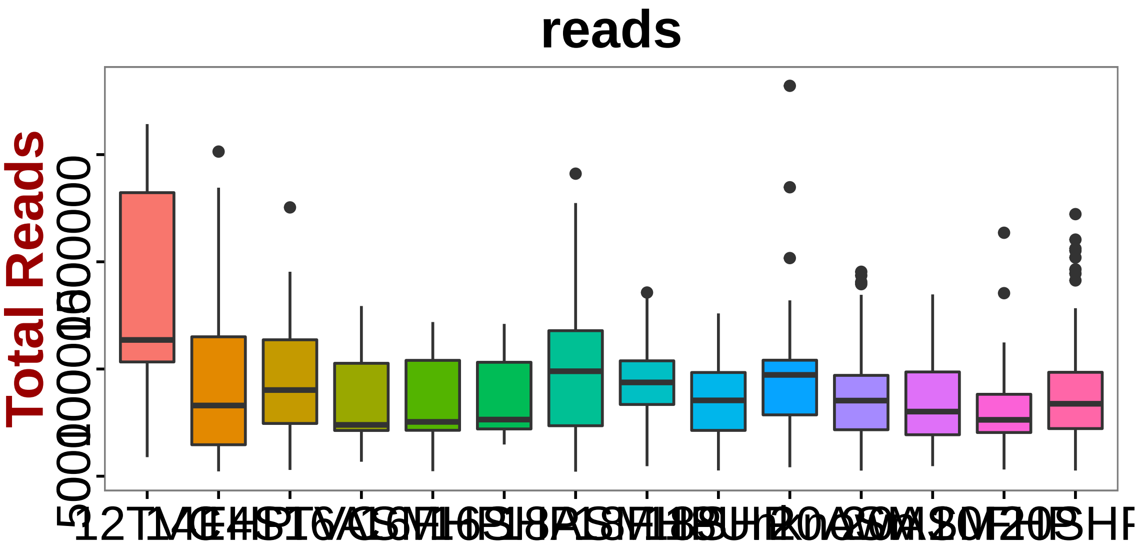


```
boxPlot.FPKM(hpfall.cluster, "map.rate", name.y = "Mapping Rates", name.x = "",
    ratio.plot = 0.1)
```

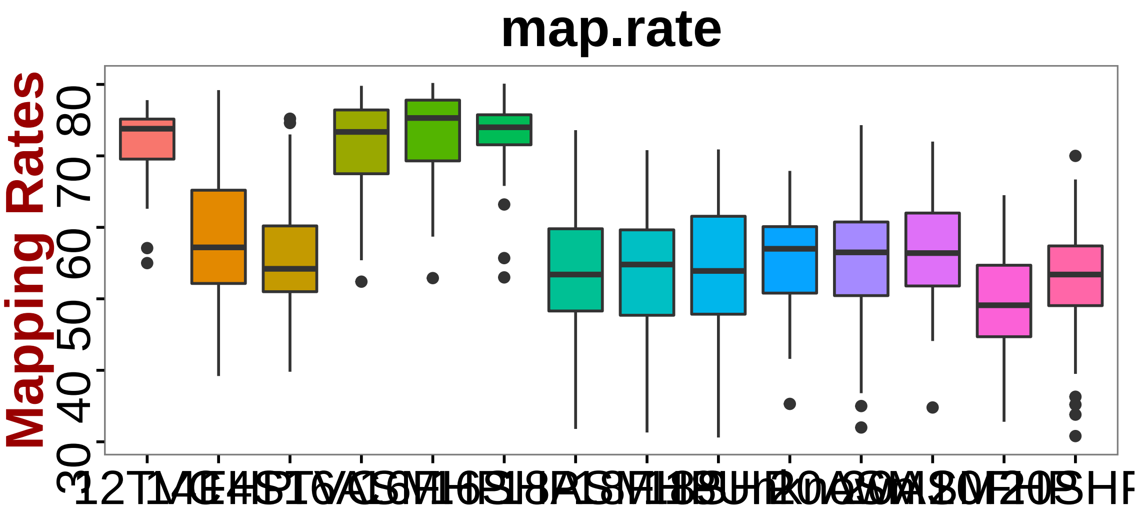


```
boxPlot.FPKM(hpfall.cluster, "nGene", name.y = "Number of Genes", name.x = "",
    ratio.plot = 0.002)
```

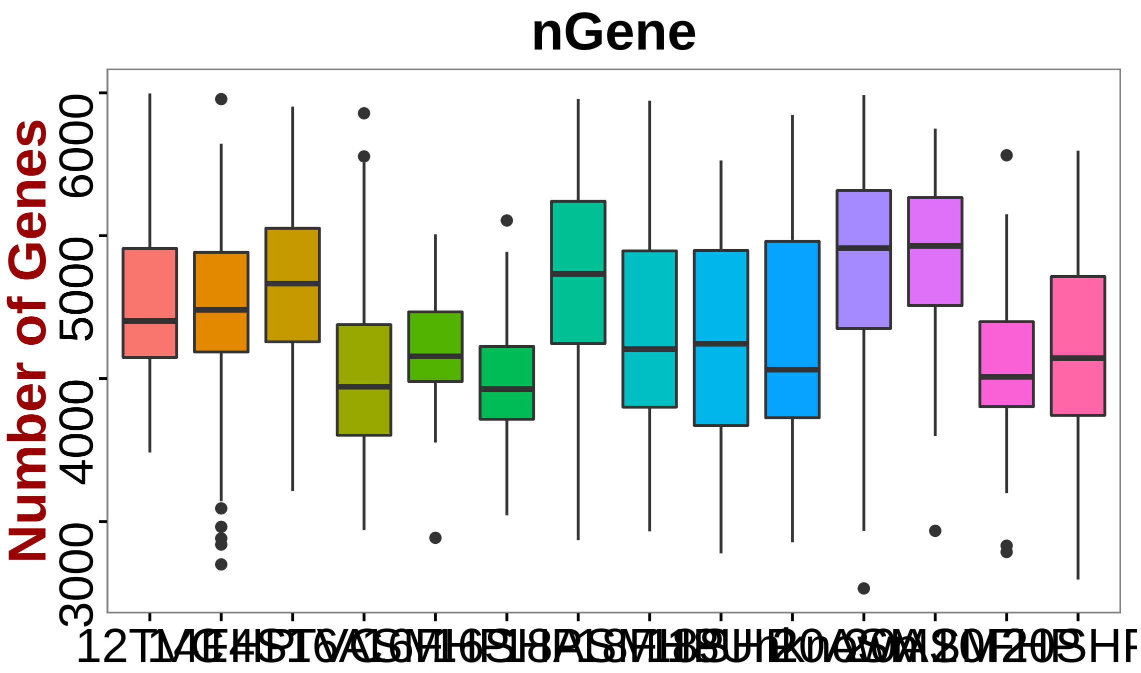


```r
save(hpfall.cluster, file = "hpfallCluster.Robj")
save(hpf12, hpf14, hpf16, hpf18, hpf20, file = "hpfall.Robj")
```
