## Supplementary material for "A single cell transcriptional roadmap for cardiopharyngeal fate diversification": Markdown_Preprocessing

### Preprocess

*Xiang Niu May 11, 2016*

1. Load packages

```
library(Seurat)
library(pbapply)
library(RColorBrewer)
source("functions.R")
col = colorRampPalette(rev(brewer.pal(n = 10, name = "RdBu")))
```

2. Merge Data

```
# read data
hpf12.data = read.table("hpf12.txt", sep = "\t", header = T, row.names = 1)
hpf14.data = read.table("hpf14.txt", sep = "\t", header = T, row.names = 1)
hpf16.data = read.table("hpf16.txt", sep = "\t", header = T, row.names = 1)
hpf18.data = read.table("hpf18.txt", sep = "\t", header = TRUE, row.names = 1)
hpf20.data = read.table("hpf20.txt", sep = "\t", header = TRUE, row.names = 1)

# change the column names
colnames(hpf12.data) = paste("hpf12_", colnames(hpf12.data), sep = "")
colnames(hpf14.data) = paste("hpf14_", colnames(hpf14.data), sep = "")
colnames(hpf16.data) = paste("hpf16_", colnames(hpf16.data), sep = "")
colnames(hpf18.data) = paste("hpf18_", colnames(hpf18.data), sep = "")
colnames(hpf20.data) = paste("hpf20_", colnames(hpf20.data), sep = "")

# pool all cell types
hpfall.data = cbind(hpf12.data, hpf14.data, hpf16.data, hpf18.data, hpf20.data)
dim(hpfall.data)
```

```
## [1] 15287  1796
```

```
# transform data to log scale
hpfall.data = log(hpfall.data + 1)

# setup Seurat object
hpfall = new("seurat", raw.data = hpfall.data)

# Take all genes in > 3 cells, all cells with 2k < detected genes < 6k, use
# an expression threshold of 1 log(FPKM)
```

```
hpfall = setup(hpfall, project = "allhpf", min.cells = 3, min.genes = 2000,
    is.expr = 1, names.field = 1, names.delim = "_")
hpfall = subsetData(hpfall, subset.name = "nGene", accept.high = 6000)
hpfall
```

```
## An object of class seurat in project allhpf
##  14864 genes across 1182 samples.
```

3. QC of data

```
############################## hpf12#######################
hpf12summary = list.files(path = "./summary/hpf12summary")
reads12 = NULL
map12 = NULL
cellName12 = substring(which.cells(hpfall, "hpf12"), 7)
hpf12summaryName = gsub("cufflinks", "", cellName12)
hpf12Names = NULL
for (i in hpf12summaryName) {
    hpf12Names = append(hpf12Names, grep(i, hpf12summary))
}
hpf12summary = hpf12summary[hpf12Names]
for (i in hpf12summary) {
    cell = read.csv(paste("./summary/hpf12summary/", i, sep = ""))
    reads = grep("Input", cell$Reads.)
    map = grep("overall", cell$Reads.)
    totReads = as.numeric(substr(as.character(cell$Reads.[reads]), 22, 50))
    mapRates = as.numeric(substr(as.character(cell$Reads.[map]), 1, 4))
    map12 = append(map12, mapRates)
    reads12 = append(reads12, totReads)
}

# Add data infor to Seurat object
[, c("map.rate", "reads")] = NA
[hpfall@ident == "hpf12", "map.rate"] = map12
[hpfall@ident == "hpf12", "reads"] = reads12
############################## hpf14#######################
hpf14summary = list.files(path = "./summary/hpf14summary")
reads14 = NULL
map14 = NULL
cellName14 = substring(which.cells(hpfall, "hpf14"), 7)
hpf14summaryName = gsub("cufflinks", "", cellName14)
hpf14Names = NULL
for (i in hpf14summaryName) {
```

```
    hpf14Names = append(hpf14Names, grep(i, hpf14summary))
}
hpf14summary = hpf14summary[hpf14Names]
for (i in hpf14summary) {
    cell = read.csv(paste("./summary/hpf14summary/", i, sep = ""))
    reads = grep("Input", cell$Reads.)
    map = grep("overall", cell$Reads.)
    totReads = as.numeric(substr(as.character(cell$Reads.[reads]), 22, 50))
    mapRates = as.numeric(substr(as.character(cell$Reads.[map]), 1, 4))
    map14 = append(map14, mapRates)
    reads14 = append(reads14, totReads)
}

# Add data infor to Seurat object
[hpfall@ident == "hpf14", "map.rate"] = map14
[hpfall@ident == "hpf14", "reads"] = reads14
############################## hpf16#######################
hpf16summary = list.files(path = "./summary/hpf16summary")
reads16 = NULL
map16 = NULL
cellName16 = substring(which.cells(hpfall, "hpf16"), 7)
hpf16summaryName = gsub("cufflinks", "", cellName16)
hpf16Names = NULL
for (i in hpf16summaryName) {
    hpf16Names = append(hpf16Names, grep(i, hpf16summary))
}
hpf16summary = hpf16summary[hpf16Names]
for (i in hpf16summary) {
    cell = read.csv(paste("./summary/hpf16summary/", i, sep = ""))
    reads = grep("Input", cell$Reads.)
    map = grep("overall", cell$Reads.)
    totReads = as.numeric(substr(as.character(cell$Reads.[reads]), 22, 50))
    mapRates = as.numeric(substr(as.character(cell$Reads.[map]), 1, 4))
    map16 = append(map16, mapRates)
    reads16 = append(reads16, totReads)
}

# Add data infor to Seurat object
[hpfall@ident == "hpf16", "map.rate"] = map16
[hpfall@ident == "hpf16", "reads"] = reads16
############################## hpf18#######################
hpf18summary = list.files(path = "./summary/hpf18summary")
reads18 = NULL
map18 = NULL
```

```r
cellName18 = substring(which.cells(hpfall, "hpf18"), 7)
hpf18summaryName = gsub("outcufflinks", "", cellName18)
hpf18Names = NULL
for (i in hpf18summaryName) {
    hpf18Names = append(hpf18Names, grep(i, hpf18summary))
}
hpf18summary = hpf18summary[hpf18Names]
for (i in hpf18summary) {
    cell = read.csv(paste("./summary/hpf18summary/", i, sep = ""))
    reads = grep("Input", cell$Reads.)
    map = grep("overall", cell$Reads.)
    totReads = as.numeric(substr(as.character(cell$Reads.[reads]), 22, 50))
    mapRates = as.numeric(substr(as.character(cell$Reads.[map]), 1, 4))
    map18 = append(map18, mapRates)
    reads18 = append(reads18, totReads)
}

# Add data infor to Seurat object
[hpfall@ident == "hpf18", "map.rate"] = map18
[hpfall@ident == "hpf18", "reads"] = reads18
############################## hpf20#######################
hpf20summary = list.files(path = "./summary/hpf20summary")
reads20 = NULL
map20 = NULL
cellName20 = substring(which.cells(hpfall, "hpf20"), 7)
hpf20summaryName = gsub("cufflinks", "", cellName20)
hpf20Names = NULL
for (i in hpf20summaryName) {
    hpf20Names = append(hpf20Names, grep(i, hpf20summary))
}
hpf20summary = hpf20summary[hpf20Names]
for (i in hpf20summary) {
    cell = read.csv(paste("./summary/hpf20summary/", i, sep = ""))
    reads = grep("Input", cell$Reads.)
    map = grep("overall", cell$Reads.)
    totReads = as.numeric(substr(as.character(cell$Reads.[reads]), 22, 50))
    mapRates = as.numeric(substr(as.character(cell$Reads.[map]), 1, 4))
    map20 = append(map20, mapRates)
    reads20 = append(reads20, totReads)
}

# Add data infor to Seurat object
[hpfall@ident == "hpf20", "map.rate"] = map20
[hpfall@ident == "hpf20", "reads"] = reads20
```

```r
############################################################### Plot mappint rates from all t
plot(NULL, xlab = "Mapping Rates", ylab = "Density", xlim = c(0, 100), ylim = c(0,
    0.1), cex.lab = 1.2, font.lab = 2)
points(density(map12), type = "l", col = brewer.pal(5, "Set1")[1], lwd = 3)
points(density(map14), type = "l", col = brewer.pal(5, "Set1")[2], lwd = 3)
points(density(map16), type = "l", col = brewer.pal(5, "Set1")[3], lwd = 3)
points(density(map18), type = "l", col = brewer.pal(5, "Set1")[4], lwd = 3)
points(density(map20), type = "l", col = brewer.pal(5, "Set1")[5], lwd = 3)
legend("topright", legend = c("hpf12", "hpf14", "hpf16", "hpf18", "hpf20"),
    lty = 1, col = brewer.pal(5, "Set1"), cex = 0.75, xpd = T, inset = c(0.1,
        0), bty = "n")
```

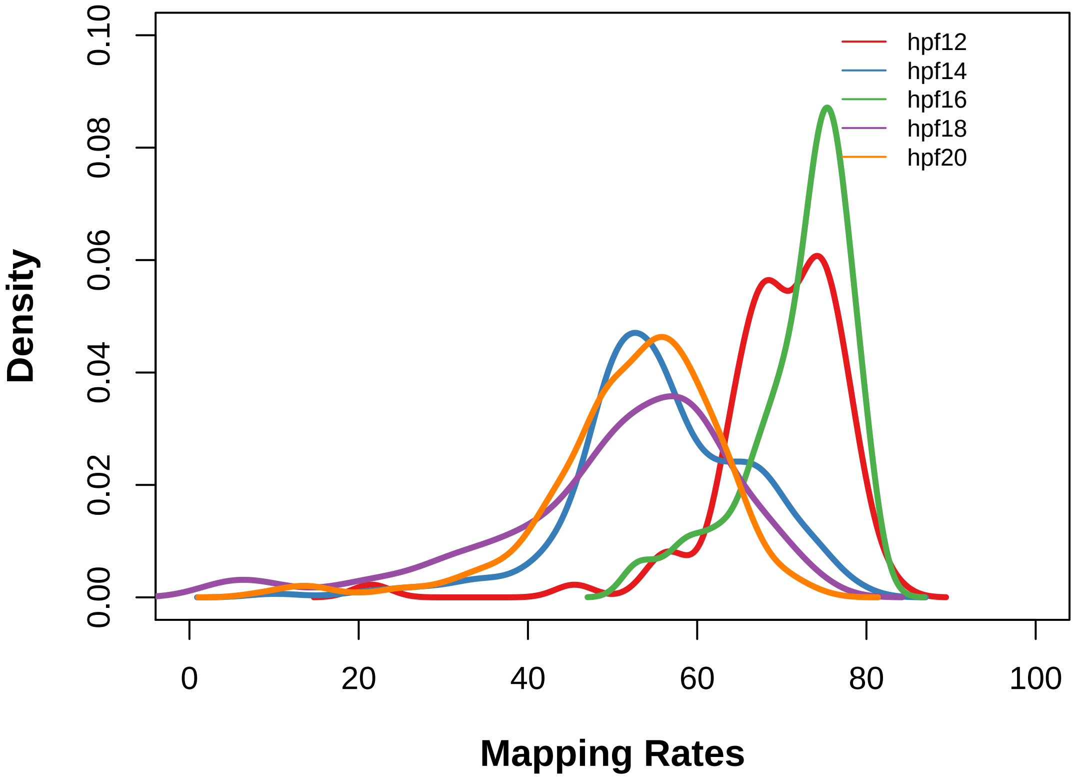


```r
# Plot total reads from all data
plot(NULL, xlab = "Total Reads", ylab = "Density", xlim = c(0, 3e+06), ylim = c(0,
    3e-06), cex.lab = 1.2, font.lab = 2)
points(density(reads12), type = "l", col = brewer.pal(5, "Set1")[1], lwd = 3)
points(density(reads14), type = "l", col = brewer.pal(5, "Set1")[2], lwd = 3)
points(density(reads16), type = "l", col = brewer.pal(5, "Set1")[3], lwd = 3)
points(density(reads18), type = "l", col = brewer.pal(5, "Set1")[4], lwd = 3)
points(density(reads20), type = "l", col = brewer.pal(5, "Set1")[5], lwd = 3)
legend("topright", legend = c("hpf12", "hpf14", "hpf16", "hpf18", "hpf20"),
    lty = 1, col = c("red", "blue", "yellow", "green", "black"), cex = 0.75,
    xpd = T, inset = c(0.1, 0), bty = "n")
```

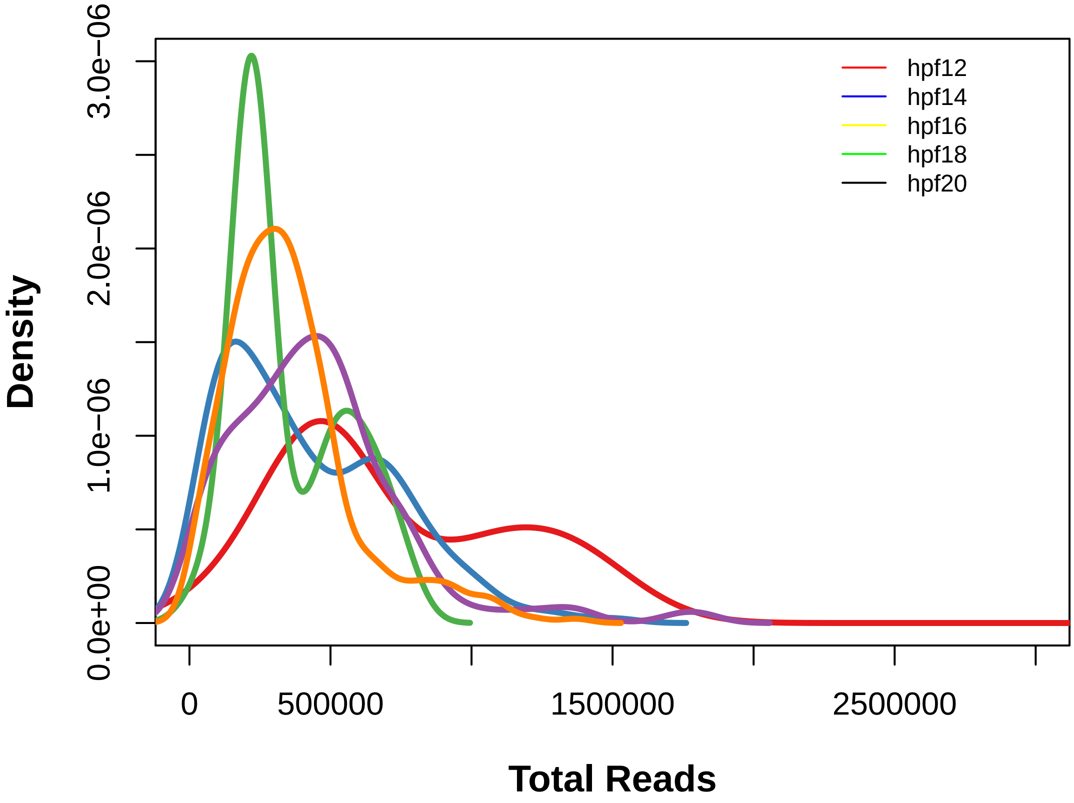


```r
# Remove cells with mapping rate < 30%, reads > 2*10^6
boxPlot.FPKM(hpfall, "map.rate", name.y = "Mapping Rate(%)", ratio.plot = 0.06,
    name.x = "", name = "Mapping Rates")
```

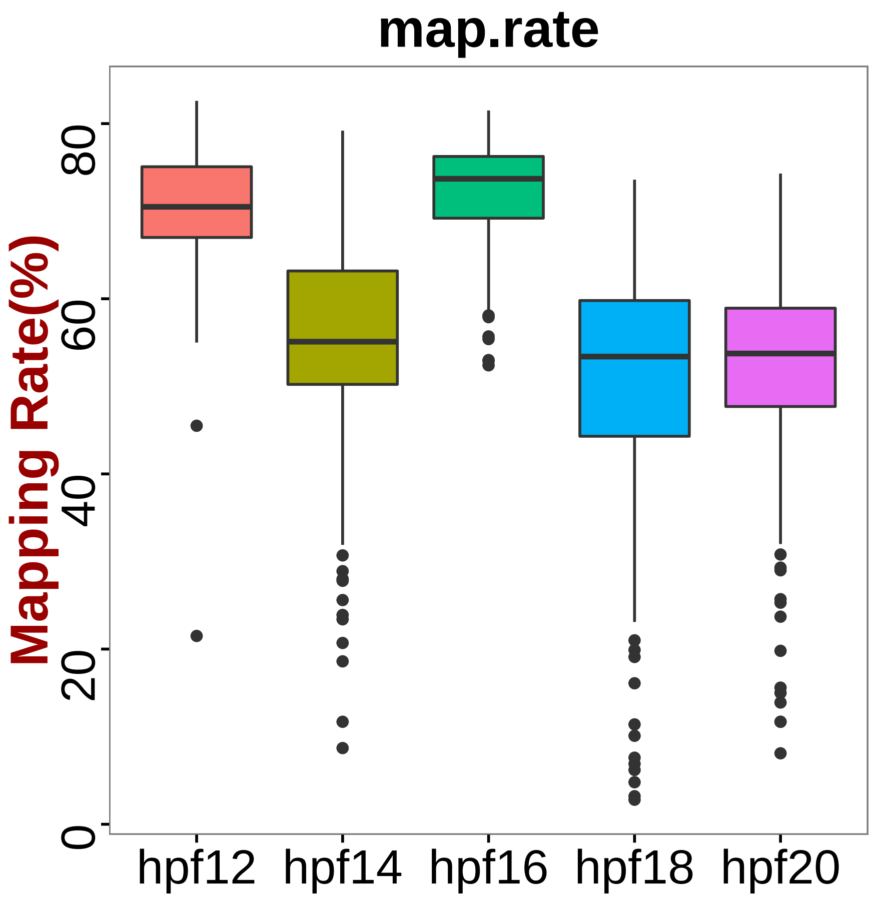


```
boxPlot.FPKM(hpfall, "reads", name.y = "Total Reads", name.x = "", ratio.plot = 1.2e-06,
    name = "Total Reads")
```

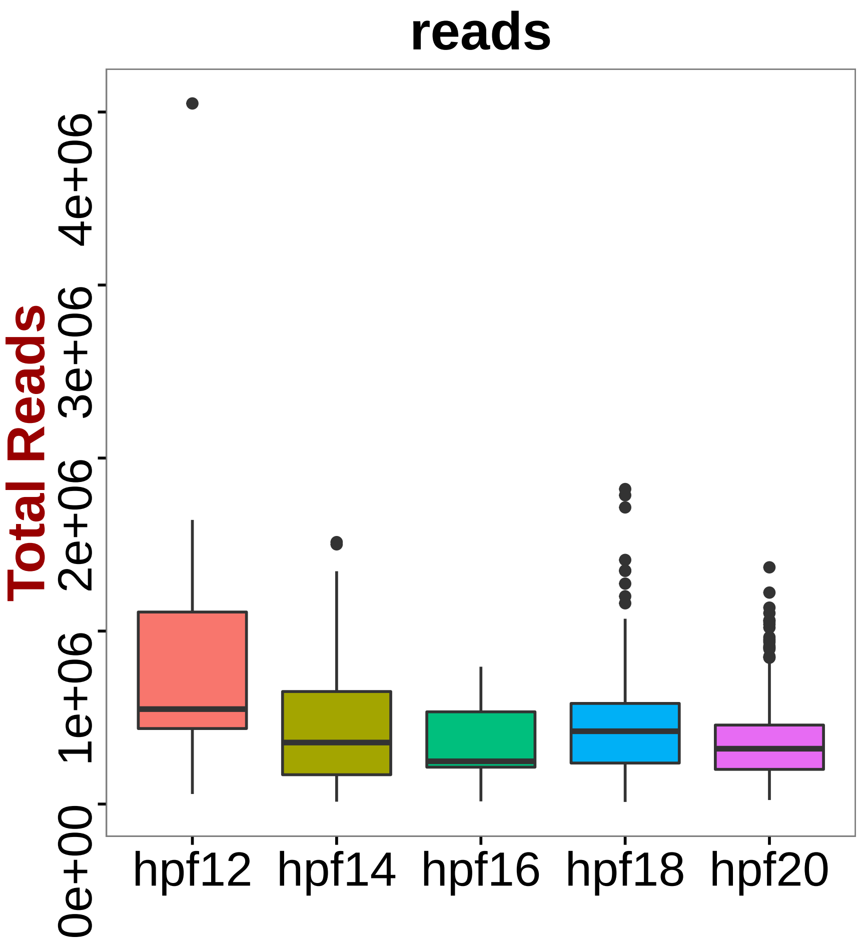


```r
hpfall = subsetData(hpfall, subset.name = "map.rate", accept.low = 30)
hpfall = subsetData(hpfall, subset.name = "reads", accept.high = 2e+06)
hpfall
```

```
## An object of class seurat in project allhpf
##  14864 genes across 1138 samples.
```

```r
boxPlot.FPKM(hpfall, "map.rate", name.y = "Mapping Rate(%)", ratio.plot = 0.1,
    name.x = "", name = "Mapping Rates")
```

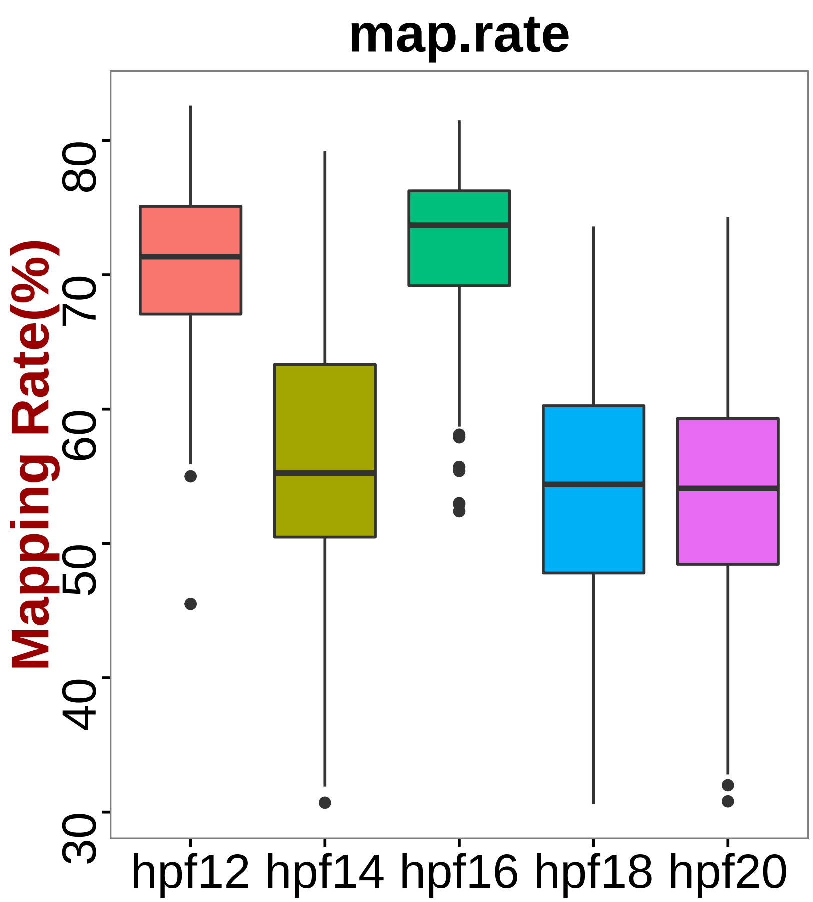


```
boxPlot.FPKM(hpfall, "reads", name.y = "Total Reads", name.x = "", ratio.plot = 3e-06,
    name = "Total Reads")
```

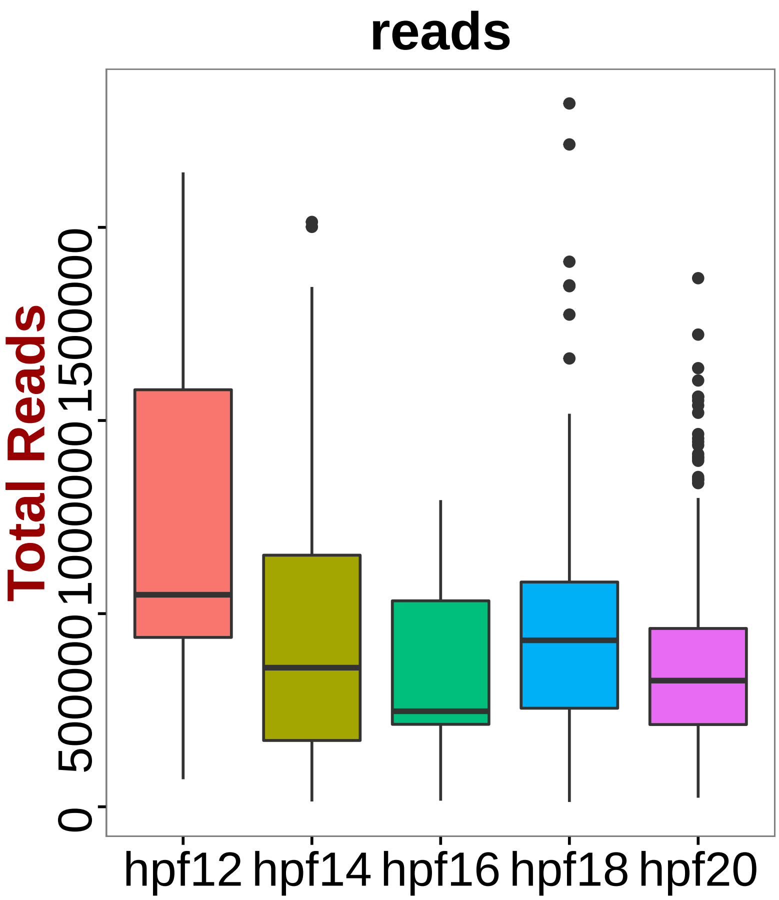


4. Remove Contamination and Batch Effect

```
# PCA on all the genes
hpfall = pca(hpfall, pc.genes = rownames(hpfall@data), do.print = F)
pcScree(hpfall, rownames(hpfall@data), 10)
abline(h = 0.5)
```

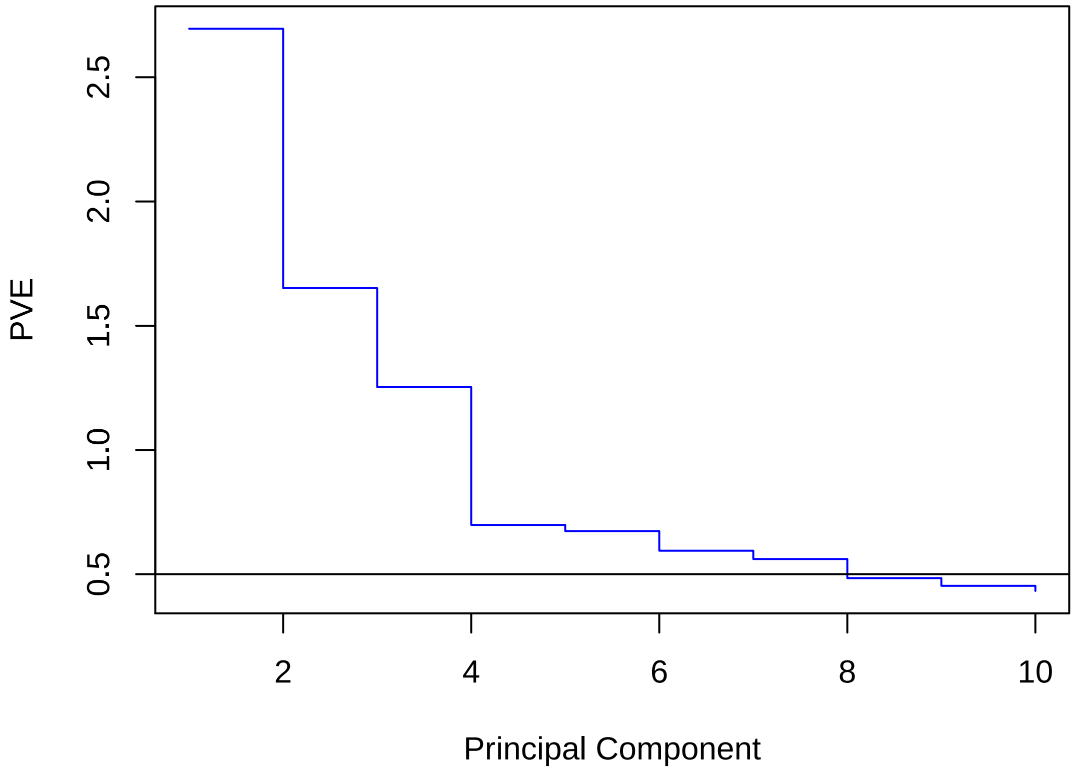


```
# Top 7 PCs are selected (PVE>0.5%)

# PC1 Population of potential contaminated cell type
doHeatMap(hpfall, remove.key = T, slim.col.label = T, genes.use = pcTopGenes(hpfall,
    1), cells.use = pcTopCells(hpfall, 1), col.use = col)
```

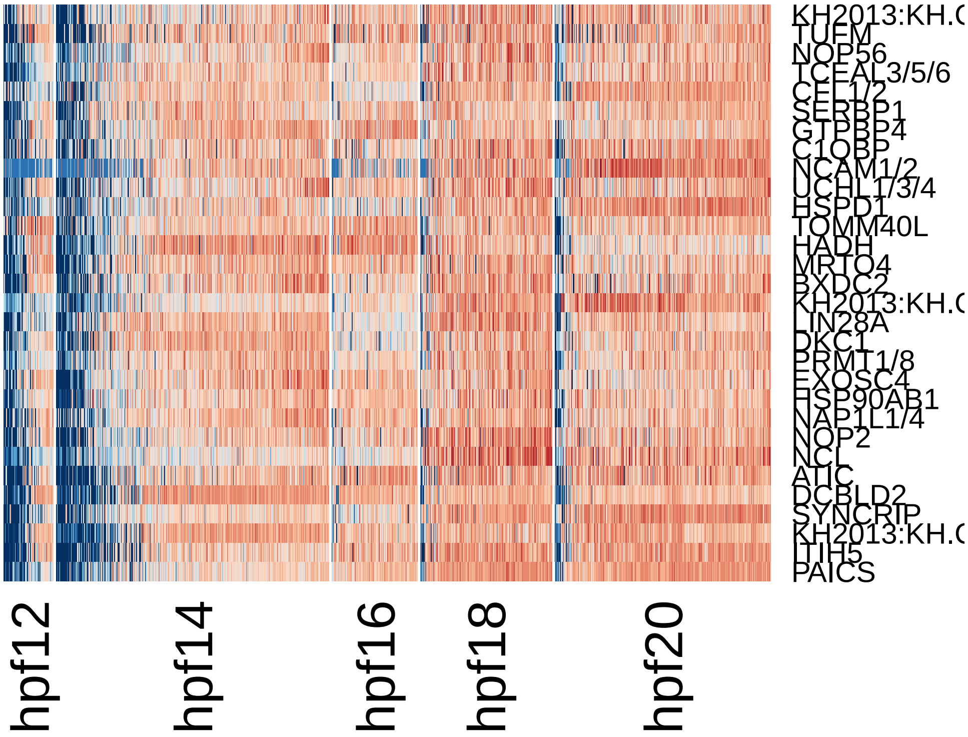


```
pca.plot(hpfall, 1, 2)
```

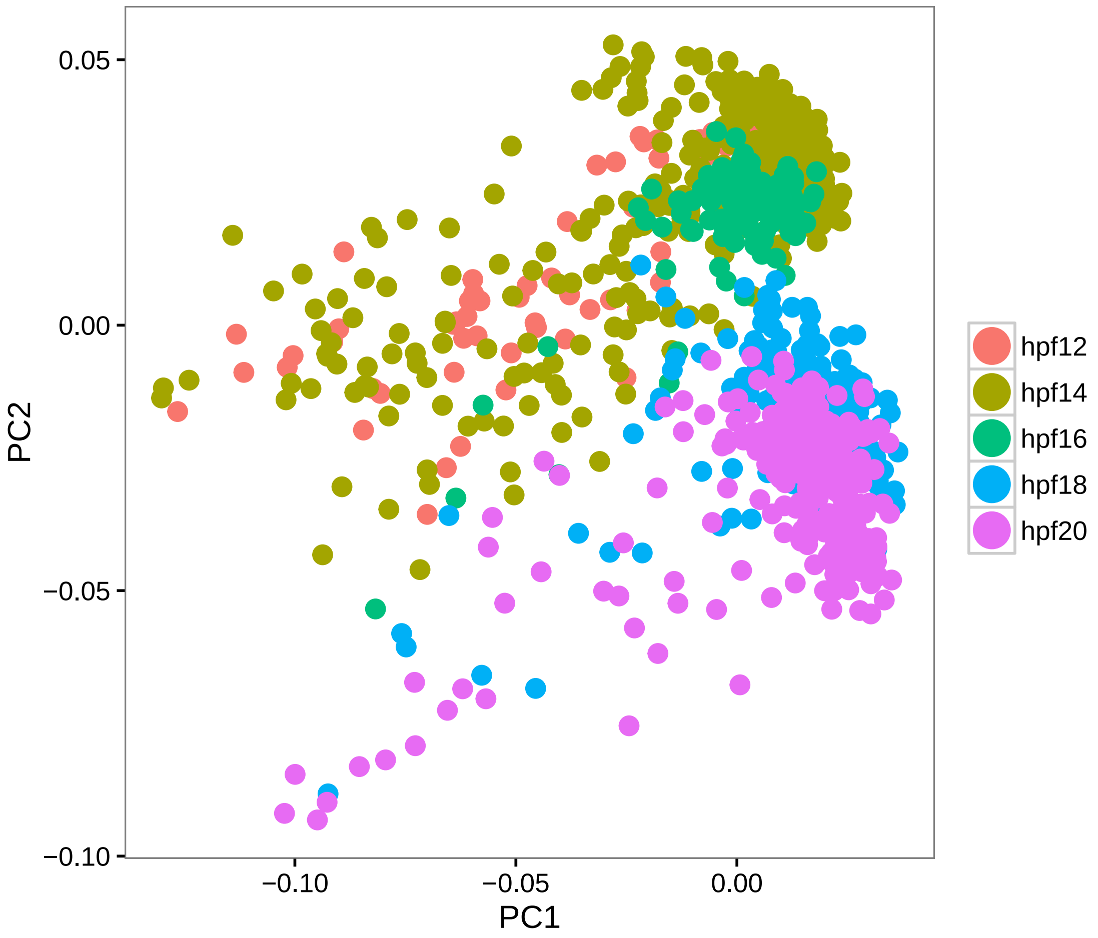


```r
# PC2 Batch effect due to library preparation, it separates hpf12/14/16 with
# hpf18/20
doHeatMap(hpfall, remove.key = T, slim.col.label = T, genes.use = pcTopGenes(hpfall,
    2), cells.use = pcTopCells(hpfall, 2), col.use = col)
```

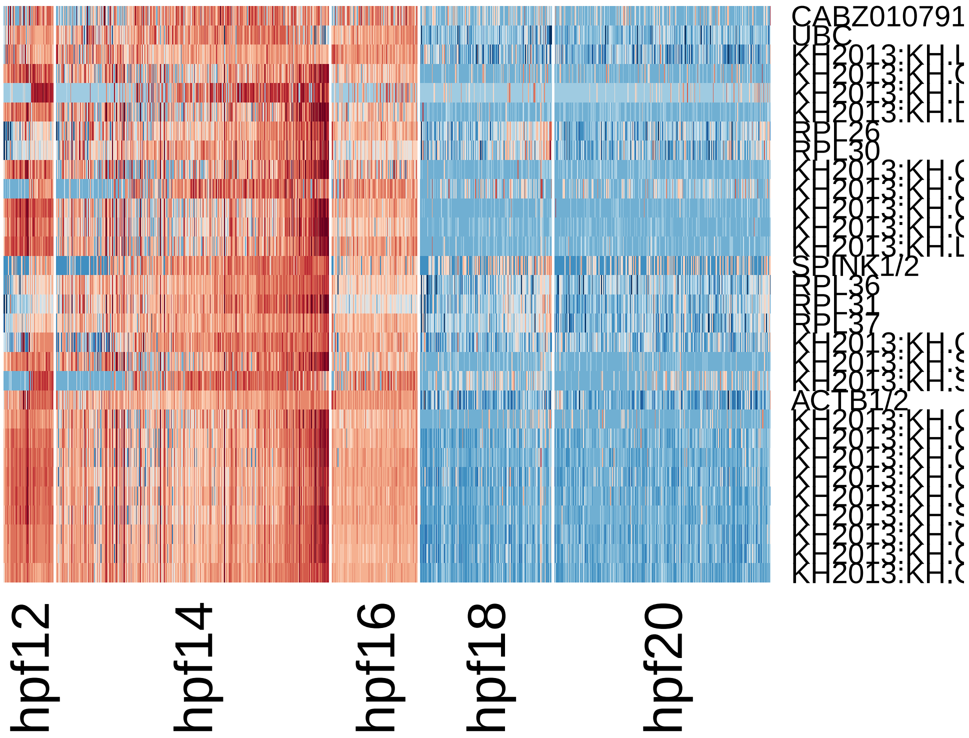


```
pca.plot(hpfall, 1, 2)
```

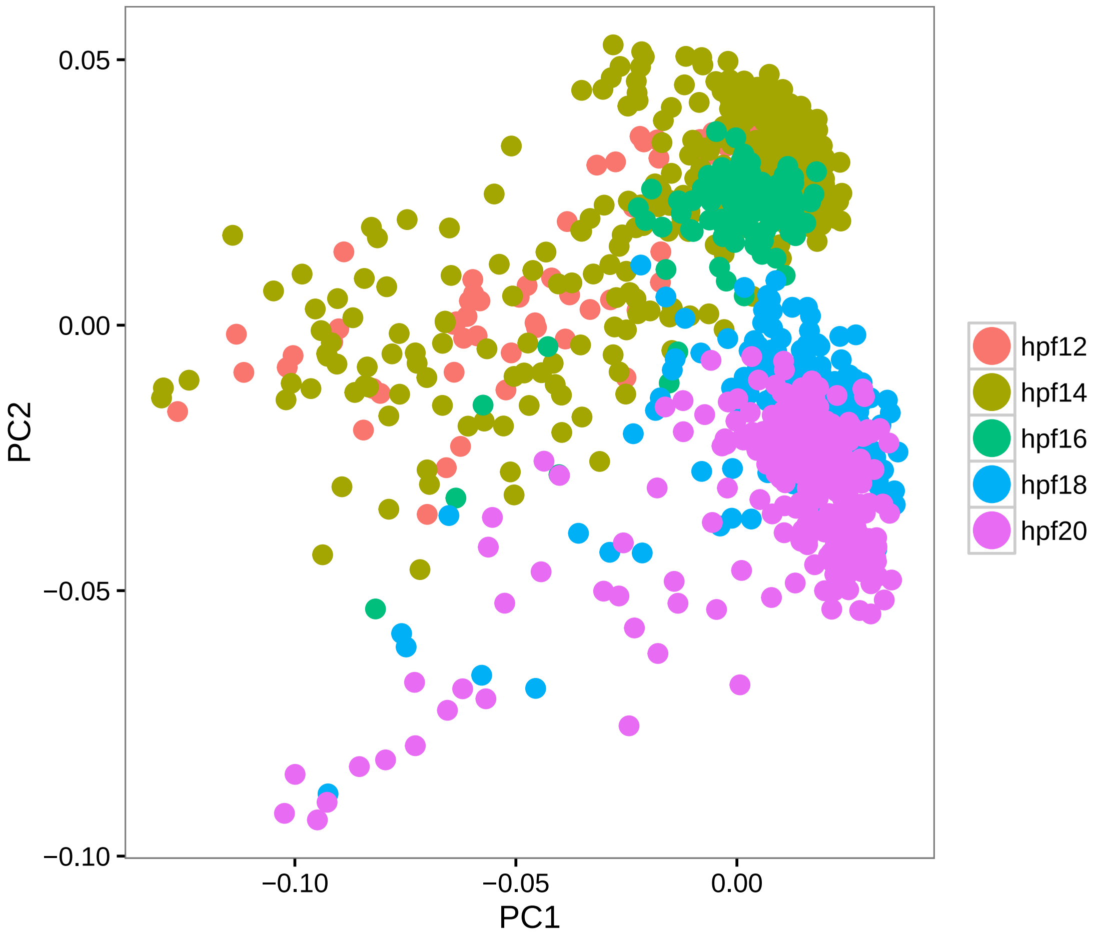


```r
# PC3 Mesenchymal contamination
doHeatMap(hpfall, remove.key = T, slim.col.label = T, genes.use = pcTopGenes(hpfall,
    3), cells.use = pcTopCells(hpfall, 3), col.use = col)
```

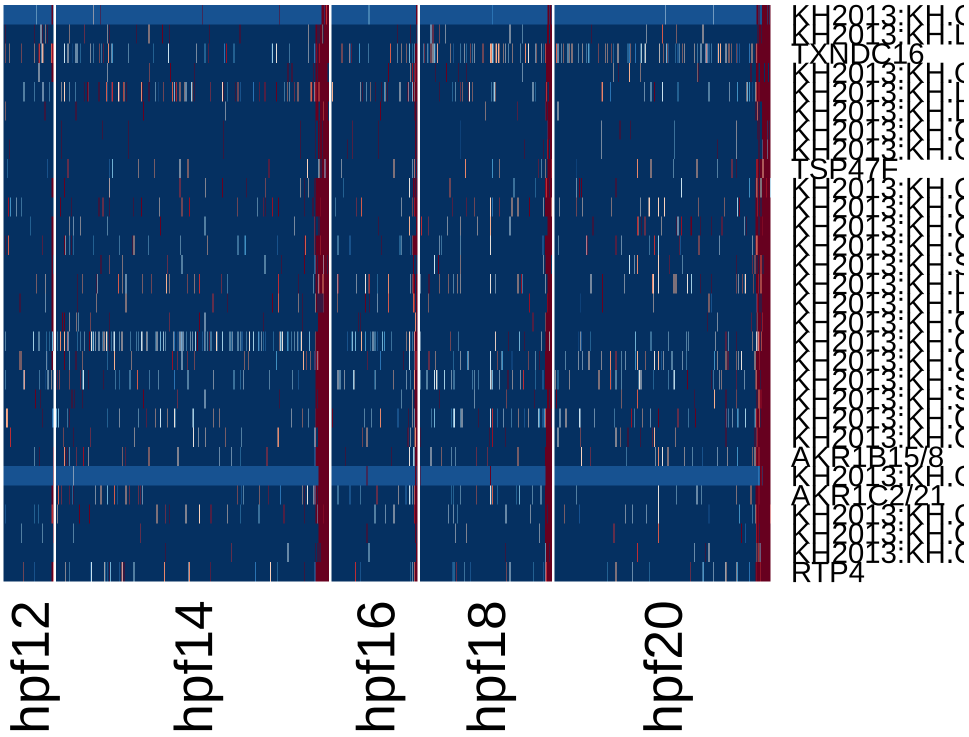


```r
pca.plot(hpfall, 1, 3)
```

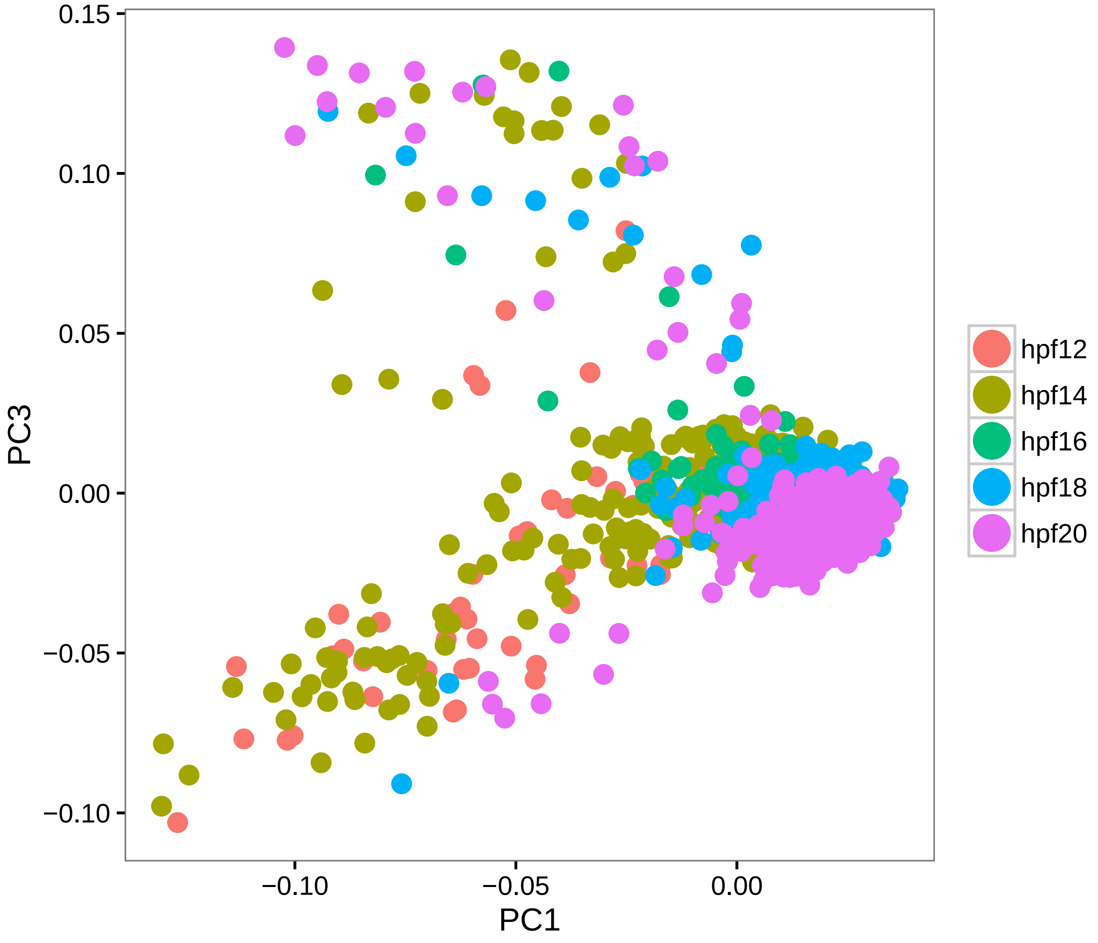


```
feature.plot(hpfall, c("RTP4", "TWIST1"), reduction.use = "pca", dim.2 = 3)
```

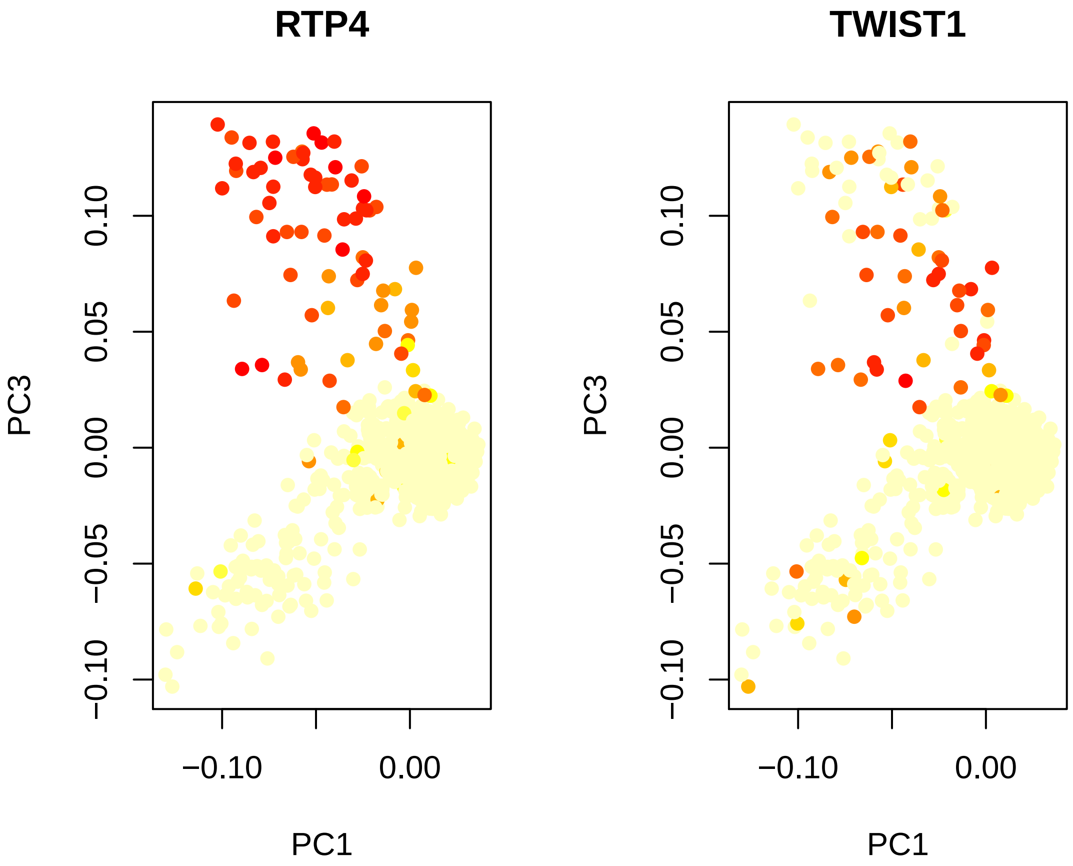


```r
# PC4 Heart vs Muscle split
doHeatMap(hpfall, remove.key = T, slim.col.label = T, genes.use = pcTopGenes(hpfall,
    4), cells.use = pcTopCells(hpfall, 4), col.use = col)
```

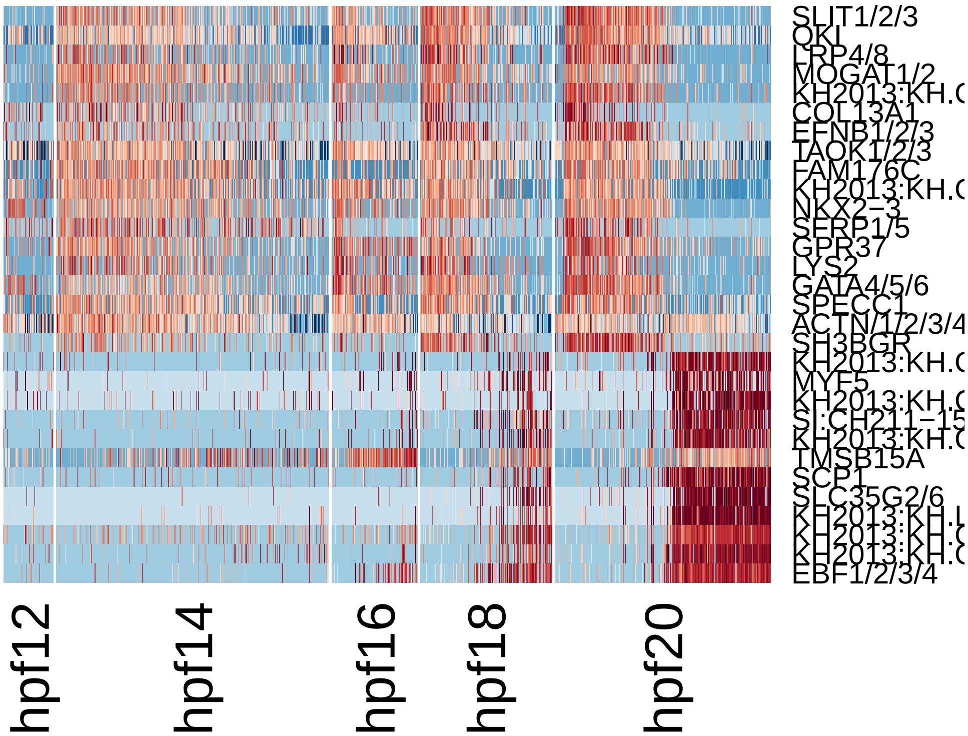


```
feature.plot(hpfall, c("NKX2-3", "EBF1/2/3/4", "TBX1/10", "GATA4/5/6"), reduction.use =
    dim.2 = 4)
```

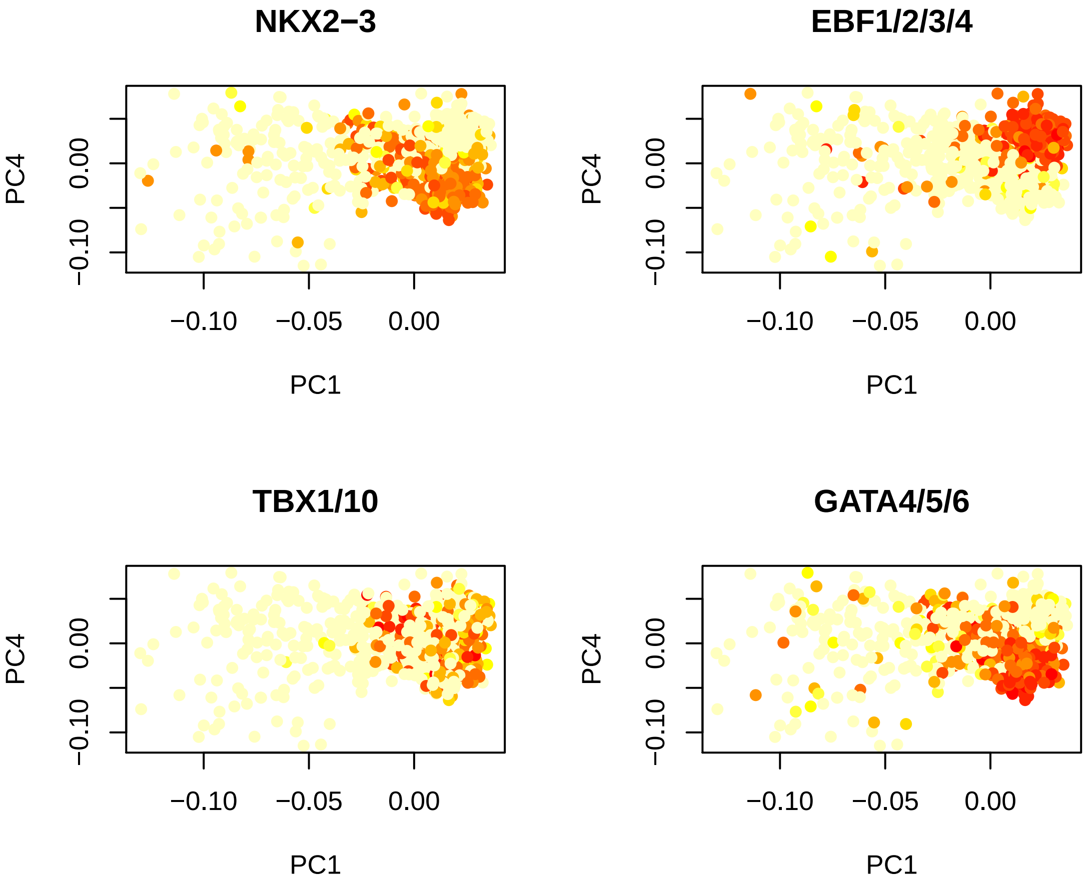


```r
# PC5 Another batch effect
doHeatMap(hpfall, remove.key = T, slim.col.label = T, genes.use = pcTopGenes(hpfall,
    5), cells.use = pcTopCells(hpfall, 5), col.use = col)
```

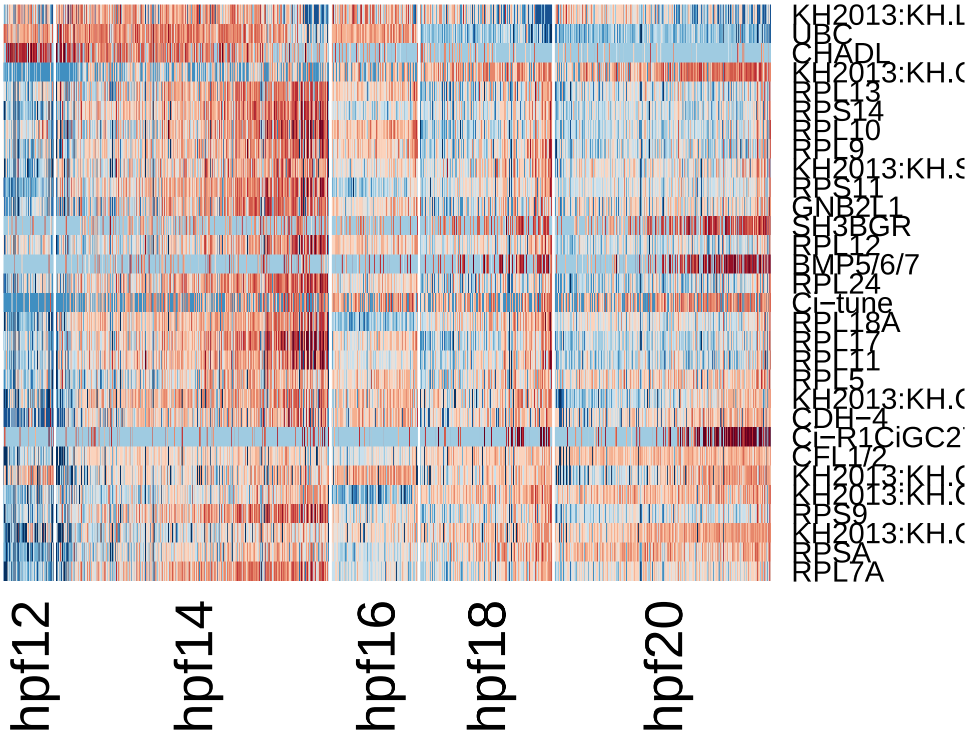


```
pca.plot(hpfall, 1, 5)
```

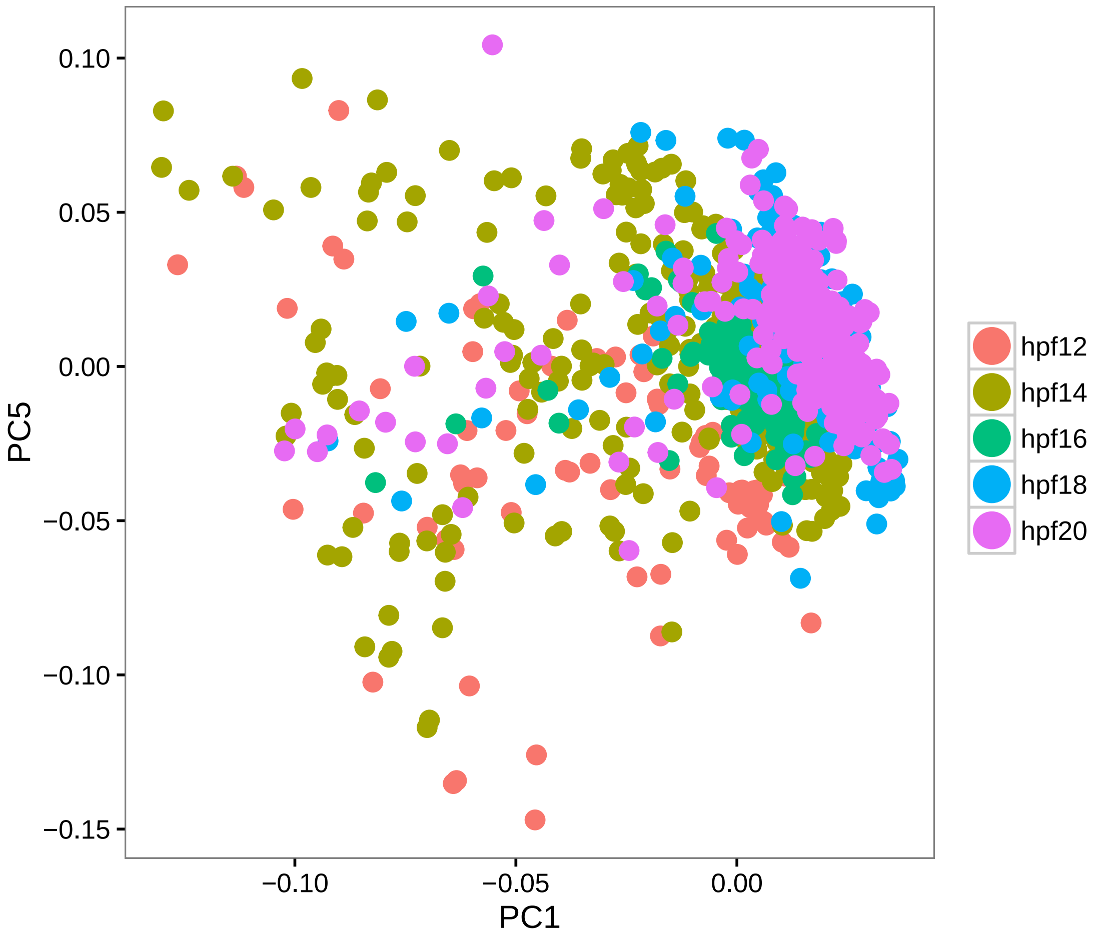


```
# PC6 Saperation by heart vs muscle
doHeatMap(hpfall, remove.key = T, slim.col.label = T, genes.use = pcTopGenes(hpfall,
    6), cells.use = pcTopCells(hpfall, 6), col.use = col)
```

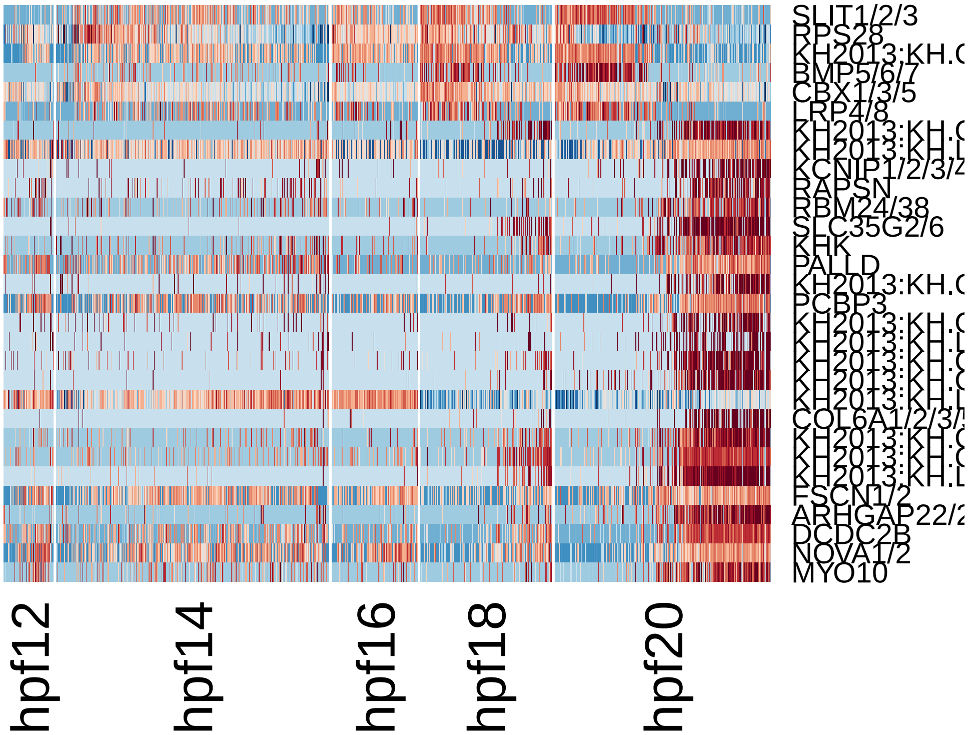


```
pca.plot(hpfall, 1, 6)
```

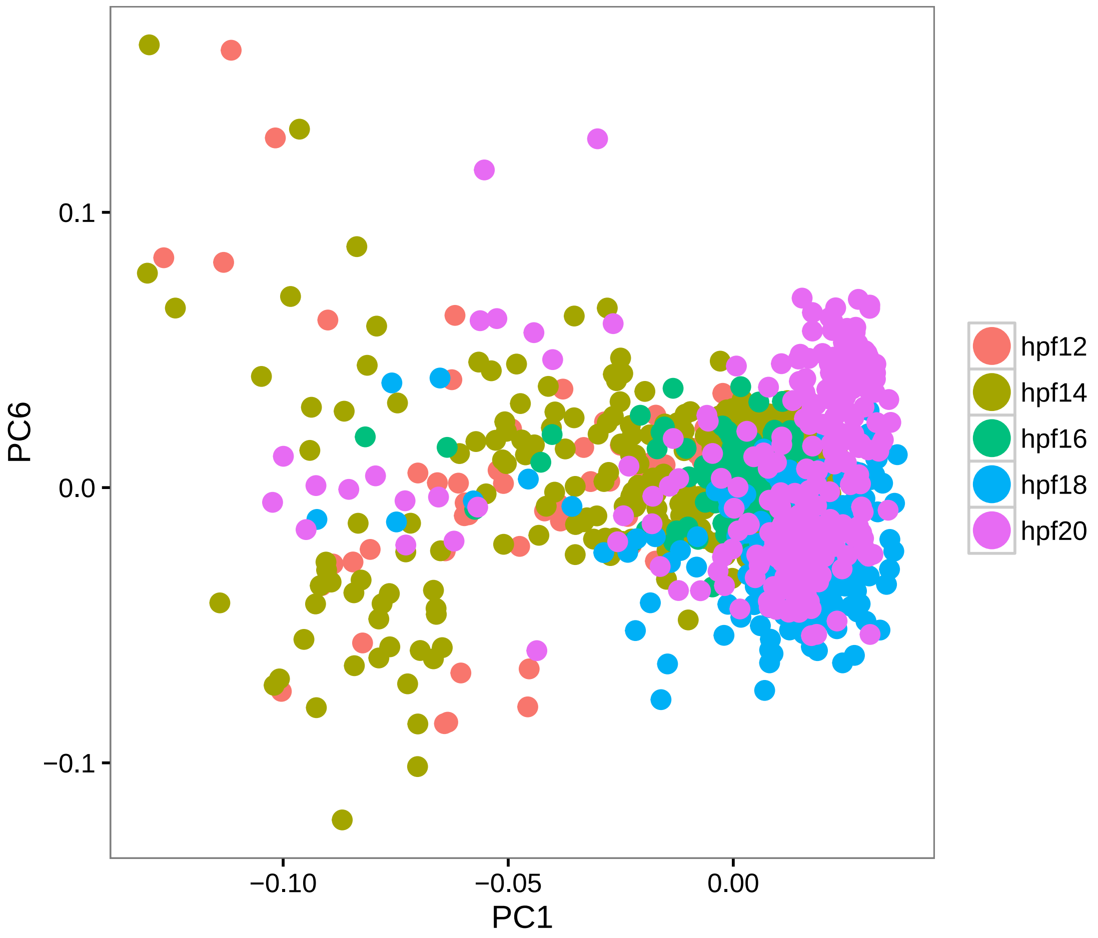


```
feature.plot(hpfall, c("NKX2-3", "EBF1/2/3/4", "TBX1/10", "GATA4/5/6"), reduction.use =
    dim.2 = 6)
```

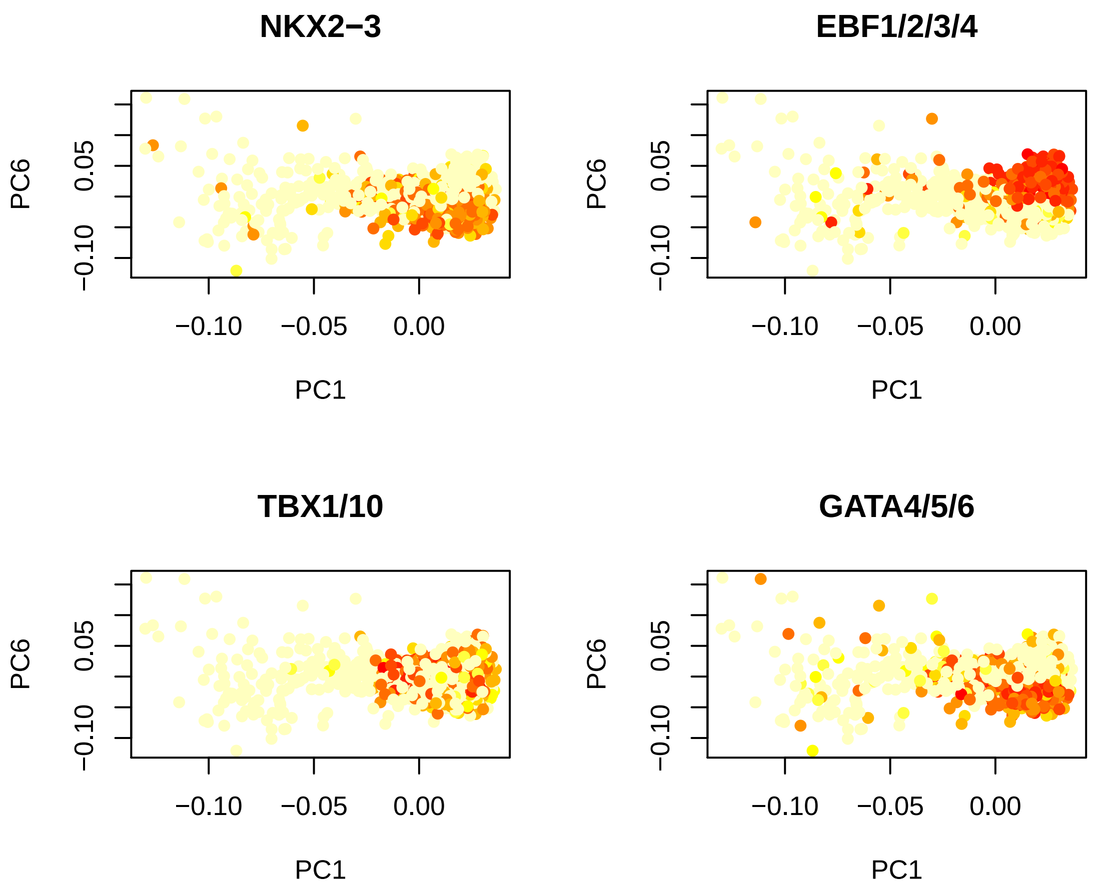


```r
# PC7 Batch effect marked hpf14
doHeatMap(hpfall, remove.key = T, slim.col.label = T, genes.use = pcTopGenes(hpfall,
    7), cells.use = pcTopCells(hpfall, 7), col.use = col)
```

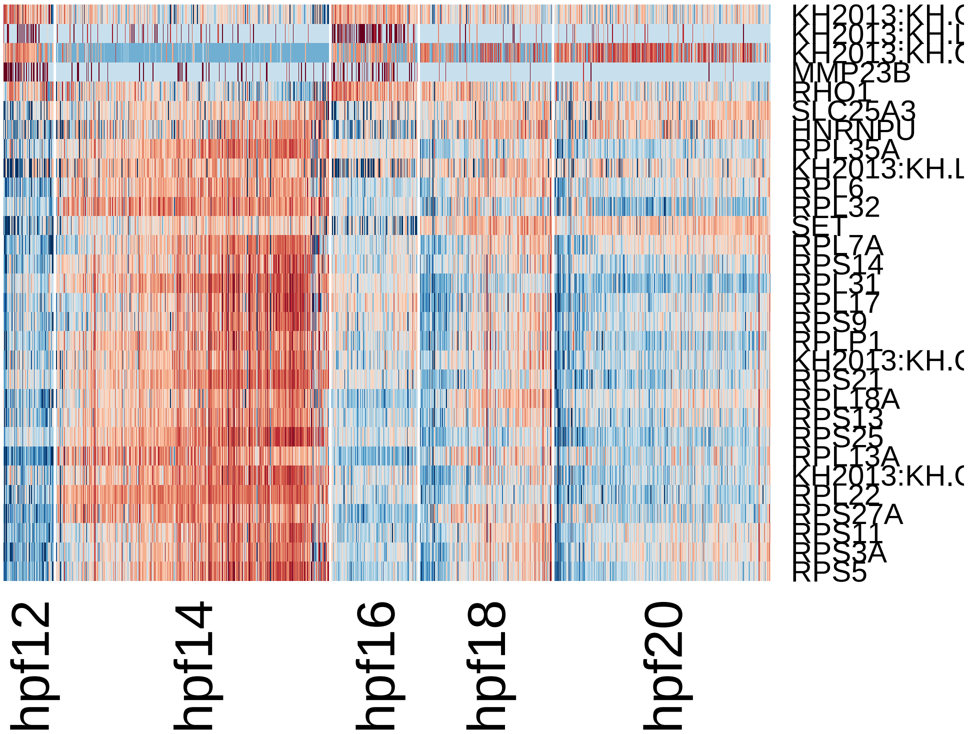


```
pca.plot(hpfall, 1, 7)
```

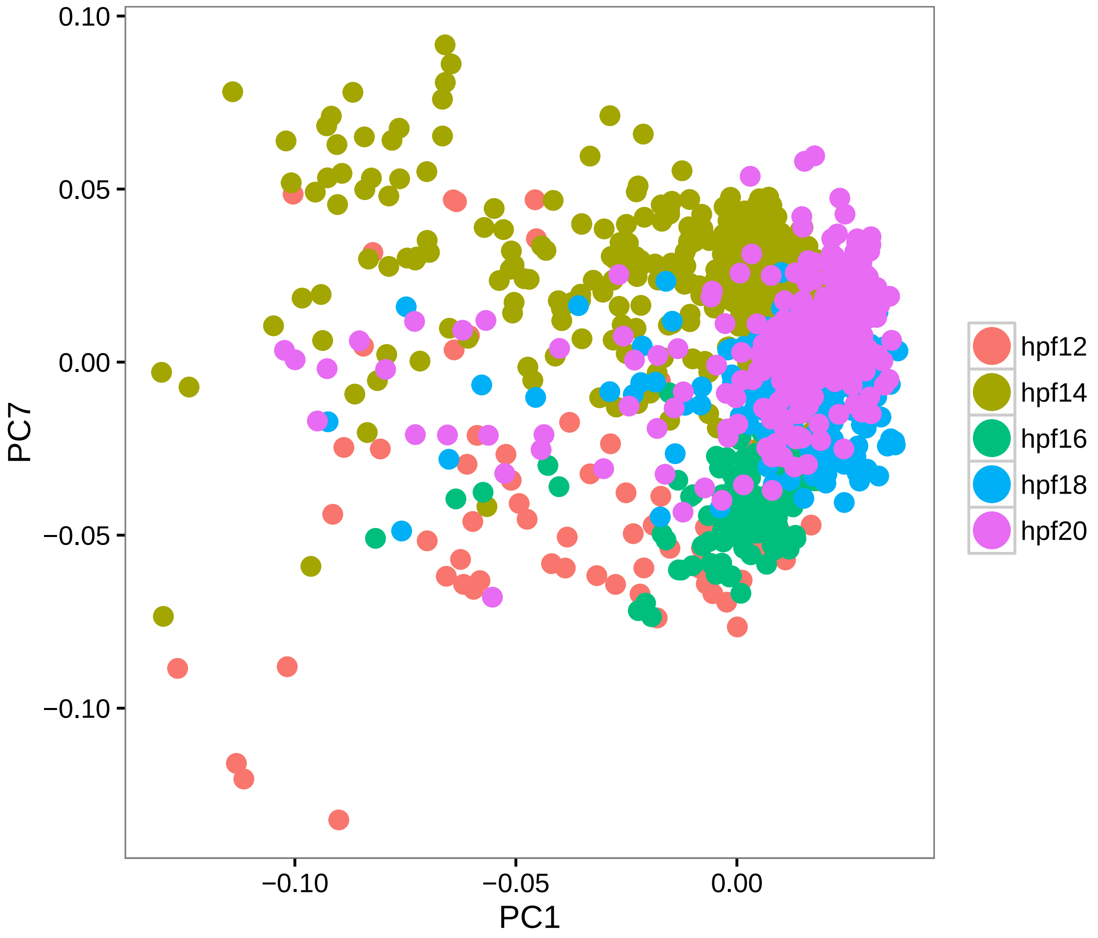


```
# Remove batch effect
hpfall.remv1 = RegressOut(hpfall, c("PC2", "PC5", "PC7"), do.scale = T)

# save object
save(hpfall.remv1, file = "hpfall.remv1.Robj")

# Rerun PCA with all genes
hpfall.remv1 = pca(hpfall.remv1, pc.genes = rownames(hpfall.remv1@data), do.print = F)
pcScree(hpfall.remv1, rownames(hpfall.remv1@data), 10)
abline(h = 0.5)
```

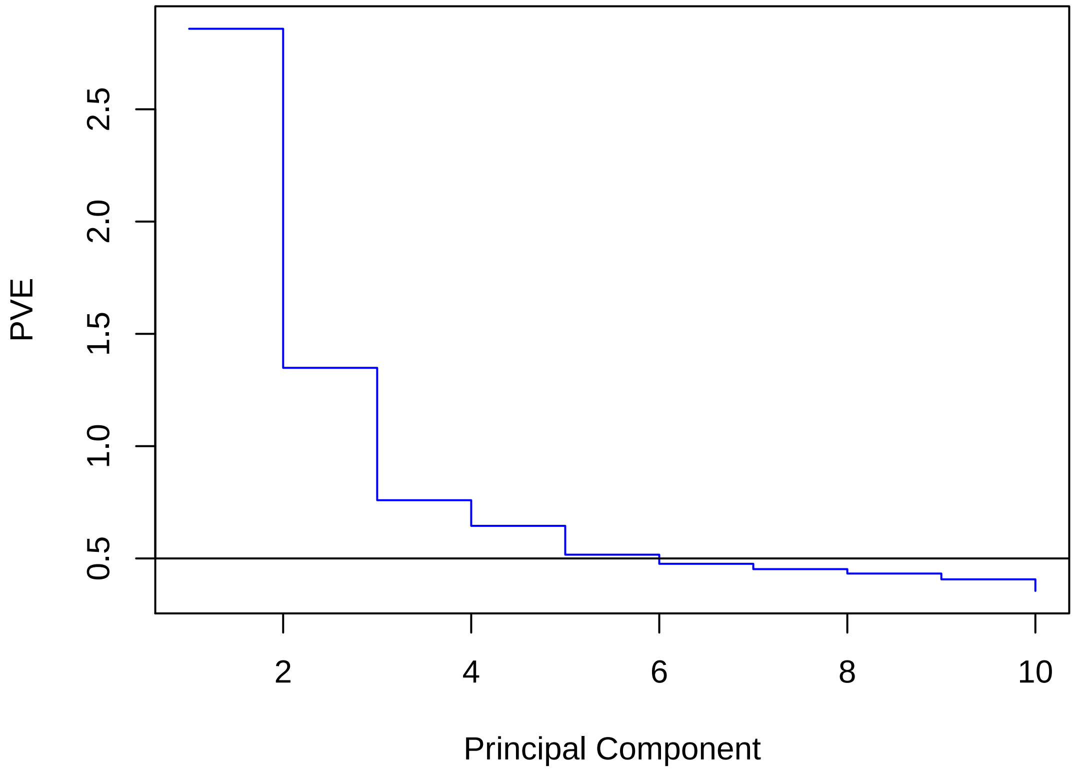


```r
# Top 5 PCs are selected (PVE>0.5%)

# PC1 Population of potential contamination cell type
doHeatMap(hpfall.remv1, remove.key = T, slim.col.label = T, genes.use = pcTopGenes(hpfal
    1), cells.use = pcTopCells(hpfall.remv1, 1), col.use = col)
```

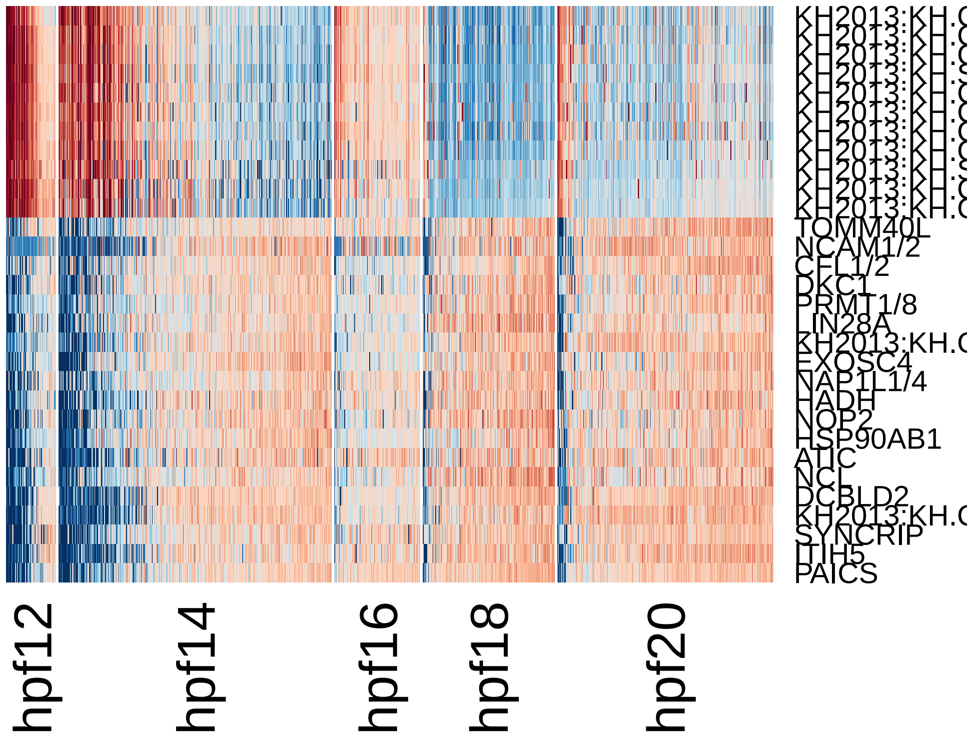


```r
pca.plot(hpfall.remv1, 1, 2)
```

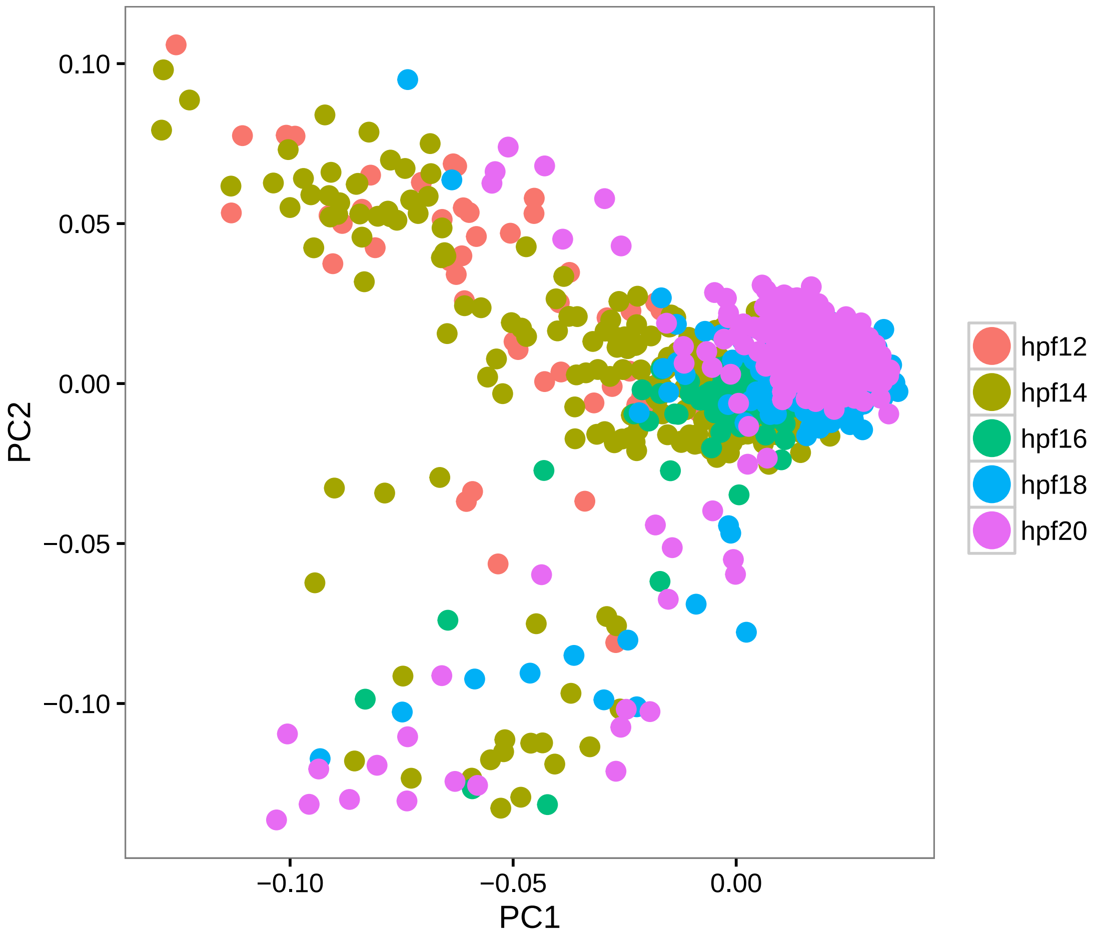


```
# This group of cells also do not express known TVC lineage markers
feature.plot(hpfall.remv1, c("NKX2-3", "GATA4/5/6", "EBF1/2/3/4", "TBX1/10"),
    reduction.use = "pca")
```

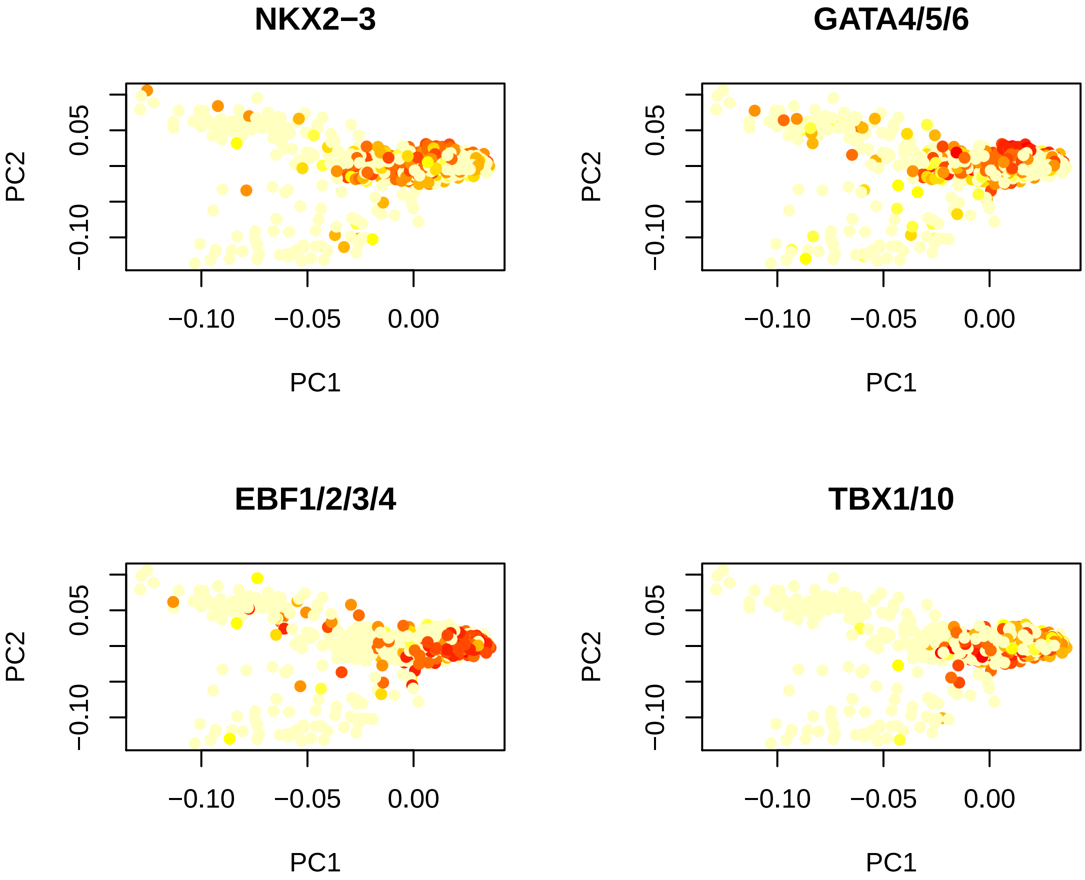


```r
# PC2 Mesenchymal Contamination
doHeatMap(hpfall.remv1, remove.key = T, slim.col.label = T, genes.use = pcTopGenes(hpfal
    2), cells.use = pcTopCells(hpfall.remv1, 2), col.use = col)
```

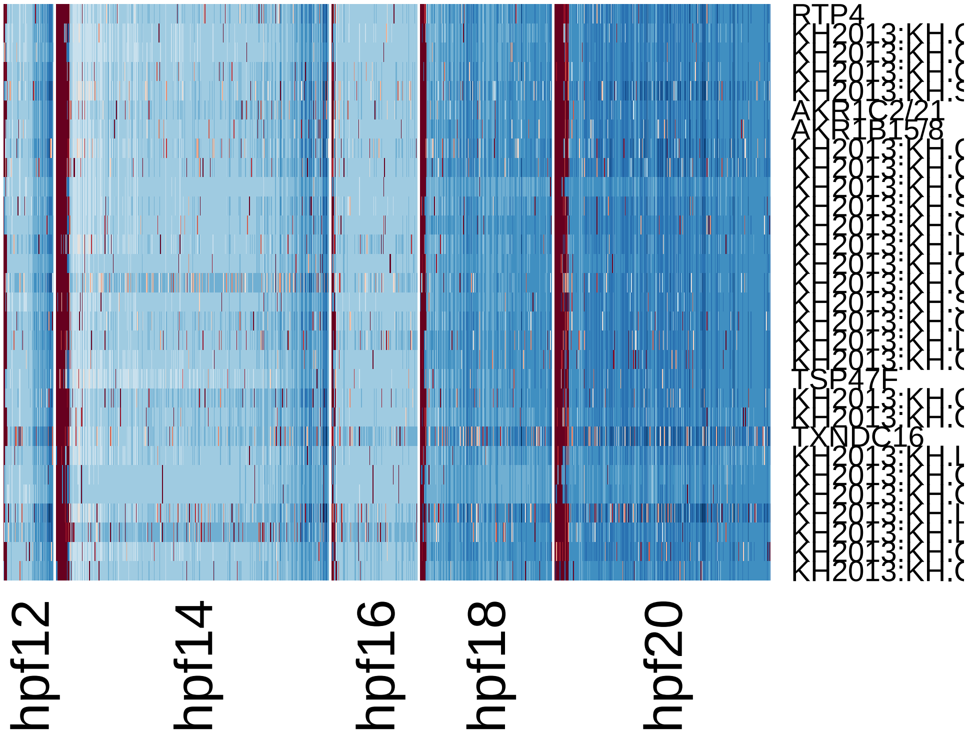


```
pca.plot(hpfall.remv1, 1, 2)
```

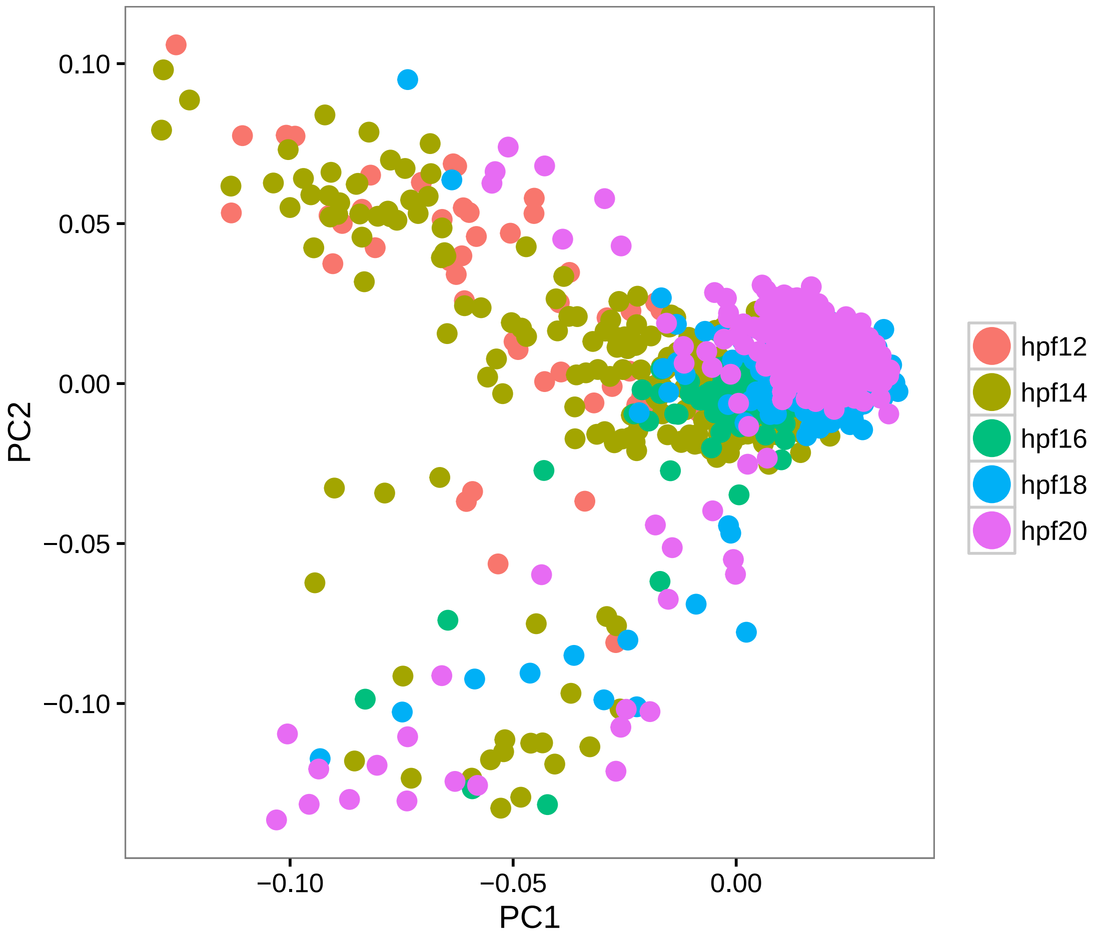


```
feature.plot(hpfall.remv1, c("RTP4", "TWIST1"), reduction.use = "pca")
```

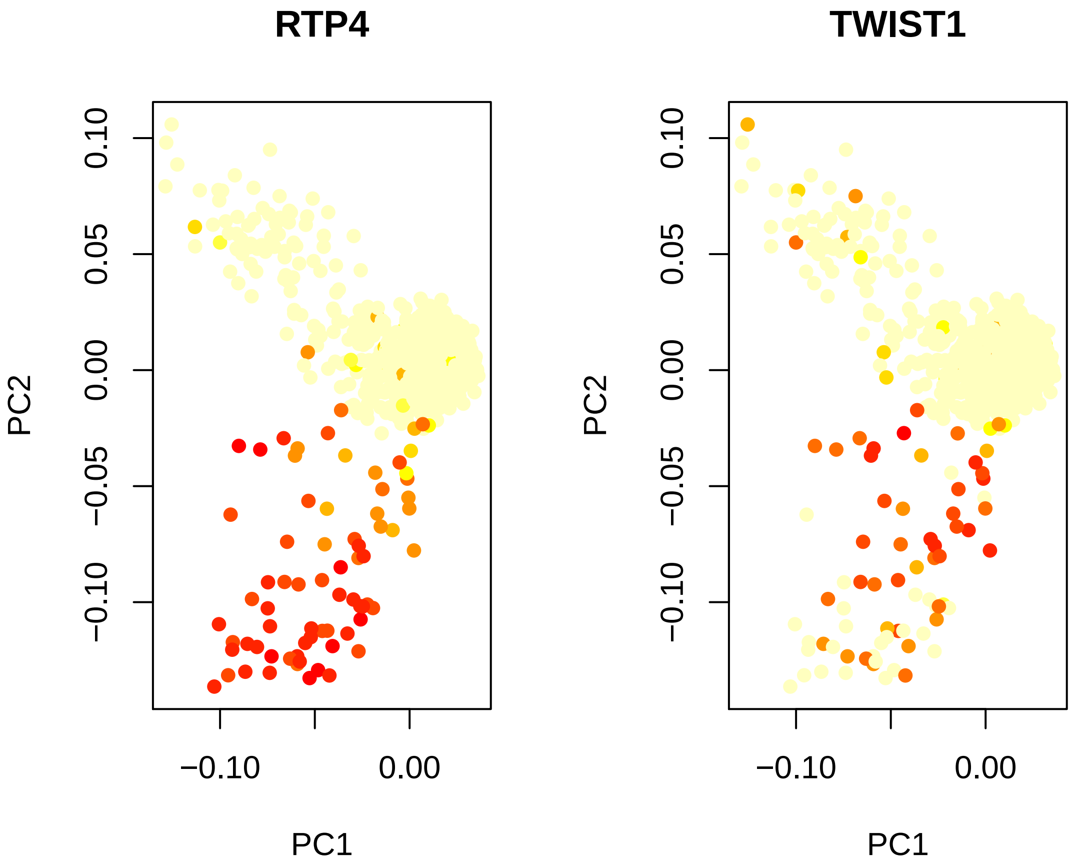


```
# PC3 Heart vs Muscle split
doHeatMap(hpfall.remv1, remove.key = T, slim.col.label = T, genes.use = pcTopGenes(hpfal
    3, do.balanced = T), cells.use = pcTopCells(hpfall.remv1, 3), col.use = col)
```

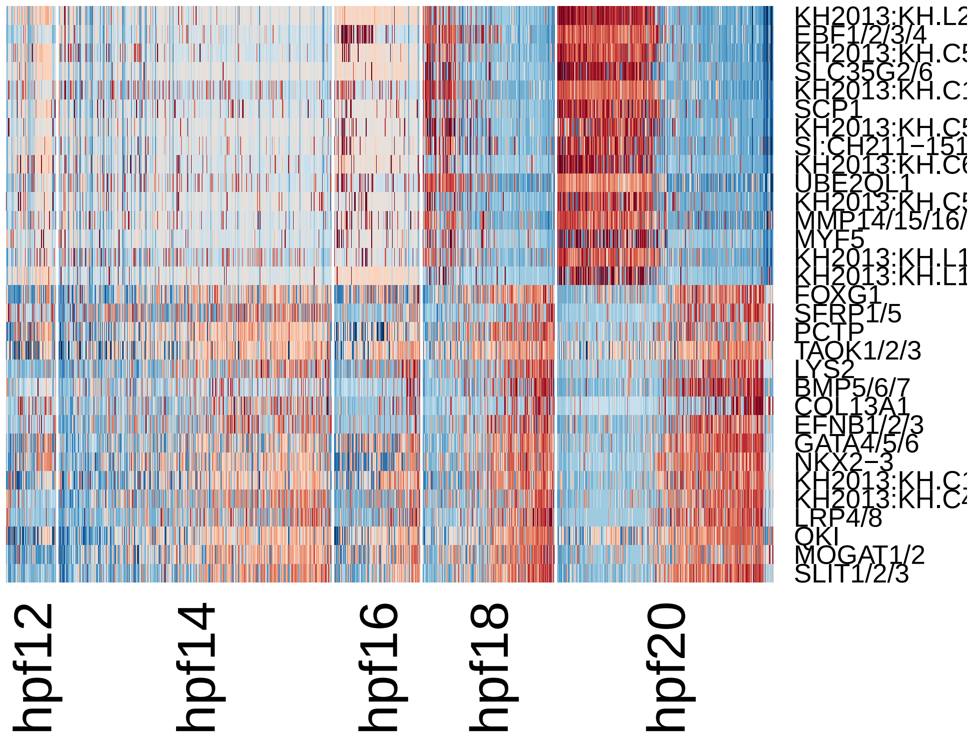


```r
pca.plot(hpfall.remv1, 1, 3)
```

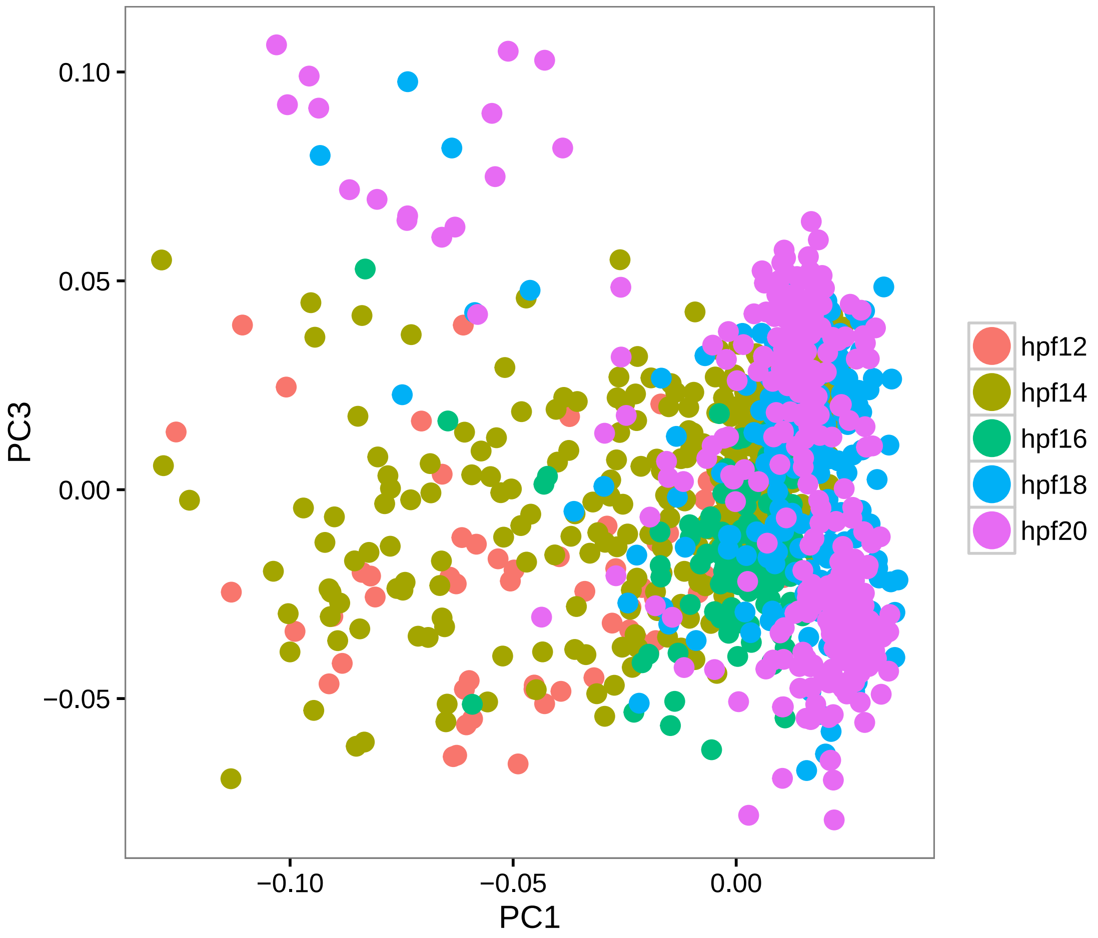


```r
feature.plot(hpfall.remv1, c("NKX2-3", "EBF1/2/3/4", "TBX1/10", "GATA4/5/6"),
    reduction.use = "pca", dim.2 = 3)
```

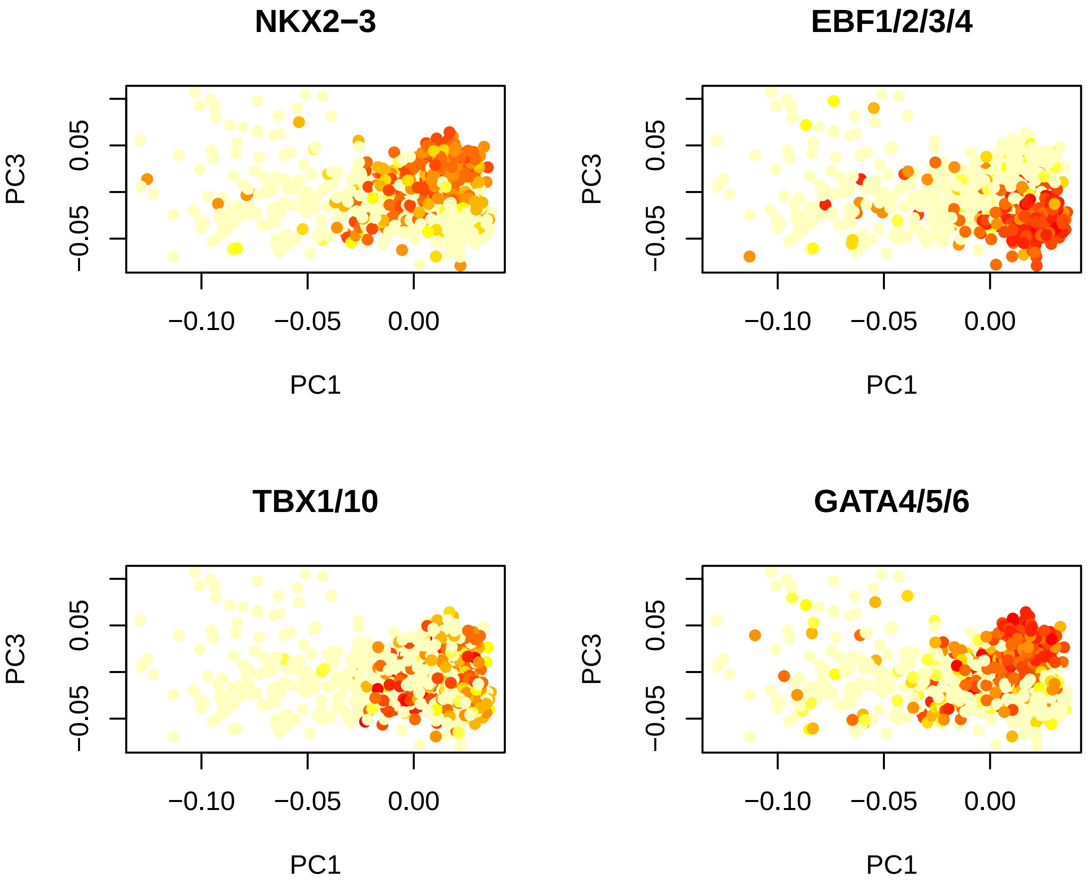


```
# PC4 Saperation by muscle/heart (MYO10, NOVA1/2) and technical noise
doHeatMap(hpfall.remv1, remove.key = T, slim.col.label = T, genes.use = pcTopGenes(hpfal
    4), cells.use = pcTopCells(hpfall.remv1, 4), col.use = col)
```

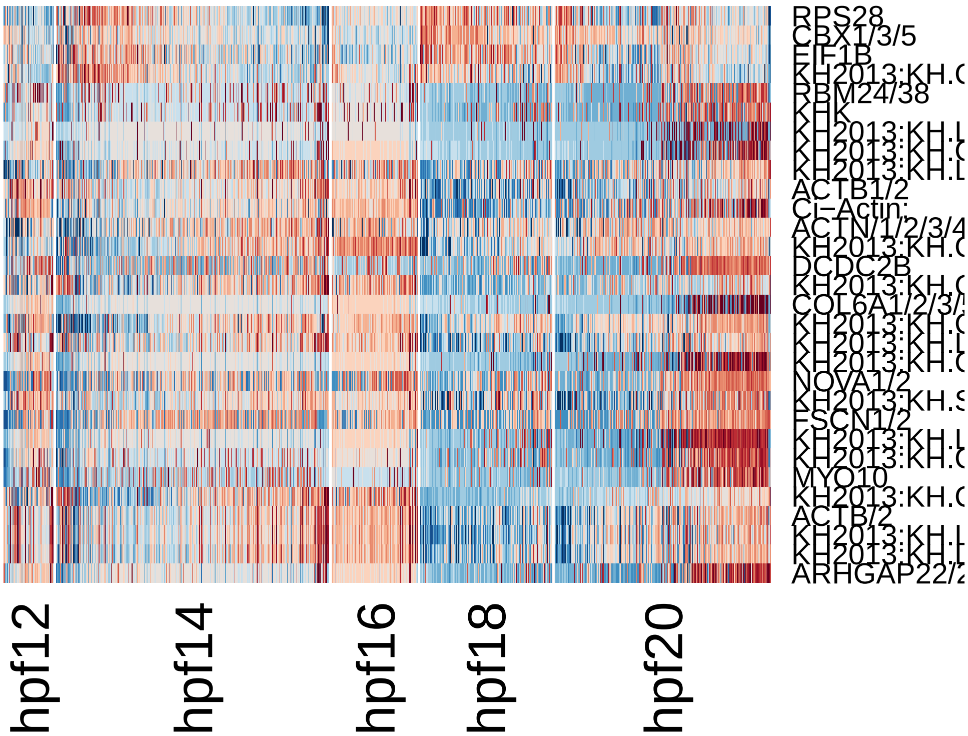


```
pca.plot(hpfall.remv1, 1, 4)
```

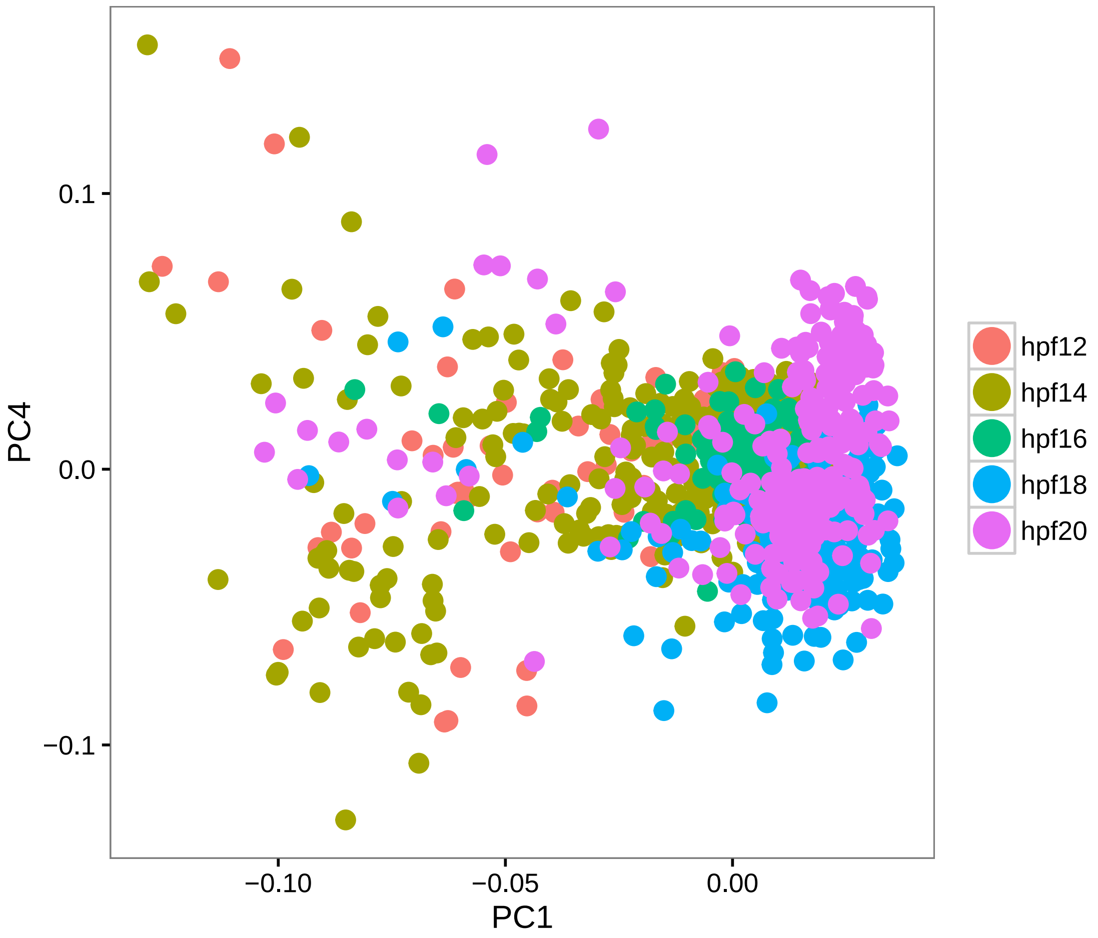


```
feature.plot(hpfall.remv1, c("NKX2-3", "EBF1/2/3/4", "TBX1/10", "GATA4/5/6"),
    reduction.use = "pca", dim.2 = 4)
```

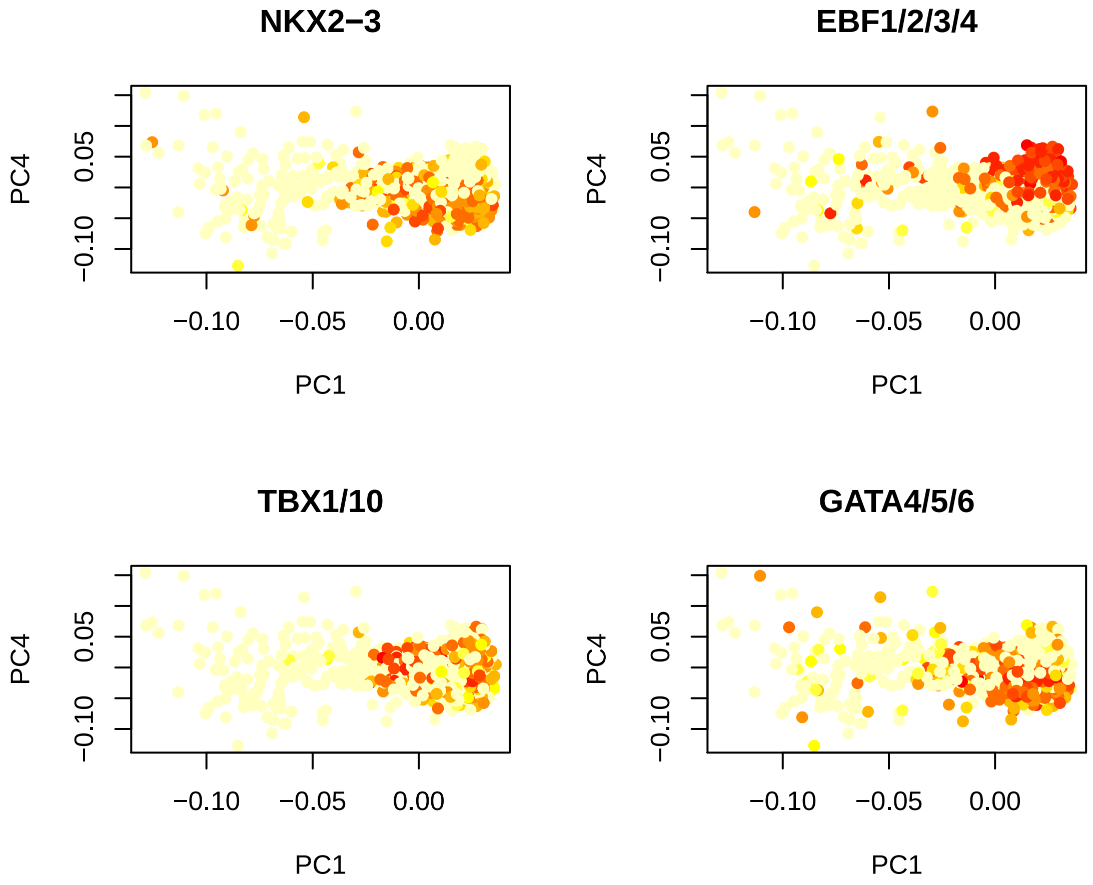


```r
# PC5 Saperation by technical noise
doHeatMap(hpfall.remv1, remove.key = T, slim.col.label = T, genes.use = pcTopGenes(hpfal
    5), cells.use = pcTopCells(hpfall.remv1, 5), col.use = col)
```

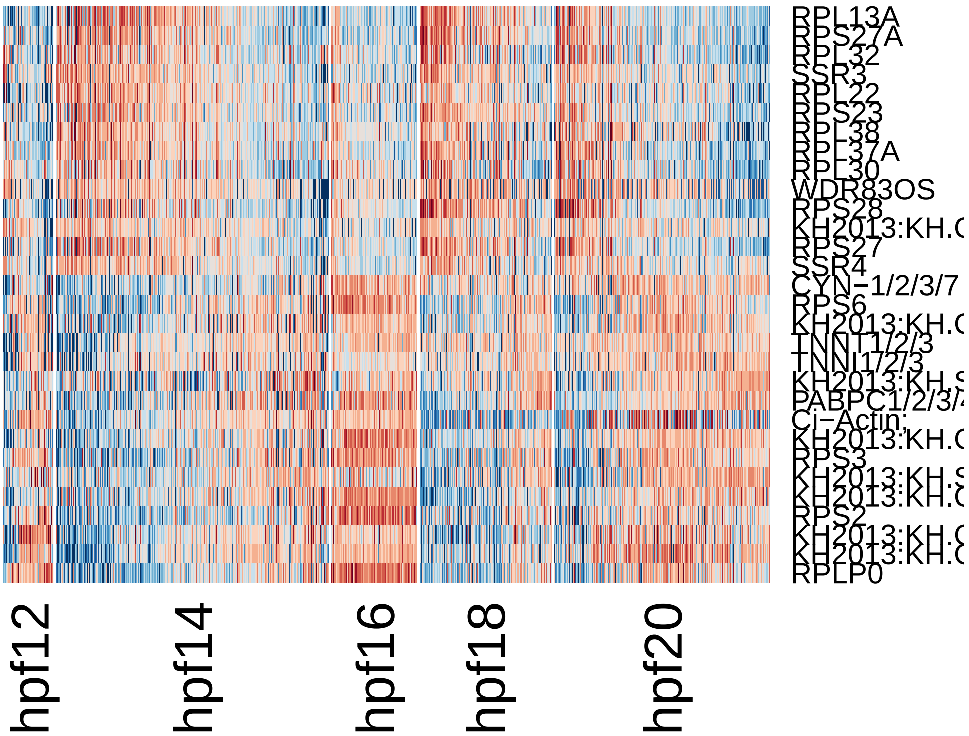


```
pca.plot(hpfall.remv1, 1, 5)
```

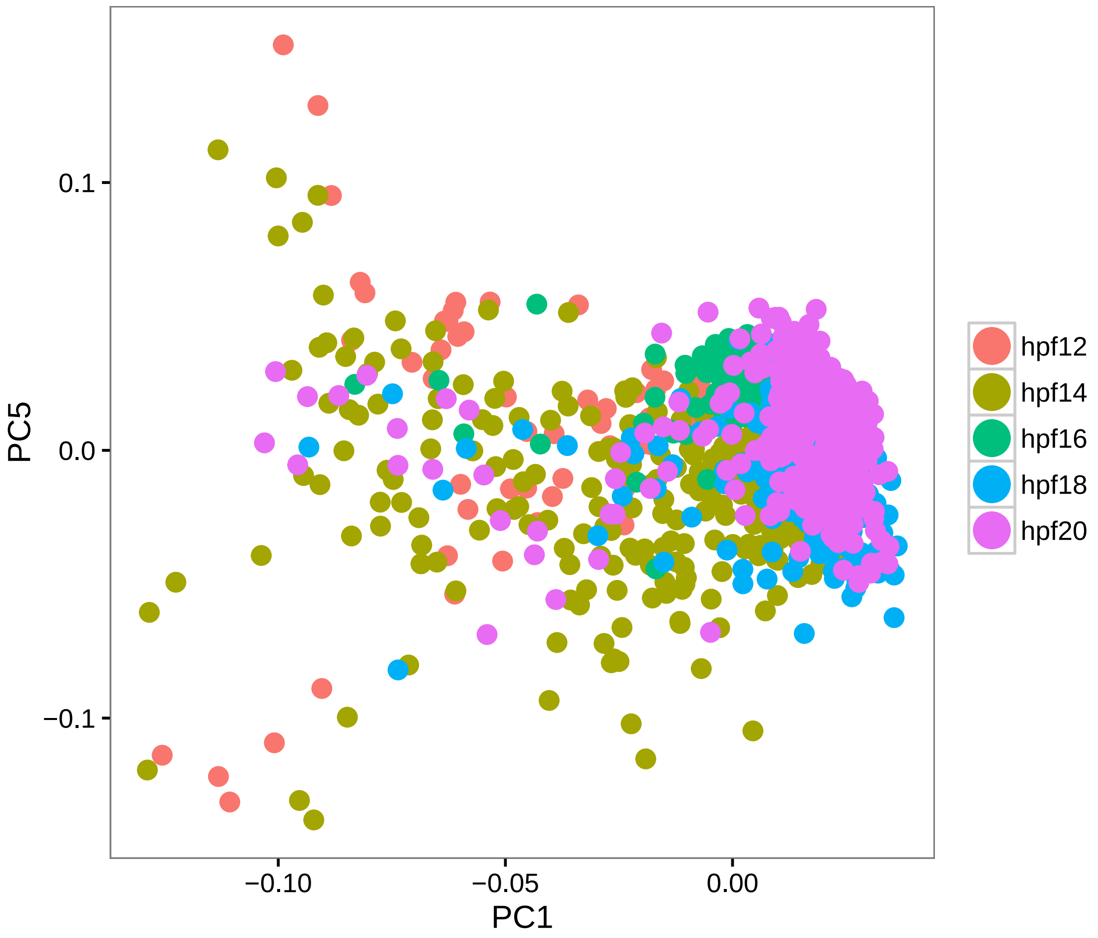


```
feature.plot(hpfall.remv1, c("NKX2-3", "EBF1/2/3/4", "TBX1/10", "GATA4/5/6"),
    reduction.use = "pca", dim.2 = 5)
```

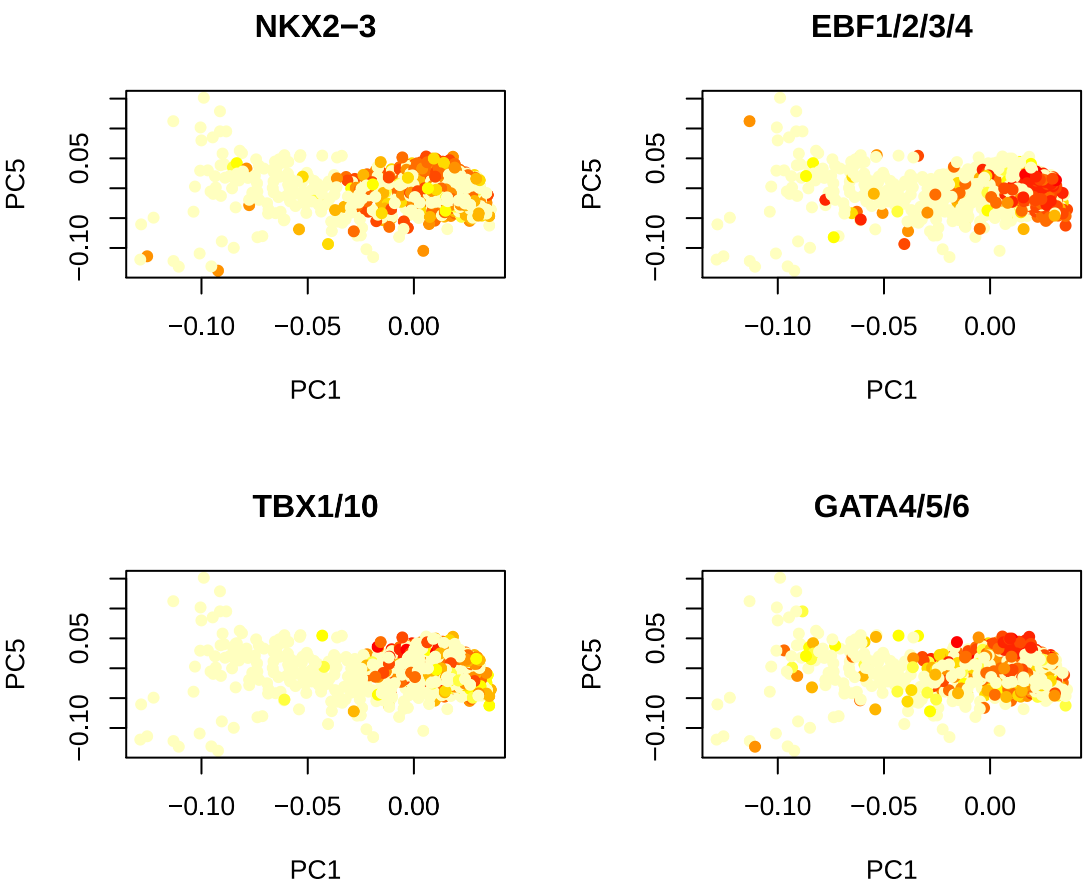


```
# Run tSNE using PC1-3 as input (spectral tSNE), we get distinct point
# clouds
hpfall.remv1 = run_tsne(hpfall.remv1, max_iter = 2000, dims.use = 1:3)
tsne.plot(hpfall.remv1, do.label = T, label.pt.size = 1)
```

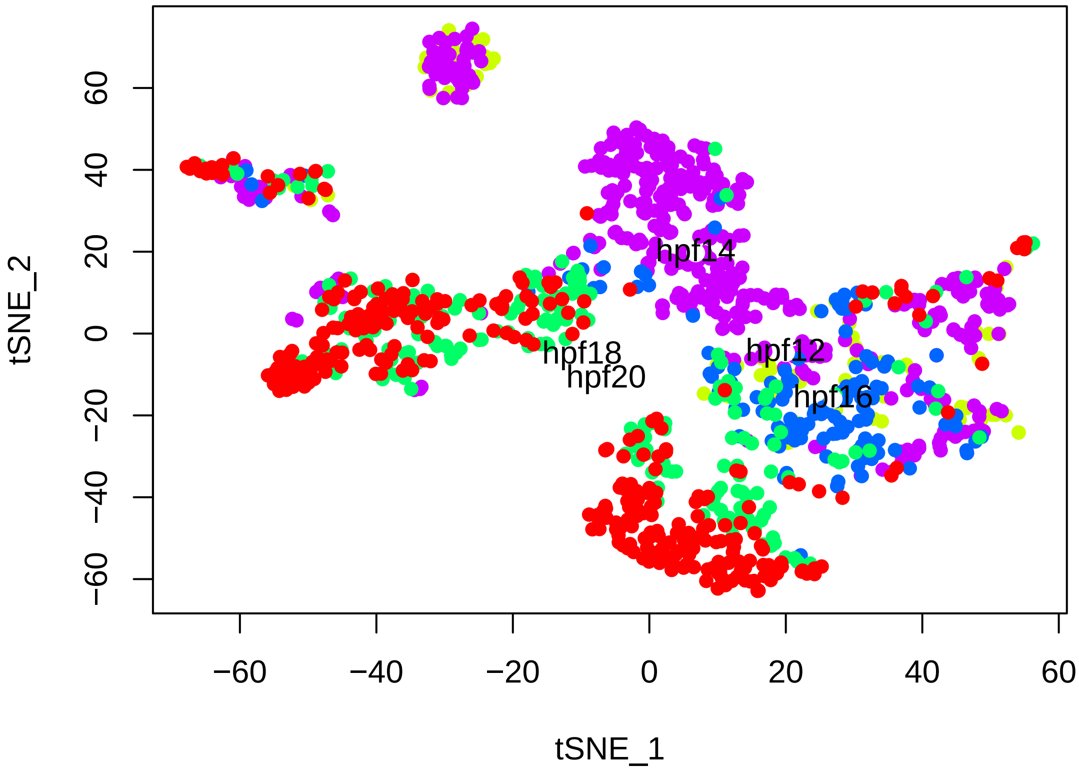


```
# Find cell clusters using Modularity optimization cluster detection.
# k.param = 15 for optimization of detecting small subset of comtamination
# cells.
hpfall.remv1 = FindClusters(hpfall.remv1, pc.use = 1:3, do.modularity = T, resolution =
    prune.SNN = 0.1, print.output = 0, k.param = 15)
```

```
## [1] "SNN : processed 284 cells"
## [1] "SNN : processed 569 cells"
## [1] "SNN : processed 854 cells"
## [1] "SNN : processed 1138 cells"
```

```
tsne.plot(hpfall.remv1, do.label = T, label.pt.size = 1)
```

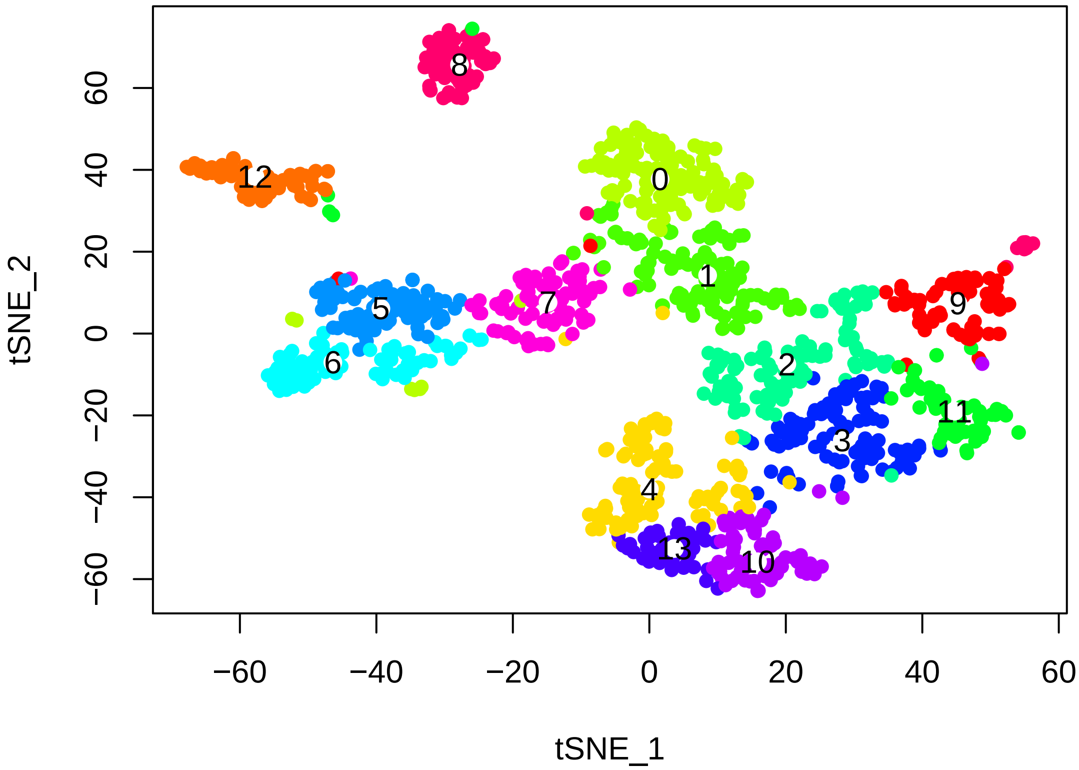


```
# The non-TVC contamination cells are located in cluster 8,9,11,12
doHeatMap(hpfall.remv1, remove.key = T, slim.col.label = T, genes.use = pcTopGenes(hpfal
    1), cells.use = pcTopCells(hpfall.remv1, 1), col.use = col)
```

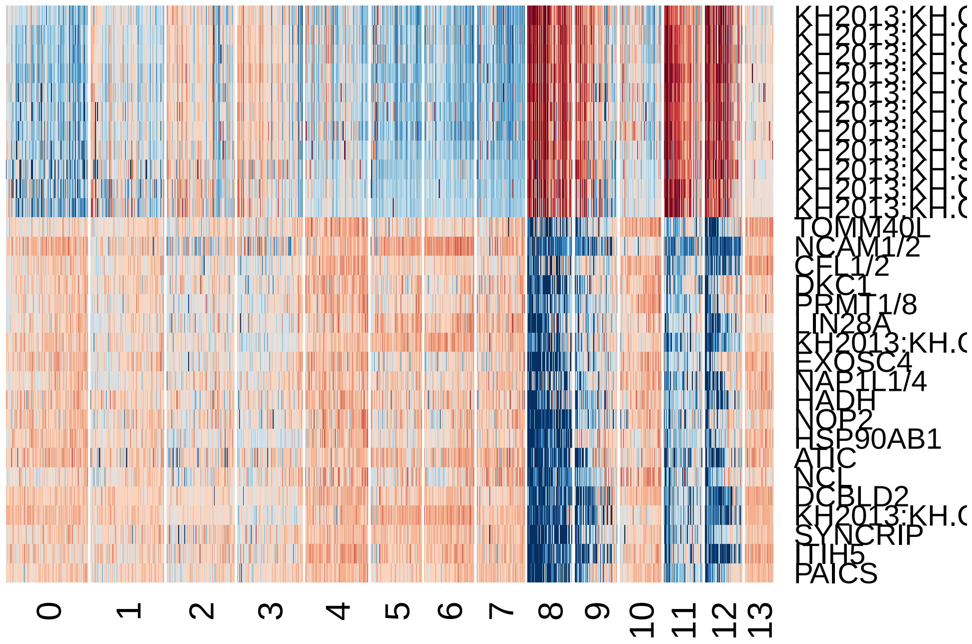

```r
# The mesenchymal contamination cells are located in cluster 12
doHeatMap(hpfall.remv1, remove.key = T, slim.col.label = T, genes.use = pcTopGenes(hpfal
    2), cells.use = pcTopCells(hpfall.remv1, 2), col.use = col)
```

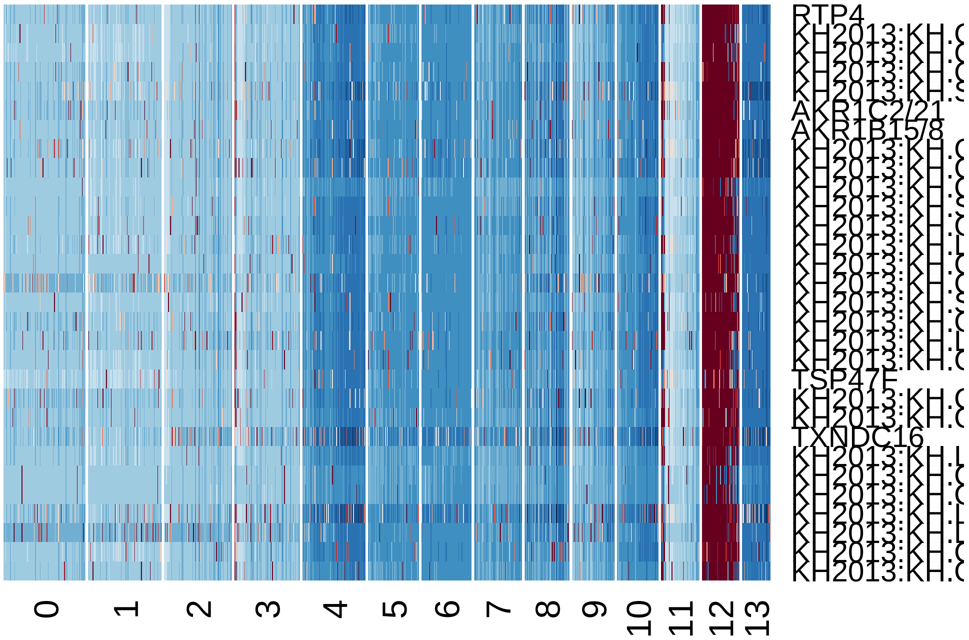


```r
# Find markers for these cells Find cluster markers using ROC test with
# thresh.use = 1, min.pct = 0.5 The ROC test returns the 'classification
# power' for any individual marker (ranging from 0 - random, to 1 -
# perfect). Though not a statistical test, it is often very useful for
# finding clean markers.

# Find markers for mesenchymal cells
mesen.marker = find.markers(hpfall.remv1, 12, thresh.use = 1, test.use = "roc",
    min.pct = 0.5)
head(mesen.marker[order(mesen.marker$myAUC, decreasing = T), ], 20)
```

```
##                            myAUC avg_diff power pct.1 pct.2
## AKR1B15/8                  0.998 5.261399 0.996 1.000 0.044
## KH2013:KH.L122.14_JMJD5    0.997 4.569135 0.994 1.000 0.241
## EGLN3                      0.997 4.312634 0.994 1.000 0.305
## KH2013:KH.C1.439_CG14331   0.996 3.798124 0.992 1.000 0.582
## KH2013:KH.C14.521_JMJD5/7  0.996 5.603266 0.992 1.000 0.065
## RTP4                       0.996 4.621945 0.992 1.000 0.038
## KH2013:KH.C1.561_TUB1/3    0.994 4.178222 0.988 1.000 0.048
## KH2013:KH.L18.67           0.994 4.708396 0.988 1.000 0.238
## KH2013:KH.C4.120           0.993 3.867655 0.986 1.000 0.501
## KH2013:KH.C6.268_JMJD7     0.991 4.448726 0.982 1.000 0.205
```

```
## AKR1C2/21                  0.989 5.118831 0.978 0.983 0.056
## TXNDC16                    0.988 5.751290 0.976 0.983 0.133
## KH2013:KH.C5.228_JMJD5     0.988 5.183405 0.976 0.983 0.035
## KH2013:KH.L154.6_JMJD5     0.985 3.895723 0.970 0.983 0.305
## KH2013:KH.C4.618           0.982 4.715708 0.964 0.949 0.101
## KH2013:KH.S936.1_JMJD5     0.981 5.585791 0.962 0.966 0.076
## KH2013:KH.C9.326_JMJD5     0.980 5.946468 0.960 0.966 0.066
## KH2013:KH.S936.3_JMJD5     0.980 4.861421 0.960 0.966 0.126
## KH2013:KH.C3.921_JMJD5/7   0.979 5.033655 0.958 0.966 0.042
## KH2013:KH.C1.30_CG14331    0.977 3.816522 0.954 0.983 0.359
```

```
write.table(which.cells(hpfall.remv1, 12), "mesen.cellname.txt")
write.table(mesen.marker, "mesenchymal.marker.txt", row.names = T, col.names = NA,
    sep = "\t")

# Find markers for the other group of cells by comparing to mesenchymal
contam.marker = find.markers(hpfall.remv1, c(8, 9, 11), 12, thresh.use = 1,
    test.use = "roc", min.pct = 0.5)
head(contam.marker[order(contam.marker$myAUC, decreasing = T), ], 20)
```

```
##                                                myAUC avg_diff power pct.1
## SDC/1/2/4                                      0.897 2.491357 0.794 0.924
## CTNNB1                                         0.890 2.178370 0.780 0.960
## FARP1/2                                        0.861 2.957062 0.722 0.763
## KH2013:KH.C6.26_GJA4                           0.835 1.883192 0.670 0.949
## CD151                                          0.825 2.937783 0.650 0.722
## KH2013:KH.C11.378_CLDN1/10/14/19/2/3/4/5/6/7/9 0.822 2.707400 0.644 0.838
## FOXH1                                          0.821 2.963121 0.642 0.727
## KH2013:KH.C7.805_ID1/2/3/4                     0.803 2.472959 0.606 0.833
## KH2013:KH.L154.1_LRP5/6                        0.796 2.110253 0.592 0.631
## MAP1LC3C                                       0.792 2.678988 0.584 0.657
## LAMB1/2/3/4                                    0.792 1.649389 0.584 0.828
## Ci-Col12A1                                     0.791 2.145317 0.582 0.823
## KH2013:KH.L20.63                               0.791 1.213074 0.582 0.995
## KH2013:KH.C1.738_CG9550                        0.790 2.475260 0.580 0.717
## TIE1/2                                         0.789 1.132390 0.578 0.995
## CDH1/13/15/2/3/4                               0.786 2.108102 0.572 0.707
## ARVCF                                          0.785 2.861593 0.570 0.631
## CFL1/2                                         0.784 1.644946 0.568 0.828
## KH2013:KH.C14.71                               0.782 1.112152 0.564 0.970
## ADD1/2/3                                       0.781 2.614835 0.562 0.636
##                                                pct.2
## SDC/1/2/4                                      0.407
## CTNNB1                                         0.508
```

```
## FARP1/2                                      0.169
## KH2013:KH.C6.26_GJA4                         0.763
## CD151                                        0.153
## KH2013:KH.C11.378_CLDN1/10/14/19/2/3/4/5/6/7/9 0.407
## FOXH1                                        0.153
## KH2013:KH.C7.805_ID1/2/3/4                   0.508
## KH2013:KH.L154.1_LRP5/6                      0.119
## MAP1LC3C                                     0.119
## LAMB1/2/3/4                                  0.441
## Ci-Col12A1                                   0.458
## KH2013:KH.L20.63                             0.983
## KH2013:KH.C1.738_CG9550                      0.237
## TIE1/2                                       0.983
## CDH1/13/15/2/3/4                             0.237
## ARVCF                                        0.102
## CFL1/2                                       0.610
## KH2013:KH.C14.71                             0.898
## ADD1/2/3                                     0.153
```

```
write.table(which.cells(hpfall.remv1, c(8, 9, 11)), "contam.cellname.txt")
write.table(contam.marker, "contam.marker.txt", row.names = T, col.names = NA,
    sep = "\t")

# Remove contamination and mesenchymal cells
hpfall.remv2 = subsetData(hpfall.remv1, which.cells(hpfall.remv1, c(0:7, 10,
    13)), do.scale = F)
hpfall.remv2
```

```
## An object of class seurat in project allhpf
##  14864 genes across 881 samples.
```

```
# Restore hpf labels
hpfall.remv2@ident = factor(substring(hpfall.remv2@cell.names, 1, 5))
names(hpfall.remv2@ident) = hpfall.remv2@cell.names

# Gene detected in each time point
boxPlot.FPKM(hpfall, "nGene", name.y = "Number of Genes", name.x = "", ratio.plot = 0.00
    name = "Gene Detected")
```

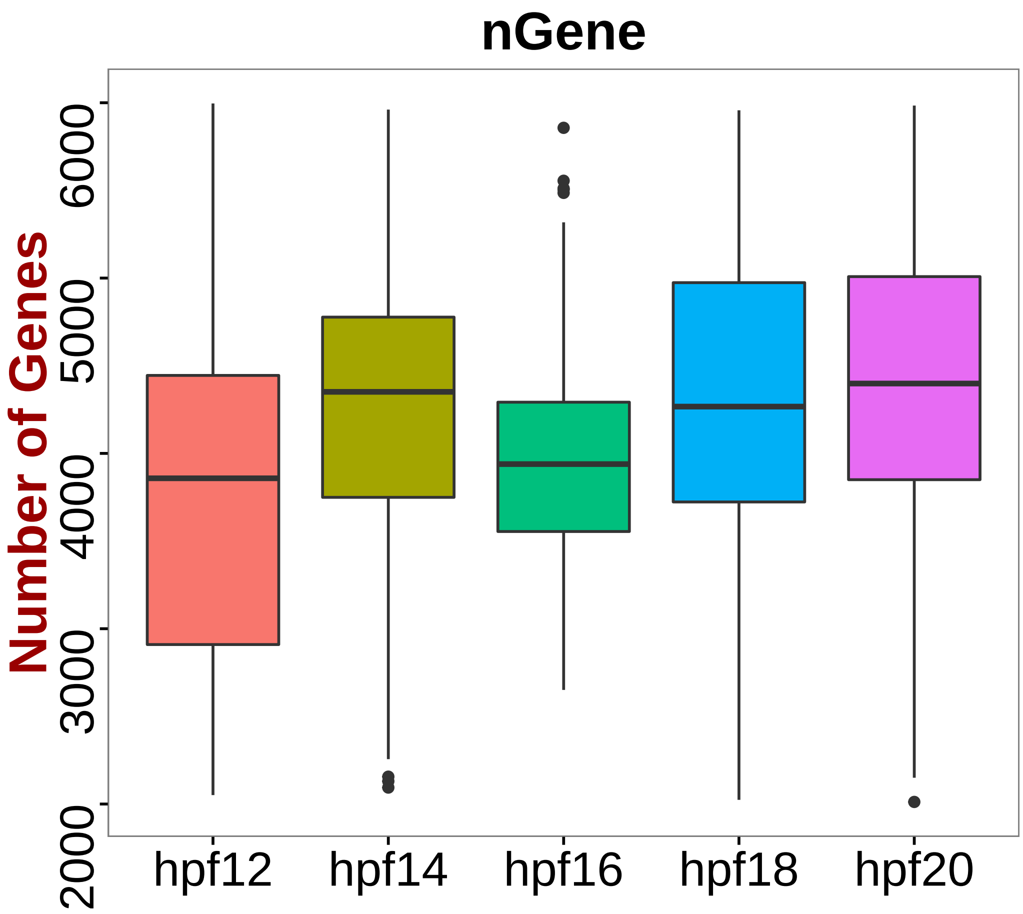


```r
# Contamination cells are located
save(hpfall.remv2, file = "hpfall.remv2.Robj")
```
