## Supplementary material for "A single cell transcriptional roadmap for cardiopharyngeal fate diversification": Markdown_Trajectory

### Trajectory Analysis

*Xiang Niu June 3, 2016*

1. Load Data and Packages

```
library(Seurat)
library(diffusionMap)
library(princurve)
library(pbapply)
library(RColorBrewer)
source("functions.R")
load("hpfallCluster.Robj")
col = colorRampPalette(rev(brewer.pal(n = 10, name = "RdBu")))
```

2. SHP Trajectory

```
# Select all shp markers
all.marker.files = list.files(pattern = "markers")
all.markers.shp = c()
for (i in grep("TVC.markers|STVC|SHP", all.marker.files, value = T)) {
    my.txt = read.table(i, sep = "\t", header = T)
    all.markers.shp = unique(c(all.markers.shp, rownames(subset(my.txt, power >
        0.3))))
}
length(all.markers.shp)
```

```
## [1] 742
```

```
# Retrieve SHP trajectory
shp = subsetData(hpfall.cluster, which.cells(hpfall.cluster, c("12TVC", "14STVC",
    "16SHP", "18SHP", "20SHP")), do.scale = F)
shp@ident = factor(shp@ident, levels = c("12TVC", "14STVC", "16SHP", "18SHP",
    "20SHP"), ordered = T)

# Store real time label
[, "real.time"] = as.numeric(substr(shp@ident, 1, 2))

# Calculate distance matrix for diffusion map based on selected shp
# trajectory markers
shp.dist = as.matrix(dist(t(shp@scale.data[all.markers.shp, ])))
```

```r
# Run diffusion map and return top 50 dimensions
set.seed(1)
shp.diff = diffuse(shp.dist, maxdim = 50)
```

```
## Performing eigendecomposition
## Computing Diffusion Coordinates
## Used default value: 50 dimensions
## Elapsed time: 0.31 seconds
```

```r
# Diffusion map eigen values
plot(shp.diff$eigenvals, type = "o", col = "blue", ylab = " Eigenvalues", xlab = "Eigenv
    main = "Diffusion Map Eigenvalues", lwd = 2, cex.lab = 1.5, cex.main = 1.5)
```

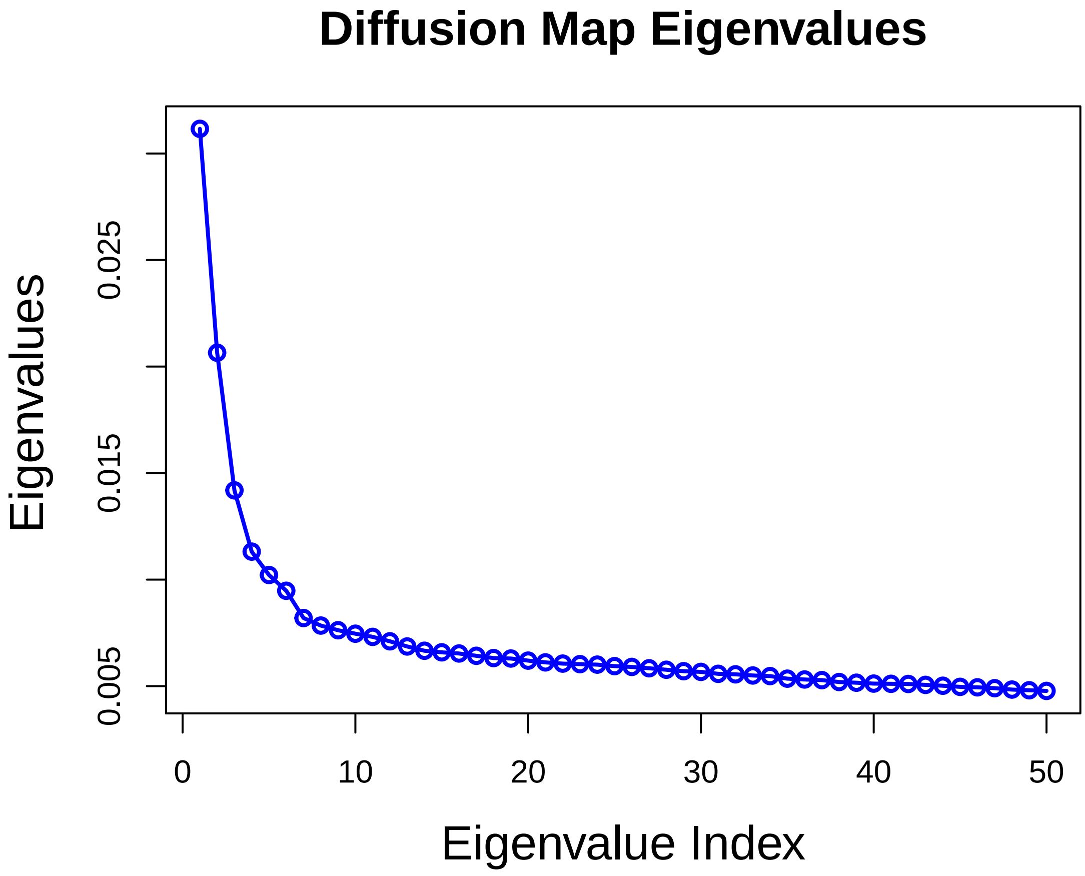


```r
# Save first two diffusion map coordinators
shp@tsne.rot[1:2] = data.frame(shp.diff$X[, 1:2], row.names = shp@cell.names)
colnames(shp@tsne.rot) = c("tSNE_1", "tSNE_2")

# Visualize top two diffusion map components
tsne.pseudo(shp, do.label = F, label.cex.text = 1, name.y = "Diffusion Map Coordinator 2
    name.x = "Diffusion Map Coordinator1", label.cols.use = c("green", "yellow",
```

```
        "orange1", "orange4", "orange4"), label.pt.size = 1.5, xlim = c(-0.1,
        0.05), ylim = c(-0.05, 0.05))
legend("topleft", legend = c("12TVC", "14STVC", "16SHP", "18SHP", "20SHP"),
    col = c("green", "yellow", "orange1", "orange4", "orange4"), pch = 16, cex = 0.5,
    pt.cex = 1)

# Fit the first two diffusion map components with principal curve
shp.princurve = principal.curve(as.matrix(shp.diff$X[, 1:2]), start = as.matrix(shp.dif
    1:2]))
lines(shp.princurve$s[order(shp.princurve$lambda), ], lty = 1, lwd = 4, col = "purple",
    type = "l")
```

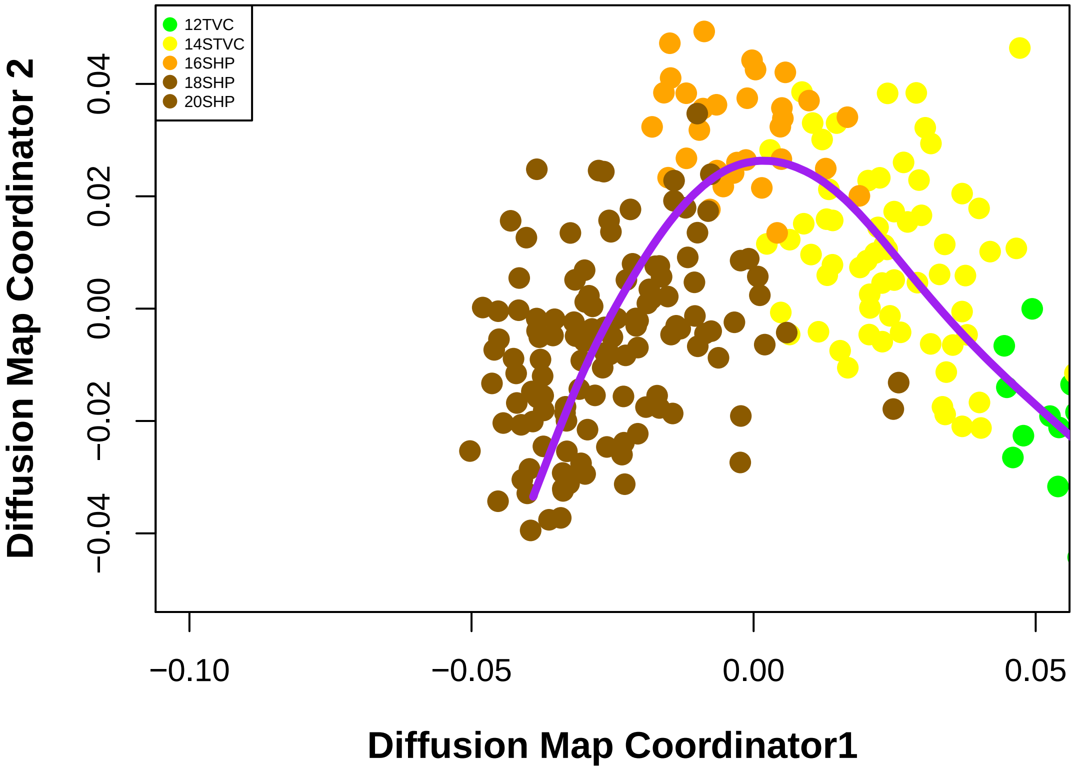


```
df = data.frame(shp.princurve$s[order(shp.princurve$lambda), ])
colnames(df) = c("x", "y")
ggplot(data = df, aes(x, y)) + geom_line(size = 1.5, colour = "black") + geom_density2d
    bins = 6) + scale_colour_gradient(low = "darkgray", high = "white", 3) +
    xlim(-0.084, 0.08) + ylim(-0.05, 0.05) + geom_point(data = data.frame(shp@tsne.rot,
    color = shp@ident), aes(tSNE_1, tSNE_2), size = 2, color = c(rep("green",
    table(shp@ident)[1]), rep("yellow", table(shp@ident)[2]), rep("orange1",
    table(shp@ident)[3]), rep("orange4", table(shp@ident)[4]), rep("orange4",
    table(shp@ident)[5]))) + theme_classic() + theme(legend.position = "none",
    axis.title = element_text(size = rel(1)), axis.text = element_blank(), axis.ticks =
    axis.line.x = element_line(colour = "black", size = 0.5, linetype = "solid"),
```

```r
    axis.line.y = element_line(colour = "black", size = 0.5, linetype = "solid")) +
    xlab(label = "Diffusion Component 1") + ylab(label = "Diffusion Component 2") +
    geom_line(size = 1.5, colour = "black")
```

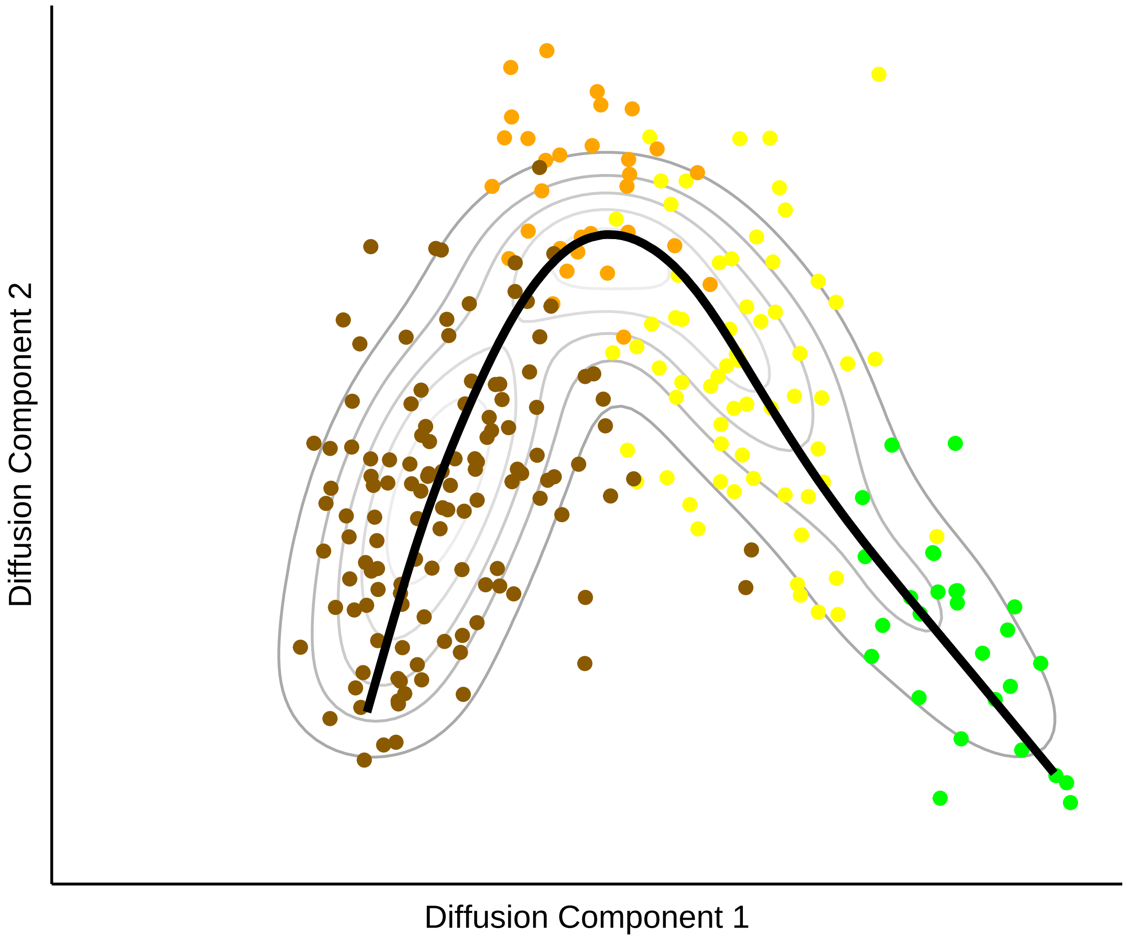


```r
# Pseudotime score is assigned to each cell as projection on principal curve
[, "pseudo.time"] = shp.princurve$lambda

# draw pseudotime cell density
plot(NULL, xlim = c(-0.05, 0.2), ylim = c(0, 100), main = "", ylab = "", xlab = "",
    axes = F)
lines(density([which.cells(shp, "12TVC"), "pseudo.time"], adjust = 2),
    type = "h", col = "green")
lines(density([which.cells(shp, "14STVC"), "pseudo.time"], adjust = 2),
    type = "h", col = "yellow")
lines(density([which.cells(shp, c("16SHP")), "pseudo.time"], adjust = 2),
    type = "h", col = "orange1")
lines(density([which.cells(shp, c("18SHP")), "pseudo.time"], adjust = 2),
    type = "h", col = "orange2")
```

```
lines(density([which.cells(shp, c("20SHP")), "pseudo.time"], adjust = 2),
    type = "h", col = "orange3")
```

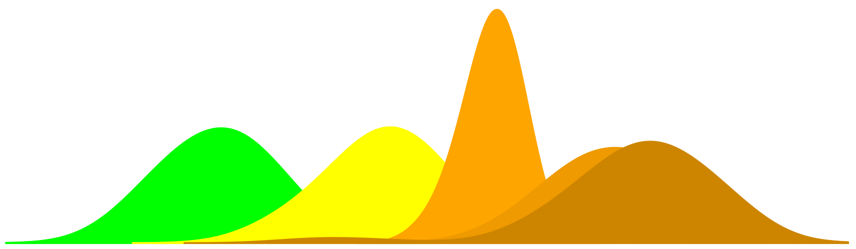

```
# Scale pseudotime into 0-1 range
shp.new = scale.pseudo(shp, "pseudo.time", do.scale = T, do.PC = T)

# Visualize markers on diffusion map
gene.viz = c("HAND/2", "HAND/1/2", "TBX1/10", "EBF1/2/3/4")
feature.plot.pseudo(shp.new, gene.viz, name.x = "SHP Trajectory", name.y = "",
    pt.size = 0.75, nCol = 2)
```

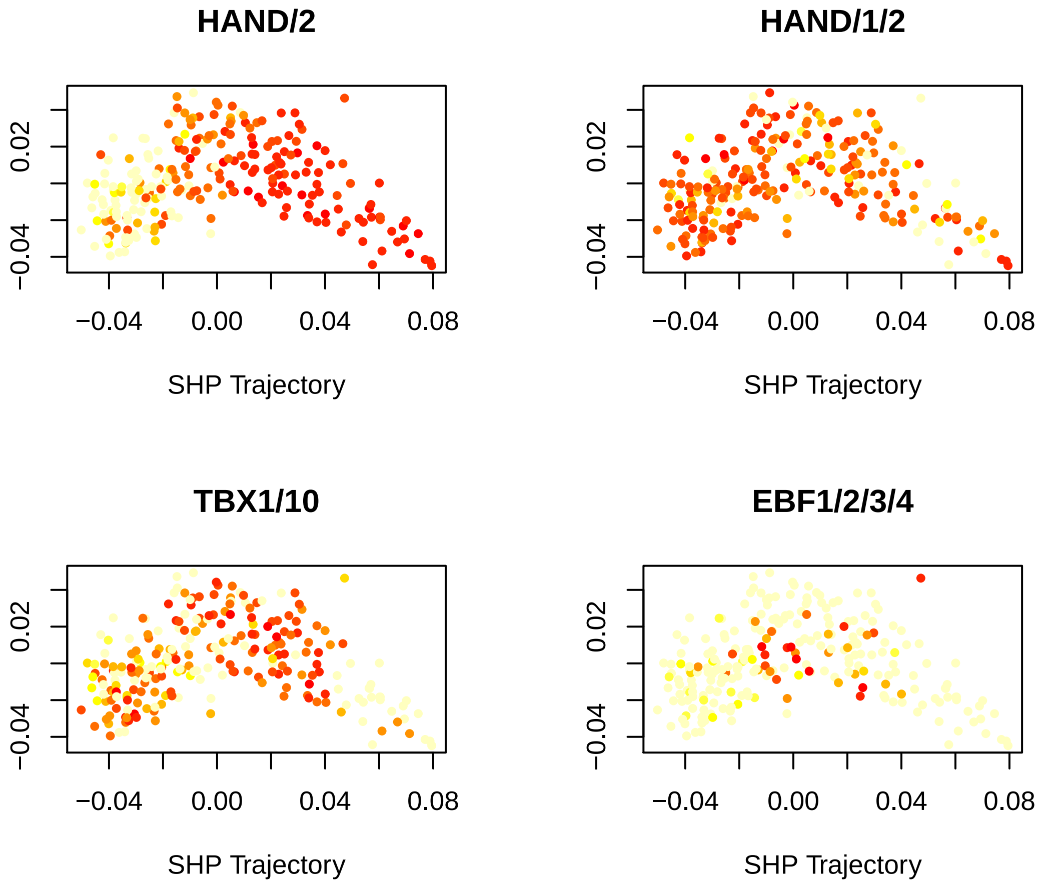

```r
par(mfrow = c(2, 2))
for (i in gene.viz) {
    genePlot.pseudo(shp.new, gene = i, col.use = c("green", "yellow", "orange1",
        "orange2", "orange3"), do.spline = T, name.x = "SHP Trajectory", cex.use = 0.8,
        cex.lab = 1, font.lab = 2)
}
```

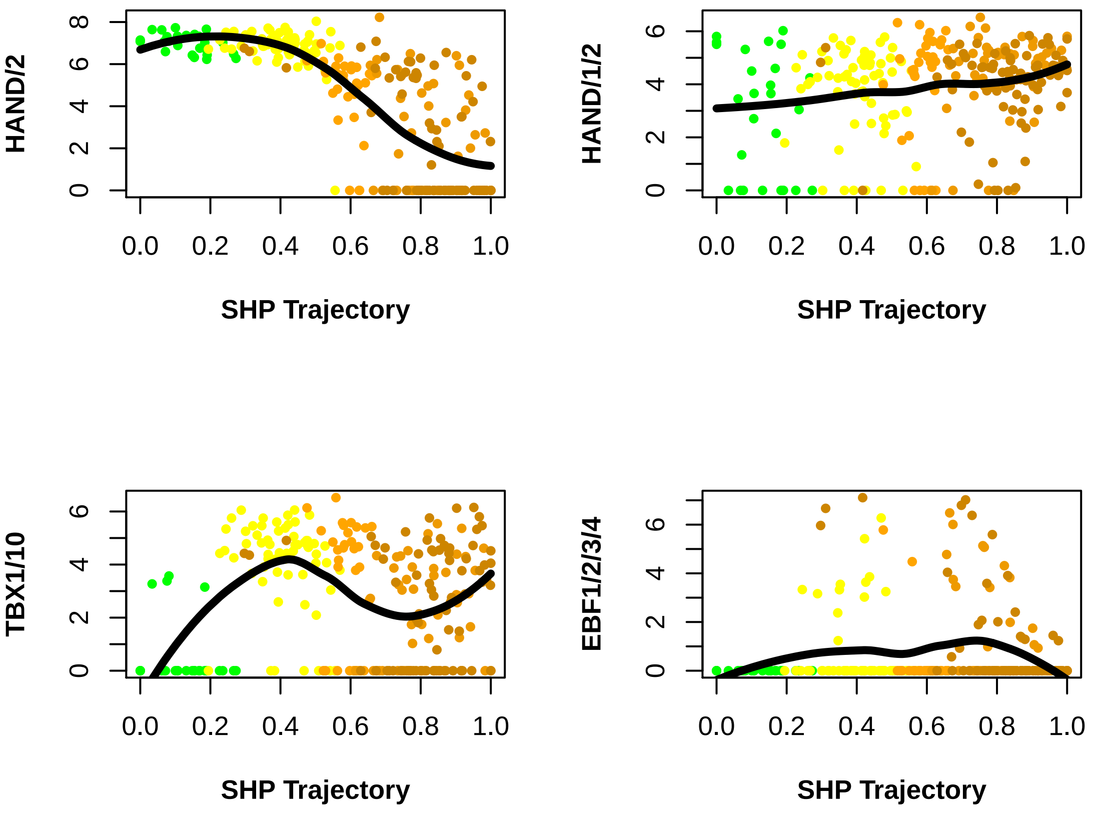


```r
# Pseudo time scores correlate with real time
vlnPlot(shp.new, "pseudo.time", size.x.use = 2)
```

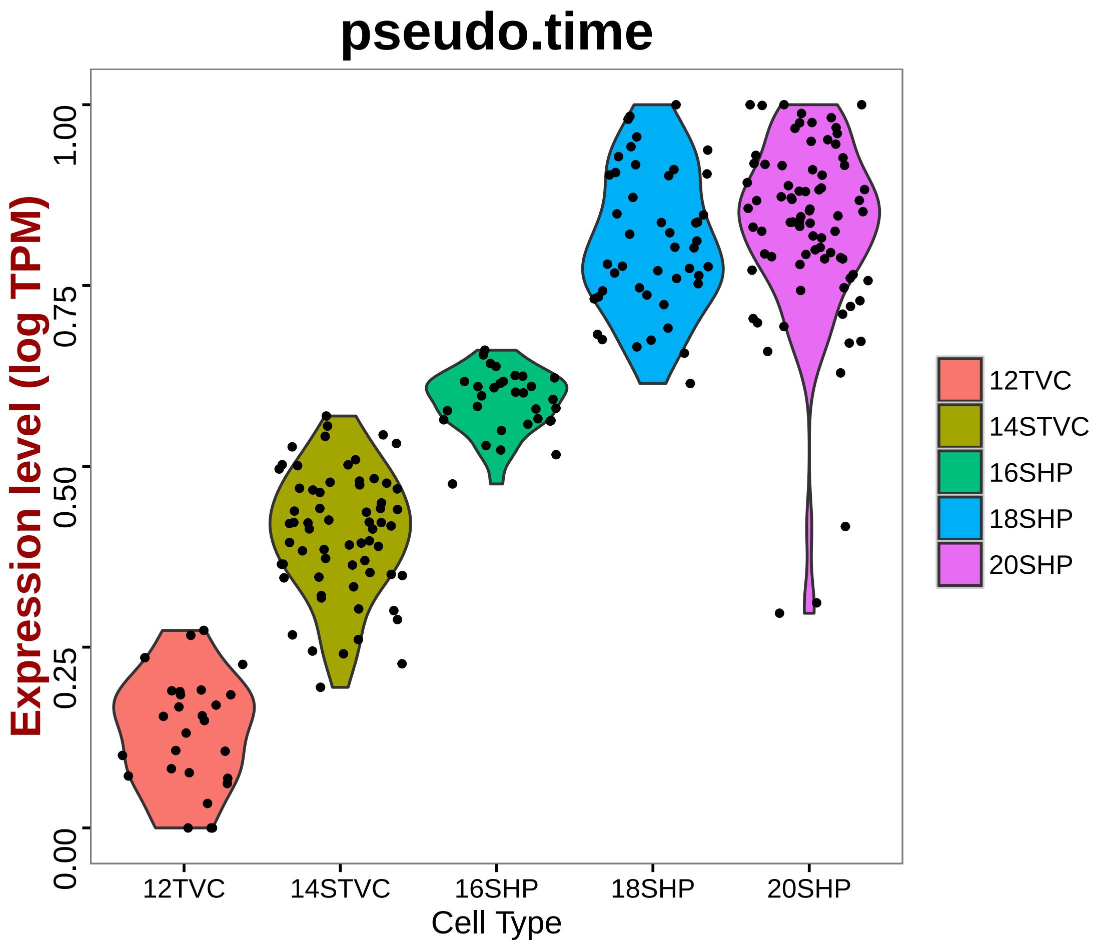


```
plt = vlnPlot(shp.new, "pseudo.time", size.x.use = 2, do.ret = T)
plt[[1]] + geom_boxplot(width = 0.25, outlier.size = 1, aes(fill = ident)) +
    geom_smooth(aes(group = 1), method = "loess", size = 1, se = F, col = "black",
        lty = 16) + annotate("text", 1.2, 0.9, label = paste("Correlation",
    round(cor(shp@data.info$real.time, shp.princurve$lambda), 2), sep = "\n"),
    size = 6, fontface = "bold")
```

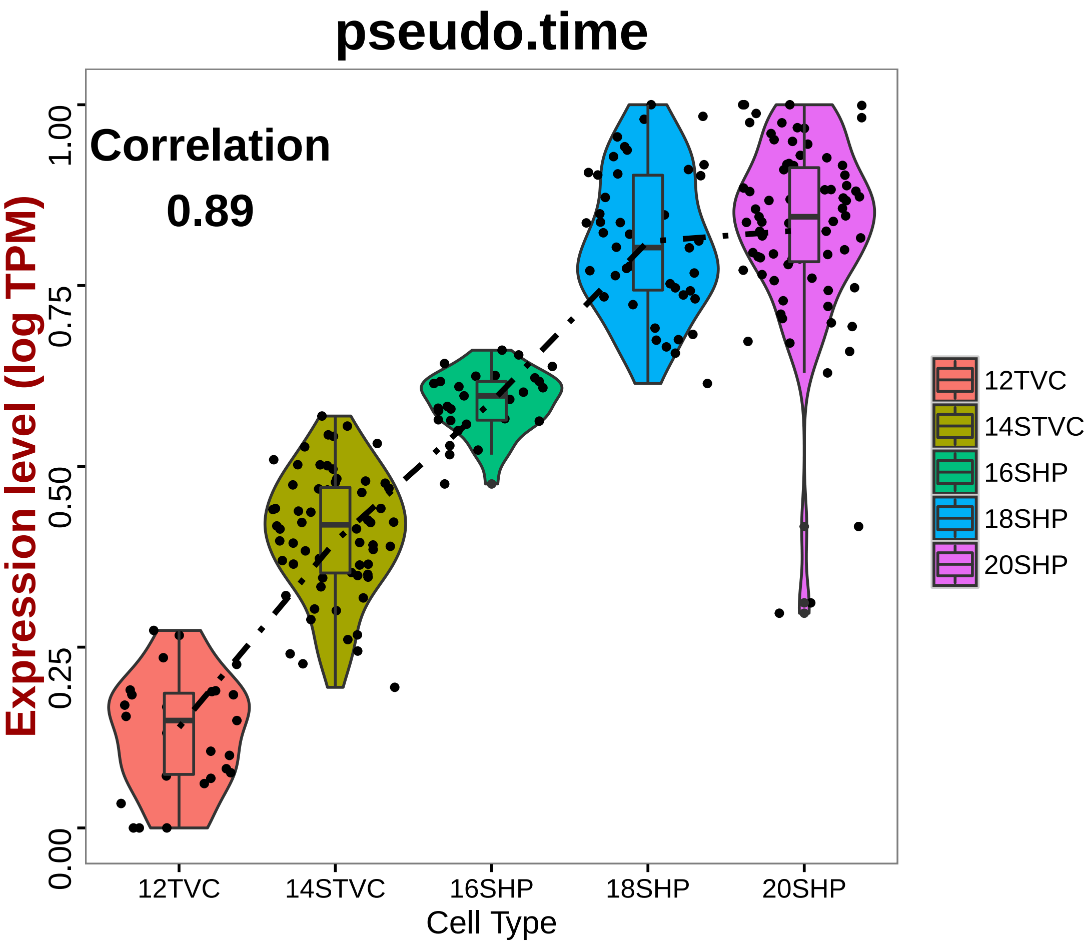


```
# Align genes by their induction time
shp.marker.ps = pseudo.gene.cluster(shp.new, genes.use = all.markers.shp)
doHeatMap(shp.new, genes.use = unlist(shp.marker.ps), remove.key = T, slim.col.label = F
    order.by.ident = F, draw.line = F, cells.use = pcTopCells(shp.new, 1), labCol = F,
    labRow = F, col.use = col)
```

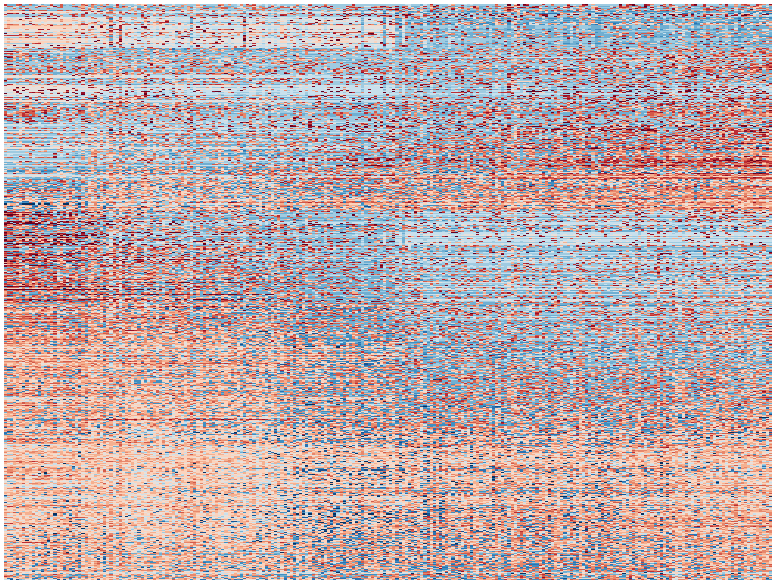

3. FHP Trajectory

```r
# Select all fhp markers
all.marker.files = list.files(pattern = "markers")
all.markers.fhp = c()
for (i in grep("TVC.markers|FHP", all.marker.files, value = T)) {
    my.txt = read.table(i, sep = "\t", header = T)
    all.markers.fhp = unique(c(all.markers.fhp, rownames(subset(my.txt, power >
        0.3))))
}
length(all.markers.fhp)
```

```
## [1] 811
```

```r
# Retrieve fhp trajectory
fhp = subsetData(hpfall.cluster, which.cells(hpfall.cluster, c("12TVC", "14FHP",
    "16FHP", "18FHP", "20FHP")), do.scale = F)
fhp@ident = factor(fhp@ident, levels = c("12TVC", "14FHP", "16FHP", "18FHP",
    "20FHP"), ordered = T)

# Store real time label
[, "real.time"] = as.numeric(substr(fhp@ident, 1, 2))

# Calculate distance matrix for diffusion map based on selected fhp
# trajectory markers
fhp.dist = as.matrix(dist(t(fhp@scale.data[all.markers.fhp, ])))
```

```
# Run diffusion map and return top 50 dimensions
set.seed(1)
fhp.diff = diffuse(fhp.dist, maxdim = 50)
```

```
## Performing eigendecomposition
## Computing Diffusion Coordinates
## Used default value: 50 dimensions
## Elapsed time: 0.561 seconds
```

```
# Diffusion map eigen values
plot(fhp.diff$eigenvals, type = "o", col = "blue", ylab = " Eigenvalues", xlab = "Eigenv
    main = "Diffusion Map Eigenvalues", lwd = 2, cex.lab = 1.5, cex.main = 1.5)
```

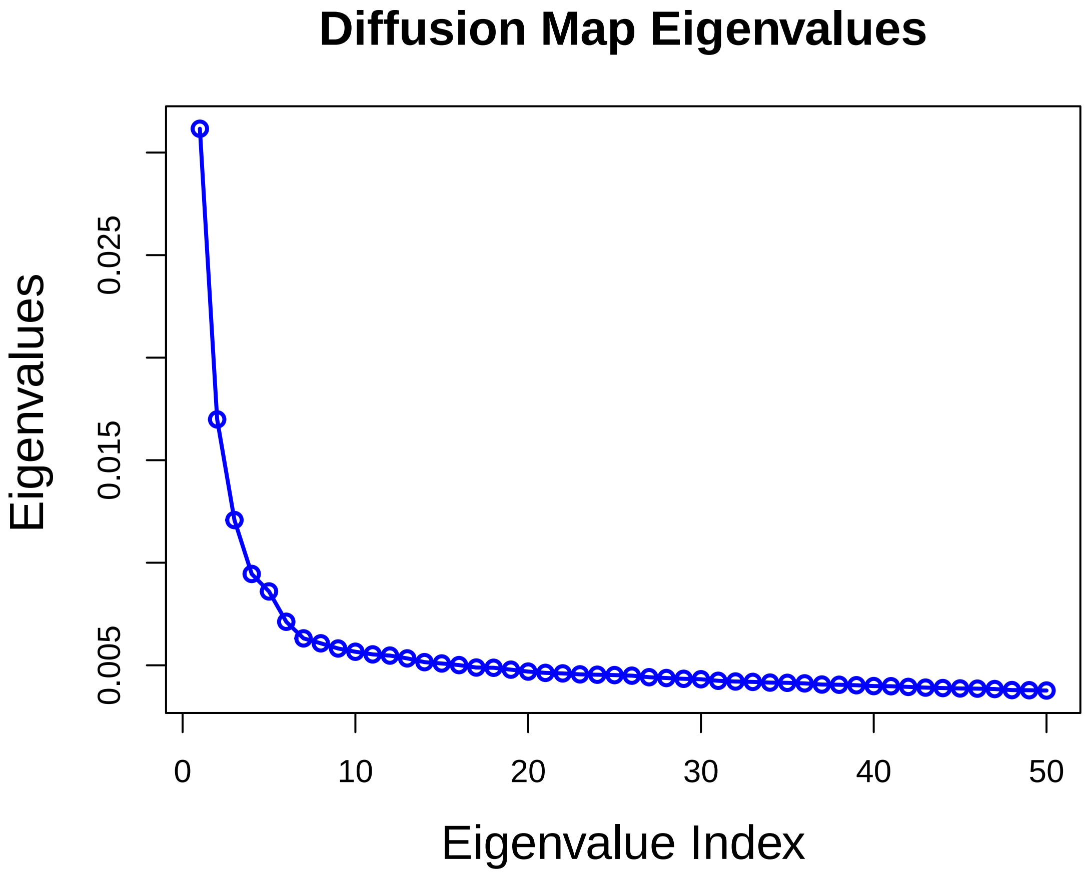


```
# Save first two diffusion map coordinators
fhp@tsne.rot[1:2] = data.frame(fhp.diff$X[, 1:2], row.names = fhp@cell.names)
colnames(fhp@tsne.rot) = c("tSNE_1", "tSNE_2")

# Visualize top two diffusion map components
tsne.pseudo(fhp, do.label = F, label.cex.text = 1, name.y = "Diffusion Map Coordinator 2
    name.x = "Diffusion Map Coordinator1", label.cols.use = c("green", "red",
```

```
        "red", "red", "red"))

# Fit the first two diffusion map components with principal curve
fhp.princurve = principal.curve(as.matrix(fhp.diff$X[, 1:2]), start = as.matrix(fhp.dif
    1:2]))
lines(fhp.princurve$s[order(fhp.princurve$lambda), ], lty = 1, lwd = 4, col = "purple")
```

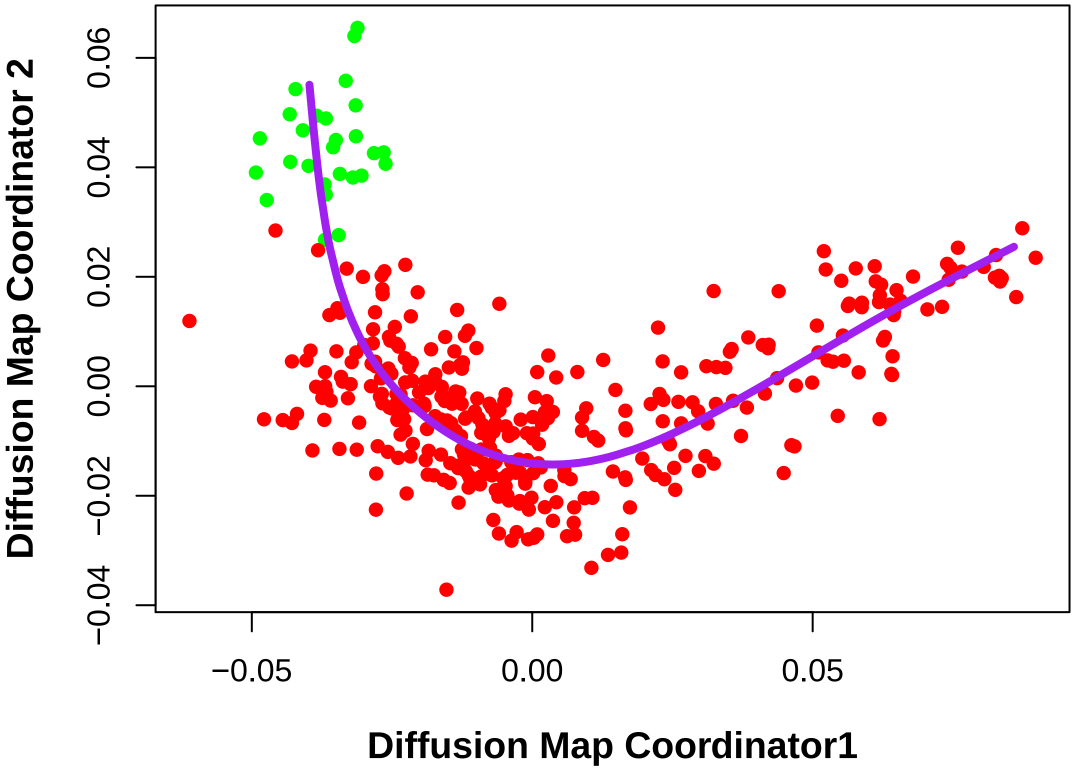


```
df = data.frame(fhp.princurve$s[order(fhp.princurve$lambda), ])
colnames(df) = c("x", "y")
ggplot(data = df, aes(x, y)) + geom_line(size = 1.5, colour = "black") + geom_density2d
    bins = 20) + scale_colour_gradient(low = "darkgray", high = "white", 3) +
    xlim(-0.07, 0.1) + ylim(-0.07, 0.045) + geom_point(data = data.frame(fhp@tsne.rot,
    color = fhp@ident), aes(tSNE_1, tSNE_2), size = 2, color = c(rep("green",
    table(fhp@ident)[1]), rep("red1", table(fhp@ident)[2]), rep("red1", table(fhp@ident)
    rep("red4", table(fhp@ident)[4]), rep("red4", table(fhp@ident)[5]))) + theme_classic
    theme(legend.position = "none", axis.title = element_text(size = rel(1)),
        axis.text = element_blank(), axis.ticks = element_blank(), axis.line.x = element
            size = 0.5, linetype = "solid"), axis.line.y = element_line(colour = "black"
            size = 0.5, linetype = "solid")) + xlab(label = "Diffusion Component 1") +
    ylab(label = "Diffusion Component 2")
```

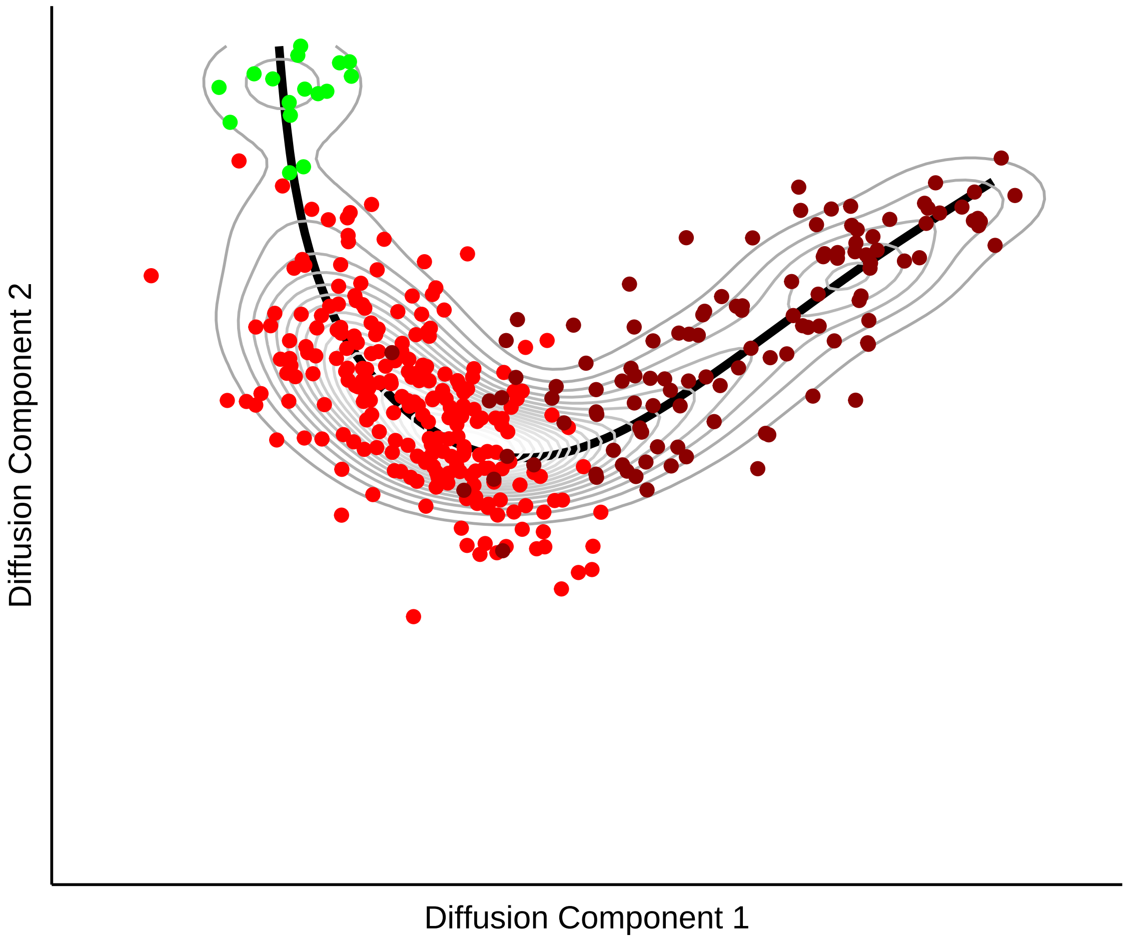


```
# Pseudotime score is assigned to each cell as projection on principal curve
[, "pseudo.time"] = fhp.princurve$lambda

# draw pseudotime cell density
plot(NULL, xlim = c(-0.05, 0.2), ylim = c(0, 100), main = "", ylab = "", xlab = "",
    axes = F)
lines(density([which.cells(fhp, "12TVC"), "pseudo.time"], adjust = 2),
    type = "h", col = "green")
lines(density([which.cells(fhp, "14FHP"), "pseudo.time"], adjust = 2),
    type = "h", col = "red1")
lines(density([which.cells(fhp, c("16FHP")), "pseudo.time"], adjust = 2),
    type = "h", col = "red2")
lines(density([which.cells(fhp, c("18FHP")), "pseudo.time"], adjust = 2),
    type = "h", col = "red3")
lines(density([which.cells(fhp, c("20FHP")), "pseudo.time"], adjust = 2),
    type = "h", col = "red4")
```

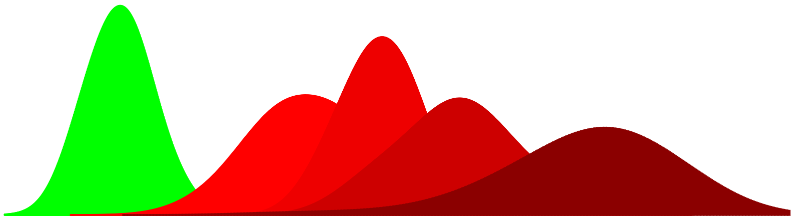

```r
# Scale pseudotime into 0-1 range
fhp.new = scale.pseudo(fhp, "pseudo.time", do.scale = T, do.PC = T)

# Visualize markers on diffusion map
feature.plot.pseudo(fhp, gene.viz, name.x = "Fhp Trajectory", name.y = "", pt.size = 0.7
    nCol = 2)
```

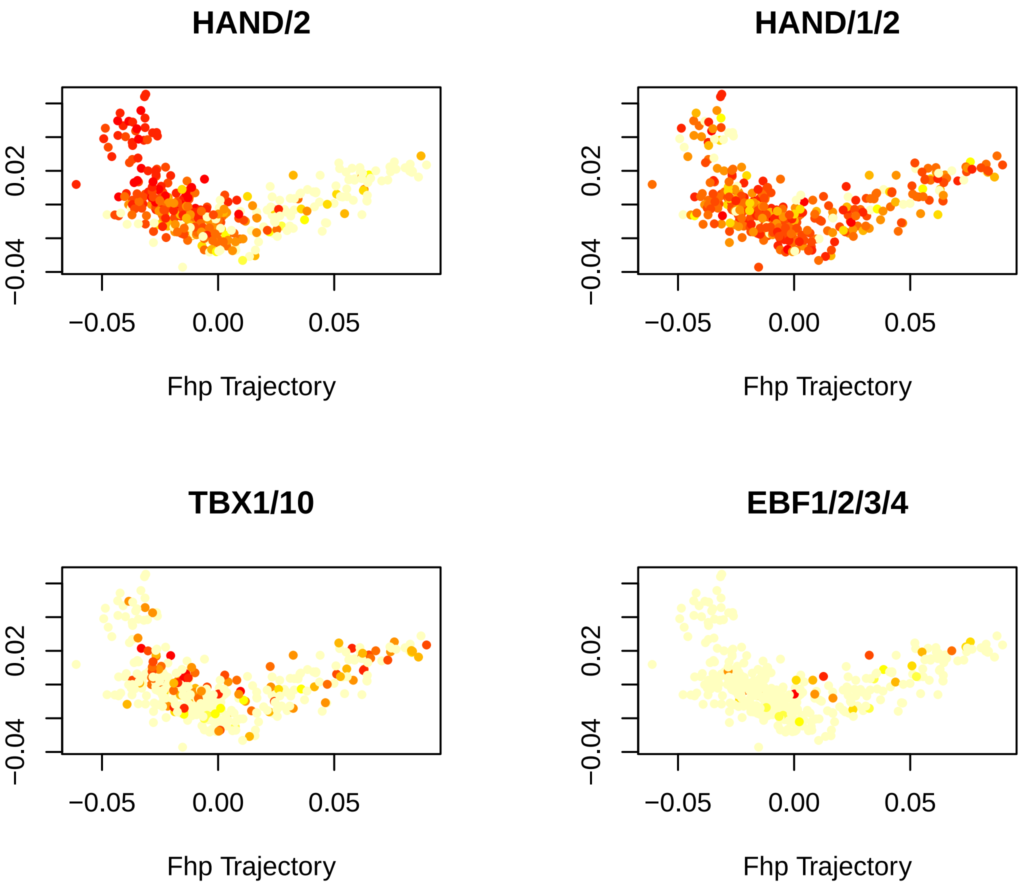


```r
par(mfrow = c(2, 2))
for (i in gene.viz) {
    genePlot.pseudo(fhp.new, gene = i, col.use = c("green", "red1", "red2",
        "red3", "red4"), do.spline = T, name.x = "FHP Trajectory", cex.use = 0.8,
        cex.lab = 1)
}
```

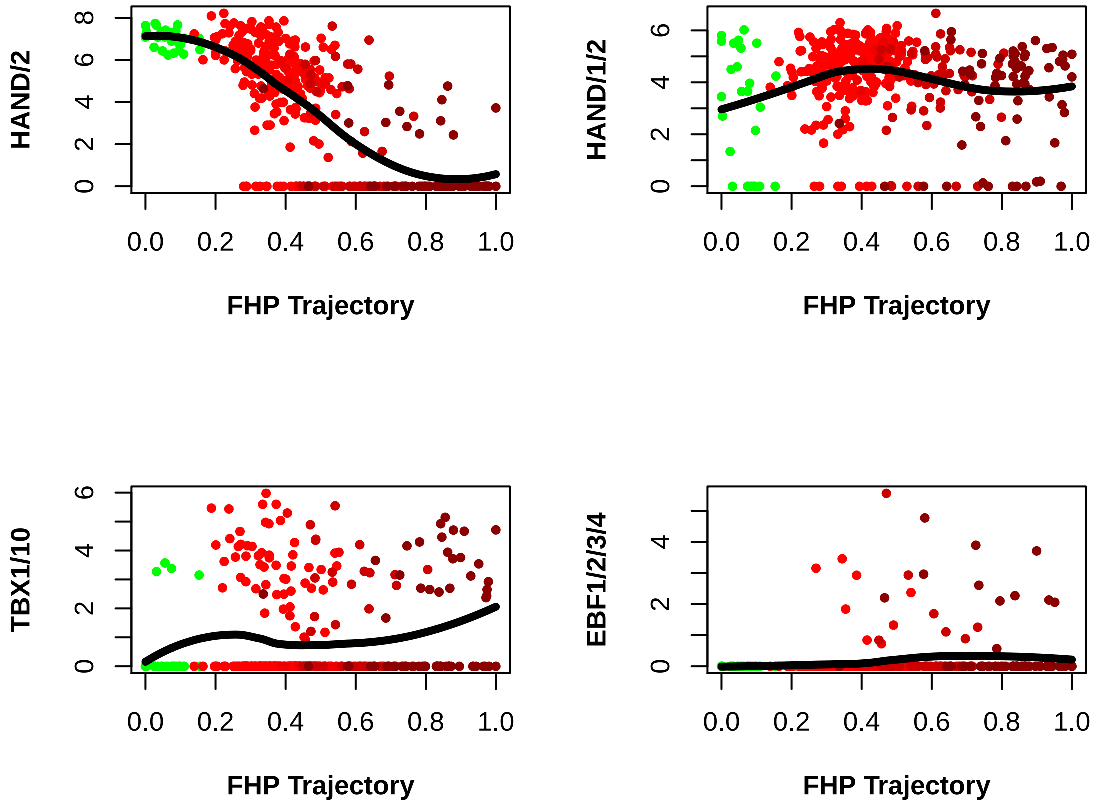


```
# Pseudo time scores correlate with real time
vlnPlot(fhp.new, "pseudo.time", size.x.use = 2)
```

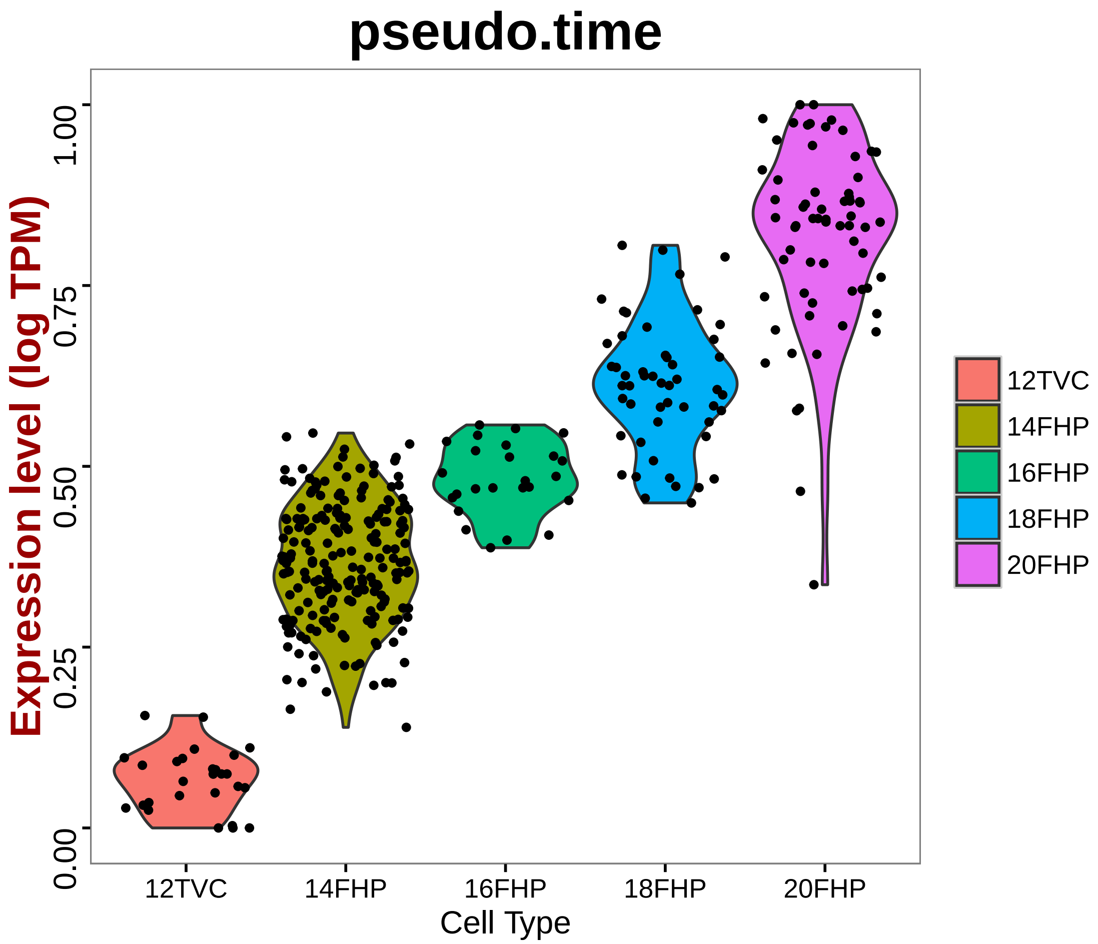


```
plt = vlnPlot(fhp.new, "pseudo.time", size.x.use = 2, do.ret = T)
plt[[1]] + geom_boxplot(width = 0.25, outlier.size = 1, aes(fill = ident)) +
    geom_smooth(aes(group = 1), method = "loess", size = 1, se = F, col = "black",
        lty = 16) + annotate("text", 1.2, 0.9, label = paste("Correlation",
    round(cor(fhp@data.info$real.time, fhp.princurve$lambda), 2), sep = "\n"),
    size = 6, fontface = "bold")
```

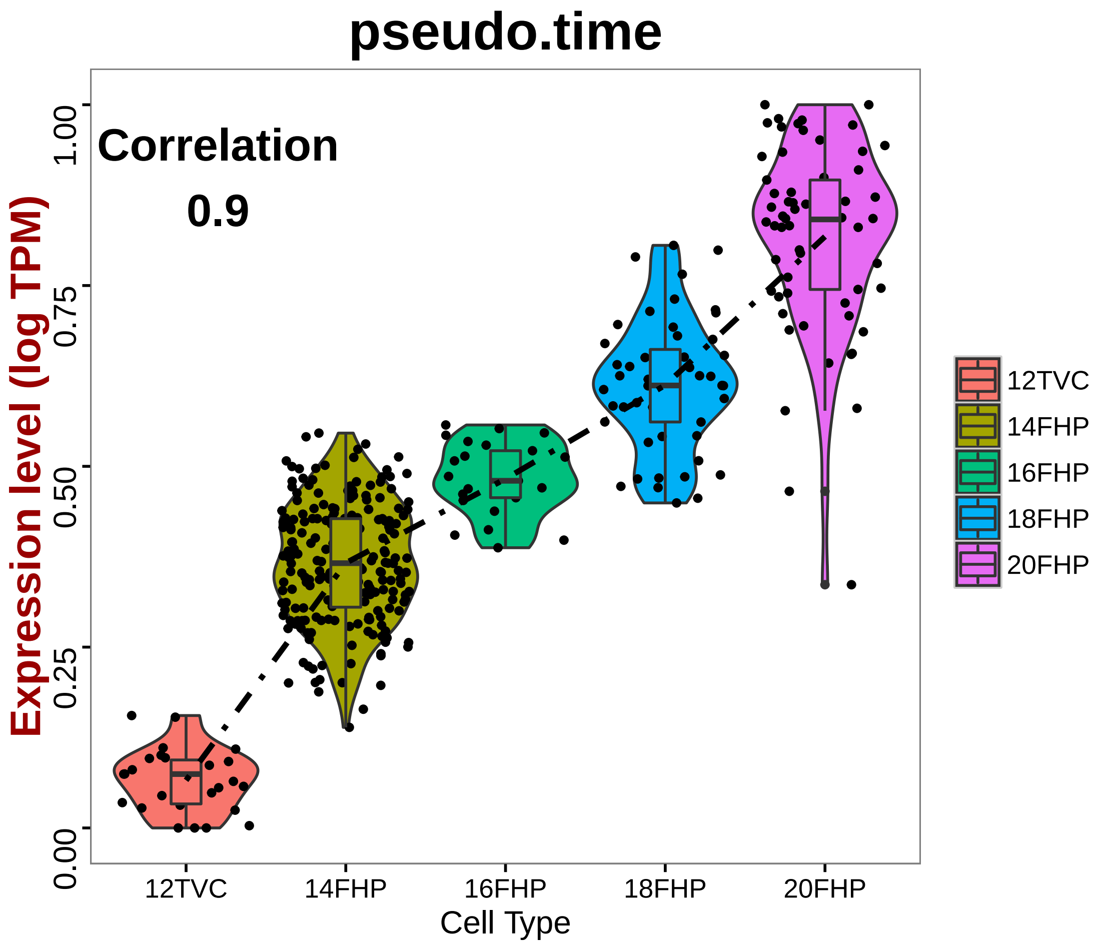


```
fhp.marker.ps = pseudo.gene.cluster(fhp.new, genes.use = all.markers.fhp)
doHeatMap(fhp.new, genes.use = unlist(fhp.marker.ps), remove.key = T, slim.col.label = F
    order.by.ident = F, draw.line = F, cells.use = pcTopCells(fhp.new, 1), labCol = F,
    labRow = F, col = col)
```

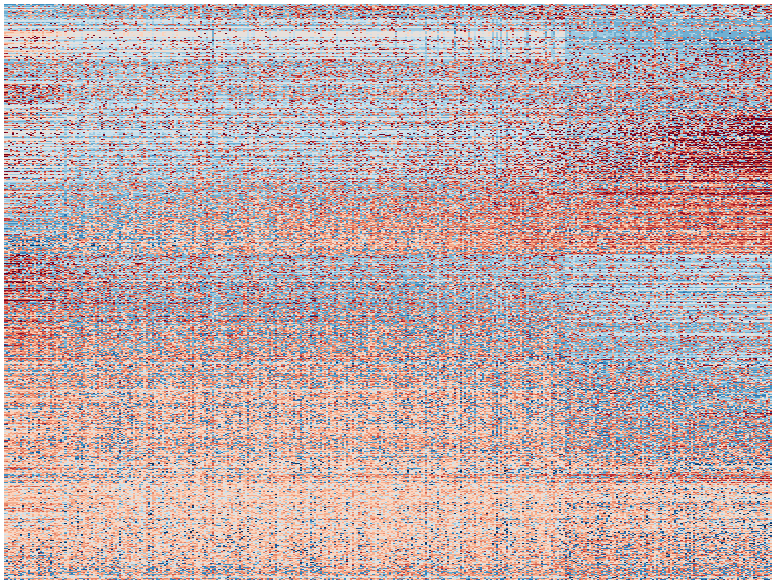

4. ASM Trajectory

```
# Select all asm markers
all.marker.files = list.files(pattern = "markers")
all.markers.asm = c()
for (i in grep("TVC.markers|STVC|ASM", all.marker.files, value = T)) {
    my.txt = read.table(i, sep = "\t", header = T)
    all.markers.asm = unique(c(all.markers.asm, rownames(subset(my.txt, power >
        0.3))))
}
length(all.markers.asm)
```

```
## [1] 926
```

```
# Retrieve asm trajectory
asm = subsetData(hpfall.cluster, which.cells(hpfall.cluster, c("12TVC", "14STVC",
    "16ASM", "18ASM", "20ASM1", "20ASM2")), do.scale = F)
asm@ident = factor(asm@ident, levels = c("12TVC", "14STVC", "16ASM", "18ASM",
    "20ASM1", "20ASM2"), ordered = T)

# Store real time label
[, "real.time"] = as.numeric(substr(asm@ident, 1, 2))

# Calculate distance matrix for diffusion map based on selected asm
# trajectory markers
asm.dist = as.matrix(dist(t(asm@scale.data[all.markers.asm, ])))
```

```r
# Run diffusion map and return top 50 dimensions
set.seed(1)
asm.diff = diffuse(asm.dist, maxdim = 50)
```

```
## Performing eigendecomposition
## Computing Diffusion Coordinates
## Used default value: 50 dimensions
## Elapsed time: 0.996 seconds
```

```r
# Diffusion map eigen values
plot(asm.diff$eigenvals, type = "o", col = "blue", ylab = " Eigenvalues", xlab = "Eigenv
    main = "Diffusion Map Eigenvalues", lwd = 2, cex.lab = 1.5, cex.main = 1.5)
```

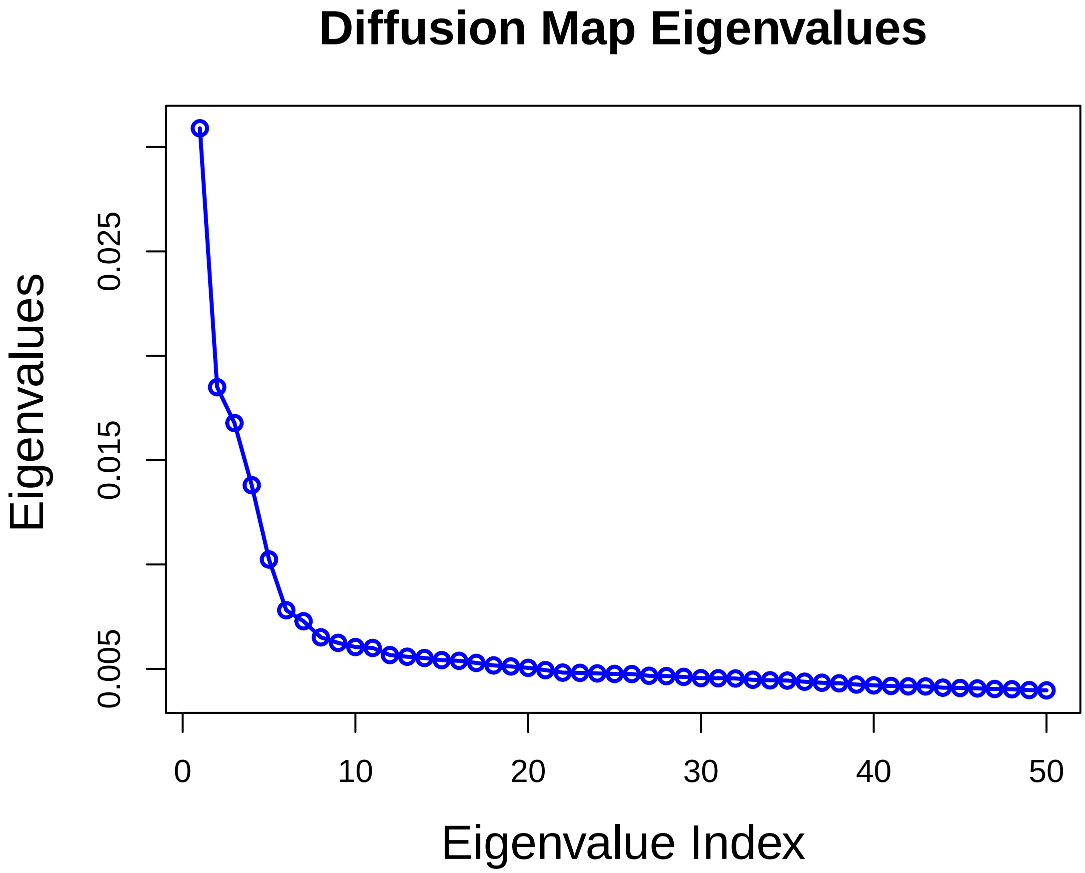


```r
# Save first two diffusion map coordinators
asm@tsne.rot[1:2] = data.frame(asm.diff$X[, 1:2], row.names = asm@cell.names)
colnames(asm@tsne.rot) = c("tSNE_1", "tSNE_2")

# Visualize top two diffusion map components
tsne.pseudo(asm, do.label = F, label.cex.text = 1, name.y = "Diffusion Map Coordinator 2
    name.x = "Diffusion Map Coordinator1", label.cols.use = c("green", "yellow",
```

```
        "blue1", "blue2", "blue3", "blue4"))

# Fit the first two diffusion map components with principal curve
asm.princurve = principal.curve(as.matrix(asm.diff$X[, 1:2]), start = as.matrix(asm.dif
    1:2]))
lines(asm.princurve$s[order(asm.princurve$lambda), ], lty = 1, lwd = 4, col = "purple")
```

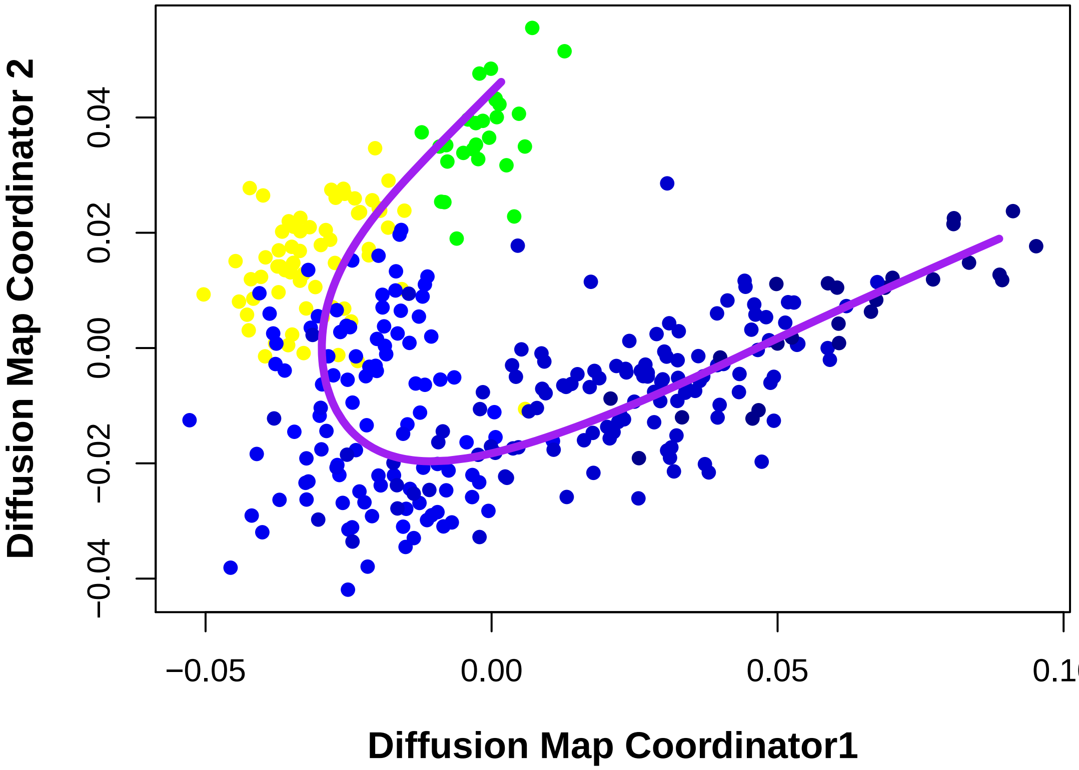


```
df = data.frame(asm.princurve$s[order(asm.princurve$lambda), ])
colnames(df) = c("x", "y")
ggplot(data = df, aes(x, y)) + geom_path(size = 1.5, colour = "black") + geom_density2d
    bins = 6) + scale_colour_gradient(low = "darkgray", high = "white", 3) +
    xlim(-0.06, 0.1) + ylim(-0.06, 0.05) + geom_point(data = data.frame(asm@tsne.rot,
    color = asm@ident), aes(tSNE_1, tSNE_2), size = 2, color = c(rep("green",
    table(asm@ident)[1]), rep("yellow", table(asm@ident)[2]), rep("lightblue",
    table(asm@ident)[3]), rep("blue1", table(asm@ident)[4]), rep("blue4", table(asm@ide
    rep("blue4", table(asm@ident)[6]))) + theme_classic() + theme(legend.position = "no
    axis.title = element_text(size = rel(1)), axis.text = element_blank(), axis.ticks =
    axis.line.x = element_line(colour = "black", size = 0.5, linetype = "solid"),
    axis.line.y = element_line(colour = "black", size = 0.5, linetype = "solid")) +
    xlab(label = "Diffusion Component 1") + ylab(label = "Diffusion Component 2")
```

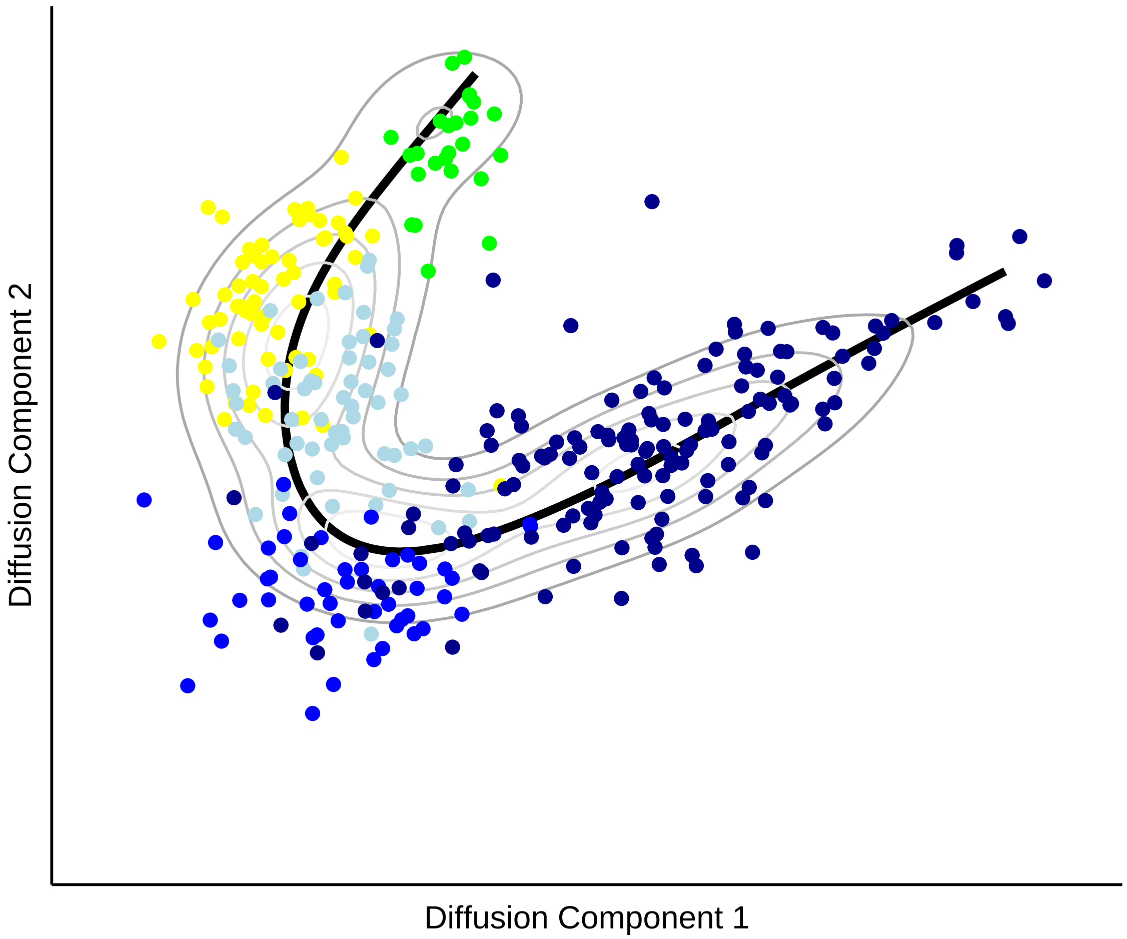


```r
# Pseudotime score is assigned to each cell as projection on principal curve
[, "pseudo.time"] = asm.princurve$lambda

# draw pseudotime cell density
plot(NULL, xlim = c(-0.05, 0.3), ylim = c(0, 100), main = "", ylab = "", xlab = "",
    axes = F)
lines(density([which.cells(asm, "12TVC"), "pseudo.time"], adjust = 2),
    type = "h", col = "green")
lines(density([which.cells(asm, "14STVC"), "pseudo.time"], adjust = 2),
    type = "h", col = "yellow")
lines(density([which.cells(asm, c("16ASM")), "pseudo.time"], adjust = 2),
    type = "h", col = "blue1")
lines(density([which.cells(asm, c("18ASM")), "pseudo.time"], adjust = 2),
    type = "h", col = "blue2")
lines(density([which.cells(asm, c("20ASM1")), "pseudo.time"], adjust = 2),
    type = "h", col = "blue3")
lines(density([which.cells(asm, c("20ASM2")), "pseudo.time"], adjust = 2),
    type = "h", col = "blue4")
```

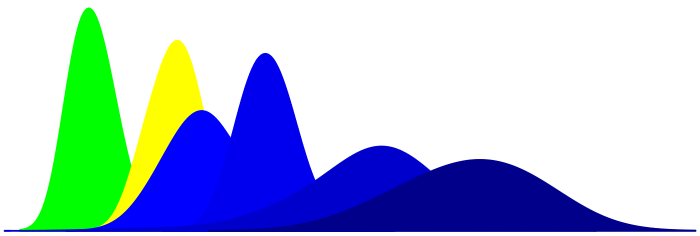

```
# Scale pseudotime into 0-1 range
asm.new = scale.pseudo(asm, "pseudo.time", do.scale = T, do.PC = T)

# Visualize markers on diffusion map
feature.plot.pseudo(asm, gene.viz, yaxt = "n", name.x = "asm Trajectory", name.y = "",
    pt.size = 0.75, nCol = 2)
```

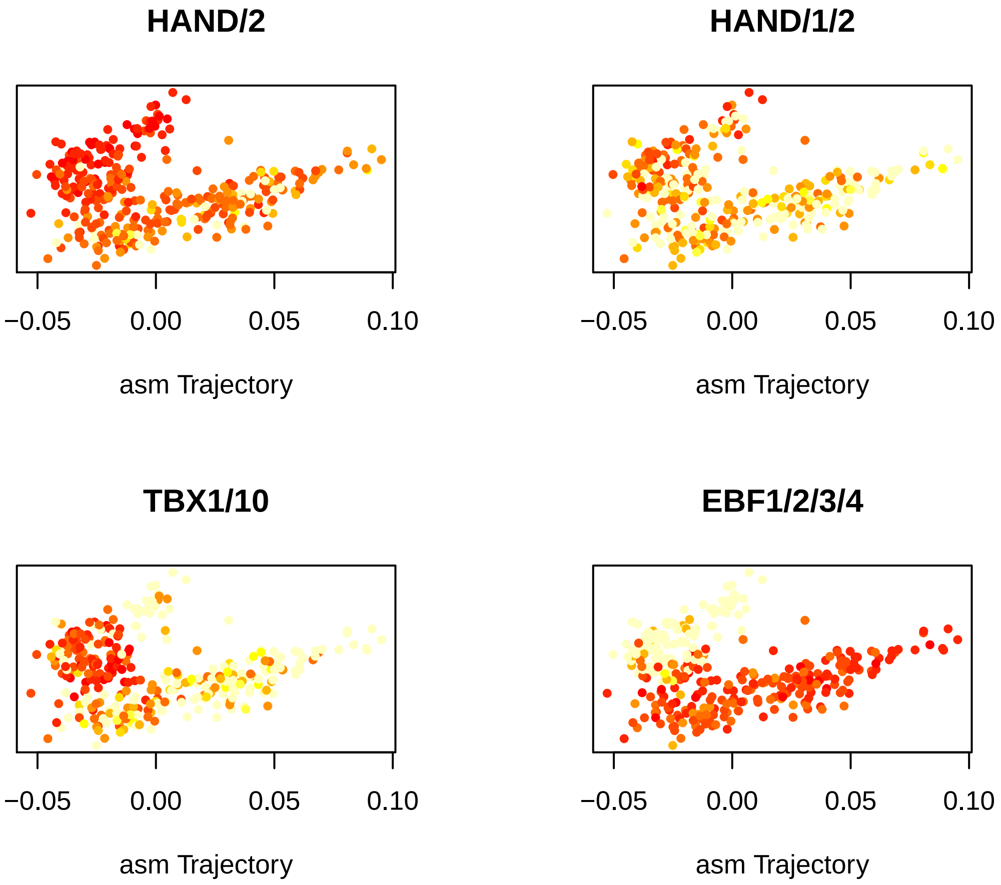


```
par(mfrow = c(2, 2))
for (i in gene.viz) {
    genePlot.pseudo(asm.new, gene = i, col.use = c("green", "yellow", "blue1",
        "blue2", "blue3", "blue4"), do.spline = T, name.x = "ASM Trajectory",
        cex.use = 0.8, cex.lab = 1)
}
```

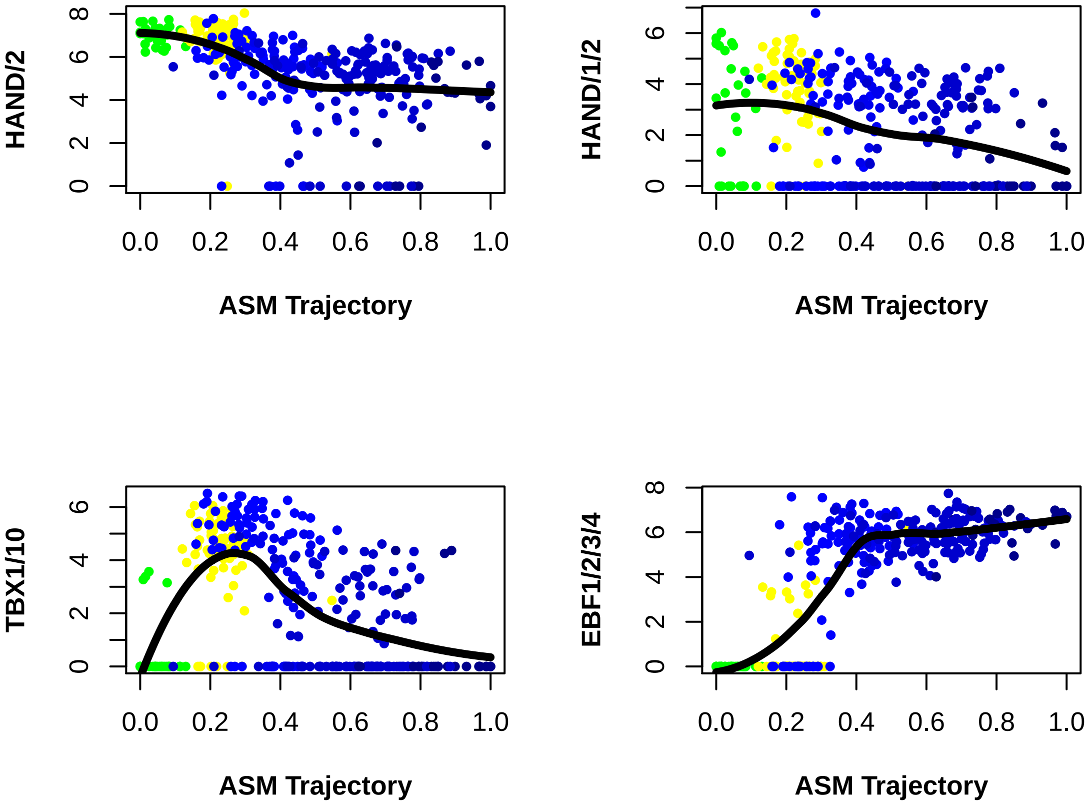


```r
# Pseudo time scores correlate with real time
vlnPlot(asm.new, "pseudo.time", size.x.use = 2)
```

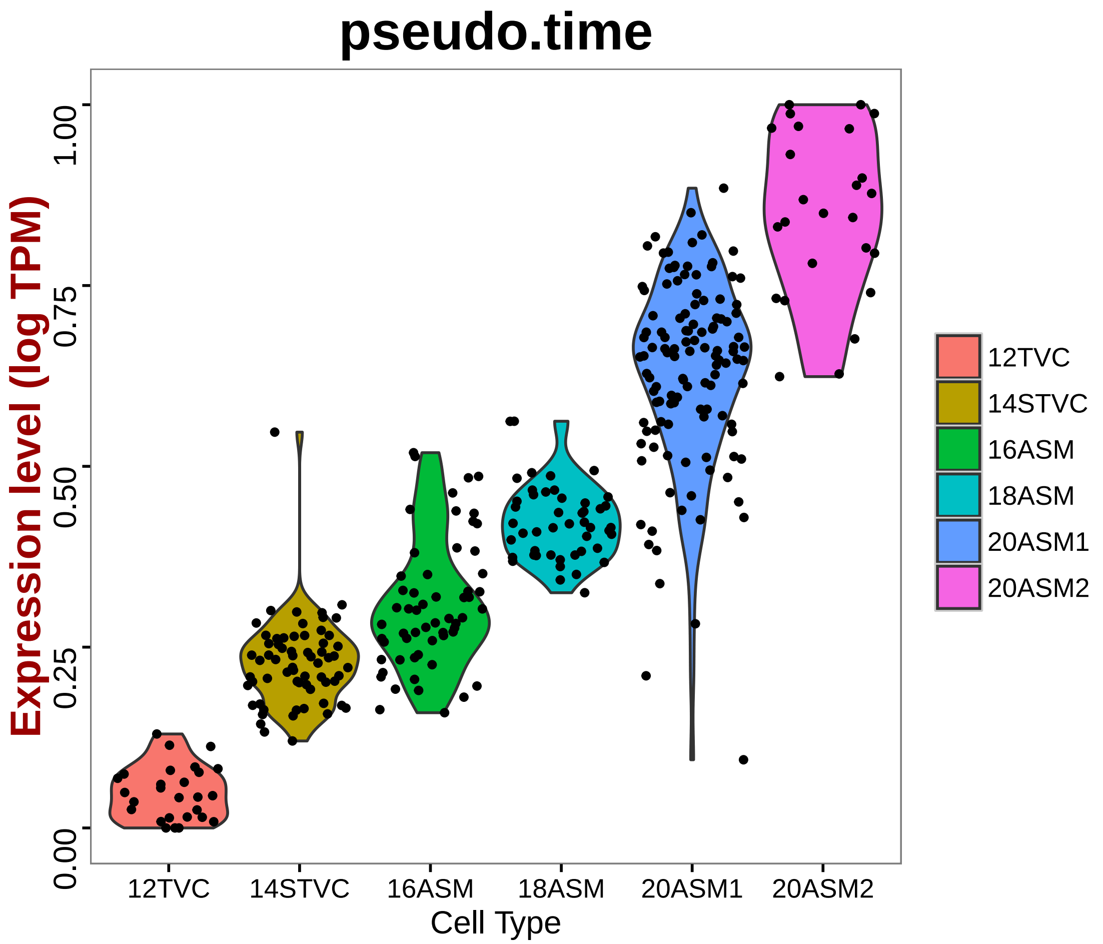


```
plt = vlnPlot(asm.new, "pseudo.time", size.x.use = 2, do.ret = T)
plt[[1]] + geom_boxplot(width = 0.25, outlier.size = 1, aes(fill = ident)) +
    geom_smooth(aes(group = 1), method = "loess", size = 1, se = F, col = "black",
        lty = 16) + annotate("text", 1.2, 0.9, label = paste("Correlation",
    round(cor(asm.new@data.info$real.time, asm.princurve$lambda), 2), sep = "\n"),
    size = 6, fontface = "bold")
```

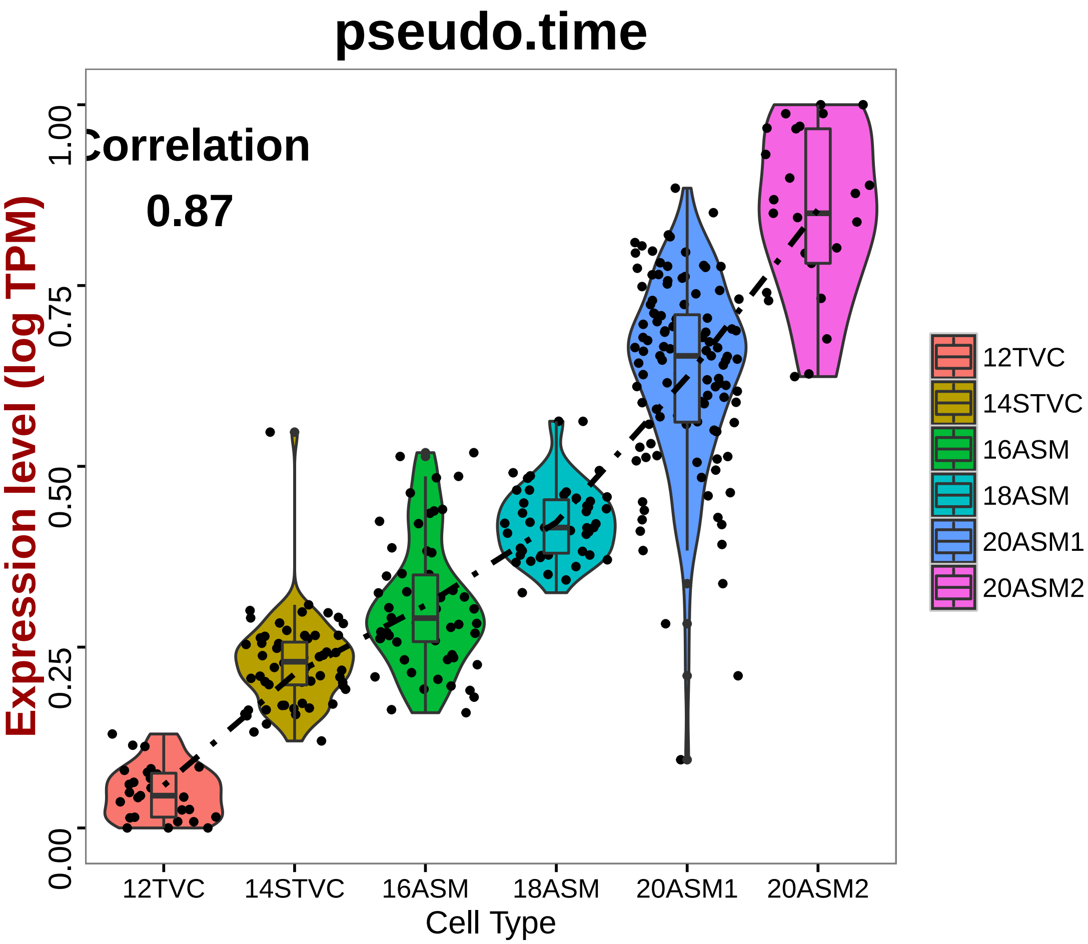


```
asm.marker.ps = pseudo.gene.cluster(asm.new, genes.use = all.markers.asm)
doHeatMap(asm.new, genes.use = unlist(asm.marker.ps), remove.key = T, slim.col.label = F
    order.by.ident = F, draw.line = F, cells.use = pcTopCells(asm.new, 1), labCol = F,
    labRow = F, col.use = col)
```

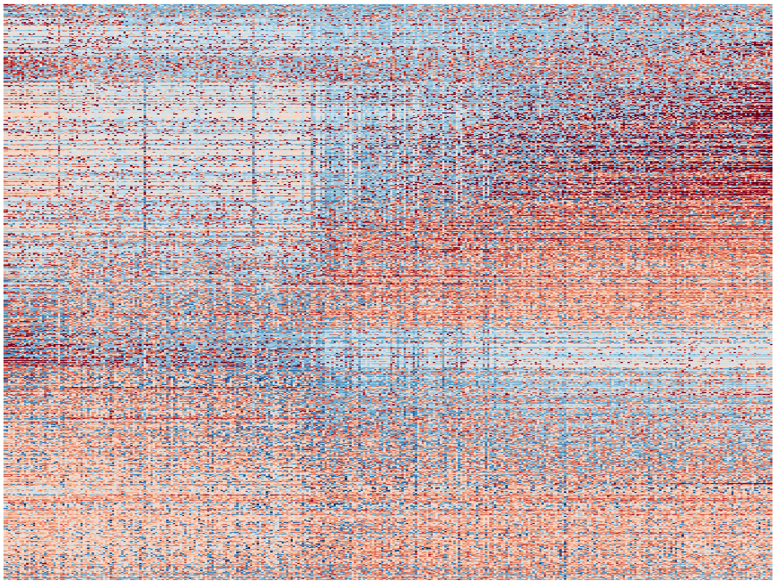

```r
# Save objects
save(shp.new, fhp.new, asm.new, file = "traject.Robj")
```
